# Comprehensive analysis of proteins associated with light responses and stress tolerance in the gametophyte of the fern *Dryopteris affinis* ssp*. affinis*

**DOI:** 10.64898/2025.12.05.692567

**Authors:** Sara Ojosnegros, José Manuel Álvarez, Francisco Vázquez, Naiara Goya, Valeria Gagliardini, Ueli Grossniklaus, Luis G. Quintanilla, Jaume Flexas, Alexis Peña, Helena Fernández

**Affiliations:** Area of Plant Physiology, Department of Organisms and Systems Biology, University of Oviedo, 33071 Oviedo, Spain; (S.O.); (J.M.A.); (F.V.); (N.G.); (A.P.); Department of Plant and Microbial Biology & Zurich-Basel Plant Science Center, University of Zurich, 8008 Zurich, Switzerland; Global Change Research Institute, University Rey Juan Carlos, 28933 Móstoles, Spain; Research Group on Plant Biology under Mediterranean Conditions, University of the Balearic Islands - Agro-Environmental and Water Economics Research Institute, Palma, Spain

## Abstract

The fern gametophyte, despite its crucial role as the pioneer stage ensuring the successful establishment of the fern, has remained largely neglected as a subject of research, being particularly evident at the molecular level. In this study, based on an RNA-sequenced gametophyte transcriptome from the apogamous fern *Dryopteris affinis* ssp. *affinis*, 1,160 proteins functionally associated with light responses, transport processes, and stress mechanisms were identified. Light-responsive proteins are linked to photosynthesis, photorespiration, xanthophyll metabolism, and photomorphogenesis. Among them are several HIGH CHLOROPHYLL FLUORESCENCE (HCF 136, 173 and 244), which participate in the formation of photosystem II, and ENHANCER OF VARIEGATION 3 (EVR3), which is involved in chloroplast biogenesis. Regarding photomorphogenesis, the protein list includes the photoreceptors PHYTOCHROME B (PHYB), CRYPTOCHROMES 1 and 2 (CRY1 and 2), PHOTOTROPIN 2 (PHOT2), and UV RESISTANCE LOCUS 2, 3 and 8 (UVR2, 3 and 8). Likewise, related to transport, 659 proteins were found either in membranes and cytoplasm, moving a wide range of molecules, such as PERMEASE 2 and 3 (AAP2 and 3), SUCROSE TRANSPORTER 4 (SUC4), AMMONIUM TRANSPORTER 1 (AMT1), MECHANOSENSITIVE ION CHANNEL PROTEIN 1 (MSL1), among others. Furthermore, our results comprised an extensive arsenal of proteins to battle against both biotic and abiotic stress, or involved in plant immunity, such as BRASSINOSTEROID-SIGNALLING KINASE 1 (BSK1), NON-RESPONDING TO OXYLIPINS 7 (NOXY7), and CALLOSE SYNTHASE 12 (CALS12); whereas α-MANNOSIDASE 2 (GMII), PALMITOYL-MONOGALACTOSYLDIACYLGLYCEROL Δ-7 DESATURASE (ADS3), and FORGETTER 1 (FGT1) are linked to salt, cold or heat stresses, respectively. Analysis of protein–protein interactions revealed MODIFIED TRANSPORT TO THE VACUOLE 17 (MTV17) as the most connected protein, with 19 interactions, mainly supported by text-mining and database evidence. These findings contribute to expanding the limited knowledge at the molecular level of ferns overall, paving the way for future experimental investigations.

## 1. Introduction

Today’s ferns, a clade of nearly 12,000 species (Nitta et al. 2022), are descendants of the first vascular plants, which appeared on Earth approximately 470 million years ago (Harrison and Morris 2017), thus constituting a valuable genetic legacy. Since ancient times, their only interest has been in ornamentation and traditional medicine, but there is currently a growing interest in investigating its biology from diverse perspectives. The most studied aspects are physiological (Racusen et al. 2002; Salmi et al. 2005; Bushart and Roux 2007; Salmi et al. 2007; Salmi and Bushart 2010; Eeckhout et al. 2014; Tosens et al. 2016), reproductive (Lopez and Renzaglia 2014; Valledor et al. 2014; Suo et al. 2015; de Vries et al. 2016; Grossmann et al. 2017; Rivera et al. 2018; Wyder et al. 2020; Fernández et al. 2021; Ojosnegros et al. 2022, 2023, 2024), ecological (Wang et al. 2010; Dhir 2018; Sareen et al. 2019), as well as pharmacological, with promising discoveries of components with antimicrobial (Nayak et al. 2013), anti-inflammatory (Dion et al. 2014; Huwae and Silahooy 2024), anti-diabetic (Chen et al. 2015; Lamichhane et al. 2019), or anti-cancer properties (Soare and Şuţan 2018; Pratoko et al. 2024). On the other hand, molecular studies on ferns are still scarce. This may be due to their high number of chromosomes and large genome size (Manton 1950; Barker and Wolf 2010; Sessa and Der 2016), which hinders the acquisition and interpretation of molecular data. Indeed, the largest genome (160.45 billion gigabases pairs) of all known eukaryotes has recently been discovered in a fern species: *Tmesipteris oblanceolata* (Fernández et al. 2024). To date, the complete genomes of only seven fern species are available, namely *Salvinia cucullata* (Aragon-Raygoza et al. 2022), *Adiantum capillus-veneris* (Fang et al. 2022)*, Adiantum nelumboides* (Zhong et al. 2022), *Marsilea vestita* (Rahmatpour et al. 2023), *Alsophila spinulosa* (https://fernbase.org), *Azolla filiculoides* (https://fernbase.org), and *Ceratopteris richardii* (https://fernbase.org).

The life cycle of ferns consists of two independent generations: the sporophyte, which forms spores through meiosis and is typically diploid, and the gametophyte, which produces gametes through mitosis and is generally haploid. In addition to sexual reproduction, ferns also exhibit asexual reproduction by apomixis, which comprises two processes: diplospory (generation of unreduced spores by altered meiosis) and apogamy (formation of an embryo from somatic cells of the gametophyte) (Liu et al. 2012; Grusz et al. 2021). Ferns are the terrestrial plant clade with the highest prevalence of apomixis, with ∼10% of fern species showing obligate apogamy (Liu et al. 2012), mainly due to non-functional gametes or limited water availability, which is required for fertilization (Grusz et al. 2021). The fern *Dryopteris affinis* (Lowe) Fraser-Jenk. ssp. *affinis*, hereafter referred to as *D. affinis*, produces non-reduced spores that develop into diploid gametophytes. These gametophytes produce male but not female gametangia, which makes them obligately apogamous (Fraser-Jenkins 1980). In addition, the gametophyte of this species shows other characteristics, such as ease of *in vitro* culture from spores and rapid growth, while whole living individuals can be directly observed by light microscopy (Menendez et al. 2006; Rivera et al. 2018), which makes them a good experimental model (**Fig. 1**). During growth, gametophytes go through several morphological phases. Following spore germination, the gametophyte initially elongates unidirectionally, producing a uniseriate cell chain (filamentous phase; **Fig. 1a**); then, it shifts to two-dimensional growth (spatulate phase; **Fig. 1b**); and finally, the establishment of a notch meristem gives rise to a heart-shaped gametophyte (cordate phase; **Fig. 1c**). Beneath the notch meristem, a central dense region becomes apparent, marking the site where the future sporophyte will develop. In addition, a cluster of elongated cells arranged cylindrically can be observed extending from the embryo to the base of the gametophyte (**Fig. 1c**), which we speculate may act as a sort of rudimentary transport system to nourish the developing sporophyte.

**Figure 1.**
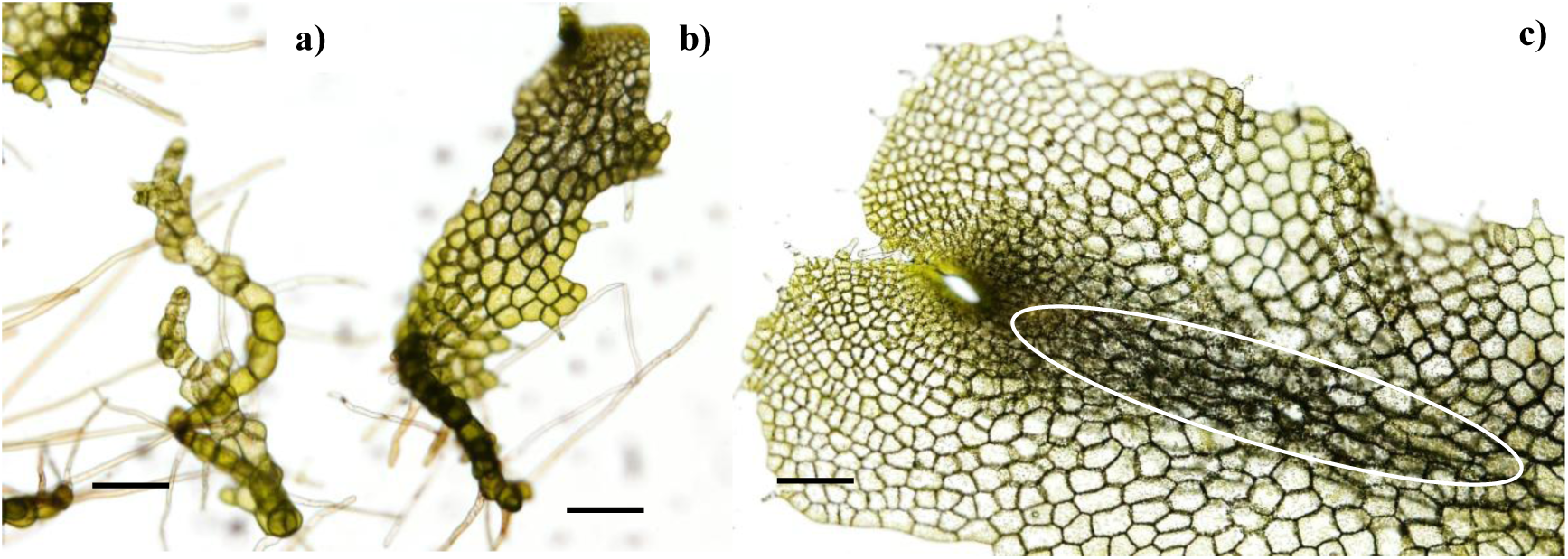
Developmental phases in the gametophyte of *Dryopteris affinis*: **a)** filamentous, **b)** spatulate, **c)** cordate, showing the apomictic embryo in the centre, and elongated cells, resembling a rudimentary transport system (white circle). Images were taken with brightfield microscopy (SCT Olympus BX-61). Bar = 200 µm.

In a previous work, gametophytes of *D. affinis* were analysed by RNA-sequencing, resulting in the first transcriptome of this species (Wyder et al. 2020). These data were compared with sequences from *Arabidopsis thaliana*, as this fern is not yet sequenced, finding numerous annotations of proteins associated with different aspects of plant development, in particular gametogenesis and embryogenesis, including a sort of proteins involved on apomixis (Ojosnegros et al. 2024). In the present work, the stringency was increased by lowering the E-value threshold to less than 10^-100^ so that homologous proteins could be included. Taking this into account, the study concentrated on annotations associated with light response processes (photosynthesis, photorespiration, xanthophyll metabolism, and photomorphogenesis), then on cellular transport, and finally on biotic and abiotic stress responses, thus extending earlier published annotations. In this context, special attention has also been given to the interactions among the annotated proteins and to their potential biological significance.

## 2. Materials and methods

### 2.1. Collection, asepsis and “in vitro” culture of spores

Fronds from sporophytes of *D. affinis*, collected in Turón forest (Asturias, Spain, 43°12′10′′ N - 5°43′43′′ W, 477 m a.s.l.), were put between paper sheets in the lab to release spores from sporangia, and the resulting spores were stored at 4 °C until use. To start the *in vitro* culture of spores, they were previously sterilised. For this purpose, 5 mg of spores were immersed in distilled water at room temperature overnight for hydration, then centrifuged at 2,500 rpm for 5 minutes to remove residual sporangial material, and finally washed with a solution of NaClO 0.5% for 10 min. After sterilisation, the spores were rinsed three times with distilled water, centrifuging between washes at 2,500 rpm for 4 minutes. Subsequently, the spores were cultured in Erlenmeyer flasks that contained 50 ml of liquid Murashige and Skoog (MS) medium (Murashige and Skoog 1962), supplemented with sucrose 2% (w/v), and pH adjusted to 5.7. These flasks were placed on a rotary shaker at 75 rpm inside an incubation chamber at 25 °C, under a photosynthetically active radiation intensity of 70 μmolm^2^s^-1^ and a photoperiod of 16 hours of light. Three weeks later, spores started to germinate and were transferred to pots containing 20 ml of solid MS medium supplemented with agar 0.7% and put under the same conditions as mentioned before for one month. About 50 days after starting spore culture, the gametophytes had passed through their developmental phases and part of them reached the cordate phase. At this point, spatulate and cordate gametophyte were pooled into 100 mg samples, frozen in liquid nitrogen and stored at -80 °C until use.

### 2.2. RNA extraction from gametophytes and transcriptome generation

The methods for carrying out the RNA extraction and sequencing are extensively explained in Wyder et al. (2020). In short, for RNA extraction, previously frozen gametophytes were homogenised with a Silamat S5 shaker (Ivoclar Vivadent, Schaan, Liechtenstein) twice, for 10 and 5 s, respectively. RNA was isolated using the SpectrumTM Plant Total RNA kit (Sigma-Aldrich, Buchs, Switzerland) and any remaining DNA was removed with the TURBO DNA-free kit (Life Technologies, Carlsbad, USA). Afterwards, the RNA quality was checked with the Bioanalyser Agilent RNA 6,000 Pico Kit (Agilent Technologies, Waldbronn, Germany). For transcriptome generation, libraries were prepared with the TruSeq RNA Sample Prep Kit v.2 and sequenced on the Illumina HiSeq 2,000. The Transcriptome Shotgun Assembly project is available at the European Nucleotide Archive under the accession number PRJEB18522. The *de novo* assembly of the transcriptome in FASTA format and the transcriptome annotation were deposited in the Zenodo research data repository (https://doi.org/10.5281/zenodo.1040330). The highly similar contigs assembled by Trinity were collapsed using Corset 0.93 (Davidson and Oshlack 2014) with a distance threshold of 0.3. BUSCO v 2.0.1 was used with the Embryophyta odb9 dataset to assess transcriptome integrity.

### 2.3. Computational analysis

As the *D. affinis* genome is not yet sequenced, the transcriptome dataset obtained was compared to *A. thaliana* sequences with the Geneious Prime software v 2023.2.1 using the Araport11 database. For this purpose, a BLASTX was done. Once the possible homologies were displayed, the results were filtered, selecting only transcript annotations with more than 450 bp and with an E-value of less than 10^-100^. Then, repetitive annotations were eliminated, as well as those annotations with a purely metabolic function, which were not the objective of this work, using those that remained for the analysis with the String program v 12.0 and Uniprot database. Both were used to study the Gene Ontology (GO), including the three categories: biological function, molecular function, and cellular components, and Kyoto Encyclopaedia of Genes and Genomes (KEGG), to infer the possible function of the proteins in *D. affinis* gametophytes. String program provides with data such as protein count, strength, and false discovery rate (FDR). Protein count represents how many proteins in your network are annotated with a particular term, respect to how many proteins in total (in your network and in the background) have this term assigned. Strength indicates the magnitude of the enrichment. It is calculated as the base 10 logarithm of the ratio between the number of proteins in your dataset annotated to a given term, and the number of proteins expected to be annotated within that term in a random dataset of the same size. FDR describes how significant the enrichment is, and it is noticed by the p-values correction, for multiple testing within each category using the Benjamini–Hochberg procedure. Only categories with FDR < 0.05 were considered significantly enriched.

Plots recording the functional enrichment analysis were performed using the *clusterProfiler* package in R (R Core Team 2023). Gene–category interaction networks (cnetplots) were generated to visualize the genes associated with each significant biological or molecular category.

The String program was also used to understand protein-protein interactions, always selecting a high threshold (0.700). In this sense, interactions can be of different types: (a) experiments: proteins that have shown to have chemical, physical or genetic interactions in laboratory experiments; (b) databases: proteins found in the same databases; (c) text-mining: proteins mentioned in the same PubMed abstract or articles from an internal selection of the String program; (d) co-expression: protein expression patterns are similar; (e) neighbourhood: protein-coding genes are close in the genome; (f) gene fusion: in at least one organism orthologous protein-coding genes are fused into an only gene; (g) co-occurrence: proteins that have a similar phylogenetic distribution; and (h) homology: proteins that have a common ancestor and similar sequences. These interactions are assigned scores close to 0 if they are weak, or close to 1,000 if they are strong. In this work, the figures included were elaborated using the next softwares: Cytoscape v 3.10.3, R v 4.3.2 (R Core Team 2023), and RStudio 2023.06 (RStudio Team 2023). Moreover, some *D. affinis* transcripts were compared using BLASTX against fern species with sequenced genomes deposited in FernBase. In nearly all cases, this comparison yielded the identification codes of highly homologous proteins, which currently represent the only available information.

## 3. Results

Our results revealed a substantial set of homologous proteins derived from the gametophyte of the non-sequenced fern *D. affinis*, which exhibited a maximum E-value of 10^-100^ when blasted against the model organism *A. thaliana*. After excluding purely metabolic proteins — those associated with the metabolism of biomolecules — but retaining those involved in photosynthesis, photorespiration, and xanthophyll metabolism, which are one of the focuses of this work, approximately 3,000 protein annotations remained from the total transcriptome. The high degree of homology was notable, as more than half of these proteins showed an exact E-value of 0. Of these 3,000 proteins, we placed particular emphasis on those related to light, transport and stress, representing 1,160 proteins, thoroughly describing and illustrating their main characteristics in this section. The groups of proteins studied and the percentage of proteins that were present in each group can be visualised in **Fig. 2**. The group with more proteins was transport, with up to 45% of the proteins studied, followed by the response to biotic stress with 11% and photomorphogenesis 7%.

**Figure 2.**
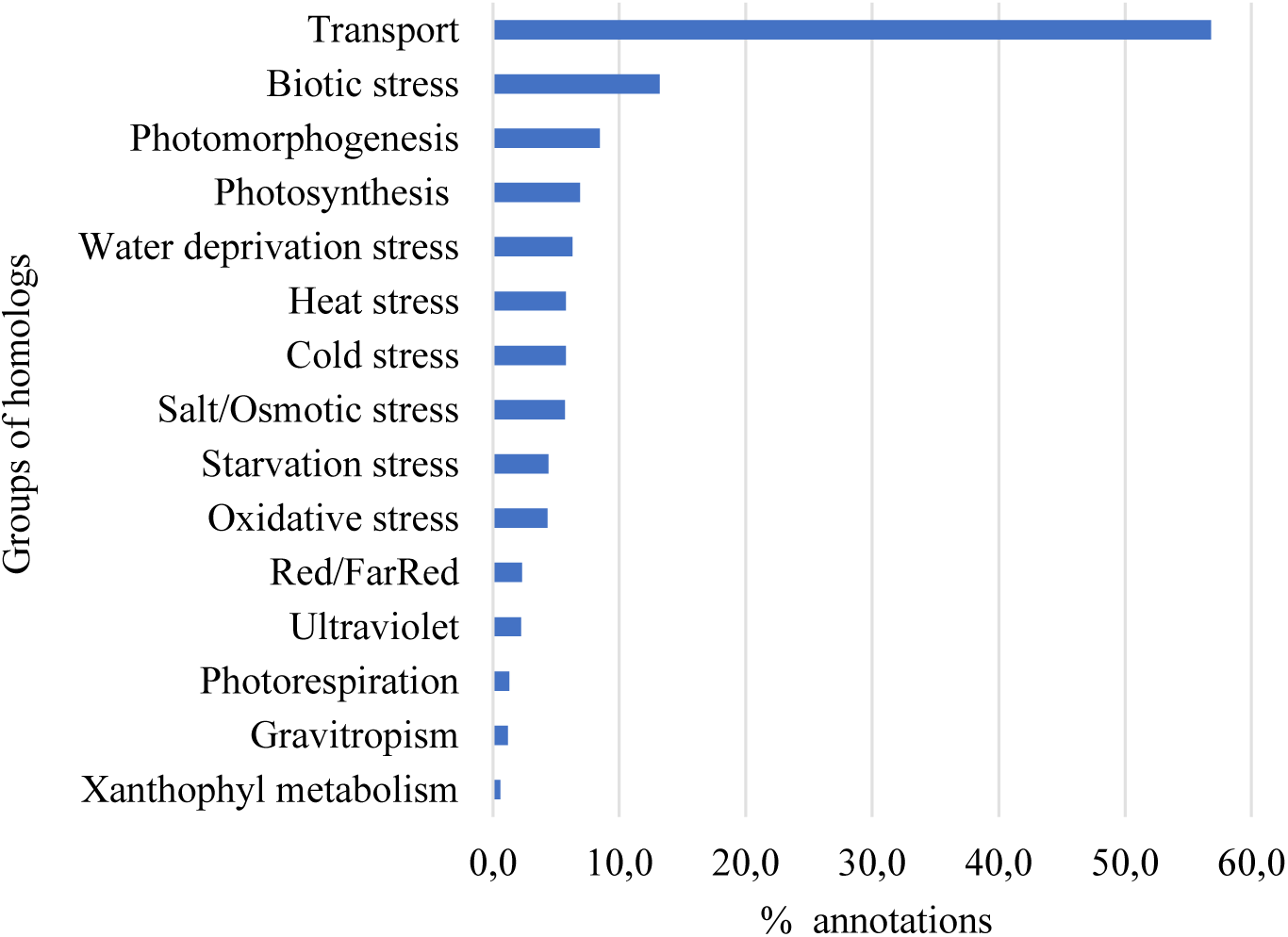
Bar plot showing the percentage of the selected groups of proteins extracted from the gametophyte of *Dryopteris affinis*.

**Table 1** provides the protein count, indicating the number of proteins in each group relative to the total assigned to each category, along with their strength and signal, determined by the String program.

**Table 1.** Functional enrichment of the protein dataset, in the gametophyte of *Dryopteris affinis*, highlighting light-dependent processes, transport functions, and responses to biotic and abiotic stresses. Columns show protein counts, enrichment strength, and adjusted significance (FDR).

| Biological function | Protein count | Strength | FDR |
| --- | --- | --- | --- |
| Photosynthesis | 54 of 250 | 0.71 | $4.43 \cdot 10^{-61}$ |
| Photorespiration | 16 of 53 | 0.85 | $1.51 \cdot 10^{-29}$ |
| Xanthophyll metabolism | 9 of 15 | 1.15 | 0,000407 |
| Photomorphogenesis | 23 of 67 | 0.97 | $3.24 \cdot 10^{-9}$ |
| Response to red or far-red light | 50 of 210 | 0.75 | $1.9 \cdot 10^{-16}$ |
| Response to blue light | 17 of 90 | 0.65 | 0.0023 |
| Response to UV | 30 of 113 | 0.8 | $3.79 \cdot 10^{-10}$ |
| Transport | 663 of 2,659 | 0.77 | 0 |
| Response to the bacterium | 103 of 473 | 0.71 | $1.52 \cdot 10^{-33}$ |
| Response to nematode | 25 of 82 | 0.86 | $3.01 \cdot 10^{-9}$ |
| Response to wounding | 38 of 228 | 0.6 | $4.74 \cdot 10^{-8}$ |
| Response to water deprivation | 71 of 368 | 0.66 | $5.68 \cdot 10^{-60}$ |
| Response to oxidative stress | 68 of 434 | 0.57 | $2.26 \cdot 10^{-82}$ |
| Response to salt stress | 86 of 444 | 0.66 | $3.52 \cdot 10^{-166}$ |
| Response to starvation | 52 of 207 | 0.77 | $4.79 \cdot 10^{-18}$ |
| Response to cold | 96 of 391 | 0.76 | $1.12 \cdot 10^{-34}$ |
| Response to heat | 72 of 254 | 0.83 | $1.28 \cdot 10^{-28}$ |

‘Protein count’ indicates the number of proteins from the dataset annotated to a given biological function, together with the total number of proteins known for that category. A high count in broad categories such as photosynthesis or transport highlights the large contribution of these proteins to the dataset. ‘Strength’ represents the degree of enrichment, calculated as the ratio between the observed and expected protein frequencies; values above 1 reflect strong overrepresentation, even if the absolute number of proteins is small, as observed for xanthophyll metabolism. ‘FDR’ (false discovery rate) corresponds to the adjusted probability after multiple testing correction (Benjamini–Hochberg), with extremely low values (e.g., 10⁻⁶¹ for photosynthesis or 10⁻¹⁶⁶ for salt stress) indicating very robust associations. Together, these parameters show that the dataset is particularly enriched in light-dependent processes (photosynthesis, photorespiration, photomorphogenesis, and responses to red, blue, and UV light), as well as in transport proteins and stress responses to both biotic and abiotic factors.

### 3.1. Proteins associated with light: photosynthesis, photorespiration and xanthophyll metabolism

The proteins associated with light were categorised according to the Gene Ontology (GO). Regarding the biological function, two main groups are deemed: one involved in the processes of photosynthesis, photorespiration and xanthophyll metabolism, while the other plays a significant role in photomorphogenesis. As for the first group, the molecular function of most of the proteins was linked to the binding of molecules such as ions and nucleic acids, followed by catalytic and oxidoreductase activities. In terms of the GO classification of cellular components, many of the proteins were found within the cytoplasmic space and plastid organelles, as well as in the cell membrane. The most prevalent protein domains included chlorophyll A-B binding and aldolase-type TIM barrel.

The enrichment analysis identified several biological processes significantly associated with the corresponding genes. To illustrate these associations, gene–category interaction networks (Cnetplots) were generated for this group of proteins (**Fig. 3**). Each network displays the relationships between enriched GO terms and their corresponding genes. Only categories that remained statistically significant after multiple testing correction using the false discovery rate (FDR < 0.05) were retained, ensuring that the visualized processes reflect robust functional associations.

**Figure 3.**
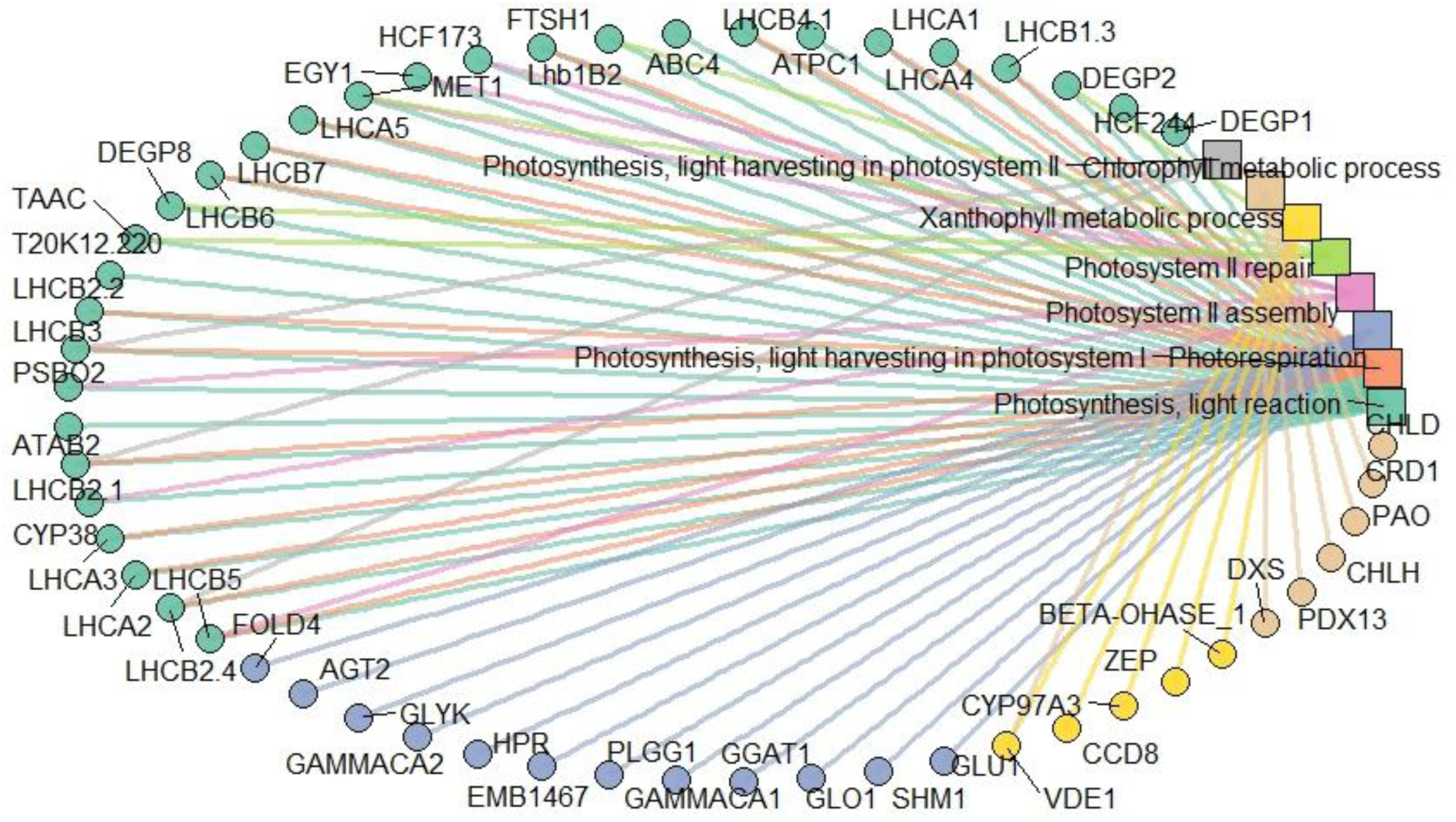
Circular gene–category interaction network (Cnetplot) for photosynthetic processes. Only significantly enriched categories (FDR < 0.05) are included. Node size of categories is scaled according to significance (−log10 FDR), while node colour distinguishes different categories. Circular nodes represent genes, and square nodes represent GO terms.

Upon examining the biological functions of the *A. thaliana* homologue proteins devoted to photosynthesis, it was evident that they primarily participate in the electron transport chain. Key proteins in this process are PROTON GRADIENT REGULATION 6 (PGR6), a kinase that regulates the function of plastoquinone, β-carotene, and xanthophyll lutein homeostasis, LEAF-TYPE CHLOROPLAST-TARGETED FNR 1 (LFNR1) and 2 (LFNR2), which manage electron flow to fulfil the plant’s requirements for ATP and reducing power, and K (+) EFFLUX ANTIPORTER 3 (KEA3), which promotes photosynthesis when the chloroplast ATP synthase activity is low, by reducing the pH gradient across the thylakoid membrane. In addition, proteins involved in the formation, assembly and stabilisation of photosystem II were identified, such as HIGH CHLOROPHYLL FLUORESCENCE 136 (HCF136) along with two auxiliary proteins: HIGH CHLOROPHYLL FLUORESCENCE PHENOTYPE 173 (HCF173) and 244 (HCF244), and CYCLOPHILIN 38 (CYP38). Besides, a significant number of subunits of the light-harvesting complex, which captures and transfers excitation energy to photosystems, were present, including LIGHT-HARVESTING CHLOROPHYLL B-BINDING 2 (LHCB2), PHOTOSYSTEM I LIGHT HARVESTING COMPLEX GENE 1 (LHCA1), and LIGHT HARVESTING COMPLEX PHOTOSYSTEM II (LHCB4). Associated with this, a protein necessary to maintain the efficiency of the light-harvesting complex, SUPPRESSOR OF QUENCHING 1 (SOQ1), was annotated. Likewise, three subunits of the magnesium-chelatase complex involved in chlorophyll biosynthesis were found. In support of a proper chloroplast development, two proteins are reported: ENHANCER OF VARIEGATION 3 (EVR3) and FTSH PROTEASE 1 (FTSH1). Furthermore, proteins related to various other categories, including carbon fixation, as well as regulators of photosynthesis and stomatal movement, were also noticeable.

Within the group of homologs involved in photorespiration, various enzymes were identified as key players in the photorespiratory core cycle, including HYDROXYPYRUVATE REDUCTASE (HPR) and HYDROXYPHENYLPYRUVATE REDUCTASE 2 (HPPR2). Additionally, the regulation of amino acid biosynthesis and catabolism during photorespiration involves GLUTAMATE: GLYOXYLATE AMINOTRANSFERASE 1 (GGAT1). Proteins associated with protein homotrimerization, biosynthesis of hydrogen peroxide, and tetrahydrofolate interconversion were also observed.

Regarding xanthophyll metabolism, several homologs involved in its biosynthesis were detected in the transcriptome, including β-CAROTENOID HYDROXYLASE 1 (BCH1) and LUTEIN DEFICIENT 5 (LUT5). In addition, VIOLAXANTHIN DE-EPOXIDASE 1 (VDE1), a protein that regulates the concentration of zeaxanthin in chloroplasts as part of the xanthophyll cycle, was also identified.

### 3.2. Proteins associated with light: photomorphogenesis

The enrichment analysis highlighted key biological processes related to the genes of interest. In particular, photomorphogenesis emerged as significantly represented. Among the proteins reported here, a considerable number were associated with this process, which describes the effects of light on plant development. It is well-established that plants respond to various light wavelengths, including red, far-red, blue, and UV. Ion and DNA binding, as well as catalytic and transferase activities, were the predominant molecular functions, mainly associated with the nucleus and cytoplasm, and being WD-40 repeats and PAS the most abundant domains.

The analysis of enriched GO terms of biological functions in this group of proteins associated with photomorphogenesis (**Fig. 4**) reveals proteins involved in processes mediated by continuous far-red light, such as LONG AFTER FAR-RED 3 (LAF3), and by red and far-red light, such as PHYTOCHROME B (PHYB). In addition, two transcription factors implicated in phytochrome signalling were identified: VASCULAR PLANT ONE ZINC FINGER PROTEIN (VOZ1), which acts on phytochrome B, and PHYTOCHROME A SIGNAL TRANSDUCTION 1 (PAT1), which acts on phytochrome A. Besides, several serine/threonine-protein phosphatases and kinases are linked to the regulation of phytochromes. For the response to blue light, notable proteins implicated included CRYPTOCHROME 1 and 2 (CRY1 and 2), PHOTOTROPIN-2 (PHOT2), and G-PROTEIN COUPLED RECEPTOR 1 (GCR1). Additionally, ADAGIO 1 (ADO1), a component of an E3 ubiquitin ligase complex involved in the blue light-dependent circadian cycle, was identified. Regarding the response of gametophytes after being exposed to UV light, several homologs were identified involved in repairing DNA damage, including UV RESISTANCE 2 and 3 (UVR2 and UVR3). Related to the response to UV-B specifically, homologs that confer resistance and tolerance to this wavelength were found, such as HOMOLOG OF REVERSIONLESS 1 (REV1) and WD REPEAT-CONTAINING PROTEIN CSA-1 (CSA1). Also, some proteins focused on root protection against UV-B were found: ROOT UV-B SENSITIVE 1 (RUS1) and 2 (RUS2). Likewise, the numerous subunits of the COP9 signalosome complex, relevant in photomorphogenesis, are highlighted. Particularly, seven components of this complex were identified in the transcriptome. Moreover, several additional proteins were detected, including ROOT PHOTOTROPISM PROTEIN 3 (JK218), a signal transducer involved in the phototropic response; DAMAGED DNA BINDING PROTEIN 1A (DDB1A), a component of the light signal transduction machinery; and EMBRYO DEFECTIVE 168 (EMB168), an E3 ubiquitin-protein ligase. The latter two proteins are implicated in the repression of photomorphogenesis in darkness. Besides, a protein related to the repression of photomorphogenesis in light, SPA1-RELATED 3 (SPA3), was found.

**Figure 4.**
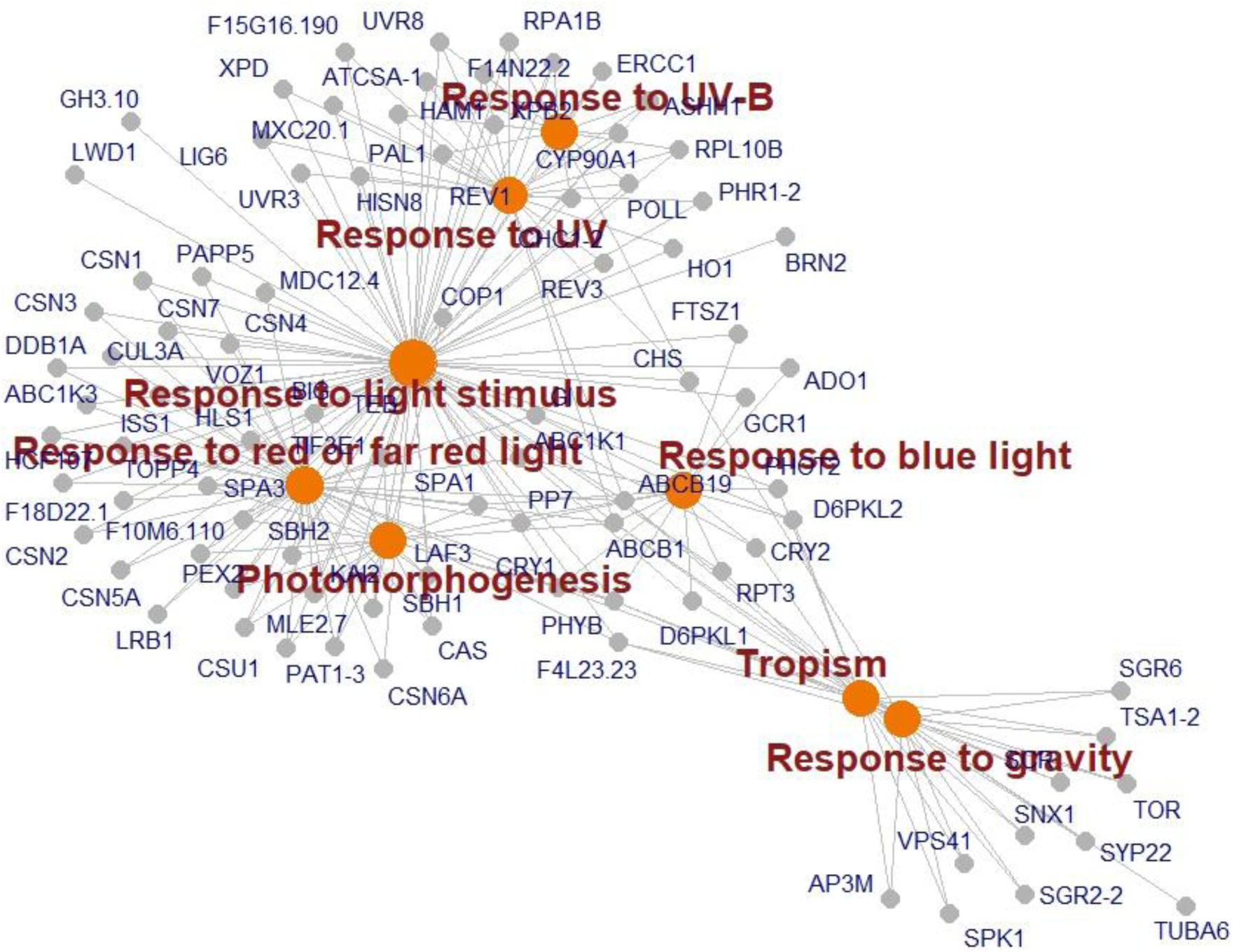
Gene–category interaction network (Cnetplot) related to photomorphogenesis. Only significantly enriched categories after false discovery rate correction (FDR < 0.05) are included. Grey nodes represent genes, and orange nodes represent GO terms in the gametophyte of the fern *Dryopteris affinis*.

As well as photomorphogenesis, gravitropism also affects plant growth, in which the gravity modulates its direction. In our protein set, some homologs related to this process were identified, such as those implicated in the amyloplast sedimentation in the endodermis during shoot gravitropism, where they act as statoliths, named SHOOT GRAVITROPISM 1 (SGR1) and 2 (SGR2). As it is well known, the gametophyte of ferns lacks endodermis.

### 3.3. Proteins associated with transport

In this work, a total of 659 different proteins associated with transport were found. Functional enrichment analysis revealed that several transport-related GO terms were significantly overrepresented (**Fig. 5**). The dot plots highlight the main biological processes (BP) and molecular functions (MF) associated with transport (**Fig. 5 a, b**), including a strong enrichment for ATP-binding cassette (ABC) transporter activity, being the primary protein domains the ABC transporter transmembrane region and the mitochondrial carrier. To further explore these associations, an interaction network of ABC transporter genes was constructed using Cytoscape (**Figure 5c**).

**Figure 5.**
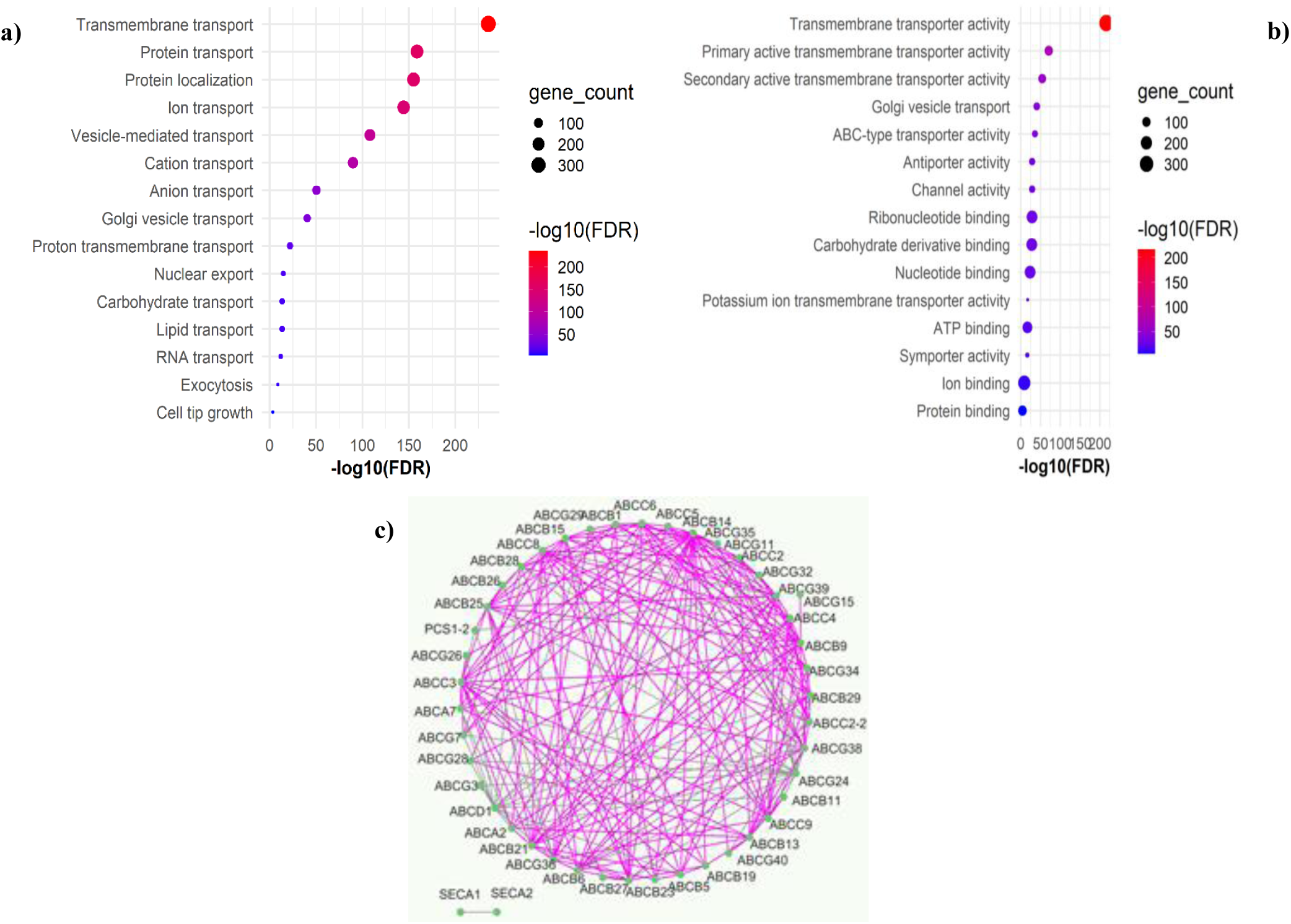
Functional enrichment and interaction network of transport-related proteins identified in the gametophyte of the fern *Dryopteris affinis*. **a)** and **b)** Dot plots showing significantly enriched GO terms related to biological process, and molecular function, respectively. **c)** Interaction network of ABC transporters generated with Cytoscape, where nodes represent ABC transporter proteins (mainly from the ABCB subfamily) and edges indicate functional or predicted associations among them. Pink colour refers to protein-protein interactions coming from experimental evidence.

The classification of biological functions points out that the most abundant categories were associated with transmembrane transport, cellular localisation, protein and ion transport, and vesicle-mediated transport. These proteins bind to a pleiad of compounds like anions and cations, hormones, proteins, lipids, carbohydrates, nucleic acids, nitrogenous bases, etc., and move them through the cell. In terms of cellular components, nearly all the proteins were localised in the cell membrane and the cytoplasm.

Notably, as shown in **Fig. 5c**, a significant number of ATP-binding cassette (ABC) transporters were identified. Specifically, the gene count revealed a total of 47 different proteins belonging to this family, which were type A, B, C, D, E or G. This group included members involved in hormone transport, such as ATP-BINDING CASSETTE B1 (ABCB1) and G40 (ABCG40); in fatty acids transport, such as D1 (ABCD1); as well as those responsible for ion transport, like C5 (ABCC5), which regulates K^+^ and Na^+^ levels in the cell.

As mentioned above, the proteins involved in transport encompass a long archive, which is briefly commented. Then, some examples of transporters of different compounds, such as AMINO ACID PERMEASE 2 (AAP2), LYSINE HISTIDINE TRANSPORTER 1 (LHT1), and CATIONIC AMINO ACID TRANSPORTER 1 (CAT1), with affinity, as its name suggests, for cationic amino acids. Other important transporters included an ATP/ADP/AMP exchanger in the inner mitochondrial membrane: ADENINE NUCLEOTIDE TRANSPORTER 1 (ADNT1); a protein involved in ammonium uptake from the soil and transport to the shoots: AMMONIUM TRANSPORTER 1 (AMT1); other relative that facilitates ammonium transport between the apoplast and symplast of the cells: AMMONIUM TRANSPORTER 2 (AMT2); and a sugar proton-coupled antiporter located in the vacuole: TONOPLAST SUGAR TRANSPORTER 2 (TST2).

Another important group of transporters are represented by all those involved in the endocytosis and exocytosis traffic. In line with it, multiple subunits of the clathrin-associated adaptor protein complex and the exocyst complex were reported. In terms of nuclear import, components of the nuclear pore complex, such as NUCLEOPORIN 155 (NUP155), were identified. Furthermore, a protein essential for the transport of adenine, guanine and uracil was also noted: PLASTIDIC NUCLEOBASE TRANSPORTER (PLUTO). Another remarkable result was the obtaining of various aquaporins, i.e. channels in the cell membrane that facilitate water permeability, among them NAMED PLASMA MEMBRANE INTRINSIC PROTEIN 1;2 (PIP1;2) and 2;1 (PIP2;1). In the gametophyte of *D. affinis*, some mechanosensitive ion channels were also observed, such as MECHANOSENSITIVE ION CHANNEL PROTEIN 1 (MSL1), which opens in response to stretch forces in the membrane lipid bilayer.

Following with ion transport, some additional examples can be given, such as a boron transporter from roots to shoots: BORON TRANSPORTER 1 (BOR1); a calcium-dependent protein that catalyses the import of ATP co-transported with metal divalent cations across the mitochondrial inner membrane in exchange of phosphate: ATP/PHOSPHATE CARRIER 2 (APC2); a K^+^/H^+^ antiporter that regulates K^+^ uptake and homeostasis in the cell: CATION/H+ EXCHANGER 17 (CHX17); and a homolog needed in copper import into the cell: HEAVY METAL ASSOCIATED PROTEIN 51 (HMP51). Finally, it is worth emphasizing the identification of numerous transporters for molecules such as nitrate, magnesium, zinc, sulphate, sucrose, citrate, and phosphate.

### 3.4. Proteins associated with stress

The proteins related to stress were categorised into two groups based on the type of stressor: biotic stress (resulting from bacteria, nematodes, or viruses) and abiotic stress (resulting from factors such as water deprivation, oxidative stress, salt stress, starvation, cold, or heat). The classification of molecular functions predominantly featured binding and catalytic activities, while the cellular component classification was predominantly represented by the cytoplasm and the cell membrane. Identified protein domains included chlorophyll A-B binding and the DnaJ central domain. The set of proteins extracted from *D. affinis* gametophytes related to biotic stress is shown in **Fig. 6**.

**Figure 6.**
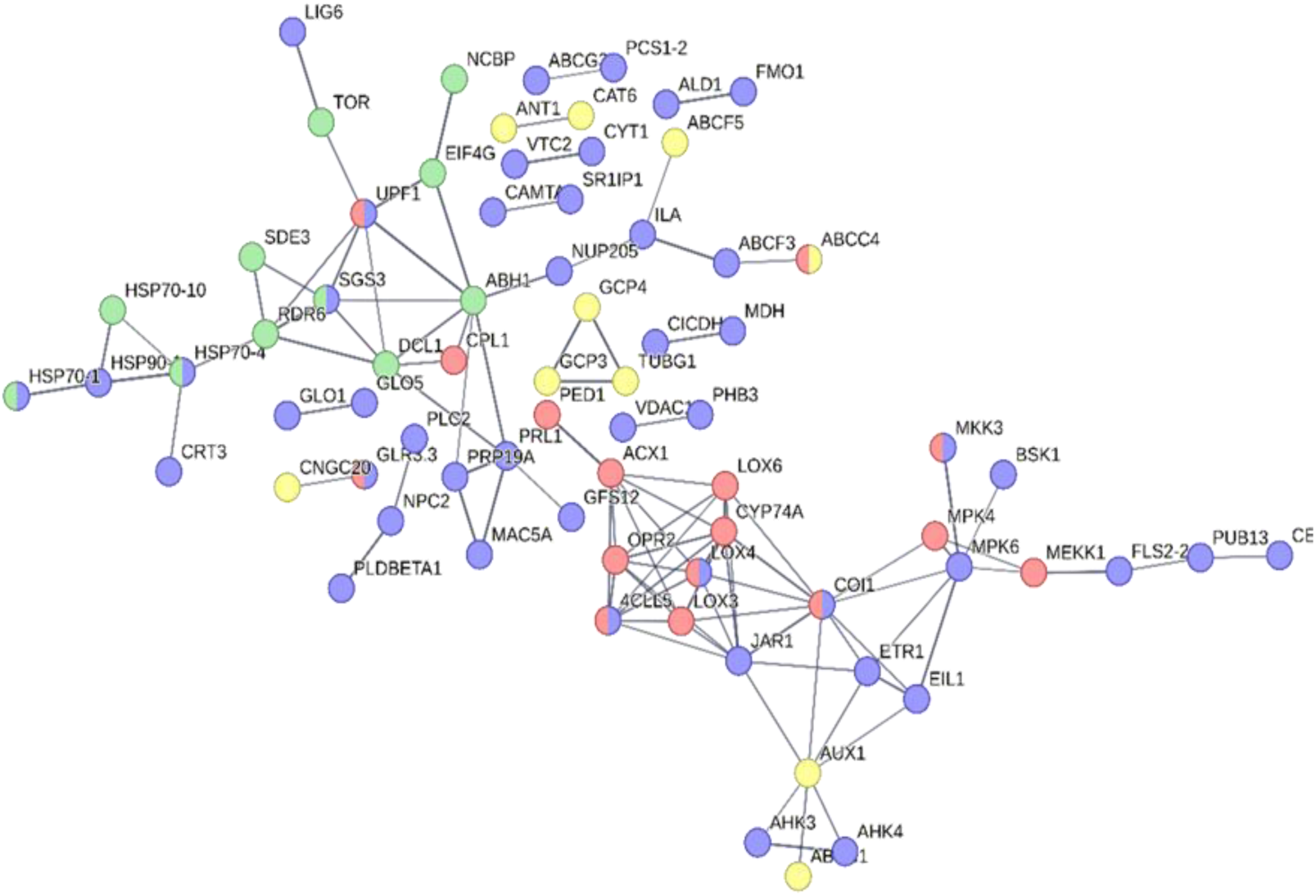
Network of proteins obtained from gametophytes of *Dryopteris affinis* related to response to bacterium (blue balls), nematodes (yellow balls), virus (green balls), and wounding (red balls), provided by String program v 12.0.

Many *A. thaliana* homologous proteins reported were involved in resistance to both bacterial and fungal attacks, including CERAMIDASE (ACER), AMINOTRANSFERASE ALD1 (ALD1), L-TYPE LECTIN RECEPTOR KINASE S.4 (LECRK-S.4), GUANINE NUCLEOTIDE-BINDING PROTEIN SUBUNIT β (AGB1), and β-GLUCOSIDASE 42 (BGLU42). Additionally, two proteins involved in regulating defence against bacteria — HOPW1-1-INTERACTING 1 (WIN1) and 2 (WIN2) — were detected, along with a homolog of the receptor FLAGELLIN-SENSITIVE 2 (FLS2), which recognises bacterial flagellin, a pathogen-associated molecular pattern that triggers plant defence responses. This was complemented by a protein involved in fungal defence that also restricts bacterial growth: LYSM-CONTAINING RECEPTOR-LIKE KINASE 1 (LYK1). It is worth mentioning homologs that play roles in defence against insects, such as SIGNAL RESPONSIVE 1 (SR1), and against nematodes, such as EMBRYO DEFECTIVE 2728 (EMB2728).

Related to biotic stress, some proteins involved in plant immunity were annotated, such as BRASSINOSTEROID-SIGNALING KINASE 1 (BSK1), NON-RESPONDING TO OXYLIPINS 7 (NOXY7), SERINE/THREONINE-PROTEIN KINASE/ENDORIBONUCLEASE IRE1A (IRE1A), PLANT U-BOX 59 (PUB59), SUPPRESSOR OF CPR5 44 (SCPR44), and ASPARTYL PROTEASE APCB1 (APCB1). A few proteins essential for the induction, establishment, and activation of systemic acquired resistance, such as TRANSCRIPTION FACTOR TGA2 (TGA2), FLAVIN-DEPENDENT MONOOXYGENASE 1 (FMO1), and SUPPRESSOR OF FATTY ACID DESATURASE DEFICIENCY 1 (SFD1), were also identified. Additionally, a homolog involved in chitin-triggered immune signalling, PBS1-LIKE 34 (PBL34), and another regulating the hypersensitive response, RESPIRATORY BURST OXIDASE PROTEIN F (RBOH F), were found. Several proteins linked to immunity signalling, such as MITOGEN-ACTIVATED PROTEIN KINASE KINASE KINASE 1 (ARAKIN) and PBS1-LIKE 39 (PBL39), were also observed.

In addition, a protein related to cell death as a defence mechanism was included: PLEIOTROPIC DRUG RESISTANCE 8 (PDR8). E3 UBIQUITIN-PROTEIN LIGASE NLA (SYG1) was associated with general defence responses, while CALLOSE SYNTHASE 12 (CALS12) was linked to callose formation after wounding. Similarly, other homologs that regulate the biosynthesis of metabolites responsible for defence, such as GREENING AFTER EXTENDED DARKNESS 1 (GED1); related to disease resistance, such as NRL PROTEIN FOR CHLOROPLAST MOVEMENT1 (NCH1); and to non-host disease resistance, such as ORNITHINE-Δ-AMINOTRANSFERASE (Δ-OAT), were notable findings.

Transitioning from biotic to abiotic stress, we observed that *A. thaliana* homolog proteins associated with various stressors, such as water deprivation, oxidative stress, salt stress, starvation, cold, and heat, were well-documented. Some networks can be seen in **Fig. 7**.

**Figure 7.**
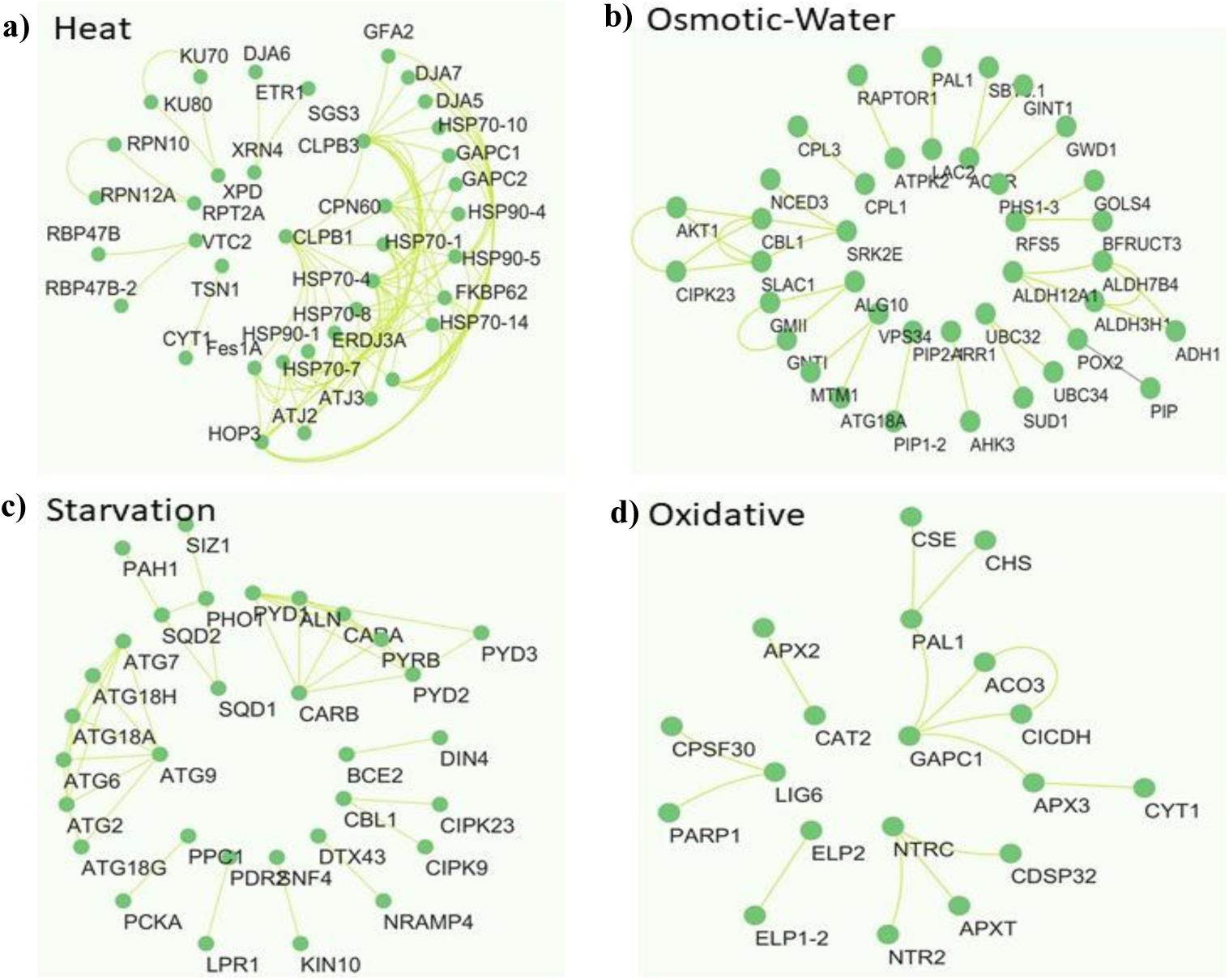
Networks of proteins extracted from gametophytes of *Dryopteris affinis* associated with the response to stress by: **a)** heat, **b)** osmotic/water deprivation **c)** starvation, and **d)** oxidation, provided by Cytoscape v 3.10.3. Green colour refers to protein-protein interactions coming from database evidence.

Focusing first on water deprivation, several proteins involved in the response to dehydration were identified, including one which enhances its tolerance by improving ion retention, called PYROPHOSPHATE-ENERGIZED VACUOLAR MEMBRANE PROTON PUMP 1 (VHP1), and another which regulates drought responses, named CALCINEURIN B-LIKE PROTEIN 1 (CBL1).

The gametophyte also contains homologs related to oxidative stress. Some examples were an enzyme which restores the functionality of proteins that have been inactivated through methionine oxidation: METHIONINE SULFOXIDE REDUCTASE A4 (MSRA4); two proteins involved in resistance to oxidative stress by intervening in reactive oxygen species production: ACTIVITY OF BC1 COMPLEX KINASE 7 (ABC1K7) and 8 (ABC1K8); other that contributes to maintain the reactive oxygen species homeostasis in mitochondria: ALKALINE/NEUTRAL INVERTASE A (A/N-INVA); and LIPOCALIN IN THE PLASTID (LCNP), which prevents thylakoidal membrane lipids peroxidation, conferring it protection against oxidative stress. Other noteworthy achievements were a protein that helps plants to perceive singlet oxygen in plastids, avoiding any possible photooxidative damage that can take place: EXECUTER 2 (EXE2); the protein Γ-GLUTAMYL TRANSPEPTIDASE 1 (GGT1), which prevents oxidative stress by metabolizing extracellular oxidized molecules of glutathione; and GLYCERALDEHYDE-3-PHOSPHATE DEHYDROGENASE C SUBUNIT 1 (GAPC1), responsible for plant adaptation to conditions generated by oxidative stress. Other noteworthy findings included a protein that intervenes in redox homeostasis, named GLYOXYLATE/SUCCINIC SEMIALDEHYDE REDUCTASE 1 (GLYR1), and another that promotes oxidative stress resistance, called NEUTRAL CERAMIDASE 1 (NCER1), along with several ascorbate peroxidases that eliminate hydrogen peroxide.

Homologs associated with salt stress resistance and tolerance included α-MANNOSIDASE 2 (GMII), PRE-mRNA SPLICING FACTOR SR-LIKE 1 (SRL1), PROTEIN PHOSPHATASE 2C GROUP 1 (PP2CG1), and BEACH-DOMAIN HOMOLOG A1 (BCHA1). Moreover, two proteins essential for generating responses to salt stress were DEAD-BOX ATP-DEPENDENT RNA HELICASE 3 (RH3) and MA3 DOMAIN-CONTAINING TRANSLATION REGULATORY FACTOR 3 (MRF3); whereas PROTEIN KINASE 2 (PK2) is involved in plant adaptation to high salinity conditions.

Regarding starvation, the *D. affinis* gametophyte exhibited a range of *A. thaliana* homologous proteins: PEROXISOME UNUSUAL POSITIONING 2 (PEUP2), essential for autophagosome formation under nutrient-limited conditions; 14-3-3-LIKE PROTEIN GF14 PSI (GRF3), which regulates nutrient metabolism; 2-OXOISOVALERATE DEHYDROGENASE SUBUNIT β 2 (DIN4), required during sugar starvation; SOS2-LIKE PROTEIN KINASE 17 (PKS17), conferring tolerance to low potassium conditions; and PROTEIN KINASE 6 (PKS6), required for potassium homeostasis under limited availability of this ion. Significant findings were observed in an activator of the amino acid biosynthesis pathway required during adaptation to amino acid starvation: GENERAL CONTROL NON-DEREPRESSIBLE 2 (GCN2), and in a protein which alleviates the stress generated during copper-limiting periods: KIN17-LIKE PROTEIN (KIN17). In addition, we detected PHOSPHATIDIC ACID PHOSPHOHYDROLASE 1 (PAH1), an enzyme involved in galactolipid synthesis in the endoplasmic reticulum, required for membrane lipid remodelling, as an essential adaptation to phosphate starvation.

For combating cold stress, the fern gametophyte relied on diverse proteins, including several calmodulin-binding transcription activators that contribute to freezing resistance. Related to this, we annotated 3-HYDROXYISOBUTYRYL-COA HYDROLASE 1 (CHY1), involved in cold signalling and tolerance; SENSITIVE TO FREEZING 2 (SFR2), which also plays a role in freezing tolerance; and ENOLASE 2 (ENO2), a regulator of cold-responsive gene transcription. In addition, we found Δ (8)-FATTY-ACID DESATURASE 2 (SLD2), an enzyme required for sphingolipid formation, necessary for cold resistance; other proteins crucial for maintaining chloroplast membrane integrity during low temperatures, such as β-KETOACYL-ACP SYNTHETASE 2 (KAS2); and other needed in chloroplast biogenesis in frigid conditions: PALEFACE 1 (PFC1). The set of proteins associated with cold stress is completed by LONG-CHAIN BASE KINASE 2 (LCBK2), which is implicated in reduced root growth under freezing conditions.

On the other hand, the response to heat stress involved several noteworthy homologous proteins, such as a FORGETTER 1 (FGT1), which facilitates the acquisition of heat stress memory; HEAT INTOLERANT 4 (HIT4), a protein needed for basal thermotolerance, regulating transcriptional gene silencing; and SERINE/THREONINE-PROTEIN PHOSPHATASE 7 (PP7), which confers thermotolerance to high temperatures. Likewise, a high number of chaperones and heat shock proteins expanded the annotations.

After performing a general analysis of the entire set of proteins related to light, transport and stress, approximately 180 proteins were selected (see **Table 2**). Their sequences are provided in **Supplementary Table 1**. The sole criterion for selection criterion was the availability and abundance of information.

**Table 2.**
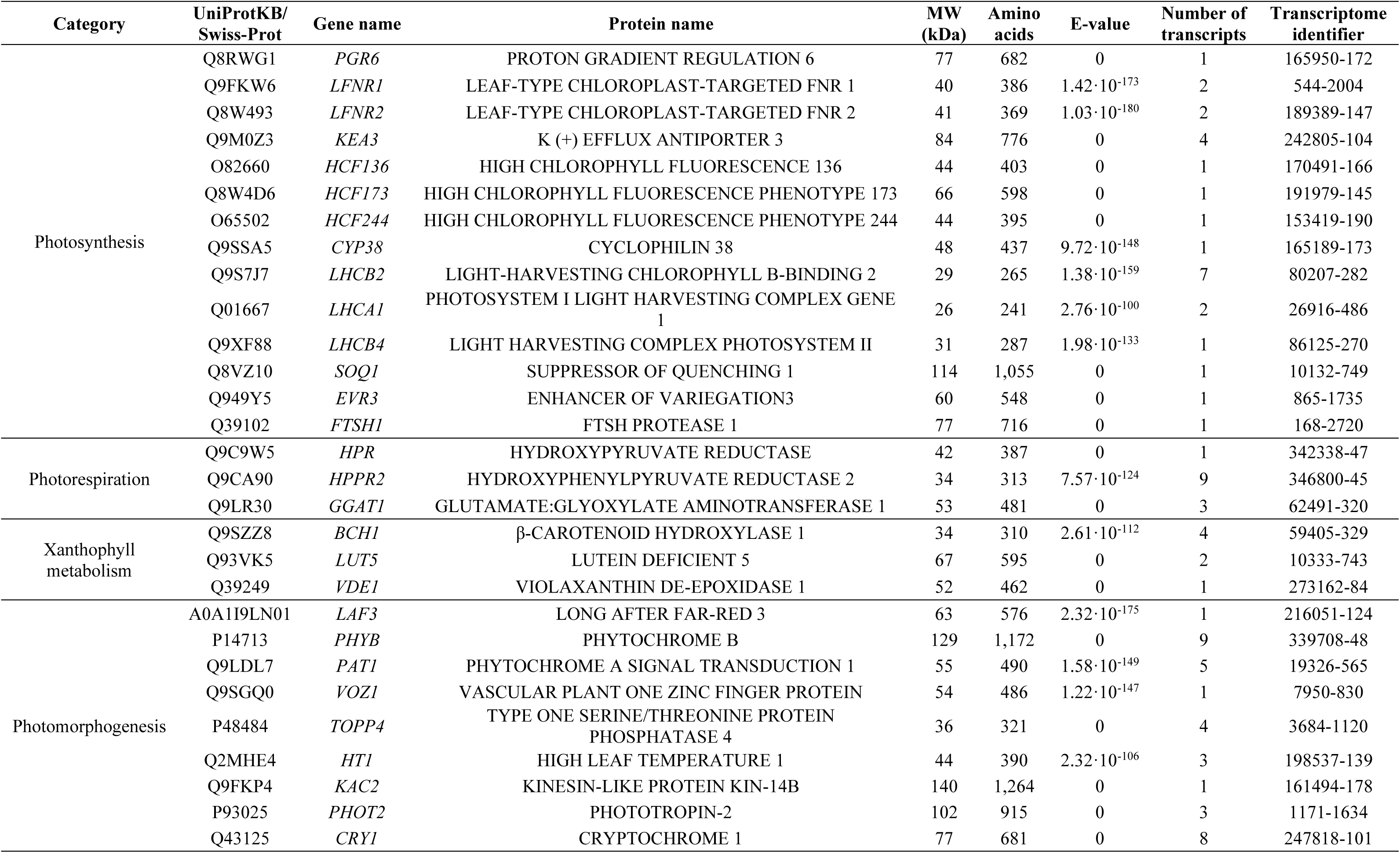
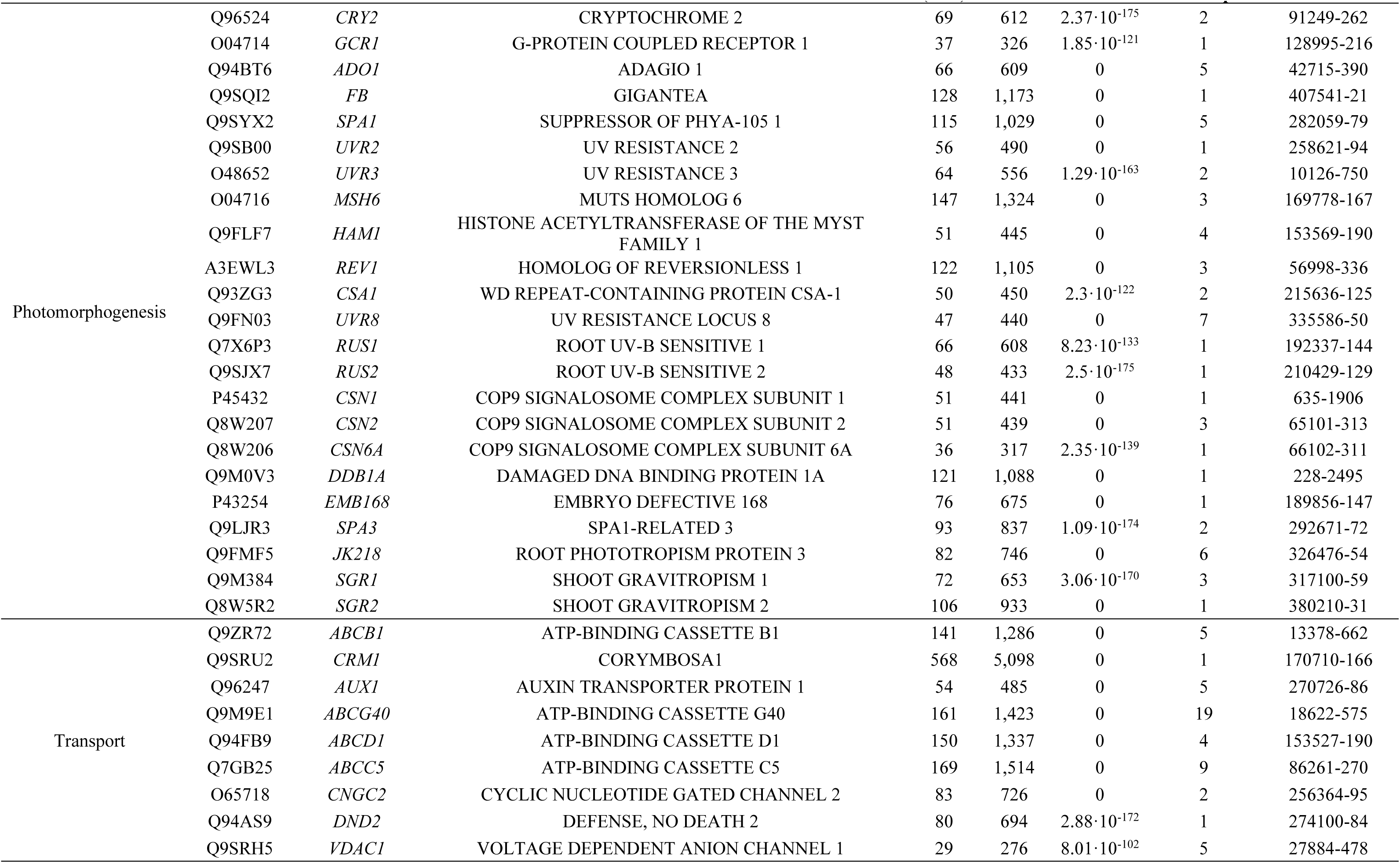
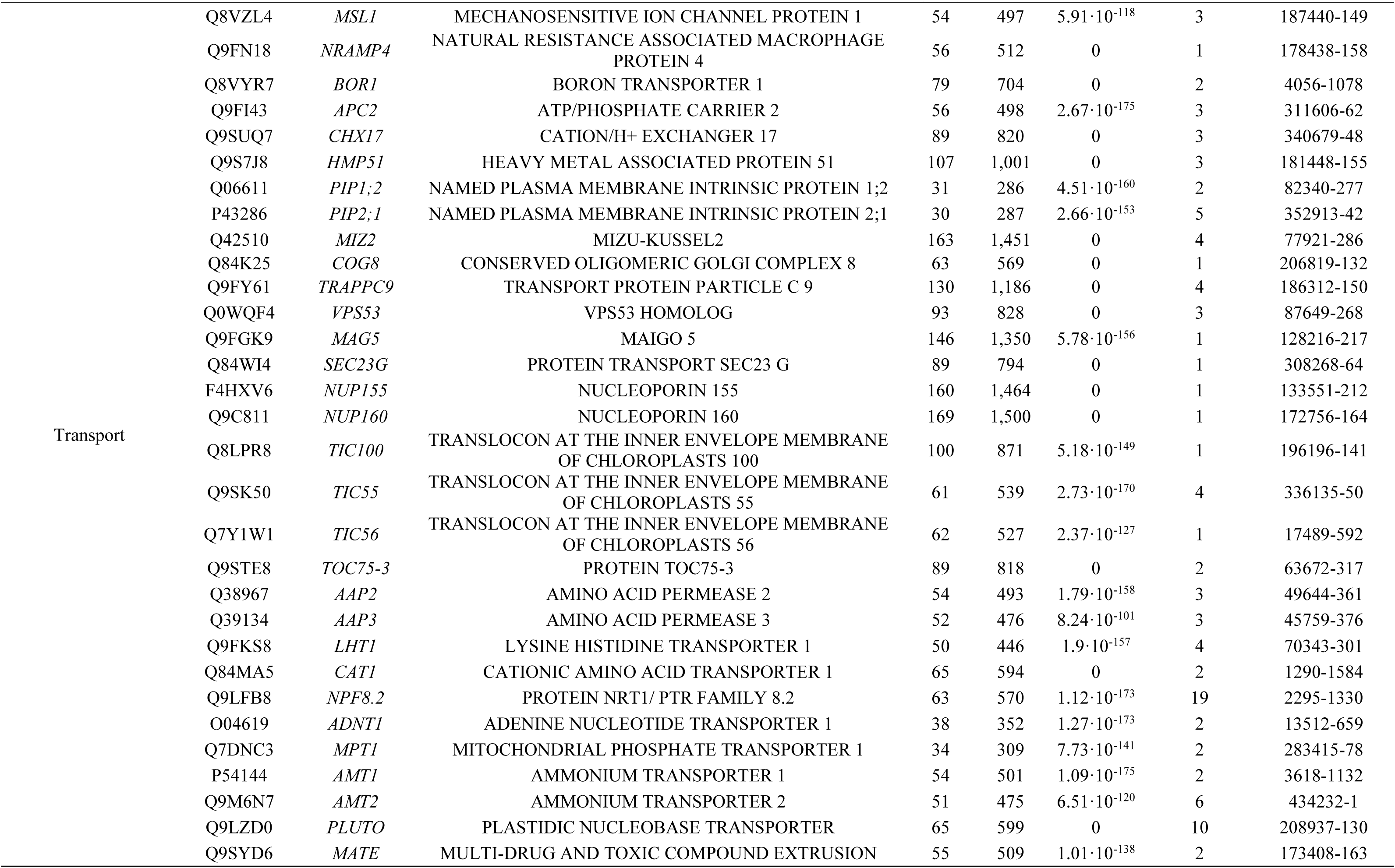
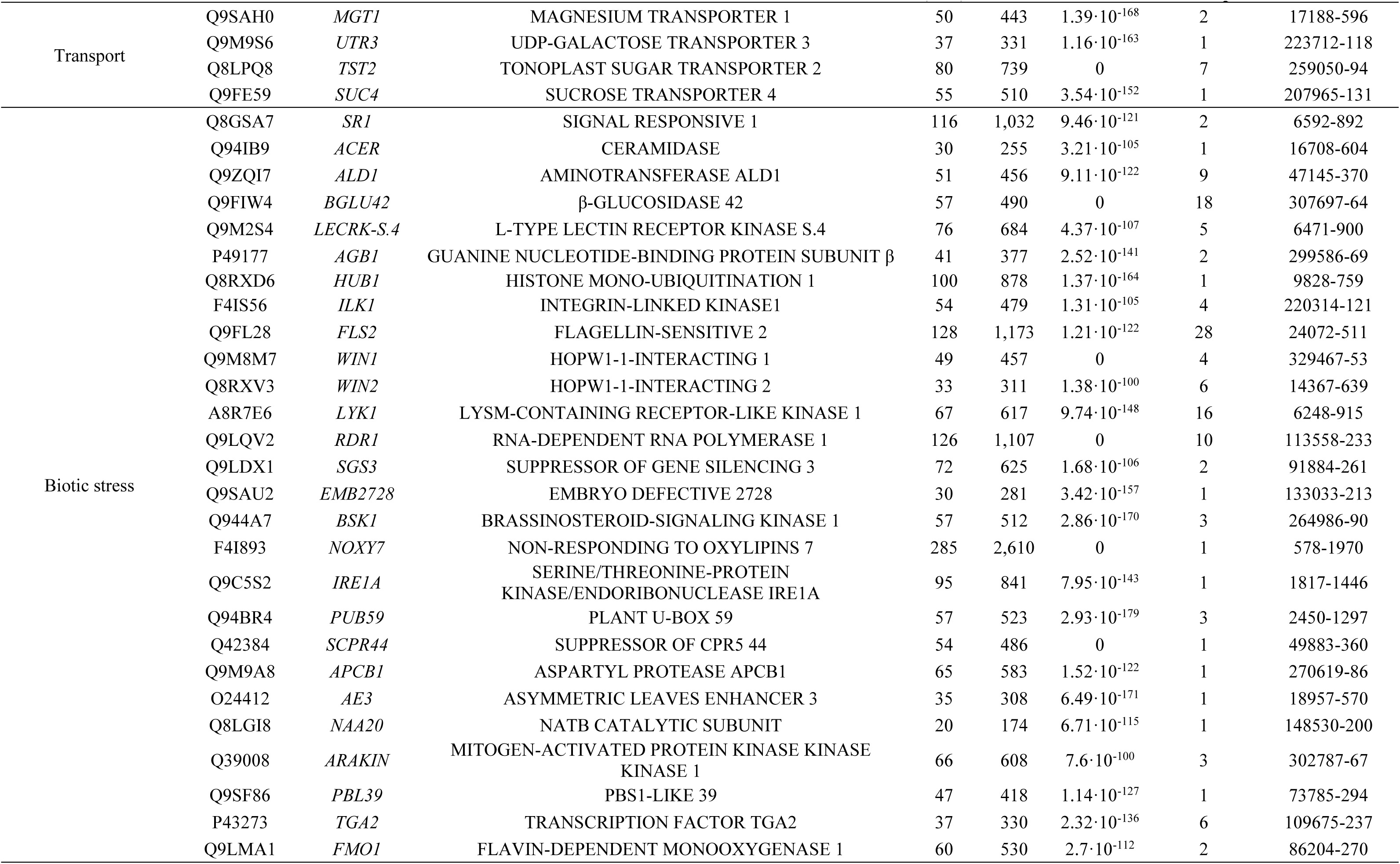
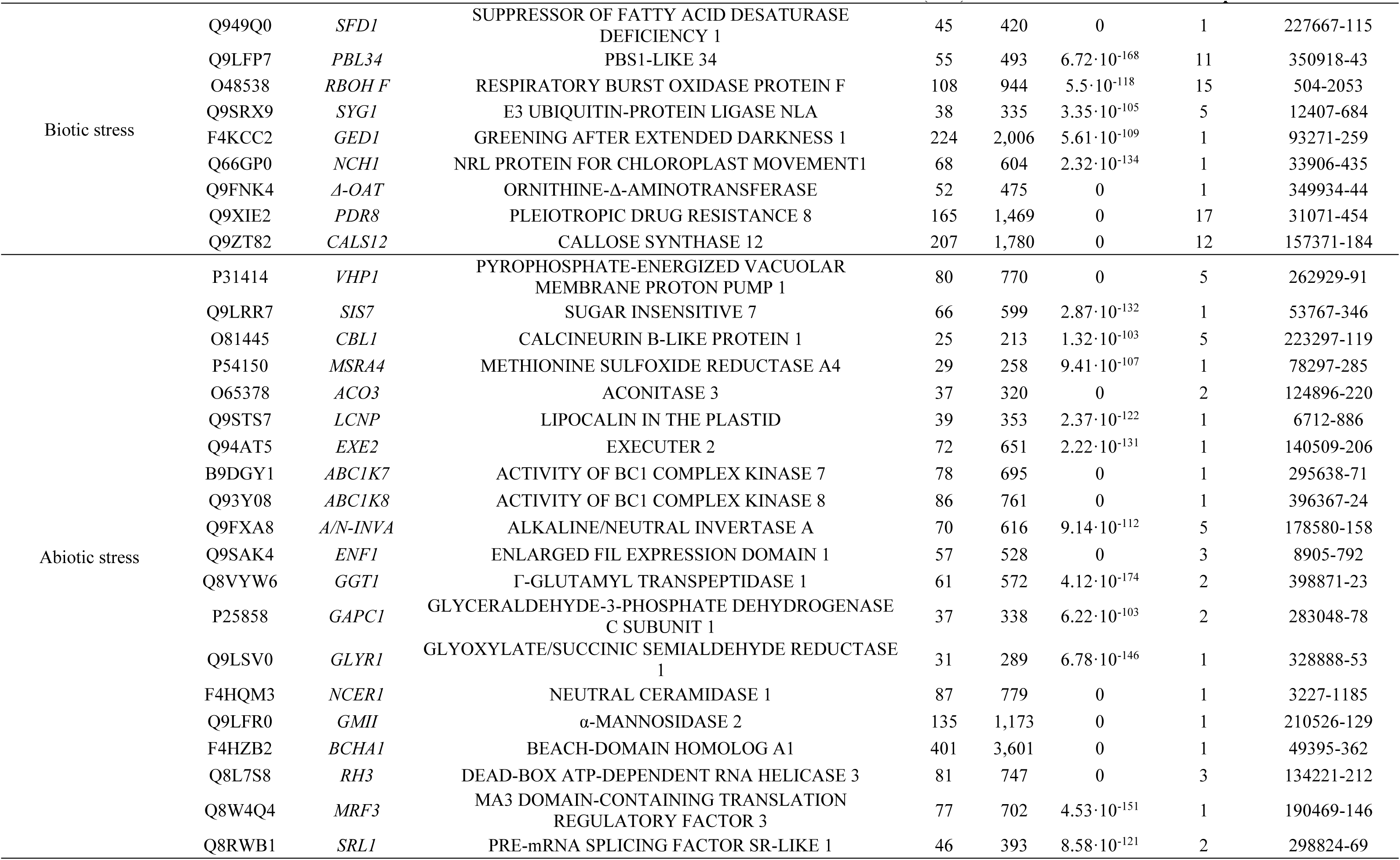
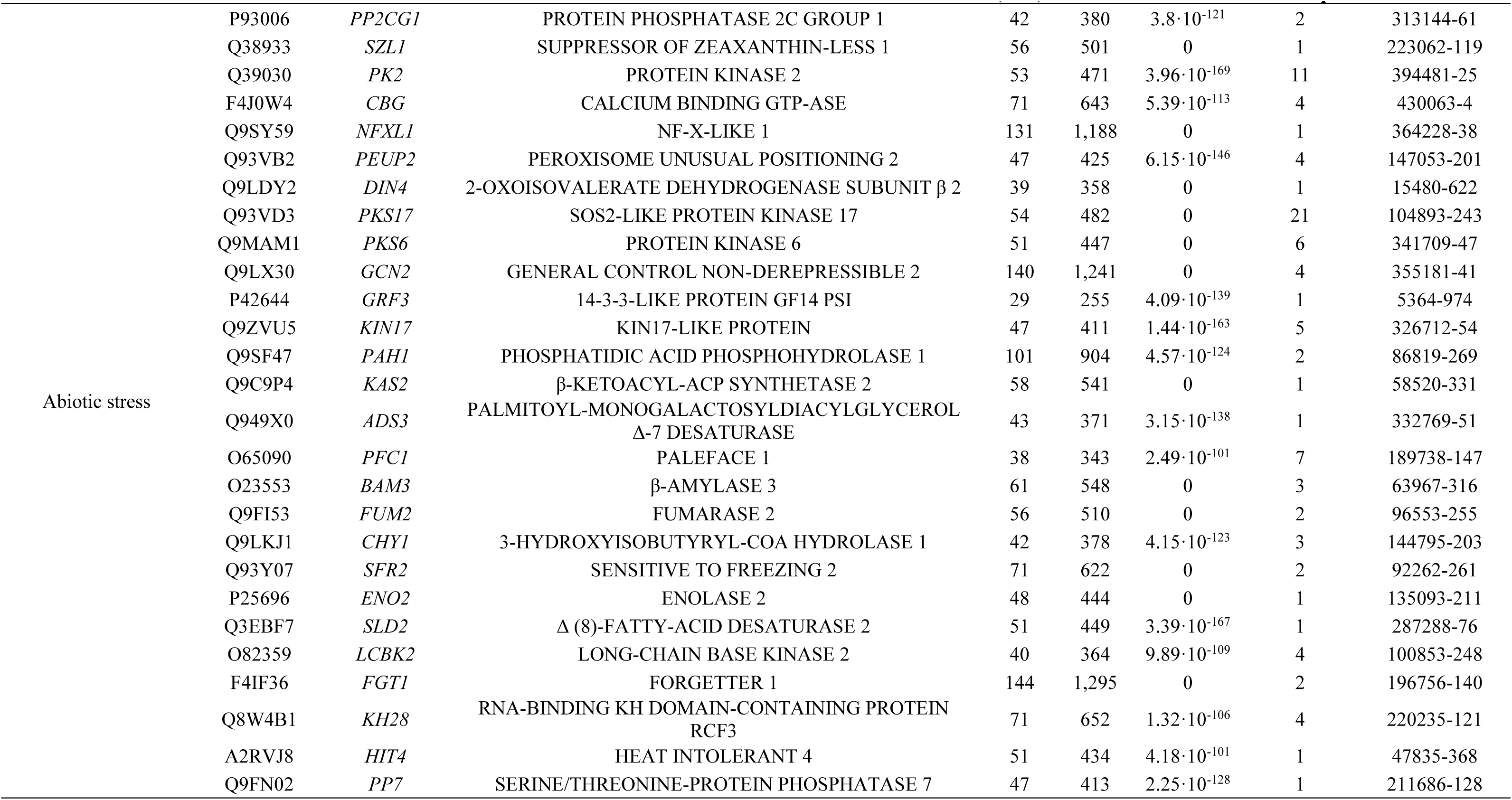
Selected proteins found in gametophytes of *Dryopteris affinis* associated with light, transport and stress.

### 3.5. Protein-protein interactions

The interactome of the 1,160 proteins studied associated with light, transport and stress was analysed using the String program. A network consisting of 1,160 nodes and 3,168 edges was obtained, which surpassed the expected number of edges (1,327). The Protein-Protein Interaction enrichment p-value was less than 1·10^-16^. A comparison between some types of interactions (neighbourhood, gene fusion, homology, co-occurrence and co-expression) in each group of proteins studied is represented in **Fig. 8**. Neighbourhood interactions peaked in photorespiration, whereas gene fusion in abiotic stress, especially heat. In homology, co-occurrence and co-expression interactions, the highest rate was found in those proteins associated to photosynthesis.

**Figure 8.**
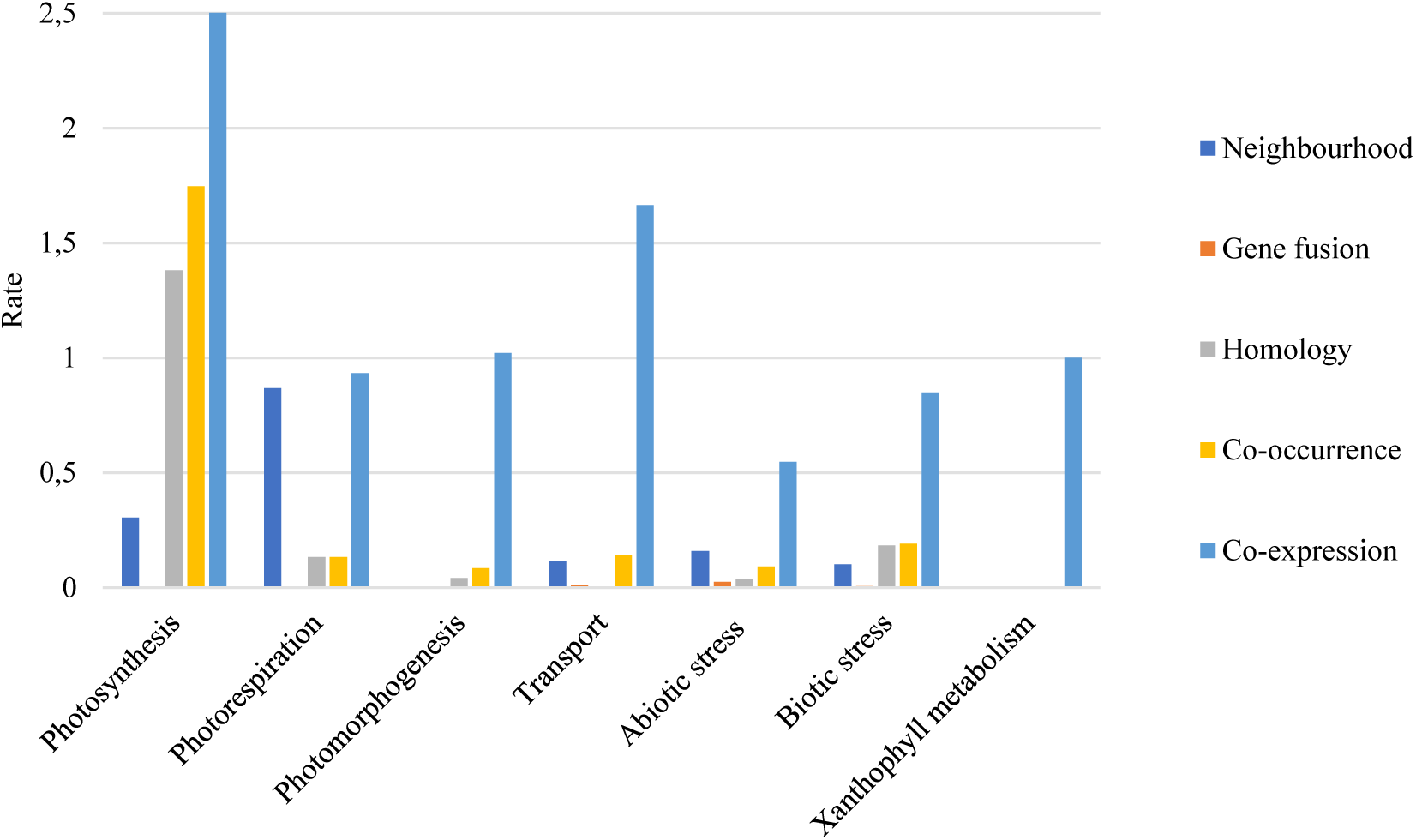
Estimation of the protein-protein interaction types in the selected groups of proteins extracted from the gametophytes of *Dryopteris affinis*.

To further investigate this initial analysis, the interactome was thoroughly examined. It revealed that 135 proteins had an interaction strength score above 999, where 1,000 represents the maximum possible score and 0 the minimum. Some examples of proteins with the strongest interactions were PHOTOSYSTEM I LIGHT HARVESTING COMPLEX GENE 2 (LHCA2), SUPPRESSOR OF PHYA-105 1 (SPA1), and EMBRYO DEFECTIVE 2817 (EMB2817). Looking at the type of interaction in each group of proteins, the analysis revealed that the strongest interaction in photosynthesis came from experiments between the proteins PHOTOSYSTEM I LIGHT HARVESTING COMPLEX GENE 1 (LHCA1) and CHLOROPHYLL A-B BINDING PROTEIN 4 (CAB4); in photorespiration, from databases, between GLUTAMATE: GLYOXYLATE AMINOTRANSFERASE 1 (GGAT1) and GLYCERATE DEHYDROGENASE HPR (HPR1); in the response to UV, from experiments, between HOMOLOG OF REVERSIONLESS 1 (REV1) and 3 (REV3); in xanthophyll metabolism, from databases, between LUTEIN DEFICIENT 5 (LUT5), VIOLAXANTHIN DE-EPOXIDASE 1 (VDE1), and β-CAROTENOID HYDROXYLASE 1 (BCH1) with each other; in the case of transport, from text-mining, with the proteins EXOCYST COMPLEX COMPONENT SEC10 (SEC10) and SUBUNIT OF EXOCYST COMPLEX 8 (SEC8); in the response to biotic stress, from text-mining, with REGULATOR OF NONSENSE TRANSCRIPTS 1 HOMOLOG (UPF1) and UPF2 (UPF2); in the response to water deprivation, salt stress and starvation, from text-mining, between SOS3-LIKE CALCIUM BINDING PROTEIN 5 (SCABP5) and SOS2-LIKE PROTEIN KINASE 17 (PKS17); in the response to oxidative stress, from experiments with ELONGATOR COMPLEX PROTEIN 1 (ELP1) and 2 (ELP2); and finally, in the response to cold, the strongest interaction came from experiments between 3-KETOACYL-COA SYNTHASE 6 (CER6) and 9 (KCS9), while in the response to heat, it was from text-mining, between the annotations KU70 HOMOLOG (KU70) and KU80 HOMOLOG (KU80). Specifically, protein-protein interactions due to co-expression relationships seem to be more frequent in photosynthesis. Additionally, several cases of gene fusion were observed among several proteins associated with heat stress, which are red coloured (**Fig. 9a**). Finally, proteins with the highest values of co-occurrence interactions are shown (**Fig. 9b**).

**Figure 9.**
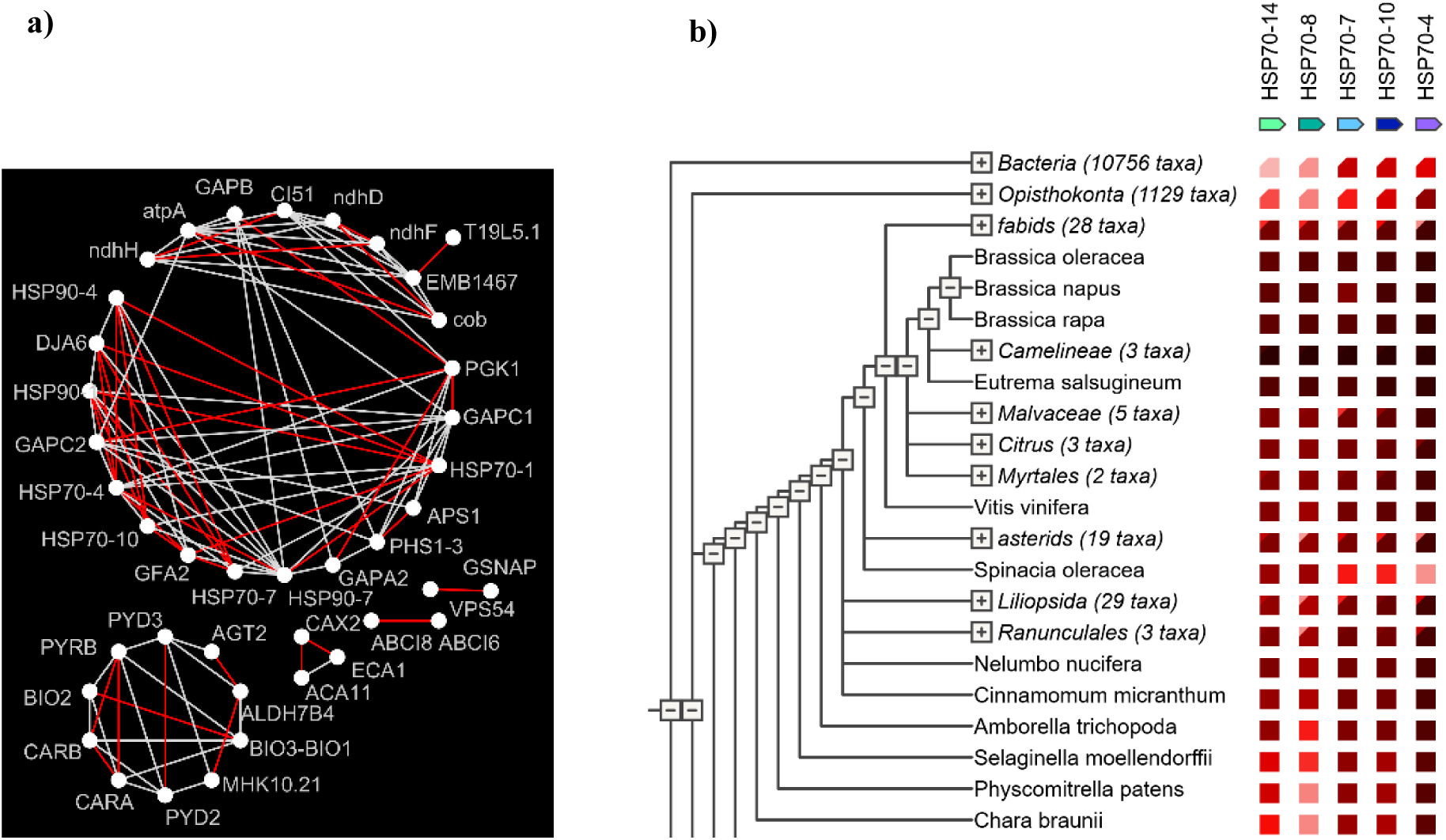
Protein–protein interaction network of a selected group of proteins from the *Dryopteris affinis* gametophyte, obtained using the STRING program: **a)** co-expression interaction in photosynthesis; **b)** gene fusion in response to heat.

As for the number of nodes, the proteins that had the most were MODIFIED TRANSPORT TO THE VACUOLE 17 (MTV17) with 19, CONSERVED OLIGOMERIC GOLGI COMPLEX 6 (COG6) with 18, and EXOCYST COMPLEX COMPONENT SEC6 (SEC6) with 16.

### 3.6. Comparison with fern genomes

As explained above, the *D. affinis* genome is not yet sequenced, so the transcriptome dataset was compared with other species for information. Some BLASTX searches were performed with the transcripts of around 180 proteins discussed here, associated with light, transport and stress against the fern species whose genomes are sequenced and deposited on the FernBase site (https://fernbase.org), showing high homology. The identification code of the proteins and, in some cases, their name, are provided in **Supplementary Table 2**.

## 4. Discussion

Despite the peculiar genetic traits of ferns, characterised by a high number of chromosomes and a large genome size (Manton 1950; Barker and Wolf 2010; Sessa and Der 2016), advancements in next-generation sequencing techniques have enabled researchers to explore these non-model plants. This has generated valuable insights applicable across various fields, including biology, biotechnology, and medicine (Jiang et al. 2024). In this study, the transcriptome of the gametophyte of *D. affinis* was meticulously examined through RNA sequencing, contributing to a broader understanding of this fern species (Grossmann et al. 2017; Wyder et al. 2020; Fernández et al. 2021; Ojosnegros et al. 2022, 2023, 2024). Following a comparison of the fern transcripts with *A. thaliana* sequences using Geneious Prime software, proteins with high homology related to light, transport and stress were analysed using the String program and the Uniprot database. Approximately 180 of these proteins were selected for discussion in this section, focusing on their roles and presence in the gametophyte of *D. affinis*, as well as their interactions. The selection of homologous proteins was guided solely by the availability of evidence for a potential role in genome regulation—for example as transcription factors or signalling molecules—rather than by their well-known roles in catalysing fundamental metabolic reactions. To facilitate a better understanding and analysis, they were categorised into the following groups: light (photosynthesis, photorespiration, xanthophyll metabolism, and photomorphogenesis), transport, and stress (both biotic and abiotic).

### 4.1. Light

#### 4.1.1. Photosynthesis, photorespiration, and xanthophyll metabolism

Photosynthesis is a complex process, and because of its vital role in plant life, it is one of the most studied processes in plant physiology. However, until recently, data on land plants other than spermatophytes, —i.e., bryophytes, lycophytes, and ferns— have been limited (Tosens et al. 2016; Carriqui et al. 2019; Gago et al. 2019; Roig-Oliver et al. 2025). Ferns offer an opportunity to examine photosynthesis in two generations of life cycle, sporophyte and gametophyte, which are genetically similar but morphologically distinct (Hagar and Freeberg 1980). Both generations depend on photosynthesis for their growth. While the sporophyte has developed large fronds with stomata and intercellular air spaces, the gametophyte is composed of a single cell layer across most of its surface, lacks stomata and vascular tissues, and forms a rudimentary cuticle. Thus, it is more exposed to atmospheric exchanges or variations (Hagar and Freeberg 1980; Watkins and Cardelús 2012). All this suggests an apparent vulnerability of the gametophyte and highlights a significant gap in understanding the physiological bases that directly impact this stage (Krieg and Chambers 2022).

Efforts are being made to investigate the physiological mechanisms by which the gametophyte of ferns copes with light, revealing strong similarities to those used by the sporophyte of ferns and seed plants (Farrar et al. 2008; Fernández-Marín et al. 2012). This work contributes to increasing the list of homologs involved in photosynthesis in the gametophyte of *D. affinis* (Ojosnegros et al. 2023). Our results point out how well the gametophyte of ferns is provided with proteins involved in securing the good course of the photosynthetic process. Among the new proteins observed, there are PROTON GRADIENT REGULATION 6 (PGR6), a chloroplast kinase that ensures the correct functioning of the electron transport chain by controlling the plastoquinone, β-carotene and xanthophyll lutein homeostasis (Martinis et al. 2014); two LEAF-TYPE CHLOROPLAST-TARGETED FNR 1 and 2 (LFNR1 and 2), which regulate the electron flow to meet the demands of the plant for ATP and reducing power (Lintala et al. 2007); and K (+) EFFLUX ANTIPORTER 3 (KEA3), which promotes the photosynthesis when the chloroplast ATP synthase activity is low, by reducing the pH gradient across the thylakoid membrane. The gametophyte of *D. affinis* accounts for some homologs associated with the photosystem II formation, assembly and stabilisation, including HIGH CHLOROPHYLL FLUORESCENCE 136 (HCF136), two HIGH CHLOROPHYLL FLUORESCENCE PHENOTYPE, 173 and 244 (HCF173 and 244), and CYCLOPHILIN 38 (CYP38). In line with it, the capture and transference of excitation energy to photosystems was well represented by structural subunits belonging to the light-harvesting complex, such as LIGHT-HARVESTING CHLOROPHYLL B-BINDING 2 (LHCB2), PHOTOSYSTEM I LIGHT HARVESTING COMPLEX GENE 1 (LHCA1), and LIGHT HARVESTING COMPLEX PHOTOSYSTEM II (LHCB4). The appropriate function of photosynthesis in the gametophyte could be favoured also by the presence of other relevant homologs such as SUPPRESSOR OF QUENCHING 1 (SOQ1), necessary to maintain the efficiency of the light harvesting complex (Brooks et al. 2013); and ENHANCER OF VARIEGATION3 (EVR3) and FTSH PROTEASE 1 (FTSH1), both involved in the proper formation of chloroplasts (Lindahl et al. 2000; Li et al. 2012).

Photorespiration, another key metabolic process in plants, remains relatively underexplored in ferns; it involves the light-dependent uptake of molecular oxygen and the release of carbon dioxide from organic compounds, thereby resembling respiration and counteracting photosynthesis (Peterhänsel et al. 2010). Previous results with the gametophyte of *D. affinis* revealed numerous homologs of proteins related to photorespiration (Ojosnegros et al. 2023). In the present study, we identify additional homologs, including HYDROXYPYRUVATE REDUCTASE (HPR) and HYDROXYPHENYLPYRUVATE REDUCTASE 2 (HPPR2), both of which participate in the photorespiratory core cycle. The first catalyses in *A. thaliana* the NADH-dependent reduction of hydroxypyruvate into glycerate (Cousins et al. 2011), while the second does the same in the NADPH-dependent reduction of glyoxylate and hydroxypyruvate into glycolate and glycerate, respectively (Timm et al. 2011). A homolog that intervenes in the amino acid content during photorespiration, GLUTAMATE: GLYOXYLATE AMINOTRANSFERASE 1 (GGAT1), was also found.

To close this first section, we refer to the xanthophylls, which are indispensable molecules in ferns that protect against light and oxidative stress, and contribute to photosynthesis at wavelengths where chlorophyll is ineffective (Latowski et al. 2013; Van Wittenberghe et al. 2021). Although our studied fern shows a great affinity for shaded sites (Salachna and Piechocki 2021), it seems to be well-equipped to cope with an excess of light. At this point, many homologs were detected in the *D. affinis* gametophyte transcriptome implicated in xanthophyll biosynthesis, such as β-CAROTENOID HYDROXYLASE 1 (BCH1) and LUTEIN DEFICIENT 5 (LUT5). We also identified the homolog of VIOLAXANTHIN DE-EPOXIDASE 1 (VDE1), which participates in the xanthophyll cycle by regulating zeaxanthin concentration in chloroplasts (Hieber et al. 2002). Fernández-Marín et al. (2021) discovered that this last protein is activated in wintergreen ferns under freezing and dark conditions, functioning as an adaptation and survival mechanism to stress. The proteins identified here add to those previously reported (Ojosnegros et al. 2022), where ZEAXANTHIN EPOXIDASE (ZEP), an enzyme of the xanthophyll cycle that converts zeaxanthin into antheraxanthin and subsequently into violaxanthin, was detected in the gametophytes of this fern.

Our results reinforce the central role of photosynthesis and photomorphogenesis in the gametophyte of ferns. The cnetplots provide a clear view of how multiple genes converge on fundamental mechanisms that sustain plant growth and development in this still underexplored phenotype.

#### 4.1.2. Photomorphogenesis

Fern gametophytes are very sensitive to light throughout their development (Wada 2007). The gametophyte of *D. affinis* is known to contain numerous proteins associated with photomorphogenesis (Fernández et al. 2021; Ojosnegros et al. 2022, 2024). In the present work, more proteins have been annotated, such as the *A. thaliana* homologs of LONG AFTER FAR-RED 3 (LAF3), implicated in the response to continuous far-red light, and the photoreceptor PHYTOCHROME B (PHYB), which triggers different morphogenetic responses depending on whether it is impacted by red or far-red light (Salome et al. 2002). In spores of the fern *A. capillus-veneris,* exposure to a short pulse of red light induces germination, whereas far-red light inhibits this response, being a reversible process, which indicates a phytochrome dependence (Wada 2007). To date, no studies have been conducted on phytochrome in *D. affinis*, but it is likely to have some similar activity. The idea is reinforced by the findings of the homologs of PHYTOCHROME A SIGNAL TRANSDUCTION 1 (PAT1), and VASCULAR PLANT ONE ZINC FINGER PROTEIN (VOZ1), both transcription factors key in the signal transduction in the phytochromes A and B, respectively, and also the homologue of TYPE ONE SERINE/THREONINE PROTEIN PHOSPHATASE 4 (TOPP4), implicated in the regulation of phytochromes, and whose function is the control of phytochrome B (Yue et al. 2016). On the other hand, it is striking that a homolog of HIGH LEAF TEMPERATURE 1 (HT1), a serine/threonine/tyrosine kinase required for red light-induced stomatal opening, was annotated in the gametophyte of *D. affinis,* which lacks stomata (present only in the sporophyte). The detection of stomata related proteins in the gametophyte of a fern is an intriguing finding, since extant fern gametophytes are generally reported to lack functional stomata (Krieg and Chambers 2022). Rather than reflecting a direct role in stomatal regulation, the expression of these proteins may indicate the persistence of conserved genetic pathways inherited from the sporophyte generation, or their recruitment into alternative physiological functions such ion transport, stress signalling, or cellular differentiation. Fossil evidence (Clark et al. 2022) shows that the common ancestor of land plants, prior to the divergence of bryophytes and tracheophytes did bear stomata, suggesting that part of the underlying molecular machinery has been retained through evolution, even though stomata are no longer present in the gametophytic phase of living ferns. In line with it, our findings highlight the evolutionary continuity of regulatory networks across generations, despite morphological differences.

Likewise, *A. thaliana* homologs related to radiation in the 400 – 500 nm range (i.e. blue light) were also analysed. Our results highlight homologs of KINESIN-LIKE PROTEIN KIN-14B (KAC2) and PHOTOTROPIN-2 (PHOT2), both required for the chloroplast avoidance response under high-intensity blue light in *A. thaliana* (Suetsugu et al. 2016). This movement of the chloroplast, which is towards the light when it is weak to absorb light, and away when it is strong to avoid chloroplast photodamage, contributes to achieving a higher photosynthetic efficiency, key to plant survival (Wada 2013). Some studies (Kagawa et al. 2004; Wada 2007) have been carried out with PHOT2 in *A. capillus-veneris* gametophytes, to try to understand the effect that blue light has on their chloroplasts. These studies demonstrate that the phototropin family of blue-light photoreceptors, to which PHOT2 belong, is necessary for the response to blue light in ferns. Associated with this nature of light, the gametophyte of *D. affinis* is provided with other interesting proteins, such as the photoreceptors CRYPTOCHROME 1 (CRY1) and 2 (CRY2), G-PROTEIN COUPLED RECEPTOR 1 (GCR1), implicated in its signalling; and ADAGIO 1 (ADO1), a member of an E3 ubiquitin ligase complex that in *A. thaliana* participates in blue light-dependent circadian cycles (Takase et al. 2011). CRY1 and CRY2 were also detected in gametophytes and sporophytes of *A. capillus-veneris* (Imaizumi et al. 2000), and frond of *Asplenium yunnanense* (Zhou et al. 2013). These cryptochromes present the same DAS motif in ferns as in angiosperms. Recently, it has been reported that CRY1 originated from a common ancestor of seed plants, whereas CRY2 originated from a common ancestor of land plants (Lu et al. 2025). Additionally, the homologs GIGANTEA (FB) and SUPPRESSOR OF PHYA-105 1 (SPA1), found in this research, had been reported controlling circadian rhythms in *Arabidopsis* (Huq et al. 2000; Ishikawa et al. 2006). Undoubtedly, our results contribute to emphasising the importance of continued research into photomorphogenesis in non-flowering plants, which may reveal key insights such as the adaptive evolution of timekeeping mechanisms in the plant kingdom.

Fern gametophytes are particularly vulnerable to UV radiation because they lack a thick cuticle. To protect itself from this wavelength, the gametophyte of *D. affinis* harbours homologs of proteins that repair damaged DNA after exposure: UV RESISTANCE 2 (UVR2) and 3 (UVR3), MUTS HOMOLOG 6 (MSH6), and HISTONE ACETYLTRANSFERASE OF THE MYST FAMILY 1 (HAM1). We also identified other interesting proteins related to the response to UV-B light, such as HOMOLOG OF REVERSIONLESS 1 (REV1), WD REPEAT-CONTAINING PROTEIN CSA-1 (CSA1), and UV RESISTANCE LOCUS 8 (UVR8), being the last one a photoreceptor well recognised (Davey et al. 2012). Moreover, ROOT UV-B SENSITIVE 1 (RUS1) and 2 (RUS2), focused on protection against UV-B on roots, were also annotated in this work. It is important that plants defend against this radiation because it alters primary and secondary metabolism (Sánchez Correa et al. 2023). As outlined above, light is essential for the life of ferns; however, it must be within an appropriate intensity range, as excessive solar radiation can damage tissues and lead to photoinhibition and pigment photo-destruction (Takahashi and Murata 2008).

Plant photomorphogenesis involves many signalling molecular structures, such as the COP9 signalosome complex. As it was mentioned in the Results section, several members of this complex were annotated, including COP9 SIGNALOSOME COMPLEX SUBUNIT 1 (CSN1), 2 (CSN2), and 6A (CSN6A), involved in repression of photomorphogenesis in darkness by regulating the activity of the signalosome complex. In addition, with similar functions, we identified the homologs of DAMAGED DNA BINDING PROTEIN 1A (DDB1A), a component of the light signal transduction machinery, and EMBRYO DEFECTIVE 168 (EMB168), an E3 ubiquitin-protein ligase. By contrast, SPA1-RELATED 3 (SPA3), which is implicated in the repression of photomorphogenesis in light (Laubinger et al. 2006), was also detected in the transcriptome.

An important aspect of the plant response to light is phototropism, the directional growth towards the light. It is surprising that the gametophyte of *D. affinis* appears to contain the homologue of ROOT PHOTOTROPISM PROTEIN 3 (JK218), also named NON-PHOTOTROPIC HYPOCOTYL 3 (NPH3), a protein which has been considered specific to seed plants. It may function as a scaffold protein that assembles the enzymatic components of an NPH1-activated phosphorelay and is therefore involved in the signal transduction of phototropic responses (Motchoulski and Liscum, 1999; de Carbonnel et al. 2010). Fern growth is also regulated by gravitropism, in such a way that gravity modulates its direction: upward in the stems (negative gravitropism) and downward in the roots (positive gravitropism). In our protein set, we detected homologs of SHOOT GRAVITROPISM 1 (SGR1) and 2 (SGR2), which in *A. thaliana* are involved in amyloplast sedimentation in the endodermis during shoot gravitropism, acting as statoliths (Saito et al. 2011; Cruz-Ramirez et al. 2013). Amyloplasts are observed in the root cap cells and the cells above the root apex in ferns, but they are accumulated in the cells at the root apex only in seed plants (Zhang et al. 2019). Thus, although amyloplast sedimentation is a characteristic linked exclusively to seed plants, becoming a gravity-sensing mechanism new in the evolution, we think that the existence of these proteins in *D. affinis* gametophytes probably means a related function. The findings of SGR1 and SGR2 complement those obtained in previous works (Fernández et al. 2021; Ojosnegros et al. 2022).

### 4.2. Transport

Like angiosperms and gymnosperms, ferns require transporting proteins, as transport is indispensable for life. These proteins are involved in moving hormones, ions, or other molecules throughout the plant, in cellular endocytosis and exocytosis, and in the import and export of proteins and nucleic acids in the nucleus, mitochondria, and chloroplasts. However, the studies in ferns devoted to increase our knowledge, and specifically in the gametophyte generation, are scarce. What we know about transport in the gametophyte of ferns? In the first place, early ultrastructural studies demonstrated that primary plasmodesmata are abundant in fern gametophytes and that their frequency and distribution change during development, thereby regulating the intercellular movement of solutes and signalling molecules (Tilney et al. 1990; Bartz and Gola 2018). On the other hand, rhizoids function as absorptive structures, facilitating nutrient uptake and ion transport into the thallus, a process comparable to the absorptive roles of rhizoids in bryophytes (Racusen 2002). Regarding transporter proteins, aquaporins have been identified in fern gametophytes, suggesting their involvement in water and solute transport. A study in the gametophyte of the xerophytic fern *Cheilanthes lanosa* suggested that aquaporins play a significant role in water balance and desiccation tolerance, highlighting their importance in the physiological resilience of fern gametophytes (Diamond et al. 2012). Additionally, research on *Pteris vittata* gametophytes has identified a vacuolar arsenite transporter (*PvACR3*), which is crucial for arsenic detoxification, underscoring the role of transporter proteins in metal homeostasis and stress responses. Recent proteomic analyses in fern gametophytes of *Diplazium maximum* reported some proteins related to transport and trafficking associated to environmental adaptation (Sareen et al. 2019). From another perspective, transport in the gametophyte is intimate linked to plant regulators. For instance, auxin transport within the gametophyte has been reported to be a crucial aspect of its development, coordinating cell differentiation and the transition to the sporophyte stage, suggesting a role to be involved in meristem regeneration (Whitter et al. 2023).

After this brief overview of the relevance of transport in the fern gametophyte, we now turn to a discussion of some of the key advances contributed by this research. In this new study, nearly 700 homologous protein annotations were associated with transport processes in the gametophyte of a fern, highlighting the extensive repertoire of transport related proteins in this developmental stage. To begin our analyses, we focussed on hormone-related transporters in the fern gametophyte, uncovering key proteins that may mediate phytohormone distribution and signalling during its development. Nucleotide sequences reflect homologies with proteins implicated in auxin transport, such as the *A. thaliana* homologs of ATP-BINDING CASSETTE B1 (ABCB1) and CORYMBOSA1 (CRM1), which regulate polar auxin transport (Geisler et al. 2005; Contreras-Cornejo et al. 2009), and AUXIN TRANSPORTER PROTEIN 1 (AUX1), which participates in acropetal and basipetal auxin transport (Swarup et al. 2004). Other homologs are related to abscisic acid transport, such as ATP-BINDING CASSETTE G40 (ABCG40). As it was commented in the Results section, numerous members of the ATP-binding cassette (ABC) family were noticed here, demonstrating the great role that they have in transport in the organism. In addition to the functions mentioned, homologs from this family involved in fatty acid transport, such as ATP-BINDING CASSETTE D1 (ABCD1), and ion transport, such as C5 (ABCC5), were found, the latter regulating K^+^ and Na^+^ levels in the cell (Nagy et al. 2009).

Continuing with ion transport, its transfer through membrane channels accounts for almost 30% of the energy consumed by a plant cell, underpinning all aspects of its biology, from mineral nutrition, cell expansion, and organ development to pathogen defence and senescence (Blatt 2024). In the gametophyte transcriptome, we detected several *A. thaliana* homologs of ion channels, such as CYCLIC NUCLEOTIDE GATED CHANNEL 2 (CNGC2) and DEFENSE, NO DEATH 2 (DND2). The former is permeable to potassium and calcium ions, while the latter is permeable to potassium and sodium ions (Koehler et al. 2001; Balague et al. 2003). Another ion channel reported was VOLTAGE DEPENDENT ANION CHANNEL 1 (VDAC1), which allows the passage of small hydrophilic molecules across the mitochondrial outer membrane (Tateda et al. 2011). Apart from these findings, in the *D. affinis* gametophytes, various mechanosensitive ion channels were also noticed. Among others, we highlight the homolog of MECHANOSENSITIVE ION CHANNEL PROTEIN 1 (MSL1), which opens in response to stretch forces in the membrane lipid bilayer. NATURAL RESISTANCE ASSOCIATED MACROPHAGE PROTEIN 4 (NRAMP4), a vacuolar transporter of iron, manganese, and cadmium required for intracellular metal homeostasis (Lanquar et al. 2010); BORON TRANSPORTER 1 (BOR1), which transports boron from roots to shoots; ATP/PHOSPHATE CARRIER 2 (APC2), a calcium-dependent protein that catalyses ATP import co-transported with divalent metal cations across the mitochondrial inner membrane in exchange of phosphate (Monne et al. 2017); CATION/H+ EXCHANGER 17 (CHX17), a K^+^/H^+^ antiporter regulating K^+^ uptake and homeostasis; and HEAVY METAL ASSOCIATED PROTEIN 51 (HMP51), which participates in copper import into the cell, were other proteins identified. We must also point out the aquaporins, i.e., channels in the cell membrane that facilitate water permeability, such as NAMED PLASMA MEMBRANE INTRINSIC PROTEIN 1;2 (PIP1;2) and 2;1 (PIP2;1).

As for endocytosis and exocytosis, they are crucial processes in cell functioning, transporting material in and out of the cell, respectively, and fulfilling their antagonistic roles with spatial-temporal coordination. In *D. affinis* gametophytes, representative examples of *A. thaliana* homologs associated with these processes include: MIZU-KUSSEL2 (MIZ2), which participates in endocytosis; CONSERVED OLIGOMERIC GOLGI COMPLEX 8 (COG8), required for intra-Golgi protein trafficking; TRANSPORT PROTEIN PARTICLE C 9 (TRAPPC9), a subunit of the Trafficking Protein Particle Complex II necessary for the transport of proteins from Golgi to the growing cell plate during mitosis (Naramoto et al. 2014); and VPS53 HOMOLOG (VPS53), a component of the Golgi-Associated Retrograde Protein Complex, key in retrograde transport from early and late endosomes to the trans-Golgi network (Wang et al. 2011). Further homologs are MAIGO 5 (MAG5), essential in the protein transport from endoplasmic reticulum to Golgi; and PROTEIN TRANSPORT SEC23 G (SEC23G), a component of the Coat Protein Complex II, which intervenes in the formation of transport vesicles from the endoplasmic reticulum (De Craene et al. 2014). On the other hand, concerning nuclear import and export, several components of the nuclear pore complex were identified, such as the homologs of NUCLEOPORIN 155 (NUP155) and 160 (NUP160), both involved in the transfer of mature mRNA from the nucleus to the cytoplasm (Roth and Wiermer 2012). Examples of chloroplast transport proteins are TRANSLOCON AT THE INNER ENVELOPE MEMBRANE OF CHLOROPLASTS 100 (TIC100), 55 (TIC55), and 56 (TIC56), all responsible for the import of protein precursors into chloroplasts (Kikuchi et al. 2013); and PROTEIN TOC75-3 (TOC75-3), which regulates the insertion of proteins targeted to the outer membrane of chloroplasts.

Other proteins related to transport, such as amino acid-proton symporters, were reported. Some of them are the *A. thaliana* homologs of AMINO ACID PERMEASE 2 (AAP2), with specificity for histidine, arginine, glutamate and neutral amino acids, and also accepting large aromatic amino acids like phenylalanine and tyrosine (Fischer et al. 1995); AMINO ACID PERMEASE 3 (AAP3), with preference for γ-aminobutyric acid, tryptophan and neutral and basic amino acids (Fischer et al. 1995); and LYSINE HISTIDINE TRANSPORTER 1 (LHT1), which has a predilection for histidine, lysine, glutamic acid, alanine, serine, proline and glycine (Svennerstam et al. 2008). Other annotated proteins included CATIONIC AMINO ACID TRANSPORTER 1 (CAT1), which has affinity for cationic amino acids; and PROTEIN NRT1/PTR FAMILY 8.2 (NPF8.2), which transports di- and tripeptides.

More cellular transporters of ions and organic compounds, included in the list of proteins obtained from *D. affinis* gametophytes, are the followings homologs to those of *A. thaliana*: ADENINE NUCLEOTIDE TRANSPORTER 1 (ADNT1), which is an ATP/ADP/AMP exchanger in the inner mitochondrial membrane; MITOCHONDRIAL PHOSPHATE TRANSPORTER 1 (MPT1), which transports phosphate groups from the cytosol to the mitochondrial matrix; AMMONIUM TRANSPORTER 1 (AMT1), involved in ammonium uptake from the soil and transport to the shoots; AMMONIUM TRANSPORTER 2 (AMT2), which transports ammonium between the apoplast and symplast; and PLASTIDIC NUCLEOBASE TRANSPORTER (PLUTO), required for adenine, guanine and uracil transport. Other noteworthy homologs were MULTI-DRUG AND TOXIC COMPOUND EXTRUSION (MATE) and MAGNESIUM TRANSPORTER 1 (MGT1), which transport citrate and magnesium, respectively, and are both required for aluminium tolerance (Deng et al. 2006; Liu et al. 2012); UDP-GALACTOSE TRANSPORTER 3 (UTR3), essential for transporting UDP-glucose from the cytoplasm to the Golgi and endoplasmic reticulum; TONOPLAST SUGAR TRANSPORTER 2 (TST2), a sugar proton-coupled antiporter in the vacuole; and SUCROSE TRANSPORTER 4 (SUC4), which mediates sucrose import. All the information gathered here contributes to expanding our precedent publications about transport proteins (Ojosnegros et al. 2022, 2023, 2024).

To end up with this part, one of our concerns, which we are unable to fully explain, is the observation of elongated cells between the basal and apical regions during embryo development. It is tempting to speculate that these cells might intervene in the nutrition of the developing embryo, acting as mediators of resource allocation. We wonder whether such observations may be related with the foot set up to mediate nutrient transfer from gametophyte to embryo (Conway and Di Stilio 2020).

### 4.3. Stress

In addition to light and transport related proteins, we analysed stress associated proteins in the gametophyte of the fern *D. affinis,* providing new insights that complement and expand upon previously published data on stress responses in this developmental stage.

#### 4.3.1. Biotic stress

Several studies have shown that ferns have developed evolutionary innovations to protect themselves from insects, pathogens and other predators, and to avoid potential diseases (Fürstenberg-Hägg 2013; Goswani et al. 2016; Shukla et al. 2016). Concerning insects, ferns are rarely damaged by them, specifically 30 times less than angiosperms (Chen 2022). This may be due to the production of certain substances, such as ecdysones, which cause developmental abnormalities in insects or are toxic to them (Chen 2022). In the *D. affinis* gametophyte transcriptome, an example of a protein implicated in the response to insect attack was detected: the *A. thaliana* homolog of SIGNAL RESPONSIVE 1 (SR1). Ferns also exhibit effective mechanisms against bacteria, fungi and virus infections. Associated with the resistance against bacteria and fungi, several homologs were identified, such as CERAMIDASE (ACER), AMINOTRANSFERASE ALD1 (ALD1), β-GLUCOSIDASE 42 (BGLU42), L-TYPE LECTIN RECEPTOR KINASE S.4 (LECRK-S.4), GUANINE NUCLEOTIDE-BINDING PROTEIN SUBUNIT β (AGB1), HISTONE MONO-UBIQUITINATION 1 (HUB1), and INTEGRIN-LINKED KINASE1 (ILK1), the latter being involved in responses to bacteria-derived pathogen-associated molecular patterns (Brauer et al. 2016). Related to this, we identified FLAGELLIN-SENSITIVE 2 (FLS2), which constitutes the receptor that recognises the bacterium flagellin, an elicitor of the plant defence to pathogen-associated molecular patterns (Macho et al. 2014); and HOPW1-1-INTERACTING 1 (WIN1) and 2 (WIN2), which regulate bacterial defence. This was complemented by LYSM-CONTAINING RECEPTOR-LIKE KINASE 1 (LYK1), which, in addition to being involved in restricting bacterial growth, also functions in fungal defence (Shinya et al. 2014). Likewise, in the response to the virus, the *A. thaliana* homologs of RNA-DEPENDENT RNA POLYMERASE 1 (RDR1), which participates in antiviral silencing (Garcia-Ruiz et al. 2010), and SUPPRESSOR OF GENE SILENCING 3 (SGS3), needed for natural virus resistance (Liu et al. 2019), were annotated. In parallel, EMBRYO DEFECTIVE 2728 (EMB2728), which participates in the response to phytoparasitic nematodes, was reported. This information expands what is currently known about how *D. affinis* gametophytes may cope with biotic stress (Grossmann et al. 2017; Ojosnegros et al. 2022, 2024). Many of the biochemical compounds produced by ferns under threat have been used by humans since ancient times for medical purposes, providing health benefits. Today, modern approaches combine multidisciplinary technologies with identified extracts from plant parts, enabling the production of targeted medicines (Goswani et al. 2016). Moreover, some ferns generate unique secondary metabolites that have not yet been discovered in angiosperms or gymnosperms (Goswani et al. 2016). Climate change, together with the rapid evolution and mutation of phytopathogens, make it difficult to fully understand the resistance pathways of each fern species (Chen 2022). With this study, we advocate for advancing efforts to obtain high-quality fern genomes and their associated metabolomes, as they presumably hold great potential.

Plants, unlike mammals, lack an adaptive immune system of defence cells that move throughout the organism toward the infection site. Thus, plants rely on the innate immunity of individual cells and signals emanating from the infection zone to fight off biotic stress (Jones and Dangl 2006). In the list of proteins obtained from our studied fern, plenty of *A. thaliana* homologs related to plant immunity were detected, such as BRASSINOSTEROID-SIGNALING KINASE 1 (BSK1), NON-RESPONDING TO OXYLIPINS 7 (NOXY7), SERINE/THREONINE-PROTEIN KINASE/ENDORIBONUCLEASE IRE1A (IRE1A), PLANT U-BOX 59 (PUB59), SUPPRESSOR OF CPR5 44 (SCPR44), ASPARTYL PROTEASE APCB1 (APCB1), ASYMMETRIC LEAVES ENHANCER 3 (AE3), and NATB CATALYTIC SUBUNIT (NAA20), as well as some devoted to immune signalling: MITOGEN-ACTIVATED PROTEIN KINASE KINASE KINASE 1 (ARAKIN) and PBS1-LIKE 39 (PBL39). We also recorded proteins involved in induction, establishment and activation of the systemic acquired resistance, such as TRANSCRIPTION FACTOR TGA2 (TGA2), FLAVIN-DEPENDENT MONOOXYGENASE 1 (FMO1), and SUPPRESSOR OF FATTY ACID DESATURASE DEFICIENCY 1 (SFD1), respectively (Johnson et al. 2003; Koch et al. 2006; Lorenc-Kukula et al. 2012). Other examples are PBS1-LIKE 34 (PBL34), which participates in chitin-triggered immune signalling (Luo et al. 2020); and RESPIRATORY BURST OXIDASE PROTEIN F (RBOH F), which regulates the hypersensitive response (Desikan et al. 2006).

Similarly, other notable discoveries in our dataset were homologs of E3 UBIQUITIN-PROTEIN LIGASE NLA (SYG1), associated with general defence; GREENING AFTER EXTENDED DARKNESS 1 (GED1), which regulates the biosynthesis of defence-related metabolites (Vicente et al. 2019); NRL PROTEIN FOR CHLOROPLAST MOVEMENT1 (NCH1), linked to disease resistance; and ORNITHINE-Δ-AMINOTRANSFERASE (Δ-OAT), involved in non-host disease resistance. In addition, the gametophytes also contained the following proteins: PLEIOTROPIC DRUG RESISTANCE 8 (PDR8), known for its role in cell death as a defence mechanism (Johansson et al. 2014); and CALLOSE SYNTHASE 12 (CALS12), associated with callose formation following wounding (Jacobs et al. 2003). Callose, a polysaccharide deposited on plant cell walls, acts as a scaffold to facilitate the incorporation of other cell wall components (Galatis and Apostolakos 2010). Kalmbach and Helariutta (2019) reported that some ferns only produce callose when injured. It is unknown whether *D. affinis* exhibits this trait, but in *in vitro* culture, unintentional injuries during manipulation may have triggered callose synthesis, leading to the detection of its homolog in the transcriptome.

#### 4.3.2. Abiotic stress

As mentioned above, the two generations of the fern life cycle are morphologically distinct, which causes each to respond to environmental changes in markedly different ways. In the case of the response to dehydration, the sporophyte has waxy cuticles on the fronds and stomata that can open and close depending on variations in the atmosphere and the availability of water in the soil (Krieg and Chambers 2022). The gametophyte lacks these anatomical features and is poikilohydric, meaning that it cannot regulate its water content (Kessler and Aros-Mualin 2025). Thus, the gametophyte can only rely on its ability to tolerate and recover from modifications in external water content (Krieg and Chambers 2022), so it is in a constant state of equilibrium with the surrounding environment (Pittermann et al. 2013). Although this may seem insufficient, the gametophyte is not the delicate structure it appears to be, but a robust organism that can cope well with abiotic stresses (Krieg and Chambers 2022). Among the proteins associated with the response to water stress documented in *D. affinis* gametophytes are the *A. thaliana* homologs of PYROPHOSPHATE-ENERGIZED VACUOLAR MEMBRANE PROTON PUMP 1 (VHP1), which confers tolerance to water stress by increasing ion retention (Li et al. 2005); and SUGAR INSENSITIVE 7 (SIS7), which catalyses the first step in abscisic acid biosynthesis from carotenoids in response to low water content (Ruggiero et al. 2004). This hormone is positively correlated with drought resistance in plants (Wang et al. 2019). We also identified CALCINEURIN B-LIKE PROTEIN 1 (CBL1), which regulates responses to drought.

*D. affinis* gametophytes must also cope with oxidative stress, which alters their physiological and metabolic balance. Therefore, they harbour proteins that play that role, such as the *A. thaliana* homologs METHIONINE SULFOXIDE REDUCTASE A4 (MSRA4), which restores the activity of proteins inactivated by methionine oxidation (Gustavsson et al. 2002); ACONITASE 3 (ACO3), which confers tolerance to oxidative stress; LIPOCALIN IN THE PLASTID (LCNP), which prevents thylakoidal membrane lipids peroxidation, conferring it protection against oxidative stress (Boca et al. 2014); and EXECUTER 2 (EXE2), which helps plants to perceive singlet oxygen in plastids, avoiding any possible photooxidative damage that can take place (Kim et al. 2012). Additional proteins associated with oxidative stress include ACTIVITY OF BC1 COMPLEX KINASE 7 (ABC1K7) and 8 (ABC1K8), involved in resistance to oxidative stress by intervening in reactive oxygen species production; ALKALINE/NEUTRAL INVERTASE A (A/N-INVA), which contributes to maintain the reactive oxygen species homeostasis in mitochondria (Xiang et al. 2011); and ENLARGED FIL EXPRESSION DOMAIN 1 (ENF1), which prevents the accumulation of reactive oxygen species (Fait et al. 2005). Depending on their concentration, these compounds can act as signalling molecules in basic plant processes such as response to stress, programmed cell death and ion transport, or be toxic to them (Ali et al. 2023). Additional homologs in the oxidative stress set include: Γ-GLUTAMYL TRANSPEPTIDASE 1 (GGT1), which prevents oxidative stress by metabolizing extracellular oxidized molecules of glutathione (Ohkama-Ohtsu et al. 2007); GLYCERALDEHYDE-3-PHOSPHATE DEHYDROGENASE C SUBUNIT 1 (GAPC1), key for plant adaptation to oxidative stress; GLYOXYLATE/SUCCINIC SEMIALDEHYDE REDUCTASE 1 (GLYR1), which intervenes in redox homeostasis; and NEUTRAL CERAMIDASE 1 (NCER1), which promotes oxidative stress resistance.

Another stressor faced by *D. affinis* gametophytes is the salt content. Salachna and Piechocki (2021) demonstrated that this fern does not tolerate salinity well and contains numerous proteins responsible for responding to salt stress. Accordingly, proteins required for this function were detected in our transcriptome. This set is represented by the *A. thaliana* homologs of α-MANNOSIDASE 2 (GMII), BEACH-DOMAIN HOMOLOG A1 (BCHA1), DEAD-BOX ATP-DEPENDENT RNA HELICASE 3 (RH3), MA3 DOMAIN-CONTAINING TRANSLATION REGULATORY FACTOR 3 (MRF3), PRE-mRNA SPLICING FACTOR SR-LIKE 1 (SRL1), and PROTEIN PHOSPHATASE 2C GROUP 1 (PP2CG1). Other proteins that caught our attention were SUPPRESSOR OF ZEAXANTHIN-LESS 1 (SZL1), which improves salt tolerance by increasing the synthesis of carotenoids (Chen et al. 2011); PROTEIN KINASE 2 (PK2), specific for plant adaptation to high-salinity conditions (Lee et al. 2017); CALCIUM BINDING GTP-ASE (CBG), playing a role in calcium signalling during salt stress; and NF-X-LIKE 1 (NFXL1), which provides resistance to salinity. Nevertheless, the negative effects of salinity on this species can be mitigated by growing in shaded locations (Salachna and Piechocki 2021), as *D. affinis* usually does.

Starvation–stress–related proteins detected in *D. affinis* may reflect both its natural understory habitat, characterized by low light and nutrient scarcity (Azevedo-Schmidt et al. 2024), and the overcrowded one-dimensional growth of gametophytes in the *in vitro* cultures used to obtain earlier transcriptomes (Wyder et al. 2020), as also suggested by Ojosnegros et al. (2024). In the present study, additional homologs of *A. thaliana* related to starvation include: PEROXISOME UNUSUAL POSITIONING 2 (PEUP2), required for autophagosome formation upon nutrient deprivation (Nakayama et al. 2012); 2-OXOISOVALERATE DEHYDROGENASE SUBUNIT β 2 (DIN4), crucial during sugar starvation; SOS2-LIKE PROTEIN KINASE 17 (PKS17), which confers tolerance to low potassium conditions; PROTEIN KINASE 6 (PKS6), needed for potassium homeostasis under low conditions of this ion; and GENERAL CONTROL NON-DEREPRESSIBLE 2 (GCN2), an activator of the amino acid biosynthesis pathway indispensable during adaptation to amino acid starvation. Further results in the starvation set comprised: 14-3-3-LIKE PROTEIN GF14 PSI (GRF3), which regulates nutrient metabolism; KIN17-LIKE PROTEIN (KIN17), which alleviates the stress generated during copper-limiting periods (Garcia-Molina et al. 2014); and PHOSPHATIDIC ACID PHOSPHOHYDROLASE 1 (PAH1), involved in galactolipid synthesis in the endoplasmic reticulum, required for membrane lipid remodelling, as an adaptation mechanism to cope with phosphate starvation (Mietkiewska et al. 2011).

Fern gametophytes, despite their tissue simplicity and lack of a cuticle, are quite robust and can inhabit warm or cold regions beyond the distribution range of their conspecific sporophytes (Firoozi et al. 2025), thus demonstrating their sturdiness (Kessler and Aros-Mualin 2025). In the *D. affinis* gametophytes transcriptome, some proteins identified were related to the response to cold, such as the *A. thaliana* homologs of β-KETOACYL-ACP SYNTHETASE 2 (KAS2), fundamental for cold resistance by maintaining the integrity of chloroplast membranes (Hakozaki et al. 2008); PALMITOYL-MONOGALACTOSYLDIACYLGLYCEROL Δ-7 DESATURASE (ADS3) and PALEFACE 1 (PFC1), both needed in chloroplast formation at low temperatures (Hugly and Somerville 1992); and β-AMYLASE 3 (BAM3) and FUMARASE 2 (FUM2), required for the accumulation of maltose and fumarate, respectively, under freezing conditions to protect the electron transport chain (Li et al. 2009; Dyson et al. 2016). Other cold-stress-related proteins identified were 3-HYDROXYISOBUTYRYL-COA HYDROLASE 1 (CHY1) and SENSITIVE TO FREEZING 2 (SFR2), both indispensable for cold signalling and tolerance; ENOLASE 2 (ENO2), which regulates cold-responsive gene transcription; Δ (8)-FATTY-ACID DESATURASE 2 (SLD2), critical for sphingolipid formation, which is required for cold resistance (Chen et al. 2012); and LONG-CHAIN BASE KINASE 2 (LCBK2), implicated in reduced root growth under freezing conditions (Dutilleul et al. 2012).

On the other hand, homologs implicated in the response to heat were also identified, such as FORGETTER 1 (FGT1), which contributes to acquiring heat stress memory through the control of heat-induced gene expression (Brzezinka et al. 2016); and RNA-BINDING KH DOMAIN-CONTAINING PROTEIN RCF3 (KH28), which regulates heat tolerance. Further examples are HEAT INTOLERANT 4 (HIT4), which plays an accentuate role in basal thermotolerance by regulating transcriptional gene silencing (Wang et al. 2015); and SERINE/THREONINE-PROTEIN PHOSPHATASE 7 (PP7), which confers thermotolerance to high temperatures (Liu et al. 2007). These findings add to the set of proteins associated with responses to biotic stresses annotated by our laboratory (Grossmann et al. 2017; Ojosnegros et al. 2022, 2023, 2024). Likewise, it should be borne in mind that the gametophyte is the generation in which reproduction occurs —asexual in the case of *D. affinis*, or sexual in the most species of ferns. Therefore, understanding how gametophytes respond to high temperatures will help predict how climate change may affect fern reproduction and, consequently, population dynamics (Chambers and Emery 2018). Moreover, as shown above, ferns exhibit traits that make them a potential source of candidate genes for stress tolerance and resistance. As suggested by many researchers, including Chen (2022), such genes could be used in crop transgenesis to improve food production, which in the long term would benefit both animal and human health.

### 4.4. Protein-protein interactions

The interactome of the 1,160 proteins related to light, transport and stress was analysed, as well as the total network generated, which, as mentioned in the Results Section, had a Protein-Protein Interaction enrichment p-value of 1·10^-16^, or in other words, a high significance. This result means that the network has significantly more interactions than expected, i.e., the proteins have more interactions with each other than would be expected for a random set of the same size. In this network, the proteins with the most interactions were MTV17, COG6 and SEC6.This is probably because these three proteins are involved in molecular transport, a key process in plants and, by extension, in fern gametophytes. Specifically, MTV17 is involved in the retrograde transport from early and late endosomes to the trans-Golgi network (Guermonprez et al. 2008), while COG6 is indispensable for the correct functioning of the Golgi, and SEC6 is a member of the exocyst complex that contributes to the docking of exocytic vesicles with fusion sites on the cell membrane during secretions (Wu et al. 2013).

As for the pairs of proteins that exhibited the strongest interactions in each group, LHCA1 and CAB4 were highlighted in the photosynthesis set based on experimental evidence. As explained above, both belong to the light-harvesting complex and share the same function: capturing and transference excitation energy to photosystems in *A. thaliana* (Wientjes et al. 2011). Thus, it is likely that they show stronger physical or genetic interactions in laboratory experiments than other proteins. In photorespiration, GGAT1 and HPR1 stood out in the database evidence. In this case, the first regulates amino acid content during photorespiration (Verslues et al. 2007), while the second is essential to the photorespiratory core cycle (Cousins et al. 2011). Turning to photomorphogenesis, the proteins REV1 and REV3 had the strongest interactions in the UV-response experiments. The first participates in the resistance to UV-B and the second to UV-C (Nakagawa et al. 2011). As for xanthophyll metabolism, the strongest interaction was between LUT5, VDE1 and BCH1 with each other in databases. This is likely due to their key roles in xanthophyll metabolism, including biosynthesis and the regulation of zeaxanthin levels in chloroplasts (Hieber et al. 2002; Kim et al. 2009). In transport, SEC10 and SEC8 were notable in the text-mining interaction. As commented above, they are components of the exocyst complex, which mediates the docking of exocytic vesicles at fusion sites during secretion (Drdova et al. 2013), and this may cause them to be mentioned in the same PubMed abstract or articles in the String program. The same occurs in the response to biotic stress with UPF1 and UPF2, both necessary for plant defence and wound response (Shi et al. 2012; Merai et al. 2013), which notably showed the strongest text-mining interaction. Also, in the response to water deprivation, salt stress and starvation, the strongest interaction was found between SCABP5, an essential regulator under water- and salt-deficient conditions (Li et al. 2013), and PKS17, which confers tolerance to low potassium conditions (Hashimoto et al. 2012). Different results occur in the response to oxidative stress and cold, in which the proteins rose above the rest in the experimental type of interaction. ELP1 and ELP2 belong to the oxidative-stress group, as both are vital for signalling this stress (DeFraia et al. 2010; Nelissen et al. 2010), while CER6 and KCS9 are part of the cold-response group, as they are required for generating a response against freezing conditions (Joubes et al. 2008). Their shared activity may explain why they interact strongly in physical or genetic experiments. Finally, regarding the response to heat, KU70 and KU80, which participate in the plant’s defence against this type of stress (Liu et al. 2008), showed the strongest value in the text-mining interaction, i.e. they are mentioned in the same abstracts or articles.

A notable aspect of the current research is the considerable number of potential gene fusion events, identified by the String program, among homologs of proteins associated with heat-stress responses. Gene fusion may lead to the formation of novel proteins that enhance a plant’s ability to withstand elevated temperatures, for example by improving protein folding or scavenging harmful reactive oxygen species. It appears that heat shock proteins (HSPs) are particularly susceptible to fusion with other genes, a point emphasised in the Results section. Furthermore, numerous HSPs have been documented in the gametophyte of *D. affinis* (Ojosnegros et al. 2024), prompting us to ask why gametophytes of this fern accumulate such high levels of HSPs, and whether these proteins contribute to heat-stress resilience. Additionally, research has explored the use of gene fusions to improve heat tolerance in various plant species. For example, in tomato, a fusion of trehalose-6-phosphate synthase and phosphatase genes was employed to enhance heat tolerance (Lyu et al. 2018). Among other strategies, the development of gene-fusion technology offers promise for creating climate-resilient crops, potentially helping to mitigate the impacts of climate change on agriculture. On the other hand, regarding protein co-occurrence interactions, the graph shows that several heat-shock proteins are conserved across the phylogenetic tree provided by the String program.

In conclusion, although the study of ferns has been hampered by the lack of high-quality genomes, RNA-sequencing has proven to be a powerful methodology to identify and characterise thousands of proteins in the gametophytes of the apogamous fern *D. affinis.* From its transcriptome, carrying out an exhaustive and detailed work, numerous *A. thaliana* homolog proteins associated with biological processes of photosynthesis, photorespiration, xanthophyll metabolism, and photomorphogenesis were annotated and discussed. Also, many homologs implicated in transport and response to biotic and abiotic stress. According to the String program, it was revealed that the proteins with the highest number of interactions had a role in transport, and that most of the interactions were based on text-mining and database evidence. Likewise, *D. affinis* transcripts exhibited high homology when compared to fern species with sequenced genomes deposited on the FernBase site. We have attempted to expand the little existing knowledge on this species, although the real function of these proteins in the gametophyte has yet to be verified. The information generated contributes to a better understanding of ferns and may serve for future applications in other areas, as deciphering their genetic diversity could provide a significant boost to progress.

## Acknowledgements

This research was supported by the University of Zurich, the University Research Priority Program Evolution in Action, and the Program for Mobility of Excellence at Oviedo University. We would like to express our sincere gratitude to Dr. Stephan Wyder for his great work on transcriptome performance and his humility. Also, to Hanspeter Schöb (University of Zurich) for his kind administrative and logistic support, and the Functional Genomics Centre of Zurich for sequencing and generous assistance with all the analyses.

## Author contributions

Conceptualisation: Helena Fernández and Ueli Grossniklaus; methodology: Sara Ojosnegros, José Manuel Álvarez, Valeria Gagliardini and Helena Fernández; formal analysis: Sara Ojosnegros, Francisco Vázquez and Naiara Goya; writing - original preparation: Sara Ojosnegros, Francisco Vázquez, Naiara Goya and Helena Fernández; writing - final review: José Manuel Álvarez, Luis G. Quintanilla, Jaume Flexas, Alexis Peña, and Ueli Grossniklaus. All authors read and approved the manuscript.

## Data availability

The *de novo* assembly of the transcriptome of the *D. affinis* gametophyte in FASTA format and the transcriptome annotation are available in the Zenodo research data repository (www.zenodo.org) with the identifier https://doi.org/10.5281/zenodo.1040330. Specifically, the sequences studied in this work are provided in the Supplementary Table 1.

