## Supplemental Table 1 for "Comprehensive analysis of proteins associated with light responses and stress tolerance in the gametophyte of the fern *Dryopteris affinis* ssp*. affinis*"

**Supplementary Table 1**. Nucleotide sequences of proteins found in gametophytes of the apogamous fern *Dryopteris affinis*.

**PROTON GRADIENT REGULATION 6 (PGR6)**:

TTTTTCTGGAAAGCTTTCAATTGCATTTTTTTAAAGAGTTTGTCTTGTAAAACCTTTCATGTACTTACTTTCATTACATTGTGGTACACTACTTTCATGTGGTACAATATTGTGGAAGAAATGTTCATGGAACCAATAGGTAACATGAGTTGTATTTACTAATATACATCAATACCAAACCATTTAAGATGGTTTGGTATTTATGTTCATGAAAATGTGTCTACAGCCCTTGCCAAGCCAAGGTTGCAACAGCAAATATCTACACAAAAGCTATAACTTGGAATATCTATACAAAAGCTATTACTTGGAATACTTCATTACTAGATTACAAATCAGTGTTCACTTTTGCCGCGCCATTAAAGGTTCACCATTCTCTGCTAGCACATCCCCGAGTTGCCCTCTACGCAAATTTGTCATCCGGCTCCAAGTGCGTCATACATCCATCCAATTTTTTAAAAACCCAAACACCTATGTAGTTCTTATACATATTTAGGAGTAAAATGCCCCTGAAGAAAGCATTTACATCCTTGACAACAAGGATATCCTTTCCATAAGACCTTGACTTTGACATGCCTCTCTTCAAAAGAATGCTGCTTTCACATAAAGAATGACACACACAAGATGATGTACTCCATGCAGAAAATCATTGAACCATTATGAAGAGCAAGAGTAAAGCTTGCATGACCAGCTCAGCGGTCCACAAGAACTGACTCGCTCCATGATAGCATTAGATCCCTTGTTGCAGCAGGTATTCCTTGCACAACATCGACAACTAAATTTGCAACGTTGAACTCCCCTTCAACAAGTTTGTAGATGTCTAACAGCTCTCGAATGTGTAGTCTAGAATCCTCTGTGAATGCCATCAGCAGCTGCCGCCTTATTCCAGCATCGGTGAGGACAACTCTTGCACCATCCTTTATTGTGTCAGTAAGATCTAACTTCGCTCCACTGGTTGTTGATGACCTCATCACGTCCCGGCGGTTCCTCGCTTGTAAAGGAGTCATCATCTCAACCTGCTCTCTTGCAAGAACAATCAAGTTCTCTAAGCGTTTCCATTGAAATGCTCCATCCTTGAACAGAACCTGAACAAGGCGTTCTCGCAAGGCTGGGTCAGGGTCAGTCAATAAACGCTTGGCCACATAAGGGTAGGCCACCTCGAGAAACTTGAAATCTGGTTGAAAAGTGAGACAAATTCCCTCTTGAGTCAAAAGAGAGCGAATAACTAATGAGAATCGTTCAGGAATGCGTATGGGATATTGATACACAAGCTGATTAAATTTGGCAGTCACAGAGCGGAAGTTGAAATCCGACAAACCCTTCACGGTTGAATTTTGCCAGATTGCTTCTAAAGCTGGTACAATTGGTGTCAAATCTGTTCCAGGTGCAAGAAAACCCAACCTAGTGAAATCATTGGCCATCTCCACATAATCTTCGTTTACCGCATGGACAACTGCATCAATAAGAATCTGCTTGTTTTGCTGACTTAGTACTGCAACATTTCCAAAATCAACATAAGCGATACGTCCGTCTCGCATGGCAAAAATATTTCCTGGATGGGGATCACCATGAAACAGTCCAAACTCAAGCAATTGCCTTAGAGCTGCACTTACACCCACTGTTATAAAAGCTGGTACATTAATTCCTGCATCTTTGATGGCCTGAGGATTGGTGCAGCGTATTCCATCAATCCACTCCATTACAAGCACTTTAGAACCACTTAGATCTTGGTAAACATCTGGTATCTTGACTGTGGCATCATTTTTGAAGTTCTCCAGAAAATCCTTAATGTTGCGTGCCTCAAGAGTGTAGTCAAGTTCTTCAAGTAGTTTTTCACCAAACTCATCAACAACTAGTTCTGCATTGCAGCCCAGCTTCTGTAGGCTTATTCCATTCAAGAAGGAAGCCAATGTTCTAAACAGAAACAAATCTCTGTAGATTATGGGTTCTATCCCTGGACGCTGAACCTTGATAGCGACCTCTTTCCCAGAGCTCCGAAGTGTGGCCCTGTATACTTGTCCTAAGCTTGCAGCAGCCACGGTTTCTGATGATATTATGCTAAACAACTGCTCAAGCGGTGCACCTAACTGTTCCTCAATTATCTCAAAAGCCACCTTATTAGGAAAGGCAGGCACATCATCTTGTAAAATGCACAGCTCATTCATGTAATCTTCACGAATGATGTCTGGCCGATTTGCGAGGACCTGCCCAGCCTTAATGAAAGATGGTCCAAGGTCACATAGTAAATTCCTTAACTGTGCAGCTCTGTAGGCCACCATTTCCTCACCTTTGCCAATAAGACAATCATAAGAGAGGCTGCCCCAATAGTAACCTAATTTCAAAAGGATCTCGGCGCCACGCGAAACCACAGAAAAGAAAGCCCCAGGAGAGTCTAGAACCTTCTGTCGAACCCCTTCTGGTGTGTACTTCCTAAAAGGGATGCAAACTCCTCGTTCCAAGTCGAGGACATCCATGGCTGTGCTAGATTTCTCCCCCATGGGAGTGTTTACAAATGCATTCTCTAAGGTATCTCGACGCCGCAGGAGAGTACGCTTTGCAGAAGTGGCACTTCTGGTCTTTAGAGGAGCAGAAGAGACTGCAAAAAAATCTCCACTTCTCCGAGTTGGGCTTAGGAATTCGGCAGAATTATTGCAATTTGCACATAGTCTTGCTCCAAGGAAGCTGTTTCTATAGCCATGGCCAACATCCTGTGCATCCGAGGTAAGTGGAGACGCACGAAGCCCTAGGAAAGAATCTGAAGCTGAAGCAGCCATGGCCATCTCCAAGGAATACACAGAGACTAAATTGCACAAAAAACACCTAGAAGAAGCTTTCCATGTAAGCTTCTAAGATGTATGTGCACAGAACAAAACGAGGTTTCTTCACGTGGCTTCAAAGCTTCACGGAAGATTTCCCTGAGATGTACCTCCTTGTCTGCGGAGATGACGAGCACGCAACACACAGAGACGAATTCAAACAAAAATGCGCAATGTCTGTGTGAAGTTGCTCCACAAGAAGGCGATTAGCTGCAGAAAACCCTAGCTGCCTATGGAGATGGCGTGTCGTGCTGATAACAAGAGGGATTGCTTGAAATAATGTGAGGCCTGCTGCAATCGTTTCCATTTTTTGTG

**LEAF-TYPE CHLOROPLAST-TARGETED FNR 1 (LFNR1):**

ATTTATATTAGAAAGAGGGGCTGACAAGTGTAATTAGGGGAGAGAGCCACGTGGCCTTACATAAGAGACGGCTACCTAGGATGGAGGGCGTGAAATTAATTACAAAGTGTCGCATCATCAACGACGACGTTGGGGCGGGGAGAGTAGGCTCCGAAGCTGAGGAGGAGAGGTCGCGCCGGACACTCCATGGCCGTTGCAGAGACATCTCCAAACAGGTGGGAGGAATTCAACAGTCTCCATTCCACCACGTCTTGAGGCTTCTCATCCTGGGTGTTTTGCAGGTGCCAGGGGGTTCTTGGTTGGAAGCGATTGACGGGAGCAGAGCAGAGCAGAGCATAACAAAGCCAAAGCACACTGCTTCTCCTCTTGCTCTGCCTTTATACCTTCAGTATGTTCCTGCTTGTCTGAGCTGTTGCATGTTGTGTTTTCCAAGCTTTCAGAACCTTTAGGTACACTGTCCTAAGAGAAGGATTGTATGAGCTAGATTGTACCCGTCTTTACAAATCTGAGACCAAGTCCACGAGAATTTAACGAAGTAGGATGAAACTCAAGGATGAAACTCAAGTTGAAGCTCCACACACATACAGCCGCCGTTGTAAAACGATGGCCTGCTGAAAAGAGGGACCTCTTCTGGGTGTTGCAGACTTTGCATTTCTAGTTTTTTTGGTTGTTATGGGCGTGTCATAATGTGCTTGTACTTAGATACCAATACTTGTATCGCATGATGTGCTTCCTGAATAGTTGCTGCGCAGCAAATTATTCTAGGATGCCCTCTGTTGTGAGATAATAACTTCCATGAACTACTTTGAAAAGAAGGCCTTGTGGATCCTCCAGCGATTTGTTTAAACCTCTTGTGATGCAATCTGAACCTCTACAAACCTATTTTATTAGGCCTGGGGTCCGCATGGCTTTTTATACACGAGTACAGCTGTGAAGATTGTCTCATCAACAACCAAACTCATACCATCAGCTTCGACATTTTACTAACATTCAGCTACTCCACTCCAGGGGTAGTGATGGTCTACAAGAATATGATTATCTTATAGATCAATTGCAGAAAAGGCCTTATGCTACAGAAGAAGATTCTAGTCAATGTAAATGATGGTTTTATCCCGGTTAAAATCTTTATGAAGATCAGGGGCCTACGGAAGAAGATTCAACCTAATATAGTACAATGTAGCATCTAATGCGTAAAGTCTGTGACAAGTAAGGTTAGCAGAGCACCATAAACCTATGGGATGCACCTTTTCTAGCAGAACATGAGAAATATACCAATTGATAGAATTCAATTAAAACAAAACTGTGTAATTGCTTGTAAAACAGTACAACTGTTTGTAAACTGTGTGTAGCAACAAAGCAGTGTGCCAGTTTTCTTCAGCCAATCCTTTCGTAAGATGGAAGGCTGTACCCCTCATGATAGCAGAATTGCCTGGCTATATTAGTGGCTTAATTCTGCACCTCTGTACTCTCCACATCGAGGCAATTGAATACTTCATGGCATATATATAGCCCCTGTTTTCCTCAAAACAAAAGCAAGAGCAGGGAGGTGAACTTTGCCTGATAAGTCTTGGCCCCGTAAGAATATGCAGTCTAGCCACTCACATGATCGGAGCCAAGACGGCCCCGTAAGAATAGGCAGTCCAGTGACTCACATGATCATAGTCTAAAAGGGTTACTGCCATATTTCTGAATAGCAAACATTTCCATATACCAGAAGAAGAGTAAATAAGCCAAGAGGAAAACTAACTGTTACTCAGGACCAGCAGGTAAGATTGTTATCACAGCAAAAAATGAGCAAACAGTAAGCAAACTATTAAGGCAACAAGAAAAAGAAAGCGGAGGACTTGGTTTACTCTCGCAATCTTCATCGCAATGTCGTCCAGGCAAGACCAAGAAAAGCCAGGAATAAAGGACAGGGCCAAGGTAGCCCCTACTTACAAGAACAACAATCATAGTGAAAAGTTGTAAATGAACTTGATTATGATAGGGTAGATTCTAGAATGTGCTTCTTCAATGGAATTGCTCGCCCTTGAAGGTTTGCCCCAACTGCCAGGATGCAGGGGCAACATCATAGGAGATGAGGCTGGAGCCGTCCTTGAGCAACACTTTGAAAGAGAGGCTCTGCCCCCTCATGTCCTGTGGAGTTCGCCAATTGGAGCCCCAATTCCGTTCAAGGGGAATCCATCCTGTCTTCGACCCCTTCACTAACAAACTTTTCACCTCTCCTGTGCCACCCACATTTGTCAGCAGCACTAAATTGTATCGCGAGTTGCCGTTGATGGTAAACTGGATCCCCCCTTGACGGGTGCAGGGCACACTCCGATAGAGGACAGGAACAGGATTGCCGGTGCTGTAGCTGGCGATCTCCGAGAAGGCAGAGTTGGTGACGTCAAAATGCTTGTTGGGTGGGTTGCACCACGCATTAGGATGCTCGGAGTTTGGAGGGCACAAATTCGTCGCAGTTACCACTACGCTTTTCCCAGCTGCACACCGCTTTGCTGCATCGGGAGCGGACATGTCACACTTAATCTCAAAGCAGCTGCCACATCCAGACCCACTATTGAAGAGGGAGGCACTCACTGCCGCTGTGTTGCTCCCATAGCCACTGTCATAGCGGTCTCCATAACCACAAGCACCACTGTTAATGGTGCCGGAGTCATCTCCATAAACAGTAGCCTTTGCTGTTAGCCATGTCGCTTCCGTCACTGCTGCAGCAGCTGCTAACTGCATGCAGCTGACGAGCAGCAGTAACACAAGCAAGTCGCTAGACGCCATAGCTTGCATGCCAAGCTCTCTCAGCAGATACAAAATTCAACCCAGAAGCTGCCACGCGACGCCCGAGCTTCACGGGAGAAGAAGAGGACACGGCGCGCTCCTATGCTTTGCTAGCACAAACAGCGGGAAATGGGCACTAGAGAGAAAGGAAGCTATTCTATATTTAATGCTCCTGAGTCCTCCCTTCTGTATGCAATAGTTTTGTAGGTGAGACTCCGATGAGAGAGAGAGAGAGAGAGAGAGCATGGCCTCGCAAGCTGCAACAGTGCCCTCCTCCTCGGCTGCTGCTTCCTCTTGCCTCGACCGCCTCTCTGCAACGACCCTCCGACCTGCAGTGGATCGCGTCTCATTGCCTGCATTCAAGTCATTCTTCAATGGCACGGCCCTTTCTTGTGAGAACAAATCTAGCAACTTCAGGAATGGGGGTGCTGTCACTGTGAAAGCGTCGGTTGCAACGGAAGCACCTGCCAAGGTTGTGAAGGAGTCCAAGAAGAATGATGAGGGTGTGATTGTCAACACATTCAAACCCAAGAGTCCGTATGAAGGTCTATGTCTGCTCAACACCAAGATTGTTGGGGATGATGCTCCTGGTGAAACCTGGCACATGGTCTTCAGCACAGAAGGTAAGGTGGACTACAGGGAGGGTCAATCTATTGGAGTTATTCCTCCTGGTTTGGATGCTAATGGAAAGCCTCACAAGCTTCGCCTTTACTCTATTGCCAGCAGTGCAATTGGCGATTTTGGAGACAAGAAAACTGTTTCGCTTTGTGTGAAGCGGCTTGTCTACATCAATGACAAAGGGGAAGAAGTGAAAGGGGTGTGCTCCAACTTTCTATGCGATTTGAAGCCTGGGTTACCTGTTTCTATAACTGGACCTGTTGGAAAGGAAATGCTTATGCCAAAGGACGAGAATGCAACCATTATCATGCTTGCGACCGGAACAGGTATAGCACCTTTTAGAGGGTTTCTGTGGAAAATGTTTTTTGAGGAGCATGCTGATTACAAGTTCAAGGGACTCTCCTGGCTCTTCTTGGGAGTGCCTAACAGCAGCACTTTGCTGTACAGAGAGGAATTCGAGCTTATGAAAGAAAAGTTTCCCGAGAACTTCAGGCTAGATTTTGCGGTCAGTCGTGAGCAACAGAATGAGAAAGGAGAGAAGATGTACATCCAAACACGGATGGCACAATATGCTGATGAGCTGTGGGAGCTTCTGAAGAAGGACAACACATATGTGTACATGTGCGGTTTGAAGGGGATGGAGAAAGGAATTGATGATATCATGACAAGTCTTGCTGCCAGAGATGGTTATGACTGGTTTGAGTACAAGCGAGAGCTGAAGAAGAACGGGCAATGGAATGTGGAGGTGTACTAAATTTTACTCGAAGAGCTTCAAGGTGGGGATGTCCTTTTTTGTACAAAATCCTTTTGGCAATGGGCTTCATTAATAACCAAATGAAGCTCATTCTGCCTCTCTGTACCATGATCCATCCCTAGTCCTATGATCTAGCGCCAATATAGCTTCTTCCAAGAGCCTCTCTTGCATTTCACTGCATGTACCATTGTATGTGTTCATAAAGTCAATATCTACAACAAACTCTTAAGACCTGCCATTTGGCTTGAAGCAATTCGGTCTCAAGTGCTGTGTGTTGGTGATACTAATTTTAATGATGTAAGTTTTAATATTGACTTTGAGATGGTCACACACTGAGCCACACCGATGTTTCGTTAGGGAAGTGAAATTGTAGTAAAAATTTCACTATAAGGTATGCATTTCCG

**LEAF-TYPE CHLOROPLAST-TARGETED FNR 2 (LFNR2)**:

TTTTTTTTTTACCTGAAATGAAATGCTTTATATATTACACAGAGGACAAGACATCAACTGCAGCTGTTGGGACAAAATGCATCTCCAAATTCGACAGCAATGGTTGTCACCTTTCGAATGCAGGTCTACATTAGTACAATCCCAAATAGGGGTGTTTGGGCTCAGCATCAAAGGATGGAGTGTATACAAGGGAGAAACAGGGTAAGGTCAGCTCGGTAATACACACATACATATCAACACCAACTTGATGCCCTTCTTCTGGCCTATCTTTGCAATTGAAAATTAGCAGCGCTTCTCTTCCATTAGTCTGCGTTCTAATCACCATGATCTGTAGCTGCAATCCAGCTCGACAACATGTGCCAATCCTGTCCTCCTGTTCATGTCAACGAATTGATACTTATTAGTAGACCTCCACATTCCATTGTTCCCTCTTCTTAAGCTCCTTCTTGTATTGCAACCAGTCATACCCATCTCGAGCGGCCAGACTGGTCATGATGTCGTCTATGCCTTTCTCCATCCCTCGAAGACCACACATGTATACGTAAGTGTTATCTTTCTTCAAAAGCTCCCATAGCTCATCAGCATATTCAGCCATCCTTGTTTGGATGTACATCTTTTCACCTTTAGAGTTGCTTTGTTCTCGACTAATAGCAAGATCATATCTGAAATTATCTGGGGCAATATTTTTCATCTTCTCATACTCATCGACATAAAGAAGCGAACTGGAGGTTGGAACACCTAAGAAGAGCCAAGCAAGCCCTTTGAACTTGTAGTTTTCATGCTTCTCAAAGAACATCCTCCATAAAAACCCTCTGAATGGAGCAATGCCTGTACCAGTTGCAAGCATGATCACTGTAGCATTGGGGTCAGTGGGCATTAACATCTCCTTCCCAACAGGGCCTGTGATTGACACATCACTTCCAGGTTTCAAGTCACAAAGAAAATTTGAACAGACACCCTTCACTTCTTCACCCTTGTCATTAAGGTAGACAAGCCTTTTGACACAGAGGGAGACTGTTCTGTAGTCCCCAAAGTCACCAGGAGCACTGCTAGCAATGGAGTAGAGGCGAAGCTTCTGAGGTTTACCATTGGCATCCAGACCCGGTGGGACAACACCTATGGACTGACCTTCACGGTATTTTACCTCGCCTTCAGTACTGAATACCATATGCCAGGTCTCGCCAGGAGCATCATCTCGGACAATCTTGGAATTGAGAAGGCATTTACCAATGTATGGGCTCTTGGGTTTGAATGTATTCACAATCACACCCTCATCATTTTTTCTAGATTGCTTCTTTACAGGAGGGGGGGCAGTCTCTGTTTCCACAGAGGCCCTTACTGTGGTACCTTTACCTCCTGCCAATCTTGCCTTCAAAACTGAGGAGTCCAGGCATAGCCTGTTCCCACTTAAGAAACCCTTGTTGATACAGAAAGTGACATTGGAGGAAGCAGAGGGGGAGTCGTTAAAGGGCTGCTGCAAGCAGCGCCCATGTAGAGCAGAGGAAGATGCAGAAGAAGCAACGGCAATCGAGGAAGCCATCGCCTTCTATAGCGTCTCTCTTCTGTGCCAGCCTTCTTCTTCAGAACTAGCTTGATTGCTCGTGCTCCCTATCCTTGTGTGGCTCTCACTTTATGGATAGGCCAGCCCTCTCCCTCCACAAATCACATTTTTGAATGCTGGTATAGTTTTAGACAATTCCCACCATTATTCTTACCACAACACATGCCTCGCTATTATTTAAGCTAATGCCATTGTGGGGCAGACTCCAGAGAACTTGCTAGCCCCTACACTGTGCCCACAATCACATGTCTAGGGT

**K (+) EFFLUX ANTIPORTER 3 (KEA3):**

TTTCTACATCGTCTCTCGTTTTTTCAGCTGATCCATACAGCCCACCTCGATTTTTCAAGCATTACCGACCTCGGGCAAGCTTTGTATCAAGCGGTGCCCTCCGTCGCCAAACCCTCTTCTCCATCGCCCAAATGCTTCCAGTTTCTCAGCTTCTTCCCCATCGCCAAGATAACTTGCTTCCAACAAACAAAGCGCTTGCTTTATGACCCACACACGCCCCCATCGCCACTCATTTCTTCGCTTCTTGCCAGAAAAACAAGCTCAGCTACAAGTCCTCAAGACATGAACGTATGATGGGCAGCTTGCAGAACCCTCCACTAATGTGTCTTGGGTCTGGTCTTCTCCATATTGCATGCACAGCTCAACTGAACCGCAACATCAAGTACCCAATGGAAATTACTTCATGCAGAGCGCGTCGTTCAGTTCATATTCATGCAGCATTACCGGGTTTCTGCAAATGAAGCCTCTTCTCTTACACATCTCCACAAACGCGTCTACTCATCCTTTACGCAAGCTATGGGAGATTTGAAACCTGTGTCGGCTCATAAAGCTAGTGTCACACATACGTTACGTAGTCAAAGGTGGCACAATGTTTCAATATCTAGTTCTTTACTGGAAAAGAGGCAGCAAAGGTTTTATGAATGTATTTCAAGAAAAGTCAGTTGTAAAGCAACAATGCACCTGCAAGCCAGCACCAGGCATCTTAATGCCAGGCACCCTACACTAAACTGTGGCAGATGGTTTCGTCAATGCAGAAAGTCAAATTGCTTTGCAATGCGAAATCGGTCTACTAGACATCTATGGTTGGTCCCAAGAGCTGAAGTTAACCTTGCAGCTGGAGTAGATGTTATTAATGATTTGGGTCTAGATACTCTAACTTTTCTAGCAACAACTGTTGTGGTTGTTCCTACATTCAAGGCTCTGAAAGCTAGCCCGATTTTGGGGTTTTTTGTCTCAGGAGTTGTTCTAAACCAGTTGGGTTTGATAAGGAATATATTAGATGTCAAGGCCTTGTCTGAATTGGGCATCCTTTTTCTTCTGTTTGAGATGGGACTTGAGCTTTCCTTGGCACGCCTTAGAGCACTTGCCAAGTTTGCTTTTGGAATGGGTCTGGCGCAGGTTTTGTTTTCAACTTTAGCTTTTACTGCTTTTGAGCTTCCACCAAATGGTGCTATTGGGACAAAAATCTTGGAGTTCTTGTTTCATTCAAGGCCAGATCTGGTGAATATTCGGACAATAGATGAAGCGGTTGTCATTGGTGCAGCACTGTCTATGTCTTCATCAGCCTTTGTGCTTCAGCTTCTTGCAGAGAAGGGTGAGCTTCCAACTCGTTTTGGTTCAGCCACACTTGGTATTCTACTCTTTCAGGACATTGCTGTGGTTCCCCTGTTGGTCATTTTACCAATTTTAGAAAGTGGGACCTTAGGGAAGGAAAGTGTCTGGCCACTTTTAGCTGCTGAGAGTCTAAAAGCTCTTGGTGGTCTTGGTATTCTCTCTTTTGGGGGTCAAATAGTTTTGCGTCGTATCTTTGAGGTTGTGGCTGATTCAAGGAGTGCTGAGGCTTTCAATGCTCTGTGCCTCTTGACAGTGACTGGGACATCACTAGTCACTCAAAAGCTTGGTTTCAGTGATACACTTGGAGCTTTTCTTGCTGGCGCTTTGCTGGCTGAAACGAACTTTCGCACCCAGATTGAAGCAGAAATTCGACCGTTTAGAGGCTTGCTGCTGGGTCTATTTTTTGTCACAACAGGGACCTCCATTGATATTCAGCTACTGCTGCGGGAGTGGCCAAATGTGCTGACTCTTCTAGGTGGGTTGATTGCAATAAAGACAGCAATAATCACTTCCATTGGGCCGTTTGTAGGATTGTCTCTACCAGAAAGCGTTCGCATTGGGCTTTTGCTCTCTCAAGGGGGAGAGTTCGCGTTTGTGGTGTTTTCCCTTGCCAATAGATTGGGAGTGCTGCCTTTGGAGCTGAACCGTTTACTTATTATTGTTGTTGTTCTTTCCATGTCTTTGACACCTCTTCTGAATGATTTGGGTCAGAAGGCTGCACAAGCTATAGAAGCACGTCTTGATATCCCTCGAGATAATTCGACGACTGCAGAAACATATGAGGCGAATGATCCTGTTGTTATTATTGGGTTCGGACAGATGGGCCAGGTGCTGGCCAATTTTTTGTCAACTCCATTGGCGAGTGGTCTAGGTGGTGATTCTGCTGGATGGAGTTTTGTAGCTTTTGACTTAGATCCTGGCCAAGTGAAGGCTGCTCGAAAATCCGGATTTCCAGTCCTGTATGGTGATGGCTCACGTCCTGCAGTGCTGCAGACAGCAGGCATCACTATGCCAAAGGCAGTCATGGTTATGTACACTGATAAGGAGAAAGCAGTAGAAGCGGTGCAACGTCTGAGACAGGCTTTCCCAGGGGTACCTATATATGCTCGTGCAGAAGACCTTTCACACTTGCTTGAGTTGAAGTCAGCAGGAGCTACGGACGCTATCCTTGATAATACAGAGACCAGCTTGCAGCTAGGATCTGTCCTCCTTCGAGGATTGGGTGTCTTGTCTGATGATGTAAGCTTTTTGACTCGAGTCATGCGCGAGTCTCTGGAGCTGCGTGCCAGAGTGGCACTTGACAGGCAGGAAGAGAAGGATGGAGCCTGGCTGAACTCCTTCCAGGTTAAAGTAAGGCAATCATGGGATGCCCGTATGCAAAAGCAGCGTCTCAGAGAACCAAGGCGCCTTTCTCCCAAACTAAAACTGGAAGGGATTATTGACTCAGAAGCTGTGTCTGAGAATGAGCACACATCAATCACTTCCAATGGCAGTTTATCAGAGCACAAGGCAGAGCCAGAAAGCAGTTCAGAAGGTCATGATTTAATAGACAGTGCAAACAGCATGGCAGCACAGAATTCCTTTGAGCTTTCAACAGCTAATGTTTCCCAGACTCCCTTGATTTCATCCAGTGCAGACAATGAAGTAAAGGTGCCCTGGCACCAAAAAATGAGTTTTTCAAAATATGAACCAAGTTCTGGATCTGTAATTCACCCAAGTCCAGATGCTAGAGGAGTAACATGGTGTGCATTGACTAATGAAGAAGGCTCTCAAGATGCTGATGAGGACCGTCCAGATGCGGACGTGATTGATCGAAGTGTTGAATACATAAGTAAACTTGAATAGTTTCAGAGGCCCTGGCTTATAGAATTGTGAGTGTTTAATAATGTGCACACCTTTGTCTGAAGTTTCAAAGACAACTCTGATTTTTCTTGTGAGTTGCAGCTTTGTACAAAACAGTATGTCTGAAGATAGTACCGTCTGTCCAACATTACAGAGATTTTTTTACTTGCTTGGTGGAGATTCATTACTAGGGTCTAGTAAGAAGCCCTTTCTTAACAAATTTGGGCTCATCTTCATCATCCATTATTCCCTTTCTTGTGGGAACCAAGGCCATCACCAGGTGTACCCCGACTACTTCAAAAAAGTGGATTACGAAACAAGTGTCAATGGAAGTTCAAGTGACAGCAAATCAAGGCATTAATTGGAGTTGTACTTCTGTAGTTTTGTGTATCCTTGGCTCTGCCGCTCAGTGATTACCTTTAACATTTTGTAACTGCATTTATTGAAGTTGCAATATTGCAGCTGAGAAGCCTGTATTATTGGCTTTTTGTGACTCATTGAGCAGTTTCAACATCTTGTAACTGGCTCAGAATATATTTCTTTTGGAAAGTGTATACATTGCATGTGCATATTGACAATTGTAACGTGTAATCTGTGCAATCAATTGCTTACCAGTTTAAGATCAAAAAAAA

**HIGH CHLOROPHYLL FLUORESCENCE 136 (HCF136)**:

AAGAGCGTTCTCTAACTTTAGCTCATGTTTTTTATGTCCTCATAATCCTATACAAATGAGTGATATTTGGCATGCTACGTGGGAGAGAATTGGATAAGGTCCCAAACTTCTCATTTTATACAAGTGCATTGAGACGAAGGGATGGCGACGGCAATGGCTGGTCTGCACTTGAAGTTCGCCATAGCCAGCTCAAGCAGCGAGCTTTTCTCCTCAACCCAGCGTCTACTGGCGTGCGCGCTGCCCTCCCTCTCGCAGCAAGCGCGACCTTCAACAGCTCGTTCCTGCCTGGCCAACAACAGCGGCGATGGTGAGGCCATTTTCAGCAAGAGGCAGCTGCTCGCGGGCTCTGCATCGCTCTCCTGTGCGCTTGCTGCCTCGTCTTTGGTGGGGCTTCCCACCAGGGAGGCTCTAGCGGAAGAGCTCCTTTCTGAATGGGAGATGGTGCCCCTGCCCATCAACTCCGGAGTTGTGCTCTTGGATATCGCTTTTGTCCCCGAGGAACCTGACCATGGCTTCCTCTTAGGCACAAGACAAACACTTCTGGAAACAAAGGATGGAGGAAAATCATGGGCACCTCGATCTATACCATCTGCAGAGGATGAAGATTTTAATTACAGATTCAATTCTATAAGCTTCTGTGGCAAGGAAGGCTGGATTGTTGGGAAGCCATCAATTCTCCTACACACTGCTGATGCAGGCACGAGCTGGGAGCGAATCCCTTTGAGCTCGAGGCTTCCTGGCACTCCGGTGCTTATCAAGGGAACGGAGGAAAAAAGTGCGGAGATGGTCACAGACGAGGGAGCAATCTATGTGACATCCAATAGGGGCTACAATTGGAAAGCAGCCGTAGCAGAGACTGTTTCGGCCACATTAAACCGTACAGTTTCAAGTGGAATTAGTGGCGCTAGCTATTACACTGGCACTTTCAACGCAATAAATCGCTCAAGTGCTGGAGACTATGTTGCTGTATCAAGCCGTGGCAACTTCTATTTAACTTGGGAGCCTGGACAGGAATATTGGCAACCGCATAACCGCACAAGTGCTCGAAGAATCCAAAGCATGGGCTGGCGTGCGGACGGTGGACTATGGCTTGTTGTGCGAGGAGGAGGGTTGTACCTCAGCAAAGGTTCTGGGGTAACTGAAGATTTCGAGGAAGCGCAAATCCAGAGTAGAGGGTTTGGAATTTTGGATGTAGGATATCGATCAAAAGATGAAGCTTGGGCAGCTGGAGGAAGTGGTATATTGCTAAGAACGGCTGATGGTGGAAAATCCTGGATTCGTGACAAAGCAGCAGATAACATTGCTGGAAATCTTTATACAGTAAAGTTCATAGACGATAGAAAAGGTTTTGTGCTGGGGAATAATGGCATTCTCTTGCGGTACCTCGGTTGAGGGGATGTGACCATGGGCTACAACATTTGAGGAAGATGGTTCCCCCAATTTTGTATTTATATGTAAAAAGATCTGGCTGGCTAAAGTTTTGATCTAGATAGATAGAGATGTCTTTTTCAAGGGGAATGATTTTTTATTTGTTCTATTTAGATGAGATGGGGTTTGAAATTGTGAAAAATAGATTTTTGCTTCATGCAAAGAAGCTTAAAAATAAAAAGAAGATGCATCGCATACTGCTGCTTGGTACGATCATAGTCGCACAAATTGATGGCCAAGCTTATTCTTAGATTGGAAAAACTAAAGCTCCTAATTTAATAAATTTCTACAATTATCCGAGAAGGTTAATTTGCTAAAAAAA

**HIGH CHLOROPHYLL FLUORESCENCE PHENOTYPE 173 (HCF173)**:

GTTTTGAACAAGCGAATTGGAGCCATAAAATTGTGAAGAGAAATGGGGAGCCTCTAGTTCTTGATCACGCCCTCACAATCGCTCTCTCTGCTTATCACTCACATCAAACTTTATTGCAGAGTTGCAGCGTTCGGTATGACGCCCATCTTTCTCCTTTGATCTGTGTTTTCCTCGCTATGGCTTCCTCTCTTGTTGCTCGCCTTTCATCCTCACCTGCCTCTACCTCCTCTTCCTTCTTCCCCTTCCAACCCATTTCACAAAGCTTGCCCCTCCATACCCACCATCGCCAGCTCAAATTCCCTTTGTCCGTGCATGCTTCTGCAGGAGATGAACCCCCTACACAAACAAATGGCCCAGAGCCTTCATCAGCATCATCATCTCCATTCGATGCTCCCACATCTACTGTTGCTACATCCAGTAAGCAGCCCCCACCAGTCTCTTTGAGTTTGGAGGATGTGAATCCCGTGGGTCTTGGGCGGCGATCACGTCAGCTATTTGATGAAGTGTGGCGTCGCTTTACGGAGCTTGGACAGCTCACTCGCTCCCCCCGGGATGATGAAGAGTATTCCAGTGTTTTGATCGGCGGACCCATGTGTGAATTCACCATCCCTGGTGCTGAATCCACCACCGTCCTTGTTGCAGGTGCTACCAGCCGTGTTGGTCGAATTGTCATCAGAAAGCTTCAGCTCAGAGGCTACAAAGTGAAGGCCTTGGTACGAAAAGCAGACCCAGAAACTCTTGAAGGGCTACCGCGTTCTGTGCAAATTGCTGTTGCTGATCTCGGTGATCCTTTGACACTCAAGAGCGCCGTGGAAGGTTGCAGCAAGATTATCTTCTGCGCAAGAGCTCGCTCTAGTCTCACTCTAGATCTTAACAGGGTGGATAGGCTGGGTGTCTACAATTTGTCTAAGGCATTCCAGGATTACAATCACAGAATGGCACAAGCAAGGGCTGGACGAAGTAGTAAAAGTAAGCTTTTGATTTCAAAGTTTAGTCAACCAAAGGAGAAAGATGAGTGGACGGTACACCAAGGAACTTACTTCGCTGATGATATCGCTGTCAAGTTTGATGGGGGTATGAATGCCGGACTTGAAATGGAGGAGGATGGGAATGCAGTCTTCTCAGGTTATGTCTTCACGCGAGGTGGTTATGTAGAGCTGGCAAAGAAGCTAGCGCTTTCTCCTGGTGTTTCCCTTGATAGATATGAAGGTCTGGTCCTGTCTGTCTGTGGGGATGGAAAAATCTACAGCTTAATTTTGGAGGCATGCTCCATGGATGACCCTTCTCATAAGATTGTCTACTTTGCAAGGTTCGCCACGCGTCTGGGTTACAGTAGGATAAGGATACCATTTTCTTCCTTCAGGCCTATGAATCCCGAAGACCCAGCTCTAGATCCCTTTTCTGTACATACCTTGGCTATTCGATTTGAACCTCGAAAGCAGAGGCCACCCCCTGTTCGTAATGCTGTTCGTAATGGAAGTGCCACATCAACTACAGCATCTTTTCGAGATCTCAACAGCTTTAAGCTGATAATTGACTACATAAAAGCCTTGCCGGGTGGGCAAGAGACTGATTTCATTCTGATCTCTTGCACTGGAGCTGGCCTCAGCGATGAAGATCGTGAGAAAGTCCTCAAAGCCAAACAGGCCGGAGAGGCTGCTCTCAAGAACTCTGGGCTTGGATACACCATCATCAGGCCAGGACCATTACAGGATGAACCAGGAGGGCAACGAGCTCTCGTCTTTGACCAGGGAAATAGGATTTCACAGGGTATCAGTTGTGCTGATGTAGCTGACATCTGCGTCAAAGCTTTACACAATCCATCTGCAAGAAACAGAAGCTTTGATGTATGCTATGAGTATGTTGCTGAGCAAGGACAAGGATTATATGAACTTGTTGCACACTTGCCAGACAAATCAAATAATTATTTGACACCAGCACTGTCCGTTTTGGAAAAGAACACATGACATCTATGGTGCTTGGGCGACGATCATTGCTGTATCAACAAACAATCAACATCTGGACAACACTTGGTATCTGTATCCATAGAGTTGTTGTGGTGGAACAAGTGGTTGTGATTTTGTGCAAGCATAATCACACCTACATGTTTGTGAAACCAAAAGAATTCTGTCCGCACTGAGCGATATTTACCTCAATTCATTTGTAGCTCTGCCTCATTTTCACAGTGTATGTATATAAGGGCTATCTGTACATACAGGCTCAGCCAAGGAGGGATTAACAGGTCAGATGCCAATTCTATTGAGGTATTTGTATGCATCAATGTGGGTAGTATGGGTTTACTCCCACTTCCATGTATTCTTCCAATTCTGCGTCAGCAGTATATACACCCGGCAAGCCAATATTGAATGGAATTCAGGTTCTGAGATCAATTTGAATGTAGCTGGCAAGTTAATAATTAATGGAATATATGTTCTGACACCAATTTGAGTGCAGATTTACGAGACTGCCATGTGAACAATGTTTAATGCTTGGTACTCATTCCACAATATGAACCAGCGGCACCTGTTTTCGACCTTGCAGTAAAGAGTATACTGGCATAAGTGACAAGTATTGCCTTCATACAGTATCTTCATCTTTATGGTTGCCAGACAAATGAATGCAGAACTGGTTATGTACATTGACAATACATGCATATGAGAGCTGACACATCATCTGATTGTTGCAGTGCTAGCTTGAACAAAATACTCCAAACAAGAGAAGCCTCTGGTGTTAGATGTTAGGTTTACTAGCCTTAAATTCTCTCATCCCAGGCAGATGGTATAATTCCATCTGATTAAGCCATGAAAGTGATATCAAGGCCAGAGAGGAGTTCTCATGGTGGAAGGACAACCTAGTGTAGAAATGTAATTCTTGCACTCATTCACATGCACATCCACACTCTCTCATTAATCCAGTTGATGGATGGTTCAGATTAGTAGCTTTGGAGATAGCCACCCACACCACTTTCAATCCTCATTTACATGCAAAGGTAATGAATTAGACTTTTGAAACCAGGATGGATGACGACAACTTTCAAACTGTTGTCCCCCAAGATTTGGGCTAATGGTAAAGCTTTGGCCTCGTAACCTCTAGGTTGTGGGTTCAA

**HIGH CHLOROPHYLL FLUORESCENCE PHENOTYPE 244 (HCF244)**:

GCGTTATCCACTCTTGAACCACATCAAACATGAGGGTGGACAGCCACCCAACTTGAAAATGTGAGTAAAATACGCTCTTCCTTACAGCGGCGCTAACCCACACACACGCTAGTGGGCGCTTGAAGGGGAGCAAGCAGAGAGGAGGGAGCTATACGTTGCCCCTGTGTTTGCGTGATTATTGTTGGTGTGTTCGTGTGTATTGTGAGTGAGTGTGTGTATTGTGGGTGTGCGTGTATTTTGCAGCTATGGCAGGCCTGATGGTTGCCCAGGCCCCCACCTTCCATAACCTCTCATCGTCTCCCCCTGCCCACTCCCCCCCTCTGGCTTCTTCAACAGGATCGCCCTCTGAGCTCTCCATCAACTCTGCTTTCTGGGGCTCCATCAAGCTTAACTGCTCTCCTTCCCCTGCTTTTTGCAAAGCCTCCACCAAAGCAGATAGGCGAGCAGCCAGTTTATCAGTGAGATGTGATGCCAACAAGGTCATGGCAGCAGCAACAGCCATGTCAGGGACTCCTGTAAGGTCAACCAGCATATTAGTAGTAGGGGCCACTGGGACTTTGGGTCGTCAGGTCGTGAGAAGGGCTTTGGACGAAGGCTATGAAGTTAGGTGCCTTGTACGACCCCGCCCTGCACCTGCTGATTTTTTGCGCGACTGGGGTGCCACTGTTGTGAATGGTGATCTAAACAAACCAGAGACATTGCCAGCAGCCCTTGTAGGAATACATACAATCATCGATTGTGCAACAGGTCGACCTGAAGAGCCTATACGGACGGTAGACTGGGATGGAAAAGTGGCTTTGATACAATGTGCGAAGGCTATGGGAATCCAGAAATTTGTGTTTTTCTCTATCCACAACTGTGAAAAACATCCGGAAGTGCCCCTAATGGAGATCAAGCGCTGCACAGAGAAATATATTGCAGATAGTGGCCTCAACTACACTATTGTGCGACTGTGCGGCTTCATGCAGGGCTTAATTGGACAGTATGCTGTACCTATATTGGAAGACAAGTCCGTTTGGGGAACAGATGCTTCAACTCGAGTTGCTTACATGGACACCCAGGATATAGCACGGCTAACTCTCTCAGCATTGAGGAGTGATAAGGCAGATCGAAAGCTCTTTACTTTTGCAGGTCCTCGGGCTTGGACAACACAAGAGGTGATCAGTCTTTGTGAACGGCTGGCTGGTCAAGATGCGAAGGTGACTACTGTGCCCGTCGGAATATTAAAGTTAACAAGAGCAGTAACTCGCTTGTTTCAGTGGACCAATGATGCGGCGGATCGGTTAGCTTTTTCAGAGGTGCTAACGAGTGATGTCGTCTTTTCTGCTCCAATGACGGAAACGTACTCCGTCCTTGGTGCTGATGCCAAAGACACCACCACTCTTGAGAAATATCTTCAAGAGTACTACACAAACATTCTCAAGAAGCTCAAAGACCTGAAAGCTCAGTCCAAACAAGGAGATTTCTACATCTAAATGTAAATGTGAGACTTGTACATCAAAAACCCTGGGTTCTTGTCATTTCCCTTGTACTTGTGTAAGGACACACATCATTGTACTTGTAAGGACACATCATTGTACTTTTGGACATTGGGTTAGCATATTGTTTCCGAGTTTTCTCTTTTTCTGAAGAGGCTCTAAGCCGACGAACTCCTTCCGTGTTCTATACTTCTGATGTATCATCATGCTTCCTCCAAACCTGTAATTTTGTCAGCTTGTCTTGATGCATCATTTGAATAGTTTTTTGAGCTAGCAACTTCTTTTGTGAAAAAAA

**CYCLOPHILIN 38 (CYP38)**:

GTTTTTTTGGAGGAGGGTTCCGGAAGGAGGGCATGTGGCTAGTTTAAAGCAATGGAGTGGGGACTTCTTATCCACTATCTGGCTGAGGTGGCCAAATGCACAAATGAGAAATTTAAACATGCAAGGCGAGTGATGGTGGAAGCAGAGGAGTGTGCTTGATCGCTCTCTCTGCTAAGCCAGCACACGAAAGAGGAAGCCAGGGTTAGGATTAGGTTTAGGGTTCTACCATGGTTCCCGCTACAGTTCTTGGAGCTCCCTCTTCCTGGGTTTCTGCAGCAGGTCGCGCCTCTCCCTCCTGTGAAGGAGCCGGCGCGCAATCTTTGCTCCCCACGTGCTTGCGATGGAGCGAGCATTTCTCTACGGAGATTTCTTTGTTAGGTCGAAGCAGCTTCAGCTGCAGGAGCTCTGCTAGAAGGACCTCGCCTTTCACAGCTTTCGCCCACTCCAATCAGGGGAAGGAAGAGGAGGAATGTTGTTTAATGAGGGTTGTACAAAAATGGGCAGTATGTTTGATGGCATCAGTAACGTTGGCTTCCGTGCAACCCTCATCCTTGCCAGCACAGGCACTGGTTACAAATCATATCTTTGATGATCTCGCTATACTTATTTCAGGGCCACCTGTGAAAGATCCTAATGCTTTGTTAAGACATGCTCTACCTATTGACAGTAAGCCGTTCAAGGACATACAAAAGTCACTCGAAGATATTACAGACAACCTGAAGCTTGCAGGAGCAAAGGCATTTGATCCAGTTGGACGGAGTGTTAGACAAGCTGAAAGGGCGCTTAGCCAGAACAAGTCGGAGATAATTGCCAATCTGGCTCCAGCAAAGAAGGAAACTGGCCAGGCTTTATTGGATAAGCTCAGTGTAGGCCTTCAAGAGTTGCAGAAGATTGTGGAGGACAAAAGCAGGGATGAGATTGCACCAAAGCAGAGGGAGCTTTTGTACATTATGGGAGACTGAAAGTGTTCCTGGGGCGTTGCATTTCTGCAGTGTGGAAGAGGCTATGGTGGACAAGTTCCCATTTGAGGTACCGGAGGAGTATGCCAACAAGCCTCTATTGAAGGGTAGAGCAACAGTGGAGATGAAAGTTAAGGTCAAGGACAACCCGAATTTGAAAGATGTGGTTTTCCAAATGGTGCTGGATGGCTACAACGCCCCAGTCACTGCAGGGAATTTTGTAGATCTCGTTGAAAGGCATTTTTATGACGGCATGGAGATTCAAAGAGCGGACGGCTTTGTAGTGCAGACTGGTGATCCCGATGGACCTGCTGAGGGCTTTGTAGATCCTAGCACGGGGAAGATCAGGACAATACCCTTAGAAATAATGGTGGAAGGGGACAAGCAACCTATTTACGAAGCTACTTTAGAGGAGCTAGGGCGCTACAAGGCCTCGACAAAGCTCCCTTTTAATGCTTTTGGGACCCTTGCTATGGCACGAGAGGAGTTCGAGGACAATTCAGCTTCTAGCCAAATATTCTGGCTTTTAAAAGAGAGTGAACTCACTCCAAGCAATGCCAACATCTTAGATGGGCGATATGCTGTGTTTGGCTATGTTACTGAAAACGAAGAGCTTCTTGCTGACTTGAAGGTAGGAGATGTCATTGAGTCCATCAGAGTTGTAGCTGGACTAGACAATCTGGTCAATCCAAGCTACAAAATCTCTTGAGTTGCATTGCTAGCATTTTGGGTGGCAGTACATTCCTGTAAGATGCAGTATATTCTTGTGAGATGCGGTCTATTTTAATGTGTTGTGCCTCTCTTAGAGGGATGGCATTTTTGCTTGAAAAATTAAATTCTTGAAGAAGTGTGTTTTTGCAAAGCTTTTCAATTAAGCATCAAGTGTGTTTTTGCAAAGCTTTTCAATTAAGCATC

**LIGHT-HARVESTING CHLOROPHYLL B-BINDING 2 (LHCB2):**

TTTCAGGTCTAATAAGATGGCGATGAACATTTCAGGGGAGCAGTATCGAAGCTTGAGAGAGAGAGGAAGGTAGAATGGCCACCATGACCTCTGCATCTGCGATTAGCAGCTCTACATTTGCAGGCCAAACGGCATTGAAGCCCCAGAGTGAACTCTCGAAGAGAGTCGGCAATGTGGATGCTCGCGTTACCATGCGGAGGACTGTGAAGAGCACTCCCGAGAGCATCTGGTATGGCCCTGACCGCCCTAAGTACTTGGGCCCTTTCTCTGAGGCAACCCCTTCGTACCTCACCGGTGAGTTTCCTGGCGATTACGGGTGGGACACCGCAGGACTTTCTGCAGATCCGGAGACTTTTGCCAAGAACAGAGAATTGGAAGTGATCCACTCGCGCTGGGCGATGTTGGGTGCCTTGGGATGTGTCACCCCTGAGCTCTTGGCCAAGAACGGTGTGAAATTTGGAGAGGCTGTGTGGTTCAAGGCTGGCTCACAGATCTTTGCCGATGGTGGACTTGACTACTTGGGCAACCCGGGCCTTGTGCACGCACAGAGCATTCTCGCTATCTGGGCATCCCAGGTTATCTTGATGGGCGCCGTGGAGGGATACAGAGTTGCAGGAGGGCCTTTGGGAGAGGTTGTAGACCCCATCTACCCAGGCGGGCAATTTGACCCCCTGGGCTTGGCTGATGACCCAGAGGCATTCGCAGAGTTGAAGGTGAAGGAGTTGAAGAATGGGCGCCTTGCCATGTTCTCCATGTTTGGGTTCTTCGTTCAGGCCATTGTGACAGGCAAGGGCCCCATCGAGAACTTGTCCGACCACTTGGCCAACCCCGCTGTTAACAACGCCTGGGCATACGCCACCAACTTCACCCCCGGCAAGTGAAAATTTGCTCCTTTGTAGTCTATCGCACTCGGGTAGCCGAAGCTTTCGCCTCCCTGCTTGTACCATAGTCTTTAAGTTGATTGCCTTTTCTTTAGTCTTTTTTCTTTTTCTACGACATTGTTTGCCAATTGTGAATGCGGCCCTTTTGCAGCAGAAACGTTTACACGCATAATTAGATTACACTGCACCAGTTTATTATTTCATCTAGTGTTTGTTGGTTGTATTTTGGCCCTTGTAAATGAGCTGCGCTATTGCACCTCCAACCACCCTTTGGAGGTGCATTAAGCCAATCACTGCGCTGCAACACTTTGTTAATCGTGCGTGAGAAGCCGGGGCGCATGGAGAAGCCTGGGGGCATGGATAAGTCTGAACAACCTAATGGGAAAAAAATCCAATTTAAAATGAAGTTCAAAATAAAATGTGCACGAGTGCCTACCGTTTGTAGAAAA

**PHOTOSYSTEM I LIGHT HARVESTING COMPLEX GENE 1 (LHCA1):**

GAGAGAGAGAGAGAGAGAGAGAGAAGACTGACAGGCAAGATAAAGCTTGATTTACAGTCACTCAATGAGAAAAGCATCCAATTGAAACTAAATTGCATAATGCTACACAACCGTGGAAGACTTACTGAAACCGGAAGTAGCCCGGTCGGTAAAGTGTTTCAATGTTAAGTTCTAGGTGTAAGCTCGAACGTGTGATGGCACTACTCCGAATGATGCTCAATTTTGGTTATGGGTGGCCATTCGAGACCTCTCCATCAAAAAATAAAAGACCGCCCACGACATAGTATATAAGCACATTTAATCTTTTAAAAATTCCTTCTTTTCTTATGGTTAAGCTCCAAAGTTATGGGAAAAACTTCCTAGTAATTGAGGGCATTTAAAGGTAGAAAGACAAATTTTCAAGTACCACTCTCAAAGAACCAACTCGAAAATAACCACATACAAATGAGGCATATGCTCACACACTGAATGAATAAACCAACTTCTCGTCAGATATTAAGAACAAATGTCGAAAGCCATGTGCACTTCAAACACTCGGTGGGTCATGTGAGTGAAGCCAATTTACAATACACTCCTGGGAATGATAATGTCTGCGATATTCTTGTGCCATGGATCGGCCAGGTGGGCAGCAAGGTTCTGGAGGGGGCCTGTGCCTGGGTAAGCAGCTGCCTGAATCGAGAAGCCCACAAATGCCAAGATGGCCAATCGACCATTTTTCAGCTCCTTAACCTTGTACTGCTTGAAGGCTTCAGGGTTTTTGGAGAAGCCAAGAGGGTCGAAGGGGCCACCAGGGTACTTTCTTTTCTCAATGTCTGTCTCATCATTTCTTTGGGCCTCAGCGAAGGCAATGGCAAAGATTTCGAAAGCGAGGATGAAGGGCAAGTTCCCCCATGGCACAGGGTTGCCCAAGTAAGTGGCTTGGCCTCCCGGAGTTGCCGCCCACTTCTGTGCATCTACCCAATTTCCAAGACCCAAAGCCTCGGGCACAAGTACACCGGGCACAGCGAGCATTGCCCATCTGCAGTGAATGAGCTCCGACTCTTTGAAGCGTTCGAAGTTCTCTGGAACCTCAGCCAAGCCTAGAGGGTCAAACCCAAAATCTCCAGGGGAGGAGCCATCGAGATGGGCAGGTCGAGGCTGGCCGGGAAACCACTCAGAGGAAGCCATGAAGACGCGAGAGCCATTGGAGCAGGCGAGCGCGGGTGTGAAAGAGGAGGCGCCCAAGAACTTGGACTTGGAGAGAGGCAGAGAGAGGGCGGGCACCAAGTGTGAGCACGCCACGCTTGTAGACGCCATTGCAGCTTCTAGACCCGCTTCACGCTTGCTCTGCGCACCCACACTCGCGCGATCTAGCCTCTCTTGCACCTTCACCCGCGTCAACTTTCACTCCCTCTTCCTTATTTTCCGTGGATATCTTCTTGTAAGATAGGCCCAATCTCTCATGGGCGCTTCTCATTGGCTCCTTTGTGGATGTATGCTCTCTCTCCACCACCAATACTTTTATTTCTTACCTTAGCTCTCTCAACCAATATTATTGTTTATTTTCTTGACAGGACCCTCCTTATTGGCATTGTGG

**LIGHT HARVESTING COMPLEX PHOTOSYSTEM II (LHCB4):**

GGAAAAGTGGGCAATGGGAGGGTGCCAAGTAAGCAAGGAGGAGATAGAGCATGGGATTCTCATCCACAAACGAGAAAGATAGGGTGTTGTATAATATGGATAGTGTGTGTTTAGAGTGTGCACGGTGAAGCATACTCATATCTGATCCGTTGGTGCAGAGAATTGAAGAAGCTCGCGAAAGAGAGAGAGAGAGAGAATGGCAGCGACTTCTCTGGCTTCCTCTTTCTTGGGTACTCAGGAGCTCGTGAAGAGCTTCGGTGTGAGCAGTGGGTCGCGCGTGGTCTGCAAGCTGGGCTCCAAGAAGAGCGCAGCTAAGCCAGCGCCCAGAAAGAGCAGCGCAGGAAGCAAGGATAGGCCTTTGTGGTACCCAGGCTGCAAACCCCCTGCGTGGCTAGATGGCTCTCTTGTGGGTGATTATGGGTTCGATCCCCTCTCTCTCGCCAAGCCCGCAGAGTACCTGCAATATGACTTCGATTCCCTCAACCAGAATCTTGCAAAGAACCCTGTCGGCGGCCTTATCGGAACTAGGGTTGACAACCCTACCAACATCAATGCTACCATCCAGCCCTACGCTGAAGCCTTTGGTCTCCAGCGCTTCCGAGAATGTGAGCTCATCCATGGCCGCTGGGCCATGCTCGCTACCCTCGGCGCCCTTTCCGTCGAGGCCTTCACCGGTGTTACCTGGCAAGATGCTGGAAAGGTGGAGCTTGTTGAGGGAGCTACTTACTTCGGACTGCCCCTACCCTTCTCCATAACATCCCTTGTATGGATTGAAGCCCTCATCATTGGATACATTGAGTTCCAGAGAAATGGGGAGCTAGACCCGGAGAAACGCATCTATCCAGGTGGCAAGTTTTTTGACCCCTTGGCTCTTGCTGCAGATCCAGAGAAAAGAGAAGTGCTCAAGCTTGCAGAGATCAAGCATGCCAGACTTGCTATGATTGGTGCCCTTGGTTTTGCTGTTCAAGCTGCTGCCACTGGCAAAGGGCCTTTGGACAATTGGGCTACCCATTTGAGTGACCCCCTGCACACCACCATCATTGATACCTTCTCTAAGTGAGCAGGGTCCATCTCTCTCTCAAAGCATCCTATGCCTCCCTAATGTACTATCCAGTGTAACTAGTTTTTTGAATGAATACTGAATTTGTTTACTTACCAACATGACAATGTAGGGATGTGGTCTGATGCATGCAGCCTGAAGCATTTGTGGTTGCATTTAAAATGAATTGCTACCTTGTTTTTGTCCATGTTCAAAAAAGTTCAGTGCAGAATTTTTGGTAAAGAGCCATGTGTAATAAGTTATTCATGTAATTTTTCTACCAATTTGGGCTAGCTTGCGTCCATGTTTGAGTAAAAAAA

**SUPPRESSOR OF QUENCHING 1 (SOQ1):**

GGCCACGTAGGATATCTGTCTTGAATCATACAACGTAGTCGTTTATCCACAAGCAGCAGTGCATATATCATACATGTTGTACTCGCGCACTGCCACCCCTTCTCTTCCCAAGCTTTCCATGGCTTCTTCCTCCTTATCTCCTTGCTTCTTGTCCTCGTCATCTTCCTTCCTTCGCCACCGCTCACCCTCCTCTGCACCGTGCCACCCTCTTCTACAGCATCTCTCGTACCCATCACGGCCTTCTTCGCCTTTCACATCCTGTTATGCGCCCATGCATCGCATGTGTTCTTCCACGCAGCTCATTTGCGCTTTCTCAGCTTCGCCTGCTGGTGCCAGCACCTCCAATGACGAGCGCACGAATGCCTCCGATCGAGTCTCTGCAGTGCTTTTCGACATGGATGGTGTCCTTTGCAACAGCGAGGGGAGGTCGCGCTTGGCCGGCGTGGACCTTTTTGCAGAGATGGGTGTGCAGGTTTCTGAAGCAGACTTCTTTCCTTTCATGGGAACTGGTGAAGCCAACTTTTTGGGGGGTGTGGCTCAGGTCCATGGAGTAAAAGACTTCGATGTTTCTTCTGCAAAATCTCGTTTTTTCCAGATATATTTAGAGAAGTATGCAAAGCCTAACTCTGGTCTTGGATTTCCTGGAGCATTGGATTTGATTATGCAGTGCAAAGCTGCAGGTTTGAAAGTGGCACTTGCTTCTAGTGCGGATCGACTCAAAATTGATGCTAATTTGGCAGCTGCGGGTCTACCTTTTTCTGTCTTTGATGCCATTGTGTCAGCAGATGCATTCAAAAATTTAAAGCCATCTCCTGACATTTTTTTAGCAGCTGCCAAAGCCCTTGATGTTCCAACTATTGAGTGTGTGGTTATCGAGGATGCTTTAGCAGGTCTCCAGGCTGCCAAGGCAGCAGGAATGAGATGCATTGCTGTCACTACGACTCTGAGCGGTGATAAGCTTGTCCCAGAAGGTCCATCTCTTATCAGGAAAGACATCGGCGACATACAGCTGGATGACATCCTCAAACTGCAGGGATCTGGTAATTCTGAAACATATTCTCAGTGTGAAGTTGTCCCTACGCCTTCAGCACCAGCTTTAGGCATTCTTGACGAAGAGGTCCAACTTCCAGCTGGGTCAGCTGTGACTCGTAGGAATGCCCTGAAATATGGAAGCCTTGCAGTTGCATTGTCGACCTCTTACATCGCTCTCACACATTGGAAGGCTATGTCCTACACATCCCCAGAGGCTCTTCTCAATGCATTTGTTGGCCTGTTTCAACCATCAATAGCAAGTGGGGCGGATGAGTCAGATCGTGCTGCAAAATACTTGAGCTACATTAATGATTTGGAAACTCGTGGGGGAGGACAGCAGGTACCGGAGTTTCAGTCTCAATTGGAGTGGCTTAACACAACTCCTCTTAAATTAAAAAAGGATCTGGTGGGAAAGGTGGTGGTTTTAGATTTCTGGACATATTGCTGCATAAATTGCATGCATATTTTGCCAGATTTGGAGTACCTAGAGAAAAAGTACAAGGGTAAACCGCTTGCTGTGATTGGTGTTCACTCTGCCAAGTTTGACAATGAGAAGGATTTGGGTGCCATTCGCAATGCTGTTTTGAGATATAATGTGACACATCCGGTTGTAAATGATGGTGATATGGTGTTGTGGCGAAAGCTGGGTGTAAACTCATGGCCCACTCTTGTGGTTGTTAGTCCTCGAGGAAAAGCTCTTGCTGTAGTGTCAGGGGAAGGCCACCGCAAGGATCTGGATGAGTTAGTGAGTGCTGCTCTTCAATATTATGGAAACAAAAAACTCTTAGATGATCGGCCTTTACCCGAGCTTTTAGAAAAAGACAAAGACACACGCTTGGTTTCCTCACCCCTGAAGTTTCCTGGCAAACTTACTACAGACCTTGCCAATCAGCGACTTTTTATTTCCGACAGCAACCATCACAGAATAGTGGTTACTGATCTTGAGGGGAACTTTATACAACAAATCGGTGGTACTGGCGGTGAAGGCCTACAAGATGGAAGATTTGAGGAAGCTGCCTTTAACAGACCTCAAGGTGTAGCTTATAACTCAAGTAAAAACGTTTTATATGTTGCAGATACAGAGAATCATGCGTTAAGGGAGATTGATTTTGCCAATGAAACTGTGCGCACACTAGCAGGAAAGGGGGTGAAGGGATCTGATTACAGAGGTGGAAAACAAGGAACATCACAGGTTCTCAATTCCCCTTGGGATGTCTGCTGTGATTCTGCCTCAGGCACTGTCTATATTGCTATGGCAGGGCAACATCAGATTTGGCAGCATACTTTATCAGATGGTATCACAAAAGTCTTTAGTGGGGATGGCCGCGAACGCAACCAGAATGGTAAAAGGGGTCAACAAACATCATTTGCACAACCGTCTGGCCTCTCTTTTTCTGCAGATATGAAAGACCTTTACGTGGCAGACAGTGAGAGCAGCTCTGTCAGAAAAGTTGACTTAGCTACAGGAGGCTCAAGCTTACTTGCAGGAGGGGACCCAATTTTTGCAGATAATCTCTTTCAGTTTGGTGACAAAGATGGCACTGGATCTGCAGCTCAATTTCAGCATCCGTTGGGTGTACTTAACGGTGTTGATGGCCTTGTCTACATTGCTGACTCTTATAATCACAAGATTAAAACTATGGATCCTAGCAGCAAGACGGTTAAGACAATAGCAGGCACGGGAAAGGCTGGGTTCAAGGATGGAACAGGCTTAGGATGTCAGCTATCTGAACCAGCTGGACTTGCAATTGGGCTTGGTGGGAAGTTATATGTGGCAGACACAAACAATAGTGTCATTCGAGTTCTAGATACCAAAGGCAATAGCGGTCCATCAATCTCTACTCTTGAATTAAAGGGTGTTCAGCCTCCCCAATTGAAGACAGCTGGAAAACCAAGACGCCTTCGACGTAGATCTTCAAGTGACATGGATGTGATAAAGACAGAGTCAGTTAAAGGCAGCAATGGAAGTATCAAGATAAAAATTGCTTTGCCACCTGGATATCATTTCACTAAGGAAGTCACTAGTAAATTTGAAGCAGATGCTGAACCTGATGGCAAGGTCGAGTTAGACCCATCTGCAGGTACATTTGATGCCAATGGACAAGCTGTTATCGGATTCTCACGATCTCAGGACGCATTGGCAACTATACAAATCAACTGCAAGGTGTATTACTGTGAGGAGGATGAGGTGTGCCTCTATAAAGGGCTTGCCTTTCAAATACCTTGTGATGAGGTTTCCTCATCATCTTCAGTGGTTTTGCCACTGACATATAATGTAAAGCCACCGGTTCGACAAACACTATAGTAGACATTTCAAAACGCCTAGATTATTTTGGTAGGTATTGAAGGGAAGTCTGATTCTTTTGTATAATATCCATTATGTAAACCAAAAGAAAGCCTTTTGTAATCATATTCAATGTAAGAACACCATTATTGCGTCATGAAAATCGACAATGAAAAAACTCCATCTCTGATGTTGCGCAAAAACCGTGTTAATCTATCTGATGTGGTTGTGGC

**ENHANCER OF VARIEGATION3 (EVR3)**:

AGAGAGAGAGAGAGAAAGAGAGAATGGAATCAAAGAAACTACTTTTGTTGACCAGCTGGATCTTGCTTCAAGTTATTTTACAAGGCCCTCTGCCGAGTGACCTGAAAATTGTGTAATTCGTACAAGGAGATCATCAATGCTTATCTGCTATTGAGTGTGTATAGTAGAAGACTGCGATTACTCTTACGCAGCTCATCTTGGGGTGCAATTAGCTGTAACTACGTTGAAATGGAAAAACACTACCCAGGTTGAACAATGTGAGCTATGTACATTTACCTGCCTTAGATAAGAGGGGATCCCAATCCGATACCAAGCTCTTCAGCAAGTCCATCCCAAAGAGGTAACAAAGTGAGAAGAACTAGTATAAAAGCCACCGCAGATGCACCCTGCCTCCATGTCCCCACCTCAGTAACATCATTCAAGCATGGCTTCTCAGGTTCTCTTTGGCAGATCAAAATGTAAAGCCCCCATGGAAGCGAAAGAGGTCCTCCTAGCACTCCAAGACCTAGAAGTAAGTATGTCCCCAATGCAAAGCCTGTCAAAGCGCCCTTCCCAAATGCTGCCTGCATTGCCCTTCCACCATCTAAACAACCAACAGGAAGAAGGTTGAATGCTGTTGTGGTTAATCCGCACCAGCCTGCAAGGACCAGAGGATGAATGGAGACCGTAGCTGCATGCATGGCATCATAGCCAAGCACACTTCTGCTAAAATACCCTAGTAGTAGTGATCCTTGAAATAGCAAGCTGGGCACTTGTACCAAATCGCTGGCAGCTTCGGGGCTTGTAGAGAGCAAAAGCCCCACCCCGAGCATGGCAAAAGACAGGAGTCCACCTGCCAGGGGGCCCGCTAATGAAATGTCAAATTTTGCAGTGCGGTCAGGAAGGATTGATCTGAACTGTGTGATGGCCCCAAAACTACCAAGAGTTATGTTGGGAATAAAATACGGAATTCCAAGCTTGACTTTTTTTCGAGAAGCAGCAAGAAAGTGTCCAACCTCATGGAAGAGTTGTACTCCAAGAACTCCATATGTAAGAGGCATAGCAGCCTCCACATATGGAATCAAAGTGTTCAGATCTGGCGGTTCCACAACCTCTGGATTTGTGAAATACCGCACAACTTCAGGCGGCAACTTTGAAAGCTGTGATGCGATTCCAAGCTCCAGACAAGAGCCTGCTGTTGTGACAAAGAGTACAAAAGCTATCACAAACTGCCACAAGGTTGTGGACTTCGGTTCAGAAACTTCTCTACGTAGTAGCACAAAACTGACTCGAGGGCCTCCTCGAGGATCTAATTCTTCTGAATTTGGTTCCTCCACCATAAACAGAGTATACTTAGAGCCAGTCACCTCCCGCAATTGCCGTTGAAGCTTTGCAAAGACTTCTTCCCTTTTGCCTCTTAAATTACCTAGAAACAGAATTCCTGCACCTAGATCCCCAAAAGGCTCTTGCCCAGTTACCCAAAAAGTGCTATAGCCAAAGACTCTCTCCTTCAACAGCTTCACATCTTCTGGATCTACTTTCTCAGGACCAAGCAGATCCAAAAGTTTAAAGGCATCCATTTGTACATTGGCATGTTCTGGTGCACGGGTGTGGTCTGTCTCCTCAGAAGATGGTGTCGAAGGCTTTGGTTTGGACTCATCAGGCTCTTCTGTCGCCACCGTCGTAGAGTTTACGCTCTCTTGCTCATCATCAGGCCCAGTGGCATGACACCGGAACCTGCTCTCATTCTTAAATTTAAGGCCGTCGAGATTTCTGCAAAAATTTCTAAACCGAGAGCCCCCATTACCGCAAAGCTTGCAGGTTGCAGAAACTCCTAGAGATGCCTTTCTGAAGCTGAACTGTGCTTGTCGTTGCAGCTGATGGTGAGAGCAGGCGATGGTTGTGATGGGTTTTCTCAATTTTATCAAGCAGCAGCATCTGGCGGGAGCGTCTTCTGAGCTGGATATGGTGGAGGGCAATCCATAATAAGCAGAGCAGAGAGAGACGCAGGAAGACAAGGGATGAAAGCAGAACTCGGCGTCTGCAGGCTTGCAAGGTCTGTTCAGCGATGGAGCTCGGAGAAAGAAGGGGGCACCACCAGACAAGCTTGTGGTAGCAGCCATGTCTATGCAAACCTTCCTTGAAGGCCTGCTGCCTAAGAAGTTGATCAATGGATTGTCTTACATTTTGGTCACACTCCTATCA

**FTSH PROTEASE 1 (FTSH1)**:

CGTTGCTCTTCCCATATACACCAGGGGGTCAATGATGACTACATATCCTCCCAAAACATTTCTGCATGTGAGATGATAGCATTTGAAAATATACATACCATCTGCAACTACTAAAAACCATTTGGGCCCATCAATTCAGACACCTAAAAATGATCACTCAATATGACCTTGTCTACAAAAAGGTTAAAACTGCATGATCAAATCCCATTGAGCTTGCCAACTATCAGGGTTTGTAGATACATGATGCATTAAAAGAATGATGAGGCCGTTTCCAACATATACGTGGAAACTTACAAGACCAGAAACATTTAAGGCTGGTTTTGGCACACAAGGACCCATCAAGTTTGAAATTTGGCAAAGGGGCTCAAGTACTAGAAGATAAATCTTTGTTTGAGATTTAGTACACATGACCAGCATTTAATTTTCAATAATAATTTGCTCTGGGTACCTTTTTTCTAGGGCCTAGTCAAACAGAATACACAAAATAACACGAGTAATTTCTCTACATATAACATTATGAGCTAAATCATTGACACAGCATTAAGCGACGAACAGCTCAGCCTTGCTGTCCATGAAGAGGCTCATGAATTCCTCACCATCCACACTCTCCTTTTCAATTAGCAAGTTGGCCAGCTTGTGGAGAATGTCAATATGCTTGGTAATGATGTTTTTCGATCTAGAGTAAGCCCGTTCTACAAGCTCCCGCACTTCAGAATCAATCACATCCGCAGTGGCCATCGAATAATCAGATTGTTGAGAGACTTGTTGTCCTAAGAAAGGGTTTCCTCCTCCTCCTCCAAGTGCAACCTGCCCTATCTTCTTGCTAAAACCAAACCTCTCCACCATCTGCCTTGCAACCCTCGACACTTGCTGAAAATCGTTAGATGCACCAGTAGTCACATTATCTGCCCCAAAAATGACCTCTTCTGCCACCCTTCCACCCAAGGCTACTGCCATCTGATTTTCCAAGTAGCTTCTACTGTATAGCCCGGACTCCAATCTCTCCTCACTGGGAGCAAAGAAGGTAAGCCCTCCAGCACCACCACGAGGAATAATGGAGATTTTGGCCACGGGATCATATTCTGGCATAAGCGCGCCAACAAGAGCATGCCCAGCTTCATGGTAAGCAACAAGTTTCTTTCTCTCTTCCGACACCACAGCATTTTTCTTCTCAGGCCCAGCAATTATTCTTTCAAGTGCATCCGCAATCTCATCCTTGCTTATCTCTTTTAAATCTCTTCTTGCAGCTAGGATTGCTGCCTCATTCATGAGATTCTGCAGATCAGCCCCAGTAAAGCCAGGAGTCCTCCTGGCAATTTTATCAAAATCAACATCTTTGGCTAACGCTTTTCCCCGAGAATGGACCTGTAGAATCTTGACTCGACCTGCTACATCTGGTCTGTCTACTGTAACCTGCCTGTCAAAGCGACCAGGTCTAAGAAGAGCAGAATCAAGGACATCGGGCCTGTTTGTAGCAGCAAGTACTATAACTCCACTGTTTCCAGTAAAGCCATCCATCTCAGTCAGCAGCTGATTGATGGTTTGTTCTCGCTCGTCATTGCCACCACCGAGACCTGCACCTCTCTGTCTGCCCACAGCATCAATCTCATCAATGAAGACAATGCAAGGAGCCTTGGCCTTGGCCTTTTCGAATAGGTCTCTCACACGGGAGGCTCCCACACCAACAAAGAGTTCTACAAACTCAGATGCCGCACATGAGAAGAAAGGAACACCAGCTTCCCCTGAGACAGCTCTGGCCAACAGCGTCTTTCCAGTGCCAGGAGGTCCAACAAGGAGGCAACCCTTAGGGATTTTAGCCCCTAGAGCAGTATATTTGTCAGGATTCTTCAGAAAATCCACCACCTCTTGCAGCTCGAGCTTGGCTTGATCAGCGCCAGCCACGTCACCAAATGTCACTCCCGTTTCGGGAACCTCTTGAAACTTGGACTTAGAGCGACCAAAGTCCATGGGGCCACCAAACCCACCCGGGCCACCTGGACCACCCTGAGCACGCCGGAAGAGCAGGAAGAGACCACCAAATGCAAGGATGGGGAACAGAAGGTTACCTAGAATGTTGAAGAGGCTATTATTGGCCTCACCTTCAGAGACAGATATGTCGACCCCATTCATGGCAAGGATGTCAACAAGGTCGGGATCATTGGGAAGTGTCACATTCGCCCTCCTCCCATCCACTGCAGTGAGTTGCAGAGTGGTTCCATCCTTGCTGAACCGCACCCTCTCCACCTTACCCTTCTTTACTGCATTCAGAAACTCACTGTAGCGCCACTGAGTACCCTCTGGAAGATCTGACTTCACCGGTGGTGCAGAAAGCGAGCTTCTAGTGAAAGGATTTAAGTTTTGCTCCGCTTGCTGTGCCTGCAGGAGCTGTGGGGGCTGAGAAGGCGGGGGTGACATGTCTGCAGCTATCGCGGGAAGTGACGAAAGGGCTGCCAGGGCAGCAATGGCGGGTAGCTTGAGAGAGGGACATGCTGTTTCCGATGGTTTCATAATGGTAGCTCGCGTGACAGAAGCTCGACAGGTCCGGAGGGGCCGATGAAGAGGGATGAGAAGGGAGCTGCCATGAAAGCTTGAGCGTTCCAAAAAGGGGTGAAAAGGCAGAGAACTTGCAACCGCCATGGCAGCTGAACAAGCCATCTTCGACTTCGAGTTCGTGCACGCTTGCTCCTCCAAAGAAGAAGAAAATAGAGCAGTCTATCTGCC

**HYDROXYPYRUVATE REDUCTASE (HPR)**:

TTTTTTTTTTGCAGATAAAAATATTGAATCTCCATTTGAGCAAAAGTGCACTGAATGTGGTGTGGATGGTCAGAGCTCTCTAGCTGTCTGATAGCAGAATTTCCATAAGAAGATTTCTCTACATTCCTTATTTGGACAGTTTAATGTTTCGAAGCTGTCCTTTTGATTCGGATAACACTGACACACGGATATTTTACAGTACCATATGCAGCTACATAAGCTTCGATAAAGATTAGTGGATTGGATGCTGAGGAACATCACAAACTACACAATTTCAAACATTGCATTCACATCACAAATTGGAAGCTGCCGAGGTTTTGAGACCTAAGTTTTTGGCATTGACAATGCTTGGGCAAGCAGCAGGCGGCTTGGAATCTTCTTGGAGGAAAGGGGCAACATTGTTGGCATCATGCCAGATAGGATACCCTTTCAGCTTGCCCAGAACATTTAAAGCTGCTAAAGTAGCCATGCCCTCACGTGTCCACTTAGAAGCAGATGCAATATGTGGCACAACAACTGCATTGGGGAGCTCAGCAAGCCCAGGCTTCATGAAGGGTTCATCCTCAAATACATCAAGCGCAGCTCGAAACTGTGGATTGGCTTGCAAATGTCGAACCAACGCAACTTCATCAATAATTGGTCCTCGACTTGCATTCACTAGTATTGCCTCCTTTTTCATTACAGCAAGGCGATCCTTGTTGATGAGATGAAAAGTGGTTTTATCCAAGATAGGATGCAAGCTCACCACATCAGACAACTTGAGCAAATCTTCAACTGTCTGGACTCGCTTCCATGTAACTGGACTTTCCCCTTGGGACTCGAGGAATTGGCCATAAGCTGTAACAAATTTTTCAAGTCGAGTTGCTTGATAAAGGTCATAGTAGACAACATTCATCTTAAAGCCTTCAACCATCATCCTTGCATAAGCAGAACCAATCCTTCCAGCACCAATAATGCCCACAGTCTGGCCCTTTAATAAATTTCCAATAAACAGATTTGGAAGCCAGCCATCATATTTTCCAGCTCGCATGAAGGAATCAGCTTCAACCAGCCTTCGTGAAGCTGCTAGTGTTAATGAGGCCGCTAGCTCTGCTGTCGTTTCCGTGAGCACACCTGGTGTGTTGCCTACAGCTATGCCATTGCGTGTTGCTGCCTCCACTTCAACATTATTGTACCCAACAGCCATGTTGCTGAATGCATGGCCTCCAGCTCTCTTCAAAGTAGAAAAGAGAGCATCTCCCCATTCCTCAGTAAGCTGGCCGATGACACCATCGCACTGGGTTCCAATCAGACGTTTGATGTCATCAACACTAAGGATAGTCTTCTTTTCTGTGCAGATCTCAACGCGGCAGCCACCATCGACAAGGAGGTTGATCCAGCGGGAGCCAGGCATAGGCTTGGTGCTGACGACGCGCAAGTGGCCATTAGGATTGAGGACTTCAACAGAGAGAGGGGAGGAAGACATAGCCACAAAGCCCTTGGCCTGGCAGGAGCGTGAGGGAGCAGCTACCAGAGAGACGAAGCGGAAGAGCAGGAGGCTCAATCCATCCCGCACTTTAATATCGCACTTGTTGAGATAGGGTAATAAAGGGCTCATTCTCAGCCTCACCTGCATTTTGCCCCCCTCTTTACTTTGCCACCTTACATTATCACCCTCTTATCTCTTTAACTCCAAAGTTCGCG

**HYDROXYPHENYLPYRUVATE REDUCTASE 2 (HPPR2)**:

GAGAGAGAGAGAGAAAGAGAGAGTAAGAGAGAGAGAAAGAGATGCCTATTAGAGGAATGCCAACCAAGGCAGTATTCCTCCGTTGGTTTACTTTACACGGGAGTTAAAAGGGGCTGCCCAGAGAAGTGAGCTTCAAGGTTGGAAACAACTAGATCTGCCATACCCCTCCTGGTTTCCCAAGTGGCACTTCCAACATGAGGAGTCAAAACAACATTATCCAAGGACATGAGTTCAGAAGGAACATGAGGCTCATCCTCAAACACATCAAGCCCGGCACCTCCGAGACGGCCCTCCACCAATGCCTTGACGAGATCAGCCTCATCTACAATAGGACCTCGGGCAATGTTTATGAGATATCCATTTGGACCCAATGCATCCAGGACTTCCCGGTTGATGCTGCCCCTGGTTTCTTCAGTCAGGGGGCAAGCAAGTATTAAAACATCTGATTTGGCAGCAAGCTCTGTCACAGTAGGATAATATGTATAGGGTATGCTTGGCTTTGAAGATCTGCCTTGATATGAAATTGGGCACCCAAAAGCTTCAGCACGCTTTGCTATGGCCATACCAATTCTTCCAAGCCCCACAATTCCTACTCGCTTTCCACTTGCCTTTGTGGTCAGTATGAAGGCATTTTTGACAGGCCAGCAGCCCTGTCGGACAAACTGGTCAGCAGCTGGGATTCGGCGCAGGCACGCAAGCATAAGGGCAAGTGCGGTGTCAGCCGTGCAATCAGTAAGGACGTCGGGGGTGTAGGTGACGGCAATGCCACGTTGGCGACACTTGGGAAGGTCGATTTTGTCAAGGCCGACGCTGTGGGATGACACAATTTCGAGATGTGGGAGTGCATCAATAAGGGGGGCACTTGCGCCCACAACTGTATTGCACACGAGGCCTCGGATGGAGGCTGCGTGTGCGGCAACAAAGGCGGCATGGTCAGGAGCTTCCCAGAGCTTGAAGACAGAGAAGCTGTTCTCCAGCGATGGTACCATTGTGTCGATCATGGGCTGCGTCAGCAGCACCCCAGGCCGAGCCATCCCACACCCACACAAAGAAGCAGCCTCAGCCTCAGGCTTTTTACTGGTTGGTTTTCGGCGCTCCTCCCACTCACACACTCCTTTTAATGTGCATGGGTAGCGAGGC

**GLUTAMATE:GLYOXYLATE AMINOTRANSFERASE 1 (GGAT1)**:

TTTTGAACATAGTGCCCTGGGCCTCACATTAGTTGGGTGTACACTACTTGCCATCCATTGCTAAGAAAAGACAAGATCATGACACGTGGAATTGAAATTGTTTAAACATTCTTTTTGGGCTGCATGTAAGGTACATCTGATTTAGATGTCATGAATGCCAACTGACTACACTAGCTAAAGGTACTACTGTAGTCTGTAAGCTGAGGAGCTGACATTTGAAGACAGTTGTTCTGAAAACTGCAAAGTAGTGAATGCAGCATTGTCTTTATTTCCCAACTTGGATGATAAAATAAAAACCTGGCCTGCTTGGCATTAATTACATAAGATGAATAGTGCCCTGTCTGCTTGGCATCGACTATACAGCATGAATAGTGTTAACCTCAGGGTAACAGCACTCATAATAGAAGTCCAGAAGCTTCAATACATAGTAGCCCAAATCTACCCGATAGTTTGTTGCCGTTTTTCAGCAAATCGGTATGCGATTCTTTGCAGGACACACGGTTTTTCCATAGCCTTCTATATGACAACAAGGTCCATCTTTTGTGCACTACCATAGTGTCAGTATAGTAGAATACTGGGACGGAGTAGCAAATGAAGAAATACTTGTCCTTCTAATTATGAAATAGCCCCCTCATAATTTAGATAGAGTCCTGGATGGCCCCTCATACTGTGCCATAAAATTCTCATTGAAATTTTTGAACAAGTCCATAACTTCTGGCATATCTTCCTCGGCTGGTAGAATCGTAGTCCTTAGATGAAATGTACCGTCTTTTTGGCCAAAGCCCGATCCGGGAACTGTTGAAATTCCAGTAGCCTCAAGCAGCCTTAAGCAGTAAAACACATCTGGTGCTTTACCTGCCTTTTTGGCAGCTGCTATGGCAGCCAGTGGCAGCTTCACTTTTGGAAACGAGTACATTGCACCTTCAGTGAAATTACAAGTAATGTTTCTGCAACTATTAAAGCCATCTGTCATCATATGAGCACGCTTCCGTAAAGACCCCAAAGTTGCCTCGCTCTCTTGGGCATATATAGGATAAGAGGTCTCTCCTGGCTTTGGGGGGTTGACCATCAGTCCCACAAAAATTTGACCTGGTACATTCGGACTGAGGGCAATGGATGCCACTTTGTACAGTTCATCCACTGTCTTTGGGTGAATATTTGTCAGTTCAAAATATCCGCCACGCTGACCGCACTCGCCCCAATAGCCTTTTGAGACCGTATGAAAAGATATAAGCTCTACTTCTTTGCTGAAGGGAGGACCCATATCCATCAAAACCTTCTTTGCACTTATGAATGGTCGTTCATCCTGATATATGTTCAGCTGGTACACTTCATCAGCCAGCATCACAATCCTCTCTTGATGACAGAACTGAATGAGCTCCTGAAGATTAGCTTCTGAAAGACATTGCCCTGTTGGGTTTCCAGGGTTGATGAAGACCAAGGCACGCAGAGTTATTCCCTTCTTCCGGGCTTCAAAAACTTGGTGTCTCAGATCTTGAACATCCAAACCCCAATTAGCTTCCTCATTTAGATAATAAGGGACTAATGTCCCTCCAACCAGTTGAATTGTTGCAGAATATAGAGGGTATTGTGGTATGGGACAAAGCACCCCATCCTTGTCATCCCTAACGATGATATTGAGAAGCTGCATCACGCCTTTACTAGCCCCATCAGTGAGAAATATCAACTCTGGATCGCTTGGATATCCATCTCGTCTCAAAATAAAGTCTGCAACATCTTTCCGTATACCAGGTAGTCCTCGTGAATCACTGTAAGCACCTAGACCCCCCGATGTGTAAGATAAGTAGTGCTTTGCTCTAGAAATGGCATCAGCAGGGAAAAAAATGTCCACGTTTGGATCATCCATAAGAAAAGGGGCTTGACAAAGTGCAATCACCTGACGAGGAAATGTCAGTGGCTTTTGCCCCAGGCCATGGGGGTTGCCTACATTGGTGAAAATAATCTTTTTGCCATCCTTTTGCAATTCAGAAGCTCGGATGTAGAGCTCTCCACGTACTGCATACACCATCTTCTGTATGTTGGCATTGATAGTATCCGGATCAAGCACTTTCTTGGGCGCCATTGAAGCACAACTCTCTCACACGCGCGCACACACACACACACACATACACA

**β-CAROTENOID HYDROXYLASE 1 (BCH1):**

GGAAGTTGCTCAATGAGAGCGCTCCACGTGGGCGAGGAGATTGCGGCGGCTGTAGCTGCTTGTTCGATCGAGCCATCGCTGCAGAGAATTCGTTATGAAGCAGCCCACATTTCCGGCAGACATTCCTTCGTCATTTCCAAGCTTTTCTGCACTTGGGCTCCAATCAGAACATGGGTTTCAAGTCCCCTTTCTTATCCCCTCACATTTCCATCTCTCCTTCCACAGCTCTATTCCGCCATCGCCTCTTCTTCCACAGCAGAAGAGCCCTTCATTTTCACCCCCTCCTCCCTAAGCAGAATGCAGTTATTAGGATGGTTGTTGGAGAGCGCCATTCTACCGGCATCTTTGAATCTGCAGAGTCCTTGGCTGCTTCGGAGCTTCCCCCCCCAGCCGCGTTAGTCGAATCAGTTGTGCCCTCTGTGGAGCTAGGAACTGCGACAAGAATATCTGAAAAAAAGAGCAAAAGGAAAGCTGAGCGCATGGCTTATCTCTTTGCTGCTGTCGCGTCAACTGTCGGCATTACGTTTCTAACTGCGGGCGCTGTCTACTATCGCTTTGCTTGGCAAACCGAGGGCACTGAAATTCCTTTGTCTGAGATGTTTGGAACATTTTCCCTAGCAGTGGGTGCATCTGTCGGTATGGAATTTTGGGCTCGATGGGCACATCGCGCTTTGTGGCACGACTCTTTATGGAGTATGCATGAGTCCCACCATCGACCTAGGGACGGGCCCTTCGAATTAAATGATGTTTTCGCCATTATCAATGCTGTTCCTGCCATCGCCTTAATGGCGTTTGGTTTTTTCCACAAAGGGCTAGTTCCTGGGCTGTGTTTTGGAGCTGGCTTGGGTATCACAGTCTTTGGTATTGCATACATGTTTGTACATGATGGTCTAGTTCATCGGAGGTTTCCTGTCGGTCCAATTGCAGATGTCCCTTACCTCCAAAAAGTTGCTGCTGCACACCAACTTCATCATGCAGACAAGTTCAATGGTGTACCTTATGGACTTTTCCTTGGACCCAAGGTTTCTATGTAAACTGAATTTCAATTACTGATTGAATGCTGTTGAAGTGGTATTGCAAGTTGGAGGAGCTGAGGAGCTTGAAAAGATGATACAATTGAGGAAGAAAAGGGCCAACCATTGAGTCTTTAATGTGGCAGGATTCGAGTTTTTTCGAAAAAACTTGGCAATGGATGATATACATAGGGTGGGCTTTGCACAAACTGTTTGCTCTTTCATTGAGTCCTTCATGACTAAAATTCTTGTTTCCAGTATAAGATTGGTGAGGAGTACCCAATGGGTACACGCATGCTAAAACGATGGAGATCAAGAGCAGAGCGGAAGGGTATTCGCGTGTACGATTAATCAGATCCATGTGCTCACTTGTTTTAAGAGAGAGAGAAGCCTGACATTATGCTCAATAGTAATTTATTCAGATGTAAAGAAACAACCATAAAACAAGTTTTTTCAAATAATTCTAGGTTTGAAGCCAAAGGAATGCATCGAATGCAGAACAGGTCTTGAGTTTTATCTTTATCTTTGCTTTGAGATCGAG

**LUTEIN DEFICIENT 5 (LUT5):**

GATTTGGGTACCCTAGATCAATGGCTGAGAACTTGCCCATGGAGGCATGAATCACGAATGATGTAGCCAGGTGTCATTCATGAACTGAGGGTGTTATTTTTCCACGTTGCTCTCTGAATTTCTGAGTGAGAGGTACTTACGACTGAGGTAATCGTCGAGGTCGCCATGGAAGCTGAGAGGTAGCAGACTCAGATGATAGCTGCTTGAAACTTCCTTCTGTAGCTCATTAAAGCTTCAATGGCGAGCTTGCAAGCTGCACCAGCTTTACAAGTCGTGCCTCCTCTGTGTAGTCTCCACCAAGATGGTTCTTCGACTTCGTGCCTTCGTTTCCTTTCAGACAGGCGAATTAAGCCTCTCTCCATCTCCATCCACACTTCCATCTTTCCGAGCTTCCTCCCCTCCTCCTCCTTTACATCTACTTTGTCATCTTCATTTGTTACAAATGCCTCCTCCAAAGAAAATGGTTCTCAAGACGAGGCCCCTGATGCGGCCATTCAAAATCTTCAAAGGTTACTGGCCTCAAAGAACGAGGGGGAGCTTAGAGAACGCATACTGTCTGGCGAATTTTCAGTCGGCTCTTCCAAAAGCAATATGCGGAAGGCACGCGAGGTTTTAATGACATGGGGGCAGCTCGGGCGCAGCCTTGCATACTTATTGGCAAATTTGGAGCGAGGTTACCTGTTGCGGCAGGCCAGGAAGATGCCTGAGGCAAGGGGGAATGTCAAGGCAATTACTGGACAACCCTTTTTTATACCCTTGTATGACCTGTATTTAACGTATGGGGGCATCTTCAGGCTTACTTTTGGACCCAAGTCGTTTGTTATAATCTCTGACCCTGCAATAGCAAAGCGTCTGTTGAAAGACAATTCAAAGGCTTATTCGAAGGGTATTCTTGCAGAGATTTTGGATTTTGTAATGGGAAAGGGGCTCATACCTGCAGATGAGGAAGTATGGCGTGTTCGACGACGGGTGATAGTTCCAGCTCTGCATAGAAAGTATGTTTCGGAAATGGTTGGCTTATTCGGTAGGGCAGCACAAAACCTATGCCAGAAGCTTGATGTTGCTGCTAGTAATGGGGAGGATGTGGAAATGGAATCATTGTTTTCTCGCCTGACGCTGGATATAATTGGAAAAGCAGTCTTCAATTATGACTTCAACTCATTGTCGCTGGATACAGGAATTGTAGAGGCTGTGTACACTGTTTTGCGAGAGGCGGAAGACCGAAGCATAGCCATCATCCCATATTGGGACATTCCTATAGTCAAATATCTTTCACCAAGACAGAAGAGGGTAGCAGCATCCCTTCAGTTGATCAACAAAACTTTGGATGATCTTATTGGTATTTGCAAGAAAATGGTTGAGGAGGAAAATGTGAACTTCAATGAAGAGTACATGAATGAAAGAGACCCAAGTATTCTGCATTTTCTCTTGGCTTCTGGTGATGATTTTTCAAGCAAGCAGCTCAGAGACGACCTAATGACTTTGTTGATTGCTGGTCACGAAACATCTGCTGCAGTTTTAACATGGGCTTTCTTTCTTCTGGCAAAGAACCCTGCTGTTTTGGCCAAAGTTCAGCGTGAGGTGGATCATGTACTTGGGGACAGAACACCCAACATTGAGGATATGAAGAAGCTTAAATATACCACTCGTATCATCAATGAGACATTGCGGCTGTATCCACAGCCACCTGTTTTGATTAGGCGATCATTGGAAGAAGACATTCTCGGGAAGTACTATATATCTAAAGGAGAGGATATTTTCATTTCTGTATGGAATCTGCACCGCTCTCCAACTTTATGGGAGGATGCCGAGGAGTTTAGACCAGAAAGGTGGCCTGTAGATGGTCCCGATCCAACTGAAGTGACAGAAAACTACAGATATCTACCTTTTGGGGGTGGTCCTCGCAAATGTGTAGGAGATATGTTCGCTACCTTCGAGAACATCACAGCTGTAGCCATGCTGGTTCGGCGTTTCAATTTTGAATTGGCTTTAGGAGCACCACCGGTGGGGATGACCACAGGTGCCACAATCCATACAACTGCCGGCTTAAACATGACTGTAACTCACAGATTAAAGCCTGTAGTCATTCCTGTAACGGGAGGAACAACACTAACTACCATAGATGAGAATGGCTCAATAAAAAGCTATACGCCCAGCAATGGGACTGCAACACTTTATACATCTCAGAAGGAAAATCAGGGAGCTCAAAAAATAGGACCTTTGTAAATATAATCGGTAATGTGGTAATCAATTTTCTTTCCTGCAATGTTGTGCGTTTCTTTGGAAAAGGGAGTTACGAGAGGAGCAAATGCAATGATTGCAGCCTGCTCCCTTGTGGGGAGTTCAAAGATAATGTCTTGAGTTTCACGGATGATGAATGCCTATGAATGTACCTGAGGGGGGTGCTATGAATGAAGTATCATCACAGTGCCTTTGCTAGTCACATACTGCAATGGGGGCATAAACTTGTTTTGAAATGGTTTTATAGCTTTGAACAACTCAATCATCCAACCTCATTGGGCCAGTGTCAAAGTTGGGCTTGGTTGATACTCACAAAGGGTCTTAACGTAAGAAATCTTGAAGCTTCCAGCAGCACAGCTCATGCAAGGAAACTTTGAGGACCAGACTGATTTTTCAGTGAACAGAACCCTCCCTCGGCACCTTGTCATTACTGTTGTATAACGATTACTATCTTCACAAACCTATTTTTTAAACTTCAGTCAAGCCGTGTACTCCACCCCTACATTCAAAATTTACTGTACCTCCTTTGGTGAACTTTGGAGTGAGTGCTTTGGTAAATATTTTGATTCAATATAAAGCAATTCAGGAAAGGCTGGCAATTGCATGTGCTGCCCCATGTGAAAGGCCAGAGTTTTAACATTTTATATTGGCCTTCAAATGCCGAGTGCAGATTGCCTTTTGTGGTTTTGGAATGTCGAAGTCGAGACAGATTGCCTTGAGTATGTTCATCTGCTTCTCTACTTTGTGCAACAAGTTTACAATGTAGTCATTTTTGTGTCACTTTGGTGACTTAATAGAACTGTACCCATCTGAAAGCCCCAGTACCAGCATTCTACTTTGAACTGAGAATCATGCAGCTATGCATTTACTTTTTCCTGATAATTCTGTACAGAACCTCAACCATTTGGTCAATGCTCAATAGGATGGGTTGATGCTAGGCTTCCTGTGTCCGGTGTCGCAATTGCTGTGCTGCTCGGTAATTGAAGGAAACTGTTTTTTGTATCTCAGGATTTTATGTGGGGCGCGCATCAACTCTGCCTTGCAGGCTCTTTTGTTTTGG

**VIOLAXANTHIN DE-EPOXIDASE 1 (VDE1):**

ATGGAATGCTCTCGAAACAAAGCATCCTAACCATGAGCTTGGACGCAGCGAAACTTGCAGAATGGCGCCCCTCTCCAGCTCCTGTGGCGCTTCGTTACACTCTGAAGCACGATGGGCAAGCCTTCCCTTACTTCATGATCATGGTGGGCATTATCGGCATTATCATCTTGGAAGCAGCACAAAAACAAGGACTCTGCATTTTGTGAAACCTCTCAGAGTCATAGCTTCCAATGAGGCAGCATCTTCCGCAAAGGCCATCTCGCGCGCTTGGCCTGTGCTACCCTCACAGCCCATCTCCCTTCTCGCAGGTCTTCTTCTTGGGCTTCAATTGTGGACGGCACCCTCTAGCCTAGCTGTGGATGCCCTCAAAACATGCACTTGTCTTCTCAAAGAATGCAGGGTAGAATTGGCACGGTGCATTGCCGATCCTGCATGTGCTGCAAATGTTGCATGCTTACAAACTTGCAACAATAGGCCTGATGAGACAGAATGCCAGATAGGCTGTGGAGATGTTTTTGAGAACAGTGTGGTTGACCAATTTAATGACTGTGCCGTATCGAAAAAGAAGTGTGTGCCTCAGAAGGCGGATGAAGGAACGTTTCCAATCCCACAACCTTCAGCACTTGTCCAAAAATTCAATACCACTGATTTCAGTGGAAAATGGTATATCAGCAGTGGTCTCAATCCAACCTTTGACACGTTCGATTGCCAATTGCATGAATTTTCAACATCTCCAAATAAGGTGACTGGCAAGCTTACATGGAGAATAAAAACTCCAGATGGGGGCTTTTTCACACGCTCTGCAATGCAGAATTTTGTGCAAGACCCAAAACAACCAGGGATACTTTACAACCATGGCAATGAGTTCCTACATTACCAAGATGACTGGTACATCTTGTCTGCCAAGATGGAGAATCAAGCTGATGATTACGTCTTTGTTTACTACAAGGGGCGAAATGATGCATGGGATGGCTATGGAGGGGCGGTCGTCTACACACGGAGTAAAGTACTTCCTCCAAGCATTGTCCCTGAACTGACAGCAGCAGCCAAGAAGGTTGGGTTAGATTTCAACAAGTTCAGAAGAACCGATAATACGTGTGGGCCCGAGCCTCCCCTTTTTGCAAGACTAGAGAAGAAAGTTGAAGAGGGGGAGCAAGCACTTGTGCAAGAGGTGAGAGAAATAGAGCAGGAGGTTGTGAATGTGGGCAAATCTGAAATGACATTATTTGACAGGCTGAAGAAAGGGATTTTAGAACTGAAGAAGGACCAAGATTATCTTCTTAGAGAGCTGGGAGATGAAGAAAAAGCCCTCTTGAGTGACTTGAAAATGGAGGCAGCAGAGGTGGAGAAGCTTTTTGGAAAGGCACTGCCCATAAGGAAACTGAGGTGAAGCCAGTATATTTACACTTACCAAATTATCATGGTGCAACATCCTTGTTCTTATTGGGTCTTGGTTATCAGCATCAAAGTTTGATACTGTTGGAATCGGATGATCACGAACAAATGGTATCTTTTGTATTATCATGTTCATGAGTAGACGTGTTCTATCATTGACGAGCATATTTCATTGTGTATAATCTTGATGTACATCTCCAAGTCCTCCTCACTTGTTACCAATTGTCGCAGTTGACACTCCAAGTAGGATGACGAGATGGAATTTGCAAGATTGAGCAATTGTTTCTCGCTTTTGAAGTTAAACTCGGGCAAGAGCATATTGTAGAAGAAACCTTCAGTTTGGTGGCTAGGGTGGAAGTTTGTGAATCTTACAACTTGTTGATTTTTCTCGATGTGTTGGTCATTAGCATTAGTCCCAATTGTTCTCTTATTTGTTGGCGATTACTTTTGTATTGTCTATGATACTCACAAAAGGTTAAGTACTCAAGGCATGATGGTCTTGTGATGTATATATCAATAGAGCACATTACTAGGTTATGATGCCTTGAAATCATCCATTGTCTCATGGATGGTGGTGAAGAATCAATAGATTTTACAATTTCAC

**LONG AFTER FAR-RED 3 (LAF3)**:

GGGGCAAAGTATATATCAATCATGAACTTTGAATAAAAAGAGAGAGCAAAGCTAGAGTGTTCTATGATATTTACAATTATGAAACATTTACTTGCTGTCCCGATCATCATTCTTGCTCAAGAATAACTAAACACTGAAAGCTCATAAAATGGAATGCTATGGTTGATTAAACTTTTGAAAACACTTTGATTCCTCCCTTGTAAGTTGCTAAGACACTTATTTGATCTAAGTTCTTAAAAATGTTTCCAGAAAGCACAGCAAAATCTGCAAATTTTCCTCGAGATAAGGACCCAATGTCATCATCCATAAAGCTAGCATATGCTGCAGACGTTGTGTACCCTTCTAAAGCTGCCTTGACATCAATGCATTCTGCTGGTATCCATGAGGAATTCCAGCCTGCAGGTGTCCTTTCTTTAGCTGCCTGCATACCTCCTAAAGGATTAAGATGTGCCACAGGCCAGTCTGAGCCAAATACAAGAGTTGTCTTGGCCAGCAATGACTTGAACAAGTACGATTCCTTGAGAGCTCTCTCTTCTCCAAGTTTTCTTGGAGCAATGTAAGCATCATCTAGCAAGTGCTCAGGCTGCATTGAAGCTACTATGCCTTGAGACCCAAAAATCCCAAGAGCAGTTTGTGATAAATGTTGTGCATGCTCAATCCTTAACCTTCGATCACGATGACCATTTGTAACGTTCAGAGCTTGATATAAATCCAGTATCCTATCATTTGCAGCATCTCCAATAGCATGAATCGCTATTTGCAATCCTCTTTTGTCTGCCTCTAGCATATCTCTCATGAGGGTTTCTGGATTGGTCACTTCCAGCCCAAAGTTGCCTGGATCGTCATTATATGGCTCATTGAATAAAGCAGTCCCAGATCCCAGTGATCCATCTGCAAATGCCTTCACACCTCCAACTCTTAGATTTTGGCTTAGCTTTGGGCCTGTTTGTTTTATTTGCTTATCAACATCGGACCATGTCCCAAGAGGAAAGAAAACATGTACTCGTAAAAGAAGATTACCCATTTGATCTGCCCACTTGTAGACGTCATGGAAATCATCCCAAACTTTTGTTGTTGGCAATCCAGGGAAAAACCTTCCAAAGTCAACAACTGCTGTAACTCCCTTAGATAACGCCTCCTTGCTTGCTCTAGTAAGAGCATCTCTTCGTTCTTGAGCTGTAGCTTCAGGAATACAGTCGACAACAAATTTCATTGCTGAATCAACTAGAATGCCAGTCAGCCCACCATCTTTTCCCCTGATGAAGCTCCCACCCTCAGGATCAGCCATGCCCTCTGTTATCCCCGCAAGTTGCAAGGCCTTTGAGTTTGCAAGTGCCATGTGACCATCCATTCGATAAATCCATACAGGAACTTCTTTTGTAAAAGAATCAATCCATGATGCACTCGGCAACTCTCCACCCCATTTTTCATTGTTCCATCCACCTCCTAGAGCCCACTGTCCACTCTTTACTTCTTGCACACCTAGCTGAACTTGCTTAATAAAGTCACTTTGGCTGCATGTATCTTGAAGTGTGATTTGCAGCATCTGAAGTCCCCCAGAAATCAAATGAACATGTGAGTCTATGAATCCAGGTACTACAAATCTTCCTTTCAGATTGTGATATTTGGTATTTCGGCCAGTCCTTGCCTTAACCGTGTGGAGGGATCCAATGTCAAGGATGCGGCCTTTGGAGAGGGCCATGGCGTTGGCCCATGCCATTCTTGGGTCACCCGTAAAAATGGTAGCATTTGTAAAAACTACATCAGCAACAGCTTCGGGAGCGGGAGCTCGGAGCAAGAGGTGGGTGGCGATGGCGAAGCAAAGGGGCAACACGAGTAGCAGGCAGAACCGTGGTGCATCCATTTGCCCAAAGCTGGCGATATAGCAGTTTATGTGCGCATTTGTGGGATATTGATACTACTTAGAGAGAGAGCAAGCATTTTCAAATAGATTGTGGAGTATGTTTGTGGAGAGAGAAAGAGAGAGTGATGTCCAAGCACTCAGGAGCGCAGCCCTAGACACGCAACTGCTTTGCTCTCGTCCGCTGCCCGAGCTTCCCTGGTTTTGGATCTTCACCTTCACTAGCCAGGTCCAAGCAGCCCACCCAAGTTTGACTTCTCAAGGAGTGTCTAAGTTGTACTCCTTCATTGCTTAACATTTTTGCCTTACTTCTTGCAACTTGTGCGGAGTTGCTATGCCAACTGCACTTTTCCTCTTTCTTTCTCTTCAAAGCAACATGACAAGGAGAAGCCTCTTAAACATGACCACCATTAATTAGTGGATGTTCACTTGTATCTATCCCTAGTTGAGTTGACAATCTATTACTTGAACCATCGAGTAG

**PHYTOCHROME B (PHYB)**:

CCCCCCCCCCCCCCCAAAACCCCCAATTTTCGCTTGCTGATCGCTCGCCCAATTCCTGCAGCATGTCCTCCAAAACCATGACCTACTCGTCTTCTGCTGCAGCCGCTCGCAGCAGCGGCTCTTCTGTCGTCCGCTCCAAACACAATCGTCGCGTTGTGGTTGCTCAGACCACTGCAGATGCAAAATTGCATGCAGTGTTTGAGCAAGCACAGTCGGAGGGTGATACAGACGGCAGCTCCTTCGACTATATGCGCTCCATTGAGGATGCCCGAGACTCTGTTTTGAGTGAGAGAGTTCCTGCCCAGGCGGTAACAGCATATCTCCAGCGAATGCAGAGGGGTGGCCTCATTCAGCCGTTTGGATGCATGCTTGCAGTTGAGGAGGGCAGCTACAGAGTCATTGCTTTTAGTGAGAATGCTGCGGAGATGCTGGACCTCATGCCACAATCAGTACCAAGTATGGGCACTCAGAATTTCTTGCTCGGCATTGGCACGGATGCCCGCACCCTTTTCACGCCTTCAAGTGCTGCTGCTCTGGAAAAAGCCTCGGGTGCCGTTGATGTCTCTATGCTTAATCCCATCACTGTTCACTGCCGCAGCTCTGGTAAGCCCTTCAATGCTATCGTTCACCGTATCGATGTTGGTTTAGTCATTGATTTCGAGCCAGTTCGGCCATCTGACGTAGCCGTTTGGGCTGCTGCCGGGGCCTTGCAATCCCACAAGCTCGCTGCCAAAGCTATTTCTAGGTTGCAGGCACTTCCAGTAGGGGATATCGATCTCCTTTGCGACTCTGTAGTGGAAGAAGTAAGGGAGCTCACAGGGTATGATCGCGTGATGGCATATAAGTTTCACGAGGATGAACACGGCGAAGTTTTGGCTGAGATACGGAGGTCCGACTTGGAGCCCTACTTGGGATTGCACTATCCTGCTACCGATATACCACAGGCATCTCGATTCCTTTTCATGAAGAACAGGGTAAGGATGATATGTGATTGTCGGGCAATCCCTGTAAGGGTTATACAGGACAAGGAGCTAAGGCAGCCTCTGAGCTTAGCGGGTTCTACATTACGAGCGCCCCATGGATGTCACGGTCAGTACATGGCGAATATGGGATCAATTGCTTCACTGGTTATGGCTGTGGTAGTCAATGATAACGATGAGGATCTGAGCAACAGACCTCATCAGCCCAAGATGCGGAGGCTTTGGGGACTAGTAGTTTGTCATCACACCACGCCCAGAGCTGTCCCGTTTGCACTCAGATCAGCTTGTGAGTTTTTGATGCAGGTATTTGGTCTTCAGATAAACATGGAGCTGGAGCTTGCTGCTCAGATGAGGGAGAAGCACATTTTGCGCACACAGACTCTTCTCTGTGACATGCTTCTCAGGGATGCACCGATTGGGATTGTGTCAGAATCACCCAACATCATGGACCTTGTGAAGTGTGATGGAGCTGCTCTGTATTATGGGAAGAAATTCTGGGTGCTTGGAACAACTCCCACGGAGGCTCAGATCAAGGATCTTGCAGAGTGGCTTCTCGATGTCCACAGAGATTCCACTGGTCTGAGCACGGACAGTCTTGCGGATGCTGGATATCCAGGTGCAGCTGCGCTAGGGGATGCTGTTTGTGGAATGGCTGCGGCAAAGATTACCACCAGTGACTTCTTATTTTGGTTTCGGTCTCATACGGCAAAGGAGATCAAGTGGGGGGGGGCAAAGCACGATCCGGAGGACAGGGATGATGGCAGGAAAATGCACCCCAGGTCTTCTTTCAAGGCCTTCTTAGAGGTAGTGAAAAGGAGGAGCTTGCCTTGGGAAGATATGGAGATGGATGCCATCCATTCTTTGCAATTGATCCTTCGAGGATCGTTCCAAGACATAGATGACAGTGATACTAAGACAATGATCCATGCACGCCTAAACGATCTCAAGCTTCATGGCATGGATGAGTTGAGCACTGTTGCAAATGAAATGGTACGTCTTATAGAAACAGCTACGGCACCTATATTTGCAGTTGATGCAGGTGGGTTTATCAATGGGTGGAATGCCAAAGTGGCAGAATTGACAGGTCTTCCAGTGGACGAGGCTATGAGTAGGTCCTTGGTGAGAGACGTGGTGGTGGATGCATCTATGGAAACTGTAGAGAGAGTGTTGGATCTTGCACTGCAAGGGCAAGAAGAACAAAATGTGGAGATTAAGCTCAAGACATATGGTGATCAAGTGGATAAAGGGCCAGTAATTTTGATTGTGAATGCATGTTCGAGCAGGGATTTTACAGACAATGTGGTTGGAGTGTGTTTTGTTGGTCAAGATGTGACGGGTCAAAAGGTAGTAATGGACAAGTTTACTCGCATTCAGGGTGATTATAAGACTATAGTACAGAATCCCAATCCTTTAATACCGCCCATTTTCGGGGCTGATGAGTTTGGGTATTGTTCTGAGTGGAATCCAGCAATGGAGAAGCTGTCAGGTTGGAAACGCGAGGATGTTATTGGTAAAATGTTGATCGGGGAAGTGTTTGGTTCAGAGTTGGCCTGCTGTAAGCTGAGAGGGCAGGATTCTATGACCAAATTCATGATAATTTTGAATGCTGCAATGGGCGGACAAGACTCAGACCGGTTTCCTTTTGGCTTCTTTGACCGCCACGGTAAGTATGCAGAGGCCCTTCTGATTGCCAATAAAAGAACTGACAGTGATGGGACGATCACTGGAGTCTTCTGTTTTCTTCATACTGCTAGTCCAGAGTTGCAACAAGCCCTACAAGTTCAGAAGCGCTCTGCACGGACAGCCTTAGATAGATTAAAAGAAGTCGCTTACATGAAACAAGAAATTAGGAATCCTTTGTATGGAATCATCTTCACACGAAAGCTGCTGGAGGGCACTAGTCTTTCCGACGAGCAAAAGCAAATAATGGAGACGAGTGCTTTATGTGAGAAGCAATTGCAGAACATTCTGGATGAAGACAATTTTGAGAAGCTTGATCAAGGGAATGTTGACCTGGATACATTAGAGTTTACCATGGGCACGGTGATGGATGCTGTAATAAGCCAAGGAATGATTAGATCCAGGGAGAAAGGACTACAGCTCATAAGAGAAACCCATGTCGAGATTAAGAACACATGTCTTTTTGGAGACCAGTATCGTCTACAACAAGTGCTGGCTGACTTTCTTACCACAGCTGTGAGATTCACATCGTCATCAGATGGCTGGGTGGGCATCAAAGTCGTGCCAACCATAAAGAACATAAAAGACGGATTGCATATAGTGCATTTTGAGTTCAGAGTGTCACATCCTGGTTCTGGAATACCGGAAGACCTTGTCCAGCAGATGTATGACCGATCTCAAGAAATTACACAAGAAGGGATGGGACTTAGTGTATCACGGAAGCTTGTTAAATTGATGAATGGAGATGTCAAGTACATTCGGGAACCCGGAGTTTGCTACTTTCTAGTCACTGTGGAGCTTCCTCTTGTTCAAGATGATGACTGATGCCAAGCTTCAGGACGAGGACTTGGGCCTAAGTTCAACATCTCAGCGATTTCCAATGGCAAAACATCGTGTCGTGTGTGCAAGTGACAATTATCTTGTCGCGTATGCACTTTCACCGCTTGGTGCATAGTGCCAAGCATGTTTCAAACTACAGCAGCAGGTGCCCTTGTGTGTATATATACCTGTTGTGTTTCAAACTACGGCTTGCAATCGTGCACTGCGGAAAGAGAGAAAGGCCGCAATGCACATTAGATGTTGCATCATCAAAATATTCAAACTGTTCCTATCAAGGCAATCGAAGCCCAACTATGCACACATACTAGATACCTTTACCAACTACGTGTGTAGATTGAAACATCTTCTCTGTTTTTTTGTCAAATCTACTGCCTGGTTTCCCCAACATACTCTTGGTTGAGAAGAGGGTGGCACATGAAGCCATGTTCCTGCTCATGTGCAAGTAAAAGTGCAGATGAATAAGAGGTGACTTAAACTTTTATCCTGACTATGATTTGGGTATGTGAAGTATTCCTCATCTCTAAAAGAAGAAAGATGGTGTGTCACATGCTTCTACGCAACCTCTCACTCAACCGTGGGCACAGAATTAACACTGTTATGCACAGATTAGATATATGCGGAAGCTGTACTAATATTTGGAAGTATGGTTGCGGAAGCTGTACTAATATTTGGGAGTATGGTTGCATTGATGATGGGATAGAGGCAGCATTGCATGGTCAGGTCAATGAAGTACCAAAAGTCTGTTTGTTGTGTGCTTTTACTCCAGAAGTTGATTTTGTAATAGGCCTTGCACAGTTAATGTTATTCCACCACTAGATGCTAAAATCAATTATTTAATAGTTCCACTGTTAGATGTCTTTTGTTTAGTATGTAGAGGTGGATCTCTGCCACAGGTTGTTCGTCCTCATTCTAAGGGTGCATCGTTGCCGTTCCTTCATTTTCTCCTTGGTTTTTTACAAATAAACATGTATTCTGAAGTGGGTACAAAATGTCAATTCACAACAATTTGCTAGGATAGCACTCGGAAATAACTATTTCCATGTAATCGATGCAAGATTGCAAATCATGTTGTATTTCATTTGATGGTCCTTGGAAAGGAGATGTTAAATTTCCTTCTGTCTAATTAACGAGAGGTACGAGGATGTTTGTGAAAGAGGTGTGCATCAATAGAGGTTAACCTGCTAAGGTATTTGGTGAGAACAAGTTTTTTGCGGCTGCTACCAAAGATGAGGTTGATAATGGGTAAAATGCCATTCTTGCAATGTGCCCTTTCTAATTGGACTCCATCAGGATGTTCGCTTGTTATGACTCAAACCTCCATTGATGGAGGGCTACTCATTTCTATGTATCACAGAGCCCAATGAAGCCTAGAGGACCTTGTAGGAGCAATAGGTGATATGGTGTCTTTGATAACCTAGAGGGTGATGTCTACCAAGATCGAGAGTATTGCTCTCCACAATTTGTTGTATTTGATAGGGTTTCAGCCCTTCGAACA

**PHYTOCHROME A SIGNAL TRANSDUCTION 1 (PAT1):**

AATAAAATGCAGCTGAAACAACCTCAAATTCACATATGACTCACTGACAATCACAAATCGGGCGAAGGAAAAACACGATTGTCCCACAGTGAAGTATAACAGGCACAGATGCTATCTGGCAAATTAAGCCTATATAAGAAGCCACTGCTGTATCAGTCACATTGCCCTTCAAAATGCTTATAAGAACGCGCAGAGTTCAAACATGAAACTTAGCATCTGTAATTGAAAGAGCAAGCTGACTTGTAACTCCCAAACAAATTACTGTGAAACTTAATAGCTAGCCAAAGTAAGAGTGCAAGCTTGGACATACTGCCTCCTCTTCCTGTATACCCTGCAGCAAGACTCGTGTGTGCTGTTATATCATACACATTCTACATACATCGTTCAGCAATTTTGAAGGTGGAATAATTGCGAGCACTTATCATGCCACTTGAGCTAAACAACGGAGATGGCCCATGAGTACCAGCAACTAAAAACCAACTTGCTGAGCAGCAGATCTGCTCTGCCTTGGCCTGAACAGAACCAAGGATCCTGCCCGCACATATAGAAAAAATAGCATACATGCACTTAACCCTTACAAAATTCTAAATAGCCATGGACTGGCTAGCCAAACTAAGTCCTTAAAAAAAAATTAAGAAAAGATTCTGTACCTGGACCTCTTAGCGGTCCTCTTCAATCCAGAATACAATGTTCTCTGGACCAAAACTTCGTGCCTAGGTCATTTCTCCAGCCAGCTGAAAGTATAAGCTTACACTAGCTACAAGCTAGAATATACACTTTCATTCATGATCTAATGCCATGCGGAAGCCGCCATCAAATTTCTGTTCTGCCATCCCAGTACAAGAATCTCGCCGTCTTCCTTCAGCCTGTATTTCTCATTGTAAGACTCGAGGAGGGCCTTGATGGTGTTGTTTACATACAAGCTTAGAGGGTGAGGCCTAAAGCCAGCCATTAACATCCGCGCTCTCCACTTACCTGCAAGCTCATGGCGTTCAACTCGTTGTACACCTTCGCATGCTATGACGTTGACAATGTCGCGAGCCAAGCAATGTTCCTCAACATTGACCCTCTCTTTGCTCTCTCTTGGGAGCGCTACATCAATAGACTCGAAAACAGCTGAGTAGTAATCCAAAGTTTCCATGAACCTTGCCAGAAAAGGGGTTGTGTTGGTGTTGGCATCTTGTTCTACCAAGGTTAGAACTTTTGGGTTAAGACTCTTAGCCATTCTAAGAAGACGATCACGATGATTATCAAGGCAAACATTCTCATCCGGCATATGATGAAGCTGAAGAGCAAAGTTCATTGCCAGAGCCTCTCCAGGCCTTCTTTCTATCATATGCGCCTGCAAACCAGAGAGCACTGTTGAGATAGTGCTAAACTGAAATGGAACTCCGAGGCTACTTGCAAACTTTTCCAACCTCTTCTTCACAGCAAGCATACCACCCAATGGATAAGTTTGAGCTGTAGGATCATCCACAGTACTAATCCGAATATGAGGAGTACCTCCTGGCCTACTCCCTAGTGCTTGTATGAGAGAGAACCACTGTGTTCCTTGATCAATCTGGAAATCAAAAATATGAACCATGGGCTCATCTTTAAAGGCTTCCGCTATTGCACCATTTGCCGCCATATAGGCGAACTTAATGTATGGGCAGATTTCGTAAAGCTTCTGCATGGCGGACAAAACTTCAATTGCAGGAGCTTCCCAACACTTCAAAGCCTTGTAAATGGTGGAGCCGGAGGCATGAAGCCTTGCTACAAGCCCCTCAGACATGTAGGCAGCAAGTCTATCTAAGGGATCTCCATACACAGATGCAGTTTTCCGAAGATCTTGCATTGCCAGCTCAGCTACATCCATCTTTTCTTCGGATATTGCAGCAGCACATGTGACGAGAAGGTTCTTGACACGGCTGCCGAGATTCAATGCTCCCAATGACTGCTCAAAAGTAGACAAGCCATCCAATGGAGATGTCACTGCACATTTCTCCTGGTCTGTTTCTTCTGCGATTCCTAGATTTTCCTGAAGAAGATCATCAAGAAGCTCTACCCAAGAGCCTCTATTCTCATCTAAAAGCTCGTTCATATCTGCATCGTCTGCATCTGGCTCCATAAGTGCATTTTCCAGCTCATGCAACTTGTACACCAAGGCTTTACTATCACACTCAGGAGTATCCTTGGCAGATGCTAAAGCTGAATCTACTACATTTAAGTGCTTATCGGAAAATGTAAGAGTAGAATCAGGACTCTCAACAGAAGACGATGCAATTCTTTGAGATACTTGATTGTAATGTGAGCAGCTAGAAATCTCATTTTTGTACATATCGCTATCATATTGATAGTCAGAGCTGGAATGCGACGGAAGCTCTGAGCTAACTGAAGATTGCGGAGGAGATAGAGACGAAATGGGTGAATTAACATGTCTAAATTGTCCCTGCCCATATTCTGCTTGTTGCTTACTTCCTTCCCTCATGTTGTTATTCATGTAAGCCTGTGCCACTTGTCGGCCATTCCGACCCTGTTGCAAACTGGGGGTAGAGGAATCTGAAGTTCCTCTTGAAACCTCTGAAATGCGGCTATGCTTGTACGTGTAGCCCATACTCATGTTGGCTATGACTGATGGTCTTGAGAACAATCTCCTCAGTAAGTGAATGAGCGCAATCCGAGCACTGCACAGCTGTGTTGCAGCTGCTGTAATCTGAGTGGTTGAAAACCCTGTTGCACCTACTAAAAGTAGGTGCAAAGATGCTGTATAATCCTCTTACTAAACCCTTACTGAAGAGCTTAGCACCAAAATAGGCAAATATGTATACTAGAAACCCTGTCGTGCCTAAATTGTGAAGCTGCTTTTCCAGATAAACCGTCTTACTACACCCTGCTGCACCTGCTAAGGTAGTTAAGATGCTGTTATTGAAAGGCCTGTCTCAGCTGCGGGTATACTTCATGCAAGAGATTGTACAAGAAAAGGTATAAGCTGTACCATAAAGATATCTGTGCATGTGTGCTGGCTGTGCAAGCTTCACCTCAAATATCCATGCTTCCGGCTGCTGGGCCGTGGTTCCCCATTGTCAGTATCAAGTGCCCACTAAGATGCAAAATCCACCATTAAAGCTCTCCGTCGATTCTTCAAGTGCTCGATAAGACGAAAAATCCACCATTGAAGCTCTCTTTCACATGCTCTGGCAAGGCCGCAATGGTCACATCAAGGGAAAGCGTGCAAGTAGTGGAGTGTCACTGTTGTCTGCAAAATAGAGGGCAACCCACAGAGGAGGACTCCGAGAGACAATCCAAATGGGTAAGTGCGTTAGGAAAACGAAGCAACGCACACACACACACACACACA

**VASCULAR PLANT ONE ZINC FINGER PROTEIN (VOZ1):**

TTCTTAAAGGTTTCGAACTTTTCCCTTTGCGTGAGCTTGCCTGTGCGCGTCATCTTCGTGGGTTTTTCTTTCCCGGCGATTTTGGGCGATGGGGAGCCAAGGCAGGCACACCAGGCCCTCCACGCAGACCCAGTCCTTCAAGGCCAAGGCTAAGAATCGAGTTGACGATCTGCAGGGCTTGTTTTGCAGCTTGCAACAGGCCCGGAGAGAAAACAGAGGCGTTGATGCTGCCATGCTTGAAGAGCAGGTCAACCAGATGCTTAGAGAATGGAAGGCGGAGCTCCATGAAGCTTCCCCTGCTACCTCCTTAGCTGTAGCAAGTCCAACCTCATCTGAACTATCTTCTGACATGCAAAGGCTTCTTCAATTGACCGAGGAAGATGATGATGCCACCAGTAGCTTGGCAGCTTGTCTGGCTAAGCCTATGTATGAGCAACGGGGGCGCAGCCCACAGGCATTCAGCAGCAATGAAGTGGAATACAATGCCAGTTATGTGGGAGTTCAAGAGCGCAGTTTTGAGCAGGAAATGTTGGAAAGACTGCTGAGAGCAGAAACACAAGCAGAGTGTTCACCACTCTGTCATGACGATAATTTATGTTTACAAGAACAGCTGACAAGCTCTTCTTTGTGTTTGCAAGATTCTCTCTCACACAAGTTCTATTCAGATGGTGCTGGTTGTCCTCCAGATGAACCATTTTCATCAGTAATGTCTCCCCCTCCTGCCGCTTTCCTTGGCCCAAAATGTGCTTTATGGGATTGCCCCCGACCTGCAACAAGATCAGAATGGTTACTGGACTACTGTAGCACTTTTCATGCAACCTTGGCTCTCAACGAGGGTGCAGGTGGACAATCTCCTGTTGTTCGACCAGGTGGAATTGACTTGAAAGATGGACCCCTGTTCTCATCATTATTTGCAAAGAGTCAAAGCAAAGCCGTTGGTATTCCTGAGTGCGAAGGTGCAGCTACTTCAAAGTCGCCATGGAATGCTTATGAACTATTTGATACCTCCATGCTTCAGGCTGAGGTGATAAGGGAATGGCTTTTCTTTGATAAGCCTCGCAGGGCCTTTGAAAGCGGAACTAGAAAGCAACGTTCTCTGCCAGATCATAATGGCCGTGGCTGGCATGAGTCAAGAAAACAGGTCATGAAAGATTTGGGTGGGCTTAAGAGGTCGTATTACATGGATCCTCAACCGACCCCACAGTTTGAATGGCATCTGTATGAATACGAGCTCACTGACTATGATGCGTGTGCCTTGTATAGGCTGGAGTTAAAGCAAGTGAACACAAACAAGAAAGGCAGTAGGTCAAAGGCACAGAATGATCCAGTGACTGCTTTACAAAACCAAATTGGCAGACTGCCAGCTAATCTATCGGCGCAAGATGGTAAGGTCTGGACGCCTGAAATGATAACAGAGCCAAAGAATAATAAAAATAGACCAAAATCTACCCTTAATGCTGCTCGCTCAAGCGGGCAGTCGGCTGATAATGAGGTTTGCACAAGCTCTTTTGATTATGCAGTGGAAGATATGAGCAACTATCTGCTTAGTCCATAAGGTTACACAAATGTCCAGACGCTGCTCGGTACTAAAATTGCTCATTTAAGCTCTTTTGGCAACAATTGTTTTCCTTGATTGGCAATTGCGTCCTAGGATGGTGTTGTAATCCTGGTCTAAGATAAGTTGCATGATAATGTATATGTAAATATCAAGGGATGCGGGTGGGGTTGTGAGCCTGAAGTCTCTCCCCTATTTTACAAGTTTCAGAGATATATCTGTCCTTCACTAGTCGTGCTTTTTGGCTAGTCTGCTGGAATACAATTCTGTGGTTTGTATATTAAAAAAA

**TYPE ONE SERINE/THREONINE PROTEIN PHOSPHATASE 4 (TOPP4):**

CACAAAGGCTACAATACAAACAGTCGAAGCTGGAGACTTCATAAAGTGATTGCAACGAGCCTCTCCTCTATACAGTAACCAGGCAAGGCAAAACAAAATATTTCACTTGAGAAGATAAGACTAATCAATGTGTTTCTTGGTGAGGACAAAGCCCACCGTGCTTCATACATCGGACAAAAAGCTTGTTAGCTGGTGTTATGAAAACTTTTCATTGGCCCTACTAAAGAATGAAGGGGAAGGAACAGACTACAGTCATGATCGTTTCAATGACAGCTGCTCAATATTACTCCATACCTCGTGGAAAACAGGCCAATAGCACAATTGCTGCCTCATCTCATCTTTATTGAGGAGAATAAGAGTAAGAATAGTCCTGCAATACTTCAAGATAGAAAATGGTTCCAGATGTACTCACTCAATAGGGATGCCGTGTGCACACGTAATCCAAGGAAACGGATACTTGTAGGCATTTATTACTCGGATCAATTGAGTGCAAATGCACTCAGGGCGGTACTAAAAATGACCAAATGCATGTGTCTTGAAAATGCACAATCTACAAACACTCTCAAAATCCACAGAATGAAGAAGATACATATTTCACAGACTAGCACAATATCAGGAAAACAAATGGATTTTTCGTTATACAGCAAAATAAACGCAAATAATACCACCTCTAGATCCATTTGAGACTTTAAGACGCAATAACACTCTGATCTCCCCAGTGCTCACATAAAAAAACAGATGAAGTGTACATGATTGCTCTTATGAAGCTTCCATTGCTTAAAAGCAACCCCATGTGCCACCAGCATCAAGTCCAGGTCAAATAGCTTTCTTTGGCAGAATGTAAACTTTCCAAACTAGCTCATGTGCCAATCTGCAGCGCATCCACAGTTTCTCTTGATTTTCACGGAGAACTAAGCTTTGCCACCCCTTGCTCCACGTGGTGGTGTTCCTGGTTTTGCTGCGTTGACAGATGTATAAGAAAATTTTGGCTTCTTTTCAGCAGGTTTGAGGATCTGAAAAGAGCACATAAGTGTCTCATCAACACTCATCATTGCTCCAGCATTGTCAAATTCCCCACAATAATTAGGCGCTGAAAAGATGGTAACGAGCTGTCTGTCCGCAAAAAACTCATAGCCATCCTCTACCACCTGATGAGCTCGGCATACAAGGTCAAGATCATGCTTTTGTAAAAATTCGTTTACTTTGTCTGCACCAAATGTAAAAGAGACACCTCTATCATTTTCACCCCATCCTGATATCTCTCGCTCAGGGTCGGCCCATAGGAGATCACAAAGCAGCCCTAGATCAGGCACATCAGTTGGCCGAGGTAATGCTCGGATTTGATCCAAATTGTTCAATTCTGGTGAAAGTCCACCATGCATGCAAAGAATCTTCTCATCAATCAGAGCAGCCACAGGAAGACAATTGAAACAGTCAGTAAAGACCTTCCACAACCTGACATTGAACCTCCGCTTGCACTCATCATAAAACCCATATATGCGGTTAATGGAAGCACACTCGTGATTGCCTCTCAGCAGAAAAAAGTTTTCTGGGTACTTGATTTTATAGGCCAAAAGAAGGCAAATTGTCTCCAAGCTTTGCTTGCCCCGGTCCACATAATCTCCTAAGAAAAGATAATTTGCCTGCGGAGGAAGTCCACCATATTCAAAAAGCCTTAACAGATCTGAATACTGGCCATGAATATCACCGCAGATCTTGATTGGCGCTTCAAGTTCCAGCAGGTTAGGCTGTTGCATAAAGATGGCCTTGGATGTTACGCAGAGCTGTCGGATCTCAGCCTCTGAGAGCTGCACCTGCTTGCCAGGCCGCGAGTCCCGTACTTCCAAGAGACGGACAATGATTTCATCCAAAGCAGCTGGGTCCATCAAACCCTAGAAACAGAGCAAGGACAGAAAAAGCCCACAGAGAAAGACGACTGAAGCACCGTCCTGCACGCAGAAATTTGCGCCAAGAACTCACCCACAGGGGCAAATCCAATGTGATCAAAATGAAGCAGAAAGCCCACCTTCTCTGTGAAGAAATAAGACCTAGGAAGACAGAGCAAACCCAAGAAGGATAGATGAAGAAAAATGGACGCTTGATCTAATCACACGCAGAAGCCCCTGCTTCTGCGTAGAGAGGGCGCGGGACGGGGCAAACCACCCATCTGTTGTTGGGTAATGTCGATTTCAACGGCCGCCTCCCGACGGCACTCTCTTCATACGTCACCATCCATCCCATAATACATTTATGCCACCTATTCACATTCGCCTCACCTTTGAA

**HIGH LEAF TEMPERATURE 1 (HT1)**:

ATTTTATGTGAGGATGAGGAAGGTCGTTGGCTATGGCGCTCGATTTTCTCTTTCTCTCTCTCAAGAACCGGTCCCCGCGCAGCAAACAAAGCTCCGCCATGGCCGCTTCCTCTGCTACTTCCACTTGTCGCCCGTGACCCTTTTTTCCTCACCGCCCCCAGTCCATTTCTTCATCTCTTTTCTCAGCTTCTGAGACGACAAGCGTGCTTGAATGACAAAAGTCTCCGTGTTCGGTCTCCATTTGTAAAGGATGAAAGTCTTCATTCATTTTTTAAGAGATTTGCTGCTGCAATCTCTTCATTTTGCACCTCTTGGGCTACCCTGTACAGTACAGGCGGAACTGAAAGACCAAACTTGCTGCTTTAGAAGCTGGAATTCTAATTGATTACTATCAGGAAGCAGAGAGAGCAAAGGCCAAGCAGATGCCTCCTTCAAGGGAGAGCAGTGGGGACAGCACTCTTGAATCGACATGGGTGGGCCGTACGCATTTCTCAAGGTCGATTTGCTACCGCCTCAGCAGCAACGCTCTTTTAGGCCTGTCGTTACCTCGTTTTGATGATTCCAAACTCAAGGGCTTTGTCAAGCTGGTAGGAGAAGACCCAGACACGTTGGCATCGTCGGGCTTGCTCTCAAACGGAATTCTTGCGAGCTCAAATGGCTCAAAGCTAAGTGATGAAGGTATGGTGGCTTCGGTTTCTGTACAGAATTTACAGCCTCTTTACACATCAAGCTCTGTGGATATCTCACCTGAGAAGATTGTTGGAGGGCAGTTATGTGTGCATCATCGTTCTCTCTCTCTTCCATCACCGGGTAGCCGACGTCATCATGAAGGTGAAGAGAATTTGTACTCGAGTGGAGAGTATTCAGATGATGGGACTAAGTTTGATGATGGTAGCCTATCAGATGTGGCGAGGGCTGCCTCTTGGACTGAATATTTTGCTACCCCCGGTAATGAGTTTGCTGTTGAGAAAGCGGAGGACTGGATGGTAGACTTATCTGATCTCTTCTTGGGCCATAAGTTCGCATCTGGGGCTTACAGTCGCATTTACCTTGGCAAATACAAGGACTCTGACGTGGCTGTCAAAATGTTGAGAAAACCTGATAATGATCAAAAAATTTCGAAGCGCCTGGAGCGCCAGTTTAGTTGTGAAGTTAATGTCCTCTCTCGAGTCCATCATCGCAATGTCGTAAAGTTTGTTGGAGCATGCAAGAAGCCGCCAGTGTGTTGCATCATCATAGAGTATCTTGCACTTGGTTCAGTGAGAAGTTATCTGCACAATCAACAACCATTTGCGCTATCATTGAAGGTAGCAGTTGACATGGCACTTGATGTCGCACGAGGCATGGAGTATTTGCATTCCCAAGGTTTGATACATCGTGATCTCAAGTCGGAGAATCTGATTATAGCTGATGATTTGTGTGTCAAAATTACCGACTTTGGCGTTGCATGCTTTGAGTGGGAATCCTCTGTCATGACTGATGATGTGGGTACATATCGATGGATGGCTCCTGAGATGATCACCAAAAAGCACTTTTCGAAAAAATCAGATGTGTACAGTTTTGGCATTGTTTTGTGGGAGCTTCTTACTGGCAATGTTCCTTTCTCGGATTTATCACCTGTTCAGGTTGCTTTTGCTGTAGTACACAAGCATGCACGACTTCCAATTCCACTCAATTGCCCCAATCTCCTTGGCCATTTAATGCATCAATGCTGGCATTCGGATCCGGACAAGAGGCCAGACTTTACCAAAGTGGTCGGAATGCTAGAAAACATGCAGGAGTCAATTGTCTTGAATCGCGACATGAGTGTTTTTCACGGGCGCCATAAGCATACACATGGCTTGTTGCATTGCTTTGGTCGTTGTGCTTCAAATTATGTTTTATGACGGGCCAGCAGCGTGTGCTTGTCACGTTTGCATTCGCCATGTCTGAATCAAGAATTTCTCGTAGGACGATTGAAGGAAAGTTTTCAAAGAGATGAGTAGGAAATTTCTTGTAAAAGGCTCATGTAAATAACAATGAGCTTCTGAGACAGGCGATTTGGTTGGGACGAATATGCAACATGAAGTAATTGGTCTATTGTATGTAGGCCTTTTGTATGGAAATTGTTTAGGTCCTAGTGTAGCAATGAATTTCTTGGCAAGGAGCGAGCTTGCCCAATTAATTCAGTCAAGGAATCATTATTTTCGCGTACAGCAAGCTTGTTTATTCACAGATTGATGCGACAAATCGGAAAGTTTAAAGGTTGCTTTTGCAGCAAGAGCAGTGCTTTCTGCTTGTATTTCATGCTGAAGCTGAGTACTCTCTACATCTAAGTTTGTAGCGATGACATGCATGGCTATAGGTCGATGCAGTTCTACCCTCATTGGCTGGCCTTGCTTGAATAAGCTTGCGGTATTAATATTGAGAATGGGAGATGTTCTCTTCAAGTTACTAAACATGTATTCAAAAAATATGAAGGCTTCTTTATTTTGTACATGCAGCATACACCAAAAGATGAGAGAACATTTGTGAATCAATTTAGTTTTGAAAATTTAAAAAAGTGCTATCGAAGTCTAGTGCCCAAATTTTTTTTGGGTTTACATGTAAAATTATTGGCCAAAATTCT

**KINESIN-LIKE PROTEIN KIN-14B (KAC2)**:

CACCAATAATAATAACAACCCATTGAAAAGTTCATGTATAGCCATGATCCCAACAAGTTCTTCACATGAACACAAATCCACATCAATTTGTAAGCCCAGCATTACTATAGAGCTTTTCATTGTCTTTTCATTTACAAATAAACAAGAAGTATAATTGGGCTACAACAGATGTATAGGTTCTCCTTACAAAGAGGTCTGCCCATAGAAAATTCAATATTGATACACCAAATCAGCTACTCGTCAACCTGATGCCCGGTTGAGATTACGGAATGAGGGGAAAGAACAACACATTCTCCACCTCTGAATGATAAGGAGCTAAGATACAAAAGAAAGAAAAACCACAAAACTATAACACCATCACATACACCACAACAGATTTAGCCTGTATCTAAGTGAAAACTTTTACTCTAACACCACTTCCCGCTACGCGTCTGCTATCCAGCTGCAACCAGTCTGTTATACTGGCCTCTTGAAAGATAAAGAGGCAGTTGATAATTCTCGTGAGTGCTCTCTATCAAGCTCAGGAACAGAGCTTTTTAGAGCAGGTACGCGGATAGCCAACGTCTTGTAAAGCCGAGCATACCGTGCTCGCGTACCATCAGCAGTCCTTGCCAATGCTAACTGACTAAATGCTTCAGGTAGCCGAGTAAGCACTTGCTTGATCTCCTCCAAGGATAATCCAGCCAGTGTTGAAGGCGCAGGCACAATTGCCACCATCTCCTTCCCAGTCTTATCCTGGACTTTTGCCTGTGCTTTTACAACAAGCTCAAGAGAAGATGTGGCACCACATTTAATCACAAATGGGTTTGCGGACCCATTGTTGAACTGCAAGACACTCCATTGCATCACATCACCATCTGCTCCGATGATGTTTTGGTCCCCCCAGTCACCTGACATAGCTGACAATTCTCCTGCACCATCATCCCCAGAAGCTCTTGCCCGTGTGTGGTGAGGACCATCGCGTTGCTCAGCTATCGACTCTACAGCTCTTCTTGCCTCCAGTATGTGCTTTTGAGCACAAGGATGATCTATTGTTAAAAGAGATGGCATTCGATCTGATAATGAATGAAGGGCAGAAAGCATCATTTGCTTTCTGGCAGCGAAAATTGCCTTTACAGGAGCATCTCTCTTGATCTCTTCAGCTTGTTTAACAATGTCCTCAAGTGACATAAGGGATGCTATTCTTGCATCTTCACTTGCAGTGGATGGGTTCCTGATGGGAGACCCACCCTCTTCTTTTGTTAATAGAGCAACGTCAGACCTCATTTGCTGTAAAACCATTCTTCGTTTGTGTTCTACAGACTCCAAGGCAGAACGCAATTTCGCAATTTGGCCCAAGTCATCTGCTTCTTCTGTTGCAAGTGTGGCGGCCACATCCTTCAAGTGTTGCAACTGGGACGTATAAACCCGCTTGGTGTAATCTGATAAGAGTTGTCCCAAAGCATCAGTACTAGGGCGCTGCGGCACGCCAAGACCAGACATCCAGCCATCCATAATTGCCGTTGATAACAGCTCTAGCTGCCCAGCAGTGCCACTAACTGCATCACCTCCAGATAGAGAAAGGAAATCAAAGTTTTCAGCAAGCCAACTGCGAATTTGCCGTTGCAATTCGCCTGCAGGTGTGTGCACAAACAGTGCAGCAAGAGCCTTGTTGCCAATGGCAAAATTGCGAGCTTCTTCAGTCGTTTCCCTCAGCTTTCCACCAGTAACATGCTGCCTCCAAGTCTCCTGGTCAATCCCACGCAACTTCAGAACAATGGATGAAAACTTTGACTTTTTCTCATGCTTGATGTGGACCTTGAATCCTTGTATGTGTTCATCAATTAAATCTTTTCGCACATTGGACTCATACTGAATTGGTGACCTTGCTGGACTACTCCCTCTTGAACCTCGGCTTCTGCTAGATCCTGAACCTACCTTTTCCAGAAAACGTTCCACTGGTGGCACTTTAATAGACTGTAATTCCGGAGCTCGAGAGAGCAAAGACCTTATGTAAAGAATCCTGACCCTAGAGACCAGCATAGTATCCATCACTCTTCGGGGTTCCATCCTTCGGACAAATGCAAAAACAGCATTACGAATTTCTGCCAGCATCTCATGTTCTCTAGCAGCACCGGCTTTGATAACTGCAGCCAGGACCAGCATTAGGAGCTTGTTTGCTCCATCTGAAGCTGCTGATCCCTCTGACTGCTGGTCCGGATTGAAGTCGTTCAAAGCATTGGTCAAGTACTCGCCTGCTGGAGTTGTTTTGACTTTTTCAGAGCTTGATTTTACAAGCGCCAAGTTTCCAGCTATGGCAAGAGCACCAGATGAGGAATCCAGACTCATGGCAGTTGATGACCTTACATCTGACGTAGCAGGTGGTTTTGTGCCTGTATTCCTGTTGGGAGCATCTAATTGAAAATCCTCAACACCCCTTGGCTCCATCTTCGGGACCGCTGGTGAGTTGGTAGAGGCACTTCTATCAGTGAGCCGTTCAAACAACTTTTCATTCTCCTCATGCAAGCGCTCAATAAGCTCATCACGTTTGGACAATTCTTCTTCCAGCCTTTTGACCACCTCACTCTCCACTCGATCCTTCTGCTGCGAGTCTGTATCATCCGGCTTGCGTACTTGCCTAAGATTGGCTCTACCAAAGATATCTGCAGCAGGTTGAGAGCGAGCTTCTTGAAGCTGCTGGTTCAACTCCTGCACTTTGCCCTCAAGAAGCTGAACCTTCTTAATGTACTCTTGCTGCTGAGTTTTTTGATCTTGTTCAGACTTCCCAAGCTGGGCAATCTGAGCCTTCAGTTGCACATTTTGCTCCCTTTCAATCTTCTCCTTCTCAGCAAGCACAGCAGTCTGACTCTTATGATCAGCCTGAAGTGTAAAGGCCACCTTCCAGGCTTTTTGAACTTCATTGAATAAGAGAAGGCACTGATCGTCTGCTTCTTTCAGAGACTTCTTCAATTCTATTGTTTGCTGCTGTGTTTCGTTCAGTTCCTTTTCTTTCTCATATAACTCCTTGCGTGCTTCATTGGCCATGTCGCGCCACTTCTTTATGGTGTCACGATTGCCAAGACTTAGTTCCACATTGCGCGCCCTGGCAGCGAAATGCAGTGATGCAAGTGTTTCCTGGATATCAGTCTGGCTGGGACTAAGGTTGACAATCAGTAGAGTTTTCGCATCACCTCCTAGCGAGTCTGCAAGGGTTTGAGTAAGCTTTGAGTTTGCATAAGGTACATAATCCTTTTTAGCTGTTAAAGCTGAGATCACATCACCGAGCGCTGAGAAGGACTTATTTAAATGAAGTAACTCTGTGAGTCTATCACCAGTGGCCTCTTCTTTTGAAAGCCTCTCACTGGCAGCCATGTCTACCATTGACAGCTTGCTATATTGATTCTCACCAGAAAAACTGTTGGTGTAATGAATGTGGATTGTTAAGACCAAGTGCGCACGGTTAACAGCATCTTTCCCTCGATTCTGTGTACCAGTCTTGTATATTCTAGCAAAATCTGTTGGATTCTCTACTCTCTCAAGAGCAAGCTCTATAAATTTTCCATTATAACCCATCTGCGCAGTTCCTGATGAGCGAATGGAGTTGTATAGGAGATCTCGAACCTCGTTATTAAGTTCGAACATAGTCACTGAAAAGGAGAATTTTGAAGCGGATGTCATGTCGTTATTAGACAAGTCAAAAAGTTCTTCAAAAGCCCTAAGAAATACACCTCGATCGCTATTTGATCCTTCCATGGTATGAGTTTTCCCTGATCCACTTTGTCCATAAGCGAAGATGCATGCATTATAGCCATCCAAGGCAGACTGCACAAATGGCTGAACATCTTGGAAAAAATCTCCTTGACCCACATGCGGACCATACACTCGATCAAATTCAAATTCTTTTTTCTGTGTGGTCCCTGCATCAATCCCCAACGATGATGTGTTTATTCGTATCAAAAAATCATCTGGAAAATCGGTTATTGCTGGGCCCTCATGCTCAAATTGTGGACGCACTCGACAATATACTCGGATGTTGCCTTTCACGCCTAGAAGATCATTGAAAAGCTTTTTCTTCTCTTTCCGCAGCAAAGATAAGTGAGCTTCACCTTCTAGAACAACCTGATCTAGTTTATGAGCTTTCTCAGCCAAAACACCGAGGTAACGTGTAACCCTGACAATTTTTGCATTTGAATACTCTTGCAAATCACTTGCTTCTTGTCGTAGCTCCAGCATTTCCTCTTGTGACACCAACACCTGCTCATGGAGCTTTAAAAGCTTGCTGGCGAGGGCAGAAGGGACAGCTCGCTGGGTCATGTAACTGGGTTGTGGAGCGGCTCCAAGGATGGGTAAAGAGAGACGTCTAGACGCAGCGTTGGGACTGAGAGATGATGCTGCAACTCTATCCAAAAGCACAGCAGCATCCTCATCAACATCATCATCCTCGTCATCGCTGACTGGGTGTGCCCTGCTTGTGTCAACCGACCTGCGAGGATCGTAGCTTGGCAGTTCCCAGCCCCATGAGCTTCGAAGTGACGCATCTCCCATCTTTATACACTTGAAAGGCAATCAGATGAGGCTAACAAAACCTGCCCTGTAGGCTAAAATGAGGATAAACTCTCACTGCAATCACTTGCAGAGACCAAATGTGTGTATGCAATACCAAAACCATATCCTGCACATGTACAAGTTACCTCAAAACCATGGCTGGCAGTGTAAAAAACCTGAAGAAGAAGCCTGTAGGAATCAAACACAAAGCTTCACCGGCGTTTGGCAGTAGTGTTCGAGCTTAAAAGCTTTATATTGCAGTCCAAAAGCCCTAAAATGTAGAGGATGCAGCGAGTGTGCAAAACCCAATTGCAGAATGCAAGCTGAGAAACCCTCGCCTTTTTGTATAAGCTGCTAAGGAGTTGAATGCAGGGGATGCTGTAGTTGCACCCAACAGCTGGGAAAAGAAATACAAGGTAATCTGATGCAATAAGGGGTGTGTATGTGGGTGCGTGCGTGCGTCTGTGTGTGTGTGTGCGTGAGTGAGTTTGTTTGTGTGTGTGTATGTGAAGAATCTGCTCTCCCTGCGTCGGCAATGGTCAAGAGAAAAAAA

**PHOTOTROPIN-2 (PHOT2)**:

TTTTTTTAAAGCAGAACCTTTCCTTTGCATAATTGAAAAAAAAAGAAAAGACAATTTGGGTGTATCCTATCCACAGGTGGGAATAAATGAAAAAAAAAATTTAAGCTTTTAGAGTAACCCATGTTTAGTTGTGAAGGTGCATAAAGGTGGCGTGGAATGGGAACACCTTTTTCCGGCTGCGTGGATCCGCCACACCGAAGTCCCAAGTGCACCACGTCTTGCCCCGCGTGTCGTAGGCAAGTGCCGGTCGTGACATGGCCTTCTCTGTACACAAGCATATGAAGGCAGCTACAACGCCCACTTTAAGCCTGGCCACAAGCGGCGGGGTGCGGGAAGCTGTGAGACGCATGTAATCCTTGGCAGGACAGGCGGAAACATGCTCCCACCGGCCCTGCGCCTCCTCCAGCTTCCAGATCGAGATCCCCGCAAAAGGCTCAAGAAAGGAGGCCACAAACAGACATCCCTCCACCTCCACCAGCTTCCCAAAAGGCGCCCGGGGAGGGTGCGGCATATTCTCCACCTCCTTCCCGCTCTTGATGCAGAAACTGGAGTCCTCAATTTTCTCCAACCCCGGCGAGCTCCTCCACGCATGAATCACTCCCTTGCACACCACAGGCTCCCGGTTCGCCGACGGCGCCTCAACCTCCCACTCCATCCGCTTCCATCCCGACGAGCTGCTGCTGTAGATGAAGATACGGCACTTGCCGGCCAAGGTTTTGGCCAGCAACACCACCTCGTACTCCCACTCCCTCCTCCGATCCTCCACAACCTGGATGAAAGCCCTGTGCAGCACCTCGAATTCCTCAAACCGCAGCACCGGCAACGCCTTCCACATCCGCATCACCGGATTGCACACGTAGAGGTACGTCTCCCCGTAGCCTTGCGTCCGCTTGCTGAAGCACAGCAGTCCCTTGTCCGACGCCCAGAGATCCAGTGGGAAGGAGCCGCCTCCCGGCAGACACGACGGATCCATCTTCCGCGGCAACAGACACGCATGCGGCAGCTGGAGCAGCAGCTGCTGCTGCCCCACCAGCTCCGCCCCCCCGAAGCAACACAAGCAAGCTTTTGCAGTGCCTGCCGCGGCAGCAGAGTCACACTCCCTTGCGAAGGCGGGCTCTGAGAGCATCGATTGCCATCTCCTGCACACGCACATGAACCGCCGGTGGCTCAAGATCGGCACTCTCACCAGCACCATGCGCAGCACATCCTCCGGCAGCCGGCTCCACACGTCCCCGTCCATCTCCGCCTGTGCCTCGCTGCACGCCTGCTGCGCACCTTCAACCTCGTCCTCCATGAAGAGGTAAAGGCACTCCCTCCCCGCCATCGCCATGGCTGCAAGCGCGAGTCACATTGTGGAAACACAAGCGCTTACACGCGCTTCATGAATCATGAGATCGGACGGGGACAAACTTTCAAGTTTTTAAAAAAAAGTCAGACCAGAGCTCCTGGACACTGAGGCTGCACCGTGGCACATGCAAAACGATAAGGAAGCACGTGGTGGCCAATGGTCCATATTTCGTTTCCAACAAAAAAGATAAGGAAGCACGTACGAGCTCCGACATATATGACGTGACTCTCGGCCCTCCCGCTCGGTACGCATGAACAGCGCAGATTGCGCAGTGAACAGCGTCTTGGATCTCTTTAAAGAGCAGGTTGCAACTTGCAAGGCGGATTCAAGCACAAGAGGAAGAGCGCATAGGTGTTGATCAGCAATGTCCGCCAAGACGAAGCACAGCGTGCGGGTGGCGCAGACGAGTGCGGATGCCAGGCTTCATGCGGCGTTCGAGGGGTCGGGCTCGGGAGGCGGCGCCGGGGATGGCGGCTCCGGGGTGTCCGACCACTCCAAGTCGGGGCAGCCATCATCCAGCGGCCGCACCGGGGTGGCTCCGGAGGCCATCACGGCGTACTTGCAGCGAATGCAGCGCGGGGGCCTGACGCAGGCCTTTGGGTGCATGTTGGCCGTGGCGGGGAACAGCATCCTGGCTTTGAGCGAGAATGCGCCGGAGATGCTGGAGATGGGGGTAGGGCTGGGCACGGACCTGCGGATGCTGTTCACATCGGGGAGCACGGCGGCGCTGGAGCAGGCGGCCAAGGAGGAGGATCTGAACAGCGTGAACCCGGTTGTGGTGCAGAGCTGCGGCGGGTCCGGGAACCGGTTCTACGCGGTGTTGCACAGGATTGAGGATGTGGCGGGGCTGGTGATGGACTTGGAGCCGATTGAGGGCGGTCTGGGTGTTCATGTGGAAAAGAAGTCGAGTGCAGCGATGGCTGTGAAGCCTATTGCACGGATCCAGTCGTTGCCGGGAGGGGACATTGGGCGGCTGTGCCAGGCGGTGGTGGAGGAGGTGCAGGAGATGACAGGCTACGATCGGGTGATGGCCTACAAGTTTCACGAGGATGAGCATGGGGAGGTGGTGGCAGAGATGCGGCGCCCGGACCTGGAGCCTTACCTGGGCTTGCACTACCCCTCCACGGATGTGCCGCAGGCGTCCCGGATGATGTTCATGAAGAATGGGGTGCGGATGATCGGGGACTGCACTCTCCCGCCGGTGCGGGTGGTGCAGGCGAAGGAGCTGGCCCAGCCCATCAGCCTGGCGGGGTCCACTCTGCGGGCACCGCACGGATGCCACGCGCAGTACATGTGCAACATGGGCTCTGCGGCGTCGCTGACCATGGCGGTCGTGATCGACGACTTCGACGACGATAGCAGCTTGAGCGGGAGCAGGAAGCTGTGGGGACTGGTGGTGTGCCACCACACGTCTCCCAGGAAGGTGTCGTACCCCCTGAGATGCGCGTGCCAGAAGCTGATGGAGGCATTTGGGGTGCATCTCAACATGGAGCTGGAGTTTGCTGCACAGCTGAGGGAGCACCACATTCTTACGACGCAGACGCTGCTGTGTGACATGTTGAGGATGATTCGGGGCACGCCCATTGGGATTGTGTACAGGTCGCCCAGCATCATGGATTTGGTGAAGTGCGATGGCGCCGCTTTGTACTATGGGGGGAAGTTGTGGCCTCTGGGGACTACGCCCTCTGAGTTCCAGGTTCAGGACCTGGCGGAATGGCTGCTGGGTTCCTCGGAGGAGATTGCCAGCAGTGGAGTGACTTGTACCGACAGTCTGGCGGAGGCAGGGTATCCTGGTGCTACCGCGTTTGGTGATGCGGTGTGTGGCATGGCGGCTGCTAGGATCACGCCCAATTCCGATTTCCTGTTTTGGTTCAGGTCGCATACTGCAAAAGAGGTGTTCTGGGGAGGAGCTGAGCATGACCCTCAGGCCAGGGATGATGATAGCCGGCTGCTGCCCAGGTCGTCGTTCAAGGCGTTTTTGGAGATTGTGAAGAGGAGGAGTTTGCCGTGGGAGGAGGTGGAAGTGGATGCTATCCGCTCGCTTCAGCTCATTCTGAGGGAGGACCTGGAGGAGTTCTGTGCTGCTGCTGCTGCCGGAGCTGACGTGGAGGATGTGGATAGCCTGGTTCCTTTGTCCATAAAGCTGTCGTTGAAGGAGAGCGAGGAGATGGGTGGTGCGGAGGATTCGAAGAAGCTGGAGAGGGCAACCAGTGCAGCTGCAGGAGGAACAGGAGGAGGAGGGTGGGAGAAGATGTCGCTGCCTTCCAGTCTAGCGCAGGAGTGGATGGAGGCAATTCGAGGTACTGGAGATGGCGGGGCCAGTGGCGGAGTTCCCTTCGATTGGGATGCCATCAGTGTGTTCCAGCAGACCAGCTTTGTTGTTGTGGATGCTCTCAAGCCAGACTTGCCCATCATCTTTGCCAGCACAGGGTTCTTCAACCTTACCGGCTACACCTCCAGGGAGGTGATTGGAGGCAACTGCAGGTTCTTGCAAGGGCCTGACACCAATCCTGAAGACGTTGCCAGCATCCGGGATGCTGTGGTGCCACGAGGCACAGGCACCTTCTGCGGCAGGCTGCTCAATTACAGGAAAGATGGCAGCAACTTCTGGAACCTGCTCACCATAGCTCCCATCAAGGACGACACTGGCACCATTGTGAAGCTCGTCGGGGTGCAGCTAGAGGTGAGCAAGTACACGGAGGGATCCAGAGCAAACAGGCTGCGGCCCAACGGACTGCCGCAGTCCCTCATCAAGTATGACGTCCGGCACCAGGACAAGGTTTCCGCCTTGGTTGCGCAGATAGTGGCTGCTCTGACCAAGCCATACAAGGTGGAGCCTCCCAGGCCGAGCTACGCCATGCGAGCCTCGCTCACTGGGCAGACCATCCAGCCCTTATCCCCGGGACGGGCTGCTGCTGCGCGACCCTACTCTGCCTCCGATGTGCCCCAGACTGCTGCCATACCCAGGGAGGGCGGAGGTAGAAGGCGGCACCGGTCCTCCACCTTCCTGTCCTTGCTGGGGATGGAGGAGAAGGATTCGGAGGAAGATCAGTTCCCGGAGCCGGAGCTTATCATGATGGACAACGCTTCAGTTGGCCGGCCCGGAAGCTTGGACGATCGGGAGCGCACCAGGAGGGGCATCGATCTGGCCACCACGCTGGAGCGAATCGGGCATAGCTTCGTGATCACAGATCCCAGGTTACCGGACAACCCCATCATCTTCGCATCGGACCAGTTCCTGGAGCTGACAGAGTACAGCAGAGAGGAGGTGCTCGGCGAGAACTGCCGCTTCCTCCAGGGCCGGGACACGGATCTCAAAGCGGTGCAGCTCATCCGAGATGCAGTCAAGGAGGGGCGCGACATCACAGTGCAGCTGCTCAACTACACGAGGGGCGGCAGGCCGTTCTGGAACCTCTTTCACCTGCAGGCCATGAGGGACAAGAAGGGCAACCTCCAGTACTTCATCGGAGTGCAGCAGGAGACGGACACGCTGGACAGGGTGGAGCAGGAGAAAGCTGAAGTGGTGCGCGCCACTGCCCAGAATGTGGATGTGGCTGCCAGGGAGCTTCCCGACGCCAACCTCACCCCGGACCATCTGTGGGAGCGGCACTCCAAGGTGGTCACCCCCCTCCCGCACAGCAAGATCAACAGCCCCTGCTGGTACGCCATCCGGAAAGTGCAGCGGCGGCTCCGGCGAGGGGAGAGGCTGGGGCTCAAGCATTTCCGGCCCATCAAGCCGTTGGGATCGGGAGACACCGGGAGCGTGCACCTGGTGGAGCTCCGGGGAACTGGGCAGGTGTTCGCCCTCAAGGCCATGGACAAGTCCATGATGCTGCACCGCAACAAGGTGCACCGCGCCAGAGCGGAGAGGGAGATTCTAGGCATGATGGATCACCCATTCCTGCCCACCCTGTACGCCTCCTTCCAGACCAAGACCCACGTCTGCCTCGTCATGGATTTCTGCCCCCGAGGCGACCTCTTCCTGCTGCAGGACAAGCAGCCCGACAAGACGCTGAGCGAGGAAGCTGCCCGCTTCTACGCCGCCGAGGTGGTGGTGGCCCTTGAGTACCTCCACTGCATGGGTGTCATCTACCGGGATCTCAAGCCCGAGAACCTGCTGCTGCAGAAGAACGGCCACATTTTACTCACCGACTTCGACCTCTCCTTCCTCACCTCCTGCCACCCGCAGCTCATCACGAGCGGCCGCGGCAGCCGCAGGCGCATTAAAAAGAGGCGAGCACGGGTCATCTTCTGTGCAGAGCCGCATGTCTCCTCCAACTCGTTTGTGGGCACTGAGGAATACATTGCTCCTGAGATCATTTCCGGGCACCCTCACAGCAGCGCCGTGGACTGGTGGGCATTGGGAATCCTGCTCTACGAGATGCTCTACGGCCGGACCCCTTTCTGCGGCCGTAACAGGCATAAAACCTTCATCAACGTGCTGAACGACGAGCTCACCTTCCCAACCAGCATCCCTGTGAGCGTTGCGGGCAGGCAGCTGATAGCGGGCCTGCTTCAGCGGGATCCTGCCAGACGGCTGGGCGCTTTTCGGGGGGCAAGCGATGTCAAGAAGCACCCCTTCTTCCAAGGCATCGACTGGCCTCTCATCCGAGGGAAGAACCCACCAAATAATCTGCACAAGCTTAATGCCACCACTTTCCACGAGTTCTGATCAGGAGAGCTTGTTTTCTCAAAATTGTTCATTTTTCTTTTCTATATATAGTGAAAGAGCTCTTTTTGTAGAAATTTAAGAGCTTGAAAACATATTCTTAAATGTGTCTACCCTTCAATTAAGAGTTTGGATAATTTATTCATTGTATGTGAGCAAGGACACTAACCTTAGATTTATCTTATTCACTATGTGTAAAATACTAGGAGTGACAGTCCTCATTTATAAACTTGCCATTGATTCAGGCTTTGCATACTTCATTTGCTGTGCAGCATTGCCTTGAAGTTCAGTTTTCAATCTTTCAAGTTCCAGTCGAGAGACTCTTAAGCATGCAAGCACAGACTTGAGATTGTTTATTTCTTGTGTTTGAGAAAGTACCTCTTCATCCTTGCGAGCAAGCTGCTCCAATAAGACCTGATTCTCTTTCGCAACGCCATCCTTCTCCTTCTGTAGAAATGATACGTTAGCTTTAAGTTCAGAGTCCAACCTGCTTGCTTCATCTTCCATAGACTTAAGCCTCTCCCTCAGCTTTCTTATTTCCTTATTGTATCTTTCAACCTCTGCTTCAGATGAAGTTGCTCTTTCCAGCGCTCTCACCAAATCCACTTGCAAAACCTCCACGTCTGAAGAGCCCTGATGTTGTCCATTCACATTCTCTAGTGCACATAGTTGAGCTCGTAATTGTGCTGAGTCTGCCAGGGCGTGTACCAAATGCTTCTCCTTCTTCTCCATCTCCTCTTCTAGTGCTAACAAGCGGGCCTCGTCTTGTTGTGCCTCTTTCAACACGTTCTTAAGCTCTCGAGTAAATAGTTCCTTGAGAGAGGAGATGCGCCATCGCTGGAATGCAAGTGCGGCATCAGCGGCAGAAGCACTAGTGCTGAGGGTAGCCTTAAGATCAACGGCACAGTCGAGGGCAGTCACACCCGAGAAAGCAGAAGCTTCTGGTTGTGCAGAAATGTGGACAGGGTCTGGCGAAGATGATGAAAGAGACGTGGCTGGTTGGCCATAGTCATCCTCTCCATTGTTGGAGCTCTGCTCGTCGCTGTAGTTAAGCACGCTGCCCAGCCTCGTCAATTTGCCAAGCTCATCCCGCACCCTTTGAAAATTAGGGTTTGTATTACGAGGGCGCTGCACGGCAGAAAATTCTTGAGGCGTGACAAGTGCCCCCATCCCACGAGTAGGCGAAGGACTTTGAGAAGGTGCTGTCACGAACAAAGCTTTGCTGGCCCTTCTCGACTTCACCATTTCCCCAGAACAACAACGGGGGATGACAGTTGAGTGCAACTCCAAAGCTCCCATGCAACTAATTG

**CRYPTOCHROME 1 (CRY1):**

GTTTGCGCAGCTTCAATGAGGGTTTTTTAGGGAAGACGATGCTTCTTCTGTTGAGAAGAAATTAGGGTTTCTTCGAGCTCGCGTTGAGAAATCAGGGCTTCAGTCTATTAGGTGTCCTTCTTGCTTTAGCAGCAAAGCCACGCACCAGCTTTTCAGTGTGGTTTCCTCCTCGCCATAGCCAAAATCTTCTCCAGCCAGCTATATTTGCTCCTCATCCTCATATCCGATTCCCGCTGCCCTTCTTGCCAATCGCCCCCTCTCCTGCATCGAACGTCGAGCTTCCTCACTCCTCATTCTCCTTTTTATGCAATTGCCGAGACCCCCGCTTACATCGCGAGGTCATGCCGGGAGACCCGCGCACTGAAGTTTGAGCATTACCCTCACCCTCCTCAATGAAGTTTGAGCTCGCCCTCTTCCACACCTCCCTAAATGCGCCATCGCAGGTGTAATCGAAGCAGAGACTTTGCTGAATGCAATAGAAGCCTTAAGAGAGAGGGTGTGCAAATTGGGTTTCTTTTTTTTACTGCCGAGGGCTTGTGGAAGACCCACAAAAGTAGCACAAAGCTTGCCAATCAATTTTTACTTTGCTCATGGAAATGAGCGAAGCATTTATCAGCTTGATGTAAGGGCTGAGAGGCAGGTCTTGAATCTTGACAGTGCCAAACAGGGCTTGTTTGAAACAAAAATCTATGTAGAGCCAACTTCCCAGTATTTGTGTTGAAGTAACCAAATCGCCGATGGCAAAAGCAAAAGCGTGCACCATAGTGTGGTTCAGACGAGATCTCCGGCTAGAGGACAATCCTGCGTTGTCGGCAGCTGCAAGGGCGGGCTCTGTGGTGCCTGTTTTCATCTGGTCGCCTGAGGAAGAAAGGCAGTTCCACCCGGGACGAGTATCGCGATGGTGGCTCAAGCAGAGCCTCGTCCAGCTCGACACGGCCCTACGCAGCTTAGGTGTTTCCTTGGTTTTGATAAAGACCCATGACACCCTTTCCGTCCTCCTAGAGCTTGTCCAGACAACAGGCGCTAGCCAAGTTGTTTATAATCATCTATATGATCCCATTTCTCTTGTTCGAGATCATCAAGTTAAGCAGGGCCTCACAGAAAACGGCATTCAGGTCCAAAGTTTCAATGGTGATCTACTGTACGAGCCATGGGAAGTGCTTGATGATCATGGAAAACCTTTCACATCCTTTGAGAGCTACTGGATGAAGTGTCTTAGCATGCCCTTTGAACCTGATTCTCCCTTGCTTCCCCCAAGGAAGTTAATGGCATTCCCAGGCAACTTGTCTTCTTGTACATTGGAAGAACTTGGGCTTGAAGATGAAACAGAGAAAAGCAGCAATGCCCTTTTGGCAAGGGCTTGGGCACCAGGTTGGAGTAATGCTGACAAAGCTCTGGAAGCATTTCTTGCTGGTCCGTTACTGGATTATTCTAGAAATAGGCATATGATAGATGGGCCCACAACTTCCTTGCTCTCACCTCACTTGCACTTTGGGGAAGTCAGTGTTCGGAAAGTGTTTCATTCTGTTAGAAGATTGCAAATTTTATGGGCTAAAGATGGTTCAATGCTAGGTGAAGAAAGCATCAACTTTTTTATGAGGTCTATTGGTTTTCGGGAATATTCAAGGTATCTTAGTTTTAATTTCCCATTTACTCATGAGAGGTCTCTCCTTGCAAACTTGAAGTCGTTCCCTTGGAGGGTTGATGAGGGGTTTTTTAAAGCTTGGAGGCAAGGCCGCACTGGATATCCATTGGTGGATGCAGGAATGAGGGAATTGTGGGCTACAGGGTGGCTTCACAACAGAGTACGAGTCATAGTATCCAGCTTTTGTGTAAAATTTCTTCAACTACCTTGGAGATGGGGCATGAAATATTTCTGGGACACCTTACTAGATGCAGATTTGGAATCAGACGTGCTTGGTTGGCAGTACATATCTGGAAGCCTTCCAGATGGCCATGAGCTGGACCGAATGGATAATCCTCAGACTGAAGGATACAAGCATGACCCACTTGGTGAGTATGTTCGGCGCTGGCTACCTGAACTTGTTCGATTGCCCACAGAATGGATTCATCATCCATGGGATGCTCCTCCAGCAGTCCTTCGAGCTGCTGGTGTTGAGCTAGGGTCTAATTACCCTCGGCCCATTGTAGAAGTAGCAGCAGCCAGGGAAAGACTGCAGCAAGCAGTTGCACAAATGTGGGAGCAGGAGGCTGCTCTAAAGGCTGCTGGAGTCGTTATTGATGAGGAGGCTTTAGATGAGAATTTGGAGGTGAAGGTCGTGACTCGGCCTTTAGTTTCGACACAATCAGTCAAGGCAGCCATGGCACTCAATCCAGCAACTGAAGTTCTGCAACTTCCTTCAAGAAGCTCTCACCGGGATCAAATGGTACCTTCCTGTGCCAAGCAACTGGAAGTGAAGACTGGTCTTCCACTTGAAGGTCTCAATTCGTCAGTCTTACCTGAAATTCCAGTGTCTTCACAGCAGGAGAATTTGTTAACGTCTACTCCGACTCCTGAAACAAGAAATATATCTCAAGCCTCACCAGAGAGTTGTGGTAAGTTGATGAACTGTGCTGAGGCTCCTAGGCCAACATATACTTTGGCTCCGCCTCCAGTCTCTGACATGAACTCCACAGCAGAGTCTTCCTCTTCTCCAAAAAGGCTAAGTGCCGAAATAAGAAAGGATGCATCCATGATGTGGGTGCCTGCAATCTTGCATCAACAGCCGTCAGACTTTGGATGTGTGCCTTATGCAGGGGATGCAGGACAATTGAGAAAACATTTGCGCTCTTTGCAGAATTCAAGGCCAGATAATAAGGTGGAAGAGGGGGCCCAACTTTCTGGTGTCCAGTGGCCGCGAAGAACCAAACGGAAGGCTAGAGCCTAAATTCTGCCAATGTTTAGGAGTCTTCGCGGTATAGTCGAATTCTCCTAGTGTCTTGCAGAGTGTTTGGCTTTCAGCGTGGTTGGTATACCAAAATGTTTTCTGATGTGTACAAAGCTCTCAGAGAACAAGCCGTTAAATGCCTTGTGATCAGCAGGTGCAAGTGTTGCAGCAGAGTCATCAATGATCACACCGCCCCATTCCGAGCTTTAATGAATGATTAGTCATGATTAGTCACAGTAAGCCATGGTATGGTGTTATCCCCGTCTCTTGAAGAACGTGGCGAATAAAAGGGAATAAGCTTCCTCCCGTAGATTTCTTTGGGGTTGTGTAAAGTTGTCAATTCCATTTGCACTCTAAATTTAAGTGAAAATCTGTTTTGTTGTTTTTCCCTCGTTCCATTATCTAGGAGGTGCGTAAAACAATGAATGCCAAGGGTTTTGTGGCATTTTTGTATGAGGAGCTGCAGTTGTAGTAGCAGCACGTTGAAGTCTGAAATGTGCAGAGGCTGCATAGCAAAGCCTGGAGTTGATTACCTATCGGGTTTTTTCTGATACTAGTAGCACCTGGCTCATGACTGTTATCTACTACCTTTATTGTATTTTTCAGGAGGCTTTCAGGCCAGAAAGAATATGCAATAGATAGCCCTTGACTATGGTAAGCAATGAAGACATGGAATAAAAGGTGGTTGATAGTCTCAGAACTGAGCAAATACAGATGCTTGTATTAGCACAGTAATTGTTATCTCACTTTGCACTCAATCAGTCAGATGATGCAAGGCTGCTGTCCAGAGTATGAACAGTTGAATAAGGATGTTACCAGGTTTAGTGACTTTGCCGCTTGAGTATGAATGTACTCTTGCATTGGCTTGAAAAGAATGGCTACTTGTCTGCCCTGCTTCATTGAAGAAGTGGATTATATATAACATAATACCTCTTTAGATTGTGTGTGATGGTACTTTTTTGCGCTGTGAGAAATTTGAGATGAGAGCAACTTCAAGTCGGTTTAAAGCATAGTGAACAACAATACATATAGGACGATAATAAGATGTAATTATCAGGTTGGATACTTTGTTACTGTT

**CRYPTOCHROME 2 (CRY2):**

GAAGAATTTAGAAATCAAAAGGTTAGTGTTGGCTGAAAGCGTCACGTGAGTGTGTAGCACGTGGCAGAAGCTGGTTGATTGCACTATGATGTATTCACATGGTTATGGTGGGCTCACCGCCACGTGATTAACTCTTAAGTATGCGCCTCCCTAAAACCATCCTCAGCAAAGCAAAGCAAAAAATCACACAGTATATATATGCTCTCATGAAGATCAAACCCTCTTAATGGTATCTTCAGATCTGTGCATCTCGCCATGGCTGCAACCATTGTGTGGTTTCGCCGAGATCTGCGCTTGGATGATAACCCAGCACTCAAGGAAGCATCGCGCAGCGGTGGAGCCGTGCTGCCGCTTTTTATCTGGGCTCCTGAAGAGGAAGGCCAGTTTCAGCAGGGCCGCATGTCTCGATGGTGGCTTCAGCAAAGTCTCATTCAGCTCGCCCAATCCCTTGAAAGGCTTGGCACTCCCTTGATCGTCCGAAGGTCCTGCAGTTCTTTGCACGCCCTCATCGACATTATTGCACACTTCCAGGCATGCCAGGTCTTTTTCAATCAGCTCTATGGTATGTGGCGCATTGGAACTCAGCTGTGTGTGTGCATTTTGTTTAACTTTTTCAAGCTTCTCCTCTAGGGAGCGTGACGTGCTGTGCACACAGAAACCACAGCCATTTGACATGTATGTGGAACACCTTAATTCGATGAAAGTAGCTGTAATATTTCTTTTTGAACTGGTTTATTGAATGATGAATACACATTCATAACTTATTAATCATGGGACATCCGGTGATCTTGTAAGACAGCATGGGCTAGGTTACATGCCTTCACACGGAGTACATTGCTCATTTGTGCTTCGACATGATGCATTGCAGCCAATCCCATCTGGCCACGGCATTTTGCATGTCATTTCATCCAATGTTTGTCTCATGAAAGATTGACACAGACGTAATGGGCTACCAGATGGTAAAACTGCTTTGCAAACAAATCTATGTAGTGACATCGTCTTTGATGTTGAGGATTGCAAGGGGACTTAGGGTTCATGAAATTACAAACATATGTTACCCCACGGCATCCTGTACACATGGTGCGATTATCATTGTGATGCAGTCGGAATCCAATGTGCACTGTAATACATTATAGAGGTTTCATAAGTAAAACCTTCATGATCGAGTACATAATTTGAAACATCATTGCCAACAGATGCACACTGCAAAACTGCTTTTGAGCATGGTTCTCTTGGAAGACAAAATGTTTTGTAGAGAGCAACAGCGCACACTACAACTTAGTAACAACCAGTTGAAAGGGAAGATAAATAACTTTATTTTTATAATTTATAGGTCTTTTCAACAATGGTAGTACATAAAGTAAGGCTTTTACAAACGGCTTATGCTTGGTTCATTAAGGGATGTCCTCCTATGCATCAGGTTATGGCTTCTAGACGTTGCAGGTGATGTGCATTAATTCCGTTCCGGTTGATTAAAAATCCTTGTGTTTGGCATAGCTAATCATGAGTTTTGAATATGAATGTGCAGATCCCATATCATTGGTGAGGGACCACAGGGTCAAAGAGGGATTGTTGCAAATGGGTGTGGATGTGAGAACCTTCAATGCTGACCTGCTGTATGAACCATGGGAAGTGAGCGATCCGTCAGGCAAGCCATTCACCACATTCAATGCATTCTGGCAAAAGTGCCTTAACATGCATGTTGAGCCAGAGGCACCATTGCTGCCTCCTAAGCGCTTGCTCGGATACACAGGGCATGTTCCTAGCTGCACGATAGAGGAATTGGAGCTTGAAAACAGTACAGAGAAAGATAGTAATGCCCTCCTAGCAAGATCGTGGACACCAGGTTGGAGTAACGCTGAAAGGTCGATGGAGACTTTTGTGGAAAACCATCTATTACAGTATGCCATCAATCAGCACAAGGTGGATGGGTCCACCACATCGTTTTTGTCACCTCATTTACACTTTGGTGAGTTGAGTGTGAGGAAAATATTCTACCATCTCCGTGTCAAATCAGTAATGTGGGTGAAAGAAGGACATCCAGGTGGAGAGCAAAGTGTCGATTTGTTCCTAAGGTCCATTGGATTCCGAGAGTATTCAAGATACATATGTTTCAATTTTCCTTTCACACATGAGAAACTTTTACGGCAGGATAATAGAGCTTTTCCATGGCGTGTTGATGAGAGCTTATTCAAAGCATGGAGGCAAGGCAGGACAGGGTATCCCCTTGTAGATGCGGGCATGCGCCAGATATGGGCAACTGGGTGGCTACACAATCGCATTCGGGTTATCGTGTCTAGCTTTTCTGTGAAGTTCTTGCACCTACCTTGGACATGGGGCATGAAATACTTTTGGGATGCTCTTCTAGATGCCGATCTGGAGAATGACATCCTTGGATGGCAGTACATTTCAGGTGGTCTCCCAGATGGGCATGAACTCGACCGGATGGATGATCCACAGATGGAAGGCTATAAGTATGACCCACATGGGGAGTACGTGAGGAAGTGGCTGCCGGAGTTGTCAAGGCTACCCACGGAATGGATCCATCATCCTTGGGATGCTCCCCCAAGGGTCCTCCGAGCCGCAGGAGTAGAATTGGGATCCAATTATCCCCGCCCTTTAGTGGAATTCGCAGCGGCCCAAAGCCGCTTGGAGAAAGCGCTACGGGAGCCATGAATCCAGAGGTTTTTACGTGTCCCAAGTCATATTGAGGATCCTCAGAAAGAAATTTTCACACCCTTTATGAGTCGGCGTTACAAAAAGGGAGGATCCTTCAGAGGGGATGTTTGGTGTTGGTACTGTTTTGATTGATCCATCCATGGCACGGTCATGGAGTTTTCTTACATTACACTTCGTTCATCACCAAGGAAGGTACGCATCTACCTGCTGCTTGGTCTGAGGTTCACGAACGTCGAAGTCCATCAAATAAAGTGTATCGAATGGTGCTTTTATTTCTATTGGGCAGCATCAAAGACCGTGGAATTCTAACCTTTAAATGCGCCCCTCAAAGATGGCATTTTGTAATTCTGCAGGTGAGCTCTTCTTATAAACCATTGCATGTGGCTTCACGGCATCCTCCAGCATTGATGTGATGTTAACCCCTGAATGCACACAGACAAGCCAATCACAGAGATGTCCTGTAGATAGATGCAGGAACTTGTACCTACAAAGATGTGCATTCTTCGACATAAATGTACACAATTATGTGCATTTGCATTCTAACAGCACACACGCAGACTCGCAAATAATGGAGCCATGCTAACAGACTCGCAGACGCACACACATGCATTTGCATTCTAACAGATTTTTACAATTGTAATGATCAAGAAAAGTGCAGCTGGAATTTCACAAGAGTTCAAAATGACACCTTGGCATTTGGGAAAGGATAAGCAGCCTTCACAAACACGGTGTGCAGCTCGGCGTTTCAAGTCTATCAACACTGTTCGAAGTCTCATGAGTAGAAGATACTGTAGCTGTTTCATAGCAACCACAATGAAGGCTATCAAAATTATATGTATGCAGTCTACTAGTACTATAAAAGGCTTGTTCGAAGCTGTTGCCAATAATCAACTTTTATCTGCATTGTGGGTTTGCTTTTGATCTCTTCCAACTGATTGATTGTATTGGATTATTGTGAAGCCTGTTTTAAGG

**G-PROTEIN COUPLED RECEPTOR 1 (GCR1)**:

AGAAAAAAAAAATGTAGCTTTCAAATAAACGGCTATGCATGAAAATATAATACAGCATGTGATTGTGAGAATCTGGCTGAGCATATGTGATGTGTATATAAGGTATAATTTACATGGGCTAGCATGGGAAATGTGTGTGTGCAAAAGAGTACACAAAAAGGTTTCCACAATGTTTACATGTGACTAATGAATGTAAAAACCCCCATAGGGTTGGTAGGGCCAACAGCTCCCAATCTACATAGGACAATTTTGCAGATTGTTTCTAGTTTTGGTCATATTACACTTTGTCTTCTTGACTATCAAGCTGTGTCAACTCATTCTCTTGAATTTCTCGAAGGCGCATGAAACCAGCTCTCCTCCAATATAGTTTCTCTGGCCAATATTGCTCAGTTTTTTCTTGAAGTGTCCGGCGTACGGAGGAATTCAACCCATACGCAATGGAATTAAAAAGCCCCATCAGTGCAGAAGTTCCAACGTGCAAGCAATACAACCAAAACAATTGCTGATGAGGGGCCACAAAATCATGTATTCTATTGATTGTTCCAAATGCCCAAGATCCTATCAGAATTAAGGGATAATAACCCCATCTGTTCATAGCTTTCATGTCAGCCTTCACTTCTACTTGTCGTTGCCGATCAGACATTTCCACAGCCATTCTAGCAGCATAATTCAGCATCCGAATGACTTGAAAGTATATTACTCCATTGAACGAGATCGCTCCCCACAAGGGAACATAGAAAGTTAAGAAGCGAACAACCTTTCCAGCGTTAGTTTCATTTTGAACCCAGCACCAAGCACCTGCGTGCCCATAGTCGTTTCCTATTGAGGGAATGATTGCCAAGATCAGAGAGGTACCCCAAACATAAAGATGGAAGATTGGACCAAGTTCTTCTACATCTGTCTTGTGTTTAACCACCGTTCGATGGAGAGTAAAGGCTATTGTGGTTGTCCAAAGAAACGATGCCACGGAAAAAAATTGTGTAGTGTAGCCTTGAACATAGCACAGAAACCCCTTCCCTGGATCGCCGAGAAGATTGAAAATGCTGCAAAGCAAGTCAGAGAGGGAGAGGAAGAAAACGAGCTGGAAAGAGAACTTGCGAAGCTCTTTGAAGAGAAGGTAGCAGAGAACGATGAAGCCTGATCCGGCAAAGGAGAAAGCCGACGCAAACACGTTAATGCTAGTCAAGGTTTGTGTCTGCTCTGCACTTAAGCCTGTGCTCATGTTGCCTCCCTCCCTCTCTCTGTCACAATACGCTGCACACCTCCATCGCCTACCTTCTTTGCCATCCACAGTTGAGTTTTTGTCCGAGAGAATCTTTGGAAATGCTTTTCGAATCTTCCTCAAATAACGAGACCCACAAATCAAG

**ADAGIO 1 (ADO1)**:

AGAGAGAGAGAGAGAGAGGTCTGAGGATATATTATGCAAAAAAAATATACATATATATCTGTTTTTCCCTCATTTTTGTTGTGTGTGTGTATGTGGGTTTCCCTGCTTCTTCCGTCCTCTCTATTAGTGTGCTCTCTCTCTAGACTTTGAGTATGTGTGTATGTATCCCTTCTCTCTATCTTCCCTCTCTCTGTGTGAGTGTTGTATCTTCTTCTCTCTAGTTTCTTGAGAGTGTGAGAGTGTGTGTGTGTGTAATTGGTGCGTGTCGCGAATCATCGCGCTGGCCATCGCCATGGCCAACCTCTGCTTCATCTTGCGCGAGAAGGGCTGAGCTCCAGTTTGTAAGCTTCTCTTCGCTCCTGCTAGCTTGCTTAAGAATCTGGCTTGTTTACAATGGGCGATTGGAAGAATATTGATGAGGATGCAGATATGGTTGACATTGATTCATGTCTCATCGATGAGGAGGACGATGATTTCACAAATGATGATGTGTACCATCACCCTCAGTATGATAGTCATGGGGTTATGCATGGTGATGATGACCCATGCTATCCCCCTCCATACCATCAGTATGACGCTCGGGTGCCTGCGGAGGATGATGACGACGACACCCATCTGAGCCCAAAGCTAGAAGCTCTTGTGGGTGTGTACAATGCATCCCTTCACTGCGACCAGGATGGTGCCCGTATGGGCGTGTCTGATGATGATGCACACAACTTCCATTGCTACCACCATGACGATGCCCCGGTAAGCGAGCATGATGAGGAGGTGGGCATCTTTCCATATCCGAAGCCAACTGAGATGCTGTTTCGGACCAGCCCTTGCGGCTTGGTGGTGACAGATGCGCTTGATCCTGATCAGCCTGTAATCTATGTCAATGCTGTCTTTGAGATGGTCACAGGCTACAAGGCTGAGGAGGTCCTTGGTAGAAACTGCCGATTTTTACAGTATAGAGGGCCCTTCGCAAAGAGGAGACATCCATTGGTAGACACAACTGTTGTAAGAAAGATGCAAAAGTGTCTTGAGAACGGGCTAGAGTTTCGAGGAGAGCTTCTCAATTTCAGGAAAGATGGGACTCCGTTGATGAACAGATTGTGTTTAACTCCAATCCATAGCGATGATGGAACCATTTCTCACATTATTGGGATACAAGTGTTCATGGAAGCAAATATTGATCTAGGACCTGTAGCATGTAGTTTGTGGAAAGAGTCCCTTCACACAGCTGAATGGTACACTGAAGAATATCTACCTCACAGAGAGAAAGTTACTGCTTCCGACCAGCCTTCTCGCCAGCATCGTTGTGGACTTTTGGACCTGAGTGATGAGGTTTTAGCTCACAAAATTTTGGCACTATTATCACCGAGGGATGTTGCTTCGCTTGGTTCTGTTTGCACTCGGTTGTTTCAGCTTACTAAGAATGAAGACTTGTGGAAGATTGTATGTCAAAATGCATGGGGTGTGGAAACTACGGCAATCCTTGAAGCTGTGCCAGGAACGAGGAGGCTGGGTTGGGGGAGACTAGCAAGAGAGCTAACCTCTCTCGAAGGAGCAGCTTGGAGGAAGATTGTTGTAGGAGGGGCTGTAGAACCTTCGCGGTGCAATTTCAGTGCGTGTGCCGTTGGAAATCGACTGGTTCTCTTTGGTGGTGAGGGCATCAACATGCAGCCTATGAACGACACCTTTGTACTTGATTTGGGTGCGGCAGCACCAGAATGGCGACACATGCAGGTTAGCTCGCCTCCTCCTGGTAGGTGGGGGCACACTCTTAGTTGTCTGAATGGTTCATGGCTCGTAGTTTTTGGAGGATGTGGGAGAGAAGGGCTGCTCAATGATGTGTTTGTTTTGGATTTGGATGCACAGCAGCCTAAATGGCGGGAAGTAGCAGGGGGAGCTCCTCCTCTTCCCAGATCATGGCACAGCTCTTGCACACTTGACGGGTCAAAACTAGTAGTATCCGGGGGGTGTGCAGACTCCGGTGTGCTATTAAGTGACACTTTTCTGTTGGACTTAACAATGGACAAGCCCATGTGGAAAGAAATACCAGTGTCATGGAGTCCACCTTCCCGCCTTGGCCACTCACTGTCCGTATATGAAGGCCGAAAGATTCTCATGTTTGGAGGCTTTGCGAAAAGTGGGTCCTTGAAGCTAAGATCAAGTGATGTTTACTCGATTGATTTGGGTGAAGAAGAGCCTAATTGGAAGCATTTGAGCAGCAATCCTGGGGGTTCTGTACCTCCTCCAAGGCTTGATCATGTTGCGGTTAGTCTGCCTGGTGGGCGGGTGCTGATTTTTGGTGGCTCTGTGGCAGGGTTACATTCAGCTTCACAACTTTATCTGCTAGATGCAACAGAGGAGAAACCTACTTGGAGGAGACTTGATGTGCCTGGGCAAAAGCCAAAATTTGCTTGGGGACATAGCACGTGTGTTGTGGGGGGAACTAGAGCTGTTGTGCTAGGGGGTGAAACAGGCGAAGAGTGGATTCTGAATGAACTTCATGAGCTCTCACTCGCAACAATGTAGGGCTCAAGTGTTGTGCTCCCAATCTGCACCAGTCATGTCTAACCACATAGATGAAAGACTTTTTGGAAGAGGTCTGAGAAGGACAGCCAGATTTGTGATAGGAAAGTAAGATGTGTTATTACAGTGCGCACATCGAAGGCATTGATGGAGGCTAGCTTATCCAGCTTCTGTTCTTACCTGCCTAAATTTTCTTGTGTGGGGAAAGTGGCAGGTCTGGTCGTGGTCTTTCCAGGGAGGCTTATTTTCTGGTGTAAATAATTTGTTAGTGTACCATGTTGTGTCTTTCCAAATTCAAAGTTGCCGAAGGATTTGGGTCTTGTAATAGGCTGACTTAACTGAGTAGGTTAGGTGATTGGTTTGAATTTTTTCTACTAGACAATGTAAGTAATTTTTGAGTGGTATCATGTAATGTAGCATTCGATAGCCGTGTTGTTGCATAAGGACGCCCAAGTTTTGGGAGCAGCGTCTTTTGATGTATGAATAATTAACTGGAGCTCTGCACCAATCTCACTGTCCTCAACACCAACCAGCAGTATAGGTATTTTTTCTAATTATGTTGTATGGAATGTGTACTTATTCGCAATTGTAATAATCATTATCTTGAATTTGGAAAAAGCTGATTTTTTTGGT

**GIGANTEA (FB)**:

CCCTCTCTCTCTCTCTCTCTCTATTTCAAGCTCTGCTGCTAGCACTTAACCCTAGCTGATTCTATGTTCAAGGCTCCTCTTGTACGCTCTCAATTTCTGCATAGAAATTTCACAGAGATCTTCTGTTTTTCAACCTCCAAAAGCTAGCTCGGTTCTGGAGCGGATTGCATCAAGCAGAGAGCAATAGCTTCATGTCTCCTCCTCGCCAGAAATGGATTCGGCGGTTGGAGTCATCGTCGCTTGTCAGGCCACCTCCACTAAACCTGGACCAAAGACAGACGGAGGTACTAGCTTATGTTGAACTCTTTGCACAATTCGCATCTGAAGCTTTCGGTCATGATATCAGCGAGCTGGTGCGGAGTCATTATCCTCATGGTGAGGCATCTCTATTGGATGATGTACTTGCCATGTTTATCCTGCATCATCCTGAACATGGCCATGCTATTCTACATGCTTTGCTATCTTGCATCCTTGATGGAACCATTGTTTACGACAAGAAAGCACCACCTTTTGGGTCTTTCATATCTTTATTCAGCCCATCCAGTGAGAAGGATATTTCTGAACAATGGGCTCTCGCCTGTGGAGAAATTTTAAGAGTTTTGACACATTATAACCGTCCAATCTCAAAGGCTGAGGGTTGTCAAAATGTTGGAAGACCTACAGATGCTTCTGGTTGTGAAGCAAGTTCAAGCAATGGACCTCAAAATCATCAAGATCTTGAGAGAAAAACTCTCACAAGGTTACTGACTCCTTGGATAACTGATAATCTACTTGCTGCTCCCTTGGGTGTCAAGAGTGATTATTTTCGCTGGTGTGGAGGAGTGATGGGAAAGTATGCAGCTGGAGGGGAGCTGAAGCCACCGATAACAGTTGGGAGCAAAGGACCTGGGAAACATCCACAGCTGATGCCATCAACTCCTCGATGGGCTGTTGCAAATGGTGCTGCTGTAATACTTAGTGTATGTGATGATGAAGTGTTACGCTATGAAACAGCAGATCTCACAGCAGCAGCGGTCCCAGCATTATTGTTGCCTCCTCCAGCCACAGCTCTGGATGACCATTTAGTGGCTGGCCTCCCTCCGCTTGAGCCATTTGCTCGACTGTTTCATAGATACTATGCCATTGCCACACCTGGTGCAACTCAGCGATTATTTCTAGGTCTCCTTGAGGCTCCACCGTCATGGGCGCCCGACGCTCTTGATGCATCTGTGCAACTTGTTGAACTGCTTCGAGCAGCTGAAGACTACAACTCAACAATGCAGCTTCCTAGAAATTGGTTGCATTTACATTTTCTGAGACCGGTTGGAGCAGCTATGGCCATGCGGTCAGGAATTGCAGCAGATGCAGCAGCAGCCTTACTGTTTCGAGTGTTTTCACAGCCAGCTTTGCTTTTCCCGCCACCTGGGCATGCACAGGATCATCAAGGAAGGCAGTCGGTTTCTGGATTCTCAAGGCATCCAGAGACAAGTGTAAGGGAACAGATGGTTATTGCTGCTGCAAAAGCCACTGAAGAGGCAACTGCAAGAGGCATAGCATCATTGATGTGTGATCATGGACCAGATGTAGAGTGGCAAATATGTACACTCTGGGAGGCAGCTTATGGACTTCTTCCACTGAACTCCTCTTCTGCTAATCTGCCGGAGCTTGTTGTTGCAACACCATTGCAGCCTCCTTCCTTGTCTTGGAATCTCTTCCGCCCTCTACTTCGAGTGCTTGAGTACTTACCGCATGGGAGTCCCTCGGAGGCTTGCTTGCTTAGAATATTCACTGCCACAGTTGAGGTGATTTTGCACAGGGCTATTCCCAGGAAAGAGAATCAAGATCAAAGAACTGGAATTTTCAGAGCTTCTCAATTTGGGATTGGTAGGTTTGGTTTGAAGCAAGTGGCCAGCACAGAGCTGCGAGCCATGGTGCATTCTTTGTTCACCGAGGCTTGTCCATCTGCAGATCTTGCGGCAAGATTGCTTTCAGTTGCTTTCACAGTTTGCTTGAGCCATGATGCAGTGCAACAGGGCAGTAGAAAGACAGAAAAGTCCCTCGTAAGCGAAGAATTAAGTGAGGAGGGAAGGGTTAGGTTGAGCACTGGGAAACAGAGGGGAGCAGTCTCAACTTTTGATTCGTATGTTGTAGCTGCAGTGTGTGCCTTGGCATGTGAAGTACAGCTACTTTCTCTAGCACCTATGTTAGGTGCATCAAAGCAAGGTTCATTCCAATCAGCAGGACCTGGAGTAGGTGGCTCAGGGAGTTTAAATAGTTTACAGGTTGGCGTGGTCTCTGCAGTGAACAATGCAAGGAGACTCCTTCGGCTTTTAGAAGCTCTACTGTCCTTGATACCACAATCATCATATGTGATTCCAATTAGTGGAAGCAGCTCCTCTGGTGAAATTGTTGCTGCTGCTGTGGTAGCTGCTCATATATCAGAGGTTCTAGGGAAGTCGAAGGCTTGTATGCATGCTTTGGGAGCCATTATGCGCTGCTCCTGGGAGCCTAAACTGTCTGGAAAGGCTGCCTCGGTGCTAGCTCTTGTTGAAGCCAATAGAAAAGTTGTGGCTGCAGTTGTTAATTCATCTGGAAGCAGTCTTTCAAATGAAGTTGTCAATGAAGAAGATGTGAAGGATTGTCCAGCTGAAAGTAAAGCAGTCATTAAAGGAAGAGCACAGGGACATTGCTTGGATCCCAAGAATGTCTTTCAAAAACTGGAGTCATGTCATGGCTGTGACACAGCAATAGATGATCTTAATGGCAGCTCTAAGAGGAATTTTGTAAGCGTTATCTGGAAGACCTGTTCAGAAAAGAGCTCTGCAGGCCTTCCCATGAATGCATCAGACGTTGTTTCGCTTTTGTGTATGGACAAGGACAATGCCTCAAATGACAACATTGGCTCTTTTGTAACAACTGTGCTACAAGAGAATAAGGATTTGTCAATAGCAGCTGTGCCATTGCTTTGCAAAAGGTTGATAACTGCACCCGAGGTCCCCACTTATGGGGAGGGCACATCTGCCAAGAAAGGATGGCGACAGGTTGTGGAAGCTCTTTGTAATATAGTCTCAGTGTATCCTGAGAAAGCAGCGGCAGCAATCATGTTTCAGGTAGAAAAGGATCTTCAACCGTGGGTATGTAATAATGATGCACCAGACTTTGTGGATGTTTGGCGAGCGAACAAGCGCATTGTATACCTTCTAAGTGAGATTCTACGGCTACATCACGCACCTGAAGCTCTGAAAATTGTCGCCAAGCGTTCTTATACCCTTGAACGGGCAACAGAAGGACTGGCAGTGGATGGTGAAGCGTGTACTATGCCACAGCTTGAGTTGCTGGAGGCTGCAGCAGTTGCTACTCAACTCTTGGTCATAAATAACAGCCCTACTGTAGAATCACTATCGATTCTTTTAAAGATGCGTCTTCCTGCAATTGTACATTGCTTGTCCCACAATAGTGCTCATGTCAGGGCATTGAGTATAGCACTTTTGCAGAATGAGGTGTACTCTAACTCAGCAAGCCTACACTTAGAGAAAGGCAAGATACCATCAGAGGAATGGAAGCTGTTAGTTGGGCAGTGTGTTGCACAAGAGATGCATTCTCTGCAGGCAATGGGAATGCCCATTTCGACACTGGTAGAGGCGGCGACTTCCTTGGGTTGCTGTGTACTCGCTTGATGCAACCTTATTTGTGTAAAGTGTTTGTTTGGAAAGTGTCCTCAAGTTTTAGTTCTGAGGGGAGCTCTAGCTTCTGAGGCATTGCCATGCTCAGGTAGTCATTGGAACTAACGCAGCCGTGCTGAAGCTATGTTCATTGACCACTTGTGTTTTGTATCTATGTGTTGTAAAGGCCCCCTTGGTTATGGGGCTGTAAAGGAACAACCCACAATTCCTCAAAGGTTGTATGGGGATTCAAACAAAGCTAACTCAGTAGAAGTGGAAAAGATGACATCCTGATACATCTTAGCTTAATGGGTTCCATAGGTGCAGAGCTTTTTCATCTTTTCTAATGATGCACCAAAGCGTGTATCGCATTTGATTGTTTCGGTTAAGGTAATTGCATGATTGATGGCGGAGACTGCACACTGTTGTATAGCTGGTCCCTCCTTCCTCGAAGGGTGAGATGCAGGGGCAGCATATTCAGCCTTGTGGATAGCATTTTTTGTTAAAGTAGTGAAGAGGGTTCATGAGCATGTGGAATTTTAACATGTTCTGCACTTCTTCTCCTGTATGATTGGATTAGTTTGTATGATTGGATTAGTTGCTCTTATTTTAAAAATTGTCCTTCTCTACTGCTGGGAAATTTGTTCTTGTTGAGAGTTCCTAGCAATGTGATCCAAGC

**SUPPRESSOR OF PHYA-105 1 (SPA1)**:

AATGGAGAGAGTTCCGAAAATCCCAATAAGACAGAGATCTGTGGAGTGTGGATACTAGCAGTCTCTCTCCGTACCACTACCTTACCTCCTCTTTCACACTGGCTATGGCGGCAACAAACGCTTGAAAACAACCCACATAGCAGAAAGAGAGTTTGTGCCTCTTGGAGATCGACCCACGTCTCGAATAAGAACCTACATGCAGTAATTGCAGGTAACAGATTTGCTACAACCTCCTACAAGCTGTAATTCTCCTCTCAGAGCCTCATCTTCACTTCAACGAAATCAACCGCAGCTTGCCATACCTTGGTACATTTGATCTCTTTGCAGAGCTCCACCTTTGCAGTGGCTATGGCTTGAAAACTTTCTTGTTGAGTTCCACCTGCACTGGCAAGAAAGTGCCTCAGGGTTTAGGGTTGCGAAGCAGAGCACAAGCAACTCACATTTTTAAGAATTGTCTTTCTCAATGCCATCACACCGTCCCGGTTGATTTGACTCTCTTTTCAGACAACTAATGTTGCAGAAACGTTTGTCTCTCACAGAGCTATGTTGTCAACCCGCTAGCTTTGCAACTCTTGAGACTACATTTTGCGATTGCCGGTCGTTACTTAGGATGTGCTAAACTGGAATTAGCTTGGCAGTCTTTCATGGTTTTGTAAGTAATTGTTGCTAGAGCTTTGGATGGACTGGATTCTTGAAGTCCTGCCAGTGTACTTTAGCTCTCATTGACAGAAGCAAGCAATACATCTGCTCTCAGGCAAGTAGTAGTGTTGATATTCTCCGAAGGCATGCACACGCTTACCAGGTGTCTTTAATGGACAAGCATGCTGTTCGGAGTCCAAATATTGATTGCAATGTCAAGATAACTGCTTCAAATGAATGTAGTGTAGCAGAGCCAAGCTGCACCAAACTCGTTGCTGATGATGCACAGGTACTGGAACAACCAGAAAATAAAGTAGTGGAAAGTTCATGCTCTCGGGGACTGCTTGAAGGCATGGTTATACATTACATTCAATCCGATCAACCGTCCAATTTGCGATTGAAAGCGCAGTCTGTTAACTGCAATCTTTTGTCTTGCAAAAGAGATCTTTCTGAAAGGATCCCTGATTCAAAAGCCTCCATCAAGCAAGCTGAGAGCTACAAGCAAGAATTTCGTGAACATCAAATCCAGCAATGGGTTTCTAAGCACAGCTACACACAGGAAACTCTCCCAAAAGATCATTGGCTTGGTGTGTATCCCCAACCAGGAGAAGGGGTCAAGAGGGCAGAAACTCCGGCCTCAGTTCAGCATGCCGGAAGCCGTCAAGGTCTGGACACGGGTGCACTAATAAGGCAGCCACAGGCTTTCATAAAAAGCTCTTCTAATGTGGATTATAGAGATTCTAGTAAGAAGCATGTCCATTTTTTTTCCAGCAATGCTCTCATTGCCGACTCAGCCACCGACTCAGCCGGTATGGATGCATCTGAAGGTTTTTTTTTTAGGAAACGAAAGTTTACAGATTTAAAAACGGGGATTCATCAGGACTTTCCAAATGAGGCAGTTGCAGCAGATCGTCAGCTCAACAGAGCAGCAGGTTTAGAACAAGCGTATTTTGAGGATGACAATAAGGCAACTGAGATCACTTTGCGCCAATGGCTTAGCAGGCCAGGTCGCACTGTTGATTGTCTTGAATCTTTAAATATATTTAAACAGATTCTCCAGTTTGTGGATCTTGCTCACATGCAAGGCGTCGTGCTTAAAAATGTTCGCCCATCTTTGTTTCTTCTTTCTTCATTGCACAGGGTGACTTTTTTAGATTCTGCAAGCTCCAGCTCTAGTTCTGGCTCATCTGAACCTTCTACAGGTGCAGCGTCTGTTCAAGCTTCTCTTCTGGCAGATACAGCATCCATGAGGGTTGACACTGCTTCCCAGACTGCAGAAAGATATTTCTTGATGATGCATACACCTCAAAGTAGGGATCTGGGGGAGCATATCCAGCCAACGGGTTTGAGAAGTACAAACGAGCATACACTTATTTCTCCAACTCAAACTAAGATTTCTGCTGACAGGATACACTTTGGCTCTGATTCTGAAGCTCGGGATTCAGGTACTGGTTGCTCTGATTGAATCTGTGCATTTTGGTGAAGAAAGCTAAGGCGTTTTTTGTTTGTAGGGAAAGCACAAATGCAAGATAATCCTAGGGACCCTGGGGAGGGTCCTTCCAATAGTAAGTCTCACCAGAAGGACATTTTCCCGGTGCGACCTACTCGGATCATGGAGGACATGTGGTATACAAGCCCTGAAGAAGTTCAAGATGGGACATACTCCTTTGCATCAGACCTTTACAGTTTAGGTGTTTTGTTCTTTGAGCTGTTCTGCCCTTTTAGCTCCTGGGAGGAACATTCAAAGATAATGGTGGATCTGCGCCATAGGATATTGCCACCCAGCCTTCTTTCTGATCAGCCCAAAGAAGCTGCTTTTTGTCTCTGGCTGCTTCATCCAGAGCCAAATTCTCGGCCCAAAACCAGAGATGTGCTACAATCAGAGATTCTTAGTGATGCTGGTGAGGCGCTTGCAGAGAGACAAGATGCTATGAATCTAGAAGAAAAAGAAGCAGAAGCAGAGCTGCTGTTGGAATTTCTTCTACGTATGCAGGAACAGAAAAGGACCATGGCATGCAAGTTGGAAGATGAGATTTCATGTCTAACTTCAGATATAATGGAAGTGAAGAGAAGACGGTCTTGTTCGCAGCGCTCCTGCACAATTCATGAAGAGCAAGATCCTTCTAGCAGGATAACACAAGAAGGAGCTTCAGGAAGAGTTCAATGGAGCTCAAAGAATTTAACATCAAGTGGCGTAGTGTTGCAGGCCGATGACACAGAACTCTTGAGTCAGGGCCTAGGTAATGAAGTTGCTGTTGATTCTGCAGAAAAGCAGAGCTCCTGGGGCTTTGATACAAAGCTTGATACAAAGCGAAAAGAGGAACAAATACGCTCAAAATCTGCTCGATTGTTGAGTAATTTTGATAAGTTTGAGGAAGCTTATTTTGCCATGCGCTGGAATGTTCAGCCTGAAGTTTTAGTAAGCCAAGCAACCCAAGAAGCCAGCAGTGGCAGTGCACCAACAGATTCTAGAAATGCTTTTGTGGAGGTGAATAAAACTTCTGAGACTGGCACTGGTGGAAAGGATTCTCTTGGATTTTTCTTTGATAGCCTTTGCAAGTACGCAAGGTATAGCAGGTTCGAGGTAAAAGCCACACTTCTTCATGGCAATTTGCTGAACACTTCCAACATGGTCTGCTCACTAAGTTTTGATCGTGATCAAGAGTACTTCGCGACTGCTGGTATATGCAGGCGAATCAGGGTGTTTGATTGTGATGCTGTATTAAATGAGAATGTAGATATACACTATCCGGTTGTTGAGATGATGGGAAATTCAAAGCTTAGTAGCATTTGCTGGAATAGCTATATCAAAAGCCACATTGCATCCAGTGATTACGATGGCGTAGTTCAGCTTTGGGATGCTAGCACTGCCCAGTCATTGATGCAATACAAGGAACACAAGAAACGTGCGTGGTCAGTTGACTTTTCTCAAGCTGATCCTACAAAGTTGGCTAGTGGCAGTGATGATGGATGTGTAAAGCTTTGGAGCATCAATCAGGAGGGAAGCATTGGAACAATCAAGACAGAGGCAAATGTGTGCTGTGTCCAATTTCCTGGAGATTCTGCACATTTGATAGCTTTTGGCTCTGCTGATTATAAAGTGTACTGCTATGACTTGCGCAGTACGAGGCTTCCATGGTGTGTCCTGATGAATCATAGCAAAGCTGTCAGTTACGTGAAGTTTCTGGATTCTGATACAATTGTGTCTGCATCTACGGATAATACCTTGAAGCTCTGGGATATATCGAGACCAAACGCAACTGGAGGTGTTTGTAAAACGCCATCACTGACTTATACTGGTCATACAAATGAAAAGAACTTCGTTGGACTCTCTGTAATGGATGGCTACATTGCGTGTGGTTCGGAAACCAATTCGGTCTTTGCCTATCACAAGTCTTTACCTATGCCAGTCGCAACCCATAAGTTTGGCTGGCTGGATCCACTTACCGGCAAGAACATGGGAGATGATACTGGAAAGTTTGTTTCAAGTGTATGCTGGAGAGCCAAGTCGCAGATTCTCGTAGCAGCAAACTCATTGGGAAATGTCAAGGTGTTGGAGATGGTATAGGAGAGTGGACATAGTAAAATACCACAAATTTGATGATTGCATACGGTGCATTGCTTTGCAGAGCTTTTTCTGTGCATCAGATTTCTATTGCATGCCACGGGCGCAAAGTACTTTATGTACTTTGTCTACTTTCATAGAGAAGAGTCATAAAGTTTTTGAAGAGTGATACATGGAAGCATGTAGTTGTTGAAAGATGATTCAGGGAAATTTACCCTAGGGTTTTTGAAGAAAACACCAGGATTTAGAGTTCCCCCACAGAGTGGTCGTGTACAAATAATTGAAGTGATCACACAGGCCGGTTTAGTTTGTAAATAAAGAGATCTGTAGAAGACACAAACTTTACTCAGAAGCGGAGTTTTTGTTTACCTATTTGTGACAAGGTGAAGCAGACGGAAGATGACTGCTTTTTGCTCTCTCAACTGAGAGGCTGAAGTTGGATCGGTATCCGTGTTTTTTTGGTTATCAGCACAGGCTGCCATCACAAGTGGTGCCTCAGTTTCACAGGAAATGCAAGAAATAGAGTTTTATAGGACATTTATTAGGACCAGCAGGAAATGGAGAAAGCGAGTTCTGTTGAGTCAGTTTTCGCCTCAAGTGAGCATGACATCTTTTGGCACTTTGGGAAGAGCCAGGGGCCATGTATGTAGGTCTGTAATGACGATGTTTATTGGCAATCATGGAAATTTATATAAGCAGGTCATGGCTAGCATGAAAAATGCAAAATTTCATGTTATTGGTGTGGCTTGCGGGTGCCCTATTCATGGCATTGTAAGAGTGCAATTTATCTTTTCATGTTATTTGTTTCAACAAAAATATTTATCTGAGGGACAGGAGCACCAACATATGTAGGAGGTTCCTTTTTCGTGCATTGCAG

**UV RESISTANCE 2 (UVR2)**:

TTTTAAGTTTATCCCTTCTTTAAAGTGTAAGCAGGAAAAGAGAGCTCATTTTCTTCTGAGCGCGGGAGGAGAGATGGCGCCAGCAGGTAAGCGCACGGGGAGCGACGGGGAGAGAGCAGCAGCAACAGGACCCGCCGCCAAGAAGCTCAAGGCAGCAAAGGAGAAAGAAGAAAAAGTAAAGAAAGAAGCAGGGGCTTACTTCTTACAACAAGACGAGGTACTAGCAAGTGCAGAGGTCAATGCAGGCAGGGTGCGCTTGCTAAAGAAGGCAACCCAAGAAGCAGGTCAAGGTGCCCCCGTTGTTTATTGGATGTCGCGTGACCAACGGTCCCGCGACAACTGGGCACTTCTCTATGCTGCTCAGCAGGCTGAAAAGCAGGGCTCTCCCCTCTGTGTTGCTTTCAATCTTGTTGATTCCTTCCTTCAAGCCCAGGCTCGCCACTTCGGGTTCATGCTCAGAGGCCTTCGCGTAGTCCATCACAACCTCTCCTCCCTTGGCATCCCTTTCTTCCTCCTGCAGGGAAAAGCAGAGGAAAACATTCCTGCATTTCTTGAAAGATGCGGTGCATCAATTCTCGTCATGGATTTTTCTCCCTTAAGAATTGGCAAATTATGGAGAGAAGGTGTTTGTAGAAGAGTCCCGTCTTCAGTTTCTGTGTTTGAGGTGGATGCCCATAATGTAGTTCCCACATGGGTTGCTTCTGACAAGCTAGAGTATAGTGCTCGGACAATTCGTGGTAAAATTCAGAAACAGTTAGCAGCGTATCTGGTCGAGTATCCATCACTCGTGGTCCCAAGTAAAACATGGACACTTGAAAAGCCCGAGGCCATTGATTGGGATTTTCTTGTTGCAGATGTCATTAAGAAAGGTGCAGAAGTGGGTGAGATCAAGTGGTGTGAACCAGGAGAAGATGCTGCATTGGAGGTGCTTTTAGGAAAGAAGAGTGGGTTCTTGACAACACGATTAAGAAACTATGCGGACGATCGCAATGATCCCTCAAAGCCTTCTGGACTCTCAGGTCTCTCCCCTTATTTGCACTACGGTCAAATATCAGCTCAGCGCTGTGCTTTGGAGGGCCGCAAGTTGAGAAGATCACATACAAAGGCTATAGATACTTTTTTGGAGGAGCTAGTCATAAGACGAGAGTTGGCAGATAATTTTTGTTTTTACCAGCCACATTATGACAGCCTTCAAGGTGCTTATGAGTGGGCTCGTAATACCCTTTTGGCTCATGCTTCTGATAAGAGGGAGCATATTTACACGCAAGAGGAGTTGGAGAAGGGGAAAACTCACGATGAACTATGGAATGCTTCACAATTTGAAATGGTATATCTGGGGAAAATGCACGGCTTTATGAGGATGTACTGGGCAAAGAAGATCCTTGAATGGACAACTGGGCCAGAAGAAGCCCTTGCAGTGGCGATTTATCTGAATGATAAGTATGAACTTGATGGAAGGGATCCAAATGGTTATGTTGGTTGCATGTGGTCCATCTGCGGCATACACGATCAGGGATGGCAGGAGCGTCCAGTATTTGGGAAGATTCGGTACATGAACTACAATGGGTGTAAGAGAAAGTTCAATGTGGATGGATACATTCAATATGTGAAGCGCGCGGTTGCTGAGTTGAAGAAGGCAGCAAAAGATCCTGCTAAGATTCCTCCATCATCAACACCATCAAAGAAGAATGTGCTGTAGATTATGCATTAGTACTTAATTTGATGAAGACTAGAGGAGAAATTTGCTCATTATATGTATAAGATCTATTGCAGGGACTGTTGTACTTAAGATTGTAAGTGAGCCAGTAGTCTCAAGGGTTTGTAATGCACATTCATTACGGTTCGATGTCATCAAGACATTTGATACTCGACCAAATTTGTAAATGCACTTTTGACCAATTATTAAGGAGTTAATCTTAGTCAAAAAAAAA

**UV RESISTANCE 3 (UVR3)**:

AAATGCCTACGTGGACAGCATTTAAACTACGCTCCCGCTCTGCAGGCGGCTGTGGCACTTGGTTGATTGCCAGCCATGGAGATCAACTCTCTCATGTGGTTTCGCAAGGGTTTGCGTTTGCACGATAATCCTGCTCTGCACAAAGCCTGCGAAAACGCCAAGCACGTGTACCCAGTCTTTGTTCTAGACCCTTGGTTCCTAGCCCCCGACCCTTCAGCTCCATCGCCCGGATCCACTCGTGTTGGCGTTAACCGCATTCGTTTCCTTCTTCAGAGCTTGAGCGACCTTGATGCTCGTCTTCAAGACAGAGGATCTCGCCTCCTCTTCTTTAAAGGAGATCCCGTTTCCATCATTCCTGATCTCCTCAAAAAGTGGAATATCAAGAGGCTTTGCTTTGAGTTTGATACAGAGCCTTATGCACAGACGAGGGATAAAAGTGTCAAGGAAGGTGCGACAGCAGCTGCAGTAGATGTCTTTTGCCCCATCAGTAACACAATCTTCAACCCTGAAGAAACTATCAAAAAGAATGGTGGGTCACCACCTTTAACCTATCAATCCTTTTGCAAAATCGTTGGCAAAGCATCAGCTCCCATCGGAGATGGCCCCTCTCTCATACCCCTGGCCAAAGATTTCAATGGGGCCAAGATAATCTCTCTTCCATCCCTTCAAGATCTTGGCTATGATGACTTGGAGCAGGAAGAGTTTACGCCTTTTCCTGGAGGAGAGACCGAAGCTTTGAAACGACTAGATGCATCATTACTGGACAAGAAATGGGTGGCTGAGTTTGAGAAGCCCAAGGGGGATCCAACAGCTTTCATCAAGCCAGCCACCACGGTCCTCTCACCCTATTTGAAGTTTGGATGTTTGTCAGCACGCCTTTTCTATGCGAGGTTGAAAGAGGTTTATTCTCAATCCAAATCTTATTCAAAGCCCCCGGTGTCTTTGGAAGGCCAGTTATTGTGGCGAGAATTCTTTTACACTGCTGGATTTGGCACTACCAAGTTTGATTGCATGGTCGGCAACCCTATTTGCAAACAGATTCCTTGGAAAGACGATGATCACCTTTTGGCTCTGTGGCGCGAAGGTAGAACAGGGTTCCCTTGGATTGATGCTGCCATGTTACAGCTGAGGAAGTGGGGTTGGATGCATCACCTTGCTCGCCATGCAGTTGCTTGTTTCCTCACTAGAGGGGATTTGGTGTGTTTCAATTGCTGTCGCTATAAATCTTCATAAGTTGCCATAAAATCCTGATAAAAAAAAGGGCAGATTGATTTACATCTAGCAGCATTTATGCATGGACACTGGACAGAAGTTTCGCAAGCTGATTCTTGTAGTCTTCTCAGTTTGTATACTGGGAGAAAGGTCGTGATGTGTTTGACAGATTACTTATAGATGCTGACTGGGCCATTAATAATGGGAATTGGATGTGGCTTTCGGCGTCGTCTTTCTTCTATCAGTATCATCGCATTTATTCGCCCATAACATTTGGAAAAAAGTATGATCCTGATGGGAAATACATTCGACATTTCCTCCCTGTATTAAAAGACATGCCCAAAGAGTACATTTATGAGCCTTGGACTGCTCCAGCAGAGGTCCAAAGAAAGGCAGGCTGTGTAATTGGCAAAGATTATCCCAAACCTGTGGTGGATCATAGCATTGCAAACAAGCAGTGTCGAGACATGATGGGCGCTGCATATGCCTTGAACAAGGGGACTGAGGGGCAACCAACCCAAGGACAAGTAGCTTCCTTGCAGGAAAAGCTGCAAGTGGAAGTGTTAAGCAAAAATGTAACTGTTGCAAGCCCAAGCAAAAGAAAGAGCCGAAAGGGAAGTAACACGATCGACAGAAGTGGCAAGCGGCAAAGAAGGATAACAGAGTACAAATGAGAACTTGGACTTGCATTGTCTTTGTAGACTCGCAATTTCCTTGTCCTACATCATGCCTCAATACAGGGCATTGATTACCCATACTCCTGACAGTAAGGGCATACACACAGATTGTGAGTGCATTTGCTATTCATGGGAGCTTGCAGAAGTGGAATATTGCTGAAGGTGAGAAAAGTTTTTGTAACCAAACAAAATTTGTAGTAACAGGTAGGTCAGTATCTGTACATTTGTACTGCTTTGCAATATAAGATAGTTGAGGGATTGTAAAGGGACTGTCAATTGTTTGCATGGGTCAGTTGCTCTTGTGGATCATCTCTTTCTACAGATGGTCAGCTACCCAATTTGTCCAGTCTTTGAAAAAGACAGCTAAGACATCTTGTGAGGCCAAAAGATTCCTCAAAACTCTTGTTGGTCCTTGTAACACATGTTATCGACAATATCACTATA

**MUTS HOMOLOG 6 (MSH6)**:

GCCAAGCTTGCAGCACGAGAGAAATGGCGCCTACCAACAGCAAGCTCGCCAAAACTCCTCAGCAGCAGCGCTCCATCTCTTCCTTTTTCACCCCAGAACCCGCCTCCGCCAAGCGTAAGAAGCCCCTCCTCCTCGTTTCTTCCTCTTCTGAGAAACCACGCCTCTCTCACCCAGAACAAGCTCGCCATAGCCACGAATCCCCTGCTTCTCCTGCATCAAAAACACCCCTTTCAGACCCCCTTTCTCGCACGCAGAGCTCAAAACAAGCTCGCCATAGCCAGGAATTATCTGCCTCTCCCGCCGAAGAGCTCATCAATGCACAGCTTTCCAACCCTCATTCTGTCATTGGTAAACGACTTAGAGTCTATTGGCCCATCGATGATACCTGGTATGAGGGTTCTGTGAAGGAATACATCTCTCCGAATCATCTGATCGAGTATGACGATGGCGATGAAGAAGCTCTATGCCTGGAGAATGAAAAGGTCGAATGGGTTTCTTGTGAGAAACCTGCGCACAGAAGAAAGCTGAGAAAGCTCAGCTTTTTGAAAGAATCCTCAAATGTTCAGGCAGAGGATTGTCTGCAGGGAAATAGCTCGAAGCGGACTCGAGGAAATGATGCAGGTATTTGCAATGTGGAAGAAAATGGAACCAACAAAAATAACAGTGAGGAAGATATGGATAATGACGATGATGATGACGATGATGATTATGATGCCAAATCTGCTAGGAATGCTGAAGACGAGTCTGATGAGTTGGATTTCAATGAGGATGAAGGTGCAAAGGCCAAAAAAACAAAAAAGACTGTGAAACATCAAGGCAGAAAGAGCACATCGAGGTGTACCTCTGCAAAGGCAAATGCACAAAATCAATCGACGCCTGGGTGCAAAGGGTCAAAGAATGTTCTTGATGAATCTACTGTCACGGAAGCAACACATAGTGTACATGAGCAACCAAATGGATCCATTGGAAATATTCATGTTCATCTCTCTGGAGAAGCAGCAGAGAGATTTGGGCTTAGAGCTGCTGAAAAGTTCAAGTTCCTTGATAAGGATCGCAAAGATGCATCTGGTAATCGGCCTGGTGACCCAGTTTACAATCCTCGCACACTTTTTTTGCCAGCAGAATTTACAAAAGGACTATCAGCAGGCCAGCGGCAATGGTGGGAATTCAAGTCAAAGCACATGGACAAAGTTATTCTTTTTAAGATGGGGAAGTTCTATGAGATGTTTGAGATGGATGCCCATGTTGGAGCTCAATGTCTCGACTTACAATATATGAAGGGTCATCAACCACACTGTGGGTTTCCAGAGAAGAATTTTGAAGAAAATGCAGGCAAATTGGCAAAAAGGGGCTATCGTGTTCTAGTTGTGGAGCAGGTTGAGACTCCTGGGCAGTTGGAGGAGCGCAACATGAAGTCAGGCACAAAAGATAAGGTTGTCAAGCGTGAAATATGTGGAATTATTAGCAAAGGAATTATAACAGAAGGAGATATGCTTACATGTTCTCCAGATGCGGCATACCTCATGGCAACATTTGAGATATCTAATGAGGAGAGCTTGAATGGTTCTTCTATCATAGGATTGTGCTTTTTAGATGCTGCAACTAGCCGCTTTCTTGTGGGCCAATGTTCAGATGACTTGGCAAGGTCACGGCTAAGGTCTATCCTCTCGGAACTGAGGCCAGTTGAGCTTGTCAAGCCAGCTGGTTTGCTTACCTTGGCAACAGAGAAGGTCTTGAAAGAGGAGACAAGAAATCCTTTGGTCATAAGTTTGACCCCTGGGTTGGAGTTCTGGGATGCAGAGAAAACAATGAATGAAATTGTGACGTTTTTTGTTGAGAAATCAGGAAACAACGGTGGAAATGTTGATGAAAAGCCAGACTGGGAAGACGGACTGCAAACTTTGCCAGAGTTCTTGCAGAAGCTTGCTCAAACCGGTGCTAAGGCAAAAGCTGCAATTAGCGCTTTAGGAGGCTGCGTCTCCTATTTGAGACAATCTTTGTTAGACCGCCAGTTGTTAGGCTCTGGCAGATTTGAACTACTTCCGGGTTCGGATTCCTTTCTGGAGCCACCATTCAAACATAGTTCAACCTTTGACAGTGAAAAAAAGGAGGAAAATACTTTGTTGCCTAACATCTTTGCAGAAGACCCAGAGCCTTGTATGAAACTTGATAGCTCCGCTTTAGAAAACCTTGATATTTTGGAGAGTAGCCATGATGGAAGCACCCGAGGGACACTTTTAGCGCAGTTAGATCATTGTGTGACAGCCTTTGGACGTCGATTGCTGAGGCAGTGGATTGTTCGACCTTTAATATGTGTTGATTCAATTGTAGACCGGCAAAATGCCATTGAGGACATTAAGGGTGTAGCGGCTGATGCAGCATCCAAGTTTAGAAAAGATCTTTCGAAACTCCCTGATATGGAACGATTGGTTGTCCGGCTGAAGTCTTACAATGAAAAAACAGGAAGAAATGCTCATGCAGTTGTGCTGTACGAGGATGCGGCCAGAAAATTGTTAAGAGAGTTTCTAGGTGCTTTGCGTGGATTTCAGATAGTCAATGAAGTCTTGTCATCGTTCTCTAGGGTCTCAAGCAAGTTTCGGTCAAGGAGGTTACAGTCATTGTTTACATTGGGAGAGGGTGTGCAAGATCTGGCTTACGCTCTCAAGTACTTTGAAGATGCATTTGATTGGCTTGATGCAGAAAAAAGTGGCCGCATTATACCACATGAAGGCGTGGATGCAGAGTTCGATTTAAGGTCTGAAAATGTTGCAACAGTTGAAGCGAAACTAGAAGCATACTTGAAAGAGCAAAAGGAGTACTTCAAAAATCAATGTATTGCATTTGTGACGGTAGGCAAAGATCGTTTTCAGCTAGAGATTCCTGACTCTTTGCAGAAGAAGGTGCCACCTGAATACAATGTGGAATCACAGAAGAAGGGTTTTTGGCGTTATTCAACTCCAGTTGTGAAACAGCTACTCGCAGACATTGTTCTAGAAGAACAACAAAAAGAGGCAGCACTGAAAAATATTTTGCAAGGTTTACTTGCTCGATTCGGCGAAAAGTACTCTTTATGGTTGTTTTTAGTGCGATTAATTGCAGAATTGGATGCATTGATAAGCCTTATGTTGGCCAGCAACACTATTGTGGGAGCTATGTGCAAGCCTGTAATAAGCAACATCAACAACTCTTTAGATAACTCAGTACCATTTCTACTTGCCAAGGGATTGAGACATCCTATTTTAGAGCACAGCCTTGATGGGGGGAGGTGTTTTGTACCAAATGATATCAACCTTGGGGGTAAACAACATGCGCCTTTCATGCTTCTTACTGGCCCAAACATGGGAGGAAAGTCTACACTTCTTCGTCAGGTCTGCTTGGCAATCATTCTAGCTCAGGTAGGTGCTTATGTCCCAGCAGAGGACTTCCATTTTTCCCCTATTGATCGTGTTTTTGTCCGGATGGGTGCAAAAGATCACATCATGGCTGCTCAAAGCACATTTCTGGTAGAGCTTTTGGAGACTTCTTCTATGCTGACTTCTGCGACGCGACACTCATTAGTAGCATTAGATGAATTGGGAAGAGGCACTGCAACATCAGATGGCCAAGCCATAGCGCACGCTGTTCTTGAACATCTTGCGCACAAGGTGGGCTGTCTAGGGATGTTCTCGACTCACTACCACCGCCTTGCAAATGATCACAATAACGATACAACGGTCGCATTGTATCATATGGCGTGTAAAGTTGAAGATGACAATGCTATTGAGAAAGTCACTTTCCTATATAAGCTTGCAAGTGGTGCATGCCCAAAAAGCTATGGTGTGAATGTAGCACGCATTGCAGGTATGCCTGAGTCAGTTCTGAGAAGAGCAGCAGTTCGATCTGCAGAGATGGAGGGGATAGAAAGTACACTAGCCATGAAAGGTTATGAAGAGAAGCTCTTTGTGGAGGTACTGCAGGTCGCAACAAAGGCAGATTGTGCAGATTTGGGTGTCATGACCAACTTACGTGACCGATGGAGATCGTTGCATGCAACTCAGATTCCTCTTTTCCAGAAATGCTGATTTCTGAGTGGCTTTTCTGTCTTCCCCAGCCTCTTAAACCACACATTTTGGATATGGAGTCCTTCAAACATCTGGAAGTTTTCTCAGGCCTGGAAACCATGGCAGCCTAACTGGGTCAATCACAGTCTAGATAAGAGATAGTCCTCAGGGTGCCCTACAAATCATGATGCGAATTCGAGAACTCATATCGTCACCATTGCCAGCTCTCATCACTTCATTAATGGCAGGGATTATTTGGCCTCGGTAAGCCCTACAGCATGTCTGCTTTTATGATCGCCATGCCAGCAGGCCCGTTTCACCCTTATTGCCTTTTCTAAGGCATTTCTCTGCCCATTCTAAAGGGTACCGTGAAAGTAGATATAAGTTATCCGGCTCTAACGGCCTGGCAGCCTGCTAGTTTGTCCAGTTGATTTTGAGACATGGACTAGCTTATCTAACGAGTCCCAACCATTTTTTGATGGCAGTTTGGGCTGGTCCTTAGTGTTAAATATTTTGTGCAGTAAACAGGTCTTATACTTGCAGCGAAAACGTACTTTCATATGGATGAAAGCAGATGTGTTTTACTTTTCTTGCTACTCTTGAGTACACCTTTTTGTATAACCTCACTTTATGTGGCCTTTTGGGGCTCCTTGATTCTGGACATGTGAGCAAGGTTGGATTTTGGTATGCATGTAAATTTTCTTCATTAACCTTTACAGGACAACCACACTTGTAACAGTTTTTAAAACTGGCGAGTTCCGATCACAAGCAATGTCAGAAAAAAATCGACTTTTAGTGGC

**HISTONE ACETYLTRANSFERASE OF THE MYST FAMILY 1 (HAM1)**:

TTTTTTTAAACAAAGACACGGAGAGTTATTCATGTTCTATTATCATGTGTTAAATGTATATACAGGCTTTATAACCATTTGAGATTGAACTGTGGCTCCCTCACCACCTTTATCTACCTTTTCTGGCCGAGTGTGTTACAAGTGCAGAAATCCTATGTCGACATCAAGTTTATGACGGACATATCAACATCGAGTTTATGACGAACCAATTCCAAGTTAAGATTACAAAATTAGGCTAGCATGGAGATATTTACTAACACGTACAAATTCTTTTAATTACACGTAAGTATTTCCTCTCCCACGTCAATTATGTCAAAAATGCCCTTCAATCTCAATATATATCTACAAGCACTATAACCTAAAGCTTTTCTACATACATTCATTGCTCTTTGAATGGTGTCCAAATCAATTTCGTGGCATCCACCTCAAGTCCCCCTTTGCCAGCAGCCTTCAAATGACGCTCCAAGAGCTTAGGATCTGCACATATGACATGTTGCCCTTTTCTATATTGAATCAGTTCAAGAGACTGCAGGGTGCTGATGACATCCTCTGTCTTGATAGCAGTCATATCACTTAGCTCCTTAATAGAGAGGTTTCCCCTATGCTTTTTTATAATATCCAAGAGAACACGAGTCCAATAGCCCCTATAACTAAGCAGCCCCAAGTCCGATAAAGGCCTTTCCGGTGTTCCCACTTTGAGCTCCTTCTTTGACAATTCATATGAAAAAGAAATCAAGAACTTCCCATAGCCCTTTCTCTGATAAGGTGGTAGCGTCAGGATACATGCAAGATTATAGCCTTCTTCAGATTGTTTTTCCTTGGAAAAATATCCAACCATGTGGCAACCGCGATCATCACATTCACACAAAATGTAAAATAAGAAAAAATCCACGTCATAATACAGTGTTTTGTGATCCAAAAAGAGCTTGGCCAGGTAGCACAGATTCTGGCCATAAATTTTGTTCTTCCTTCCATCAACCTCAAACATGGATAAAGAGCCACTACGATAGATTTCATCTCCAGGGGGGTGCTTGTAAGTGCACTTCCTCATGTGCCTTGCGAGCTGCTCCTTGCGTTTCAAAAAATTAAGGCAGAACTCACAAAAGAACAGCTTGTCACAGTTACTGTACTCTGGTGGGAATGGAGAGAAATACCAAGTATCAATCTCGTACTTTCCCAGCTCTATTTTGCTTATATTCTTGACTTTAGTAAATTCCTCATGCTCCCGCAAACTTGCAGCATCAAAATCTTCGTGCCCCTCTTCAACATGGGTTTCATCGATCTTCCGTTTCTGATGCCGAGTCATTTTCAGACTACCAGCCTTGTCATCAGCCTTCTCTTCCCCCTCCTGTTCAAGAGATTCCAACTCCAATTGATCAAGTTTCACCCACTCGTCGAGCCTCCTATTAAATTCTGTGTAATGGACATAGTATTGGTAGTCATCAGGTCCAGTGGGGAGCTTGCGCCTTTCGATAACACGAACAGGGTGATACTTGTCATCGCGCCAACGGCAGAGAACCCTAGCCCCGACTTCGAGAGGGAGGCCCATGAACTTGCGGCGTTTGAGGTGGGAGTCCAACGCAAGCTCGGCCATGGAGGGAGGCTCCAAATCCATGATGCCATTCTTTCTGATCTCCTCGCCGTTACCGAGAAAATGTATGTGCTTCTCCATGCCCAGGCTTCCATTGCTGCGCGAGAAGACCTCCACGCTTCCCATTGTCTGAGGAAACCCTAACTTCGCTTGCTCTGTGTGTCCCGAGTTGCTATAGTAAAAATATTGACAATAGAGAG

**HOMOLOG OF REVERSIONLESS 1 (REV1):**

GATACAATTGTGATGCACAAAGTGCGAAGTAAACAAACACTAAAATACATGTGTTCGTTAAGATAGCATTATTCCTAGGCAACTAAGGTACTATCTTAATTCAAGGATCACAATGGCCATATCCCTTGACACTAATGAACTTTCCCTTTTTGTTGTATTATTACAGACAATTTCAGACCAATTTACATTAGTGTACTTGTACAAATTTGTATACAAAAACAGTTTATTGCACTAATAGCTCCTTTACAAGCCCCCTTGCTGTGCTTGTTCGGATATTAGTGCATGAATATTTAAGCTCTAATATACAAGGTCCCTCCATAGACCTCTCCAGTTGCTGCCTGGATAAACGGAAGAATAGATTCATAAATTCTTTCCCAAAACTGGTCTTCACCTGCTAGACGTTTTACAACCCGAACAATCATGAAGACCTCTTCCAGGTTATATGACAGTTTTGTTGTAAGGTACTGTTTAAGCAATTCCAAGCAGCATCTGAATGCTATCGAATCTGCCAGATCTTCATGTAGGTCCTCATGCACAATGAGCTTGAAGGAAAACATTGCATGAAGCAGAACAGACGAAAAATTTTCAGGACACCTTAAATCCAAGAAAGCTGTACAGAGAAGGCGAAGTATCTCCGCCCTGCCTCCATGCAATCTCTTAAATAGGAAGACCCATTTGGGAGGCGCACCACACCAAAGGCCATCAAGTTGGAGAGAGCTAGCTGACAGCTCCGACATCTCTTCCAGATGTCTTTCAGGCATTGTCAACATTCCCGTATCATGTTTATGTCTTTGATTGTCAGGTCGATGTACAGGGAGAGATCTCAAAATTTCCTCCCTTAATTGTGAGGGCAGCTCCTCCAAAGTGGCCATATCAACCTGTGATAAAGACAATGGCACAGATGATTCTGGATCAGCTCGTGTCCCTCCTGCAGAGGAAGGCTCGGCTCTTTCTTCCATGCGGCTGCGTTCCATTCTGGAGATTGGTAAGACATCATCAAAGGCTGCAGAGCACTTATCCCTCTCTCTTTGAAAGTTACTAGCATCACCTCCATCAGCAGCAGATACAAGAGGCTCATGTACTGGAGTAGATGACCTGCTCATCACAGATGGCATACCAATTCTGGCTTCAGGCTGCTGGGACAGTGATAAATGTTGAGCAACTGCACCATTGTAGGCTTCATCCAGTTCTGCTAAAATATCTGGAGGCAAGGATACCAGGATGGAATGGTCCAACTCGGAAAATTGAGGTAAATTATGCAAAGAACTGAGATTTTGGTTCTCTTTTTCCTTTGGTTGCTTAGGTCCAGAAAAAGGGCCTTTCCGAACTTTTCCTAAGCTTAGCATCCCCTTATTTTCCTTTGGTTGGCTTCTGTCTATCCCCCAAGGGGCGACTACGTGTCCATCCTCCTTCGTAAGATGGGGTGGCTTCTCTCTCACAGTAAATGATTTCTCATTTGATACATCAGACAAGCTATCTGCTTTGGACATGCCTTTACCAGGACTCTCTTTGGACACAAACTGCAAACATGACTCCAGTGCTTTCTGGTGATGCTGACCTTCATCACTAGAAACAGTTAGGTTTGATTCCAATTTGGTGACTTGAAGGCCCATTCCTCGGATCTCTCGTACATCAAGGTGGAAAGAGGCAAAAATTTGTCTGCTGACTCGAAGAAGAACATCAACACTATTGGTAGCACAGCCCAACGTGACTGATTTGCTTAGGTTGTCACAACTTCCGCACCCCATAAATTTGTGAGGTTCTCCAGCACCTTCCTTCCTCTTTTTTATCTTGAGTGTAAATGTCCGGCCAGTAACAAGAGCGTTTTGCAAGCGCGTTGAGACCTCTTCACATAAATTGTTTAAAAAATGTTGTGCATCAGAAGCTGAGTCAAAGCGAACACCCCAATTGACTTCAGCACCTATTGACTTGCGTTCTTGATAGTTTTGAACTTGGCGATGGTCGATTCCACGAGCATAGCACCACAGCATATCACCAGTTTTGAGACCAAAACCCCTTTGCAAACCTTCCTTGGAAACACCTCGAAGTTGCATACAATTGTAAATGGATTTGCTATGTAATTTTTCTTGCAGTGACCACCCAACCCCAGGCAAATCCTCAACTGACAAGCCTGAAAGAAACTCTTCAACATGTTCAGAATCAATTTTATACAAGCCGTCCGGTTTAGCCTTGCGGGTTGCCATTCGTGCTAGCAACAAATTTCCAGCTATGCCAACACTTGCAGTACATCTTGTCGAAATGACAATTTCCTTCCGGATATTTGAAGCCAAAACTTCAGGATCTTCAGAACCTGTGACATCCAAATATGCTTCATCACAACTAATTGCTTGCACTTTTTTGCAATACTTGTGAAGAACGTCATAAAACTGATCTGCCACCTGCCCATATCCTTCAAAATCATAAGGAACAACCTCCAGTTCAGGGCATAGAGCTTTTGCATCCCGGATAAACATGCCAGACTTGACACCAAATGCTCGGGCAGCATAATTGGATGATGAAATCTCGGCTGTGCCCTGCGTATTCTCGGAATGGCACACAGCAACAGGCTTTCCAACCAAATCAGGTCTATTCCTCACTACTACAGAAACAAAAAAGCAATCCATGTCAACATGCATGACTATCCGATCCCCATTATCGGAAGAAACAGAAAAAGGATTTGGCTTCTCATGCTGTTGATCTTTCTCTAGCTTGCTACCATGAAAACGATTCTGATAACGTGTTCTCCAAGTGCCGATGAAGTGAAGTCTTGAATTCTTGAAGTAGTTTTGTACAAAGTTAGCATCACCCAGAGTGCAATGCGATCGGACACTTTCTGAAAAGCTCCCAAAATTTCTTTCTTCCGATGCACTTGGACCTAAGTCAAGCCCACGGTCTTTCTCTGAATGTAAGTCTGACTCAAGGTCTCGTTTCAATGGATATTGGCATGACATATGCGTGTGGCCCTTTTCAAGAACTTCATAATTATCAGCTGTGGTCACTTGCTTCTCAAGTGGCATTTGTTCTTGTAAATATGTGATAGGTTGTGGCAATGGTAGAATGCCTGCATGAGACAAGTGCTTGTTTATATCCTCTTTTGTGCAATCCAAAAAAGCTGACAGAGTTTGCTGATTGGGATGATTGCGTGCAATCCGCTCGTGAGAATATAATGAAATAGGTAGAACTTTCCCACAAGATATAGAATCCACAACCCAAGAAGGCTTCACGATTGGCAAGCCTCTACTAAAAGACCTACAAGCACAGATGAATCGATTGAGCTGGTGACTCTAAACAATAAACAAATAAAGAAAACAACCTATGATCTCTTATTTTGCTGTCTGGAAGCGTGCTGCAAATGATGTGGGTGACCATATCCCGAGAAAAATAATTCTCAAACATTCCCCCATATTTAATCATGAGGTTGCGCAGTTCCTGATGAGATGGTGTAGTGAAGCCATCCACAAAAATGGAAATGCCAACAAAAAGCTCTGTTAGATCTCTACATGGAGCTGCTGCATCAAGAACACGGATGTTGGTAAGAAGCTCTCCATTGCCGGATGATGAACTGAGATCTGTGGGTGCCAAAGTACTTGAGTCCTTGTTAGATGCCTCATCTTGGAATTGTAAGCTTAGCTTTCTTTTCTTTTCAGTCATGTATTTTCCAAAGTCACCAAAGGCGCTGTCGTGTTGAATTTTGTATTGTTTAAGAGAAGGAGAGTTGGAGCCCCACAGCATGCCTGTTTGTTTCTGCAACTGCCCATCCATCTGCTTCCTCTTCATGCTGCTACTTTCAACAAGTTCTTTGTTGGGCACAGCCATCCGGCCTTTCAGATTGGTAGGAAGCACTAAACCCTTAAATCTCAGATTTTGCTACGCCTCACACAGAATTCTGTACGTACAATCTCATACAATAG

**WD REPEAT-CONTAINING PROTEIN CSA-1 (CSA1):**

CTCTCTCTCTCTCTCGCTCTCTGTCTTCTGGATGGCTCAGGAAAAATAAGGAAAGAGGGGGCGCAGGAGGATGTGGCAAATGGTTGGCAAGAGAGAGATTGGTGGTGTTGCGCCTCATCGTTTTTCCGCTCTCTTGCAAAGGAAACGTCTCCATAGCATTGAACTCGCCAGCCACAAAGAAGTTGTCAGTGCGCACAAGAGTGGAGTCAACAGCCTTCAGATTGATGCCACGGATGCAAGATATTTACTTTCTGGAGCATCGGACGGTTCAGTGGCTGTCTACGATATTGAACAAGCAACAGCTTTTGAACATAATACACTAATTTGGAAGCATGAAGCCCTCTTTCTGGTCGACAAAACAGTGCAGCATGGCCACTTGTATTCTGTTTCGTGCGTGCATTGGTATCCCATTGACACTGGCATCTTTGTCACAGGGTCTTTTGATCACTTTGTGAATGTTTGGGATACAAACACAACACAGGTTGAGCTGCAATTTAATATGTCAAAGAAAGTCTATGCTATTTCCATGTCAAACCTTGCAACAACACATATGCTGATTGCTACTGGAACGGAGGATACTCGAGTCAGATTATGTGATGTAACATCAGGCGCATTTACACACACACTTACAGGCCACAGAGACAGTGTATGGGCTGTGCAGTGGTCGGCCAAAAATGAATGGGTGCTTGTAACTGGTGGTTGTGATGGGGCCATTCGCTTCTGGGATATTCGAAGGGCAGGATGCTACCAAGTTTTGGATCAACATTGCTCACAGATTGGACGGCGTCCTCCTGTATTAAAGAGTGGTACAGGAAATCATAGAAGAGAAAACAATTCGATTGCTAGTTCTTCTAGCAAATCAACTTCAACGAACGAGAGAGGTCAAAAGCGGCATCTTGGTACTTCAGCAATGGACTCTAGAACTGCACATTACGGATCGGTAACAGGACTTCAAATGACCTTTGATGGACTCTATTTGATTAGTGCAGGTCTTTGCTTTTAGGTACAGACTCAAGGCTTAGATTGTGGGATTTAGAGTCTAGCTGCAACACACTTTTAAACTACCCGTCAACACGAATTCGTGGAAATAAAGGCACACAGATGACTCTTTCACCAGACTCCTCGCTTCTTTACACACCCTCTGCAAATGCCATTCTGGTCTACGATGTTTGGTCAGGCCAATCTTACTCTTCTTTGCGGGGCCATTATGACCTAGTGAACTGCTGCACTTTTCACCCACAAGGACAGGAACTCTACAGTGGGAGTAATGACCGGAAAATTTTGGTTTGGGCCCCACCCTGTCCCGCTGATGCCGAAAAGCTGGGAAGGGAAGCTCAAGAGCAAACTGTACAGGCGGACGAAGATGCATGGAGTGACTAAGAGGGAACAGTGCAAGGTGCATCGATACCCTCGCAGCATATTGAGACCTCGGCCCTGCATGTTGGCTGGATGATTGATTTCCAAATTCAATGCTCCTGGCAAAGATCATTCAAGTCAATCTTGTTTACACTCCATTTACAAGAGTTTAAAAGTTTTCTACCTTCAAAGCCAAGATAGATTCAAGGGCCCAGATTAGCTGACAGCAAATGGAAGGCATGTATACATTTGCCGCCGTCACAAAGAGAAAGTTGGCATAGCTTACAAGTGTGCACCAAGTGGTTATTTTGCATCCTTCCAGACAAAGATGGTTTGATGAATTCTGCAATAGGGTGGAGAGCAGTCCATTGTAGCTTATTCACAAGAGTATAATAGTTGTAGTCTTTCTTGACATGTTATCAATTGCTTGATTCTTTTTAAGTTTGATAAATTCTTCAATAGGGTGGAGAGGTGCTCTAGGCGCTCCATTGTAGCTTATTCACAATAATTGTAGCCTTTTTATCAA

**UV RESISTANCE LOCUS 8 (UVR8)**:

GCTCTCTCTCTCTCTCTCTCAATCAACTCATTCTCCTGCAGCTTACAGCTTGCCATTCCTCTTTTATGCCAGTGAACGGCTGCTCTCTTCTCTACAGCCCCTCCTCATGCTGCAGCCCTTGAAGCCCTATTCTCTCTCATTGAGAGGATATCAAGCGGCTGCTGTGGTTAGAGAGTTTTCGCGTGAGCCTCTCTGTTACTGGAGAGGCTAGTTACATTGAGAATGCAGACGTTCGTTCTATCAGAGAAAAGGATCTGGGTTTTGCATTAGATTTTCTCATACATACGCACAGGCCCATCAAGCTCAAGCAGGCTGCACAGGAATGGACGAGGCGTCTTCCAGGCTGAGGCAAAGAAGAGTGGAGCTCGTCTCAGCCGGTGCCAGTCATACTGTTGCGCTCCTTTCTGGAGACATTGTATGCTCTTGGGGTCGAGGGGAGGATGGACAACTAGGGCATGGAGATGCAGATGAGAGACATTTTCCAACAATTATTAGCTCTTTGGAGGAATGTGAAATTTCTTCTATAACATGTGGGGCTGATCACACAATCGCACGATCAGATTCTAGAGCAACAGTATACAGCTGGGGGTGGGGTGATTTTGGACGCTTGGGACATGGGAACTCGAGTGATCTTTTTATTCCTCAGCCTATTAAGGGACTTAAAGGGATTGAAATCAAGCAGATTGCTTGTGGTGACAGCCACTGCCTTGCTGTGACGATGGATGGTGAAGTGAAAAGTTGGGGTAGAAATCAAAATGGGCAACTTGGGCTTGGACACACTGAAGATTGTCTCATCCCCTGTACCATCCAAGCTTTCCAGGGAATATCTGTGAAGATGATTGCCGCTGGAGCTGAACACACTGCTGCAGTGACAGAGAGTGGCAAGCTGTATGGTTGGGGATGGGGCCGGTACGGAAACTTGGGATTGGGTGATCGGAATGATCGCCTAGTTCCCGGAGAAGTTCCATCCATTCCAGGCGAGCAAATAAGTCTTGTTGCTTGTGGCTGGAGGCATACAATAGCTGTTTCAGACTCTGGAACACTCTACACGTATGGATGGAGCAAATATGGGCAACTTGGGCATGGCGACCACGAAGACCATCTTGTACCTCATCAAGTGGCTGTTCTAAAAGCAAATCGCATTGTTCAGATTTCTGGAGGCTGGAGACATACAATGGCCCTTGATGACCAAGGAAAGTTGTACGGATGGGGTTGGAACAAGTTTGGACAGGTTGGAGTGGGAAACAATAATGATCATAACAGTCCTCAGTTGGTGAAAACCCCAGTGGAAGAGCGTGTCAATCATGTGTCATGCGGTTGGAGGCATACTGTGGCCATTTCAGCTAAATATAATGTGTTCTCATGGGGTAGAGGAACGAGTGGCCAGCTAGGTCACAAAGATGCTGTTGACAGGAATTTGCCAAAGATGATAGGTTCCTTGAGCGCAGATGGGGAAGCGTGTGCAGGGGTTGAAAGCTCAACTTCTGTCGCAAATGCAGCTGCTGCAAATTGGATTTCGCCTTCTGAAAGATATGCAGTCGTTCCAGATGAAACTGTAAGAACGCTTGCCAGTCGTCTTCACCAAATAGCCAGTGTTGGTGGCCAACGTCTGGAGGGCACCAGAGAAGATGCGAGTGTGCCAGATACTGATTTTAAACGTGTAAGGACGGAGTAGATCATACTCCAAAATATGAAGGGATTCATGATTTGTGCAGACTCTTGAGGTGGTGTAGGAGTTACGGCAGGAGTCAACCGATTAATCGTGGCGAAGAGGGTCGAGCTTATATAAATGGCTCACTGCTCTCTGCTATTTATGTACTTCAAACGGATCCAAATCTGTACAGGCAAACCGCTCAATTGTTTCTGAAATCTGCTTGAGATCCAATGGTACTAGTTTGTTTGGTAAAAAGATCTCACATTTATTATACAGTGTTTCATGGAAATCATTTGTAAGTGCAGTATGGTGCTTTGTGGCTATTTGGTGGCTCTGCTCGAAACACTGGGGATTTTACATTGCTATTTGGAAAATACGCTGGGCCTTTTGAAGCATAAAAAACCTGGCCATGTTCTTTACTATTTAAAAAGTGTCTATATGTTAAGCTTACGGTTAGAGCTGTGGTGATAAAGCAGCTTAACAGTGATGCTGACATATATTTGCCACCTGTTGAATTCTTTCCTCCTGCTGCACATTGACAGTTATTGCAACGCAGTTGATTTTGTTGGAGTCGATTGCCAATGTTCTATAAAGGGCATGTCAGTAAATGATATGGGTCATTGAGGTGTAGCTATGCCTTCTTTTCCAAAATTCAAGAAGAGCTACTTAATTGTTCTTCACCATGTGCACGCGAAGCTGAGTAAAATGTATGAGGAGTTCAAATTGTTCAGTATTAAATTTGACTGCAAATCAAACCAAAGGTTACTACTAATGAAGTATGCTGGTGCCTGATGTGTCGACCGTTCCAAGAGCATAATAATGCCACCCGACCCAAACTGAAGCTCAGTATTGAATAAACCAGCTCTGCAGATACTTTCTTGCAATCTTTTTTTTTTTTTTTTTTTTGTTCTTTTTTTTTTTTTTTTTTTAATGA

**ROOT UV-B SENSITIVE 1 (RUS1):**

CTTCAGTCCATCTTAAAATTTTAACTTTTTCACAGCAATCCTTTCTCTCTCTCTCGTTGGCTATTTGGTCTAGATCATACTCTCTCTCTCTCTCTATTTGGCATACTCTCTCTGTCTCTATTCGGGATTGTTAGCAAATGTAGAGGACCACTAGAGATGCTCAAGCAGGCAGCTCTTCCAAGGAGAACAGCAGCTCCTTGTCTGGGAAAGGAGCTCATTAAGACACAGATAACAACCCATTTCGGAAATGGTGAGTGCTTGCTGGGTTGTGGTTTTCAAAACCCTAAATTCTATAAAGCTAGGGTTTTTAGCAGCACAGACTTCAGATGTTCGAAGAAAAACATCTCTAGAATCTTGAAAAACAATATTTCTTGCTCTGCATGGAGAAGCCAGGGTGTGTATGAGATATGGCTGAAGGCTTCTTCATGTAATGGTAATGAAGAACATGGAAGATGCGACGGCTCTGACAGAGAGCGTGGAAGCATTTCTTCAGCTAATGCTTACAAGGAACATGGAAGACGAAATGGTGATTACAGAGAGGGTTTTGAGGATTTAAGCATTCCAAAAGAAGCAAGCTGTGGAGGAGGTGGCAGTGGCAGTGGGGGTGCCGGAGGGCATTGGGGTAGTAGCGGGGGCAACAATGGAGATGAAGGCGGAGACTGGTTTTGTCTGCTTGTAGGCGATGCATATGCTCATCTTTTGTTTTTGCTCGGGCTCATCTTTACGCACGTGAAGATTGCAGGGACCAAAGCTTTGATACCTGCTCTTTTATTGCACCTTTTTTTGCATAAAGATTTGATTGCCACTGCAGATGGAATTACCTCACCTGTCAAGCAGTCTCGCGACACTAGTGAAGTATGCAGAGAGCAAGCAAAACCAAGCAGTTCACATGAAGGAACACCCGAACCCGATGTTGTATTTGAAGTGCGAGGTGGTACTTGGAGAAGGCTGTTGATTGATGCTGACAAGGACGAGTTTGTAGTTGATCCTGTGAAAGAACCCAGTAAAGAAGCATTTGCTACAGATGATCATGCCCAGGATAATATGAAAAAATCTTATCAAGGTATGACTCTAGAGCGTACTATCAAATGGTGCACGGACCTTGTCAGAAACGTGATGCTCCCAGATGGCTATCCGAGTAGTGTGACAGGAGATTATTTGGATTATACATTATGGAGAATGGTCCAAAACATTGCGAGCCAAATCAATGGTGTGCTCACCACCCAGGCTCTGCTTTATGCCGTGGGTTTGGGTAAAGGGGCAATTCCTACTGCAGCAGCGGTGAACTGGGTATTGAAGGATGGTATAGGTTATCTGAGCAAAATTATGCTTGCAAAATATGGTCGCCATTTTGATGTGCATCCAAAAGGGTGGCGATTGCTGGCGGATCTCATAGAAAATGCATCTTATGCTTTGGAACTGTTGACTCCAACCTATCCACATCTTTTTGTATACTTAGGTGCAGCAGCAGGAGCTGGGAGGTCTGCTGCTGGTTTGATACAGGCTGCCACTAGAAGTTGTTTCTATGCAGGATTTGCTGCTCAGCGCAATTTTGCAGAAGTGATAGCCAAGGGGGAGGCACAGGGCATGGTTAGCAAATCCATTGGTATTGCACTTGGCATTGCTATATCTAGTTATGTAGGTGCTTCTGGACCAATGCTAGTTACTGCTTTTCTTTCGGTCACTGGTGTCCATATGTTTTGCAACTTAAGATCATATCAAGCAGTGCAACTTCGCACTTTGAATCCATACCGAGCAAGTTTAGTATTTGGTGAGTATATAAGAAGAGATGTAGTTGCCTCTGTGGAAGAAGTCAACGCAGAAGAGCCCATCTTTACTGAAGTCCCTTTGCTATTATTTAGGCTGAAAGGGCATAAGGATCATACCAGCAACTTACTCTCTTTGCAAACTAAAGAGGATGCGGCAATCATAGCAACAAAGCTGAACATAGGTGTTTCATTTGCTAATGTAATCAGATCACAAAAGGAAGCTGATGAGCTGTTCAAGATCTACCAAAATGAGCAGTATGTTTTGGCAGAAAGGCAAAACACTTATCAGGCAATACTGAAGGTGGGAGCTACATCTCATGACCTCCTCAAATTGATGCTTCAAGTGTGCTACCTCTATCAGCTGCAAAATGGACCTTGGAGATCCTTCGCCGAGGGGAAGTTAGATTTCCATGCTGCCAGTCCAGTGGATATCTCCTATAGACTTACAATAGATAACTTCGATCGCTTACAACAAGAACTAGCCTCTGCTGGCTGGACCACTTCGGATGGATTGGTTGCTCGGCCTCTACCTTATAGGCTACTCGAAACACCCCATCAAAAAGATGATTATTCCTCCAAACACACAAGCATGGTGTAAATTATGAGACCATTTTTGTGCGCCAGTTGTCCCAAAGCACTGTTTATCAAAACTTCCCTCTATTCGGCAAACTATCATAGTCATTAATGTCAAGGTTAAGGTAGTTCAAGGCTCCTATACCTTTGTAACGTCTTTTGAGAATGCTGATTGTTACAGGCTTCATTTTTCAGGTCATATTTGACCAACGTCCCAAGGTACTCTGATCATTTGTTGCAGCGAATTCAATCACACACATACCTTGAAAATGTGAGTTGTTG

**ROOT UV-B SENSITIVE 2 (RUS2):**

CACTGCTCCTAGAAATAGAAGAATAGAAGTATTGTGCAGAAAAAATGCAGGAGCGGGTGAGATGGGTGGAGCGGAAAGAGAGTGTTCGACAAGAGTTCTCATTCGACGGCCGTGCCTCCGACATCCTCTGTTCCAAGACTTTGAAAGACGAGCGACATTTCCCTAACCGTATTGTTGAATCCTTTCTAAGCCGGTTCTTCCCTCTCGGTTACCCTCACAGTGTTGCAGAGGGATACTTAACTTATTCTCAGTTTCGCGCTCTGCAACATTTTTCGAGTGCAATGCTATCTGTCTTATCAACCCAGTCTTTGCTATATGCTGCAGGTTTGAGGCCAACTCCCGCCCAAGCCACTGTTGTCAGCTGGGTTTTAAAGGATGGAATGCAGCATGTGGGAAAACTAGCATGCAGCATGTGGGGAGCTCGGATGGATGCAGAGCCTAAGCGCTGGAGGCTATTCGCTGATGTCCTGTACGACTTGGGGACTGCTTTGGAGATTGTTTCACCACTTTGCCCACACTTATTTTTGGAGGTTGCAGGTGTGGCAAATCTTGCCAAAGGAATGGCCACGGTCGCTGCTCGAGCAACCAGGTTGCCTATTTATTCTGGGTTTGCCAAAGAAGGGAATTTAAGTGACCTTTATGCCAAGGGTGAGGCGATTTCCACCTTATTCAATGTGTTGGGACTTGGTGTTGGCATACAACTTGCATCCACTATGTGCTCAACCATGCAAGGGAAGCTTGTGGTGGCTCCTTTGCTTTCAGCAGTACATTTATTCAGTATCTCTCAAGAGATGCGTGCAGCTCCTTTAAATACGTTAAATGCACAGCGAACTGCCAGCTTAGTTGCAGAGTTTGTGGAGACTGGCAAGATTGCTTGTCCGGCAGAAATTAGGTACAAAGAACGCCTTGTACTGCCAGTTTGCTTAAATGTTGATGCTGGAAGTGTCGAAGCAGGTGTTCCGATCAAGCAAGCAGTTGGCAAACCTTCAGTGTTGAGGGAACTCAAAGCTAAATTTCAAAAGGAGAGGTTCTTGCTGTGTTTTAAGGACAAACAAACAGAGATGGTACTGCATCATAATGCAACAGGAGAGGATGTATTGAGAGGCTGGCTGCTAGGTGCATATGCTGCAAGGATAGCTGGAGTTGAAGGCATGTGCACCGGAGCCAAAGGTTATGGGGAGCAGAAATTGGAAGCTCTTGGGAAGGCCTATGAACTCGTGGAGAGGTCGCTTCCAATTCTGATCTGTGGTTTAAGAGAGAGAGGCTGGCATACAGATCTGTTTCTGGAGGGCTCTGGAGAGCGAGCAACCTGGTAAAGGAAAGCTCTTTTTATACTGAAGGATAGATTGCATACTTTTCAGAAGCGTACAATCTTATTCTTAATGTTTGATGAAATGCAATACCATTAGTCTTAGTGGAGACCAGATCGATGTCAGGCTCTTTGATACTTGTTTTAAGTTGTCCTCCAAAGCACCCAGTTTGTAGTTGTTGAGTTTGCATGCAACCCAAGGGGACAGTGTCAGGTCCCGGCATTGTAGAGGGCATCCCATCTGCACCCCAATAGCCACAACCTTCAGGGAATAGTCCTTTATGTGAACCAACGTGTAGTTAGCCTTGGCTGTATCAATAGATTACTCTTGCTTATGTATCTATTGCCAATCTAAAGGAATTTTGTGGGCATTGTTTGCTTTTGAAGATTATGCACTTTCTTGACCAAGACTACAATTTGTGTATCAATCAGGCGTTACCAAATAAATGAAGACATTCAAAGTGTTAGCCTAAT

**COP9 SIGNALOSOME COMPLEX SUBUNIT 1 (CSN1):**

ACTTTACGAGCTACGAAGTTTGGCACCCCTGAAATAAGACTTTAATGAGAAATCAGATGTTTGGAATCAAACAAGAGGGTTCACTAGTGCCACTACAAGTTCTAACATCACTACTTAGGAATCATTAGCCCACACGCGGCAAATTTGGCTGAGAATTTCATGTAATAGGAACCCTGCACTTCTCAAACATGTGACCTAGACCCATTATATCAAGGCCATTCCCACACAATAAAGAGTCCTTAAGACAGAATCGGTCATACTCACCTGTATTCTCATTCAAAATTTTGAAAAAAAAAGGGACCTAGACCAGAATTTCATGTAACAGGAACCCTCCACTTCCCTAAGATTCTCTAAGTAAGAAAGGTTTTAAATAATAAGCCACCATCTTAATTGAACCAAGCCAATGATTAGCCACACAAGATGTGCTGAGACGAGGCTATCAGTGTTGAGAAATGTCTACTCTTACATTATCACTTTCTCGACCCTCCTCTTACCACAATCTCATGTTTCAAGAGATTTGCACGTACGAGCATGGCACGTGTATCTCTTTCAAAGTCCGACCCTGTTTGCAGAGCTCGCTGAAAAGTTCCATTCCTTTGGTCAGCATGACGTGCATGTAAAATCTTGTTATGTGAATCGATCCGAGCCTGAATTTGATTCTCCATAATTAAAGCTGCAAGCTCCTTCTCTAGGCCAGCTACAGTTGTCTTAAAAGCATCAGCCATTCGATTCAAATCCACAGAAATAAAAGGAGTTGTATACTGAATAAGAGCCTTGTGACGAATTAGAGTGTACAGAGATTCAACATGGTCATGGAGGTGAATATCTAGCAGCAGATTTGTCTTCAGCTTCTGCAGGTAGCCCAAGCAGGAAGCATAGCGACTTGCATAGAAATCATGTATAAGTTCTCGAACTTCTGGAACTAGCTCTAGAAAATTCTTGAAGTTAATATTGTCAATAACTTTGCTCTTCAACTCTGCTCTGTCAAAGGAAGCTAAAGCACAAAGACCTCCATATGTAGCCACGTCTTGAGGAGCAATGACTTCATTATAAGTAGCACCCAGTTCAAAGCTGGTATCAACAAACTTTCTGGCAGCAAGTTTGTACTTCTGATTCTCTAAATATGCAAGGCCCGCATCAACCCGCAGCTTTGCCACAACCTGAGGCTCCTGGTTATCAGGTGTCTGCTCAGCCTTCGAAACATAATTTGACACATGATGGAATTGCCCCAGGTTGATGGCAACAAGGATGACATTGAGGCACATAGATATGATATGCTTAGATGTAGTGCAGTAATCACGAGTTCGAACAAAGCACTTAAACGCTTGTTGCAAATCGCCTCGTGCATTGTGAAAGTCTCCCAAATCATTGTATCCCATTCGAATACTTTCTTTGATCAAATTTGTCTTGTAGCCGTTAAGCTCAGTTTCAAGGCGTTCGAGGCGCTGGGCAGACCGACGATCGACAGTATCCACCCACTCCTGGTCAAGGAGGAAAGTAGGGCCAAGGCGGGTATTGATTTTCTCACAAATCTCTTTGTAGAGGGAGGTGCTTTCGCCTTTGCGAATCTCTTCGTGGGCCATGCGCAGAGCCTCTAGCTCCATAGCCTGGTTGCGGCAGTGCTCAGCAATGAAGAAGAGGCGCAGGATCTTGGTGCGACCCTTGTAGAGGCTTGCATAGGCTTCTACGTCCATTTGGTCGCCTGAGACAGCAGGATTGTGATCTTCTGCTGCAGGAGATAAGCCCAAGGAAGAGGGCTCCATCTGCTTGCCCTTACGCTGCTCCTCTTCTTCTTCCTCTCCCTCTGTGTTTGGAGGATCCATCGCCTCCATATCGTCGTCTGAAAACATCTCGCTTTCCCTCCTCAGCTTCGATGCTTCGAATTTCAAGAATTTTCTATAGACAGA

**COP9 SIGNALOSOME COMPLEX SUBUNIT 2 (CSN2):**

CATGCCTTGTACACACTAGTCAACGTGACCTCAAAAAGTTCTACGGTGATCTCTTCTAATTACTCCAGAAGAAGACTCACTTATATAGCAGTAAAACCTTGTAATGATAAAAAGGTAGCATACAGGGTAGTCTTTCAAAAGCACATAGCGGAAGAGAAAACCTCCGCAATGTGCAAATTAATATGGGAGAGGGCAACCTGACAAACACCTTTCTTGATTGCATAGAATCTCAAGGGAGCATAAGCAGCCCAGCCCCTTTACAAAAGCCGATAATTTCCTCTGAAAGACCACAAAATTTTTTACTCAATTACACCAAGCTGACAATTCAATCAACACCACCTTTTGCATATGCAGCGGTAGTACTCCAAGCCAGGTTAGCCCACCCGTGTTGTGACCACTTGGCACAAACCCCTTAATTGAGTGTTCCACTTATCGATTGCTGTATACTTCTTTGGGCCTTTTGACCTGTCCCCGAGTTCAAGGAGCTGGTTGACCTGATCTATATGACCATGTACTCGATTATCCAAAATTAGCGAGACAAGCAGCTGCTCCACATCCTTTTCAGGAATATTCAGCTCCTGCGAGATGAATGGAATCCGTATCCGTGTATATGGTTTGATAAGCTTCAAGAGAACTTGTGTCCGAATGTTCTTCAGCAAGTCCTCAATGTAATTGCGAATGAATGGGTCGTCCATTATCGTTTTCCGATTACTCTTAATTATTTTCTCAAACTCCAGTATATCATTCCGCTGGTATGCTGCAACGAGATTGGTCATTGCAAGAATTTCAGGATCATTCTTGTACGGTTTTGCTTCTTGACCATCGAAGGGATTAACTTGAGACTCCATAAGCATGTTCGCAAGTACTAGGTACTTGAGACATTGGATGCGCCGATGCGTTCCTGCCTCATCGTAGTTCTTGAAAGCATCAAAAAAGTCTGTTGCAGCATCCGCCCATTGCCGTTCTGCCATGTTCATCTTTCCCCCACACTCGTGAATGATCCCCATGATACGTGGATGTGGAATTGCAGATTTGATGGTTAGCGCTTGCTTATACAATTGCTTCAATTTCTTGTTATTTTTTGTCTCTGTGTACATCTGTATCTCAATGGCATACACTTCCAATAATTGTGTGCCTTTCTTCTGATCATCTGAACCATCCTCTCTCCGGCAGGACTTGTGAAGCTCCTTCAGTATCTTGTTCATGCGACCATACTCTCCCATATCAAACCACAATTTGCAAAGCTTTAAATTTGTTTTGAACCATAGCCGCTCATTTTTGGCATCCTCCAAAGCCCTCAGAGTTGTCTGATAAAATTCTTGAAGTAATTCCATGTTCTGACTAGCACTTCCTGATACAAAGTCCAAAATATTATTTATACATTTCTCACTGTAGTTTCGTGTCACCGCTGATTTGATGTACATCAACATCTCTCTGTAAGCATCCATCATTTGCTTAAAATTGGCAAGACGATAGTGTAGTTTGACTGTCTGCTTCAAAGCCTTGAAACCCCATTCACCCTTCTCCCCCTCCATCTTGACCACCTCCGCAAAACCAGACAAGGCACCTTTAGGGTCCATTTCCACCAAACCCTTTGAATTGTAATATTGGTTTTCAATATCAACATCCTGCTCTTCTGGCTCCTCATCTGAGTACTCAAAACCATAATCTTCCATATCTGCATCTGAACCCATAGCTTGAATTTGCTGAGAGTCCGAGACGATCGACGAAGAGAACTGTTTTGTGTTAAGCGAGTTTTCGCAGCAGACAGAGTGTATGTTATTTCTGATTTTTGCTTCGGCCGGCCACAGCTGAAGCCCAAGTTGTGGGTTCAAATCTCTGAAAGCCTTGGATGATATTTGTGCCCTTGAGTTGTGGGATAAGGGGGGTCCCAATCCTATTTGTTCACCCGCCCTTGGGCTACTAGCATTTTGGCCTATTTCCTAGGGATTTGTAGGACCTAGTTATGAACCCTTCTCTCTCTAATGAATGTTGCCAACATTATAAAAACTCCTTAAAAGCATTGACATTAATATGGGTCTCCTATGTGTTCCTCCTCAAAATAGGCATTAAAAATATCTTTAACAATTAGTCAAAGTAGCCAACCCAATATAAAGGGCCAAATAAAG

**COP9 SIGNALOSOME COMPLEX SUBUNIT 6A (CSN6A):**

GTCTCAAGCTACACCATCGCTTCCATGTCCACGTGGACTGCACAGCAGAGGTGTATTGCATACATGTTGTGTGAATATCCACTTGGAGCTATTTATTATTTTCACACAACTTCGTTTGCGTAGGCTATTCACATAACTTCGTTTGCGTAGGCTTTTAATTAAAAAATGCCACGTCAGTGGCATTTTTTAACACAAACGAAATTTCTCTCCTTGAGTTTGTATCTCTAAATTGGTAAATTGTAAGAAGAAGCCCGAGAGCCTGGGTATCAACCGCGAAAGCAGCAGCATCGCAGCTCTCGCAGCCTACAGCTTCCCTAATCAGAGATCGCCATGGCTTCTAGCAGCGGCCTTACCTTCAAGCTTCACCCGCTCGTGATCGTGAATGTCTCGGATCACCACACGCGCATCAAGGCGCAGGAGAGCTCTTCTACTAGAGCCCCCCCTCGTGTCTTTGGCTGCCTCCTTGGATTGCAGACCGGCCGCACTGTCGAAATCTTTACCAGCTTCGAAACCAAGCTCTCTGAGGACTCCTCTGACTGCCTTGATCGTGATTTCCTCTACCAGAAACAAGAGCAATACAAAAAGGTCTTTCCAAAATATGACATCTTGGGATGGTATTCTACTGGAAGTGATATCCAGGAGACTGACATGCAAATGCACAAAGCTCTGATGGACATCAATGAAAGTCCTGTGTACATGCTTTTGAATCCAGCCATCAATCATGCCCAAAAGGACTTGCCAATCACCATCTATGAGAGTGAATTACATGTTATTGATGGAGTGCCATCACTGATATTTGTCAAATCAAGCTATACCATTGAGACAGTAGAAGCAGAGCGAATTACAGTTGATCATGTGGCTCATATCAAACCTTCTGATGGAAGCACTGCTGCCACACAATTGGCAGCTCATTTGAGTGGCATCCACAGTGCGGTCAAAATGCTGAATTGCAGGATTAGGGTTCTTCACAATCTCCTGATCTCTATTCATAAAGGGGAAATACCTTATGAACACTCGCTTCTGCGACAAGTTTCTAGTCTTCTTCGTAGACTGCCTGCCGTAGACTCTGCAAAATTTCAAGATGATTTTTTGATGGAATACAATGATACTCTGTTAATGACTTATCTTGCTATGATGACAAAGTGCTCAAGCACTATTAATGAGTTTGTGGATAAGTTCAACACAGCTTACGACATCAAGCAAACCAGACGAATGGGCAGGATGCCTTACATATAGAAGAGTATAGTAGTGAAACCTAATGGGACAGTCTTTTGCACTCGGCTCTGTATACAAGGATAAATATGGCTGGCTTGCATACATAGTGACACATCTCTTGGATTGGTGTGGGCATAGATTTCCGCATCTCACCTAATATTATGATCTTTTCGGGCCAAAAGTTTCGGTGTCAATTTATAGCAATCAAGTGGCAGAACATGGACCTTTTGGCAATGTGCAAGGAAAAGGCTCCACCGAACATTGAAATGTATAGGTGACTGCATAGGAGAGCCCATAGAAGCAAGTTTGGTAAAAGTCAGCCTTGGGCTCTGCAGTCATGGGCCAGTACAATTTTGATACCTGCTTATGGTTTACATGTTTTGTTTTAGTCATGGCATGTGCATGTAGTCTATATAGGTCGTGGATGCAACTAGTAAATGCTCAGAGGAGGTTCATTAATCGTCTTATGCGCTGGATATGCCTCTATTCAATTTTATGTAGTTTCATATACTTTAACTCGCAGTACACGGCAGTACTTTTCTTCATTTTCTTTTTCTCATCAATGACATATCTTATAAGAATTCCCATGGACCATTTTTTATGAAAAAATCATCCCAAATTCCTCATATTTGGTGCCTTGGGCAGATTAAGCTTGGTAGTGTAGGGTCTCATTTGCATTGGCATTAACCATGTTGTTAACTTTTACTTCACTTTCGAACATTACAAGAATATGTAGCTGCAAGTAACATTGCATGCATAATCATTTTCAAACATGTAACTTTTGATATCTCTTCTCTGTTGTTGATTCATACTTGGGCATGAAAAA

**DAMAGED DNA BINDING PROTEIN 1A (DDB1A):**

GGCGAACGCACTCGCGCGGGCACGCACAAGTCATTTCTCTCTCTTTCTTGTCTCGATGAGAGAGATTACTCCGAGCTTATAAACAAGTAACCTGAAGTTTCCCTCGCTAGGGACTCTGTAGGAGAAGGAGTAGCTGCAGAAGATGAGCATCTGGAATTATGTGGTCACGGCGCACAAGCCTACCAATGTCACACACTCCTGCGTGGGCAACTTCACTGGCGCCCACGAACTGAACCTCATCATAGCAAAATGTACACGAATCGAGATACACCTGTTGACGCCGCAAGGCTTGCAGCCTATGCTGGATGTACCAATATATGGACGTATAGCAACACTCGAGCTTTTCAAACCACCAGGGGAAGCACAAGATTTTCTATTCATATCCACAGAAAGATACAAGTTTTGTGTTTTACAGTGGGATGCAGAAGCAGGGGAACTTGTAACCAGAGCTATGGGAGATGTTTCTGATCGAATTGGAAGAGCCACAGACAATGGCCAGATTGGCATCATCGACCCTGATTGTAGACTTATTGGTTTGCACCTTTACGATGGGTTATTCAAGGTCATACCAATCGACAACAAGGGTCAACTGAAAGAGGCATTTAATATTCGCTTAGAGGAACTACAGGTCCTTGATATCAAATTTCTTTACGGATGCGCTAAACCCACAATTGCAGTTTTATTTCAGGACAACAAAGATGCACGCCATGTCAAAACATATGAGGTATTGCTCAAAGAGAAAGACTTTGGAGATGGTCCTTGGGTGCAAAACAATTTGGACAATGGGGCTGGTCTGCTTATTCCTGTGCCTTTGCCATTGGGAGGTGTCATCATTATCGGGGAACAAACTATTGTGTACTACAATGGGTTTGTTTTCAAGGCCATTTCTGTACGACCGGCTATCACCAGAGCTTACGGAAGAGTCGATGCAGATGGGTCTCGCTATCTCCTAAGTGACCTCACAGGGATGCTACATTTACTGGTGATTACTCACGAGAAGGAAAGAGTTACAGGCCTTAAAGTGGAGCGCCTTGGAGAAACGTCAGCTGCTTCAACAATCTCTTATCTGGATAATGCTGTGGTCTTTGTTGGGTCTAGCTATGGAGATTCACAGCTTATTAAGCTCAATCTACAACCTGATTCCAAAGGGTCATATGTAGAAGTTCTTGAGAGCTTTGTCAACCTTGGGCCGATCGTGGACCTGTGTGTGGTAGACCTTGAGAGACAGGGACAAGGGCAGGTCGTTACTTGTTCAGGAGCATTCAAGGATGGATCTTTGCGCATTGTTCGGAATGGTATTGGCATCAATGAACAGGCATCTGCAGAACTTCAGGGCATTAAAGGCATGTGGTCATTGCGGGCTGCTTCATTGGATGCATTTGACATGTTTTTGGTGGTTAGTTTTATAAGTGAGACACGCATATTGGCAATGAACTCAGAAGATGAGTTAGAGGAGACTGAGATCGATGGATTTGATTCAGATGCCCAGACCTTGTTTTGCCAAAATGTCTTGTATGATCAGCTCATCCAGGTGACAGCATCTTGTTTAAGACTTGTTGATGCAACTTCGAGGAGGCAGCTTAATGAATGGAAAGCCCCTCCTGGTTTGTCAATAAATGTAGCCACAGCAAGCTCCTCACAGGTTTTACTCGCAACGGGTGGAGGCCACCTTGTCTACATAGAGATCGGCCGCGGTCTTCTGATGGAGACGAAACATGTGCAGCTTGAGTATGAGATTTCGTGTCTGGACATCAATCCTGTTGGTGATGACCCAGACCGAAGTCATTTGGCAGCTGTTGGCATGTGGACAGATATTAGTGTGCGGATTTTCTCCCTCCCAAGCCTTGATCTCATAACAAAGCACAGTTTGGGAGGTGAAATAATACCTCGTTCAGTTCTCTTCTGTGCTTTTGAGGGGATTCCATATTTGCTTTGTGCGCTAGGAGATGGCCATCTGTTCAATTTTATCCTAGATGTCCATACTGGGCAATTGTCAGACAGAAAGAAAGTCTCTTTGGGAACACAACCTATTACTTTGCGGACGTTTCGCTCAAAAAATGCTACTCATGTCTTTGCAGCCTCAGATAGGCCAACTGTTATCTATAGTAGCAATCGAAAACTTTTGTACAGTAATGTAAACCTAAAGGAGGTCAATCATATGTGCCCATTTAACTCAGCTTCGTTTCCAGACAGTTTGGCGATTGGAAAGGAAGGGGAACTCACAATTGGTACAATTGACGATATTCAGAAATTGCATATTCGAACAGTCCCTTTGAATGAGCAGGCTCGGCGCATTTGTCACCAAGAATCATCTAGAACGTTCGCAATTTGTAGTGCAAAGTACATGCCAAATATGGAGGATATGGAAACACATTTTATTCGGTTGCTGGATGACCAAACATTTGAAAATATGGCTTCTTTTCAGCTAGATGCATTTGAAAATGGTTGCTCAATAATGAGCTGCTCATTCACAGATGACAACAATGTTTACTTCTGTGTCGGGACGGCATATGCTATTCCAGAAGAAAATGAGCCCTCAAAGGGCCGTATATTAGTATTCCTTGTGGAGGATGGGAAGCTACAGTTGGTTGCTGAGAAAGAGACAAAAGGCGCGGTCTACAACTTGAATGCATTCAATGGAAAGCTCTTGGCAGGTATCAACCACAAGATTCATTTGTACAAGTGGACACAACGAGAGGATGGTACAAGGGAGTTGCAATCAGAGTGTGGCCATCATGGACATATTCTGGCACTCTTTGTACAGTCTAGAGGAGATTTCATCGTGGTTGGAGATCTAATGAAGTCAATTTCACTGCTGATATACAAGCCAGAGGAAGGTGCAATCGAGGAAAGAGCCCGAGATTTTAATGCGAATTGGATGTCTGCTGTCGAGATTCTAGATGATGACATTTACCTTGGTGCAGAGAATAGTTTTAATCTGTTCACAGTGAGAAAGAATAGCGATGCTGCCAGTGATGAGGAGCGGGGCAAGCTTGAGGTTGTTGGCGAATATCATCTAGGAGAGTTTGTAAATCGCTTTCGGCATGGTTCTCTCGTCATGCGTCTCCCAGACAGTGAATCAAGTCAGATCCCCACTATGATTTTTGGCACAGTGAATGGTGTTATTGGAGTGGTTGCATCTTTACCACAAGAACAGTATATATTTCTTCATAAGCTTCAACAAAATCTTGTCAAAGTGATCAAGGGTGTTGGTGGCCTGAGCCACGAACAATGGCGCTCCTTTAGTAATGAGCGGAAAACAGTAGATGCTAAGAACTTTCTGGACGGTGATCTTATCGAGTCCTTTTTGGATCTTAACAGGACCAGAATGGAAGAGATATCCAAGGCAATGGCTGTTTCTGTGGAAGAGCTTTGCAAGAGAGTGGAGGAATTGACTAGGCTTCATTAAGTTAATGATATATTTCTGCAGCCTAGGCACTTGGGGATAAATTGTGCTCTAGCAGGCAAAATCAGCTGCAGGTTGTAAAAAATGTTGGCACAAAGCCCCTTCTTGGGGGTGAAAGCATTTTTCTTTCATCTGGCTGGTTTATTAGGTCTTGCAAGATCAGGAGGCAGCAGTGAAGCATAATTTTATAGAGCAGTGAAGCATAATTTTATAGAGTTCAAATGTAACTGCTTGGGCTTAACCCAAGAGCATTTTTTTTTCTGCCTACAGGGCTTGTTCTCTTGCAATGCTCATCTGCTCTGGTAGTGATAACAGAACATTTGCTGCCCCCTTGTGTTTCAATATATGGCTTTTACTAAATATGGCAATAAAGCATTTTTTGTCTCTCTGGGAGTCTCGTTTCATTCATGCTCTAAATTATAGACATTGAGTGAGCATACACGCCGTTCAAAGGTACTGTCTCATCATGTTCTAAAAGGCCAATTTGGATAGCCTAAAAATTGGTTTTCTCCTGCATATCTGGAGTTGTAAATATTGAGTATTCCAAACAGACATTAAGAGAACTATTGAAGCGCGTACGTAAGTAGTGTGGTGTAGGGAAAGAAGTAAAGTAGTTTTTGCGTGATGCAGTGTTTGCTCGATGCTTGTACTTGACATGAGAACACTGGAACAAGGG

**EMBRYO DEFECTIVE 168 (EMB168):**

GGACACACTAGGAGGAAATAAGGACACGAGAGCAAAATAGAACAGAGCATGAAATGCGGGAACGATTTGGGTGGTGGAGGCACTAAAGCCTTGTGTTTTGATCACGTTTCCTTTTGTTTTCCACTGACGCTACTGCCCGGAGGGCAATCCGACGCGCCCTTGTACCTTGTCACAATCCCTTTTCTATTGGGTCTCTGCGCTACGTTAAGGCAATCCTCGTCGTTGATCTGGGTCTTCAGCAAATCGTCGCCTTTTCTCTCTGTATTACCCATATCTGTATTACCCAGATTCTTTACTCTTTTCCTGTGTGCAGAGAATTGCAGGCCAGCAATGCCTCTATACAGGTCTTTTGATTACTCGAAAGCAGGTGATCTTTGAGTCCACAAGAGCTTATGGGTGTTTTGTATTAGACCAGGATGAGACCCAGTACCCATTTTTCCCATACCTTTGATCACATCGCTACACGAACGATTACCTGTCTTACCTCTTACAGGTCAAGGGCAGCTCCAATTAGGCTCCTACAGCATGCAACACTGATGTAGATTTTCTCACAATTGCCACCATGAACCCAGAAGGTTCGGTTGGCGTTCAGGTTCCATCTGTGACTGGCGGGAAGGGCGAGCCCTTACGGTCTGAGGACACTGATGCCGGGCAGTCGAAACGTGACCTAGATAAAGACATTTTATGTCCAATATGCTTCCAAACCATGAAGGATGCTTTCTCCACTGCTTGCGGACATAGCTTTTGCTACATGTGCATTATGACCCATCTCAACAACAAGAAGAATTGCCCTTGTTGTGGACTCTATCTAACTAACACACAAGTATTTCCAAATTTCCTTCTCAACAAGCTTCTGAAAAAGGCATCAGCATGCCAACTTGTAAGCAGTGCTACTCCCTTGGAACACTTTAAACTATACCTTCAAGAGGGAGCGGAGGTTTCGGTTAAGGATCTGGACTCACTCCTCTCTCTTCTTGCTGAGAAAAAAAGGAAGGCTGAGCAAGAGGAAGCAGAGACAAATATGGAAATACTTCTTGATTTTTTACACAGGCTTCAGCAACAAAAACAAGAAGAGTTGAATGAGCTACAAGCTGATCTGGAATTCATTAAAGAAGATGTCTCAAGTGTTGAGAAGCAGAGGCATGAACTTCTGAGAGCCAGGGAACGGTATTCGGTGAAGCTGCGCATGTTGTTACAGGGCTCAAGTCCTCATGTGACCAATGCAACTCCTAGCACGGGCAATGCAGTCTCTATGCCAAAAACTGAGGCAGGATTGGCCATGATATCAACGGCTCTAAAAAAAGGAGATGGCAAAGAGAGAGCTTCTTCCATAACCTTTCAAAAAAGAGATCAACCATTTGGCGGATCCACAGACTGCATTTCCAACCTTCAAGGCCCAACACTTTCTCCGAGCATGGCAATCGCAAAGAAGAAAAGGGTTTTAGCGCAGTTTGAAGACTTACAAGAGTGCTACCTGCTCAAAAGGCGGCGGATGGCACCCTCCCAGCGGCCGAGGCAGCTGGATTTGAATTCCAAGAAAGACAAAGATAGCAATGACAAAGATATTGGCAGTGCAGGCCTCGAGGACTTCCAAAACATTTTGAATGCCTTCACACGTTACAGCCGCCTTCGGGTAATAGCAGAAGTCCATCAAGGCGATTTGTTCCATTCTTCTAACATAGTTTCCAGCATTGAGTTTGACAGGGATGATAAGCTGTTCGCAACTGCGGGTGTGTCAAGGCGCATCAAAATCTTTGAATTTGCGACGGTCGTAAATGTCCCAGCAGAGGTGCATTTTCCTGTCGTTGAGATGTCGACAAGATCAAAACTTAGCTGTCTTAGCTGGAACAAGTATTTAAAAAATCAAATAGCGAGTAGTGATTATGAAGGCATCATAACCGTTTGGGATGTAAATACACGGCAGAGCATCATGGAATACGAAGAGCATGAGAAACGCGCATGGAGTGTTGATTTTTCACGAACCGACCCTTCGATGCTAGTCTCTGGAAGTGATGATTGCAAGGTGAAAGTTTGGTGTACAAAGCAGGAATCGAGTGTGCTTGATATAGATATGAAGGCCAATATTTGCTGTGTCAAATACAACCCTGGGTCCAGCCATTACCTGGCGGTGGGTTCAGCTGATCATCACATTCATTACTACGATTTGAGAAATACTCACACTCCATTGCACATATTCAGTGGACACAGGAAAGCGGTCTCATATGTGAAGTTCGTTTCACCTGTGGAGCTTGCCTCTGCATCCACGGATAGTACTCTACGCCTCTGGGACGTGAAAGAGAACTCACATATACGGACCTTGAAAGGACATACGAATGAAAAGAACTTTGTCGGGCTTTCTGTAAACAGTGACTACATTGCTTGTGGCAGCGAGACAAATGAAGTGTGTGTCTACCACAAGGCGATTTCAAAACCAGCAGCATGGCATCGGTTTGGGTCTTCAGACCTTGATGATAGTGATGACGATTCTTCCCACTTCATAAGTGCAGTTTGTTGGAAGAGTGAAAGCCCAACCATGCTGGCTGCCAACAGCCAAGGTACAATCAAGGTGCTAGTGCTGGCTCCATAATTGCTTGATTCAAGCCAACAGATTCTTCAATTTTGTGCCGTGTATGCTCATGCGCGATCTAATGCCAAGATGAAAACTATGTGCTAGTGAAATTCTTGTTTTTGAGGCTGCTCTACCTATCCCAATCAGGGTTCGAAAGGGCACATGATGCCATTATATGTTTCATCTCACTTGCAGCATGCAATCCCCTTGCCTGTCTGCCTTCTAAGGCTCCTTGTATTTGTGTTTCTATTGATGAGTAATGTTTTGTCCAATTGGCACACAGATAAATAGCCAATTTCCTGGCTTGTTGTGCAAAATGTTCACAAAAATTCAAGCAGCGACTTTGTAACCTGTAACCTTGTTATTCAAAAATTGCACAATGGGCATTACCACATCATGTCAGTATTTTGTT

**SPA1-RELATED 3 (SPA3):**

CTGTCTCAGGCTCCAAATGGCATAGCTGGCATATCCTATCCTCTGCAGGTATCCCCCTAAATCTGCCAGTTTCGACCACGAGAGAGAGAGAGGGAGAGATATGGACATCCATCCCTTCAAGGACATCTGATAAAACTGGAACAACACAAAGTACGACCCCTGCGCAATCATGACATGCACAAATCGGGAAATGGTTAAGCAAGTTATGGACATCCCAGTCAGCCATCCAAGGAAGCGTAGTCCAGGACGAGGAAACATAAAGAGGAGGTGCAGGACCATTTCTATGCTTCCCCTCCCCCCCCCTTAAGTATCAAAAATAAATTACATTTTTCTCCACTAGTTAGCATCCGCAGGTTGACCTTGTGGTGAAGCAGACCCCCGCTAAGGCAGGAAACAAAAATCTTGCATCAACATTTTAAAATACAAATTTAGAGGCATGGGCAAGAAATTAGAAGTCCCATTATATCAGCACTTGCACTTGTAATGCCTTGCACATTGTCGATTGTTTGTCCTCAAAACTTCAGCGTAAAAAAATCACAGGTCCAGGACAGCAAACTTTCTTACTCAATTTTTGTGGAAGATTGCACTCTCTGCCTGTATTATGGCAACATTGTACAAGCTTCTCAAATAAAAACGTATTGTACAAACTGATCAACAACAAAATTGTAACTCTGCTGTGATTCGTAACATAAAATCACCAATCTTGGCTGATCACTTCACAGAATTTGGACTGGACTTCAACCACCATCAAACAGATATGGGTCATCCGGATTGTAACCACTTCCTTTACATGCTAACCCTAACTAGAACAAACTGTTAGTATACTGCCTATTGAAGGAGTCCGAAATTTTACTGTACAAAATGCCTTGCTGGAATGTTCCAGCATATTATATATATGGGTCTGTAGGAAAAAATTATCATTACAGGTACAAGCTTTGTGGATGGAGCCAGATTCCTTGCAGCAAAAAGGGCAGGTCTGTACTTGCAGAAAACCCATCTGTTGCATCTCGAGGTAAGCCCCATCTGCTTGGTGCAGCAAAGGCTGCCAATCCACAATCTTCATGTCGAATTTGCAAATGCTGTAAGCATATCATGACTGTAGGAGAAAAACGGCATCGTAAGTATGCAGGTAAAAACGCATTGCTGAGAGGCCTTGCACCGACGATCGAACAAGATGCAGGTAAAAACGAATTGCTGAGAGGCCTTGCACCGACGATCGAACACTGGCTTCTGCCTACCAGGTATATAAATACAGATTTGAGCCTATCAATGTGATAGTAAGGGAACACTGGCATCTGCCTACCAGGTATTTAAATACAGATTTGAGCCTATCAATGTGATAGTAAGGGAAGGGAAGACAGAAGGCATCCCAGCACAATCTTCCACTTTTGTAACATCTTGAGACTGGTCCTCTCCATAATACATATGCCATTAATCAAGTCAGCTGTAATCACTAATGCCGTATATGTTTCCACTCACCTTGTGTAAAGTAAAGCAATTCGTATTATATTATCCTAAACCATCTCCAATACCTGAATATTCCCCACTGAGTTAGCTGTCACCAAAATTTGAGATTTCTTTCTCCAGCAGACACTTGAAACAAACTGATCCCCAATATCTTCAAGCTCTCGACCTGAAATTGGATCAGCATTCTTGAACTTGTGCGACAGCATGGTCTCCGGAAGAGACTTGTGGTAAACAAAAACCTCATTTGTTTCTGAACCACAAGCTATGTAGTTTTCCTCTGCAACTGAAAGTCCAACAAAATTCTTCTCATTGGTATGACCAGTGTAAGTGGTTATGCAATCACTCCTTGATCCAAGACTAGCACCATACCCTCTAGCATTTATGCTGGTTTTAGTCAAATCCCAAAGTTTCAGAGTGTTGTCAGTTGATGCAGACACAATGCCTGTCGGGCCCACGAATTTTATGAAAGAGACAGCCCTTTGATGGCCCGTCAAAATGCAAACTGGGATTCTGATGTAACGAAGGTCGTGAAAATACACTTGGCTATCTGCTGACCCAAATGCCAGAATATGGGAATGATCCGGGGAAAACTGCACGCTACATATGTTGGCTTTTGTCTTGATTGTCCCAACACTTGTACCCTGATTTGTGCTCCAAAGTTTAACACAGCAGTCATCACTCCCACTTGCTAGTTTTGAAGAGTCACTATGAGAAAAGTCTACGGACCAAGCACATTTGCCATGCTCCTTGAATTCCATAATTGTTCGACCAGCTGTTACATCCCATAACTGAACAGCCCCTCCAAGGTCACTGGATGCCAAGTAGTTTTTGCGAAAATTATTCCAACGAACACAGCTCAGTTTTGTTCTACTGTTCATCTCGACCACAGGGTAATGCATGTCATTAGACTCTTCCAACAAAGCATTACAGTCGAAAATTTTTATTCTTTTTGAAACTCCTGCTGTTGCAAAGAACTCGTCGTCGCTATCAAAGCTCACAGAGCACACCATATTTTTTGTGTTCAAAATGTCACCATACTGCAATGTAGCCTTGACGTCAAACCTACTGTACTTCATGAATTTGCAAAACTCATCAAAAAAGCATCCCAGTCGCCCGCCACTATCACCTTGTTCTGCTTTGGAACCCCTGAAGCTGTTGAATACTCTTGGCATGCCCATTTTCTCTGCACGTGACCCTTGCCCCAAACCACTTTGCCTGATGACGGCATTTGTCAATCGAGAGCCAGAATGATGCTCCCTCTTTATTAGTGAAAGGTAAGCTTCTCTCAAATTGGTGAAACTCCTCTTAAGCTGAGAAGACTTCGCAAGCACAAGTTCCTGCCTTCTACCACTCAACCCACTTCCGCTACAGCCAGAACCTAGGACTTCTTCATATGTAAAACTTTCCACAAGCCTGCTACTGTTGTTATCAAAACCAGCCTTCCGTAGTCTGTTCTTCAAAAGTAAAGCCTGCCATCTCTCAACCTCTTTGGTGTCTTCATCTAGCAAACCAAGCTCTTTAGCTAGCTTCTGAATGACGACACTCTTTTTTCGCTGCATCTGTAGGAGAAAGTCCAATGTTGTTTCTGCCTCCACCAGCTTCTCTTCCAGGTTCATTACAGCCTCTCTTTCTGCCAGCTTGTCTTCAATGTCATTAAAAACATCTGCACGCAAAATATCCCTTGCCTTAGGTCTACAGTCTGGATTTGGGTGCAGCAATAACAAACAGAAGGCTGCTTCTTTAGGATAATCCAACAGAAATGACGGTGGAAGCAGGCGATGTCGGAGGTCAGACATGGCCCTTGACCACTCTCCTTGAGATGCAGAGCTATAGAATAGCTCCAAGAAAAGCACTCCCAAGCTGTACACATCTGAAGGGAAGTCTGATGTCCCACCATTAATCTCCTCTGGACTTTTAAACCACATCAATTCCAACGAAAGTATCCCTTTCAAAGGAAACTTTTGGTCAATACGTTGATCTCCTGTATGGTCACTGGAAGAAGAGGTAAAGGAGCTGCCCCCTTGCTCTTCTGCACTATGTTTATCTTTGCCAAGTCGAGATTCTTGTCTTGCATCATCAGCACTCCCATGCCTAGAACTCATGCTTTCTTCATCAAACTGTATGTCCCTCGGTTCACAAACAGGGATGGCTGCAGCAAGCTCTTCTTGCGTCTCATAATCCATTTGCTGGGCAGACGCTGAATCATTTACAGTAGGCCTATCCATACTTGGGCCAATTAGTTTACCATCCATACAACCTGTAGCAGTAACTCTGCCAGTAGACTCTGCTGATGAGCATGAGTTTGTTCCCGAACTGGCTGACTCTATAATAGAAACACGATTCATTGACGAAAGAAGAAACGAAGAGGGACGGACATTCCTCAAGACAACTCCTTGAGAGTGAGCAAGCTCCACAAACTCTGCAATCTGTTTGAAAATGTGTATATTTTCAACTTTGTCTATGACTCTGGCTGGGGTACTAAGCCAGGCCCTGAGAGAAACACTAGAGTCATGATCTGCACGCCCACTGCTACTAGTATCTGATAAGCTCTCGATATCGGGGGTTCGCAAGTAATATGCATTTTGCTTCATATCTTCAAAATCGTGTTGACCCTCCATACCAGTGGCTGAAAGATTTCCAACTAAACTCCTTTGAGTGTAAAAATCCACGTTTACATCTGGTAAAACCAACTTTTTCTCACTTAGAACTGCAGGCTTATTTCCGAGTTGTTCCTCCTCAGGGTCCATTGAATGATCGGTAATCTTGTTATGTGAATACAGGTTCAAATTGCCAGAGTCCTTAGTCGTAGGGCTCTCTTCTTTGCCTGCTGATTGTGATGGTGCATTGGACAGTTCCAAATCTGATGCAGCTGGTTCCTTCTCACATGCTTCCGAGTTTTTTTCGCCCAGTAAGTGTGAATCATTCGGTAGGAGCAATTTGTTTTCCTTCATGGAAGTGTCTTTACTATGACTACTTGATCTTCCTTCATCAGCTGCTCCATCTTTGAAGCTGCAACTGGGAGGAGCATTTGAATTCACCATTGTGCGAGTCACAGGGTCTATAACACCTGCTATGATAATAAAATGAAGGTCCTGGACATACTTATTGATGGTTGCCCTTAAACCTGCTAACAATCCAGTATCAGAAACAGAAACTTTGTTGCTGTAATCAATCTTTTGATTACTAATTAACAGATTATTTGGCAGAAAATCATTGATGACATGACAAGGTAGAGGCCAAACAACTGTCATTCTGGATATCTGGAGTTTGGCTCCAGGAATGAAAACTGCATTATGAATCTTGTAAAGAACTCTCTAGTCTCTCACGCACTTTGCTGACCACGCTTCCTCGGCGTGCTTTCTATTGAGCCTGACTTAAGGGGAGCATGATTGCTCAGTAAACTGTACAAGAAGGTGGAAATGAATATCCATCAAATCTCAGCCATCACAAATAACACTGCGAGTGCGCCATATGATCGCTGTGAACTCGCAATCTTGAGAACCATGCGAACTGCACGATCATAATAAACTGATCAGAAGAGCTGAACGATCATCACAGCAAAAATGCAACGAGAGTAAAGTGTTGTCGAAGCAAGCGGCCATGGCAGCTAGAAGTTGCGCTGCCAGGAAGAGTTAAGTGCTGTCCAAGATCATAAGAAGTAAAGAGCGAAGGCGAAGCGAATGATCGGAGACAATTGTAGTGCAGGCAGAAAGAATGAAGTGGTGATGGAGGGCACACAGAAAGGGAGCACGAAAGAGAGAGAGAGTAAGGGGGAAGCGATGAAAGGAGATTGGAAGGAAGCCATGGAACCTGCAACGGAAGCTTAAACCCTAGGAAAGGATGGATGGATGGCAAGACAGCCAAATTGGGAGATAGGAAGAAGGGACCCGTATTGGCAGAAGCTCTCAGACCACACAATCGCATTTTTCATTCATCAGACCTTAG

**ROOT PHOTOTROPISM PROTEIN 3 (JK218):**

GAAAAACTCACGCTGGAGTAGATTTTACACAGGAGCGACTTGTACGAGAGGGGGGGCACATCACGCCACTACAGGAGTAGCTGGACCTGTAATAATCAATCAAAAGGGGCTTTTCTTTGAGTGAGCCCCACTACGTGTGGCTAAGTGGGGTAGTTACAGACAGCTGCCGGTTATGCAAGACTTGGCTGAGCGCTGCAGTGCGCTGGTAAACCCGACAGCTGTCGGTCACTTAAATGATGTATGAATATATTATTTTGAAAAATGAAATAGAGCCACCGTGGGCGGGCGGAGATGCGGTAAGAGGCCGGATACCGACAGCTGGCGGTTGCAAAGACTGCCGGTCAGACCAACAGCTGCCTCAAGAGATGGAATTGCGCCACCGGCGCGGCTGTTTTTGAAGATGAACAGGCGGCGCTGCAGGGGCACCGCCGCTATGTGTGCTGCTGTCGTCAATAGAGAAACGGGACGTAGGTTGGTTCTGGTGATGGAAAAGATTCCCGGGCGCTCGCACAAGCTTCTTCAGACCACTACTCCAAGGAAAACCAACGCCGCTCAAACGGCGGGGGGCGGGAGGCGCCGGCGGCAGGGTGGAGGCAGCGGTGCCATTTCCAACGGTGCCATGTTTAGAGGAGCGCTCAAGCTGGTGTTGAAGCGCGGTGTAATCAGCTTGGAGGTCAGCGTATTTGGACTTCATTTTCTCGACGTCCAGCTTGAGGAGGCGGATGTCACGGCGGGCTGTAGACCACGGGGAGGAGGCGGAGGCGGCGTCGTCGGATAGGGCGTCGGTGGGGACATGGCAAATGTCCGGAGGCAGGGGAGAGTCGTAGAAGGCTTGTGGAGGATGAGGCAGAGGGGGCCTGTGCGAGAGTGCGCTGCCCGTGATGGCATTGCGTAGCTTTACCTGCTCGCAGAAGAGGACCTGCACCACCACGCGCAACGGCAGCCTCTCGTTCTGCGCCGCGTGCATGCATGCGTCAAGCGATAGCTTCTGGCAGTCCATCAGACGGCACAGCTTCTTCCTGTCCACTTCCGACGTCGCCGGGTGTGCCTTGAGGTAGGTGTCGATGGCTCGATAGAGGCCATCATCACAGGCACGTGCGTACTCCGGGAGCTCCTCGGCCAAGGCTTGGAACTTGGGCACCGACAAGTTGGCATCTCGTGCCACCTCTGCCAAATAGCTATCTATAAGCTTTGCCACTTTTATCTTGGCATTTTGTGAGCTCATGCCACCGCCACCGCCGCCGCCTCCCACCCTTTGGTGCTCACTCATCCCATTGTAGCTGCACATCATGTTTATGTTCATCAAAGGACTTGCTTGTTCCTGCATTAAGAAATTCTCCACTACCCTCTGCACCAAATCGACATCGAATAACGTCTCATTTGTATGATTGAATGATGGTATTAGTAAATCTGTCAAGCTTGCTTGATCCAGTTGCATTCCCACTCTTCTTTCCAGCTCCTTCCTGCACATTACACTTGCATTCAACATGTTGGCAATCCTTAGCATCCTCAATAGGAAGTTGCAGGTCAAGGAGTCCTTCTGTGAGGGTAGAATTGTTACTATACTCTCTAGCATCAGCCTATTTTTGCTTTGCAAATTTGTGAAGGCATCTTTTTCAGTCGCACTAGTGACGTTAGTTACAACCGGAAGGCCTGAATCAGGTGTCGGAGTGCTTTCGATGCTTGATTCGATGTCTTTGTCGAGTCCCGGGATCCATTTCAATGCATATTGCAAAACAGCAGTGCCCACCAGATCAGATCTCATGCCTTTTGATTTAATAGCAGAGATTACCTTCACAAAATATTGCACACTTAGGCTTGACACATCCTCAACCCACCAATCCTGAGGGACTACTTTCTTTAGATCATTTTTTAAACCATTCCATACAGGGCTATCCCCTACATGGTTATTGGGAATACTAAAATGGCTGTTTATGCCATTCTCTTGTACTCCCCCCCTTCCAGTAAAAGACCAACTAATCCCTCTCGGATCCGTACACGCTTTCCACGCGATCGACTCGGTACATCTTCTTACAATTTGCAAATTCTCAGTCCATGGACATAAACTCTCACAGCTTTGCAGCACTGTGATAGAATCTTTCCATGAAGCTAGCACAACAAAACTTAAGAAGGCCTCTGTCTTCGATATGAGGTTTCCATCCTCTAGATCCTCTGTCATTTCGAGATACTCAGCCGCGCACCTCAATGATGCAATGCAATTGGCTGTTAGATCCACAGCAACCCCATAGCAGAACTTGGATGCCAATTCAAATGCCTCGGGACCTCCTGGCATGTCCGTCAGTCGAATATGATCCTTTTGCGTGTCTCGTGACTCGAAAACTAGCCGATTTAACCGCCCACTTCTTGATAACAAAGGGAACTTGTGGAGATTGAATGTGATATCACCTACTTCCACCACAAGATCACTTTGCAACTCTGTAGCAACAAACCAGGCTTGACCCTTTCTCTCAAAGCCATCAGCTTTAGTGCCCATTTTCATGGGGCTCATCATGTAGCTTCTGTTGTTCATTTGGATTTGGCCATCAGCAGAGCCAAGCAGCCTATTCACATGCCCTTTCAATTGCAATTCACTATCAGAGCTGAGCAGCCTGTTAACATTAACATGGCCCTTGAGGGGATGCTCTGTGTCACTTGAAGAGAGAAACCTGTTTACTTGGCTCTTTGCTTGCATTTCATTATCTGACCCGAAAATCCTGCTCGCTTGACTCCTCGTATACTCTTGCGTTTCGCTATCGGAACTAAGCCTTCTGCTGCTAGCTAATTCCCTCATGCTTGGCATTATGATCACCCCTTTTGCCTTACACAGTTTTGTAGAACTAATTCATCAATTTCTAACCTTAATTTTCAAGGTTCCACACCTTAATTATAATGATCATAAAGAAGAATGAACCTAAGGACTAAAAATCTTCAAGAGCTTGAAGAAGACTAATTTCTTCAAGGGTCTAACCTCAATCATAAGGCTCAACCCTTGATGAGCTTGGAAAGAAATTAAATCTATTCATGTGGGCTTGACAAAGTTTAAATGTTTTGCTGGTTCAGAGACCTATGTTCGAACCCTTAATGAGCTTGACAAAGAGTAAATGATTTGTTAAATCAGAGAGCTCATCTCAACAATAAGGTTTCAAACCTTAGTAGGCTCATATATGTGGATTAAACTGAAACACAGATTTAGAAGTACTCTAATGAGCATTTTCCAAGCCAGAATGCATCCATATTGTAATAGAAAAATATGAAGGGGGGTGTAGATAAAATCTGTAAACAAAAATATTATAACTAGAGGTAAATTCCTTTTTCTGTAAGAGTTATGTGTGAAAAGAGAGTAGGGCTTGATGCAAATTAATCAAGATTAAGGAACTGCTAATTCTGTAATAAGAATGAACTAGGATCTTATGACTAGTAGTGGGTATTTTTCTGTGGGGTATATTGTATGGAAGTAATGAGCCGCATGAAACAAAAATTAGCAGAAATTGAGGTGGGGAGGCAGCCAATCACTTAGCAGCATTACCCACACACACACCCATGACACACACACGGTGCCACACCTCTCT

**SHOOT GRAVITROPISM 1 (SGR1):**

TTCAAACTTCAATATAAAGCAAGATCACACGTTGCTAAGCAATAACAAAAACACAACTGTCGTTGAATCTCTTGCTTGAAAAATTTGTGGGCAAAATTACAAGGAGTCATCTTTTCCAAAATATGAAGCAAGTGCCCACAAAGGAGGTAATGCACTTGTTGCTTGATATCTAGCAAGACAACCACACTTATTGTTGAATCAACACATGAGATGTGGGTTGATCACAACCCCAATTGCTTGGAAATAAAATTCCATCACTCATTTTTCACATTCTGATGACCTTTTCTGCTTTGTAGTGCATGCATGTTGCCCCTTGTTCTTTGCACAAGCTTGATGTAGTTCATGTGGCAGCTACTTGCCAATGATCTTTTTTGCTTGGTAAATGATTCATGATTTTCTTCATCAATTTAACCTTTTGCAGAGCTTGAAATTTCATGATGCCTTCATTGAACCTACAAGGGATGATGCAGTGTCGGTGAAGACCACGCCGATGCTGTATACAAGCTAAGATCTTTCCAGCCTAGCTTGAGAAACCCTGCTTCTTCCAAGAGTGAGTATCCTTGACATGGGAACATATTGAGTAGCAGAGTAGCTTGTGTCGCCGCATTACCTGCTAATGAAACTTGCCGGAATCCAGCAGTGCTCATCACTTCCCTCCAGTTATCAAACTTCACTTCCCCTGTTCTAGCAGGACCTCCAACTGCCACTATGTTCTTCATCTCACTTGAGATCAATTGTTGCTCTACAGTATGCCGTTCCTCGCTGTTCTCCGGGTGGCTCGCCCCCAGTGAGTCAAAGAGAGCTGAATAGTAGTGCAGAGCTTCAACAAAGCGGTTGAGAAACGCGCCTGCATGGCTTAAATCTTGCTCCACTATGGTGAATACTGTTGGCTTCAGCTGCTGAAAGAGCTGCAATGTGCGGCTATCTGACCCGGTCACATCGTATAGGGAATGTTGAAGCCAATGCACAGCTAGTGCATCCCCTTTCCTAACCTTAAGAGATGTCAAGTCAAGGTTTCCAACCCTATCTGCCACACTATGAAACTCAAATGGCATACAAAGAGTACTAGCAAAATCGGATAGCCTCTTACCAGTCGCCTCTAGAGCTTCAAGGGATGTGCCAATGCCTGTGATGCGCACACACGGTGGCCCCTCTGGACGAGATGCAAGGATGTGGAAAAGTGCGGGCCATTGTAGGCCTTGCATTATATCACAATCTATGATATGTACCCTTTGATGCCCTTCGAAAGCTTCAAGAATTGCTTGGTTTGCTGTGAAGTGAGAGAATTTTACATAAGGGCATAAGCCGTTGAAGATTTGGAAAGCTGACGCGATGCTTTGGTTGCAGAACCTTTGCATGGCAGGCAAGGGAGAGCATATGCCCAAGCAGGAGTTGAGGATTCTCGCAGCCATGCCCTCTGTGAAGTATGCAGCCACCCTTTGGATGGGATTGCCATACGGTGATGACATCTCATTCAGCAAAGGGATCAAGGTGTTGGCTATCTCTAGATTGTCAGTTGCAACCGCTTCAGCACATTGCAATAGAAGAGAAAGGAGCTCTAAACCCTCTTCATCCGACGGATTCACATCAACATGTGCAGTATTAGCCACATTGCCCACCTTTGATGACACATCATCCATGTTTTTGTATGCATCCGGAGGTGGCTGTAGCATTATCTCTTGCTGTTGTTCATCCATGCTGTTGTATGCGTTTTGTGGATGCCGTAACATGATGTCCTGTTGTTGGTGGTGGTGATTTGATAAAATCCCTCGCTGTTGTTGATGGTGATCTGGCATAATCTCTTGTTGTTGGTGCTGCAGCGGGGCCATACATGCCCTTGGCCCATCAAAGTCTGTGTAATGCCCTCCTTGATTGCCCCCATTATACACAGGCAGTTCACATATTTGTGAATTGTAGTCAGCCCTGTGATTGTCACACAAGTCTAAAGACAAGCACAAGTCAGCTGCTCTGGAGTCTTTCTTTGCCTTGTAATCAGAGTTCCACATCGCCAATTCCTCATGTGTTCGTTTAGATTCGTGTCCAGTGTCTTTGATCAGATAACTGTTGTGATTGCCATTGAGAGCATTGTGATCTCCATTTTGAAAGTGATTGCGCTCACTCATTGATTGAAGCCTTTGGCATTGCTCGACCTTTGATACTTCTGCACTAAAGACTCGCCGATTCCTACCCTCCAAGTCAAGCTGCAAGGCATCAACCTCCAGGATGTTGTAGTCTCGCCTCCCTTGTGTGCGATGCAAGTGAAGACCATCAATGCGTGGCCTTTTGGATAGGCTGTTGTCGAATTCATCGATGGAGGATGAATTCTCGCATTTTGATTCAAGAAGCTCACGCGGTCTGATAGAGCGAAGGCGGGGGTCAAGGAAATGATCCATTTGGAGTGTGTCAAGGACCTGCTCAACTTGCACATTCCAAGGAGAGAGCACATCGGATAGCATGCTAAGCATGTCTTGCACAGACATTTCGGGCATGGCTTCAGCGAGATCATTCATCATGGTATACACCCATGTCCCCAGTACAGTAGAATCCTCCATAGCACATGCACCTCTTTTGTGAACTGTGAAAATAGCAATTCCTTGCTTTATGTATACATGCCAGATTAGGGACTAAATTGAATGACTAAGGATGCCTGTGTATGTGAGAGGCACTCTCTACAGAAGCAATTCCCAGATTAGTTAAAACACTTGAGAAAGTCATTCCTTTGTGTGCAGGACAAACTGTTAAACTAGAAAAGGCAAGGCTTATGCCAAGAATAGAAAAATTCCAAGACAGCTTGTTGTTATGTATGTACGTTTATCTGCAAGAAGACTCAAATTGTGTATGGGTTGGGCTTTGTATTCAGATATGCAGGTGCTCTGTATGCAATTGAAGGACCAAAAATTGCTAATTTAGAGAGAGAGAGGGAGAGAGAGA

**SHOOT GRAVITROPISM 2 (SGR2):**

AGAGAGAGAGAGAGAGAGGTTTCTCCAGAGAAGGGTTGAAGCGGCAGGCCGTGCTTGATTGTTATTGCAAACACAGATGAAGACAGAGCGTCCATTGACCAATCCAGAGTCTAGGCTCAAGCATGCTCTGAATTCCTTAGAAGGCCGAGAGAAGTACAGAGCCATGACCAACAGCCCATCAGACGGCGACGACGTGCGCTGGTACTTCAGCAAGGCCCCTCTCTCTCTTCAAGCGGTAGCGGCCTCAAGTCCATTGACAGAAGTTGTAGGAAAAAGTGACTTTCTTCGTTTTAGTGTAAATGATTCTCTGTCCCTTGAGGCAGCATTTTTGAAGGAAGAGGAAGAACTGGTGTCAGCCTGGTGGAAGGAGTATGCGGAATGCAGTGAAGGTCCATTAGAGGCTTCAGCTAAAGGGCAGCCGCCGTCAAGTGCAAAAGAGGGCACAGAATCTGAGAATTCTTATTCAAAAGAGGCTAGTGAAACCTTGGAGCAGTTCAACGACATTGAGCTTTATGAAAGTGATGAAGTGATTGGTGTCTCTGTCAAGGGTGGTCTCTATGAGGTTGATTTGTACCGGCGACGGTGCTTTCCTGTCTATTGGACTGGTGATCATAGGCGAGTGCTGCGTGGCCATTGGTTTGCTCAGAAAGGGGGTCTTGACTGGTTGCCTTTACGGGAGGATGTCTCCGAACAGCTTGAGTTTGCTTATCGAAGCCAGATTTGGCATCGACGAACTTTCCAACCCTCTGGGAAATTTGCTGCAAGGGTTAACCTACAAAGCTCCATCGAGGGACTGCATGCCTTGTTTACTGGAGAGGATGATACATGGGAAGCATGGCTTGGCATGGACTCATCTGGGATCTCATCAGTCCTTGGTTTTAAAGGAAATGGCTTTAAATTGCGACGTGGCTTTGCCCCATCTCAATCACCACAGCCAACACAGGAGGAATTGCGCCAACGTAAAGAGGAGAAAATGGATGATTACTGCTCGCAGGTACCTGTTCAACATTTGGTTTTCATGGTGCACGGAATTGGACAAAGACTCGAGAATGCAAATCTTGTAGATGATGTTGGTACATTTCGGATTACAACAACTCTTCTTGCGGAGCAGCACTTGACTACATATCAACGTAACAACCTACGAGTGTTATTCATCCCCTGTCAGTGGCGAAGAGGGTTAAAACTGGGTGGAGAACAAGCTGTTGAAAATTGCACGTTGGAAGGAGTTAGGGCTTTGCGCACAATGCTTAGTGCTACTGTACATGATGTGCTTTACTACATGAGTCCGGTTTACTGTCAGGATATCATTGACTCAGTAAGTGGGCAGCTGAACAGGCTTTATTGTAAATTTATCAAGCGGAATCCAGGCTATGCTGGAAAGGTTTCAATATATGGCCATTCTCTAGGAAGTGTTCTTTCTTTTGACATTCTTTGTCATCAAAACAAACTGACTTCAAGCTTCCCCATAAGCCAGCTACATGAGAACAGTCTCGACATTAGAACCTCAGAATCGCTGAACAGGAATGATGATTTAGTAGAACCTGAGCAGACTAATGAGAAGCCTTTGACAGAAAATGGTCATGCACAAGAGTCCCATAAACATGAGACAAATGATCAAGAAATTGATAATGAGATTGCAAGGCTGAAAGAGAAAATAGCAATGCGTGAGGAAGCGCATAATGCCAATAAGCAGCAGGATGAGATTGGTAACGAGTACACTATGGAGTCCAATAGACGAATAAATCAAACACAAGATGTGCAAGGAGACCTTGAGCAGTCTCCTGAAGGAAACACAATGGACTGTAAAATACTTGGGAACGTGGCTAAGAAAGACCATGAACCGATCAAACAAGAACCTTTTTGTTCTAGTGGGCCATCAACTTCCAATTCATTCAGGCAAACTAATAGCCAGAAAAGTTCTTTAAAGCCTAGATCCAGCTCAAGGAAGAGGTACACTCCCTACATTGAGTACACAAAGCTTGCTTTTGAGGTGGATACTTTCTTTGCCGTCGGGTCTCCAATCGGAGTGTTTTTATCTTTGAGGAATATAAGACTTGGAATTGGCAATGGAAAAGATTACTGGCAAGATGATGGAATTGACGAAGAGATGCCTGCATGCAAGCGGATGTTTAATATTTTTCATCCATTTGATCCAGTTGCTTATAGGATTGAGCCTCTAGTGTGTAAAGAGTATGTGCACAACCAGCCTGTTTTTGTGCCTTACCATAAGGGAGGTAGACGTTTACACATCGGTGTGCAGGAGTTTGGAAAAGAAATGTCTGCTCGCTCCAGGGCTCTTGTTCACTCTATTACTGTCGCCGGAGATCGTGTAGTAAAAGTTTTCTCATCAACAGCAGAACAAGATAACAAGGAGGAAGCGCAAGAACCAAAAGAGAGAACTTACGGATCATTAATGATTGAAAAATTAACTGGAGATCCTGATGGACGGATTGACCACATGCTACAGGATGCAACTTTTGAACATCAGTATCTTCAAGCAATCAGCTCGCACACGGCTTACTGGCAAGATGCTGACACGGCGCTCTTCATAATCAAGCATCTCTACCGTGACATTCCTGAAGAAAACCCCAAGTCCAAGGCTGAAGTAGGTTCAAATACAGAGAGCATGGACGATTCTATTTCCATTGAGCCAACTCTGTTCTTAAACCAAAACAATGTAGCTGGTGAAGTGGCCTTCACCTTTTCTCGCAGCGATAGCTTGACAACCACAAAGAGTGATTTAGCTCAACAAATTTGAATCACATGTGCAATCATAAAACTCTCTGTAAATAATGTAATCAGGTTAAGTATAGCAACTTCCATTTTTATGCAGTGTCAATGTGAAATAATTAATTGTGCAATATGTATAATGATTACATCTTTTTGTCTGGCTCTCCCTGGATAGACATATAAATCAAAATTTGTAGACAGGTCTTATAAATT

**ATP-BINDING CASSETTE B1 (ABCB1)**:

GCACATAAAGATTAACAAAGCATAAGTTAAACAGCAAATCCAAACCCAAGACAAACTCACCAAAGCTTTGTTTCGCATTTTGCTTCTGCTTTGAGTTTGCGGGCGCCATGCATTCTCAAGTTGCCTTGCTCAGAAGTAGTGTTTTAGCAACACCTCCACGAAGAGATAGTCAAAGAGAACAAATACATGAATGAATGAAGACAAGCAAGAGTGTTTCTAAGGACAAGTTGATAGAGATTATAACCTAACTACACTAGAGTCGCACACAGTTTGTCAAAAAGGCTGGTGGTGAATGCACCAGATAGCACATACGTTTTCCATAAAGAGGGATTGTGAAAGCTATGGGCTTTAGTGATTTTTGTTGGTGTACATTTTGCAGATACACTCTTTTTAGTTCACAAGCAGTATTTATAGGCTTAGAGTGCTTGCAAAACGGTACCGGATCTCTGAAATTGCTCTCTGAAATTAAAGGTTGTGGTCGTCACTTCAAGGTCTCGTGTTGGACGTTCACAGGAGAGAGGCAGGCTGTCAAGATCAGCTCCAAGTACCTAGAAGCTTTGCTAAGGCAGGACATTGAGAAATTTGACACTGGTTTGCAAACAGGCGAACTTATTGAAAGCATTTCCAGTGATGCAATACTCATACAAAATGCCATAGGCGAAAAGGTTAGAAGCTTGAAAAGGCTGGTATGTTATATGCAATGTACCCAGGGGGTGGCTGCAGGTGTGGGGCTGGCAGTCCCCCCACCGAGGCCACGGTTTCTGCACCTATGCGATGACTCTTCTGCAACAGCCTGCGGCGATGAGGGATAAGAGAAGGGAATGGGCACTCAGGAAAAATGCCCAACACTTCTTGTGATATGTCTGCTTCAACTCTGAAATCTGATTCACTGTTTTTATGCAATGTGTGCAGATGGGGAATGCCATTCACTTTATGAGTACATTTGTTGCTGGACTTATCATTGGGTTCGCCACTGTATGGCAAATGGCACTTGTAACAATTGCTATAGTGCCTTTGATTGCTGCCGCTGCAGCTCTACAAGCTTACATGTTAACGAACATCTCGGCCAAAAGCCAGGAGGTTTACGGAAAAGCTGGCGCCATTGCAGAGCAGGCCATTGCCCAAGTTCGCACGGTATATGCTTTTGCAGCCGAATCGAAGACAATGCAAACATATAACATGGCCCTCATCAAGGCACAGAAGCTTGGCCACAGGATGGGTTTGATAAATGGGCTCGGTCTAGGTACTATGTTCGCAACAATATATGGAAGTTTTTGTTTACTCATCTGGTTTGGAAGCATTTTGGTTATCCATAAAAAGGCCAAGGGTGCAGATGCCCTCATCACCATGTTTGTTATTATCCTTGGCGGCATTGCTCTGGGTCAAGCTTCTCTTAGCTTCTCAGCATTTGCACAAGCACGGGGGTCAGCTAAGATCTTTCAAACAATTGAGCAAACTTCCAACATGGAGGACAGTCAGCCAGATAAATATAAAAACACAAAGCTGCAAGGAGAGATTGAATTTCAAGCCGTGGATTTCTTTTATCCCTCAAGACCAAATGTTCCTGTTTTGAACAAATTTTCCTTGACAGTTTCAAAGGGAAGGAATGTCGCTATAGTCGGAAGTAGTGGCTCGGGCAAAAGCACTCTTATCTCGCTTCTCGAACGGTTTTACAACCCAACAGCAGGTTGTATCTTGCTTGATGGACATGACATAAGAACCTTTCAACTGAAATGGCTAAGGCAGCAATTAGCTCTAGTTAGTCAAGAGCCAGCATTATTTGCAGCCTCCATCAGAGAGAACATACTCTTGGGAAACGAGTACGCTGGAATGGATGAAGTCGAGAGTGCTGCAAAATTTTCTCAAGCACATGATTTCATTGAAAATCTCCCAGCTGGTTATGATACTCAGGTAGGTGACAAGGGTGCACAATTATCAGGGGGCCAGAAGCAACGGATTGCAATAGCAAGAGCAATACTCAAGAATCCGAGTATACTTCTCCTCGACGAGGCCACAAGCGCATTAGACTCTTCATCAGAGAAGCTGGTACAAAATGCTCTTGATCAATTCATGATTGGGAGAACAACCATTGTGATTGCACATCGCCTTTCCACCATTCAACATGTGGACACAATTGTAGTCATGGAAAAAGGGGTCTTGGTAGAGAGCGGTGCACATCAAGAGCTGATGTTGAAAAGACAGAGTGGAGCTTACTACAAACTGGTTAAATTGCAAGAGATTAGTCATGCAACTGCAAATGAAAATGTATTTGTGCGGAGAAATTCTGGTAGATTGACTCCTCGAATGACACCCAAGTCCACTCCTAGGAGAAGACATTCATCGCCTTACATTTCTCCCTATTTGGGAGCATTATCATTGCCTACTGAATTGTCTCAAAGTATGTTGACATATCTAGAACCGCAAGGGAACGATACTAGAAATGAGATCGGAACTATAGAAGTCACAAATGATGCTTACTTCAGAAGGCTTCTCAAGATCAATGCACCAGAATGGCCTCAGGCGATCTTTGCAATATTTGGATGCATCTTGACAGGACTTGTTACTCCCTTTGGAGCTCTGGTATTGGCTGCGGTTGCAAACATTTACTACACTAAAGATCCGCAGGAACAAGTTCGTTTGTATACAATCTATTTTGCCATAATAGCTTCAGTTGGTATGATAGGATATGTCATGCAAGAGTACTTTTGGGGATATGTAAATGAGAACCTAACAAAAAGAATACGAGAACGAATGCTTTCAACAATCTTAAAGAACGAAGTTTCTTGGTTTGACAAAGATGAAAATAGCAGCAGTCACTTAAGTGTTCGTTTGGCCTCGGATGCTGCAACTATTCGCACTACATTTGGTACCAGGCTCTCTTTGCTTTTGAATAATCTTACGGTGTTGACAGTAGCGTGCATCATATGTTTTGTCTTGTCATGGAAGATGAGCCTTGTTACTCTTGCTGTCTTTCCTTTAGGAGTATCAGCAGCTGTTGCAGAGAACATTTTTCTTAAAGGATACACTGGTGACATTGAACTGGCCCATCAAAAGGCAAGCAATATAGCAGGCGAAGCCATTGCGAACATACGAACTATTACTGCATTTTGTGCAGAGGAAAAGATCCTCTCTCTGTTTGTAGCAGAGTTAAAGCGCACTGAGAAAAAAAGCCTTGCTCGAGGACTAGTATCTGGATTCATATTTGGCAGTGTAAAATTTGTGCAATTTTCCTCGTATGCTATCGCGTTATGGTATGGTTTGCTTTTAATGAAGCACAACGAAGTTAGAGGTGGTGATGTATTTAAGGTTTTCTTTGTTCTCTTGGTGGGGGCTACTGTAGTTGGAAATGCAATTGGCGTAGTGCCAGACTTGCTGAAGGGTGGTAATGCTCTTGGACCGGTGTTCACAATTTTGGATCGTGAAACGCAAATTAATCCTGATGACAAGCATGCGCCCTGTGTTGATGGAATGAAGGGAGACATAGAGCTAAAGAAGGTGGATTTCAGCTATCCTTCCAGGTTAGAGGTGCCAGTTTTGAAAGAATTGAGCTTGACAGTGCCTGCTGGGACCAGTTTGGCATTGGTAGGGCCGAGTGGGTGTGGTAAGAGTACTGTGGTTGCACTGATTGAGAGATTCTATGATCCTGTGAGAGGCAAAGTGATGGTAGATGGAGTGGATATAAGGACATATAACTTGAAATCTTTAAGGATGCACATTGGGCTGGTGCAGCAAGAGCCGGCACTGTTTGCCACTACCATTTACGACAACATACTGTATGGAAATGCGGAGGCTACTCACGTGGAGGTGGAGGCAGCAGCAAAGGCAGCCAATGCACATGACTTTATAAGTGCACTTCCAGAGGGGTACAAAACTTTTGTGGGAGAAAGGGGAGTCCAACTTTCAGGTGGACAAAAGCAGAGGGTGGCCATTGCAAGGGCTATGGTAAAGAACCCAACCATTTTGCTGTTAGATGAGGCCACAAGTGCACTGGACACCGAGGCAGAGAAGGTTGTACAGGAAGCTATTGAGCGGTTGATGAAGGGACGCACGGTGGTGGTTGTAGCCCATCGTCTCTCGACTATTCGTAATGTGGATTCCATAGCTGTGTTGCAAGATGGCCTAGTTATTGAGCAGGGCAATCACACTCAACTCCTCTCAAAACCACTTGGTGCATACTCTCACCTAGTTGCGCTACAAAGCCAGGAAGCTTCTTCCTCCTGAAAAAAGACACTGATTCAGGCTGTCCTTACCTAAACTTGTGATGTCACAATTTATTGCATCAATTCTGTTTTTTGTTTTTTTGGCTCTCTGCTTATCTCAATCACATGAGTTTGTGGGCAGAAAGACCAAGCAAGCCAAAGGCTTAGCAGAACTCAGCACTTTTTCTCTCTATTGTGCCCTAACAGTATATAGTTAGTGTTTCCTTCCTATAGTTCCAAATGGCATTTGAACCTTATTACCCAATCTACTGAATACTAGCCCATCCCAGTGCCTATTTAACATCCTCAAAACATTGACATGAATAGACTTGAAGAGCACTTTGCAAATCTGCATATCTTGGGCTGGCAATGAAAAAAGGGATGACTTTTCCTTCCTTATTAGTCACATCAATCACTTTGCAAATGCCCTTTTGAAGTGGATAAGGGGCCCAAGCAAAGTAATAGACACTCTCTTGCAATGTGGAGCTTGCAATAGATCTTAGCCTTTTT

**CORYMBOSA1 (CRM1)**:

ATTTTCAGTTCAAGATACCAATGTGATGGTGGGTGGGCGTTTTTTTTGGGGGGGGGAGCCATGATTGGACCAGAGCTGAACATTTGTGATGTTAACCATCTTATACAACATTGATCACCCATTATAATACATTTACAAAAACCTCTCATGAGGGAAACCGGCCAAACGTCGTAATGAGTTTGAGATTCAATGGCAGTTTATGTTGCCATCAGTATATATTTCAAGTTTTCATTAACCTTCTCTTGTTTACATCAGCTTCAAAACTCGTTTTTGCACACCACCATGGGAATGTGCAAAGTACCTATACAGAGTACAAATACATAATTCTAGCAACCAGTGGTTTCTTTGGCAGATGCCCACTTCTGCACAGCTTCCATCAATCCCTAGCTGCCTTAGCAACTAGCACCCATATTTCTTGAACCTGAGTCTCTCAACTGTGACCTTAAAGTTTGCGCACTGAGTGATCATGCTCTACTCCATATTCCACCTTTTATATGCTGTACGCCATCTGTGAAGCAACCATGTGGAGCAAGTGCTCGTAACAGACTGCTTTGCTCCAGAAAGTCCTATCTTCAAGGAATGCAAATCTTGCATTCCTTCTCTTTTTGTATAGTACTACCTCATTGTGTCATGCAGTCCACGCCTTGGCAGAGAAATTATTCACCAGCTACCGAAAGCTTCACATTGTCTTCCATTCAAGACTTCGATTCCACAGCAGACAAGTGCCATTGCTGTTGGTGCTTCAGGCTCTTTCTAAGCATTTGAAATGCAATTTAACACTTTTAAGTGTATCATATGCAGGAAAATCATATAAGAAAAAGTTCTCTCATTGTAACCATATAGGCAACTCGTTGCTCAGTTCAGGGGAACTTGAATCACCTTGCCGTGTTAGAGCTGATAGCATCCTTGATAAACTCTTCACAAGAGGAATGACCTGCTAGAGCATCTGCCAAAGCTCCCATGACATCAAAAGCCTCCTGCAAATCTTCAGCACCTTGCATTTCAACTAACCACTCAAGCAATTTAGTTGAAAATCCCAGCATAGATTTTATATCCCGCAATTTTTCTTTCATTGTTGATTCCCAAGGTTCTAGAACTCCATGCAATGACCCACCCTCCCCTTTTGAAGATTCCTCTTGTACAGACTCCAATTTTTGTGATCTTACAGCTCCCAGTTTCAGATAGCGCTGCAGCTGATCAACCAGCCCCGTGTAAATAAGCATTGGCTGCACAACTGTAAAGAGCTGGTCTTTGTCTAGCACACCAACTTGTGAATTGTCCTCCAAAGATCTTGAATCTGGGGAGGAAATCGAGATCTCTGCAGAGGAGCGACTCAAAGACACGGCTGGAGAAATTGGAGATCCTGCTAGCGCCGCACGTCCATGCTTGAACTGCATGTATGCATGATAAACACCTCTCTGCAAAAATAACCGCCGATGCTGCTGCCAATCATCCAAAGACTCCAGAAACAATGATTGGACCATCATGAACTGCACTGTTTCATCAACGGCCCCGAGCCTTTGAGGACTACTTCCACTCGGGGTTTTAGATCCTGATGGAGAAGACTCAGGTGGAGATGGAGCATTTAAGTAGGTAGCTAATGCCTTGGACTGAAGACGCTTCTGGCTTACACCTCCCTGATCAAATAAGTGACAAGCCATTTGCAGCATGAAAGGCAACAATCGAGAATTGCTCTCTTTACCACCTCCCTTGCTATCCGTGCTGAACGATGCTCCTGTTGCAAAACGAGCCAGCATCAGCACTATATCATATGTCAGCAATCGCAAGCGGCTACCATCAGCTCGTCCCAAGGCATTCAAATTGTCCCAATATTGATCGACACAACGTGCAAACTGTGCAAGAGGCACAGAGGGACCTCGCAGTGGAAGCAGGTTGTTGCAGAGAGTTTCATTGTTGCGAAGAGCTGCACCCTCCCATTCCTTTTTGGGATTTTTTAAAGATGCATCAGCTCTTTTTGCTTCTAGATGGCATTGAAAATGGATGACATTGAAGTGGCTCACAGTTGTGTAGACCCACTCAGCACGTGTATTATTATTAGTGCCAATACCCAAGTTTACACGTTTGCTGTAGCAATATGTGCCCAACATGTCAGTAGGACATAAACTATAGCCTTCTCTGCAAACCATGCATGCTAAACCTGCTTCTTCCTCTCGCACATCATCCAGACCTTCTATAGATGGAAGGGAGATAACAATGCGTTCACCTCCTGAAGTCATTTCGCGCCTCATTCCCAACCCCTGAAGGAGTTCCTCTCTCTTCTGCAAAGCTCGCTTCCGCATCTCATCACGAGTTGCATCACGCAACCTCATGACCTCCTCTAGCAAAAATCCGTCACCCTTACGTTCTTTATCAGCTAGTGTATCCATAAGATTCTCTGCTCTTGCCCCTATCTCGCTTTCCCCGGAAACACCTTCGAGAGCATGAAGAAGGGGTAGAACACCTCCCTCGTTTAGACACTTCTGAGTTGCAAAATGCCCACTGGAGAGACCCTTTAATATCAAAAGTATGATAGGGACTGATGGTAACTCCAAGCCCTGGGCCCAATCCATAGTTGCTTTATAATCCCCAGAAGTATCAAATATAGCAAATGCTCTTTTAAGATGAAGTATGCATTCTGTTGTGATTCCTTTCTGCATGATAACCCCTTTGAGCTTTTCCCCTTGGGCATCATGCTTGATGGACTCGGTGACTTTGCAGAAGTTCTCAAGCGCTAACTGGTGCTCTGTTGCTTGTTGTGCAAGCCTTTCATCGCTTGCTTTCTCATTATGTTTTTTCATGAGCTGGTCAAACCCACTCCAGTCATGGAGATAAGGGTCAAAGTGTTTTACAAGAAGCTCCATTGCTGCTTGTTCTCCATAAGTTAGGTAAGGGAGGATGCGAGCAACCATGTCATTATTACGTTGTTGCTTACTTGACTTCTTTGAGCAATGTGAATGAGATAATCGCTCCAAAAACATTTGCACAACATGGGCAGCCTGGCTCCCAGCACCATCTGGAACCTTTGAGGCACTCAAAGCACTCTTTGCTATTCCAATGTCACTCTCATTTGCCTCTGTTACAAGGCACTCAACAATAAGCAATAGATGTTCGGCTGATTCGGTGGCCTCGACAGCAAAAGCCCGTCGTGCAGTATCAATTAAAACACCCAATGCACCCAAGCTCAGTAGAGCCAGCCGATTATCCCGTATCTTGCAGCAATACATGAGCAACTTCAACACTAATGTCAACTCTTCCTGGTTAGATTTTATTTCATCATCAGTCAGAGCCTGAACCATATTCAAAACGACTTCCAGACCATCACAATCTCTCATTACTCCAGCAATTGCGAACTCTACTTCTGGATCCTGCGTCTCCTCTCTATCATCCTCCAGCTCTTTAATCATTGGTTCAGTAGCTTCCCCATCTAATCCCTGTAGCCTGTAGGTGACAGTCATGGGTGGACACTCTCTCCCAGCTGAATTCCCTGCAGCAGAACCCCCTGCACCTGTGCCAGAGCCTGATAGCTGATTTTGAGTTTTTTTCCAAACCTGCTCATATACTTGAGCCACACTAAGATCCAAAGAAATAATGCTGCCGGCAACGAGCAATTCCATCCCATAATCATCCTCGATCAACCCAAGCAGATCCAGTTGATGACAGATTTTGTTCTTCACGTCACGCATAAGAGGGCCAACCTCTAAGCTAGAGTAAGGATTCTTGGTCATTGATCCTCGGATAAATTCTTCTTGAGTATGAGCCTTGTTCAACACCATCATATATGCAGGCTCAGGCTTCATTGGGCATATTATATTACATAGAAGCTCCAGAATATACACTACAGTGCGGTCCTTCTTTAGCCTTGCATGAGTTTGAAGACCAGCTATACAAGCTCGAATAAAGTGTCGCTTGTTCTCAAAGCTCTCCTCGAGTAAACCATCTAACAGATCCCTTAAAAGACGGCTACAGTCACCAATTAGCTTGGTTTTTTGCACCACAAGGCCTCGAACAACAAGAAGAGCCTCTAGAACTTCAGAAAGAAGCCCATCCTTCATAAACCGAACACGAATGTTAGGAACTTGTAAGAATTTGCCTAGTAGCTCAATAAGTTTATGGAGAATGTAACCCTGTGAAATGTCTACATGGAAAGTATGTTCTTGAGCTTCTATTCGACTCACTTCCTCAGTAATAAGTTTGCAGAGAGTTTGAAGAAAACCACGAACTGTCAAGAAAAGCCGAGCATCTTCAGATTCTATCATTTTGTAAAGTAGCTCAAAGTACTCACAAGCACTTTCACCGACTGCTCTTGTCGCAGGCAACAAGGCCATCAACAAATTCAAAAACTTTGACCGCCTTGAAGGACTCTGCAAGCAAAGCACCTCTATAAGCCCACACATTTCAACTCGAATTGCCTGAGAGCAAGCACTCAAGGCAAGTTCGCAAACCCACGAGCTTTCCTCAAATACTGAAAGCTCATTTTTCAGAGATAGCTTGCAAGCCCGTCTCTTCCATCTAAGCATGTACTTCAATGCCAAAAACTCCATGCGCTTTGGGTCTCTTCTTGCTTTTGGCATAGAAGGCCTTGTGCTCTGCGATGCAATGTACTGCCTTCGTACAAAATCCACATATGCAGCACCATCTTTCCATTCTCCATAGCTAACAAGGGGCACATCCTGCCCTCTTCTTTCTTGGTCATTATCCTTTTCAAAAGAACCTAATTGGGCCTTAGGAGCAGGTGAACTGTTTGCCCTGTGCATTGTGGAAGCTGCAGTATCAGTCTCATCAGTTCTGCCCTCTGACTTCTGTCCAGATGACGAATTTTCGCTTTCAGTAGGAGCCTCTGCTTTGGGAGGTGTGCAAGCCTGTGATATGATTCTTAAGCAAGGAAGGATGATATGCTCTGAGATTACTGGATGCCGAGCCCCAACCTGAATAGAGGTGAAAAGCAACTGAAAAACTACACGTAGCCGGGACTCCCAGAATTCATCAACCAAAGCACAGGTCTCAGAGAGAAGGTTCAACTCCTCCCGCACGCAAGTTGCAATGTCAATAGATCGATGATGATCCAGACAGTAGATAGCTTTTTGCTTTATCAAGTTATTAAGCTGCCCCACTGCTGCAACATCTCCCTCTGTGAAGGCACACAGAACAGACCTAGCCTGTATACGGGCCATCTTAGAACCCTGATGGATGTTATTCTCAAAAAGTTCAGACAAAATACCTGCTGATACTAGCTGCTGTCTACACTGTGGTATCTTTGCAAGGACTTGCAGCCACTCCAGACAATACGCAACAAAAGTGTTTGCACAGCCATAACAACGGTTGCACGGTCTTGAAGGTGCCAGCTTAGATTCTGTCTGCCCTGTACTAGGAGACCTTTTCTGTTCTAGGTAAGATGTCAAGACACGTCTAAGCCCTTGCAAATTCTGGACACTTCTACTGACTGAATCAAATGCCATTTTACATTTCTCCCCGTACAAAACCCCCAAAATAGCTATTTTCCTGTTTATTTTGAATGATGATGATCCTGGTAGGGACACAATCATCTGCTGAACGGAATCCTTTTGCTGAGAATCCATCTCTGTCTCACCAATACTAGATACCAGTTTTAACAGAGGTTTCTTGAAACCAAGCAACTGCTGGTACCTCTTATGAGCATTTTCTGATTCCAACTCAATAGCAGCAAGACCTTTCTTCATGTCTTCGTCATTCTCCATGCTCTCAAAATGAAAGCTAGGTTTTGCCATGAAATTGAACTCAAATCGACCATACTTGCTATAGCCACACTCATTACACAGAAAGGAATCCAGGTTTTCATAATTAATGTTTCTACACTGCCTGCATTGATAAGCATTCTCATGACAATTTCCACAGATTCCATGTTTGTCTGTAACATAACGACTGCAGCGAGGACACTGCAATGACTCCAAAGATGATGCTTGAAGGTTCTCATAGAATGAATCCAGCTCAATCATGAAATTGCAGGCAGTAATAGGGATAGAAAACTCAACCTTCAATTCTGTTTGACTGAAAGCAAGATAGCAGCTCTTTGCTCTCTTCCACAATGAACAGTTGTTCTTGAGCTCTGACAAATCAGTGACTGGCTTGTTATTATAATAAAGATTGAGTACTTTCACTGACTTTGAACGGCGGGCATCATGCACATTCATTGTTACACTCTGTATGGTATGACTACCAATGCACTTTACAAGAATGCGATTATCAGTAAATTTAGTCTCTGACTTCAAACTGTCAAGCTTCATTCGAGCATAAGGAACTTCTGGGCAGCTACAGGCCAAACAAGGTTCACTTTCCAGGTAGTATCCATCAAAGTCTACCAGGCTGCTGAGTGTATTGTATATTCTGGAATTAGGATGATTTGCAAGAAGCTCGTTTTGAGAGCAAAGAGTATCAAAAAACACTTTCACTGTATCGCTAGTTAGACAAGACTGAACAACAGCATTTTCTTGACTGCTAAAACCTGACCCTTGGATCCCTTTGCCCAGGATCCACGTGAGAAGATCTGTATACTCACCAATATTCTGCCCGAAAGAAGGGAGAATGGGAACTTTTTCCAGTAACTTCTTAAGCAGCAGTTCTCGAACACTAGATCTTCCATGATGCCAAGCACCAGTTAAGACAGCTTTGGCCTCTGCTCGTACAGATGCAGCATTCCATTCAAGTAAAAAAGCATCAATAAAGTGCAATAAGACATTTCCATCCTCAGCAAGCAGCAGCTCCACTGCAAGCTCCATGTCCATGAAAGTCTTTTCTGCTGAGGAATCGGTAGAATCCTCACTAGAACCTCTTTTCTTTTTCCGGGTATCAGTGCTCTGCTTTGTTCCACTGTCTCCTGTAGCATTGGTTGATTCTGCAGTCTCTCCTTTGTTGGCAGAAGACGCGCCAGATTCCTTCCCAGCGTAAAATGCTAAAGTGAGGAGCTTCAATGACTGTACCACGGACTCCTCCCCAAATGCAAAGACCCCATTCAAGAGGAACTGCAAAATGTCCGGATGCCTTGAGCAATACTTCTGCCAGTTTCGAGGTCTTGCCGCAGCTACCTCCACAATGGCTGACAAGCATTTTACAAGCTTCACACTTCTTTCATACACTAAACGAGAATGCAACCCACCTGCCTTTTGTACAAGTTTAAAGAGTTTTCGAACCTCTCTTGAGAGCTGCCAAGCATCCCTCACATTGTAATAATGCCCCTTTGATCCACAAAAATGTAGCAATAGCCTGCGGGCAAACCTCCTAATTGTCGTAGTGTGGGGATTGTTGATAAATGAACACAGCACTTCCTGCCAGCCCTCTAACTTGAGATCTGGAGTTGCCCATGATTTGCTACTCACTTGATCCTTGTCACTTGTTTTTTCAGGCCGAACCAAATTGTAAGCCAGCCGAAAAGTACTTTCTAATAAGAGCCTGTGAAACTCTCCAAAGAGATCATTCCGGTGAGCTTTGGCATAAGAGTCTGTGAAAAAAGGAGCAAAATTTCCAGCTGCAAACTCTTTTCTAACTGTCAATAAGGCACCACACACCGGACCTGTTGAGCTAACTGGACCTGATTCTAGGCTCCTTGACGAGCTTTTGAAGATCTGCCCTAACTGCTGAATGATACAGAGAAGGTAGTTCATTAGCTGTTGCTGCCTAAGCGCAGCACAAGCTCTGTCAAGCTGGGAGCCTGAATCTCCCTTGATTTGACCCTCATCTGAAGATGGTGAGCCGGTCCCTGATGATGACGGCATCACTACCACTTCCCCAACCCTTGCTGCAGACTTCTGTACCGGATGATCAGAACCAGGCTGGTGCCAATTCCGCAACATCAGGTTAAAAAATGTGAAAACAAGGACAATCACCTCTCCAAAATTTGATCTTGTCTTGATTGACAAGGGTTCACCTAGGTTCATCTCATCCATTAGCAACTTGACAAACATCTCCAAGTTTATGCTCTCTGTTCCTGATGATTCAATAAAAGGACTTGTGCATGCTGAGGCTAATCGATAAAAAAACTGCATGAGGGGTAGTGCTTGAGCGCCAGAAGTTTGTTTCATCCATCCTTTACTCTCATCTACCAAACTTTTGAGAAGCAAAGAATTAAGGGCCCTCTTTGAGGCAGATATTGTTGTGCCTTTGGATTCAACCGATGTGATAAAAATTTCATCCCTGCTCTCGACAACATCTAATGAAGAGCTCATAGGGGCAGAAGCAGATTGCATGCTTAAAGCAGTAGCCACTTGCAAGAGATTTTCATCACTGAGATCCTCGATAGACGCAAAACGGATGTCACCAACCTCTCCACCTACATTGTTGTCAATATCAATCGGAATGGCTGACATGGGATGATCCCTGGAATGAGGGGGTGGAAGCCTGTCTCCATCCATGACTTCATAACAAGCTTCACAGAGATCAAAATCTGGGCAAATGTTGCAGTGCCACCGACGCCCAACAATTGGAACTGTTGAACAGCCATCACAGCAATACTGCACAGATGCATTTCCACTATCCTCTTCCATCATTACCTGGCCATCAGGTCCCCCTGCAGCTGGACCACTTTCTGATGGCAATGACATTGATCTAGAATTATCTCCAGCATCATCGTTAGCCAGAATTGTTTGCTTCGGAAAGGGTACTTGAAGCAACCTGGAGGACATTGCCAAACTGCCTGATGCTCGGACTATCTCATGAGGTGCAAACAGCAATTTCTTGAATATAGCAACAGCAGGGGCCACAGATATTGAAGTTCCTAATCCATTGCTAGCACGAAGCGCTAGACACTCTGCATAACCATAGATAAGATCCACAGAAGGTATTATCAGGTTATTCACTGTCTCTGCATCGGGTTGTTCTGCAGTAAGCAGCTCCCACAGCACCTTCATCAAGTCTTCTATAACTGACGGACCATGGCACTCAAGGAAGCAAGCAAAATTTGACCTCCTATGCAAAGCAATCTTACAAACAGCCCGCATTTGAGCTGTGAACTCCTGGATCACCAGTGGTAATGTAGATCCACCAAGTCCCATTTTTGATGCAAGCAGTGGGCACACACGGGAAACTCCTGTAAGCCGTAGGGTATCCTTGGCCTGATAAAAAGCATCCTTCACAGGATAAAGTGCACGAAGCACACGCCTAGCAGCTGCCTGTAAAAGTGGCTGCCTATCGCTTTCAAATACAAGCTCAAGCAATGGCTTGCACCTTTTGCGACTGACTTCTAAACTTACTCCATCCACTTCTGACATTGGTTTTGTTCTGAAGGACAGGTAATAGCCACAAAGAAGTCTAAAGCAGTCAGCCACAACCTGCTCTTCTGAAGAAGCATTCTTCATCAGCTTCAACTTGCTTCTATTTACCATAGATCCAGACACAGATGCTTGAGTTTCCATGTCTAACACAGCCTCTAGCTTTTCCTTCCATCCAAAATCATCCTTAGGTCTTCCATAGACTTCAAGATTGTCAATCCTTGGTAATGAGAAACCTCCAAAAGTTGGGCCTACAGTAAGTGTGAACTCTTCATCTGCTAATAAAGCTTCTGCATTGGTAAATGGCAAATCATACCAGCAGCGGACCCCTTCTTCCAGCTTGACAGCCCTCTGAAATATGCGTAGTTCTGAAGGAATGTGGCTAGTACTAGAGTTTCCAACATGCACTCGGCAGCCTACCATGACTATGTCTGCATTTGAATTGTAGATAGTAATCTTAAAACCCGTCGGACTTGGCCCCTCCAAAAAGCCGTCATCGGAGGCTAAGCTCTGCTTTGCCCCTTCAGAGTCCAGATTTCGTAAGACATCACCACCCAACCTGATTTCATTTGTGATGCAACTAGTCTTCTCAAAAAAATCCAACGGAAAAACAGGTGCAGGTGAATTGCTTGCCCTGCTACCAAAAATTGCATCACCCAGCTTCTTCACCTGCTCAGCCTCCAAGATTGAAGTTTTTGAAGATTCAACACCAGATTCATTTCCATGTCCATGAGTGAAGACCTGGAGACTTCCATCATCATGCAAAATCAAAAGGTCATGTTTGTCTTTCAGAGGAGATTTGTATGCAGCAATTCCATCCACTCGTGGACATCCCAAGTTCCCTGTGACCCTTAAATACTGGGCTTGTATCTCAGATGACCCAAAAGAAACAGCCAAAGCCGAGTTGGCTCTCTGGCTGTTGAGGGAAACAAATAAACCTCGACCTACATCAAGCAGTTCCCTCCAATGGTGTAGACCTGCTACACTTTCCTTGCCTTCAAGGTCACTCTCAACAAGACTTGAAACTTCTGAAAATGAGGTTGCTTCCATGTCCAAACGGCCTATCAGTGATACTCCTTCCGAAAATGAAAGAAAAAGAAGCCTCAATGCTGGAGAATAATGCAAAGACACTCCTTTGTTAGCTTGATACCTTTCTGGAATCAAAATATTGTCAGTGAGAACAGAAGCCCCTAGATCAGCCCCTGCAGCAATACAATGCCTGTAAAGAATTCCTGAATAAGAGAGAACAACCAGCATGCATTGTCCTTGACCCATCGGTATGAGCGAAGCATCCACAATGTGGTCTTCAAGAATCGTGAAGTAATGAACTGGGCTGATACTATCTTGCGACAGATCATAAATTTTCACAAACTTGCTTGTGACAACCATTAGTTGCACCTGTGATCCAGAAACCCAAACTATACGGCAAATATAAGTATCTTGGAGGGCAAGCTCCACTGCTAGGCGATCAGTCACTTCCCCTCTGGAATTGATTGTGAATATTTGACAGTCTTCATACCCAGCAACAGCCAAGTAGCTTTCATTAACGCTATTGAAGACAAGATGAACAAGCTCAAACCGGACAGAGTTTTTCGACAGTGGCTTTACAGTTGTTTTGTCCACTGTCACAGGTGCACTAGATGGTGGCCCAATTACTTGTTCAACATCAAAAATAGTCACTTTATCCCCTTCACCAGCAGCCAACTTGCCTCTACTATTTGTGGAAAGAACAGATTTAAACAGGGTTCCATTCACAAGGTGAGATTTCAACTCTTTTGCATTAACATATTCAGCTTTAATTTTCATATCAAATGAGCCATTTTTGTAAAACCTTTTCAAATGCATGAAGTCACCCCTCAGACTCAACCTTTTGTCATCACCTAGACCTATTTCATCCTCAATATTAGGCATGGAGCACTTTATCCAGTTAAAATTATTCAGAAGCCTCTCACACAGAGACAGTACCTGCCCTTCCACATCCAACTCCGAAAAAACTGACAAAAGCTGGCTCTCTTCACCTTCATTGACTGACACCTTGCATAATTTCTCATTGTCAACCCATACGTCATCCTCCAGGTCTGAATCACTCTCAGATGGAGGCGGCTCTTGATCTGCAGAAGAGACAATAGAGGAAGGAGAGACATACTTTCGAGGTTTCAGGCAAAGACAACTTTGTCCTCTCACACCACCTGCCCCACAATCACAAAAGAACCTGCTAAGCCGAGAATATACCACCTTGTGCCCACGATGGCAAACTTTAGCACAAACAGAGCAGCATCCTTTTGACACTGTCAAGTCACAAGTGTAGCAAAAGTACCAATGTTGCTCCATGAAATTATTTCCACTGGATGTAAATGTGCAAACCCTTGATGCAAGAGTTTTTTCGCTGTTGCTTCCATCATCCTCTTCATCTTTATCCAGACTAGTTATTTCTCCATCAGAAGTTCCATCATCTTCATCCTCATCCACAGATACAGCATCACAATCAGCACCATGCCCACTTTGCTCCATGATAGTATGAGTGACCTCGAGAGTTTTCTTAGGCACTAAAGGTTTGCCTTTGTTGGCAGATTGAAGCGGACGGACACCAGTATTGAGCAAAGTCCTTCCTTCAGTTACTTTGGCTAAAATCTGTGCACGATGGCTGCCACTTAAATTCAAAAGAGAGATCAGAAATGCAAGAAGTGCTTTCAAACCTTCCATCTGACCATTTTGACCTGGCAATTTCCCCACTAGCTGAATTACAGACCAGAGTACATGTCTTATGTGTGATTCCTTGTTAGCTAACTGCAGCAAGAGGGAAAAATAAGACCTCCGGCCTCCATTATCCGAGGCGTTAAAAGCTTGGTCTAAGCTTAGAAGTAATGCATTTAAAAGATGCGAACGGAGATCTTTTCCATCAAATGTAAACTCGGAAGAAACAAGCCCATTCAGAAAACCTACCAAAACATCTCTTATACCACCAGAAGACACCGGCAAAATTCTCTGCTCCAACCACAAGCAAAGAGCCTTACAATCAGTTTTAATGAGCTTTTCCTGCAAATCAGACCTCAAGCCACCATGTGGCATGTCACCTCCAACCACATTTGTCAAAAACTGAAGCACCTTCACATTGACTGGCTCTTGATAGCACCCATCAAGAACATCCACGAGAAATGACAAGCGGCCCCCACCAGAAGTAAAATACTCTCTGAACACATCACATCTAGCTGGGTCTGAATAGATACTGTTGAAGACCTCAACTACATGAAGAATCAGCTCTTCTCGAATTGCATCACTTATCTCACTATTACATGAGAGTGCCTTCAGTAGTGACTCTTTCTGCTTGATCAGATGTAAATCACAGCGTTGAGCTCGCTCTTCTTTTGAAGGAATTAAAACACAAAGGACCTCTTCACATTTGAAGTGTACCGTTTCAAAAATCTCTTCTAGCTTGATCACCATAAGAACATCTAGGAGGCCTTTGACAAAAGCATTCTGTGCCAGCCCATCACAAGCTTTCAAAACATAACTGATAGTCTCCAGAATGCACTCAATCAGACACATGCAAGACATAACCATGGCACTAGCAGGATGATAGCTGCAGGGGTAGCCATGCAGCAAAGAGGGAAGAAAAATCTTTTCAACAGCGCTTAGATTCCCTGCCACAGCATATTTGGATAGTATTTCTGCATGCCAGGATGTGGGCATTCCACGTCTTAATAAAGTATGCAAAGGAGAGAAGCTAACTCCAAGCTTAGTGAACAACTGATCCTGCTTGGCCCAGCTGAGACCTCCTCTCAAAACTAAATGTATTGAGGAAGAAGTCTGATTATCACAGAGGATATCTACCACACTATCGTGAAAAGACTTTACACTGAAAGAGAGCGAGGACAACAAGGTCCTGGACAATTCTTCTAAAGAGCCACTGCTTATACAGTCGCAAACAAGCTGCTTACAAAACAGTGGTAACTCCTCATGCTCAGGCTCTGCTGAGGTATCAGCAAGATAAACTTCAAAGCCTTTCTGTAGAAAAGAAAGGAGTAAGTTGAGCATTGAACCCTGCCTTACCCTGTCAATCAGTGAAAATTTTGTTGGGCTATCCCTCAAACAAGCAGCTTCATAAGTAGACAGAGCGATCTTAAAAAATTTCCACCAGTCTGCACAATTCCCTTCAGTCCCCTTGTTTTGACTCCGGTTGACGTGGCTAAACAAACTTTGTGTTAGTCGCAAAATTAGATCCACGCCACACATATCGCCATTAAAATGACTATAGTCAGAAGGCATGCACCATTTGGACTGTCTACCAAAGCCGCGAACAATAGTCCACCCTAAGAAGAATGTATATCTCTCCACCAAAAGTTCTTCCACTTGAGTCACTGGTTTGCTCCTCCACCTACCTAAGAAACTCCTGATACTGTTGCAAACGTCCGATATTTCTTGCAGTGACTCAAACAATTCAGTCAAATCTCCATCATCTTGGCAAAATGCATTAATGTCATACAACTGAGCCAGAAAACCCTCACTTAGACTAGTTTGTCCACAGGCCGGACCTGCCTCTTTCAAGCCCTGACCTGCAATAGCTGCCATGGATGCGGGTGAAAATCCAGCTACTGCAACATGTGCACTATTTTCCTCAGTGAAGTGGCTTAAATTAACTTTCAAGTCAATAATAAGCCAACTGGGAAATGCCTTGTAATGCACACACATATATCTGAAGATCATAATCAATCTTGAAAAAAGCAAGGAGCAAGATGACAGAAAATTCCCCTCCAAATTCTGTGAGTTTTCACTGAACTCTTTAACAAGAAATTTAACATCTCTTATTATGGACAAGCGGGCGGAAGCACTACTAGAATGAAAAGCCATCAAAAGGACCTCCAGGAGAAAAATTGAAGGGAGGCTCGCAGTAGCTTCTGTTACAGGAATATCCATCCAATCACACTTCTGTATGCTGCTTGCCAATCTATCTCTAAAGCTTCTCCAAACTGCCAATGTTAGCGAATCATGTTGCAGCCCACCTTGCACTGAAGCATGATGGATTAATACTTTAGCCAAAAGCGCTGAGGTCTTACTTGCCATGGTGCACCCTACATAGCCCTTCTCTAAAATTTTCACTAGGAAATGCTCCTCTAGCTGGAAGGACAGATTTCCATTCATCAAGATTGATTTCATAAGGTCAAAGCTGACCTTGCTGAGCTGTATCATCATATCCGAATTACATTGAGCATCGACCTTCTCATTTGAACTGTGAAAAAATATTTCTAGTAGTAGATCAATGCCCTCTGCATTCATAAGTTCCTGCTCTGCTTGAAAACTACTTGTTTGACAGCCCGGACAAAAGATGCAGTCCCTCAGAAAAATCAGAACATCGATTACCGCCTGCAAAAGTGTGTCATGAAAGATACATGTTCTATTGTCTACATTCGACTGTGCATTAGAATCATTACTCGATGTCTGCCTGTTTAATTCTGAAAACAAGTTCCCAAATAAGGCCTCGTCTTCACAGATTGCAAATGGGACTTCTGATGAGTTTCCAACATTACTCTCTTTCCTGTAAAGTGAAGCCTTCTCCCTTGCAGGACTCGCTTCCGTACCTATAGAATAAAGCTGTCCGATAGCTTTCAACAGATCACCCAGCCAAGATCCATCTTCTGGAAAAACTATTTGCATCATCTCACTCTGACTTCTCTCAGCTGTAAGATCAAACTCCTGTGATCTGAGTTTCAGTGCAGGAGCCTGGAAGCAGATGCGAGCACGTCCGTGTTTTCTTATATGAATAACCGCATTGCCGTGAGCATGCATGGTCCTTATGAGACCAGAAATGTAAGATAAAACTTGAGGTTCAAGTACAGCAGGTAGTTTTTGTTGTGAGTAGCCTTTGCCTCCCTTTGATCTGACTGAGACTAGTACATCAAGGGCTGAAGCAGCCATAAATAATCTAAGGTTCTGGGTTACACAATTGCCAAAACCCACAGCCTCAGCAAGAGCAACGACACTGCTGATGAATGTTGCATAGCAACAGTTTTGATTATCTGTCTCTTCCAAGTTCAATGAAATCTCCTGCATGAGCAGCTGTGGAGTCCAAGCCACCAACTCAATTAAGCGATTAAGCAAATCCTCTGAATGAGGAAACCCTGAGGCAAACTTTGATAAGGAACACAAAAGCCTTAGCATTCTGGAGATTACAGACAATTGACCCCGCAATTGATTCTCTCTCTCTTCTCCAAAGTTGAAATTAAAATTTGCTAGCAGACCTCCACACACACCCAGAAGCTCCCTGTGGATGGCAGCATGACTAACTGCAAAATGCTGTGAAGCAGATAGCATGTTTGAAAGCTCTGGAAGATATTTTTCCACATCTGGTATTACTAGCTTCAGAGGCTGATCACTGGTTAAGAGAAGCATTATATCTTCCGAAATCTCTGTTTCCTGCTTACCATTAGTGAAGCTGTCCAGACTGCTGCTTTGAGTAAGCTCTTGAAAGGAAACATTAACAGTATCATCTGTTTCCAACCCCTCTAGCAGCTCTTTTGAAAAACTCAAAACTGGTGCACAACCTATTGTTCTTCCTTCATCAACTGCTAGCTCAATAAGATGGAGCAGTTTGTTGTAAATCAGCTCATCTGAGGAGCTGAAGCTTTCCTTGGCCTTTGCTGAGATAGCAAGATTCTTCAAAGCCTTGAGGCATGCCTCAACGGTGCTCCGGAAAATTCCCTCAACAGGCTCTGTTAAGCCAGGTTCTAGTGAGAGATGCTGGTAATGGAAGATCACAGCTTCCGTGAGGTTGAGGATGAAATCAAACTGCGAGGGGCTCCAGCGACGAAAATGGTGCTGAGAAGCTGTGCTTGTAGAGTCATTAGATTCTGCAGGTTCATGGGTCTGTCCTTCATGTCTTCTTTCCTCCTCCTCTTCTTCATTTCCTTCCTTGTGCTGCTGCAGGTGCGATGGCAGACAAGCGAGGAGAGAGGCAAGGCAACGAAGTGCCTTATTCAGAGAAGGGTCGGCGGCACACTTCGGTGAGGGCAATGGAGATGTGAAGCCTCCATGGCTATGGAAGGCGTCTATCAAGGGCTGCAACTCGGCCATCAAGAAGGCGATCTACACAGAGAACGCATATGGGTAGCGAGCTTCGCGAGCAATCAAGCTATGGCGGCAACGAGGGATTAGACAGAGAGGGAGATAGAGAAGAGGGCGCTGTTCGTGCTTGAACCCGCAAGCAACTTTTCTTGCAGAAGAGGGCGG

**AUXIN TRANSPORTER PROTEIN 1 (AUX1)**:

TACAAAGTTAAAACACTTAAGAGATTGCATTTTTGGAGCAAATTTGTACATGGTGGCCTCTTCATGATGGACAACAGATCTTCGTGTGATAATAAATACCTCTGCAGATTATACATTAGGAGACTCTGTATGCAATAGCCATACCTTCAACCTGATGAACCCTGTGGCAGTAAGCCCTCTACTCAAATGGCTTCATGCTTCTCAATAATGGGCATCGCATCGCATGCCTCTGCAAATGATAAATCAAACAATTTATTTGTATGAGCTACAGACAATAAATCCCATGCCTCTACAATCAATGTATACGAGGAGCACTCTCGGGCACGAGTCGATACATCATAAGCCTTATCTGAAACAACACTATCCTTGATGTATATTGTGAGGATGTACTTTGCTTGATGATGGGCACTTGTATGTAATGATGGGCACTTACATGTACTTTGATTGATAATGGGCACTTGTATGTAATGAAGGGCACTTAAATGTATTTTGCTGGCACATGTAATGATGTTACTCTGGTACGGGTGGGCAACAAGCCTCCATAGGTAAAATACATTCTTTGCTTGATGATGGGCACTTATATGTAATGATGTTACTCTGCATATGGATGGATAGGCACCACAAGCCTCCATAGGTGATACAATATAAACCTCGTTGTATGCAATAACTAATCATATTTGACGATGCTACTCTGCTCACAGATGAGCACCACATGCATCCACAATGGATACATTATAAGCCTGTCTAAAATAGCCATACCATCAACATAAGGCTTTTGATGCAATAAGCAACTGTCTATTTGGGTGGCGGTGGTGCAAGTGTGGGAGAGGATGGATGTGGATGAGGGCATTGGTAACACTTGGTGAACAAGCCGAAGGTGTGAATTTGCCGTGCGAAGTTTGTTATGCTGGCCCAGCCCCCCAATCCAAACCCTACAACCAATGTCCACACCACCACAAATGCATTGACAAGGTAGATAGCAGTCCAGCTGCGAAGATAAGTGGGCCCCTTTTCCACTGCATTCTCTCTGGCAAATGCACTTCTGTAGGTTAACATGTGGGCCAATGCTGGTATGATATACACAGTAAATGTAACCAGTAAAGCGCCTACTGCGGAATTGATTGGGCCAAAGAATGGGAAGATGATTGCTAGAAACCATATGGGTAACACCACTGGTAACCGAATCAATGCTCTCAGTAGAATGCTAGCTGTTGTATGTACTCCAAGTACCTTTTCCCATACAAAGTAGAGAGGTGTACACGCGAAACCAAAAGTTATGAATTGGTGAATCAACATCAGCACAATAGCTAAATCACGCGCTCCAGACTTAGGCAGCAATGAGAAAGCATTGGAGTGATCCAAAAGCTGATCTCCAAAAGCCCAATACACAGAAATGGCAGATGGCAGCGTCAATGTCCAGACATACACGGTGGCATATAGATATATGGTCTTAAATTTGCTTGGCTTCCACATAGCATGCATAATCTCCACTGTGACAGCGTGACCACCAAATGTGTAAAGGATATTGGTGGCTCCGGTGAAATAAAGAACTAGCGCACTTGGGCCTGTATGCTTCACATTTGGCACCTGTCCATGTACCAATGATGCAATAGTGAGATACCATGCTGTGTAAGTTGTCATGCCTAGTCCCAGAAACGACCAAACCCTATAATTGTGAAATGATGGAATAAACACCGTGATAGAACAGCATGCCCCGAAGATGTATGTCCAAGTTCGCTTGTCCAAGTGATCATTGATATAGTAGATATTACTGCCGCATGCGATGAGCTGTATGACTGAACCAAAGAGTAAGAATGTGCAGTTGAACCCCAAGCCGATAGCTTTCCAATAAGGCCCAAGTAGACCATCCAAGACTTCAAACCATTGTATAACATGGTTCTTGAAGTTCACATTCTCCTTCTCCTTGCGGGATCTGTACTCCAGGTAAAGCCAGGTTATCAAGTATGCGGTCCAGCTGCCCATGAGACCATAAAATATTTGGAAGACTATGCCTGATGCCATCCCAAGTTGAGAGAAAGAGTAGGGCAGAGTGAGGAGGACTTGAGCCACCTGGTTTGAAGCACAGCTAAACCAAGCATCATAGACAGAGCCACCATGCCAGAAGAGTTTTTTCACTTGGCCTTTGACAGAGTTATCGTTGAGGCCATCTTGCTCTTTGCGGTCGAGCTCCATCTCCGTGTAGTTCACGCCCGACATTGCTCCATCCAGCTCCTTCACAGTGCCCATTCTGCGGAATGGAGTAAGCTCGGCTTTAGCTTATCTATGTAAGCTCGAGACCAACTTAACAGTAATGGAAGCTCGTGCTTATTGTAGAAGAGTTAGATTGGGTCTGTGCTGCAAAACCCAAGAGAGAGTAGTGGAGTGGAAAGCGAGAGAGAGAGAGAGAGA

**ATP-BINDING CASSETTE G40 (ABCG40)**:

CGATGATGGGTAACTTAAATTATATTGATGAAGAGTTTAGATTGGACTCTAATACACATTCAACAAAGGCATAATGATGCCAAAATTCAGCTTTAGGCTTGTAATACATGTAATCAGACACCACAAATTATGTATTTGGCCTCATGGGAGGCTAGGTGCAAGTAGCGCAACATCATAATAATTTAGTGTCTTCATGTTTGGTCAAATAAAATGAATTAGGAAGGGATTACTGTATTTTTTTTACACTATGTCACAAGAACAATGGCTCCACCTAAAAAACAAGCTGCTAACTTTGTGTTCAATAAGATAGATTGGATGAGGATTGCGTTGACAACAACTTGTAGATGGAAAAGTGGGAAAAGAATTTAGTACCGAGCAAGAGCTTGCTTTAAAAGGAAAATTTTGTAATGATAGTGTGTTTCAATTTTGGAAGCTATTTTTGCTATGTAATTTTGGGTCTAGTATGGACAGTGTATTTTGGCCATCTAGAACTCAAATACCTAGTGTTGGTGTCTCCAGCAGTGTGCGGGAAAGGTCTGCCTCAGCGCCGGTTGGAAATACCGATTAGCCGGTTTGAAAAAAATCCGGCAAATTTTCTTTGTATTTCCGGGTTGAATCTTTGGCAAATCCGGGTTGAATCACACTGGTCTCCAGAGAGAGAGATAAATCCTTCTGGATAGCACATACTCATCGTTTTTGGAAGTTGAGCTTCACGATGGCAATGGAGAAGACGAAGGCGAAGAGCAGGACAATGCCGATGTGGGCGCCCACGACCGCACCCAGGAAGTCGTGCTTGATACCAAACGCTTGGTTCAGGAACTGTTTCACGGGCATGCTTCCCGCATCAGGATCCACCACATTGCTCTCCACGTCCCCTAGCTGTGACCCTATCAACCCATACACCGTCCACGCCAGTGGATCGGCCCAGTAGAACCACCTCCACCACACCGGTATGCTCTGCATTTCCCAGCACATCATGTTATGTTGATACACACGCACACGCACAATGTGTATGTGTGCATTCAGTTTAACATTCACACATACATCCATTCTTTTATTGATACAACGTCTATGCATGAAATTCAAGTATTCTCTGTGTGTTCATAACAAGTACTGCTTACACACATGATTTGTGCATATACCCTCAATTTTTAGCAGTTCAGAATGCTTACCACTCGAGGGATCATGAAGCCAGAGAACAGACTCTGGAGGCTTAAGAAGCCTGCTGTGATTATAGTCGAAAAGTACACGTTGGGGGTAATTGCTACAGTCATCAGGCCAAAGTATGTGAAGAAGAGAATGCAGCAGAAGGTGAAGTAAAGGAAGTAGAAAAACTTGGCTGCCGTCCAATGGAAGTTGATCAGTGTGTACACAAGCACCGTGTAAAGCACACTTTGTGCAAGATTATAAGGAATCTCAATACCCACCTGCAAGTGCAATCAATAGTTCAGTATTTCATGGTATGCTATGTTTTATTGCTTGTGATCATCCACAATCAAAGTGCCTTAGCAGCACCACCTGAGCGAGAGCGTATGATAGTGCAGAGTACATCCTTGCTCCCCTCTCACGATAGAAGACAGTTCGTTCGACCGAAGCAACTGGCTGGACAGTGTTTGAGTTTTGGAACCCAAGAAACAATACTGATGCATAGATGGCACCGAAGATGTTGTACAACTGTGTGGTTGTCTCTCTGCGAGGGCCCAAAGTAAGTTATAAAGATTGGGAGCTTTTTATCTATAAGAGCAAGAGAACTATTTGATTGCAAAAAGCCTTACGTTTTTTGACCTAGTCTCCAGAAAACGGTTCCCAAACTAAGTGCAACAGAAATTGTGAAAAAGAATCGCACAACATTGTAGAAAGGATTGCGCCAGTAAGATTTGTGTTGTTTCCAAAGGCATGCCATGCATTGCTCCCAAAAGGTTTGTGGATATTGGCTTGGGAAATAAAGGTCCTTGGAACCAGGAAGGGGATTACTTAGCTCTCCGATCAGTTCTTGATTGCGCTGGTAGAGAGCAGATTCTTTGTAAATCTGTGCAAAGTCAACTTGAAGGCGAGCCTCAGCTGAAGGCGAGCTGATTTCAAGCATCCATGTAGCTGGGTTGTATCCGTCGTTGATGGGTTGAACCCCTGGTATTGCCTCAAAATATTGTACGAGTTTGTTAGAGGAAGTACCAAGAGGACCTGCATAGATCACTCGCCCACCTCGCTTCAACAGTAGTAGCTCATCAAATGATTCAAATATGTCGATACTTGGTTGGTGAATGGTGCAAACCACTGTCCGCCCAGTGTCAACGGTGTTACGCACGGTTCTCATGACTATAGCAGCAGCTCGTGCATCCAACCCTGATGTAGGCTCATCCATAAAAATTATGGAAGGGTTTGCTACCAGCTCAACGGCAATAGTAAGTCTTTTTCGCTGCTCCGTTGAAAGTCCATTGATTCCAGGAAGACCTACTAATGCATCTCCAATAGGGGTGAGCTCAACAAGCTCCATGATCTCAGTAACAAACATCATTTTTTTTTGCTTGTCTATGTCTTGTGGGAGGCGAAGCCATGCCGAATATTCTAGGGACTCCCTTACGGTTACATTGGGGGAGTGAATATCATTTTGTTCACAGTATCCGGCAATTCTCGCAAACGTTTCGTGTTGCTTGGGGTACCCAGAGATGCTTATATCACCCTCAATGTATCCTCCTGTCTTTCTTCCAGCTAATACATCCATAAGAGTTGACTTCCCTGCACCACTCACACCCACTAGAGCTGTAAGAACTCTTGGCCGGAACGCCCCACTAACATCGTGGAGCAATTGAAGCCTGTCTTCGGCAACCCCCTGCTGCTTCATTTCTGCGGGCATGTCAACA

**ATP-BINDING CASSETTE D1 (ABCD1):**

AAGACATGAAGTGATGGTATATTTATTATAGCATGTTGTTTCAAAGAATTGATATACATCAAATTTACAGGTGAAAGGTCTCTACTACAACCTCCACTTTGGAGGGTTACCCTGACAGATCTATTTGCAGCTCTGGAAGCCCATATAAATTGAGGATACATGGAGAAACTATTAGTAAATGTATGGCCTATAGGGTTTCCGTTCTGCATTGCCCATCCGATTACCTGGTGAATGCAATGCCATTTGGTTTGTTGTCAAAACAGTTGGCTGCCCCCCGTTTTGATGCACCAAAGTTTCCAACCACCTTCGAAGATCAATAGCCCCCTCAGTTCTCTTCATGTTTAATGGAGCCAAGTTCCCAGCCACCTTCCCCATCAACCAGTTTCAACTCCATGGAGTGAAAAGGGATTAGTGCAGGACGTTGAGAAATTGTGACAACTGTTATTCCCATACTGCTTGCAAGCTTGTAGAGATGTTCTTCCACATCCACACTTGTTGCACTTCATTGCAGAATGGAAGTCAGAAATCCATGAGCCAAATAACTGCTAATTGCCAGAAAATTTATGCATACAATCACTATTTAATACTTGGTACATTCATCCAAAATACCATACTTGGGCTGATGGAAGAAAAGCCGAGCCATGCCAAGTCGTTGCTGCTCTCCTAAGGAAAGCATATCCTCCCAGTTGACAGTGGTATCCCATCCTCCTTCACGCTCAAGCAAGTAAACAAGTCGAACATCTTGCAAGGTTGTTAGCAGCCTCCTATCAAGTAACTGCAAGGCTTGTTCTCTATCACCTTCTCTGTATGCATGTGAAACCTTTGCAAGGGCCTCATCTAAAGTTAATGGATAGATAATCTGGTCTCTAAGTGTCCCCAATGCTGTGTATGGATGCTGCGGTACGTAAAAAATTTCTCGGCTGAGACCCGTTTTGTGATAATTGTGAACAGAGGTACTAGGCTTTGCAATAGTGCCACTTGCAATTGGCCATAATCCACCAAGCACCCGAAATAAAGAACTTTTTCCAGCACCGTTTGGACCTGTGACCAGTAAGCTTTTGTTGGGTGAAACAACAAAAGTCAATTTCCTTGCAAGTAGCTTCTGTGCAGGTGTTATGATATCCACTTCTGAGAAAGCAATGTTATCACTACTCTCTGAGTTGCTCAAAGCAACGGCATTTTGTGAAGGGGTTTTTGGTTCTTGCTGAGCCGCATGTAGCAGCTCATCTAACTCGGAAATTCTGGCTATGCCACCTGACAGTTCGAGAAATTTTCTGTAAAGTTCTAAGAGATCCCCAAATGCAAGAAAGCTCTGAGAAACAACTGAGGCTAGGAACCTCAACGCATGTGCCAATTCCCCTTGAATGACAGTAGAAGCACGGTCCCCTGAATTATCCATGGCATACAAAAGACTCAAACCCCATGTCACATTGTGCGGAAGCTGTTTGGTAATAAACTCATCAGCAATCCCAAATATCCATCTCCTCTTCAGAAGCTTTTTGGAATGCTCTAGCAGAGCAGTAAACCGTGCTTCAATCATTGCTTGTTCTCGTGGGCCACCCCCAAAGAAGGCTACGGACTCTGCATGTGTATGCAGCCTTGAATGCATAAACCTGAATAATCCTTCAAGTTGCTGTTCTTGACTTGTCAAAGCTCCAAAATTTGGTGTTATTGATCGTAAGAAGCCCAAACCAAGAAACATGTATGCATACAAAATTGCAACCCCTCTCCTTCCTGTTAGTAATTTCATTCTCCAAGTAAACCAGAGAATGTCCACCAAAGGTTTCACCATGCCTGTTACAAGACCCGACAGGTCTCCTCCAAGCTTTTCCACATCCTGGGTTATCCTCTGGTCGGCATCAATCCTTGTTTCTGATAGATGGAAAACTTTGTAAAAAGCATTGTTGCTGAAGTACAGACTAGATAAATGTTCGGTCATTCGTCTGCGCCATCCAAGGGCCAATAGTGCTGCTAAATATCTTAAGGATGGTGCCACAACTGATGATGCTGCACTCTGCAGGACACTTATACCAGTCAGCCGGATAAAAGCTGCCTTATCCTGTTCAAGTACGTACTTAACGCTTGTTCCATTCAAATGTGCAATTCGATCTGAAATCCAGGTCCGAGCAATTACAAGCAATGCAACAGCCATCAATTGCGCACCTTGTTTTCCATGTAAGCTTGGCACCAGTATCTTTGATAATGCAGCCACTCTTATTGGCATACTTTGAGCTTCATTCTTCAACCTTGGAACCAGTGGAATGGGCACAAGAGCATCTTGCGGAGGTGATCTTGCAAGCACCTCTGCAATATAGGAATCTGATGCTCTGGAAGAGTGGTACGGATCATCCTCCTTTGATGTTGAGAAGATTTTCTTCACAGTCAAGGCATCATTCTGGCGGGCAGACATAGCTGCTCGTGTACTTTCTGAATCTTCGGAAGAAGCAGGTCCAGTGCCTTTGTAGTAAACAGTGGAAGAGCCTTCCCCATCCAAGGATAACACAATGTCATGAAAAGCAACCAGCGCTGGCCGGTGAGATATAGTAATACAAGATGTTCCCATAGCCCTCACTTTGGCACAAAATCTCTCCTCCATGTCAGTGGTCACTGCACTTGTGCACTCATCGAGTATAGCAAAGGAAGGCTTGTGATAGAACAACCTCGCCATCCCCAATCTTTGTTGTTCTCCCAGCGATAACTCATCCCCCCAGTTAACTTCCATTTCTAAGGGATAGCGGTCAAGGAGATACTCTAAATTAACATTTCTAAGTAGCTCCCGCATCCCTTCTTCATTAAGCATTTCAGTCTCCTTGTCACCTGTAAGGGGATATATTAATTGATCTCGTAAGGTCCCAACTGCAGTATAGGGCCTCTGTGGAACATAGAAGATAACATTGTTAAGAGAAGATCCTACTCCTGGCTTTGCAATACACCCAGCGACCAAAGGCCACAGGCCCCCAAGAACACGAAACAGAGAGCTCTTCCCACTCCCATTTGGACCAGTAATCAGTAAATTGGATCCAGGTTCTACTTTTAATGTGAGATTTTCCACCAAAGTGTTTCCAGTAGGGGTCACTACCTTCACTCCCTTGAATTCTATATGATTTGCTTCCACGAAATAGCCAGAACTATCCTTATTTGAAACCCCCCGATTGCTGCTTGTAAGCCTGAGCTCCCTTGCAACCAGCATCAGCTCCCTGATGCGATCTGCATAGCCACTGATCCTGTGCAGCTTGCGCGAATTAAAAGCGAAGGTTCCAAGAGCTTGAAACAAGGCTATGATAACACTGGTATGATATCGCATATTGCTCAGCATTTCTGCACGTCCTAACGTCGAATGATCTGGCTTAAGATTTCCAGAAAAGAATGGTTCAATAATCAAAACCACTCCAACTGTAGCCCCCAAGTATTTCAAAGTGAAATCCTGAATCATGCCGAACCACCACTGGGTGTGCAGAACAATACTCATATGTTTCACAAGTGTTTTGAATCTCTGCTTAACAAACGAGGATTCTCTATCTTGTCCTCCGTAGAATGCTATACTTTCAGAATGTGTCTGAAGCCTTGTATGAATTTGCCTGTAGTCTCCTTCTAATTGCTGTTCTCTGGACATCAACTTTCCAAAAGGAGGGGATACGCTTCCAGTAATAAAACCAGCACCAAGCACATAAGCCAAAATCCACAGTGCATACTTGGGGCTGGCATATGAACAAAGGCGCCAAGTGTAAAGTAGTCCATCAAAACATGCTGTCATGTTTTCTTGAATAACTTCACTAAGCTCAGATGAGAACCTAGGTATATCACTAGCAATCCGCTGTTCAGGATTGCTGATTCTCCTATCCACATGTGACATTTTGTAAAACGTCATGTTCTGGAAGTAGTCAGCATGCACTCGATCTGTTAAAATTTTTCGAAACTGCAAGCTTAGCATCCCAGTGATAAACTTGCTGGTGGAGAACAATGCTGACTGAGCTAGGCAAAGGAGAAGATTTTCAGCAATAAGCCGAATAAATAGAGGAGCACGCCTGAGAAAAGCAGCCCTGAATAGAAACCCTTGAACTTTGGCCAACCGATTGCTCAAGGCAGTTCTCAATACCGCTATAGCGGCTAGAGAAAGCAATGTTCTCATCCCCTTGGTTCCCATTTGAGCAATCAATATGCCAGCAAGTGCATTGACTGACTTCATTCCAGTTTTCTTCTGCTTTTGTCCACCTGTAGAGCTGCCCCCACCCACTGATTTCTCTGATGTGATTCCACCCTGAATACCTGAAGATGGTGATCGTGTCTTTTTCCTGTTGGAGGAGGAATGTGCATATGCAAGACCACCTCCAGCCACCAATACAGTGGCTGCGACCACCAAAGCTCTCCTTTTATTGGAGGGCTGCTGGAAGACCTTCAAAATCTGCAGGGATGCCATGATTAACTACAAAAGGTTCACCCAAAAGAAGCTTGCCTCCGATTTACTTTAATCGAGATAAAATGACCATAGCCTGGAAGCTCCTCTTGTGTGATAAAAAGGAGAATGCACAAAAGCAGCCTTCTGTAATGGAGGGAAACCTGTATGTGTCCTCCCTCAATTCATGTAAAGCAAGAGAAATCTCTGCTTTCCAGTCTAAAAATGGCCTTGCAAAAGCCCAGCTTCTGCGTCAAAAATGGAGATGTGCAAGATGAATACCTGCTTCTATGTGAGGGTTACTCCCTCTCTTCTTCTTCCTATCTGTTTGTGTATGTACATGTGTACAAAACCTCTTAGTGCTGCGACGAAATGCTGCGATCCGATCTCAAGGGGTCCTTGCCATAGCCGAGCTTAGCAGGACAAATCTATGGCGGCAACTGCAAGCGAGGACTCCTTGAGCTCCTTTTCTTCTTGTCTTTCTCGTATTGTTCGTTCAAATCCTTAACAATT

**ATP-BINDING CASSETTE C5 (ABCC5):**

GCTGGCATGCAAGAGGATGCAGAAGTTGTTGAAGCCTTGAGAGTGAAGAATAAAGCATGGAAGAATCCAGAATGTGATTGATATGGATGGAAGCATGGAAACGTTCATTATTTTACAAAGCACGTATAGTGCTTGCAGAAAGCAACCTCATCCATGCGAAGCTAGATGATGTTGCATGTAAATTATTTACGCCATCAAGCCTTGCTAAGGGTACTATAGTAAACGTGTTATTAGCCATGAAGATACCTTCAGAATTTGGAATATGCTCCACGTGGACAATGGTTGGGCTCCACAGATTTATGTTAATTGGGCTGAAGTTAATTCGGGCTGTTCTCCAAAACAGTGGTCTGCTTTCAATTTTGGGAAAAAGTGAGTGCTAACTTTTCTCTCTCCAATGCTAATTTTGGGAAAGAGTGCGTACTGCTCTCTCCATGCTTTTGGCTCTCCACACATAGCGCACACAGCAAACGTAAGGACTTGTGTGTGTGGGGGGTACGAACCAGGAGGGTGTTGAAGGTGATGTTGAAGAGGGAGGGGAGAGCCCTAAGAGCCTTGTTTCCCATAGCATTCATGATAAGTCTATGGGTACATGATTTTGAAGCTCAATTATTCTCGCTCGTCATTTTCGTCGTGGCCATGCCCTCCCATAGAAATTACATGTGGAGGAGCAGGTCCTTGTTGCCAGTGATAACCTCTTTTGGGCTCGAACGCTATGCCCTCAAGGGCAGTTGGATATGGCAATCAATATGCTATGTGACTCCTACTTCATCTTACATCTACTTTTCCCTCCTGAAGGCATGCAATAATCGTAAAGCCCTTTCTCAAGCTCAGAAAGTGCATGCCCATCTTGCCCGGCATCATATCTCGTTCTCTGGCTTTCTGGGTGACTACCTGGCAATGACCCTTGCAAGATGTGGTGCTGGCGAAGATGCACTTGAATTGTTTCTTGGCTTGAGTTGCTGCATTGTCTACTATTGGACTGCCATTATCTCTGCTTACACAGAATGTGGACACGAAGAGGAAGTGCTTAGACTGTATGAGCGAATGCTAGATGATGGTGTGGAATCGGGTTCTTATACTTTTGTTTGCCTCTTTAAAGCTTTACTCTGGACCTTGTTTATGTCAAGAAACTTCATTGCTCGTAAATGTAAATTTGCGGAGGAGCTTGATACATCATGTACTTGCTTGTACCAACGCATTAGGAGAGCACACATACTCAAGAAGGAGAGAGAGAGGTGTTGTCACGAACGCCCCTGCATGAACACAAAAAGACAGAGAGAGAGAGTGCTAGTGAGCCATGTTTTTATTATTGAAAATATGTAGGCAGAGAAGACCTTAAGCAACATTTTAGTTTTAAACGTTTTTTGAAAGTGTACATTTTCACACATATGTATAGTGCTTCACAATTCAAATTCTACATCCATGTATTTCAATTCCTCCTGTACCACACTCCTACTCTTACTCTTGCTTGCTAGCCACCTCTAGTTTTAAGGGCGAGCACTGGTTGCACACACTCTCTTAAAATGGGCAATTTTGTCCCTCTCATTGACCGAGTGTCTTTACGTTTCTTCCAGCTCCGCTCGTAAACGCTTTAGCTCTGATGAAACCTAATTCGCCATAGCAGTTGCATTGAAGCGTTCAGCAATTATCAAATCTCATTCACAAATTAACAACTCTACTTTGTTGAATGCAGGCTTTGATCACAAAAAATAAGAGAGAATTTTATTTTGAAGCTGTTTCCCAATGCCACAACCCTGGGCAGCTTACTGCCATCTTACGAATTGCTCAGATTTTTCTCGGTGTCCACAAAATTTGTCATCAGATGGGCATGCATATGTAGTTGATCCATGCTTGCTTTATCTGGTGCTTTGCACAGTTCATCTTGTCTTTCTGCTAGCATTTTATCTCTCCAAGAGTTGGAACAAGATCAAGAAGAACAAGGGTTTGACAGCACATCGGGCTTCTAAGAACACACAGGTTAATACTGTTGGATTTTTGTACAAGGCAAGTCTTGGTTCAAGTGTTTTCCTCTGCGTGTCATACGTCGCAGTTTTGATATGGGGTGGCTTCTCTGCAATAGAGAAGGGATGGCACCGCTTTGTAATAGTAGAAGTAGCTTCATTTCTCCAAGTATGCTCATGGTACACGATGAGTGCTGCCATTGTGAATGCAAAAGAAGAAAGAAAAACCAGCTATCCTCGCATATTACGCGTGTGGTGGATCCTATCTTTCTTCACAGCTTCTCTCACATTCGCCTCAAGGTTTTTGGTTTGGAAAAGACAGATGGCGTTGCAGTTGGATGCATATGTTGATCTGTTTGCTCTTCCACCATGTTTGTTCCTCTGCATTGTTGCTGTCTATGGAAGGAGTGGCATTCGTATGCCTGCTCCAGATGGCGTATTTGAGCCTCTCTTAAATGGTTCCTCCGAAAAATCATCCTTTGTGAAGAAGGTGACCAATTATGCATCAGCTAATCTTTATAGCCTAGCTACATTGTCTTGGCTGGACAGTGTACTTGCTATTGGCTACAAGAAACCCCTCGAACTTGACGACATACCTCAGCTTGCACTTAAAGATCGTGCAGCCTTTGCATCTGAACTTTTTGAAACCTATTGGACCAACTTGAAAGCAAAGAATCTCATTAAGTGCCTTGCTTTCTCCCTATGGAAAGAGACACTGATAAATGCAATTTTTGCTGGGTTATGCACATTTGCTTCCTATGTGGGACCATACATGATTGATGATTTTGTGGAGTATTTGGGTGGGAGAAGACGTTTTCCTCATGAAGGGTACATTTTAACATTTGTGTTCTTTGTGGCAAAGCTTATTGAGACCTTAGCACAAAGACAGTGGTTTTTTGGTGCACAGCAATTAGGCTTGCGGATAACAGCAGCATTAAGTGCTGTTATCTACAAGAAGGGATTGTCTTTGTCAAACCAATCACGCAGAAACCATACAAGTGGAGAAATCGTGAATTATGTCAGTGTGGATGTGAAGAGAGTGAGTGACTTCACTTGGTACCTGCATGAGCTGTGGCTATTGGTTTTACAGATTTCCTTAGCTCTTTTCATCCTTTACAAGAACCTGGGCTTGGCAGCATTTGCCACACTTGGGGCCACTGCATTGGTAATGCTTGCTAATCTTCCACTCTCTACTGTGCTTGAAAAATTTCAAAGCAGGATTATGGAAGCAAAGGACGCACGAATGAGGGCCACATCTGAAGTGTTGAGGAGCATGAGGGTTTTAAAACTACAGGCATGGGAAATGAGATATTTGAAGAAGATACAAGATTTAAGGAACGTGGAAAGTGGGTGGCTTCACAGGTTTATGTATGCTCAAGCAGGTGTAGTTGCTGTATTTTGGTGCACACCAGCAATGGTTGGGGTGATCACGTTTGCAACATGTGTGTTGATTGGGACGCCTCTAACAGCAGGAAGGGTTCTTTCAGCGCTGGCCACGTTCAGGATTCTTCAGGACCCAGTATATGGGATCCCAGATCTTGTTGGAATGCTTGCAGAGACAAAGGTCTCAGTGCAGAGAATTCAGAAGTTTCTAGAAGAGAAAGAGATAGATGAAAGTGCTGTGGCACGCTCGGTGATAGCAAGTGATGAGTTGATGGCGGCAGTCGAGATACACGACGCATGCTTTAGCTGGAACTCTTGCCAAGAAATACCTACTTTAACTGTGCAGCAGCTGCAAATTGCGAAGGGCTCAAAAGTGGCAATATGTGGACAAGTTGGTTCAGGAAAGTCAAGTTTGATATCTTGTATTCTTGGGGAGCTTCCTAAGCTCTCGGGTAGTGTAAAAGTGTCTGGGAGTACAGCATATGTTGCACAGACAGCATGGATTCAAAGTGGCAAAGTTCAAGATAACATCTTGTTTGGAAATCAAATGGATAGGCAGAAGTACGATAATGTATTAAGTGTTTGCTCGCTGACAAAGGATCTAGAGCTTTTGTCACATGGTGATCAAACAGAGATTGGTGAACGCGGTATCAATTTGAGCGGTGGACAAAAGCAAAGAATTCAGTTGGCTCGTGCCCTCTATCAAGATGCAGATATTTACCTCCTTGATGATCCTTTTAGTGCAGTTGATGCACACACAGGATCTCATCTTTTCCAGAAATGCATATTGGAGTTACTTGAAAGAAAGACTGTTGTGTATGTCACACATCAAGTGGAGTTCTTGCCTGCAGCAGATAAGATACTGGTGCTTGAAGACGGGGCCATTGTTCAAGCAGGGGGATTCAATGATCTTATGCAATCTGGGGCGAACTTCTGCAAACTGATTGGGGCACACACTAAGGCTCTTGAAAGTGTTGAAAACACAGAATGTGCAAGGACATTGGAAGAAAACCTTGAAAGAGTAGACAAAGTTCTTGTGGAAGATAAGGTAGAGGTGGAAGGAAATGCCATTAAGCCACAAAATCCTGATAGCTCTAAGGATCCAGAAAATCCTGATAGCTCTAAGGATTCCAAGGCTTTGAAACAACTTGTACAAGAAGAAGAGAGAGAGACTGGTAAAGTTAGTGCAAAGGTCTACTGGTCTTTAGTAACAGCTGTATATGGGGGTGCACTTGTTCCTTTCTTGCTGTTGTCACAGGTTCTGTTCCAGGCTCTACAGATTGCTAGTAATTACTGGATGGCATGGGCCACTCCTGTAAATCTTGATGAGCCTCCAAAAGCTAGTAATAGAGTTCTTCTTAGTGTATACCTTGGGCTGGCAGTTGCAAGTGCCCTTTGCATACTTGCCAGAGCTCTCCTTCTTGCATTTACAGCTCTCGAAACAGCACAGAAATTTTTTGTCAACATGTTACAGTCAGTATTCCATGCACCTATGTCATTTTTTGATGCTACACCAGCAGGCCGTATTCTTAGTCGTGCTTCAACAGATCAAAGTGCACTGGACTTGGAGGTCTGCAATAGATTTGCGGGCCTGGCATTCAACATTATACAATTTATAGGGATAATTGTGGTCATGTCTCAAATTGCATGGCAGATTTTTGCTTTGTTCATTCCAGTAATTGGCTTCTCTTTATGGCTTCAGCGATATTATATTGCGTCTGCTAGAGAACTTGCAAGGCTCGTGGGAATTCGGAGATCACCAATAATTCACCATTTCAGTGAGTCGATATCTGGAGTGGCTACAATCAGAGGATTTGGGCAAGAGTCTCGTTTCAAAGACACGAATTTGGAGCTGATTGACAGTTATTGCAGACCCCTTTTTCACAATTTTGCAGCAATGGAATGGCTGTGCTTACGCTTGAATGTGCTCTCCAATGTTGTTTTTCTACTCGCTTTGCTACTGGTTGTAAGCTTACCTGTTGGCACCTTTGATGCAAGTATTGCTGGGCTAGCTGTGACTTATGGCTTGGGCGTAAGTTTTTTACAGTACATGGTCATATGGAACCTTTGTAACCTGGAGAACAAAATAATATCAGTAGAGCGTGTCCAACAATACTGCGTCATTCCTTCTGAAGCTCCTTTGGTTGTAGAGAATTGTCGGCCCCCCAGCAGCTGGCCATGGGCAGGAACAGTCACCCTTGAGAATTTGCAGGTGAAATATGCTCCCCATCTGCCCATGGTGCTGCGTGGGATCACATGCACTTTCTTGGGTGGGCAGAAAATTGGTGTTGTAGGCCGGACTGGCAGCGGAAAGTCGACTCTCATACAAGTCTTGTTCCGAATAGTAGAACCTGTTGCAGGAAAAGTGCTGATAGATGACCTTGACATTTCTCAAATGGGGTTACACGACCTGCGTTCCAAATTGAGCATTATTCCACAAGATCCTACACTATTCGAAGGGACTATACGCAATAATCTTGATCCATTGGAAGAGTGTACAGATGCAGAGATCTGGAAGGCTCTCGACAAGAGTCAACTTGGGGATGTTATTCGAGCTAAAGATGAGAAACTTGAAGCTCCAGTTAGTGAGAATGGGGAAAATTGGAGTGTTGGGCAGCGTCAGCTTGTGTGCCTTGGACGTGCTCTTCTGAAGCGGACACGCATCTTGGTCCTGGATGAAGCCACTGCTTCCGTGGACACAGCAACTGATGGTGTCATACAAGATACGATACGGGCAGAATTTACAGATTGTACAGTCATTACAATTGCGCATCGTGTTCCAAGTGTCATTGACAGTGACCGTGTTTTGTTTTTGAGTGATGGAAAGGTAGCAGAATATGAGTCCCCAGGACGACTCTTGGCGGACAAATCATCTTTGTTTGCAAGGCTGGTTGCAGAGTATACGCAGAGGTCAAGTAATGTTGCCCTGCGCTGACTTCTACAAATCAGCGGCTTTAGAAAAGATTTTGCTAGTTTGAGGGCTAGTCAAGCAACTTAAACTCGAACGAGCCTCGCTTAGAGAAATTCCATGTATGCAGCATCTCTGATTGCTTCTCGCCACTTTGCCACCAAAATCCTTTACAGCAGGATTGGCACTTTTCTCCACTCAAATTTTGTCAAGAAGCTTGTGTGCATGCTATTGTGCACTATGTAAAGACTGCATGCCAGCAACATCCGAAGCAAATGTAGAGAAAGTTAAAGCTTTTGATGTCAATTTGTTTTATGGAAATATTAAAAAACAACTTGAGGGCAAGACCAAGGTAAGTTACAAAAAAGAGTAGATGAGG

**CYCLIC NUCLEOTIDE GATED CHANNEL 2 (CNGC2)**:

CCGACTCAATGAACTTGCTTATTAAGATATACGAATAAAAGAATTACAAGGGAACTTACTCTAAGTGATCCTGAGGTTTATGTGAGTTAAACATTGCTGCATAGAATCGAAGGCGGTCACACTCCGACTCTGATCGAGCAAGCGGTGAGATTATTGAAGTGGTGCGATGTATGAATGGGAGCTCCATAGCCTCATCTACAGGCACCTTTTGTTTCCTAGCCTGATAGCGCCTCCAAGCAAGCTGGATTGTGACCACAGCCCATGTCCTCCACCCAGAGGAGTAATACCTTGCTGTCCTCTTCAAATTCTCATTTACAAACTTATACCGGAAATGTTCTGTTACAAACTTGAGATCTTGAGCCTCCAGCCCGAAAGCTTCTGTAGAATCCAAAGACTTGAGCGTCGCAAAAGAGGGGGGGAGCCGTTCTACAAATGGCTTTTTTAAACACCACGTCAAAAGCTCATCTCCACAAAAGTTTCCAGGGCCTAGACAGCAAATGCTAGAATTCCCGTTGCTTAAATCATGGCTGCTAAGGAGATGCCCTCGAACAATGAAAAGCATCCGCTGCACCGGATCTCCTTCCCTGATAACCACTTCATCCTTGATGTAAAGATGTGGCTTTAGACGTTCACAAATGTTTTCTAGAACCAGGTTGTCCATGTGCTCAAAAAGAGGCACCTGACGAACAAGATCCAGAGATAAATGTTGCTTGATGTCCCGTCTTAGCCCTTCAGGTAACCCGCGAACAATTTCCACTTCATCCACACCCCGTGTTGCTGCCCAATGGTGGCGTTCATACCTTCTTACTCTACGTCTGAGACGTGATGGAAGTTGCCGCCGACGCATCCAGCATTCAAGATCTCTTAGCTTCAGCTTCATGGCTTCTTTCTTAGCTGTTATTGAATGCAGGAAAACCTGAATATTGCCAATCAGTAATGTGAAAAGAAGCAGACCGCAAGTGATGACTACAATACTGAAGAGCACCTCTAAACTTTCGTTGCTTGGCTGAAGTGCATTACCAAACGAGCTTAGAGTCATCAGACCCCAAAATATGGGATACAGAAGCTTTGTAGACCATGTGTGACCAGTCACCAAAGGTACTGCCCACTGAAATATCCCATAGTTGTAGCTTTGATTGTTTGGACTATCAAAGCAGAGAGAAAGACTTGTCTGATTGTTCCATGAAAAAGCTGCATTATCCAGCAGTCCTGTGCCTTGATATGTGGTCGGGCGGGAACAGCCCAATAATTGTACAATACAACTGCTCCCTTCTTTACACTGATGTATCAGACAAGCCACAACCCGTTGACTTGCTAGCAGATACCAGCAAGCACCAGCCACATGGGCAGCAATGAAGTAAGCTGTTATATTCAAAGCAAAACCCCACCAAACTGTTCCAAAGACAAAGCCTGTTACATGTCTCATGCGCCGGGCAAGAAAGACCAAATGAACAACTTTGGGAATGTATTGAAAAAGGAATGTGGTCAAAAAAACTGTCATAACAGATCCCACATCTCCTCTGCTGATCATGTGTGGTACTACTATCCACAAGATAATCTGAGGTAGCGGCAATACAACAAAGAGGTCGAACCAGAAACCTCCGCCAGGCCGAAGGTAATGTAGAGCTATTTCGCGAGCATCCCATACAAGGCTTCCACGACCAAGAATTAGGGATTCCTTGGAGACATATGCCAGCTTCAACTGCAGCCAAATTTGAAATAGCTGCACTGCATCGCAAGCTGTTCTGAGGAATGTAACTAAAGCAGCAAGTGTACCATCCATATATAAACATGTGGGCTCTTTCCTGAATGCCAGGACATAGAAGAAAAGTGGGTCTATGGCTAGTGAGACTGAACGAGCTAGGAGAAAAGACTGGTTCAACTTTTGTATAACCCTCCGACGAGGGTCTAGTACTGCACCAAAAATCCAATGCCGCTGTTTCTTTTGAATGGGCTTGAGAGGGATGTCAGTGCTTACGGGGTATAATGAAGAGCCCGCATTGGCTTCCAAAAAGAGCGGATGATTTTGCTCACAGCTCACAGAATGAAAGCGTGGAAGCCCCTTCTGAATGCACTCATTACATGGCTCCGTGATGTCTCTCTTTGATGATGATGGCTCGGAACCATCATCACTAGAAAGAGTGCTCCCCTCTATCTTCTTCAAAGCTTTTTTCAACAAGCTGAAGATGGCAGTGAACTTCATGTCTCAGCTTCTTTATCACACAAGCAATTTACAGGTAGTGAACGTATGTTCCAAGTTGAAACTGTAATGCAAAAATCTTTGAACTATGATTCTGTTACCAAAGCTGCTGGAGAAGTAACCCAGACATGCCGAAAAGTACACCGAAAGTTGTGTACCAGATAACCAGTGAGCCAATTGAAACGAAGAGAATGTGGTAGCTCTCTCAGTTGCACCAGCTTGGAACCTCCGCTGGAATTTTAAAGCGTCCCACAGGAACATAGATGCATGGGTTTAATTAACAATCCGCGAATCTCCGACACATGAATTGTGTAAAACCGACACATGAATTGTGTAAAACAAGGGGTTTACGGTGTACTGCTAGTGGGTTGCAAGTGGGTTTTTTGGGTTGGGATTCCTAATGGCGTACACCGACTTTTCTGTGTGTTCTTGAAGACTATGTATCCAGATGAAGAAGCACGGCTTGCAAAATCGCCAATATTCTGCAGAGATTGGTGAAGGGAAATATTTGCACGGTAGGTTTTCGCGATTGCGCACAAGGGGTTTTATTTTGATCTAAGGCGAACAGAAGCGATTTGTTCGTGAGCTTGTCTCACCTGCAGAAGGGTTTCCTGCTCGAGCTTGTCTTCTGCGCGCAGCGAGAGAGAGAGAGAGAG

**DEFENSE, NO DEATH 2 (DND2)**:

TCTCTCTCTCTCTCTCTGCGCGCAGAAGACAAGCTCGAGCAGGAAACCCTTCTGCAGGTGAGACAAGCTCACGAACAAATCGCTTCTATTTGCCTTAGATCAAAATAAAACCCCTTGTGCGCAATCGCGAAAACCTACCGTGCAAATATTTCCCTTCACCAATCTCTGCAGAATATTGGCGATTTTGCAAGCCGTGCTTCTTCATCTGGATACATACTCTTCAAGAACACGCAGAAAAGTCGGTGTACGCCATTAGGAATCCCAACCCAAAAAACCCTCTAGCAGTAAACCGTAAACCCCTTGTTTTACACAATTCATGTGTCGGTTTTACACAATTCATGTGTCGGAGATTCGCGGATTGTTAATTAAACCCATGCATCTATGTTCCTGTGGGACGCTTTAAAATTCCAGCGGAGGTTCCAAGTTGGTGCAACTGAGAGAGCTACCACATTCTCTTCGTTTCAATTGGCTCACTGGTTATCTGGTACACAACTTTCGGTGTACTTTTCGGCATGTCTGGGTTACTTCTCCAGCAGCTTTGGTAACAGAATCATAGTTCAAAGATTTTTGCAATACAGTTTCAACTTGAAACATACGTTCACTACCTGTAAATTGCCTGTGTGATAAAGAAGCTGAGACATGAAGTTCACTGCCATCTTTAGCTTGTTGAAAAAAGCTTTGAAGAAGATAGAGGGGAGCACTCTTTCTAGTGATGATGGTTTGGAGCCATCATCATCAAAGAGAGACATCACGGAGCCATGTAATGAGTGCATTCAGAAGGGGCTTCCACGCTTTCATTCTGTGAGCTATGAGCAAAATCATCCACTCTTTTTGGAAACCAATGCAGGCTCTTCATTATACCTTGTAAGCACTCACATCCCTCTCAAGCCCATTCAGAAGAAACGGCATTGGATTTTTGGTACAGTACTCGACCCTCGTCGGACCCTCGTCGGAGGGTTATACAAAAGTTGAACCAGTCTTTTCTCCTAGCTCGTTCAGTCTCGCTAGCCATAGACCCACTTTTCTTCTATGTACTGGCATTCAGGAAAGAGCACGCATGTTTATATATGGATGGTACACTTGCTACTTTAGTTACATTTCTCAGAACAGCTTGCAATGCAGTGCATCTGTTGCAAATTTGGCAGCAGTTGAAGCTGGCATATGTCTCCAAGGAATCCCTGATTCTTGGTCGTGGAAGCCTTGTATGCGATGCTCGTGAAATAGCTCTACATTACCTTCGGCCTGGCGGAGGTTTCTGGTTCGACCTCTTTGTTGTATTGCCACTACCTGAGGTGCATTGTGCATCACTGAAAATGCATACTATGGTGAGGAAGTGATTGTGCCATAAATGCATTTAGTTGTTGAACAATGCAGTGATAGTGTGCTGTTGTGTACATGATGATGCATCATGTAGGTTTGCCAACATTTTGGTTCTGCAGTATTAGCAGCACTGTAGCATTGGTATGCTATGATTGATTGTGACACGCCGTGCTATGAGATGTATAACACAATGTGCAATATTCAGACATGACCTTGTCAGTGATGCATTAAATTGTAAGTGTTTCATACATCTTGATGCACTATATAGATTTCATATTCTCCATTGTCAACAGTGCCTTAGATTTAATGCACCAGCTCCATTTGCAACATGGGAATGCTTTTTGCTTACTATGTATGTTTTCCGTGACTTTCATAAATTTTTCAAAGCTTGTCTAGCAATGCATAATGATGCGCTTAGTAATTGTTTCTATGTGCTTGTACTGCAGATTATCTTGTGGATAGTAGTACCATACATGATCAGCAGAGGAGATGTGGGATCTGTTATGACAGTTTTTTTGACAACATTCCTTTTTCAATACATTCCCAAAGTTGTTCATTTGGTCTTTCTTGCCCGGCGCATGAGACATGTAACAGGCTTTGTCTTTGGAACAGTTTGGTGGGGTTTTGCTTTGAATATAACAGCTTACTTCATTGCTGCCCATGTGGCTGGTGCTTGCTGGTATCTGCTAGCAAGTCAACGGGTTGTGGCTTGTCTGATACATCAGTGTAAAGAAGGGAGCAGTTGTATTGTACAATTATTGGGCTGTTCCCGCCCGACCACATATCAAGGCACAGGACTGCTGGATAATGCAGCTTTTTCATGGAACAATCAGACAAGTCTTTCTCTCTGCTTTGATAGTCCAAACAATCAAAGCTACAACTATGGGATATTTCAGTGGGCAGTACCTTTGGTGACTGGTCACACATGGTCTACAAAGCTTCTGTATCCCATATTTTGGGGTCTGATGACTCTAAGCTCGTTTGGTAATGCACTTCAGCCAAGCAACGAAAGTTTAGAGGTGCTCTTCAGTATTGTAGTCATCACTTGCGGTCTGCTTCTTTTCACATTACTGATTGGCAATATTCAGGTTTTCCTGCATTCAATAACAGCTAAGAAAGAAGCCATGAAGCTGAAGCTAAGAGATCTTGAATGCTGGATGCGTCGGCGGCAACTTCCATCACGTCTCAGACGTAGAGTAAGAAGGTATGAACGCCACCATTGGGCAGCAACACGGGGTGTGGATGAAGTGGAAATTGTTCGCGGGTTACCTGAAGGGCTAAGACGGGACATCAAGCAACATTTATCTCTGGATCTTGTTCGTCAGGTGCCTCTTTTTGAGCACATGGACAACCTGGTTCTAGAAAACATTTGTGAACGTCTAAAGCCACATCTTTACATCAAGGATGAAGTGGTTATCAGGGAAGGAGATCCGGTGCAGCGGATGCTTTTCATTGTTCGAGGGCATCTCCTTAGCAGCCATGATTTAAGCAACGGGAATTCTAGCATTTGCTGTCTAGGCCCTGGAAACTTTTGTGGAGATGAGCTTTTGACATGGTGTTTAAAAAAGCCATTTGTAGAACGGCTCCCCCCCTCTTTTGCGACGCTCAAGTCTTTGGATTCTACAGAAGCTTTCGGGCTGGAGGCTCAAGATCTCAAGTTTGTAACAGAACATTTCCGGTATAAGTTTGTAAATGAGAATTTGAAGAGGACAGCAAGGTATTACTCCTCTGGGTGGAGGACATGGGCTGTGGTCACAATCCAGCTTGCTTGGAGGCGCTATCAGGCTAGGAAACAAAAGGTGCCTGTAGATGAGGCTATGGAGCTCCCATTCATACATCGCACCACTTCAATAATCTCACCGCTTGCTCGATCAGAGTCGGAGTGTGACCGCCTTCGGTTCTATGCATCAATGTTTAACTCACATAAACCTCAGGATCACTTAGAGTAAGCTTCCTTGTAATTCTTTTATTCATATATCTTAATAAGCAAGTTCATTGAGTCGGTCAAAA

**VOLTAGE DEPENDENT ANION CHANNEL 1 (VDAC1)**:

TTTTAGGCCGGACACGCAAAATGGACGGTTGAGTACGCGAGCCCTCGCTCCTAAATCTAGCAGGGTAGCGGGTTTCTTCGTCTCTACCATAGCCAGCAATATTCCCTGCAACCTCTTCACTTGCGGCAAGTAGATTCCTTATTAAGCAGCTATTTGAAATGGGCAAAGGTCCTGGGCTTTTCACAGAAATCGGCAAGAAGGCAAAAGACCTTTTGTATAAGGATTACAATTACGATCATAAGTTTAGCGTTACAACGCACACTTCAAGTGGTTTGGCATTTACTTCATCTGGAAGCACAAAAGGTGAAGCCTTTGTAGGCGATGTTGCTGGCCAATTCAAGAGAGATAAGTGGACCACTGATTTGAAAGTGGACACGAGCTCAAATTTATCTGCAACGGTCACAGTGGATGAACTTGCACCGGGCTTGAAGACAATTTTGAACTACACATTACCTGACCAGAAGGCAGGGAAGGTGGAAATGCAATACAAGCAGGATTATACAGCTGTAAGCACCAGTGTGGGTCTAACTCCAGCCCCTCTTGCTGAGTTCTCAGCTGTTATCGGGTCTGCAGAGGCCGCTTTAGGTGGTGAGGTAGCTGTTGATACCGCTACACGAGATATCACGAAGTACAATGTAGCATTGAGTTTTGCGAAGCCAGATTTCACAGCTTCACTGTTTGTGGTGGACAAAGGTGACACTTTGAAGTTTTCCTATTTGCATACGTTAAGCCCTCTTACCAACTCTACTGTTGCGGCTGAAATTGCCCACAGCATCTCCAAGAATGAAAATACGTTCGCTGTTGGAGGCTTCTACATGCTGGATCCATTGACGATTGTCAAGGCCCGACTGAATCACAGCGGAAAAGTGGCTGCTCTGATTCAGCATGAGTGGAGACCGAAGTCATTGGTGACCATTTCGGGAGAAGTAGATGCTGGTGCATTAGACAGAAACACTAAGGTGGGGTTGTCATTATCAATGAAACCATGAGCATGGAGTGTTTTACCAATATGCCAGTCCCTTTGAGCTAGTAACAACCTTAGATAGCTGGGCTTAGTTCTACACGATCAGCCATTATGTTTGCAGGTAGTATCTCTCTGTACTAGAGGGTTTGGGCTACTTGATTTAGTCTGTGCTTCTTTGAGTTGAATAATTTTGGCGGCGGGAGGTTGTAATTTCGCTTTTGAGTTGGCCCATGTTTTGTATTTAGAAACCAAATTATGCATTGCTCCAATAAGACAGTAGATGCTTTAGCATTTTATAGATGATTATGTCAGTGGATTGAGCAAATCAGTCCGCCTGCTATCCTGTGGTGACGATTTATGCCAGAGAGCTTATTTGTTGCATCTTTCTTGGTGCAAGTCTTTTATTTTTCTTTGAAACGAAGTGAGGGTAATGTTGTCCAGGCATGAGTGATGAAGCATTTTATACATCCAGTAGAGCTCTTTCATCTACATAGACTGATTCAAAATGCAGAAATTTTTTCATTCCTATCTTGGAGCTAAGTAAATATATGCATTTGCTTCATGGGCTGTTCCTGGGTAAAGCCATTCAGTGTAAAAAGGCTAGGTTTAGTACATTCAATTGGACATGTTCTGTTTTGTGGG

**MECHANOSENSITIVE ION CHANNEL PROTEIN 1 (MSL1):**

ACAAAGGGCACGTGTTCTCAACATATGTTGAGCTGCGATTGGGAGACAAGTGTGGCGGGAGCTCAGCAGGAGTTGAATTTCTAGCAGCAGCAGCAGCAATGGCGAGGGTCAGCCGTCTGCATTCCATCTCTCTTTCTCTTCTCGATTCGCGAGGCGTCCCTCGCCTGCTCGCGTCTTCTGCAGCCTCCGTCCCCTTCAAAGCCCCCGATCTCTCTGCAGCTTTCTCATCATCGCCATGTCTTTATAGCTACGTCGTAACTTCATCGTCATTTCCATCTCCAAGCTTCTCTGGAGCTCGCTTCTTCAGAAGCTCATCTTTGAGAGATTTCGTTGGCGTTTCATGTAGCCCTTGCTCCCATGGACTGACTCGTAGCTTCTTCTCCAGTTCACGAGCAGGCAAGGCACCAGAAGCAAGCAGTGCGCCCCCTCCACCATCACCTGCAGATCCTCCACAAGAAAACGTGCAGGTGTTTCCTAAGGAGTTCACGGATGCTGCCAATAACACAGTGATTGCACTCAAAGAGACAGCCACATCTTCTATGGCTTGGATTCATGATTCCACAGAGAATTTTCTTCTTGTCTTGCGAGAGCGATTTGATCTTCCTGAGTACACATCAGAGACGGTGACACAGCTGCTCTACACCTCCATTGCTGCTGTTTTAGCTTGGCTTGTCATGCCTAGGGTTTTACGCATGTTTCACAGATACTTTGAGGACGGATCACATCTGATATTGCGGCGCTCTGAGAAGATTCCCTATGAAACAAGTATCTGGTCTGCTTTAGAGGACCCTGCCAAGGTTTTCATTTCTGTGATAGCATTTTCTCAACTGAGCTCCCTAATTGCACCAACAACCATTGCATCTCAGTATTTGCATCAGATATGGAAAGGAGCAGCAGCTGTCTCTCTGGTATGGTTTTTACATCGATGGAAGAGCAATGTGCTGTCAAGAATGCTTGCCAATGAGAAGTTGGGAAGTGTTGATAGGGAGCGCATTCATGCTTTAGAGAAGATCTCCTCTATGGGGCTCTTTATCCTTGGTGGAATGGGTGTCGCAGAAACGTGTGGCATAGCTGCTCATTCCGTTCTTACTGTTGGAGGAGTTGGAGGTGTGGCCACGGCTTTTGCTGCCAGAGATATACTAGGGAACATGCTCAGTGGGGTTGCTCTTCAATTCATGAAGCCTTTCTCTATTGGTGATACAATCAAAGCTGGGTCAATTGAGGGGCAAGTGCTTGATATTGGGCTCACTTCCACGCAACTGTTAGATCTTGATAAATTCCCTGTTGCTGTACCGAACTCTTTTTTCTCAAGCCAGGTGATAGTGAACAAATCACGGGCAACATGGCGTGCATTCACTGTAAAAATCCCTGTGCAGCTGAATGATTTCGAAAAAGTACCTCAAATCACGCAGGAGGTAAGAAATATGTTGAAGACTCATCCAAAGGTGACTCTCCAGAACGGCGTGCCACTCTGTTATGCTTCTCGTGTTGCTGGTACATCACTGGAGATTACGTTACTTTGCAATCTGGTATTCAAGGGTAGAGAGGATCTTTTATCAGCTGAGCAAGATATCACCTTGCAGACATTGAGAATTATAAGCAATTCTGGTGCAGCACTAAGTGCTCCTGCTTGAGATGAATTTTCTTGTAGAGAGCTGTACAGATTAGACTGAATCATGTAGCTGAATTATGTGGTTTGTATGCCAAAGACTTCGGCATTTTTTTGTAGCGAGACTTGACCGTCGACAAGTTGCTTGTACAGTAGTCAGCAGGGACGCGTCCTGTACATATGCAGTTCAATAGATTTAAGCATACAATCTTCTTTACACAAGTTTTGTAATGACCAATAAACAAGCTCTATAGCTGGTATCTGGATCCAGCATGCATGTTTTCGATAGTGGAGCGATTTGTTTTTTCAGATGGGATGGGTAAAGAACAAATGTCAACAGAGTTTCAAATTTCATTTAGACACGACTCTTATTTTAAAAA

**NATURAL RESISTANCE ASSOCIATED MACROPHAGE PROTEIN 4 (NRAMP4)**:

TGGACAGTTTAAGTTTTAACTTCCAGGCTCCAGCTAGTGCGCGATGATGCGATCGTGCGCTAAGCTGTTGTAGTAGTACTATAGGGCAGCGGCAGGGCCCCATGTTGATTATCCACAGAGAGAGAGAGAGTGCGTGCAGGGCATAGGCGTGGCACGGGGCCAAAGCGTGGCGAAGAACGCTTTCGTGCTTTTGTGCTTTTGTGGGTATTATGAAGGCATGCAGGGGCGCCCTTCTCCCTTAGTAGCCCACCGCCAATGGCACCCACAACACCCAGAACAGAGGAAGACGTAGACGAGGCTCGGCAGGCTCTTGTGCGCCACTTCCGATCTCACGACGAAGAGCGCGCCTACGAAAAGCAAGATCAAGTCGACGTCCTCTCCCAAGGGGAAATCCCTGCCTTCTCATGGAAGAAGCTCTGGCTCTTCACAGGTCCTGGCTTCCTCATGAGCATCGCTTACTTGGATCCTGGAAACCTCGAAAGCGATCTGCAGTCGGGTGCCATGGCTGGTTACTCACTCCTCTGGCTCCTCATGTGGGCCACTGTCATGGGCCTCATCTTGCAGCTGCTGTCCGCCCGATTGGGAGTCGCTACTGGCTGCGACTTGGCAGAACTTTGCAGAGAAGAATACCCTTCCTGGGCTAGAATTATACTCTGGCTGATGGCGGAAATTGCCATCATTGGAGCTGACATCCAGGAGGTCATTGGTAGTGCCATCGCCTTGCGTATCTTGTCCCTTGGTTACTTACCCTTGTGGGGCGGCGTTCTCATCACGGGCATTGATGGGTTTATATTCCTTTTCTTGGAAAACTATGGAATTAGGAAATTGGAAGCTCTTTTTGGAGTTCTGATCTCTGTCATGGGAATATCGTTCGCTTGGATGTTTGCCCAAGCCCAGCCCAGTGGAAGGGAATTGTTTTTGGGTCTTCTAATTCCAAGGGTGCCATCGGGCACAGTGAGTCAGGCTGTAGGAGTAGTGGGTGCATTGATAATGCCTCACAACATTTTTCTTCACTCGGCTCTAGTGCAATCACGAGATATTGATATTTCTAGTAGCTCCCATGTGAAAGAAGCTCTTCGATACTTCTCAATTGAGTCTGCTTTCGCCCTGCTCGTTTCCTTGATCATAAACGTCTTTGTTGTATCAGTGTTTGCAAAGGCCTTTTATGGGACCGACATAGCAGATTCAATTGGTCTTGCTAATGCGGGAGAGTATCTACAAGAACAATATGGTGGTGGATACTTTCCTATTTTGTACATCTGGGCAGTTGGATTACTAGCTGCTGGGCAAAGTAGTACAATGACTGGAACATATGCAGGCCAGTTTGTGATGGCAGGATTCTTAAATCTGAGGGTGAAGAAGTGGGTGCGGGCAGTTGTGACGCGATCAGTGGCAATCTTGCCAACCGTTATTATTGCACTTATCTTTGATGGATCAGAGCAGCGTTTGGACAATCTAAACGAGTGGCTTAATGTTCTACAGTCGGTCCAGCTTCCGTTTGCATTGATTCCTCTCTTGTGTATTGTCTCTAAAGAGCAAGTTATGGGAATATTCAAAATCGGACCCCTCATTAAGGCAGTGTCTTGGCTAATAGCTGCACTTGTTTTGGTTATCAATGGCTACCTGCTTGTGGAGTTCTGCTCAACTTACATTGTTGCAACCACCACAGTGACATCAATTCTTGGAGTACTAGCAGCTGTGTATGTCATGTTTGTAATTTATCTTGTTATTCAAGCATGGTATGGACATCATATAACTGGACTTTCCAAAAGCAACAGTTTACAGTAGACTGTGAGACAATTTGTACACAAATGATTCAAGAGCCAAGGCAAAATACTGTATAGAGAATTAGGACGGATCTCAAAAGCAAGAGGAAGAAATCAGAGCCCCTTAATTTTGATTTCCTAAATTTTTTATACCTTCATATACGCTAGGTTACAAACTCTGTCAATATGCTGTTAAACTTCTCTTTATTCCTCGAATAATTGTACGCTGGCAAATTGAGATGAGCTACTTGTGTTACCTGTTGATATTCACACCAACCCTTTTTCCAATGAAAAATACATTTTTGTCATGCTAAAAA

**BORON TRANSPORTER 1 (BOR1):**

TAACTCTCTTGTACAGATACAGAGTTTCATAATATGCATACATTTAAATAAATTATCCATCCATGAGTTTGTGAGAGTGTTCTTGCTGATTTGATGAGATGCTTTTTTCAGCATGCTTTTACAATGTAGGATACTTATTCAGAGAAAAGGAGCATACTTAGCTAGGTGCCATGGGGGGCAGGGGGCCCTTGCCTCCCCCACTTGGGGCCGGCAGGTCCCACCCCCATGTTGTTGTAATCTGAGTAGTGGGATATTTGGTGGGAAAATATCGAAAACGTCTTGTAATTGTATACCCAACTAATAGAGCAGTGTTATCGACACCATGTTGCTTTCCTGGGATAAAAGTTTACAACTACGCATCTATCATATCCAGGTTTAATCTTTTCAAAACTTATGCAGTTCACATCAGGATTTGGTCAAGTTGTCAAGGCTTTATTCCTCCTGCGGCTAGAGCTATATAAACATCAGTTGTTGATAGCTATCCCTTGCATTAAAAGACACCTACTATTTTATGCAGAAGCTCGCTCTAGGCGCTCTTAGGTATTGCTAGTTTCTACAATTTACAGTTTTGAATAGTGCCATGGCTGAAGCATCGAGCAAGGAGCCTCAAAGCACCATACAAGGGACTATTACATGAATACTTACATTCGAACTTGGCTCAGTCCTCCAGTATGGAAAGCAATTGCATCCCTTTTGATGGAATCAAGAAAGATATCAAAGGAAGGAGCAAATGCTACAAGCAGGACTGGACAAGTGGTTTTACCTCTGGTTTCAGGATTTTTGCCCCGACTACGTACATATTTTTTGCATCCGCCATTCCTGTCATTTCATTTGGAGAGCAGCTTGACCGGGACACCAATGGAATTTTGACTGCTGTCCAAACATTGGCATCCACTTCATTTTGTGGCATAGTGCATTCCATCATGGGCGGGCAGCCACTATTGGTACTTGGTGTGGCGGAGCCCACGGTCATCATGTATACATTCCTGTATAGCTTTGTCAAGGAAAGACATGATGTCGGCCCCAAACTATTTCTAGCTTGGGCTGGATGGGTTTGTGTGTGGACATCAATATTTCTATTTATACTGGCAATATTGGGTGCTTGCTCCATCATAAACCAGTTCACACGTGTAGCAGGAGAGCTTTTTGGCTTGCTAATAGCCATGCTGTTCATGCAACAGGCTATTGAGGGTGTCGTAGAGGAGTTTCGAATTCCCAAAAATGCTGATCAATCTCTTGAAGAATATGAGCCTGCATGGCGCTTTGCTAATGGCATGTTTGGCATGGTTCTCTCTTTTGGACTGCTTTGGACAGCTCTTAAAAGTAGAAAGGCTAGATCATGGCGGTATGGCACAGGTAGGTTGAGAGGGTTTGTTGCTGATTATGGAGTTCCACTGATGGTCCTTGTTTGGACGGGATTGTCGTATACAACTTTAAGGCATGTTCCAACAGGAATTCCAAGGCGGCTTTTCAGTCCAAACCCATGGTCACATAATGCTTCAAGAGATTGGACTGTCATACGGGAGATGACCCAGGTTCCAGTTCTTTACATTTTTGGTGCTCTTATTCCTGCCACCATGATAGCGGTTCTCTACTATTTTGATCATAGCATTGCGTCACAGCTTGCTCAGCAAAAGGAATTCAATCTGAGAAGGCCCTCAGCATTTCATTATGATTTGTTTTTACTTGGTATCCTGGTATATCCTTGTTTCCATTCAAAGACCTAGAATTGACAATTGACACACAAAAACAGGATTCTCTATGGTTCTTGTGACAGGTTATGGCTTGTGGGCTACTAGGAATACCTCCTTCAAATGGTGTTATACCCCAGTCACCAATGCACACAAAAAGCTTGGCAACTTTGAAACATCAACTGGTTCGTAAGAAGCTGGTGAAGACAGCGAGGAAGAGCATCGACCACAATTCAACATTGGCTGACCTTTATGAGAGCATGAAGGAGGCTTTTAAACAGACAGGGACACCATTACATCTGCAGACACCATCTTTCCGGGCAATGAAGGATTTGAAGGTCAGCACACTTCAGCGGGCTGCAAGCAGTGGTTGCCTTCACACTCCTGTTGATACTAGTGTTTTTGATCCTGATAAGGACATTGATGATCTACTACCTGTAGAGGTAAAAGAACAGCGCCTAAGCAATTTGATGCAGTCCTTGATGGTGGGAGGATGCGTTGGAGCAATGCCCCTATTGAAGAAGATCCCAACCTCTGTTTTGTGGGGATATTTTGCATACATGGCAATAGACAGCTTGTCTGGAAACCAGTTTTGGGAAAGAATACTCCTCCTCTTTACTCCACCCAGTCGTCGATACAAGGTCCTTGAACAATCCCATGCTTCATTTGTAGAGGCAGTGCCCTTCAAAATCATTAGCATTTTCACCATCTTTCAGCTTGTCTACCTGGTTGCCTGTTTTGGCATCACTTGGATCCCTCTAGCTGGAGTATTGTTCCCACTGCTGATCATGATGCTAGTTCCTGTTAGGAGGTATGCCTTGCCCAAGTTTTTCAAATCAGCACACTTGCAAGAACTTGATCGTGCCGAATACGAAGAATCCGCTGCCATGACACATGATATGGCTGTCAGGGAAGCAGATGAAAGTGGACTTGGCAGTGCAATGGGGTTGGATGAGGAGTATGATGAGGGCGAAGTATTTGATGAGATTATTACTAGTGGGAGAGGCGAAATCCGTCACCGTCATAGCTTTGCTTCTTCGCTTAATGGGAATCAACATGATGATCATGAGCTAGTCATGCAATATGTGATCCAACCTCTGCAAGGCCCCCAATCACATGTGACAACTCCATAAACCATTGATTAAACTCTCATATTTACTCTCTCATCAGTACAAGATGATTCTCATACAATGTGTAAGATTAGTTTTGACTTGCTTTGTCAACATATACTATATTGGGTAATTACCAAGGCCTCAATCACATGTCACAACTCCATTGATTAGTACCCAAACACTCATATAAAGTCACCTGCATCAATAGAAGCTATGTGTATTACATAATGATTTTGACATGCAAAAATTGCAGATAGTTGTCTACATATGCCTTGGAATTTGGAATTCTATGTCGTAAGCTTGTATACGCTAATTTGGTCTATCAACGGGACTTATAACTAGCAACAAACATTCTCATATTATTTGTTCCAATAATTAAAGGCGAGTGTAAA

**ATP/PHOSPHATE CARRIER 2 (APC2):**

TGCGCATAGAGCTCACACAGAGTCGATCTGTCTAGTTATCTCTTGTGAGGATACAAAGTTTGCAGATAGCGCAAGCGGTTCAGCTCCAATACAGAGTTTGGGCATGCATATCAAGCCGACTTGATACCTTTCATCAATTTGCGCGGGAGTTTTGGGGAGCTGCGTTGGCATAGAATCTCAAGAGGACTTGATGCATCCAGTCATCCTGTTTGAGAGTTCAAAGTTTCTGGGAATATGCTATACAGAGTTCAAGAGTTGATTTTATCACTCCAGCATACATCGCCCATGCCTGCTTAGGGTTCAAGCAGAGCTGATTTATCCAGGCATTTGGCGTGAGCTTACCAGGCTGCACAGAGTTCAAGCTTCAATCCCTCATGCCATTCTTTGCAATCACACAGAGATTGCGCCCTAAGCAGAGCTGTTGCATATCCAGTTCTTTTGTGTGATCTTATTAAGTGTGAGCATGCATTTCTACACACAGGTTATTTTTTGACCAGTTTATTCATTTAGTTGGTTTTTCCATCAGTCATTTTGTTTCAGCATTGGGGGTGCTGCATACCCGAAATGAAAGAAAGTGGCAGCAATGGTATCAGAAAGAGAGTCTCACTAGCAGAAATCGAATCAACTGGCATCACCAAAACACGGCCCGTCTGTACCCCGGTCCCTAGATCTAGCCCAGTCACTCTCTCTGAGGTTCGGTTGGCCCTTCAAGAGACTCAAGAGGAAAGAGCTCATCGCATTCACAACCTCTTTGATTTTTTTGACACTTCTCAGAAGGGTTACTTGGATCATTCAGACATTGAAGAAGGGTTTAAAGCTCTTGCCATCCCATCTCAATACAAGTATGCTAGGGATTTATTGGATGTCTGTGATGCCAATCATGATGGTCGTGTCGATTTTGATGAGTTCCATAAGTACATGGATGCTAAAGAGCTAGAACTGTACAGTATATTTAGAGAGATTGATGTGCGGCACAATGGATGCATTCTGCCTGAGGAGCTTCGCACGGCATTACAACAAGCAGGTTTTCCAACATCTAGCTCTTTGATTCAATCACGTAGCTACAACATTAATTGGGAATTCAGCTAAGTGATGAGGAGCTTACCCATTTTGTAGAGCGCATTGACAAGGACAACAATGGCATCATAACTTTTGAGGAATGGCGGGATTTTTTACTTCTTTATCCTCATACAGCCACCATGGCAAACATTTATCAGTATTGGGAAAAGGTTTGTTTGGTAGATATAGGAGAGCAGGCTGTGATTCCAGAAGGGATAAGTCGTCATATGCATGCAACCAAATATTTCATAGCTGGTGGATTTGCAGGTGCTTTATCTAGAACTGCAACCGCACCTTTGGATCGCCTGAAGGTGCTTCTGCAGGTGCAAACAAATTCCATGGGAGGAAAGCCCCGTATTGTACCCTGCATACTACAAATTTATAAAGAGGGAGGAATCTTGGGATTTTTCAGGGGAAATGGCCTTAATGTACTAAAAGTAGCTCCTGAATCAGCTATCAAATTCTATGCGTATGAGATGATAAAGGATTTTATGGTAAAGGCAGATGGTCTCCACAGTAAAGAGGAGATCAGTATTGTGGGCAGGCTTCTGGCAGGTGGGACCGCAGGCGCCATAGCTCAGACGGTTATCTATCCGATGGATTTAGTGAAAACACGATTGCAGACTTATGCTTGTATTGATGGAAAGGCTCCTCCCCTGATGAAGTTATCAAAAGACATATGGCATCAAGAGGGCCCTCGAGCATTTTACCGAGGGCTTTTACCATCACTACTTGGGATGATTCCATATGCTGGGATAGATCTTGCAGCTTATGAATCTTTGAAGAATATGTCACGACAATGGATGCCTGAAGATCGTGAGCCAGGGCCGTTAGTACAGTTATGCTGCGGGATGATTTCTGGCGCATTAGGAGCAACTTGCGTCTACCCATTGCAGCTAGTTAGAACAAGGTTGCAAGCACAGCAGATGAACTCAGCTACTAGGTACAAAGGAATGACTGATGTCTTTCAGAGAACTTTCAAAGAGGAAGGTTTCCGTGCCTTCTATAAAGGCCTCCTTCCAAACCTGCTAAAAGTGATTCCAGCTGCTAGCATTACCTACCTTGCATATGAGGAGGCAAGAAAAAGGCTCTCCATTTGATAGTGATTTTGTGGGCGTATTTTCAAATTAGAGCTTGCTACTTATTTATCCTTCGTACGAAGGCATGTGCGATTATGAAACTTACTGCATGGGATGGGAATTCAGTATTGAACAAGCTGCATCACAGTAATAAGAAGGGAGGTTGCCGCTACCATGGTTCACAAGGCTGGCTTCCTCTGAAACACATTTGAGTATCTTCCGCTTTCACTGTGCATGATTATTTTTGTGGTTACATAAACAGCCTAGCTTGTCAAAGGCCATGCAGTTTTCCAAGACAATCCTGCTTAGTCGGTATCCATTCACTGCCTTTATAAAACCCTACCTTCTTGTGCTTAAGGTTGAGGCCAGGGCGTATCCAGGCGTTAATTCTTAAATAAAATCATTACCTCATTACCGTAATCTCAACATATAATAAAGAGCTTCATCCCATTGTTTTCTAAATGTATAGCTCATTGAAATAACATCTGGACAGCAATGTACTTTATATGGTTCACAGTTTGAAAAAGCCTGTGATGGTTTTACAGATTGTGGCTTAAATTTTGTCATGGATGATCATCTTCCATGGTTATTTTAAAAAAA

**CATION/H+ EXCHANGER 17 (CHX17):**

TTCTCCTTCTCCTTCTTCTTCTTGTTGTTGCTCGCGCTCTCTTCTCCTCATCCTCCCTCTCCTCGAAGGCCGCAGCTATGGCGACCTCTGCAGCAGGCAACTCCTCTTGCCCCTCTCCCATGGCTGTTACCTCCAATGGGGTTTGGCAAGGAGATATCCCCACCAATTTTGCCCTTCCTTTGCTCATCTTGCAGATCTCCATCGTCCTCTTGGTCACCCGTGTTCTCGCTTTGCTTCTCAAGCCTCTCAAGCAGCCTCGCGTCATCGCAGAGATCATTGGTGGGGTTTTACTTGGCCCATCAGCATTGGGACGGAGCTCCAGCTTCAGCCAGAGAGTTTTTCCCAAGCAGAGCATAACCGTCTTGGAAACTGTGGCGGATTTGGGGCTTCTATTTTTCTTGTTCCTGGTTGGTTTGGAGCTTGATCTTAATGCCATAAAAAAGACAGGCAAGCAAGCCCTCTTGATTGCGGCTGCTGGAATCAGCCTCCCTTTCGTTATGGGGGTTGGTGTGTCTGTGGTCTTACACCAGACAATTGCAAGCGAAGGGAAGTTTGCACCCTTTCTTGTTTTCATGGGTGTGGCTTTGTCAATAACAGCTTTCCCCGTCCTTGCTAGAATTCTTGCAGAGAGAAGGCTTTTAACAACTGAAGTGGGGCAAATGGCCATGTCAGCCGCAGCTGTAAATGACGTCGTTGCATGGGTTCTACTAGCATTAGCAGTTGCTTTGTCGGGTTCAGGGAAGAGTCCACTCATAGCAGTGTGGGTGCTGCTGTGTGGACTGGCTTACGTAGCATTCATGTTTATGGTTGTGAAGCCCTTGATGGCGAAGGTAGCTATACAGGTTATAGAAAATGAGCCTGTAAATGAGATGTATGTGGCGGTGACATTGGCTGCTGTATTGGTGTCGGGTTTCGTGACGGACTCGATTGGCATTCATGCTATCTTTGGTGCTTTTGTGTTTGGGCTTGTCATCCCCAAACATGGCCCCTTTGCAGGTGCGCTTATCGAGAAGATTGAAGACCTGGTCACCATTTTGATGCTTCCCCTCTACTTTGCTGTCAGTGGTCTCAAAACCAACATTGGATCCATCCATGGTGCCCAGTCTGGTGGTCTTCTTGTCCTTGTCATTATGACAGCATGCTTCGGCAAAATTGTGGGAACTTTTTTGGTGGCATATTTTAGCAAGATGGGTTTCAGGAAGTCCCTCACTTTGGGATTTCTAATGAACACCAAAGGGCTAGTGGAGCTTATCGTCCTCAACATCGGGAAGGATCGAAAGGTCTTAAATGATGAAACATTTGCAATTATGGTTATAATGGCTCTGGTCACCACCTTCATCACTACACCTGTGGTCATGGGGCTCTACAAGCCTGCAAGGAACCCGGTCCCCTACACTCGTCGAACACTTTATGCACCTGACGAGAAGGTATCAGAGTCTGAGCTCCGCCTACTAGCATGTGTCCATGGAATGCCCAATGTGCGTGCTATCATCAACTTGGTGGAGGCTGCACGGGGTACCCGGCGACGACCTCTTCACATGTACATTCTGCACCTGCTGCAGTTGTCAGAGCGGCCGTCCTCAATTATGAAAGTGCAGAGAGTGCGACGAAATGGGCGTCCTTTCTGGGACAATCAACATTTCAATGGCTATGACAACATTGTGGTTGCATTTGAAGCTTTTGGGCAGCTCAGTAAGGTTACCGTGCGGCCCATGACTGCAATTTCTCGCTTCGACGACATGCATGAGGATATTTGTGCCACTGCTGAAGATAAGCATGCATCAATCATCATTCTCCCTTTCCACAAGCATGTCCGCCCTATGAGTGACAGAGGCTGGGAAGGCCACAGCTCGGGACTGCGCCATGTTACTCAGAAAGTTCTACAACATGCCCCTTGCTCTGTTGGCATTCTTGTTGACCGTGGCATCGGGAACACCTCTCGCTCTTCCAGTTCTGTGGACCAGAATGTCGCCGTCCTCTTTTATGGTGGCTCTGATGATCGTGAGGCTCTTGCTATTGGTTACCGCATGGCTGAGCACCCGGGAGTGAGGCTTTCTGTGTTCAGATTTATCTGCCAAGATCCAGCAGAGCATCATGTGGACTTAACAGTGTCACAGCAGCAAACAACCAATGGCACCTCCTTGAAACAAAGCAGTAGTAGGTTTTCTCTTAGCTCCAGGCACTCTTGGGGCCTTGAGATGTTTGAGGTGGCAATTGATGATGTTGATTGGGAGGCAGAGAGCAAGCTGGATGAGAGGTGCTTGGCATTGGCCAAGGGGAAGGTGCCAGAAGGGACTAATAACAACAGCGACAAGGATACAGCAGCTACGTCATCTATTAGATATGAAGAGAGAGCCATAGCTCATTCCGTAGAGGCAGCAATGGAAATTGGAAGGATGGATGCATGGAGTTTGGTAATTGTTGGGAGGGGGCGCAGGCCTTCTCCTTTAATTTCACCTTTCACAGGGCGGCAAGCAGATTATCCGGAGCTAGGTCCAATAGGTGCGGCTTTGACGACGGCAGCTCTCTCTGAGATCAAGTGTTCCGTACTTGTGGTGCAGCAGTATGATCAGAAGCTAGCGCAAGAGGTACCCCAATCATCCAAAATAATGGACATCTCAACCCCTAAATCTGTAAGTCTTCTGTCCTCGAGTCTTGTTGTAGTCACTGGCCCTGCAGCACCCCCTCCCGATTCCTCTAGTACACCCCAAGCTCCGCGTGTGCCTTTGTAGCAGCAGCCTTTGCACATTGCATCTATATCCCCTTTACGTTCCTTAAATTTTTACCACTAATTAGCAGTCCAATCACTGTGCTATCTGTTTGGACATCTATAAATCCTTCATGCGCCTTTGTAGCAACCTTTGTATATTGTATCCGTAACCAGCAGTCCAATCCTTGACTTTTTGTTTTCACAGTGTGCACAAAGCTTATTTCTAAGTACGATGCCAAGGTTTTTAATACATCATCAAGAAGCTTTGCACATGCAGTGCAGATGGTAACTGCCAGGCCTTTACATGTATTTGTTTGTCAAAGTTTAGTTTCCTTGCTTGATTATTGTAAAGCATGGCTATGGTGATTGCAAGATCCACTCCCTCAGCATTCATGGATCCACCACCTCAGCATTAATGGAGTATACATACATGACAAGGTTGCACAAGCTGCTTTATTTAAGAGTTCTGCAAAACTAGACTTATGAAAGTTGA

**HEAVY METAL ASSOCIATED PROTEIN 51 (HMP51):**

GTCACATTCACTCTTATAATATACAACAATAACTACCCTGCTAAAAGATAAAACGTGTCAACTTGAGCAACTTTGAAGGCATAAACAACTCTGAGCACAGTGTGACCGGAAGGTCCGATGCAGACACCTTTCCTGATGGCAATGACGAAGACAGCTAGCAGCGCTGGAAGATTCAAAGGGAAGTTCATAAAGCCGCCGAAACTCTGCAGCCTGGTACACACGTGCGCAACTTTTGCATCTGACTCACAAATGGGAACAGTTTAAAAATCCTCCATACGAGTGAAATTCCTTGTATGTTGAAGAGAACCTTGCTCATCTTGAGAGACCAGCTGGCACTCATAAAACCTATAATCATGAGGCACATTAGCCTCAGCTCCCGTGGAAGTACCTCATTCATCAATACAAACCCACCGAAGGCGTCTGACTTACTCTCACCAAAGAGCCCTTGTGTGCTCTATTGATGAAAATATATGCGAAGTCTGGCTGCATCCTCCCCCCACTCTGTCAAAATCAATGGACTTTTAGCTGAACCAGGTCCAAAAGCCGAGGCCTTTTATATCGTCTCAGCATCAAGGACGAGCAAACAACGCTCACTGAAGAGAAAGCCATCGCAGCACCAGCCGCCCAAGGAGGTAATAGCACTCCTAAAGTAGGGTACAATGCTCCAGCTGCAATTGGAACTGCCAAAATGTTGTACCCCATGGCAAATGCATAGTTGAGGCGTATGCGGGAAAAGGTTTTCCGCGATAAGTCAATTGCTGTCACCACACCTTCAAGGCTGTTTTTCATCAACACAAAGTCGGCAGCTTCAATGGCAATATCTGTGCCTGCCCCTATTGCAATACCAACATCAGCCACAGCTAGAGCAGGAGAGTCATTTATTCCATCCCCAACCATTGCAACAACAGAGCCACCCGCTTGAAGGTTTTTGATTGCCTCGGCCTTCTGACCTGGAAGAACCTCAGCAATGACACGTTCAATGCCAACCTCCTTTGCAATAGCACGTGCAGTCCTCCAGTTATCACCTGTAACCATAATGCTAGAAATACCCATCCTTTTCAATGCCTCCACTACTACGGCTGCTTCTAGCTTCAAAGGGTCTGCTATCCCAAAAATGCCCTCCAACAGGCCATCTATAGCTACCAACACTTCTGATCTAGCCTTTTGTTCAATTTCTAACAGATGAGCTTCAGCCTCCGACGAGATCACGACTTCTTCGTCCTCCAAGAGTTTTCGATTGCCAACCAAGAGGCTTTTACTGTCAACAAGGCAGCGCACTCCTTTCCCTGGAATGTTCTCAAATGAGTGAACATTTCTCAACCAGCCTGTGTTCTTGACATTTGGTACATTTATGGTCTCGTCGCCTTCTTGAGATGACAAAACGCCATTCATATAATCTACTATAGCTTTAGCCAACGGATGTTCACTGCCAGCCTCGGCAGATGCAACCAGCTCAAGAAAGTCCATCAGCTTCACATCCGAAAACAGCTTGATACTTGTGACACAAGGCTTTCCTTTCGTAAGGGTACCCGTCTTGTCAAAGACAATGTGCTTGATGTTTTGTGCAGTCTCTAAGGCATCCCCACCTTTGATGAGCACTCCGTTCGTTGCTCCAACTCCAGTCGCGACCATAACTGCTGTTGGCGTGGCTAACCCCAAAGCACAAGGACAAGCAATGACCAGAACGGAGATGGCAAACATAAGAGAGAACACAAAAAGGTTGGTTCCCGGCACAAAATCCAAAGAATACAAGCCAAACTTTCCAGCCATATACCAGCAAAACCATGTGACAAATGCCAATGAAACCACTACTGGAACAAAGATGCTAGCCACGTAGTCTGCAAACTTTTGAATGGGAGCTTTCACCATTTGAGCAGTCTCCACGAGGTTAACAATCTGTGTTAGTGCAGCCTCCGATCCCACATGTGTGACCCGAAGATGCAAGACTCCATTCATATTAAGGGTACCACCAATTAGTTTGTCACCAACCTCTTTTGCTACCGGTCTAGCCTCACCAGTGATCATGCTTTCATCCACATGACTTGTACCCCAGATGATAAGACCATCAGCCGGAATCTTTGACCCCGGATATACTTTCAAAACATCCTCTCGTTCCACCAACTGAGCATCAATCTCCTTCTCCATGATAGGATTCCCACCAGCATCAAATGTGAGTAGGATGGCAGTTGTTGGAGCCAGAGCCAAGAGCTTTCCAATAGATTCTGATGTCTTTCCTTTGGCAACCACTTCCAAATATTTCCCAAACAAGACAAAATTGATCAACATGACTGTTGTCTCAAAATATGTTACTAGCCACTGTCCGGTGATGGCGGTATAGAAAATGGCAAAAACAGAATACATGTAGGCAGCAGTTGTGCCCATGGCTACCAGCACATCCATATTAGCTGACATGTTCTTCAAGGAGCGAAAAGCTCCCACATAAAAGCGTTTTCCAACAATAAACTGAACAGGAGACACCAATGCCCATTTCAACCAATCAACTACAAGAAAGGGGCCACAATGCCATGTCATCAGATGATGAAAAGCTCGAACTCGAGGGCATACCACCCAGAGAAAAAGAATAGGGACACTGAATATAAAGCTCAATCTTAACAGTTGCAAATAATGCTGGAGTTCTGTCTTGCGGTCTGGGGAATATGAACTGTAAGGATGGGGTAACGATACTTTGAACTTGCCACTACCTTTGCCTTCTATAGCATCCACAATTGACCTCAACTTTACAATTTCTGGGTCAAAGTTAACTTCTACCCTCTCTAGCAGTGGGTCCACTGCAAATTCCTTCAAACCTTGAATCTGATTGAGGATATATTGGACAGCCTCACCATCCTCCTCAGAGAACATACCTTCAATTGTGATAACCATCTTGCTTCTTTTCTCACTTTCAAAAAATTCAGCTTCAAAGCCAGCGTCATTGATAGCAGCAACAATGTCTTCTTTGCTTACAACAAGTGGGTTATACCTTATCTCCCCCATGCTAGAAGCTAGGCCCACTGCAGCAGAGCTCACACCGTCTAACGAATGCAACACGCTTTCCACAGATACTACACAAGCTGCACAGCTGAGACCTGAGATCCTGAAGCGTCCTGTTATTGTAGAGCTGCCGGTAGCTGAAGGAGGGCTAACAACTGTCTTGGATAATATTTCAGCATCAAAACCTGCATCTTCTATTGCCTCCGTAATGGCATTCTCCTTAACCTCCGAAGGATCGTACTCTACGATGGCTTTGTTTTGGAGGAGGGCCACAGAAGCAGAGGAGACCCCAGGTAACCTCAAGAGGGCCTTCTCTACAGAGGTTGAGCAGGCAGCACAAGTCATGCCCTCCACTTTGACCTCTAATCTAGCAGAACCAGTCTCCGATTTAGCAGGCACCTGACCGATCACCAGATTAATGTCTCCATTGGCACCAGTCGGAGATAAAAGAGGGGCAGATTCCAGGTCAAAATCATCAAAATCCTCCTCATTTCGATGCTTTTCTAAACTCCTCCCCTTCCAGTTCATTAATTGCAAGGCCTTTGACATGATTCCAGGCCGCAAAAGCACAAAGATGATCAACTTCCAAAGGCAATTACAGATGGATCACAGTGTAAGAGAGGCACGCTTTGCTTCTCTTCAACCAAACGTGTTAATCAGTTTCTGGGAAAATGATACAAAAGGGCACAAAATGCACCTTGATTGATGATTGGGCGTATATTCCTCTCCTTATGGAAACCTAAATCAAGCCCTAAACCTTATACACCGGCTAGCCTACAATCAAATGGGCTTTTTTATGTACAGAAACCCAATCATGCAGAAGAAAGTCTTTTACTGCTCGGTCCACAACCTGCTTCGATATTGATTATGAGGTAAGTCAAATCTGGTGACCAAGAAGGCTTGATTGCTGAGTAATCCTGAAAAATGAAGCAGAACTGCTATCTGTGGAGAGAAAATGCTGTACAATAGAGCAAATTTGCTTTCAAATCAAAGATCAAGCCGGATAGCTTTCAGAAAGACCCTTGTGCTTGTTTTTTGAGAGGAAAACCCTATGGCCTGCTCAAATGAGAAGAAGACAAACCCTCATATGTGAGCGGGACTGTTCTTCTTAAATTGCTTGATAGGCGTTGAATTCTGAAGTGGGCGATCCGACGAGGTCTTACAATCTGTGTGTGTGTGGGTGTGTGTG

**NAMED PLASMA MEMBRANE INTRINSIC PROTEIN 1;2 (PIP1;2):**

GCCCAGCCTAGCTAGCAGCTCCTTACCATCAGCATCCTTGCCTTTATCAGCTCTTCTTCTTCCCCTTCTTCACTCACTCCCTATAGTGTCTGTTATCCAAGAGGAGGCCTTCTCATTAAGACTAACATCTTTTGAAGAAGAAGAAGAAGAAGAGATAGAAAATCAGAGATGGAGTCCAAGGACGAGGATGTGAGGCTCGGCGCAAACAAATTCAATGAGAGGCAACCATTGGGCACTGCGGCACAGACCCGTGACTACACGGAGCCTCCGGCAACGCGTCTCTTTGAGCCGGCCGAGTTCTCATCATGGTCATTTTGGAGGGCAGGGATCGCCGAATTCTTCGCGACTTTGCTCTTCCTCTACATCTCCATCCAAACTGTGATGGGTTACAAGCATGGTGTCCCGAGTGCCAAGAACGAATGCCCTGGTGTGGGTATCCAAGGCATTGCTTGGGCATTTGGGGGCATGATCTTTGCCCTTGTCTATTGCACTGCTGGCATCTCAGGAGGGCACATCAACCCCGCGGTGACATGGGGCTTGTTTTTGGCTCGCAAGGTGTCATTGCCAAGGACATTGTACTATATCATAATGCAATGTTTGGGGGCAATATGTGGGGCGGGGATTGTGAAGGGTTTTCAACCAGATTTCTACAATAACAATGGTGGGGGTGCCAATGTGGTAAACCATGGGTACACCAAGGGTGATGGACTCGGTGCTGAGATCGTCGGAACCTTTGTGCTTGTTTACACAGTCTTCTCTGCCACGGATGCCAAGCGTAGTGCTCGTGACTCCCATGTTCCGTTGTTGGCCCCTCTTCCCATTGGTTTTGCAGTCTTCCTTGTGCATCTTGCAACCATTCCTATTACTGGAACCGGAATCAACCCCGCAAGGAGCTTAGGAGCTGCCATCATTTACAACCAGAAGCATGCATGGAATGACCATTGGATCTTTTGGGTTGGACCCTTCATTGGTGCAAGTTTGGCTTGTTTGTACCACGTTGTCGTTATCCGAGCTCTACCCTTCAAGTCTCGCGAAAGCTCCTAGAAACAAAAATCCTACTTGTTAATTAAGGTTTAATTAAGCTAGGTGTGTGAAGCTTGAGTGAAAGCATAAGAGGTTGCCCCAATGCTCCCTTATTGTAAAATCCTTCTTTGTTATTCTCTCATTGTATTATCTTCTGTAATAATCCACCACATCCTTTTTTGTCTAGTTTAATCAATTAATACTTTTTAATTAAAGATGACAGTTTTTGATTTATTGATAAAAA

**NAMED PLASMA MEMBRANE INTRINSIC PROTEIN 2;1 (PIP2;1):**

AGAAGGGGTGCATACGCCTCAAAATGAAACAAAAAGCAACATTTTTTTTTTACCAAATAAAAATGACATGCATACTTCACTTGCTTAGCACAAAGAAATGAAGGAAATCATAACTTTGCTGGGGTAAGGTTCCACATGTAGACAAGAATGAAGTGAAACCAGCTAGCTAATGTGTGAAAGGGATGATGGAAAAGAGTCTCAAGAGCAAAGATTCTACCTAGAGAATGCTCTAGATTCATAAGAGATAAAGATAAGAGATTTGGGTTTATGAGTGTATGTGTGTTTTTAGGTTTTGTGTTGTGTTTTTTTGTTTGTTTGTTTTTTTTTTTTTTTTAAAAGGGGTGTGGGAACTCATCAAGCTAGAATTCACACTTAATCACGATACAATATCTTTAGATGTTTGACTGGCTTCGAAAGGAGCCCATATTCTTGAAAGCTCCAGCGCGCAATATGTATTGATGGTAGCTTGCAGCAAGAGCTGCTCCAATAAAGGGACCCACCCAAAAAATCCAATGCTCATCCCAAGATTGCTTGTGGCCAAAGATGACGGCAGCTCCAAAGCTTCGAGCCGGGTTAATGCCGGTTCCAGTGATAGGTATGGTTGCTAGGTGGACCATGAACACTGCAAACCCTATGGGTAGAGGTGCAAGAACCGGGACATGGGAGTCACGTGCATTACGCTTTGGGTCAGTGGCAGAAAAGACAGTGTAGACAAGGACAAAAGTGCCAAGAATCTCCGCCGCGAGTCCGGTCCCTTTAGAGTAGTCAGAATTCACTACATTGGCACCACCACCTGATCTCAAATAGTAGGAGTGCTGGAATGACTTCACCAATCCCACCCCACATACTGCTCCTAAGCATTGTGCCACTATGTAGAAAACAGCTCTTGGAAGTGACACTTTTCGTGCCAAGAACAATCCAAAAGTCACCGCGGGGTTGATGTGCCCCCCTGAGATGCCGGCAGTGCAGTAGACGAGGATGAAGATCATGCCGCCAAAGGCCCACGAGATACCGAGCAGCCCGACACCACCACAGCTCTCGGCCTGCACTGCCCCAATCACCGTGCCCACGCACACATACAAAAAGAGCAGCGTCGCCACGAACTCCGCTATCACGGCCCGGTAGAAAGACCACAGCACAAACTCGTTGCCATCCACAAAAGGGGCTGGTGGAGGGTCCTCATAATCCTTAACTCCTTGCTCCTTCCCCATCTCTCTCTCTTACACACACACA

**MIZU-KUSSEL2 (MIZ2)**:

ATTTCCTGTGTCCTTATCAAGCAATCACCTTGCATAGTTGAAGTGACTCATTTTCTATAAATTATATCCTCTTGCCCCAATCTTCTTCGACTTCAAGCTTGATGCCAAAACATGGCTACACAACAAGCCATCTGACAATCCTACAAATAGAAATTCAGAATAAGATTTAAAATCAACTAGTCAATTAAAGAAACTCATATGTTTCCTGTAAAAAAGGAGAGGTTACATGGGCTGCCAAGCCTTCACAGATAACTTGTGATTCCAAATCAATTATAATGGGGCTTGTGCCTTTGAATTTCTTCCCCCCTAGCATTGAACATGGAACATAATTACGAAATTTTGTGCCCCTGCAAACTTCTAAAACCGATGGTCCATCTTAACAAGCTCTCAAGGCCTTGCCGAGTTATAGACCAACCAGAACTGGTCATACACATTTTGGAAACTTTACTAGCCAGACCACAAGATTTTATATGGTTATGCAGGCATCATAACCATCATGTAATGTGAGTGCTTTGTAAGAATGTTGTTACACACATGCATTCACCTGATAGTGTCTGCACCAGTGCCTAACTAATGCGATGGATTGTGGATACACATGCATGCAGAGGAAAAAAACTTATTGTCTCAAAGCAAGATAGCATGCTTTGTCCCAATAGACACTGCACAACAATCCATGCCTCTCTACACAACTGAAACCTAAACACATCCAGCATGATTTGCTAAGATCCTGTCTCCACATCTGTACATGCCTCAATTGATCTGACCTCTTGCGACTCAGACTCCAATTGAGGTAGTAGCTCTGCCTGTAAAGATGGGGATATATCCTGAACATGATGCCAAGTGAGTTCCCACAAGCTATCGCAACCTAGTGTGCTTCTCTGGACAAGCACTCCCCGAGCCTTCATTATCAGCAGCATATTCTTCAAAAGCTCTGGAATAAGCTCCTGAATTTTTTCACTATCTTTTCCTCTCAACTTGGCTTTCCTGTACCTGTTCATTCGGCTTAAGACAGCCAGCCAGAGTGCTCGAAACCCCGGTAGCAAAGAAAGTTGATCCAGGAACTGAAGAAAAGTTTTGGAAACCAACTTCATTGCATGATGCAGAGTGTTTTCCATGTTGCGGTACTCCTTGGGAGAGTGTCCAAGAGCAATTTCTAATAAATCATCCAGCATAACAAATATCACTTGATCAAAGCATTGCCCCCATATTAATGGTGGTAGGTGCATGCTCTCTGCAGCCGATAAGCTGCGCTGAAGTGATAGTATTGCATAATTTCTTACCTCTTCTCTTGACTCTCCACAAACCTTTCTCAATCCCTGTGCTAACCTCAGCCACATCTCTCCAAGCTCCTGTGATCTTTTCACTACCTCCTCACCCGCTGGAGCAGTTGACCAACGTATGAGGCATTTTGTAGACTCAGCCATGAGGTCCAAGGCCCTAAGAGACCTGTCATTTCCACCAACAGACACTTCTGCAAATGCTCTTGCAGCATCAAGACAAAGTACATAATTTGCCTGAGTAAGATGGGCCCCTTCAGCCATTATGAATGACAAGGCTTCAAAACCTGCTTCTGAAGCCTCCAAGTGGCGGGCTGTAATGGATAATAATGATGAAACAGTGCGCCATCCCATAGGGGATCTAATATGTTTGGCATTTGTCTTTACAAGCTGCATAACTTCCTGAGTAATCCTCTCACAGAAAGCATCAGCTACCCGTGCATCCAGCTTTAAAATCAACTGTAAAGAGCGGAGAAGCTCTTCTGCCAGATCCTCTTTGTATGGGAGTAGCCTCTGACAGATGCGTAAAAGCCCAAAAACTGCTTTCTCAACTAAAAGACAAGGCATTAGTGTTGTCTGGACAATGTTGGCTATGTGTTCATAGACTCCCTGCCAAAGGAGCATTATTCTGTCTCTATTGTTGAGAGCAACAGCAATCAATAGATCCAAACAAAAAACTGCCGTATCTTCATCATCAGGTGATCCACCAGCTTTCTGTGGTCGCCCTGCAGCCCACATGAGGGCTCTAACTAGCTGCAGCAATGACTCAGCTTGCAGAAATTTGCTCTCTGTAATGATGCTGTCAATGCGGCAAGCCTCTATGGCACGCAGAGTTCGCTGATGAGCAGCCAATTGCTGCTCAGTTGGTTGAGACCTTGGTTCGTCTGTTTCAAGAGATAACAGCTGACTAAATCGTCCCATCAACCCCTGGGACCGCTTCTTGGCTCCTGCTTGGGGTATCGCAATCAGTGTTGGGCCACCTGGGATTGGCCTTCCATGCATAGTATCACCAGCAACATCCATATCATCTCCAACATCACTAGCAACACGAGCAGGCAGAAGGCCAAGCTTGTGCAGCCGCAATATGCAGTCAAGTATATTCCTCCAGCTTGTACGTATATAATCCCCATATTTGTTAGCAATATTGAAAACTGTCACTGCAGCCATGCGAGCTTTTGTATCATCACCAAATGCTATTACGGGTTCTTCCACAGATGATGTAGGATTCAGTAATGTTGTGAATTTGCACAATGACACAACAAGGTCATCTAGCACATCCTCTAAGTGATGGCTTGATGATATCTTTGCTACTGCAAGAAACCCTTCCACACACATCCTTAAGACATCTTCATCCTCAGCATGATCAAAGACCACAGAAATGGCAGCTATGGTTGGTCCTGATATTATTGCAAACATGTCACGATCCAAAAATGGTCGTGAATCACATTTTATGTATGGTGAAGACTTTTGAGACCTATGAATCAGGCCAACCCATCTACTTGGTGTCATTTCGGTAACACCTGTCGTATTGTCGTATGTTATTTTGATCTCATTTCTAACAATGGAGTGGTACAACTCAGAGAGAAAATCCCGAGGTAGATCCTGGCCACCATTGATACTCCTGTTGTTCTTAATGAAGTCTTCTTCAGTCATCTTCTTCTTCACCTGAGAATTGTGCTGGTCAGTGTTTAACATAATGAGAGAATATGCCAACAAGAAAGCAGCATCCTTGTCAGCAAGTATTTGAGAGGTCTGTTCATAATATCTCCCTGCAAAGGCTTCTAGGACCCTGTGAATCTTTTGTGCCTCTCCAGGCAATCGGAATGTTTCCAAAAACGTTCGCATAGCTGCATCCAAGTTCATGCCTTGGAAATCAAAGGTCCTTGCAAACTCTGCCAGCACTTGCAAACAAAACTCATCTGGATCACCTAAGAAATCACCCAGCAGACTCTTGTCCAGCCCTGTGGTGTACCTAAAAAAGCAAGCAACACTCTGAGGATCGAGCTTTTCAGGCAGCAGACGGATTCCCTGCAAGAACTCAAGACCTTTCTTTGGATCTCTGTTAAAGTGATCTGCACCGATCATCAACCTCCGTTTGATGTACTTTTGCCGCTGAACAAATTTAACCCAACACTTGGGATCCTTGTAGTTCTCACATCTCAAGGTCCAAAAGGGGACATACTCTGGCACCTCACTTGGGTTAGCAGGCTCTACTACAGGAATGACAGAACCTGAAGAACCAGCCCGCTCTGCCATGCTGTGAATAACTGCAATTAGCCCTTCCAAAGCTAAAATGTGCATAGCTGACAAGGGACAATTGACAGGAAATGCACTCTTTGAAAGCAGATTTCCCAAGTCTTCAAAAATATTCCCACATGTGATATCACAGTCAAAATTAGCATACATCTCTGGCATGAATGTAGACTGTCTGCAGAAGTCTACCAAAGCTTCCATTGCCACCTCTTGTTGCTGATAGGTGGCCCCATACTTTCCTTGGGATAACCTTATGATGACACAAGAGAAAAAGGCTTCAAGCTGTAACTTCAAATGAGACCTGAGATGATGATACAAATTAAGTACAATGCTGCAAACCATTGAAAGAATGAGAGGATTCATATTTGAGAGCCCCAACTGCATAAGGAAGCGGAAAAGATCATCTTGCACAAGGGCTATCAGTTCAGGATGACAGCCGAATGATGAGCCGCCAAGCTCAACTGCTGAATTGATCAAACCCAAAGCAAAGAGTGGGACATCCTCATCAATAGCTAGCAGGTTTTGCCCAGTTACCATTCCAGAATCTACAATGTTTAAAAGGGAACACAAATAGCGGAATATTTCAACAATACCCACAATGCCATATGGCTCTATAATCAGATGAACCTTATTTCCCAAATCTTGATCAACAATTTGGCCATTTTCTTTTCTGCCTTCCAATAAACCACCACTGGATCTTGCAGAGACACCATCTGCCTTAGACCACAAAGCGTCCCCAGACTTCTCTGTGCCAGATAACAAATCAGCTTCAAAACCATTTGTGCTGACAGCACCATTGTTTGGGCCATCAACACTGTCTATATGTGTGCCATTTGCCATATCCCCGCTTTCGTTGCCAATCGTTGCAGAGCCAATGTTTGTCAATAGCTTGAGTGAAGGATCCAAATTGGGTAAATGGGCAAAGATTGACCTCACAATCTCATGCATGCTGTGCCGTGCTGTTCTTTGAAGCAGCTCTCCTTTGGTCACTGCCTGATGGACCACACGAAAGCAAGTGTTGACAACAAGACAGACATCCCTATCGCTTAACAAATTTGAAGCTGGATGCTTCATCACCACCATCAACACTTGTAGAATCTTCATGAGAACCACCTCCTCAGATGCCATATCCGTGACCTCAAATCGGCAACTGGTCACAGCATCTACTATCTCATGCATTGCCTCATCGACATGGGCGGTTTGCGAATCAAACACCTCTAAAGCCAGGATCTTGTATACAGCAGAAAGCGCAACTCCGGTGATGGGAGCTCCTGTTTCATCTGAGCGGACCACGTCAAGGAAAGGTGTAAGGTATAAAGATGGATTGATTTTGATCCAGGGTTGCTGCCAGCTGAATATGCTACGCCGGAGAGTTTTCAAAGAATACACCAGTGAATGTTCCAACTGGTCGTCACCACCCATGTAGCGACCCGCCCATCGAGCATTTCTCCTCATTACAGCCAAGACAGCTCCTATTTCTGCATTCACCATGCAAGCCAATGCACCACTGCTCCTGCTGGACTTCCTCATCTTAAAGATGTCCCAATTAAGAAAAATGAGACCCAAAAATCTTGGTCAGCTTCTCAAGCCAAAAAGGACTGAAATTTCTTGAATCTGTGTATAATTGGCAGCCCAAGAATGCCGAGTTCACAGAATTGTAGCCAGGTATACCATGAAGATTTGCTTCTGTAGTGGAAAACTAAGCTCTTCCTTCTGTGATTTCTTGAAATCCCTGAGAAATCCTTGGCCTTCAATGCAGTAGATCAAAGTCTCTGCACAATCAGGGAGACCTCTCAATTGTCCGATCCTTTCAGGGAAGCATGAAGTCTAACTTGATGCAGCAGGAAAATGGAAGGCGTGCTCCATGCCGTCAAAGACTCCCCTTCATGCTTGGATCTTGTAAATGCCTGTCATGTCAAGCTCCATAGATGAAGGCATTCCCAATGCACAGAGTACACAGAAATAGAAGCTCCAAAAACGCAGAAGAAACTCTTATTTCAATTGCACAGGGGCCAAAATGTGATCTGCGGATCAATCTATGGAAGTGCACAGCAAGAAATAATCAGACCTGGAGAAGAAGAAGAAGAAGAAGAA

**CONSERVED OLIGOMERIC GOLGI COMPLEX 8 (COG8)**:

TCTCGATCTCGTAGTCAGGTAGCTACACACACACACACACACGAACGCCATGATACAATTGCATGCTATCTATACGCATGCCTGTGCTTGAACGCATATGTAGCTACACATGCAGAGGCATCCATGCTTTTGCGGTTATGTTGTTGAAGAGATGGCATAAGGGAATGTTACATTTCTTCTAAGTCTCTGTATCCTCTCTTAAAAGCGGATCATTATGGACGTGCTTCCGGCATCTCTCAATAATGCCTCGCAACAAGCATATGTCTCGGAGCTTCTATCGTTCAGTTTGGACAGGTTGCACAAGGAACCTGAACTTCTACGAGTTGATTCCGAGAAAATTCGGAGACAGATGCAAGATTTGGCAGTTGGGCATTACCGAGCTTTCATCATAGCAGCTGAAGCTCTTCACTCTGTTGGGGAAGAGATGACGTCTGTAGATAAGCATCTGGAAACTCTGATCACAGAGATTCCAGAGCTTACTGGTGGCTGTAACAAGTTTATGGAATGTGGACATCAGATTTTAGAGAAGCGCAGCATGAACCAGACGCTTCTAGCAAACCACAGTGCGATTTTGGACTTGCTAGAAATTCCTCAATTAATGGACACGTGTGTCCGAAATGGGAACTACGATGAAGCTCTTGATCTAGAAGCCTTTGTCTCAAAGCTGGCAACTATGCACTCCAAGCTTCCTGTCATTCAATCATTAGTGACAGACGTCAAACATATCACTCAATCTATGCTTTCACAATTGCTGCAAAGATTACGGTCTAGTATTCAGCTGCCAGAATGTCTCAGAGTAATAGGGTATTTACGGCGTTTGGGAGTGTTCACTGAGCATGAAATGCGTCTTCAGTTCTTGAGATGCAGAGATACATGGCTTGCTGGGATTTTAGATGATTTGGATCAAGGAAATCCTTATGACTACTTGAAGAGGATAGCAGATTGCCATCGGGTACATCTTTTTGATGTTGTCACCCAGTACCGAGCAATTTTTACTGATGATACTTCAGGAAATGAGGAGAAATATGATGGAGGCCTTCTCTACAGCTGGGCAATGCATAGAATTACCTTACATTTGGCTGTACTCCAAGCTTTACTTCCAAAAATATCAGATGGAATGTCACTGCTTGAGCAATGCATGTATTGTGGCATGAGTCTCGGACGTGTTGGCATGGATTTCAGAGGCCTTTTACCTCCTTTATTTGAAATATCGGTGTACAACCTCTTTTCAAGTAACATGGCTGCTACCATTGAGAACTTTCAGCATGTGCTAGACTCCCACCGATGGATTTCTCTTCCCACAGTTGGTGTTGGCAATAGAGGGGCTGCCGATGAGAATTTGAATGACAATGCGGCTCCACCTTATGTCTTAATGGAACACCCTCCCCTTGCTGCATTTGTGAATGGTGTTCTAACAGCTTTAAATGAGCTGAGACAATGTGCACCTTTGAATCTTAAAGGTGTTTTAGCTTCAGAACTTCAGCAAGCATTGCAAGCTGTATCCGATTCGCTTCTTCGATACAGTGCAACTTGCATGCTGAGAGAAAATGAGTCGAATCTTTTCATGGCCATGTGCCGTGCTTTTATTGAGGTGGCATGTCCGTACTGCATCTCCAGCTTTCGTCGGTGTTATCCTGGTTCGAACCTGATAGATTCAAAAGATTTGCTGGAAAATCTTCGCAGGCTTGTGTTTACAGCATCATTTCCAGTTTTGCAGAACAATGGGTTTTCATCGTCTTCACCCAGGCTAGTAGTCAATGAGCATAGAGAAGGGTCAACAGACACTACTAATGTGCATGCTGTGGTGTCACTTGCAAATGGAGAAGTCCAGCCTACAAGCAACGGTTCCGAAGTCTTAGCTGGGGAGGGAGAATCACAGGATCTCGAAGATGACCATCAGCAAGTAGAGGTCAATGATGGTATACCTGCACAACAATCTCAAGCCTCGGAACCGAAGTAGTCTAGTTTGGTTTTTGTCTTTGTTCATCCTTTTCTGGACTTGACAGTGGCCTGAGGGCTTTTATTGCTGCTCAATGAGTAGTTGCAGGTGGTTGAGAATTGTGCAGCCGATGTGTGCCCTTGTTATTGCAGAAGTAACCGACAGGTTGCTGCAAATAACCCAAGATTCACTAAGCCATTCCCGTCCTTACAGTAGCTGTTCTGATAGCACGCATTCTCAAATTTTATATCAATCCACTCGCTCATTCCACCATTATATATTGAAGTTTTGTATTTTTTTTGGGGAAAAAGTTGCACTTTACGAAGAGAAACAAATTCTTGAGAAAAATAGAAATTTACTCAAGAGAGTAACAGTGTATAGTATGACACCTTTGTAATGAGGCTTGTAAAATGCGAAGAGTTGAGGGGTAATTTGAGACAATCAATTTGTTAGGGGTCTCAAGAGTTTTTGTCTCCATCTATTTGTAATGGGTTTTTAAAGTTATGCACATTGAAGGACAGTAATTATAGGGTTATACTTCCATTAATTTTGGGCTGTGCCTTTTGAAGGACACCAAAAGTGTTCCTACAAAGTTATGAAAAGCCTGTTAGAAGTTTTGTAAAGGCTAGTAAATTTAGTCATTAA

**TRANSPORT PROTEIN PARTICLE C 9 (TRAPPC9)**:

GAACAGTATAAAAATTGATACCTTCCTCAGCAAAACTGAGAACATAGTATAGAGCATTCCAAATGCTCATGGAGATCTAAGATGGTTTTCATTCAGCAAATTGATCCAAAATTCCATTCAACTAATGCATACAGTTAATAAAGAAGACAACAAATCTTGATTTAGTGAGAATTTGAAAATTAAATATTGTTGATGATTTAATACACACTGCAAACTTGTGATGCCTTACCCTAAACATTAACAATCGAAAGTAGGTTACAAAAGGTAATCCTCCGCCATCCCAGCTGCAAAAGAGTGGCCACCATTGTTTTGCACAATTCACACTTCTGTGCTGATTTGCACTCTTCTCATTGGCCATCATCTGTGCTGAATCCATCTAGATGGTGAATTGTATCATCCACATCCAAAACTATGCAAGGATGGTCGTCCTCCCTCCCCAGGACTGTGAAGGCCGTGATTGTCTCATTACAGCTTAGATCACCCCTTTTCTAAGAGGGTAAGGTATCCCCACAACGACTCAATTCCAGGGAAGACCTGGCTCAGATCTGAATCACGGACTTGTATTTACCCATAAATTATAATCAACCAACAGACTGTATCAGCTGCTAAAATGCCCTGATACATAGAGCCTAAATGGGGGTCCACTGCAAGTCAAAGATTGAGCCCCATCTGAAGATGATAACTTTAATGGAGAACCATGCAGCTCACTGTTTTGGTGTATGACAGCTGCTCCAAGCAGGGTGTACTCTCCTGGTACTAGAAAACATAAAGCAAAAGTATGCACAATTTTTCCTAGTGGTGGAACTTCCACTTCAACCCCATTCAGAGCACCAGCCCAAAGTACAGTTGCTTTCTCCCCACCTAAGCAATTCTCCCCTGCAACATCTCGGCATGTGACGCTGAAGCTTAATGTTACAGATTCTCTGGTATTGTTTCGCACAAGCATTTCAAGTGGGATCAAGTCATGGGTCTGGATGGTGCTCTTACGAATACCATTTGAGACAGCATCTTTCCCCTTCTTTGCAGCAGTTTCCATGACTTCAGAGTTCATCCCACAATCAGCACTGTTTTCAACTGGCTTTGGAGTTACTCGAAAACCAAAAGTAAGAGGATCAGGGCGTAGCACATCAACGGCGGAAGCCTGAAGTGCCTCTCGAAGGGCATCCTTTATAGCCAGTTCACCAGCGCTGTTTCTTCCTGAGTTCCATTTCACACATATTCTAGAGCTCAATTCCTTGATCGTTATGTTTAACTCTGCTCTTGTTTGTCTCTCTGATTTTGCTAGATCATTTACTCTATTAAAAAATGCTTTATTGAGAATTGGCAGCTTTAGTCGCTCTAAAGGTATGAGAACTCTTGCAGAGCACTCCCGATCAATTCTCCTGCTGGGATACAAGCATTCCACGCTATCCTCCAAAAGCTTCAATCCCTCTAAGCCTTCTCTAGTGCTTCGTTCACAAACAGTAATTTCAAAAACAACATCAGTAGGATTCCACAGCTCCAGCTCTAGCAGCCGCAATCCCCAACTACCTCTGTAAGGATCCATTTTTACAAGACTACCACCATCAATCGATGTCACAGGAGGGGTTGCATCGGTTGTATTAATTTGAGCTGGAACCTCCATAGACAACAATCGTGCATGTACTAAGCATAGCCCCTGAAGCACATGCATTTGCAAAGGAATGGTTAATCTTCTTCCGGGTGGAATTTTGGAGTTCTCACTCTGAACTTCTTCATCGTTCATTGAAGCTTCTCCTGCATAATGGATCACCAACAAAGGGCTGGTCGTATCCTTGACCACTCTTCCAGAGTTGCTCCCAGTATTATCCTGCTGACGTATTATCTCAGGACTCATAGAAAGTTGGCCAGCCTTTAGAATCACAGGGACTACAACACTGGCTCCAGGAAGCAGTGGAAGTGCCATTTGTAGCGCACCATGCCCCATTGAAACCACATGCTCTTGCTGCTTTCCTGTCAAGGACATGTGTGCTTCTACAGCAGGAACTGAACTTGTATTTGAAATACATATTTGAACTTCTCTTACTTCTCCTTCATACAACACAGCAGCACCCTGACCTCCGATTATTTGCGCAGTGAGGAGAGGAAGAGAAGGCATAACTGATATATTTGGACTCGATGCTTGGCGCAATTTGGGGCCTCCAGCATTTCTGAAAGGATCGGTTAAGCTCAGGCCCTTTGCCACATCTGATAATAGCACATTTGTGTCTTCAAATACGTGTTCTGTAATTACGTTAAAGCAGTGAACGATGCAGCCCCTTAGAAAAATTGGGCTCTCTTTAACTGGGGTTCCAGATAATGAGATTAACCGAGAGGAGTTTGATGGAAGAGTCACGACACAAGGATAGGGCTCAAATTCTCCAGACTGAACCGATAAGTATATGCTCTCAACAAGCACTGCAAAAGCACACGGGTTTGCAAGTTCCACTAACACTTGCACTGGCTCTCCAACAACCCAGACAATGTCCTGTTTGCTTCCAGAAGTGGAGTCATTTCCTTTAGCAAATGGAGTGTAAATAAAGGGACCCGAAGGAGCTGCTCCAGTCCACCATTCTTTTTTCCCCAGGGTTCTTGTCACTATTTCCATCTGTGAAGGTAAACTTGCGACAGAATGCAGCCTAACAAAAGGCAGACAAGGATCTGAACAGCGGGTTCCTGGTGGAAGCTGTCTAGCAGCTGTTGTTAAAGCACCAGCTAATCTGGCTTGATCACTTGGAGTGATAAGAGGATAATGGCCCCTCAATAAACGTGCTGCTGCACTCCACGCTGCAAGGGGATCACTTGTTCGTACAGCTGCTAACAGCATATCATTGAGTACATTCATTTGCAAAGTGCTCCACAGTCCATCTGCATCAGATGGGAGAAACTCCTTGATCTCATCATTGCTTCCTTGAGTATAGCTGGTTCATCTCTTTTGAGTCGGGATGAGTTACTTTGGATATTGTAAGTCTTCGCTCCCAATGCCAAGACTTGTAAAGCACTACTAGCTGACCATTGACCTTCTTGCTGGAGATATAGTTGAGCCACTTGTCTTGAGAAAAATGCAGCCTTTCTTTCATAACCTAGACACCCAAATATGCGAGCAACCTCAACATTTAGAATAAGACGGTCACTAGCATCCACAATACTTTTGGCTCCTTCCACAGCACCCATCAATAGATCTGACACTTCTTTTGCAAGCTCCTTTCTGCAAAGAAACCTTGAAAGCTTTAGTGTAGCTTCCAGCTCAAAAGCTATTGCCATGGCCCTTCTGTAGAGCTGTATAACCTCATGATAGCGAAACTTCACTTCCTCCTCAATATATGGATCTCTGTGTCCCCAATGATGTATCACTAGAGCGCACACACTCCCTTCAATTGCTCCTGCATGCCAGAACAAATCCCCTGTGAGCCGTGCTAGCTCTATTGCCGTATTATAATGTGCATTCGCATCTGAAGGAGAGCCTGCTAAGAGACAATAATCTCCAATGGTTTTCTGAGCTCTGCCTAACCTTCTTTTCTTTGCTTTACTCACCTCTTCAGTGCTTAGTGATGCTTGTGAATCCAAGGGGGTAGTTAACGTTGCACCCACAGTCTCAGCATGGAGGACCCAGCTTTCAAACTCCATCAAAAAATTTGCAGCTACGTCCTGCATAAGAGTTTGCATGTGCAACTCCAATGTTTGCCTGTCCGCAGGGGGCACAAATATAAGATGTTGCCTTTTCTTTTCCTCCTGCTCAACCTGATGTGCATCGCTTGGGAAAAATGCAAGCATGCGCTTGACAAGTGCGGAGGGATAAGCCTTGCAAATGGCAGTGAACTGTTCGTAGACGGAGACAATATCAGGAGAACAAGGGCAATGACAGAGACCGATGATGCCATGGATCTTGCGCTGAGCTTGAAAATCTTCCCAAGGGCTACGAGGGGCACCGCCTACAAGGAACTTGAACCAGAGACTGCCCGAGTTCCATGGCTGCTGAGCAAAAGGGCTCTTTTGGTGCTCTGTGTAGAAGGAGCTTACACTTGTAAGATCGACTTGCTTGTGGCTCACAACAAGAGAAATGTAGTCACGCAGGTAAGCACCTGGGATTTGGCCAACACCCACAACTACAATCCTGATCATGCACCCACTTTCCACACTCACATCGGGCTCCATCTTGCATGAAGAAGTGGGATCCTTTTTAGCTGCCAGGGTTATGCAAGTTTGAGTTTCTCAGAAAAGTTTAATCTGTAGCATGTTAACCTGCTTAGGATTCAAGCCAAGCTTTCATGGATATCTTCGTAATGGCACCTTCCATCTTGCACTTCCTTTCTGAGGTTATACAAATGAAAAGCTTAGTCTACATCTGCTATATGCAAACTAGAGTTTTATCTGCTACCTGCTGCGGTACATAAAAGCTATAGATTTAAGCCAAGCTTTCATAGATCTCTACTTTAATCAAGTTCCACGACTTGTTCGATGCAAGCCAACAGAAACAGGCAACAAAATTTTGAAAAAGTACGTCATAGAG

**VPS53 HOMOLOG (VPS53)**:

AAAAAAATTGCCATGACATCCCTGCACCTTGGCCGCCCAGAGTAATCTCAGTGTACTTATTTATGGTGCTTCGGCTCCAAATGACAACTGAAATTTTCTTGTTAGATCAGATTGAAAGTGCTTAGTTCTATAAAACACGGATGCGCACAAATGCACACACAAACATGATACAAAAATCGAGTCATAAACACACATTAACTGCCATAATGGACAGAGACTAAATTGCCAGTTGTAAATATGCCCTTAAATGACAAGGTGATGCTCGATTTCGTCCAGCTCACCTTGTATGCCAAACAGCACTAAATAGAAAGTGGCCCAATTACGCTTTGACAAAACAATCCTCTGAGACAGAACACCCAAGGGCCCCCCCTTGTCAAAAATGTGATGGAAGACTATCATAATCATAATGAGCACAAGCGTAAAGCAACTTTGGAATCCTGTAACAGAGGCAACCTGCAGCACTACTTGGATTCAGTGTGCAACCGCTGTCCGTTTATGTGTTGAACAGCTTTCTAAATGTGCCCTCCTTCTTTGAAGCCCCTTCTTTTGCAGACTCTGCCAAAGCAAATATTCTTTTCAAGCCTGTACTAGAAACAGCTGCAGCAGCAGCAGCAGATTGGGCCATGGCTCCCCTCCCTAATGCAGCCGCTCGTGCCATCATTGCATCCCTACTTGCAACACTGACAGCAGTGGGGGTGACCGGGGTAGGCGGCGTGGCTGGAGCTATAGCAACACTTGGGGTCACAGAGGGTAAGGGACTACTTGTAGCTGCCCCACTTGTGCCACGCCGATTGAAATAATCAAGAAGTGGTTGTTGTTCTGCCTTCTTCAAGCCCTTCAAGTCAAGCAAGCGTTGAAAATCAGTTGCAGTTCCTTCCAGCAACAAGGCACGATATGTGTCTGCAATTGACTCAACGGGTGACAAGATCACCTTTAAAAGAGCCTCAGCCTTGCCCATCTCGCGGCCAACAAATTTGTTGTAGCTTGCTGATGCAGAGTTCTGACCACCAAGGGAAGGAACTTCCAGTAATAATGTCTTCACAGCATGTGTATCCAGTAGCATCTGTTGTGCTCCCGTCTCAGAGATCCGCTTACATTTGTAGATATTGACATGAAAACGTGGGGCGAAAGATGCTGCTAGCTTGTCTAAGAAAAACTGATAATATAATGGGGACAAAAGGGCACTGATAGCAGGAACACTGCTTGAAAGTACAGCACTTATACCATTTACATATTCAGATTGATCTCCCACGCTTTCCACTGTAGCCCAAGCCATGCGCGACATTGAAGCTAGCTCAGCCTCCAGTTTGGTCTCCAAACCTTGAACTAACGTGGCTAGAGCCCTTGTTATGACTACTGAAAACTCATCCTGCACCTCTGACATGTCAACCTTGTCTGCAAACTGAAAATCAATAATCTTTGATATGTTCTCAGCCAGCTGCCCTGCTGTTTCATGACAATATTCTGCAGTGTTGACAATGTAGCAGATTACCCTTTCATCTTTGTCTGAGGTTTTGATTTGTCCGTCAGTGCCTGTTGCTGCTGCAACGATGCCAGTACCTCCCCTTGGGAGCCTAGATAGAAGCTTTGCAGCATAGGCCTTCAAGATCCTTTGAAACACCAAGAATAGATTGTACAGAGTTTGATTCCTCGTGAGAGCAGAGCAGCGCTTTAAACTCTTTTTTATAATTAAGAAGACCTGTGTGCTGCTTGAAAGGATGTTTGTTTGACTGCCTTCTTCAGCTTCCCACGATTCTGTCTGTACTAGTTTCTCTACACTTTCCATCAATGTTCTTTCCTCCAGTTCAACATATGCTGACATGTGTGGCTCAAAACAAGAGGAAATGATGCCGTAAAAGCTCAGAGGAATTGACACCGGACCTTTACTGCTATTATCCTCAGTTACACTGCTGGACTCTACATTCCTGCCAGCTTTTTCTGCAAGTTGCTTCTCATACTTCTTCCTAATTGCTGAAGCAGAGTGTTCATCTTCTCCACCTGGCTGTGCATCATCAGACTCATCCGCACTATCCACATCTTTGCTGGCAGTATGATCATTTCCTTCTCCAAACGCAGTTGCAAGATCTTTCTCAAACTCTGTCGTGCTCTTTAACGTCTGCAAGAGAGTAGCTACGTCAGGTTTTTCTTTGAGGTCATCCAAAGTTTCGGTCAGTTGCACCCTTGTAGCTTTGCAAAGCTGGATGCACAGCAGATATAGTACACGCCAAGCTGGAGGAAAGATTTTCCAAATCTCTTCATTTGTGCGCATCTGCCGTTTTATCCATGCATAACGACGTTCTACCTTGTCCAACCTTGCAAGCTTTGCTCTATTCCCTCACAAATATGGCGGTATTTTAGCAATCAGCTCCTCCCTCCCTCTCCCTTATAGATGCATCGAGCACATCAACAACCAGGCAGATGCATTGATGCATCGAGCGCATCAACAACCAAGCAAGCATCTGCCAACTGTTTCATAAGATTTCCATCCTCTTTTAAGCGTGCTGTCCCCAAGCTTGAAAAATCAGAAAACACGTGAGACTTTAGCATCTGTTTTATGCTCTTGAACTTTTCTCTTAGCTCAGAGATTTTAGGTATATCTCTGTATGCTTCGAAGTGCCCACAGAGCTGATTCACTGCCTCTAATTGGCCTGATGCTTCTTTATATAGCCGTTTGGCTGCCATGGTTTGAAGTTGATCCACAGCCGACACAAGCATTGCAAGGCGATGAAGCGCTGTAATGGTTGTGGTGATGTGCTTCTTGGCGAAGTCTAACTTTTTGATGTCACGGCAAATTTCTTGCACCATAACTTCACTTTGCTCTGCTTTCCCTTTTATTTCCTTTATTCTTTGTGTAAGCTCCTGTATGGCTTGCATTGCAGCTGCCAAGTCCTCCTTTGCTTTTGTTCCTGAAGAGCTTTGTTGGCGAACTGCAGCAAGTATCTCGGCATCAACTTGTCGAATCTTGTTTCGTAATTTTTGGATAATTGGCTCCACTGCCACCAGAGATGCTTCATTGGGGAAAGTCTGATTGATAAAATCCACAGTAGTGGTCTCCATTGAAGCAAACAATCAGAATAAGCCCTTCGCGTCTATTGAAATCTGTAGACAAATTACGGTCCCAAGTTCAAATCTTGGCATGCCCGAGCACTTGTATGTGTGAACGCGCGCGCATGTGAGCGAGAGAGAGAGAGAGAGCGAGAG

**MAIGO 5 (MAG5):**

CTAAACACGGGCATCCATGAGTGCACACACTGGCCACAATTAATGTAGAGTGCATGGAGTTTCAAGTTGCTTTGCTTTTGAACATATACAAAGTAAATTTTTAAAGACGGGAAAAGTGAACAGAATGAAACAAAAGAATTTACAACTGCTCAATGAGCCGTACACTACATGCTGCAAGAGCAGGTTTCCAGCGTTTTAAGAACAGGAGAGGTAAGGCATCTGTGTAACAGGCTTTCTTGACAGTAGGACACAGCTTGAATTTGATATTTTTGGGGTGAGCCCCAACTTGATGGCCTTGCACCTTTTATTTGCACAACCTATAAAAAGAATTCTGATTCCTCGACTTTTTGGCTCAGCTATACTAAGATGCCCTAAACTTGGAGTCCTATCACACATAATGGTTAAGGTATAGTGTCACGTTTGTAAATAAACAGTATCATCTCTATTCATTATCTTACATACAGATGTTGACTGACTGCCTCTATCCAAACATTTTTTCAAGATTGAATTATCCCATAGAAATTTATTTGTGGCACAGTAAACGTACAAGGATATTTCCGCCATCTGATGTTTTTCCTTCATGTAAATATTGGCCCTGTTGCCCAATAAAGTCGTTCGATTGGCATAGCGTGACATGGAGAAGTTCTGAAGCTTTCTGTAAGTGAAAATCCATTGTGTTTTGTCCGCAAAATCAGGATCTGCTGTATTTTGCATTGGTATGTACACTTTTTCAAGATTAAGAGCAGCCATTTGCCATCATTACCTCAAGGATGCCATATCTTATCTGCATCTGTTTGCTTTACGCTCTCTTGGAGCACAGAGCATATACAAATTTCTTTGGAAAATATTGCTATTCCAAAAAATCCTGGCTACAACTCAACCTCCTGCATATCTGTTAACATGCCTCCTTGTACACTGCCAGGCAATCTAGATTGCAAAGGTCCATCACTTTTACCAAAATGACTATTCTCAACAGCAACTGAAGATATTGTTGGTTCTATGATTGGTGGCACTGGCAGTGGTTGAAAATCCTGTGCAGCATACATATACCCATCCAACTTCATCCCGGTGGGAATTTTTTCCAATACTGCTTTGTTACCATTCGGCAAACCATGTTTTTGCATATAGGGAACTGCAGTAAAACTTGAATACAAGTCCTTTGGCAGTAGGGCCCCCTTCTCAGGTTGTATTTGACTGTTCTCAGTAGTAGTGCTTGCACTCATTGAATTTGAAAGGTCAGATGGACGCTCGCCAGAAGCTTCGGTGACAAAATCTGGTGTTGCTGTATGTGCAGGTATGAAAAATTGAGTTGGAGGAGGACTCATACTCCATCCTGTAGAAGGTACAAGAGGTAACTCTGCAGACTTGGCTGGTAATGATCCTCCTTTATTGAAGGTGTCTACGTATCTTGATCGTACTGCCTGAAGGCGGCCACGAGCAGAGAATTGATTGCCACTTGGTGGCAAGGGTGGAATTTCTGTGGATGAACTGAGCCTACTCCCAGCAAAGGAATCAGGACCCTCAAATTTAGCACCACCATTTGCATCATGCACGGCACTCCCACCATTGGATTTAAAGCCAGCAGTAATTGGAGGCGGTGGAAGAGCAACATTCTCCAATGTCTGTTCTGCGCCTTCCTCGATCCATTTTTTAAGATCTTTGTTGTAATAGAATTTGTTCGGGTCTCCTAAGTTTGCCTTTGGTTTAATCTTTATGAGCTCTATTGCTTTAGAAAATAACTGGCTTCCAAAACGCCCCAAATAACCAACATGACCCGGCTCAAGCTTGGTCCCTTTCGACGAACCTCCATCAGTTTCTGAACTTGTGTTAGTACTATCTTCTTCCTTCACTGGGCTCCAATTGAGGTTAGGTTCTGATACACTTCGTGGCGGTGCCTCCACTTGTTGATGATTGAATCTACTCGGACTCGCGGTCGGAGCTGTTGGGTCTAGCGTACTTGCTTTAACAAGCTTAGCTGAATTATCCACTCTTTGATCAATTGAATACCAGTCCCTCGGGTTCTTGTTAAGTGAAGGCTCAGAAGCTTGCCTTGGGGGGAGACCACCAATAATCCTATGTAGCGTACTGTCAAGTGTTCCAATAAACTTTCCAACAAGCTTTCCAGTGGCAGAATTTAAATTGTACCCACCCTGAGAATGCAAGCGTAAACGTTCCTCTAAAGCTGATGCAGATTGTCTACAAAACTCAATTTCCGGACCACGACCCACATTCTTCAAAGTCTTGGTGACAGCCTGACAATATCTACGAGCCTCTGAAACTTTTCCCGCTTCTACCAGCATATATGCGTACAATAACTTGTAAGGTTGAAAAGGCGAAACAATGAACTGTGGATTGCCCAAGGTTTTGGCATATTCATAAGTCTCTGTCCTTTGTATTGATGTGGGAGTAGCAAAAGTACGCTGCCACTTGAAATGATCCGCACCAACTAAGCACAATCTCGCGTTTTCTGAGAAGGGCTCGAAGCTGGCATCAGCAACTAGGTAACAAGTATGTGCCCCCGCAACCTCGCCATGATCCTTCCATAAGCAGTCTCCTAGATGGGTAATCACTTGTTCATCCCCATTAGTTGGATTGGCGGCAATAATTGACAAATTACCCATCCAGTCCCCAAGCATGCCTCCAGCGCTATCCTTCATAGAAGAATATAAACTCCGTGAATCAGGACCTGCAGCTACCACTGGAGAAGGCAAACTCTTTGTGGTAAAAAGCTCCGCTGTTTGGCCTCCAAATAGTAGAAATAATGTACGCAAAGGAGAGCCAGGCACAAACTGCTGTTGAGCCATTTGCAGCACTGTTTCTGCATATGCCTTCTCTCCCAAGTGCCATGCAAGAACCAATGCTGGGCCCCACATATGACCCTGCTGTGCAAGCTGAAGAGCTTCTTTTCTTTTCCCAGCTATCAACTGACTTTTCATTTCGAGCGATATCGCCTGAAGTTGATGTTCTGAGGGCACTGATTGCAAGCCAGCTGGAGATGCAAAGCTCTGATTATGAAGACTCTGACTACTTGTATTAGAAAGCAACCTTCCCAAATCAAGCTCGGGTCCCTCTTCTGACTGGGATTTAAGGTTGGTAGAACCATGCACCGACCGTAATTTACCGTAGTGCAAACAAGCAATCCTCAGCACACCCCAAAGTACACGAAGAGCTTCTGCACTTGTGCCATGAAGATCATCCGAAACGATAATGGCAATCTGATCATCAACCCATTTTAATAAATCTTTGGTTGCTATTCCGCCTCCTACTAAAGGACCTGCAAGACCTCCACGATTTAAAGAAGAAAAGTATGAAACCCCATCATTTAGTCCACTGTTTCCTGCAGCTACTGCCAGCTGATTCAGATCATGGAGCACAATTGTCTCGGGGATATTGCTGGATGATGCCTTCATTGTCACAAGTTTTCCACCAAAACCAAAAGCTACCAAGGGATGTGATGGGCGTTCAATTGTAAACTGCTTAGTTTCTGTAACCGGTCCATATATATTATGTGATTGATTAGACTGATGTGAGCTATGGAAGTTCAATGAAAACTTGTTGCCGAGTGTCCCGTGTGTCTCTGGTTGATTGTAAAGCTGTGAAACACCATTTGAATAACTTTGTAGTCCATATCCAGATCCATTGTTCTCAAATTGAAAAGACTCCTTCACCTTTTCCGTATTAGCACCAACAGAATCACTGGGGGTCGTAGCCACAGCTGATGGCTCCCATCCATGGAAAGACAAGTTCTGAGTTATGTCACCACCAAAGTCGTAGGCAACGCTGCCTTCATCTCGCAAGTAGGCATTCTTGTTTGTTTCGTGCTGATGTTCAGGCTGCATTACAAGGTTCCCTGCACCCTGCGATGTTTCAATGCTACTGCTATGGCCATGTAAGAAAGACTGATTGCCGTCAACTGTAACCTGCCGCCATTGTTGAGCTTGGTAGTCAAAATACCAGCCAGGATACTGTTCTTCCCATGGGACTTGTACCTGAGACTGTGAATCCCCAATGAAAGTCCTTTGTTCATATCCACCATCAAGAGTCGTTTGTTCACTCAAATCCAAACTCTGGCCTTCTACATAAGACCCCGATGAAGGCTCTTGCCCTGTATAAGCTGCGTGCACATGGACAACACTTCCAGAATCATCCATTGCCTTAGTAAATCCTGATGTGGATGTAGAGTTTTGCCCTAACTCCGTCTGTAATCCAGCAGACTGATTAAGGTCTGTGTGAGATGAAATTGCATTTGCTGTACTGGAAGTAGAATCCCAGCCTTCAATTTGTATCCATTGGTTACTTGTATAATCAAAATACCAACCCGGGTACATACTCTCCCAAGACTCTGGTACATTGGCCTGTTGCTGAGCCTCAAGACTGCCATTGATTGTATAAGAACCTCCACGAAGCTGCCCATCTCCCAATGAACTCTGATGCACTCCAGCCTCCGTGGACAACAGTTCATACTGTTGTGAAAAACTAGCTCCCTCTGTGTTATGATTGCCAAATGCTGCCTGATCTGCAAATGTCCTCCTCTCACCCCCTTGATCTATATCAGGTTTACCTACAGGCTGCTTGATAGTACCATTGTAGCTAAAAAACGCAGAATTGTTTGAAAATGCCTGAGATTGATGTTGGTTGGGAATGTCATAGATACCATTACCATGGGTGCCTGCAATGTCATAGCCTAGACTTACAGAAGCAGCAAAAATGGTATGCTCAGCCACATTCCCATTCTCAGCTTTCCATTCAAATGGTGAATTATTGTTGTCAAGTACACTCCCATTCTCAGCTTTCAATTCGAATGGTGAATTACTGTTGGCAGGAGGTGTCAAAATTGGCAAGTCAGCTTGAACGCCAAGCTGGTACCCAAAACTTTGATCAACTAATGCCCCTGTCCCTACATCCAGCTCTGAGAGAAAGTCATTCAAAGCATCGGGTCCTCCTCCCGGGGTATAGATTTCACTAAAACTACTCCATTGCAACTCTTTAACATTTGATTGAGGTGGAAAAGGCTTCTTGACTTGAGGGTCATCAGAGGGCACGAGATCCGATTGCAAATTAAGTGAATTAGCTGATGAAGTGGTGGTAGGCTCTGCACCTCCAGCAAAATGAGAAGGGTCAAGTTCATTGGGAGCGCTGCCATCGCAGAGAGCTAATTTGGGGTTGAAATCGTGATTTGAGGCAAGTGCTTGCAGATTGTTTGCTTCTGAGGGGGCATCAGCAAGTGTTAAAGCATCAGCTTCTGACACATTAAGCGATGCATCAAGCGAGGAAGTATCCGCCTCCAAGATGACAGTATTGAGATGGGCTACTGAATCAGCTTCTGACATGTCTTGGGTAGCCTCAACCAGACCATTTATGGTACACAGAGTCGGATTTATGGAAGGTGTTTTGTTGGGAAGAGCTATAGTTCCAGGCGTATCGTGTATCTGCGCAACATCATCGCTCATGCTTAGCTTGGACAAAGCCTCCCCCAAGCCCCCCTCTTCTTTGTAGCAATTCTGATGCAAGCCTTCTCCCTGCCCAAAATCGTGGGCAAGCTGATCAAAGAAATCCACATCCTCTCCTTCATTCGTGTGCAGAGCCAGCGATGGCCCTGCCAGAGCCATGGTGAGCTTAGCTAACCAGAGGAGGAGGAGAGATACAAGAAACTACACACCCCTGTGCGTTTGTAGCTTTGATCTCTGCCAATGGGAGGCGAGAACATGGAAGAAGAAAAATGCGATGCAGAAAGCATCACATCAAAAAATCCTCGTCCGAAGGCAAAGCTCGAAGCTACCTGTGTGTGTAGGTTGGTCTGCGTGGGTGTTAATGTGTGGAAACACATGAGAGAGAGAGAGAGAG

**PROTEIN TRANSPORT SEC23 G (SEC23G):**

GAGAGAGAGAGAGAGAGAGTTGAACGATGCGAGGCAACATGAGTTGAGAGGGCGGAGCGATGGACTTCTCGGAGCTGGAGGTGTCAGAGGGCTTGCGATGGACGTGGAACGCTTGGCCCTGCTCGCGCAAAGAAGCAGAGCAGCTGGTGATCCCTCTTGCCATAATGTGCTGCCCACTCTCCCCCATCCCCGATCTCGCCCTTCTCCCCTACGCGCCGCTACCCTGCTCTTCCTGCCACGCCCTCCTCAACCCGTATGCCCGGCTTGATGCTGCCTCCAACCTGTGGACATGCCCTTTCTGTTCCCGCCTCAACTCTCTGCCCCCCTCCTTCCACCCCCGTGATCATCAACATAGCCTCCCCGCTGAGCTCTTTCCCACTCACACTTGCGTTGAGTATCTTCTGCCCCGCTCCACCGACCACTATGCCATCACCTACCCGCTTCATCAGCACCTCCCAGCAGCCCCTGCATTCCTCTTCTTGCTCGACACCAGTGTTCCTGCAGAAGATCTGACTGCCTTAAAGGTGGCTATCTTACACACTCTCAGCCTCCTACCAGAGCATGCATTAGTGGGGCTCGTCTCCTTTGGAAGGCAGGTTCATGTTCACGATCTCAGCTTCACTCATCATTGCATGAAGGCCACTGTCTTTCATGGGGAGAGAGAGCTGCCTTTTGACCGGATTCAAGAGCAACTTGGGGTTGCCTTCAAGTCTCGGAAACATCCAAGCATGAAGCAAGGGGAGATGAATAACATTATTGCTCCCTTTTTGGATTGTGAATTCAATATTACAGCTATTCTAGATGAGCTACAGCCTGATTTGCAAGATTTCAAAGGAAGGCCTCAAAGAGCAACTGGAGCTGCTTTGGCTGTAGCAGTTTCTTTGCTGATGGGTTGTGTATCAAGTGCTGGATCTCGTATTCTGCTCTTTGTCGGGGGCCCTGCAACAATAGGACCAGGAAAGGTAGTAGGTACAAATTTTGAAGAGAGCATGCGTACTCATCGGGATCTTGTGAACGATAGAACTCACCATTCCAAAAAGGCTGAAAGTTTCTATAAAGGCATTGAAGGCATTTTAACCACCAATGGATTGGTCTTGGACATGTTTGCTTGTTCCTTGGATCAGGTTGGTATTCACGAGGTCAAGTCAGCAGTCGAAGCCTCTGGTGGACTTCTTGTTATGGCTGAGACTTTCAGTTCACATGAATTCAGAGCATCCTTACAACGGCTTTTTGCACGTGATCCAGATGGCCATTTGAAAATGTGTTTCGATGCAACAGTTAAGGTGATAGCTACTCAAAAAGTAAAGGTTTGTGGAGCTCTTGGGCCCTGCTCATCTTTGAGTAACAAGACCAGCTTGGTCAGTAACACTCAAATAGGTATTGGAGGAACAAATACATGGAAACTGTGCACTTTGACTGAGAAAACTCATGTGGCTTTCTTTTTTGAAGTGTGTAGTCAATTGGCAAATGCTTCACAACCTGGAACTGCTTTTATACTTCAGTTCATAACCAAATATCGAAGGAGTAATGGGCAGATGCGTGTCCGTGTTACCACCACAGCGAGGAGGTGGGTTGATGGCTCTCAAGTCGAACAACTTGTTGCCGGGTTTGATCAGGAAGCAGCTGCTGCAATCATTTCTAGACTTGGACTACATAAGGTGGTGCGGGAGGATGTTGCGGAAGTTATAACATGGCTTGATAATCTACTTATTAGGTTTGCATCCAAGTTTGGGGATTATAATAAGGAAGACCCTGAATCATTTCGGCTGTCTTCCAACCTTTCCCTGTTTCCCCAACTTATATTTTACCTGCGGCGTTCACAGTTTCTTCAAGTGAGTCGTAATACACCAGATGAAACTGCTTTTTTCAGGTTGATGCTTAATCAAGAAGGTGTGGTTGGAACTGTCATTATGATTCAGCCCACTCTATTTTCCTATACTCTTGATGGGCCTCCAGTGCCAGTCTTACTTGATATAAGCTCCATCAAAGCCAATTCTATCCTCCTCTTTGATTCTTTTTTCTGCATCGTGGTCCATTACGGATCAACAATTGCTCAATGGAGGAAACTGGAGTACCACAAAGACCCGTGCCATGAAAATTTCAAGAAGCTGCTAGAAGCTCCCGTGATCGATGCACAAGCACTTCTGGCTGAGAGAAATCCCATGCCTAGGTTTATCGAATGCAATCAGCATGGTAGTCAAGCACGATTTCTTCTAGCAAAGCTGAACCCATCTGTCACACACAGAAGTGAACATAGGGGAGATTCTGAGTTCATTTTCACAGACGATGTAAGCTTACAAGTTTTTATTGCTCATTTGCAAGAACTTGCAGTGAAGGATGATTACCTGTAAATATCTTATGCCCTTGCTCAATCATTTATCCCAACCAATGTCATTGCTTTTTTTCTGTAAGGTGTATACCCACGTGGTGCAACGTTTGTGAGTAAGTTGTATTCGTGTCAAA

**NUCLEOPORIN 155 (NUP155)**:

CTAAAAATTGTAACAAAGAGTAGATTGTGCAAAACAGGGAAATGCGGGTAAGTCATTAAGGACAGCAAAAGACGCAAAAAATAAATCTAGAGAGTGGATGGATATAGCAAAATGGTTGAAAGGAGGCAATCAACAACAAGAGACAATGTAGCATCTGCAGATTTAGAGTTAGAGAAAGAAATGACGTGGATGTCGGAAAAAATGCTGAATCCACTTTAGGAGCACGCGTAACATCGTGAATGTGCTTACAAAAGAAAATGAAATAGCTTCGTTTCGAGAAAAGAAAATGAAATAGCTGCCTGTGTTTCTCGAAACGAAGCTTGCCTTCCCAGGTTAACATACTCTTTGAACTCAAGAACTCTACTAACTAAGAATTCATGATTCTGTGAGTACAACTAATTTCTTTCTCCTCTTTACTTTGTTCCTTGTCCTCGCCTTTCAGATCTTTTAGCTGGTCCAGCACTGGGAGTTCACTTTTTGTGGCAGCGGCTCTAATGCTTTCCTTCCTTTTACAGCCTTTCTTCAGCTTTTTCTCATTCATTGCCGCAGATTCAGTTCACTATGGTCACCAATGTCCATCTTGCAAAAACTATTTCTAACCTTTTAGTGCCAACTGCCATTATGGTAGTCAAAAGTAGCTCGCATTCTTCCGTGTGCACAAATTGAAGGGCCCTCTTAACGGCCTAGCCAGAATACATCAATCCATTCACAAGTCCTCATTACTTCAACTGGCCTGTTCACCAAGAGCTCGACTTAATCAACCTTCTTGTCCACCTCGTAAAAGCTGTAGTAACTCTTCGATTAGGCTTGTTCAGAGAAAGATGAGAGGGAGTTGACGCCACACTAAGCACTCGGGTGATCTTGCTCAATAGGCCCTGAGTACGCCTCCCTAAATAGCAATATCAACTCAACCTTAAAAATCTGGGTTCCAATAGAAGGATGGACATCCTTGGCACAGTGATCTCCTAATTGTGTGCCGATGCCACGAAGGTGTCAAATCGAAGGACCTTTTGACTGTGGCCTCAATAAGAAGAAGATCCCATCAGAATTTCTTCCAGCTCCCGAAAGCCTTGATACAGAGGTTCAACCATGTTAGGTGGAAGCGAAAGTCTGCGAACCTCCGTCATATATCTGTTGGCAGCCGTAGTAAGTTTGTCTTTGACTCCTTGGTTTATGGAAGCCATTTGTCCAAAACTTAGTGCTCCTCTGAATGCCAAAGCAGTGGAACCTACTGTACTACCACAGACAGATGAGGACCATTCTCTCATAGCTGTAAGAACAGACCGCAACATACGCAGCCGCAGAAGAGGTGAAGGTACAGCAGCTCCTGTAGCCAGAAGACGATCATACACCCTTTGCACAAGCTCAGCTGCATCCTTGCAGGCAGCAATAAGGGCCCTTGGAACATCCTCATCTCCAACTGCTTCTTTACCAGAAGCCACACGGTCCTGAGCAGATTTTTCTAGATGCAGACAAATTGTGTCAAGAGGCAAACTGCCACCATCACCTGGAAAGAACCGAGGGCCAACTCGTCTCAGCACACTGCAGGCTTCTCCAACTCCACCAACCATTAGAGCTTGATCTATGAGCCTGGCCCATGTATCTTTCATAACATTGTTCTCAGAGTCCCGAGCATAATTCGAAAAGTGCAAGATGTCTAAGCATACCTCCCACATTTCAAAGGGCACAGCATAATCATTGTAAAGTTGTGAGATACTCTTTAAATCTCGCCCAAGTTCCTCCAGCTTTTCTTTGGCTGCACCAAGAACAACCTCTTCTGCAAGAAGATTCTTGCTAGGAAAAGATTCCGTCATAACCGTGTCACCTGGAACAGTTCCATGTGCTGCTTCACTTCTGGCAATCAAAAGCTCCAGTTCATCTTTAATTTTTAACTGGAAGCGCAATACGGCAAGCTTCCCTTCCAAAAGCTCTAATAAACCACCATCAACAACCTCAGTACTTGAGCTTGTTGTCCCACTTCCTATGGTTGCATTCTTTGCTTGCAAAACGGCACTGCTGAGGTACTGAAACCTCTGCTCTAGGGTCAAAACACCACCTCCATCCACCTGTCTTCTCTCTGCCAGTCGTAAAAGCACGTGGGCAGCAACAACATGCTGTCTGTGATGGACATAATAGCGTGCCAAAAGTTCCAGATATTTAGCCTCTTCTTCTGTCACTGGTTGCCTGGAATGGTTAACAGAAGATGGTGCAGACGGCATTGCAAGCTTCTTTCCACCTAATCCTGTCTTCATCATTTGCTTGCTACCAGCAGTTTGTAAGAAACAAACCAAATCTGGACCAGCAAGTTCTAGAAGCTCAGACTCTAAACCTATGTCAATCATTGTTTGATAAAGGAACTCATGGAAGGCTCGATCAGGCCACCTCACGCTCAGCTGAACAACCTGCCTTATGTACTTCTCCCTTAATGATTTGTCCAAAACAGGATGCTGCAAGGTCCCAAATCCAGACCGAAAACCGGATTGCGAGCCTTCTTCTTTTAGATGCCTTAGAGCATTCCCAACTATTTCATAGCATTGTTGACGAGCAAGAAGAGTCATTTCACGCCTTTTCTCATCAATATGATCATTGATAGCATCCTCGTTGGGATCCACAGCCTTTGCTTTTCGTAAAGGCAATTCAACGACAGCTTCATAAAACCTGATTTCTTCAAATCTCTGACAAACACTCAGAAGATCTGCCGATTCTGGAACCTTTCCAAGTAATGCTAAGGCCTCTCTTGCCAATCTCTCTCTTTGTTCTGCATCAGTAGCTTCTCTAGCCTTCTTCAAACACTCTACCGCTTGGTAAAACGTGTAGTCGGTCTCCCGATAAAAGCTTGGGCAGCCATCTCTCAGTTTCATGCTGATGTCATCCACTGAGCCTTGACCATTCAACCCTATGTAGTACTCCATGAGTGCTGCTACAAGACGGGTAGCAATTTGCTCTCCATCTTCTGACGAGCAGACCAGTTGGTGAAAAGTAAGATGAACCAGCTTGTGCCGCAGATTAACATCGAACAGCTGAGCTAACCGTGCCACATGGTGCTGACAAAGCATTTGCAGCAGCAAAAGTGCCTCTCCAGACCTCCGAAGAAGCCTGCGAATAAGCTCCATACTCCTCACCTCCATGACCGAAAGTTCTGCAGAGCTGTAAGGGACCCTCTGCCTTTTCTGTGTCAAAGGCTGTGAGCTAATTCCTTGAGGTGCATCAAAGAGATTCTTAGAAACCGAATCCAAACTGCTGTTTCCATAAAAACTTGGAAGGTCCCCAACCCCAAAAATCCGTCCATATGGTCCTCTTCGTTGATTTCTTCTTGCTCTGAGAAACTGTTCTAGCCCCCTTAACTTATCTTCTAAGGCTTGCATGGCTTCTATTGATAGCCTGCAGACGATAATGCCCCCATCCTTATCCTCAACACTAGATTCACCTTTGAGGACCATGACAGGGAGCTCCCAAATTGGCCAAAGCAATCTTGCAGTACACAAGCAAAGACCTTCGTGCGCTCCAGAATAGACAGGTGTGGCTTCTTGCACAACTTGCCCCATGTTGAATCCCATGGCTGGTGATGTGATTGGACTAGTGCTGGGACCGCTTCCCTGCATCTGAGGAACGCCTACAAGCCTGGGATCTTCAAAAGCTTCAGCAGCCCTCTCGACAACAGTTGATGGCACTATCTTTCCTTCCATGGGAGTTAGTCTTGCTGCAAGAAGTAGGCACATAGCTGCAGCCTCACCAGCTCCATATCGATGAAAGAAATCTTCCAACAAAGAATGTGACATCCTATTGTCCAGAATCTTTTGCAACACATCCACAGGTCTGTTAAATACAACTTCCATCAATCCCATATTTGTGAAGATAACAGACCTCCTCCGAGGCAAAGCATGTTGTAAAGCAAGCTCTCCACGAGCCCAAAGCCTTTTTGCTCTAGTAATTTCGCCTTCCATGCTAGTACCGTCACCAGCATCCCAAACAGCAGCTTCAGCTGTTGTTATATACTCTGGAGCTGGGAGAGAATCAGTAATGGCTAAGGTTCTTCCATCAACTCCTAACATAGAAACCGTCTCTCTCAGAGCCCTATAGCCTCCCCCAGTTGAAGCAGAGCTTAGAGTAAAACCCCCTGATGGGGTTGTCAAGTCTGTTGTTACGAGCAATAACCTTGACATGGCTGCAGGAGAGGAATCAGAAAGGAAGAGGAAACCACTAGAATAACAAGCAGATTCTACTTTTAAAGTAAGGCTTTCAGGTTGAGGACGACCGCTTAATGCTAGGCTTCCAAAAGATATGACTCCACCAGCTGCAACTGAAGGAGCTGGCCTGCTCATGACAAGCTTCAATGTAGCCGGCCTCAATGAGCTTCCAGTAGACCCATTGAGCCCTCTGGTTCCAACTGAAGAGAAAGTAGACAGATACATCCTCCGACCATCAGATGTAACCGCCACAAGATGTAGAAACTTTGATTCTGCTGAAGATATGGTAGAAATTGAAACAATCGGAGATTTATGCACCCGGGATGCAGCTCGTCTTCCAGACGGACGGCCACTTCCCAATCGTCCATCTCTTTGATCGCCTAGATTTCTCTCCTCTGCGACCTTTTTTAGGGAGCTGTCTCCAACTTTCCCCATATCAAAAACCTGTATCTTTGATTCATGAGTGCGTGTGTAAAGGATGTTTCTCTCGTTATCAACCACCATCTCTACCACTGGATCCACCGCTCCAAAGGTGAATGAGTTGGGCAAAACCCACCTTGATAACAGGCTGCCCAAGCCACTGGTATGACAGACTTTTCGGCATCGGCTTTTCCAACCTGCACCTGAACTGTAGAGTAGCTCATATATATGGCCATCTCTGCCTGCGAGGAAAATACGACCTTGATCTGTACTCTGTATGCAGGTCATTGTAACTCCATCTGAAGGGATTGTATACTCATGTAACAGTTGCATAGAGAGCTCGGCATATGGATCAGAGCCATCTCCACTTCCGGAACAGCAGACACCAAGTAATACAACCTCAACGGGAGTAGCTAATATCAAAAGGTGCTGAATGGCCTCAACAAATATGCCAGGCTTAGCTTTGGCAAGCCCAACTGCACATAAAGCTTGTTCCTCTCCACTGTACTCTGGACAAAGCCCATCCCACTTGTCAAATCTCCACAGAAACAGGCTGTTGTCAACGGTTGCCCATGCTCGCCGGATGCTAGGAAAAACACCACACAAGGCCGTCCCCTGGCCACCAGCTGCATTATATCTCTCAATTAGCACAGGAGGCAGCTCTCTGTTGCTTGCCACATCAACAACTGATGGCCACTCTTTGGGAGGAGAAGAGTAAGGGTTGCTGGAGTAGCGGGCAGCGTCGAGCACCTCATAAAGATCTGGAGATGCCAAGTCGCGCGAAATGGTCTCTGTGATGAGGCGCCCTGCCGCAGCCACATCAGGGCTGATGGTTTCATCGTCCCAGCTCATATTAAAGCGCTTGCGCGCACACACAGTGCTGAAGTGGGCTTCTTCTACAGCTTCTATTAATCTGCGTCTGCGGGTCAGCTCTCTCTCTCTCTCTCTCT

**NUCLEOPORIN 160 (NUP160)**:

GCAAATTTTTGAACTCTGTGAAGTGGGGTTTGGCGCAATTGTGGGCGAAGAGTAGCGCGTCATTCAGAAGCTGAAAAGAGCAAGATGCTACACAGAACTTTAGCAGGACGCTTTTCTTAGAGTACAAGGGGAGCGAGCACACATAGAACACAAATATACGCATTTTGAGCTTGAGGGTGTCCCACACAGGAAGCCATGGAGGTGCCTGTGGTAGGCAGTGAGCAAATGAGATGGGCACGCATCACAGTGCCCACCCAATTGCCCCCCTCAGAGCCTTGCTGCCCCCTCTCCCCCGACATGGCTGGAACTGTCTCTTTTTGTTCCCCATCTCTCCGCTTCACTATCTGGCGAGTACATAAATCATCTCCTGCTACTTTGGAGCTTTTTGAGCTGGCAGATAATGAAGCTCATATGTCAGGCCTACAACTTGTGTTCCACGAGCCATTGTCTCCATATATTGCTATTTACCCTGAGCAGGCTCAAATGAACTATGAGGAGGGGAATCTTTTCATTTGTGTTGTCACCATCTCGGGGGTTGTGTACATATTGAAAATAAGAGCTTTTCACAAGTACAGAAGTGTCTCAGCTGTCACAAAGTCGGACATGTTTCAGTTTTCCCTTGTAAGTGAACTTGCACGTCTCGATGAGATCACTTCCTTTGCTGGATTAGGAGGTAGTATCTGTGCTGGTGGGAAAAGTGGCTCCATTCTTTGTTACCCTTTTGAACATTTCAACATGGAAACCAAAGAACACTGCTTTGAGCTAAAAGACTCTGATGCTATTCTGGGGCGCTTGTGGGGATTAGTGGCGAGGGTTAAAAGTGCTGGTCCACTTCAGAGCCTGATCATGCAACTACATAAAGAAAAGACATTACTTTTTGCACTACATGAAGATGGAACATTTAGGTTATGGGATCTTCATGATCGATCTCGTCTTCTGAATTTTAGTTTGTACAAAGGAGATTTTACAGACTATATCCCAAAGATGATGTGGGTTCAGGAGCAGGATATACGTAATGGGTCTAGGCAAATTGCTTTCCTTTTCCAAAAGAAGGGGGCAACTGTGGAAAACTCAGTTCACATTTATGACTTGCAACTTGATCAAGTCGATGGAGCAGCTGGTCAAGCTACACTGCATTTCCAATTACAAACTCCTGACATCCAGGGGACCGTGCTGGACCTGAAGTTGCTCTCGGATAGCTTTTGGAAACTTGAAATGGTGGATGGACAGTTCAGCCTCAAGTGCTATGCAACAAAGAGGGAAGAGCTGCTAGAATCATGTGAACTTCTGGAGGCTGAATTGGAAGATCAGCTTTTTCAAAGCCCTGATAATGATTGGGAGGATCTTTTTCTTTTTCAACAAAGCATCAACACAAGTGCAAAGACTGAGTTGCGGCTTCCCATATCAGAACTGTTTTTCTCAAGAATCTTGCAGTCTGGAGTTTACCAACAACACACTCTTCGGAAAGTGTTGGAAAAATATAAAAGAAATATTTCCGAGACAGATTTTGTAATGTTTACAAGGGAAGGAATACGAAAGGAGATCTTGGCAGTTATAGAGTCAGAAAACACGGGGGATGACAAGATGGACCGCCTCGTGCAATGGAAATCCTTTTGTAGGGATTATCTCACATCATGGAAACACAACAACACACCATATGCTTTGATGCTTGATTATCACACAGGAAACATTGGCCTAATCAGGCGGCAGTCTCTATCTGTTAAGCGGTCTTTAAGCAACTTTGAAAAAATGTATTCAGGAGATTTCAATGAAGGCGGTTTTATGGACTTGATTTTTAAATGTACTTCAACCATTGTCAGACAGTTGGGCAAGCTCCCTTGTTCAGCTTTTGGCCAAGTGTTAATAGGGAACAAACACATCTCCTTGTCTCACATCTTTTCAGTATTTGTGCACATGTTGGATGTGGGTTATGATCAATCTACTATCAACAATGGAACTACTCATATTGGTGTGGATGTTGGGAGGAAAAAAACATTCGAGTATCGCCAGGGGCACCGGAAGTTTGAGCTACAGCTGATTCTTGATTTGCAGAACCTTAGAGACTTTGCAGGCAGCTGGGGTCTTGTCCTAGACAACATAGAGAAATATATCACTCAACTTACACAAGATACAAGGTTCTTTCAAAAAAAGAATTATCCTGAAGGTTTGGAATATGGAGTAATTAGCAAAAGGATGGTATTAAAATCAACATCCCAGCTCTCATGGACACACTTTGAAGCTGCCAAACATCTTTTTTTTTTACTTGGCTACCTTATCAAGATCAAAGGGCAGGTGGGTATCAGCAGTATCGAGGTTTCAAGAATACAGTTGGATCTTGTCGTGCAGCTCAAGAGAATACTTTTGCGCAGTCTTCTGGTTTACTGGTTGAGTACTAATTTTGCAGAAATAGCACCTCCTGAGGACTTCAGCTTACACTTATCAACTTTGCGCCTAGATAGTGTGGTCACTAACAAGCCTAACAATCAACACCTTGGAGTACAAGAGACGACTTTAGCAGAAATACTATTGTCTGCATGCTCAAAGAGTCTTTCAAAAGGGCCATTTCTAAGATCCAAAGGATTTTTAAATTCTGATTATCTTTTGTATGCTACAAAGAGTTTTACTTCCTGGCTGATGTGGGGGGAGGATAGTGATCAAAGCGAATCTCTCTCCAGTCGTTCTGTTACTCTGGCTGCAATGCTGCTACAACATGGGCAGTTTACAGCTTTGCAGGACTTTTTACAACTTGTGGATGCTATGTGCAATGAACTGGAATTGCATGAAGTCAAAAGCTTAAAGGGGGAATGGTGTGTGCGTCTCCATCTTCTGGGTTGTGGTTTACTTGGACAGACTCAAAGTAGTCTCAAAGAACCTGCAATCAAAAGTAGAGTGGAGAAATCTGTTCATTGCTTTTTCAGAGTAGCATCTGTGGTGGGGCAGGATAGGGAAGACCTTCAGGCACTGCTTTTGAAAGTTGGAATGGAGACGAGCTTTCTAGTTCATGGGGCCATTTCTTCCTGGAAATTTCATTACTATGAATGGGTCATGCAGATATTTGAGCAGAATCATCAACTTGAAGGTGCTTGCCAATTTGCTCATGCAGCTCTGCAGGAAGTGGATGAAGCAGCAAAAAGTAACATCATAGAAACCACACTTGAAATGTCAACTGCACTGTTAACCATTAAAGGACGTCTATGGGCCAATGTTTTCAAGTTTTCTTTAGATAAAAAGAAGTTTAAGGAGGCTTATTGCGCCATCATGTCAAACCCTGACAAAGAGAGCAAATATGTCTGCTTACGAAGGTTCCTCATTGTCTTGTGTGAACAAAAAGCAGCAGAGGTCCTGTGTGGCTCAGAGCTACCATATGCAGGGATGCTTGAAAGAATAGAACAAGAGCTTTTGTTGAAGGCACACCACTCGGACCTAAGTGCCAATCCAAATCCATTCAAGCTTCTGTACTCATTCCACATGCAACGGAGCAATTGGAGACGTGCAGCTCTCTACATGTATCTGTATTCTGTTCGCATGAAAGATATAGGTCTAACAGAGGCTGGTTTGTCAGAGAAGTTTGTTCTGCAAGAGCAGATTAACAGTCTTGCTGCAGCTATCAATGCTCTGCACCTGGTTGACTCGATGTATGCATGGCTCGACTTGGAGTCAGATATTTTTGGTGGTATAAGTGGCAGGCCATCTTCCAAGCGCATTCGTACTCATGCTAGTAACGATGCTGGCTTTTCGACGCAAAGGAGCGAAGCCTCACATGTTGATCTTAAATATCTACAGAAGGAGTATACACTTACACTGGCCCAATTGCTGCTGATTGAGGCTGGGGTGCAGCTGAATGTGCATGGAGGCGATTTGTCACTGGAAAATGTGCTTGTTCTACTTGTCCAGTCCGGGCTCTACGAAATGGCGTTTTATTTTCTTTTCCAGTTCTGGAAAGACTCGACTCTTAAAAGCGAGCTGGAGCGGATATTTCGAATAATGGCAGAAAGTTGGTGCATGCGCCAACAAGACAACAGTAGTCAAAAGCCATTGTCAAGCCATATGTCACTTTTAGAACCTAATAGGAAGTTGCTGTCATTACCGTCTTCAGAAGCAGAGGTCCCATTTGAAACACCAAAGAATTCTTTAGATCTTCTTACCAAAGACGCGATGAGCAACACAACATGCAAATCCTTACAATTTTGCCTTGATAAGTACGTGAAATTGCATCCCCATTTACTTGTTGTCGTGGCAGAGACAATGTTGAATATTGACAGGTTGATAGAGCTTCCTGTCTGGCTCGTGGATATGCTTAAGGGTGGAAGACAAGCTGGAGAGGAAGGTATGAGTCTGGCAGGCGCAAATCCAAGTGCTCTACTTCGTATCTACTTGGACTTTGGTCGGGTTACAGAAGCTGCTATGCTGTTTTTGGACTATGTGAAGGCATGGTCAAACCTGTCACCTGCGGATGTAATCAAGCGCAAGAAGATGTGTGGAGTGTGGTTCCCTTACACACTAGTAGATCGCTTGAAAGCTTGTCTTTCGGAGTTAAGTAAATCTGAGCAGGTTGAAAGCCTTGAGCACTTGCTCGAGAGCAGTTTGTCAAGGCATTTTGAGCAGGTTCATACGGATTCAAGTGATGTGAAGTCATTGGTGCAAAGCTGAGTCCAGTTTCTTTTTGTAAAAGCAAGTGATTACATATGGTCGGTGGGATGATTCATGATCCCTTCTCTTCTCAAATCAGAGGCAACTGTTCTGTGATGTATTAGCTTCCTGTAAAATTTGCATTTTTAAATGATGTATTATTTGGCATATCAGAGGTAACTGTTGTGTGATGTTCCTAGTCCTAGCTTTTCTTCCAGTGCAAGCTACAACACCTAAGTGCATCTGTAAAGAGGGTTAGTTTAGAGGGGAATTATCATGCCTTTTGTGAAATTGAATGTATGTTTGAATATGCACTTGAAATCATTTATGTTGAGCTTTGAAAATGTTCTCTCTTTT

**TRANSLOCON AT THE INNER ENVELOPE MEMBRANE OF CHLOROPLASTS 100 (TIC100)**:

GAATCTAGAACCTCAGGAATTATTTAAAAAAAATCTCACCTGTAGTGCACATCACTGCCACCATTAAAACCAGACCATCGATAAACAGATTGGTACACTGTTACAATCTAGAAGGGAGAGAAATGTGAATTGAAGAGGTGATATCTCTCAGCCGACACTTTCGTGCGCATTTTATATGTAGGCACAAGATTATTTCAATGCTTTGCACTTAGTGCCTACCAGTGCATCAATTAAAGGCTATTTAAAAGCCCGGAAATGGGAAGGCGCCTTTTAACGCCTTCTAGGATTTGTGATGAAATACTAACAAGATCTATTAGAGAGCCAACAAAAAATTGCCCTTGTAGAGTGTGGAATACAATCACGAGGGCATTTTATGAATCAAATGGCCAGGAGCTCACATAGATAAATATGTTTTTCAAGCTGTTCATACTACTGCTTAACAAGCCGTGATAGCTTCCAACATAGACAGCTACCATTAAAATCGAAAAAGGCCTGATTTTCATCTCTCCTATACCCTTCCAAACTACTCAAAATTGATGAATTCAGATAAAAAGGCTGAGATCATTGCTCTACTGGGAGTGCCATAGAGAGAAACCCAAACACTGGCTTCGTCCAGATGGTATCCATAGTGCTTTGTTGTTGTGACTTGCTGCCTTGTTTGCTTGAATAGTGTGCAGCACCAATACCTCCAGGCTTTTTAAGAAAACGTACTCGCCTTCGCTTGCTACATTTGGTGCCGCATACCATTTTGCAAGCAGATTTCTGAGACACACCGGATACTCTCAACCTGAGATCATCAGAATTGCATGAAGGCCTTACATGTAAAGTAGTCTTCTGTCTTGGGGTGGTAAGAACCGACTGATGTTCGCTACCCATATTGCCCAAGCGGAATCGGTGATGCCACAAGTCAGCTACTACTCTACCTACTTTAGATTTGAAAGCTTCACCTACAGCCATTCGTGGTCCCAATGTCAGAGATGCAAAGGCTGTGGGGAATATGGAGCCACGTTCTTTGGACATGTGCTTGATATCCTTCAATCCTGACGGATCATCTTCGATTCCGACAAATGCCACTTTCCCAAAACTTCTTGGCTTCTTGTCATCATCATTATCCTCCTGATCCTCCTGAGAGGTATTTGCTTTATCCTCTCTGCTTGCTTCAGCCTCTTTGCTTTCATCGTCATCCACCTCTGGACTTTTGGCCTCATCTTCTTCATGCTGCTCTGGAAAAGACTCATCTGACTGGTTTTTATCAACAGCCTCGTCTTTGGGAGCAGATTTGCCCCTCCTTTCTGCACGAAGCTTTTCTCTTTCTATCTGAAAAGCTAGATTCCGCTCTGCCCTTTTCAATTTTCGTCGAAATTCCATAGCTTGGTCCTCCTCTTTCCACTTAACTTTTAATTCCTCTCTCTTCTTCTCCAGATCCTTCCGAGAGTGCTCACTGGATTTAGCCCATGTTTCTTTGAGCTTCTGGAAAAGATTCTTCTTTTTGCTTCCAGCATCGTCTGTCTCATCTTTTCCATCTTTCATATATTCATCAAAACCCTGTGGTAGTGGTATGATAGCTTCTGGAATCACCTTACCATTTTCAGTTATGGGTTGAAAAAAGAAGCGAAGCTGCCCTTTAGCATCAGTTATCCAATTGATAATCCGCCCAGTGGGAACATGAAGAAGAACTTCTCCTTGAAGGTCTTCTGAGGCCCTTTCCACATTTTCTGGGTTGAGTTCAAGCATTGCTGACGGTGTGTTTCCCAAAAGGACAAACTCATCCTCTTGTGACCAAATATACTCCAATTCTGGTGGCATAGGAATCACAAAGGGTGCCTTCATGAAAGAGTTTAACCACATTTCTCTCTCATTGTCTAAATCTTCGACATATACTTTCCAATCTTCAGGAACCTCATAGTATGCATTTATAAATCCGGGTGCCATCCATAGGTCCTCCTCTTCATACATGTAAGGATGTGAGGGATCACAAAAAGGGCCCTTTACTTGCCGTACCATTCCATCTGGTTTATTTGCAAACATTCTTGCCTTTGCAGCTGCCACTTCCCCCAATGAAGCATGGATAGCTGACATCTCCACATTACACTCCTCTGCATCAGGTAGCAATTCCCCAAAATAAAATTTACCCCAAAGAGAACGACCATTAATTTCATAAACTCCACAACCATGAAAGCGATTATGCTTCCACTGCCCTGCATATTTGTAGTGCCCTTTTTCCGGCAATTCACCGAACTCCTCGATCCAGCCTTGTCTTTCTTCCCAGGATGGCATAGTTCTACCATGCTCCTCAAACTGTTCTTCAATATCCATTTTCAGCCATTCACGATCGAAAGGGTCCATGTAATCTGACCTCAAAATCTGACCTTCTTCCTTGGCCTTCATTGCTTCTTCAGAATCCGGTATAGGCTCATCAACAGGAAGATCAACTTCGATCATGCCATGCCCTTGCATCATGTTTTGGAACCACTCCCCTTCATACCTGCAGATGTCGAATGCTGTAGTATAAACACCCTTTCCATGTGCAAGGTCATCCCAGACGGTACCTTCATAACTAGAGCCATTCTTGAAGAGGTAACTGTACCACAGAAGAAACTCTTCTGTGCGTTCAAGCTCTTCAACAACGTCTTCAGGGGTCTGGATATCAAAGTGGTTATCAGGAATGGGGGGGTAGAATTTGCGGTATGGAACATAATAAGGGGTGCCGGGGGGCGCGGGCTTGCCCTCGCTCCAAAGCTTCTCATACTTCTCCCAGGAGGCCTCGGAGACAAAGTTAGGGTCCCAACCAACAGCACGCGTATCGCCCCAATAGTCTTCCCAGTGTTCACCCAGGTCTGCTTCAGTCCAATGCTCGTTATCTGGCGGGAAGTTCCATAGCTCCTCCTCATCCACTTGGTCATCATCCGCTTTGGGATCCTCTTTTTCTTCTTGCTCCTCCTCCTGCACCTGCCGCTTCCCCTCTACCTTCTTCTCTCCATCTCCCGTACCCTCTTCTCCTTCTTCGTCTTCGTCTGAGCTGTCTGTGTAATCTGAGTCATCGCTACTGCTATAATCATCTTCTTCTTGCACTTCTTCTTCTTCTTCTTTCTTCTCCGCTTCTGGTTTTGAGTCCGCCATAGCTGCAAGGTTCTGAAAGGAAAGAGAAGACTCTGCTGGCGGTGAGGCGCGCTCAATGCAGCATGTCCTTACTGCCCATCTTGCAAGCAGGCTTCAAGAGGGCGTATGGCTCCGTGCAGCACCTTCCTTATCCTCATTTCCCCTAAACCCTCAGCTTAATTTTTTGATGTCCCACATATGGCCTCGTGACTATCTCCCTCCATCTTGC

**TRANSLOCON AT THE INNER ENVELOPE MEMBRANE OF CHLOROPLASTS 55 (TIC55):**

ACACAAACACAAACACACACGCACATTTGATCACCTTCAATTTACTTCATGTGGGTGATGGAGATGGTGAAGATGATGGCGATGAGCTGCAAGTTTACAAGCGCAGTGCAGGCTGTGTCGCATTGCGCCCTTCTTCCTCATCTCCATCTCCTCACAAAGCTACCCTCGAGCTCGAGGTCCATCCATGGCAGTCAGAGGCTTGCCTTTGCTCCTCGTGCCCTCTCCTCACGCGAAAACACAGTATTGCTACAAGGAGAGGAAGGAGAAGCAGAAGAAGAAGAAGAGGAGAGAACAAGTTCTCCCATTGCGAAGGTCGACTACAACTGGGAAGAAGAATGGTACCCTCTCTATCTCACACGTGAGCTGCCCCGTGACACCCCCCTGGGTCTCTCCGTCTTTCACAAGCAGCTTGTCCTTCATTATGATGGCCATGGTGCCCTTCGCTGCTATGAGGACCGCTGTCCTCACAGCTGAATGTTGTGGTCTACCAAGCTGAGAAGTTCATTTATTCTCAGATTGGCAAAATTGTCAGAAGGACAGATTATTGATGGCAGACTAGAATGCTTATACCATGGTTGGCAGTTTGAAGGAGATGGTCGATGTGTGAAGATTCCACAGCTACCTGCAGGAGCTAAGATCCCCAAAGCTGCCTGTGTCCGGTCATATCCAGTTCATGATTCTCAAGGGGTAATATGGGTATGGATGGGAGACCAGGGCAAAGCTGACATCAAAGATATTCTTCAATTCAAGCAATTTGACGATCCAAACAATTCAGAAATCGCAGCAATTCACGTTCTACCCTATGATCACTCCATCCTCTTGGAGAATCTTTTGGACCCTGCGCACATTCCCATCTCACATGATCGAACAGATCCATCTGCAAAACGAGAGAATGCTCAAGCACTTAAGTTCGAGATCTTTGAAAGAACAGCGAGAGGGTTTGCAGGAAGGTGGGGAGAAGTGAATAAAGACGATTTCACTTTCAACTCTCAGTTTGAGGCCCCTTGTGTTCTCTGGAGTGAAAGGGAGAGGGTGGATAAGGACGGAAACAAAATTAAGTTCACAGCCATATTCCTCTGTAGACCTGCCGGGCAAGGACAGTCAATGCTAATAGCACGATTTGCACAAGTTAGCAAGAAGAATCTTCTTTCATTTGTACCAGAGTGGATTATTCATCAAGTTTCTAATAAGATTTTTGAACAAGACATGGGGTTCCTAGCCTCACAAAATGAAGTTCTTAACAGAGAGGGAGTAGCAACAAGCCATTTATATCTTAATCTCCGGTCATCAGATGTTTTTGTGATGGAATATCGGAAGTGGTTGGATAAGGTGGGGCATGGAATGCCACATTATATTGGCCATAAAAGCTTTTCCATGCCAGCTAAAGAAGCCATCTTGGAGGCAGCTCCTGCAGGTTTAGGGGCAGCAGCTGCATCATCCCAACCCATGAAAGGGGCCACTGGGTCTCTTTTTGCGGCAGATTCAACAAACCGATATTTTCGTCACGTTGTCCATTGCTCGAAGTGTAGGCAAGCCTTTCAAAATCTAAAATCTGCACAGAAGGTCTCCACTGTTTTTGCTGCACTTTCTTTCTCTTTAGGGGTGGTGCTCTCACGTACACCGTGGAGAGTGGCTCTTGTTGTTGTTGCCCTGCTAGCTGCTGCAGGGTCATACTCTTGCCAACTTGGCATCAATATGTTGACCACAAATGAGTTGCGGCCACATCGAAGATGATAAAAATTGTTGCCTGTTGTAAGGTAATATCTGAGTTTCAAGAAGGCTCATGAATTATTCTATTTGCTTGTCAAATATGTAATGCAATGTCTTATAAATCTAGTTCTATTATAAAAGATGACCTTTCTCATGTTGCCTTTACATAAAGACTCTTGTACACAGTTTGGAAATTCCACAAGAATCCTCTTAAGCCTAGATTTATAGCAGCTTCTTGTAATACATCTCTTACAGATGTCTCTAAATGGCTTAGCCATTGCTTTAAAGCAATTTTACCTAGTGTTCATGAACTTTGGAAGAATCTTCTTGAAGAAGTTGACATGCCTGTAACTTCTAGTTGGGTTTTGAATGACTCAATAGGAGTGATAAACCATTACTAAAATTTTCAAGCAGTAGAGAAATTATCTCCTACAAATCTTTGATTTCACAACCTACACCAAATTAAACTTGAATGATCTAAGGAAGACACTAGGCCTTTTTTTACGAAAGCTTTTGCCTAACGTTTGAAAATGATAGTTACAAAGGCTAGTAGAAAAATCAGCTTTGCATCTTGAATTTATGTGGCTGAAAAACAGAAAAGTAGATCTAGATAAAAGAAAGAAAAGCATCACTAAAGTTGTCGATAGATACTAATTGTGTTGTTTTTCTTGCTTACTTCTTTCTTTTTACATATGAGTGCGATTTAATGAAATGTCTTATCTTTGTCCATTCGTTCTTCACAAGCTTTGTAATGTAAGGGGCTTAGTAAATGACCTTTTTGTTCCCGATCCTCCTTTCAACAGTTCATTTTCATGAATGCAGGTCGAGGAATCTATCTGTGAG

**TRANSLOCON AT THE INNER ENVELOPE MEMBRANE OF CHLOROPLASTS 56 (TIC56):**

ATTTATTGCCGCCAATGTAGGTGACGAGGATTAGCCGCCGAAGGCGGCCGTGTTTGGCTACGCAGAGATCGAGGGTTAAGATTTTCTATCTCAAGGTCTCGGGTATCCTCTCTCTCCTCTAAGTGGATTGGGTTTTAGCGTGCAGAACTCGAAAGGAGTTCGCGAGTGAGAGATGGGGTTCTTCTTCTGCAGGAAGAAGAAGAAGGAGGAGCAGCCGAAGCCGAAGGAGGATACCTCTGAGAAGCGCGCAAGCGTATTCAAGGACCCGCTAAGAGTGAGGACATGGCAGGAGAGGTTCGAGCAATACCAGCTGGAAGAGCTGCACAGGCCGGACTACACCAAGCTCCCAGAAGTCGAAGCAGAGGCCAAGAACACTGGCTATTTTGCAGACCGACCATTGACACCCGCCGAGATTGCTAGACACGACGAGATCATTTCAAAGCTTCCAAAGAGTGAAACCTTCCAATTCCTTCTGGGTTGCGAGCAGAGGCGCCTCAAGGAGAATGCAGATGCCATTGCCCAAAACTCACGTCCTTATCGCCCTGAGGACAAAGAGCTTTGGCAGGCCCTCCCCCATGTTCCTGCCCCTCGTACCGGTGTCCCCATGCCTCGCATGGCCATCAAGTCCCATGCGCAGGCCCAAGCCACTTTCTGGGACTTCTTCAAGCAGTTCCATTTTGGCCTCTGGGGCTATCGTCAGCGTCCTTACCCGCCTGAGAAACCCAATGACATCCAGCAAATGCTCGGCTACAAGTGGTTGCACAAGCGTTATGCAGATTTTACGATGAGATCAGGTGGATGGTATTACAAGGATCGTCTGGGAAGAACAAGAGGTCCCATGGAACTTGTGAATATGAAAACAGCATGGGCTGGTGGCATAATTGATCGTAATACTTTTATATGGGGTGATGATATGGATGAATGGGCTCCGATCAGCATGGTCTATGGCGTTGAGGAGTGTGTGGATACGCTCGATATGAGTTTAGCTGCATCAGGGCTTGACTTCATTCACAGAATCACGAGAGGATTGCCTTTGTGGACCCCTAAAAAAGGTCATGAGAAGAAAAGCTTCAAACAGCTACAAAGTGAGGCTCTGGAGAAGAGGGAAAAAGAGAAGGCTGTTTTAAGGCTTAATGATGGAATATGGCCTGGAGAACGTCCTCCTAGCCATAGTCTCTTTTTGTGGGCAGGTGGCTCAGAGCTTACCACAATTTTGGAAGAACACTATAAGTATATGCCAGATAAATTTATTCCATATCATTTAAGGAAAGAGTTGGCAGAAGTGATTCCTGGACTTCGACCCTGGGAAGTCTTAGATGTAGAGCAGATTATGGAAAAAGTAACCTACGGGGAAAAATGGTATCGTGAACCCCTTGGTGCCTTCACAACGAGGGCTGATTATGATGAGGCTTGGTTCAATGACTACAAGGATAAGTGGGAGGGCATGTATGACGAGGTTCAGGAAACTTTTGCAACCGTTGACAAGAAAGAGTTCATAAGAAAGTTAAAGGAGCTTGAAAGGAAGAAAAGGAGGAGGAGATAAGTTATGGAAAGGTTCGTGGAAACCTTCAAAAAGTTTATACTGTTGTGGTTAATTTTCTTACGCTGCTTGAGTTTGCGGCCAAGGTGCAAAGGTAGAGTGCAATTTGTACGTGCTGTGAGAAGTTTATATCATTCATAGACCCCTTGCAGAGGCTATCTATGCCATGCATTTAGATAAGCAGCTTACTGAGTATATGCTCTCTCGTTTGTATCGTACCATGAGTCCTTGATATGTCTGTGTATTCTCGCATACCCTAGTTTACCAACAAGGGCATACACAGGGATGTGGCATCACAAACGTCTTCAAGAATCTGTTCAACACTAATGTTTTGCATCTTATGTTCACAGGCTTTTGCATTGGGGGTAAGTGGGAAGTCAAGGAGCTAGCGAGTAGCACATGCAATGAGCACTGTACCCCAATTGTCTTCACTATTGGTTTTCCAATCGAAACATGCTTTTTATTTCTGCTTAGGGGAGTCTACAAGGTCTGAGAGGCATGATTTTTTTTTTGCAATGAGGCATTATCAAGCTGTCAATTATTGGTAAAGAGTGATTGCTTCAATTTCTCTAATTTCCAAGCAGTTGTCCTAGAAGTAATGAAGCAGGACTAGCATGCAAGTCGTCAGAGCACGGTCATACCTATTTATAGTATGTTTGTTCACAAAAGTTAGTGGGAGAGTTTCAAATTTTATATTTTGAGTTTCATCCAGTGTATTTTTCTGTCATGACTGTGGGCAAGGGCGGCCGAAGTGGAAGGCAAGCGCCCTCAAAAAATTTGCACACACCAACTCACTAAGGGTTTCGAGTTGAGGACCTCTAAGCTCTACAACCCAAATAACCACATTCTGCTCTCTTGGGCTCATGTGGTGTTTTCCATACGGTTATTAAAATCTTAGTGATATATGCCTATTAAAATCGTAGATAAGTTTGTCTTTCAAGTTAGGTATGTTCTTACATTGCTAGATAGGGTCGATGAATTTGCTTAAAGTTATATACAATGGGTAAGGCTGGATTCACAAACATTTGAGTGGCCCTTGCACATGACCCATAATGTAAAAGCCTCAAGAAGCTAAAATGCAATTGTCTAATTTTGCGTGACATAATGCATCTTCTAACAGCTTGTGTGAAAAGCCAAAACCTTTTGTGCCCTATGTCTATGTTTGGGAAAGTGAACCCCAAACATCTCAAGGAAAAATTAC

**PROTEIN TOC75-3 (TOC75-3):**

TGCACATCGAGCATTCATGGATGATGTAAATCTTATCAAACAATAAGAAGCTGCGAGAAAATCATATTTCACTCTCCAAATTTGACTCTGAATCTTACATAGTGACCACCCAAATCTACATATTGAGACATGAGGGACGGCCCTTCAGGAAGCATAGAATAGTACAAAAATAAGTTGGTGCCTGCACTCTGCACACTAAATGCTTGACAACTTGAATAACAAACTACCTCATTAGAAATGCATACAAATTCAACAAGACAATTACGTAAAAATGGTGGAAAAAAAATGAAGGGGAGAGGAAATCTTACTGAGGACCATCTATATAACCCATAGAAGAAAATTGGGAGGCCCATGAATAATGAAACAATGGAAGTCCAAAGGCATCATGGTAGAAAGGTTTCGACCCTAGAACCTTTCTCCAAACCGGACAAATAATGATCCTGACCCTGTATTGCAGTCTTTTGTGTACTCAGCCCTGATTAACCCTAGCTTTACTCCCACCCCGTAAGACATGCCATTACCAGCACGTCGGAAAAATTCTGTTGGGTTTCCTCTAACATCTTTAGAGCTCCCCAGATCTGTTCCATATTCAAGAAACCCATACGTGTGTGTGTTCAAGATAGGCACACGCACCTCTGTGGCCAGCTCCAAAAAATTCCGGCACGCTCCCAATTCGCCCATGTTGTAGCCTCTTACAGAATAAGGACCCCCTAAAGTAAAAGCATCGTATGCTGCTAGGTCCCCAATGCAGCCACCATAACGACCGTGCAAAACCAGAATGGGAGGTGGAGCATTTTTCTCGCCTTCTTCTACTTTCCTTAGCTGAAGAAAATGTGTAAGTGTTAACTGATGCCTATTGAATATGGGATATTTGGTACCAATGCCGAGCCCTTGGTCAACCTGAAAAATATACCGAGCACCAACAGGTGTTCCATTGATAAACACAGTGTTATCTCGGGTCATGTTAGTCTGCAAAAAGCCAATTCGATCAATGCCCGTATCACTTAGAGTTGTAGGAGGGCCGTCCATGCTAAGGGCCCCATTTGGAAGTGTGCGGGCCCCATTTGTACAGACAGCACTTGTCTCATCCCTCGTGGTAATTTCTTCGAGCACCAATCCATAGGTGAATTTGCTTTGCCTGCTGAAGCTCTCTGATATATTAGCTTTTAAACCTGCCCGGTCGACCCAAATTGCAGGCACCTCATCAAGTCCTGGACCACCAGTAAATACTGGACTAAGCTTTCTACTGTTGAAACAGCTAGCTTTAAAAGTTCGGTTGCGAGGATCATCCACACCATCCAAATATGGATGTACGTACTCCATCTTGAAACCAAGGTCATCCTGAGGGTTGAGAAGGTTATTTGATGTTACAGAGCCAAAAAGTGATCGATTCAGCCCCTGCAAATTGCGATGCTCAAATGTCACACTTCCACCTGGCTGTATTGTTGCCAAAGTAGGCCAGCCAGAACCTCCAGGAGCAATACTCCACTCTGTGGAGACTTCAGCTGTCTTCTGATCAAGCTCTTTAAGCTTGACTTCAACAACAATGCCGCCATCGTTTTCTTCATCTGGACGTGGATTGACTTCAATGTTTGAAAAAAGACCGAGGCTATTGACACTCCGAAGTGCCTTTTTTCCTGCTTCAATGTTGAAAAGAAATCCCGCTTGTAGCTGCTCAGGGAGTTCTCTTTCTACAATCTGCATATGTGTGTTTCCTTCACAAGGAAGCCCCATCTTGTCTTGGAACTGCACAATGACCTTGGTAATGTCGCCCTCCACAACTTCGCAAACAATTTCTTTGGTGTTAAGGTTACCAAAATTCACAACCTGGGCACAAGCAAAGCCATTATCATGGTACCACTTCTGGACTTGATCTCTGATCTTCTGTAGGATTCTTGCTGTGACCCGCCTTTCTGATCGAAGCCAAGCACTGATATCTTTTTCCACATTCCTGGCAAGTATGCATGGGCGTGCCTTCTTGAGACGGAGTTTGTACTCTTCCTCTTGAGATTTCATGAACACTTCACGCTCCTTGTCAGTCAACTTTGTAATATCTACATCTGGAGGCTTCGTTTGTGGCATGAGACCCACATTTACACACCTCAACCCTTCTGTAGGTTGCCACTCGCTCTCTGAGAATGTAACTTGCATTCTGAGGGTTCCATCTATCTGCGGCAAAGCATCCATGTCAACCCTCTCAAACATCCCGCATCCTGCCAAATTCTCCAGCTCTGTTTGCAATTGTGCTTTCATAAAGACTCCACCAGGCTTCATTGTGATGAGCTCGTGAAAAGGGTCGTCGACGCCAACGGGTTTTCCAGTACTGTTGTCAATCAAGTCAATGCCAGAAATCTTGTACTTCTTGCTTGAATTGAGCTTGGTTACATTGGTCTTTGAACCTGGTTTCATTCCATGTGAATCATAGTTACTTTCGGGGTCTTCTGAGTTAGCAAAAGCCATGAAATGCTCTGCCCAATGACTCCAGGGAGACCCACCACCACCACCCCAGCCACCCAAATTGCTACCTCCTCCCGAACCCCCAAATCCAGCAAGCTGAAAAAGGCAACCTCCCGTAATTCCTGCTGCAGATCCTACAGCCGCAAAACCCATTGTTCTGCTGAGCATTCTGAAGCTTTCAGAGCTCTTCACACTGATCAGATTGCGCTTGAGGCTCCCCGCAACCTGATATGTTATGCCCTTGCTTCCTTGTGATTGTGCATTCGGTGCAGTGCAACAAGAGAACTTGTGCTGCAGTTTCAAGGGGTAATGACCCATGCAATGATGCTCCTGGGCTGTTGGGTCCAAGCTCTGTTGGGACACGGAAGCCATGGCTGCTGAGCCCAAGAGCACAATGGCTTGCCTCGCTATACTACGATCTTTCAACTTCCAACACGTAATCTGTATAGCTTTGTATATCAATGTGTCCATATATCTATGTATAATATTAAGGGAGAGTGAATTCAGGCAAGGAGAGATGCCATGTGGCAGCAAATGCCACATCCGGCATCTTTTCTCACCGGAATCCTATTTTTTTTGCTCAAAATGAACCCAAATGTCCCTTCTTTAAGGATCGGAATCCATTTGGACGTTGAAAGCTTTTGTTTCAATTTCAAATCTGAAAGCTTTATTTCATTAAA

**AMINO ACID PERMEASE 2 (AAP2)**:

GGGAGAGGGAGAGGGAGAGGGAGAAGAGAGATTCGTATCATCAGAAAATTGTGTAGGTGTGTGTGTGAGAGTAGAAGAGAGAGTGAAAGAGACTGAAAGCTTAACTCTCATGCAGCAACCATTTTCTCATACTACGCTTCTTCTCTTCCTTCACACAGCTTCAATGGCGACCTTGATTTTTCTTCAGCAGTAGCTACACTAGAAAGTAATTGCTGAGTCGAAGCTTTGCTTTGCTTCTTCCTTCACATCCTCCTTGCCGCTTTCTTCAGCCCTGGCCGCAGTAAGCTAAACAAGAAGCTTTGAGCTTGTAAACTGAAGCTTTCTTCTTCCTAACTGCAGTCTGAAGAGCCACCATAGCCGCTTTTATAAGCAGAAAGGAAGAGAGCAGTTGAAGAAGTTATCATTTTGCACTCGTACACTCCTTTCACGCAGGTTTTAGACATGGCAGATATGGAGTCATCTGGTAGCTCTAGCTCTTCACACAATGCCCCCCTCTTCAACGACTCAAAGGCCCTTCAAATATCTGGCTTTGGTGATGGAGAAGGCCTCTCTTCTGATAGAGATGAAGATGGACAGCCGAAACGAACTGGAACTGTGTTGACGGCATCAGCTCATATAATAACAGCAGTGATAGGGTCAGGAGTACTTTCGCTGCCATGGTGTATGGCTCAATTGGGATGGATTGCAGGGCCTGTGGTGTTACTCTCCTTTGCTTTGGTCACCTACTTCACTTCTGTTCTCCTGGCTGACTGCTATAGATCTCCCAATCGAGTCACGGGGAAGCGGAATTACACATACATGGATGCCGTGAAATCCAATTTAGGTGCGCCATCCTTCCTCTCAAATCGGTCATTTCTAGGCTTGTATCTTTTGGTGATCTGTTTGAAGCCTAGAAATATTCAGAATCTTTGATCCTTAATGGTTTGAGCTGTCTGATTCAATTTCGATCAGGTAAGTCATGTAAATGCTTGTGTGAATGTACGGAGAGTTTGATAAGTTAAAATCAAGAAGTTTGTGTCGCTGCGGCAGGGTAAGAAGTTGCTCTTCTATGTCTACTGGTACACGTTAGAGCTTTTTGTGGTTCAAATGTGATTATATAAACACAGCTTATTTTTTCAACTCTTTCTAGTATTAGGGCCGAACTCAGCTTACATCTTACCTGTGTGCTGACTTGCTCTGTAGCTTTCATTTTACTCATGCTGGCCAGGTCTCTCCATCTTGGTGTCACCTTCTGAGCACCCTTCTGTGCTCGCACGTCAAGGGGATTGTGTTTATCAATATTGTTTTTGCACGTTTACTCGCAGTTGCACGTGTAGTTCTGCTAACAACCTTTGTTTGGGATAGAAGGCTTCTGAAAAATTAGCGGCTTATAGTTTTCTCTGTATTTGGAGTTAATATTACACACGATCTGAACTCTTGATTCGCACCAAAGCCTGGTGACGAATGAGGAGATGCTTCACGCGATTGTCCAAAGCATGAAGCATCGCTGGTGAGATTTGCCGGTGAATTGTAAGGGACGTACATGAAAAAAGTCAGCATACTCACTACTTTGAAGCAATGACCACACTAGGGCCAGACAGCACCTCCGCCTTCCATTGCTGTTGTCAGACGTGACTAGGTTCTCACGACCAATTTACCTTTAGATCTTGTTAGGTGACGATGTAGGAGGGAGGCAGGAAGACAAGGAACCTCTATAGCGGAGGTCACCCAAGCCAACAATTTCTTGTCGCAACACCATAGTAGCCCACCCTTCGATTCTTATAGGAGTTGAGTGTCTGTTTTCCTAATCCTTCCAAATCTTGATCAACCATGGAATGCACCACAAGAATCTGAATTTGTAGGCTGGCATGCATTTTAGAAAACAAAGCTCATACCTTGTGTACCCATTTGTTAAGTTACAACTTAAAACGTCAGTGTGTTGTGACTGCCACTCAAACTCAGACCCGATTGTTAACCAGTTCAAGTAAACTAATTTGAATGTTGTCTCAGTATGCTGTATGTTGGTTTTACAGTTTATTTTAATCCATTTTTGAGATGTCATATGATTTCATTGCCAAAAGCATTGGATTACGTTTTTTGAAATTTCTTTTGCAAGGGGGTCTGCAAATATGGCTCTGTGGATTTGTACAGTATACCAATCTGATCGGAACTGCCATTGGCTACACAATAACTGCTTCGATAAGCATGGTGTAAGAATGCATCATTTTGTCAGTCAAACTGACTTAGTTTGGGCTTTAAGAGTGACTTAAATTGAATGGAGGCAGGTCTTTTTGTTTGTAATAAACACCTTTTCAGTGTACATGATGCAGTGCCATTCGCCGTTCAAACTGTTTCCATCAAGAAGGGAAGGATGAACCATGCCATATATCTAACAACCCATATATGATCTTGTTTGGAGCATTACAGATCATTCTTTCTCAGATCCCCAATTTCAATAAATTATGGTGGTTGTCTATACTTGCGGCAGTTATGTCGTTCACTTACTCATTGATAGGATTTGGGCTTGGTGTACGTCAAGTTATAGAACCTGGCCACTCTTATGGAACTCTTGGAGGGGTCAGTATTGGAGAAGTGTCAGAGGCTGAGAAGGTCTGGCTAGTGTTTCAAGCTCTGGGAGACATTGCATTTGCATATTCATTCTCCATGATTCTCATCGAAATCCAGGATACATTGAAGTCTCCGCCTCCAGAGAACCAGACCATGAGAAAGGCTACTCTTCTTGGAGTGAGCACCACCACCATATTTTATATGCTTTCAGGATGTGTAGGCTATGCTGCATTTGGAAACAGTGCACCTGGGAACATTTTGACAGGCTTTGGGTTTTACAACCCGTATTGGCTAGTGGATTTTGCTAATGCTTGTGTGGTGGTGCATCTGGTTGGAGCCTATCAGGTCTATGTTCAGCCCATATATGCATTTGTAGAGACTGGAATAAGTAAGAAGTCTCCAAAGTGTGATTTTGTTTCTAAAGAGTATGAATTTTACATGCCAGTGTATGGCACATACAGCTTCAATGTTTTCAGATTCATTTGGAGAACGGCTTTTGTAGTTGGTTGCACTCTTGTCGCAATGCTCCTTCCATTTTTCAATGATGTTTTAGGGATTTTAGGAGCCACGGGGTTTTGGCCGTTAACAGTCTATTTTCCCATTGAAATGTACATTGTGCAAAAGGATGTAAAGAAATGGACAAAACAATGGTATGTGCTGCAGTCTGTCAGCATAAGTTGCTTGATTGTTTCGATTGCGGCACTTCTTGGGTCAGTTGAAGGTGTTGTACAGGACCTGAAAAAGTATAAACCTTTCCAAACACAGTATTAAAATCTATATTGAACATTACAAAGCACATTTTTTGTACTTAAAAACACATTTTATCTAATAAAGACGACTCCTATGCTCATTTTTTTTTGTA

**AMINO ACID PERMEASE 3 (AAP3)**:

GAGAGAGAGAGAGAGAGAAGATGAGAAAAATGAGGGCAAATGTAGTGTGTGGTGATTGTGAGGTGTTAGAGTGTGTATCAATAGTGAGTTTTGAAGGGGGTGTAGCGTTTGAGATCATTTACAATGCCAGCAATGGAGCCTATAGCAGCAGCAAGGGAGATTAGCAAACAAATAACGCTCAAGCAATGCAGAGCCACCCATTTAGGAGTCCAAGGCTTGAGCTTGTTTTGAGAGATGTGCATGCTAATGGGGAAGAAGACAGTCAAGGGCCAAAATGCAGTAGCCCCCAAGAGGCCCAATATGTCATTGAAGAAGGGCACTATCATTGAGATCACCGTCGTCGTTATCACAAACAGTGTCCTCCATGTCACTCTGAACACACTCAGCTTGAATGGCCTCCGAAACGGTAGCGGTACCTTGTGCTCGTATTTCATGAATTTGTGCTCAGACCACCTGCTCATCATCCAGTTCTCCGAGAATGCGAACACCGGCTGCGTGAATACCTGCGATTAAAGAAGTTGAAAATTGAAATAGTTGGGGAAAAGACTAGTCGTACCTGGTAAGCTCCCACAAGGTGCACTACAATACAAGCATTGGCAAAGTCCACAAGCCAGTATGGATTGTAGAATCCGAATCCAGTCAGGATGTTGCCAGGTGCCATGTCCCCAAATGCAGCATAACCGATACAACCAACAGACATGTAATATAGAGTGGTGATGACAATGGCGATTAAAGATGCCCTCTTCATTGTTCTATTTTCCGGAGGTGGAGATTTCAAAGTATCCTGAAACATTACAAAGGGCACTTGTATAACATAGAGCAGCTGGTACAACCCAGCATTCAAGCGTCTTGCTTTACCATCATTTATGCGTCTTGTAGCCCCAAAGTGAATGTGTTAATTGTATATGCATGTTTACAAGAGCACACACACTATTGATGCAAGCAGTCACATTTGCTTGTGTGTGCTATTTATGTAAGCAGAATGATATGTGGTGCTTTTAACGCTGGACTAGAAGCCTGCTTATCACTCTCACCATAACTATCACAAGTAAAAGTGCTGTCAAATTAAGCCTTCAAGCTGGATATTCAAAAGCACATATTTTCAGATTGTTATGTATGCCAGGATTCTACTATTTAATCTTCTTATACATTGCAGTATTGATTATTTACCTGAATCTCAATGAGGATCACAGTAAATGAGTAGGCAAATGCAATGTTTCCAAGGGCTTGGAAAACAGACCAAGTCTTTTTCTCCCTTGAAATGATATCTGCACTAATTCCTGTGAGTGTGCCATAGGAATGTCCTTTCTCTGATGCTCTCCCAATAGACAAACCAAGGCCTATTGAAGAGTAAGAGAAGGACATAACAGCAGCTACTATTGATAACCACCAAAGTCTGTCAAAATCAGGTATCTGCGACAAGATAATTTGCACCGCACCAAAGAGCAGCATGTATGGGTTGTTAGATGTATGGCACTCTGCCTTATGACCATTTTTGTGGAAGCAGTTAGATCTGGTAATGGCCACCATACTGATTGATGCTGTAATGGTGTAGCCTATAGCCGTGCCGATAAGGTTAATGTATTGCACAATGCCGCAAAGCCATACATGCTTTTCACCTAGGTATGATTTCACAGCATTCATGTAAGTGTAGTTTCTCTTCCCTGTAATTGGATCAGGGCATCGATAGCAATCAGCCAGAAGGCGAGAAGTGTAATATGTGACAAATGCAAAGGCGAGTAGAATGGGAGGTCCGGCCACCCAGCCAATCTGAGCTATGCTCCAAGCAAGAGAGAGAACACCCGAGCCTATAACTGCTGTCACCACATGAGCTGATGCAGTCCACACGTTCCCTGTCCTTTGAACATGTCCATCATCATCACGGCTCTTTGATCTAATGAGCACAAAGGGGCTGCCAATGTCTTTGCCTACCTCCACAGCACATCCACTTCTCACGGAGTCATCTTTGTTTGTCGAGCAATCCGAAGCAGAGTCATTAGGCCTTGTGAGTTGATAGTTCATTTCATGATCTTGTGCCATGATTATGTGTATGGGATACTTAATAACACTTGCACAGAGGTCACTTATTGGCCATTATACAGACTAGCTGCTCCACCTGCCTGAATATGCTTCATTATCTGTCCACAGCACTGATATTTTGATGTAGGGTAAGATAGTTGTGAATGCTGACGACTCATGACATTGTCCGCCGTCGATGAACTTCCGTAATGAAGTGTTTCAGCTACATGGTAATCGACAAAGCCTATAATTTTGTGCAAGCTTCAAAGTAGAAGTAGAAAAAAAAACACAGTCGCAGAACAAATTATGTGCTTACAGAGATACTGATGTCAAAAAGGGAGGATATCACCATCTTTAAAAGGGAAGCAAGTAAACAGGCAGCAT

**LYSINE HISTIDINE TRANSPORTER 1 (LHT1):**

GCTCCCTTATATGTCATTTTTTGCATGGGCAAAAGCCTTATTATATTTCTCCTATATATATATGCCATATCACATGCTAAACTTGGGAGGAATCAATAAGAACCCCATAATATTTCTGGAAAATCCCTCTACCCCAAAAGCCAGCTTGTGAATCTAAGGTGCTAGAGGGTCTAATTTGTCTGCAAGATTCCCTTATTTTTTACAACCAATTTGGTGCCTCTGTGCCAAATTGTCAATAGCCAAATCTCTTCTTGTGCATGCTATTTTTGGATTCTCTAATTAAAGCCCCACTATATAATTTGGAGCCTTTGATCACTGCCCCAATAAACCTCTCTCTCCTCAATTTTACATAATGGTCCAAGAAAATGGTGCAGAAGAGAACCTTGGAAGAACCATATCTACTAATAATAGTCACATCCCCCTTGAGGATGAGAGAATCAAGAATGTGGATGATTGGTTACCTGTAACAGGTAGTAGAAATGCCAAGTGGTATTACTCAGCCTTTCATAATGTTACAGCCATGGTGGGTGCTGGTGTCCTTGGCTTGCCCTATGCCATGGCCCAATTAGGATGGGGACCTGGGATTCTGGTGTTGGTGCTTTCATGGGTGATAACATTGTACACATTATGGCAAATGGTGGAGATGCATGAGATGGTCCCAGGAAAGAGATTTGATAGGTATCATGAGCTAGGCCAACAAGCGTTTGGAGAGAAGCTTGGCCTATGGGTGGTTGTTCCCCAACAGTTGATAGTAGAAGTGGGAACAGACATTGTATATATGGTGACAGGGGGAACTTCATTACAACAATTCCAACAGCTTGTTTGTAAAGGCGATGGTTGCAACAAATTTGGTCTCTCTGTCTATATCATCATTTTTGCTTTGCCCCACTTTCTTCTCTCTCAACTTCCCAATTTCAACGCGATTGCTGGTGTCTCTTTGGCTGCTGCTATCATGTCTATCAGCTACTCGACCATCGCATGGGGGGCATCTATACATCACGGAACTAGAGATTACACATCATATGATTACAATTTCAACACAAAAGCAAATGGGGTGTTTCAAGTTTTTAATGCACTAGGAAATGTGGCCTTTGCCTATGCTGGGCACAATGTTGTGTTAGAAATCCAAGCCACTATACCTTCCACCCCAGAAAAGCCATCAAAGGTCGCTATGTGGAGAGGAGTCCTTGTGGCTTACCTTATTGTGGCCTTTTGCTATTTTAGTGTTGCTATTTTGGGGTACTGGGCCTTTGGCAATACTGTTGACGGTAACGTTCTCAAATCCTTAAAAAAGCCTACTTGGCTCATTGCCACCGCCAACCTAATGGTGGTAGTGCATGTCATTGGAGGTTATCAGGTATTATTACTAGCTTAGTTCTAAGCATTTCTTATGAAGTTCACTAATAAGCTGCACACAATCAACTTGTAAGTGTGTACATGCAGGTGGGAACATGTGTACACATGCACACATATATTTAGATACATTAATTCACATAAACACATGCACACACCCACGTGCACAGACACATTACAAGTGAGAAAAAGGAATCAGGGAACCAATTGAAATTTCTTTTCATCACTACTTGAAAATGAAAATGCCATTATTATTTTGGAAAACAATCAAGCGAGTGCGAGTGCGAGTGCGCACACACACAAAGCCATCCAGGGGTTGTGGGGCTTAAGATGGGAATAAGTAGCCAACAAATTATTTTTTATCAAAACAAACCTCAACCAGAAATCCCATATTGAAATTAGTAACTCTATTGAAGATCACAACTCTACATTACACAAGGGCATGCACTCACAAAACCCACCTAAAAGAGATTTTTTAGGCTCTCTCCAAGTGTCATCCCATATGCTAATTTTGCACTCCTTTCTTCTAAGGTTAAATAGAACATCTTTAAGAATATGTAAGTGTGTGGGTTGGTGAATATGTATTTTTCATAAAACCAAAATTATAGAATTTGGTCTCTTAAGCTTTATAAGCAAAATGAAAAATCCTAGCATAATTTTGGTGAATCTTTCTACCAAACCCACCCAAAATTTAATTGAAAAAAACCTAGGCAAGAATTTTGGTAGTTGCTAAATAGCCAAAAATTTTAGGAGACCCTATAAAATAGAGTCTTTCTACCAAAGCCTACAAATGGAAAGCTCTAGGCAAGAAATTTTGTTTTTGTGGAATAGCAAACTATTTTAGGAATCCTTGAAGGATAGCCATTGGGTTATAAATAGCTCCTAATTTTTTCCTTTATTTCTTCTACCATGTTTGTCATTGAATAAAAAAGGGTACCATTTTAGCTTAAACTACTCATGGTAGGAGCTTGCTACAGCCCTTGCATCATCCCTATATGACATTCACCTAATATTTTGCAAGCCCTTTAAACCTTAAGACTAGTACTTGCAAAATTGTATTTCTTTTTTTTTGAGAGTTGGTTAGACATTTCAATGGTAACCCTGGGATGAGGGTTTGAAACTCTATTGGCGCTCATGTACTTGCAAAATTTAACATCTTAAATTTTTGATTGGAAATAAGGTTCTCTATATAAAGAAATATTGAGAGTGGATCTTTTTTTTATGTTTAATGCCAAACTCTTTAAAATAGCATGAAAAAAGTAAGTATTAAGCTAACTAAAACTTGTGGCCTCCATTAAATGTGAATCCTTCCTTCTCAAACATGATGGAACTCCAAATATTGAACAAGGAAAGTCAAAGGGAGAAACATGAACATTATTAGATATTAAATATTGTGGAAGGCCTTATTTTCTTTTGTAACACTATTAAATATTATTGTGGAAGGCCTTATTTTCTTTTGTAACACTATTAAATATTTTGTCCTACAGAGACACGTCTCTTAGAACTTTGATCTTTTAAAGCTAG

**CATIONIC AMINO ACID TRANSPORTER 1 (CAT1):**

TTTTTTTAGGCCGGAATTATGTTTTTTATTATATATGAAATGAATTGCCTGGGCCCAAAACAGGCAAAAAACAAGCACAATACAGTCAAAAGACATTTCAAAACAGTGTCTCTCGCTCAGGGCTTCAATGTTCAATGTTGTTCATAATCCAGCATGCATCATGCATGGGCCTTGACCGACCCTACAATTTAAAGGTCAAAATGTGTGCAATTCTCTATTTTTTAAGTGAATTTACATGTCTTAAAAAGTGGACCGTACCTCTATCGAACTTTTTTTCCCAATTTTAAAATTCCCGCACTCGTACCGAATTTTCGAACCGTATTTTTGTACCTGATAGATATGATTGTAGTCCTTTTCGTGTTATTCGTTGGATTGGTCTTGTTGCATATGAGGTTGACTTACCTTCCCATGTTCGTGTTTATCCAGTCTTGCATGTCAGTCACTTGAAAGAAGTTTTGCATGTTAGTTTCTTCTCAGGATTTGATTGCAGTCGAGGATTTATCTTTTGTACCGTATGAGCCTGAAAGGATACTCGGCTTTTTGTGAAAAATCGCTTAGAAATAAGCTTATCACGGAATTTAAGATTAAATGGCGATATCGGATTCCACGTGTGAGTGGTTACCTACTCTCCAGAGGTATTTTCTTGATTTTCTTCGGAGCTCAGGAGTGAGCTCTTAGAAGAAGGGGAGTATATCATGTGCATGCCAGAGTTGACCGTGTATAATAGTGAGTACATAGGCCTAGGCTAGGGGTGCATAAGACTAGATTGGCAACCTAAGACCTTATCGTCGGTAAGCCACCAGTTTTGGGCACCCACGTCTGGATGCCCACGATTTGTATACACACATGGGCAGATCATCAATTGTGTGCAGTCTTCAGGAGAGCGCTGCATGAGTGCAGTGCAGTAGCGATCATGGTGCCGTGTGGACGCACGCCTGCTACACATGTAGCCAAGGCCGACAACAGCCTTGGGCTCTAGCAGGGTATTGCGGGTGGTATGGGATGCCAAGAGAGTAGATGCTGGTAAGATGTGCAATCTAGGAGCTCATCTTGTGCGTCGACGGCTGATCATGAATAGCGCATAAGATCGGAGGTGAGTCAAACCTGCACCCCCTACAATCTCATAGGTGTTTGATTTAGTTTGGCAGGTGTCTTGCAACTTGGGAGAGCAGGTCCATGCAAATCTATGTCCTGGAGGGGGAAGGATGTACATGTAAATGCTCCTCTAAAGGGAAGATCAGTTTGTTACCCTACATGGTCAGACAAATAAATCACTATTTCATGGGTTTTGGCATTGATAAAGCATCCATAGCATCCAGCACCACAAACCTTCTATGACAAGTTAGGAGTGTTGAACGGAGTCGATGTTCCTGTACACGTGAACATGAAGAACACACTGACAATGAGAATCTTCGATGATAAACACAAAGCTGCAAAGGACAAGCGTCTAAATCGACTACATTATTTTACAGGAAAGCACAAAAAAATCTCAGGCACTCACAATACTGTTTGGATCTGAAGAAATAACTGAGATCAAAGATTGTTTTTAATCACGTTCATCCTGTGAACAACTGCCAGTTTAAAATTATACATACAATAACATATGTACTATTTACTTGTGTTTAGATTGCTTCTTGAGCATTTCCCAAATCCGGTCCACTTTTCTGGAGTGCCTTCTCCTCAGTTTGAGGCTTCATTTCCACAGTCCGGCCCTCCTCACTTAAACTCTCTGTTTGGTCCATTGCGGTGTCGTAGGAAGCATGTAATCCGAAGAGAACGTAATACACAAGTATCAACAGTGTCCACACTCCAAACCTTGCAAATGATAGCCTATCAATCGAACCCAATAGGAAAACATTCATGATAATGGACAGGGATGGAAGCCAAGGCACAAAAGGAGTCCCCCATACTTGTGGTTGTCGAGCTTTCGGAACAAATATGGCTAGACCCAATGTAGCCAAAAACCAGATGGGTACAGTAATCACAAAGCCTATCCACCCATCATGTGTAGCCCAATAATAAGCAGTAGCCCCAGACGACAAAAGAATTGTGACAAGGAACATCTGAAATAGCCGAAGTTCCTTCACCGTGCTGTGCCCAGCAACATAATAGCGCCTCACAAGAAGAGCTACACTCACTAACATGAAGATGAAGAGTGTACTGATAGAGAGAAGATTGGCTAATACATCAAGGCTAGTGAAAAATGCAACAATCGCACTAGCTGAGAACATGACTAATGTTGCTCGTATTGGAGTCCCTGTCTTCTCGCTTACTTTAGCAAACCATGGAGGAATCATATGCGTACGCGCAATATGCGTAACATATCGTGCCTGCCCCACAGCACTCACCAGTAAAACAGTTGTCATGCCTTTCAAAGCACCCAAAGCAACCAAATACTGAGCCCATTTCCATCCCGCTCGTGCTTTAAATGCCAGTGAGAAAGCAGCATTGGGATCAATATCCTTATAGTTCTGCATCATGGATAAGGTCAATGCCATTAAGCAGTATAGAACTGTAGTGATTATCATTGAGCCCAACAGCCCAATAGGAATATCTTTTCCAGGGTTTTTTGTTTCCTCCGCCATCGTTGCCACAGCATCAAACCCAAGATATGCAAAGAAAAGGACTGATGCCGCCTTGAAGACACCACGCACTCCAAATGGTAAGAACGGTATCAAATTTGAAGGCTCCGCATGCACGAGGCCAGCAACTATGACAAACAATATAATCACCATGTTCGCAATGGATGCTGTCCAGTTTAGCAGAGAAGTAGCACGTGTGCTCCAAGTTGCTACAAGGCCAGTGATAACTAGTGCCGCTACTGCTATAGGGTCGAGAAGATTGAAATCCTGTGCCAAGCTTGTCTTTATGCGGAAATCATCTGCTTGGGTACGACCACATAGAGTAGCAAAGTAAGAGCTCCAAGACCTTCCTACAGCAGCTCCACCTATGATATATTCTAGTATGATGTTTCCAGCAGCAATAAATGCAATAAAATCCCCTAGCTCAACACGCAGATAAGCAAACGAGCCGCCAGCTACCGGGATTTCGACAGCAAATTCAGTATAGCAAAAGACTGACAGCATGGCAGAGATGCCAGCGGCAACGTATGAGAGGATTACAGCAGGGCCAACTTCTTTATTGGCTTCTTGACCTGTGAGGACAAAGATACCAGCACCGATGATAGCGCCGATGCCAAACCACATAATGTCCCACCAAGTGAGGGTCTTCCTCATGGGATTCTTGCTGCGATGCACAACGCCATGTGTCTCCATAAAGTCTGAAGATCGTGCATACACACGATTGAGAAAGCGCGCCGGTGTTGCAGCAAGTGCCATGCGGTAGGCACCCCAGCTTTCAAAAGAAGCCTCCGGCAGGAAATTTGCCTTGCAAGGAAGCCATTGTGACCCCTTCGCCTCCATAGTGATATCCAGGTGCAAACACAAACACAGAAATGACAAGGTATGTGTGTGTGTGTGCGTGTGAAGGGGAAAACCCTTCAAATTTAGTATGATAGGAGCATAGGATTGAATGGATATGAAACATGAGATATAGAATTACAGAGAAAGCCCAATGGTATTAACAGAGAGCACCTGATTGCCGCCATAGCCAGAGCTAAAAGCAGAGGATTGCAGCCGAAGTAGAAGGCTTTTTGTTGGCACACATGGTGCATGCACGCGTTTTCGAAATCTTCGGATGTTGGGTGGAGAATACAAAGAGTAGCGTCCGTCGTACCTTGGACTTCTCAAGCTGATGATGCTCTGTAGTGGTCACATTAGTAAATGGGCGATTGTGTGTGTGTTGATGCTTTTCTAGACGAGTCTTGGTTCTTAATGTCTTGACACTCGGCTAAAATTGTGTGTGTATGTGTGTGTGTGTG

**PROTEIN NRT1/ PTR FAMILY 8.2 (NPF8.2):**

GTTTTTTCTTGTAGCATCAAATTTGGTACCGAAAGAAAGAAAATATGGGGTTACAACGGAAAGCTGTGTATGTACATTGAGTCAGGCTGAAGATTAGCAACTGCGTTGCAGGCAGTTGTGTTGCAGATATGAAACGATGTGCTGGGCTCAAATCATTGACCCAAGCACTCTTCAAATACATTGATTGGCCAAAGCTCCCCTCATCAATCCAGGCGACAGTCTTAAGAACACACTGATTGGCCAATTCTGCTACTGTTGAAATATCATTCGATGCAACGCAGTACACAACCAGTGCATGATCCCCTACTGTTGTACCTTCCTCTTTCACAACACAATTATCCACACAGTAACTCAGGGGGAAAAGCTTATTTTGAGAAGACTTTTGACTTCCCATAGGCCATACATTATTACATGCCATCATTTGTGACATCCTCCATACACATGATCCAAGCAGCCCTAGAAGGTGAAGTAATGTCTTGTCCATATACCATTGTAAGAAAGGCATCAAAAACTAGACTGCTAACTCTTTGATTCAATCAAACCAAATGAAAACCATGGAGGATAAAGCATATGCTCTTGCTGTCCCCTAACAACTGAAGTCGCACTGCCAAATTGAGGCAAAACACCTTGACTACCCTCATGGCATATTCACAAGGCACTTACAACTGCAAATTGCAATGGAGGCAGTAGAACTCATCAACCACCTTTGAGGAAGAGCCCATGACACATCTTCAGCATCCCCATGGCTGTGGAGTACAAGCTGAAAAATCAAGTGCAGCACATGAGATGAATTTCAGTCGCTTATGAGAACTACGCCACTCAAGGTCCCATTAGCCTAGACATTTAAAAGAGCTTTCGGTTAACGAGGGCAGCCTGATATCTGAGAACGGCAAAAAAAGCCATGAATCTCAAGGTGCCAGCTTCTCAAATGCTTCCAGCCAAACCCCCTGTCCCCTACTTTATACAATAACAGGGTCGATGGAGCATTGGGGTTTAAAAGAGCAGCACTCTGAAGGGTGCAGCAAGACATGGAGCATTTGCCAGCCCTGTGCCTTCTTTTACCATGCAATGTTCATTAGCTAGGATCCACTCCTTAACCTTAAGTCAGCCTCATTCCTTGTAACCTCACACATGCATTATCTAACCAATCAAAATCACAGTTTCAACAAATATGCAAGGCTACAAAAAATGCACATGAGTATACAGGAGTACAACACAAATTCTATATAGATAACAGCACCAACTCGATGATGACAAGGCCCCAAGATTGACTGCACGTCTTCATTTCTACTTTTTCAAAGATATCTAAAGTTTCGAATTGATGATAGCTCATTGTTGCCACAATAGACCCTCAACAGGTTCACAAACCTTGTCTGCACATGCTAACATAACATTCCAAATGACTCTTTCCAATTAGCGTGCCGGGAACACTCAATATTTGGCAAACAAGATGTTATACATCAAAAGAGACTTTCTCACCATGAAGTAACCTCAAATTGAGGTTTATTACCTGTAGGAATAATGAGCATACGATTTATTGAAAAAATCACATGTCGACCCTTCGGATCCAAGCTAAATATGTTTATCAAGCCATGTTGTGAGCATATACAAGACTATATCGCTTGTTGTCGCAATTGACCAACAGTATCTTCTTATCACTCAAAGAGCTAAATTTTGATTAAATCTTGGCGTAGCACCATTGGCAAGGTGTTCTTTCCAAAATCATCATTGAACAGCACTGTATAATCCCATCTTTCTAAATTGTAAACTAATGAGAGCGCCTAATACTGCGGAAATTATATGTCCATCTCACAAAAGGGTGCAGGCAGTCAAAATGGGATACAATAAAAGCATAAACTTTGAAAAGATTGATCCCACCCTCTCAGCAATCTTCAACAAATGGAAACAAGGCACCTTCCCAACCGTCATCTCCGTGTTCTCCTTCAAAACCATCATCTCCTTGCTCTCCTCCAAAATCTTCACTCTTGCTATTTCCATCAGTGCTTTCAACATCTGTCACTGCTTCATCGTCTTGCAGTTCAAACTTCTCTAAATTTTTATCTATGGAAGTAGGCAGACTCTCAGAGTTCTTATTCGAATTGACCGTTTGTGTTCTCAAGCTGGGAGAAGCCTCCTCAATTGCAGATGCCATCCATCTACTGTCAGATCTAGATTCGGCAGAGTCCATGCACTTGAAACCCTTGTGGTTACTATAGCAGCCATTAGGAATGCAGCCTGCATTTATCAGCTGTTTTTCGAAAGTGTCATTGCCACCAGCTGTCACCGCATCCTTATCATGCAAAGGCACTGGAACTCTCACCCCCTCCTGAAGCACATCCAAGCGATTGACCCCAGGTTTCTCTGTGGCTTTTGAGTCCATAGCATATGTGTCTCCATTTTGGAAGGAGATATTCATAGATGGGAAGCCAACGGACGCTGGGGTTATACTTTTCTTAGATTTGAACAAGTCAATGCTGGGGGAATGTGTATTTTGCAATTTACTCCTCTCCGCATGAAAATTCTGGTGAGGCTCATGAACAAGAAGGGTCTGAAATTGCAGGTCTAGGGAACGATCTTCATAAGAATGATGGCCGTTACTAAATCCAACATTGACCCCGCTACACAGACTTCCTTCTGACCCACAACTACTTCCACTGCTAGAATACTGACTTTGTGAACCCCCAACATCTGTGCTAAACCCAAATCTCATGGTTGACTGAAGGTTCAACGGATTCAGACGGCCCTTTGGAAGGTAACTTGCATCTCTCACGTCCTCCCCGAGTTGATAATTCCTAAACATGTCAAACACAGGCACCAGATCTTTCTTTCTCACCTCCACAGCACGGCTAACAACTTTCTCAATTTCAAATTCTCCACCAGAATCATCTTCCCGGAGGACTTTATTGAATGCTGGTCCAGATAGTACATGAAAGGAAATCTGAGAACTTGCCCATATTGGTGGCTTGAGATGATGCATATGCACCTCAGCATTGGACAAGAACCATTGCTGCATCTGTTTATTGGTTGTTTCCTTCAAGATTGTGTCTAATTTCCCAAGTGCTGCTTCATTCTGAGATAGGCCTGCCTTGATCCTTGTTGAATGAGTTGAATTTCCTGCTCCATCATGAGCAACATTGAGGGCATTTGAAATATTGACCTTTTCTTCTCGCTCAACCCAATTTGGCCTGCGGCACAAATCCCACTTCTGTAAAGGTTCCACGATCACCTTGAAGTCGAGTACTTCTTTCTGAAGGCCAGCTACAATCGGAGAAGAAATACTATCTGAGTGTCCTTCAGTACAAGCTGACAAATGAAGTGCATGTCGAATAAGGTGACCAGATGGAGAAAGGATCCACAGCTGTTCCTTTAAAGTATTTTGGCCCCCCTCAGCATCAAGTTTGCGACCTCGTCCATCATGAAAAACGACTGCAACTGCTCCAAGTGAGCTGCAAGACCTACCACTTGCAGCAGCAGCAGCACCACTCATTGTACCACACCAACCACCATTTCCAATTTTTATTCTACTCACAACAGACAAACTAACAGTCGCAGGTAAACGCCCTGAAACATACTGGCTGGACCTCAGTGGACCTCTGTTGGACCACCATGGAAGTGATTGCACTGATCCAAGAGCACAGTCAACAAGACCTGATCCATGAGTAGAGGGCCCAACAGCACCTCCAAATGGTGATATAGCAAACAGATGATGTGTTCCCCTAGATGAGCTGACAACAATCCAGTGGCTATCTTCACTGAACGCTATATCTTGTATGACCGCATTAGTCATCCCTCGGTAAAGCTTGTAAAGGTGTACATGTGATGAGTTTGTGTCATATGAGCTTGTGTTGATACCATTCATATTTGGAGCTGGCAGTATTCGAAAAATGTTGATATTATGCCCATGAATAGAGGCAGTAACTAATAAAGTTCCAGTGCCATCAAAGCTTAATGCAGATAAAGGACTGTCGTGAGCTCGAAATTGAGATACCACAAGCTTGGTCACAAAATCACGCACAATAACCGTTCCTGCATGCTCTGTATGGCCACTGCTGCCCTTGAGGAATGGGCTCCCTGAGACTGGTGAGCCGCCACCCTCAGTGCAATACCTTGATAATGCTTTGTATCCTTTATCTCCCAGAGTTACAATCCCAGCAGCAAGCTGCTTACTTGATTCTTTTGCATAATGTGCAACCAAACTACCATTTGCAGGAGAAGTTGACGGGCTGACACCAGGAGAAGGGGTGAGGTGCTGGGGAGAAACCCTTCCAGTATTTGAAATAATAGGCTGATTTGCTGCATAAGCCAACCACCTTGGTCCAACAGCCAATGCACCGTGTCCAAAACTAAGCCTACTTGATCCCAGTTGAGAGGAAGGATATGTGAGCACACTAAAAGTACTCTGGAGTGTTGCTGCATTGAAGCAGTGAACTTGGGTAGCAAGAGCAACTGCAACAACTCGAGGGCTACAACGAACAGCAAGAATTGCTGTGCGAAATCTCAATGTATGTACATAAGAATGGCTTTGTAAAGAGTAAAATTTAACAAGAGTAGGAGCATTTAGATTCTCCGTTACTGGAGGACTTGCAACAACTGCACTATAGCCATTTGTAAAAGCGCCGTTATTGCGACAATTTGGTGAATCAGAGGTGACAACTAGCATTAAAGGATGGGCCAACTTGAATTTCTCATCAGATGTTTCTGTAGGCTGAGGCTGGACACGCAAAAAAATGACTGGACCATCACGTTTCGAGACGATCTCCCGCACTTCATTTGCATTTTCCACATCCCAAACTTGAAATCCATTTGTATAAGTCAATAATAGTACTTGTCGAACATTGGCTGGTGCTAGCTCTATTTTGTCAAAGCCTGCCCATTGCACCTGCTCATATTGACGGTCTTCGTCAGTGAGCGTGATTGTCGATGCAACTGACGAACCTGCAGAGCGTACAGTAGAAGCCAAGGTGCTGGCACTGCGAAAGATGGCTCGAAATGAGCTTGGGATAAAACCACTGCTGCTGCCTGTCCTCGGTGAACAGGCTTGCAGATTGGCATCATTCATAATCCAACGTCCACTTCCCAGTCCTCACAAATCTTGAGATGAAAGAAACGCTTTGAATCCAAATAAAATACAGCCTTGTGCATACCCAGGCTGTTTCTTATACTACAATGTGAAACAGCCTTTGCTTGATTATGCTAGGGAGATCGGATGCCCACTGACTGCACCATTGTAAAACCTTTTGCTTTTGCTTCTAGCTCTCCATTTTCTGATGAATGGTTTCTTGCGAATGAGTACCTACTGAAATGAACAATCTGAAGATAGGTTTTCCCAAAAGGGTAGCTTTGATGGCAGCCTTATTGCTCTAAGAAAGCTGCACTTACTAAAGTAAACAATGGGATGATTAGGAAAACCCAAATCTGGAACAGGCGAGGCTTGCAAAAATTGGCAGGTCGGTGTTGCTGCTTTCTCTATTTACACACAGACCCCTGAATAGCCTCGAATACATACAGGAAGGGAAAAGGGTAAGTCTATTGCACCCACTTCGCCTCTATAAGAAACCTGCTGCTTCAATTCACTATATACACGCGCAGCGCTTGTCGAGCTCCTCTGTACAAGTTCGTGCCTCGTTTCGGAGAAACAGAACGTCGTCTGCAGTTGGCTAAAATCTGTACAAAGAGGATGGCGATGAAGGACCCAATTTCACGGCCGAAGTAACGAAGCAGACGACGACGATGATGATGATGAGGATGACGGCAATGACGATGAGCTTCTTGATCCACTGACAGTCTTGTATGTATACCACTTTGCACATGCCACATAGAACACACAATTAACTAAACTAATGCCTGTAAGGAGCCAGTAAAAATAGTCCAAGTGGCCTCTATTTATGTTACTTGTAAACCAACTTCTTGAGCCCTTGCCAGTGATACTTTCTATGATTGTGATAAGTAAGCTGCTTAAGAAGCTCCCTACACCGATAGCACACAAGTACAAGGCCATGCCAAAGCTGCGTGTCGTGTCGGGTACCTGGTCATAGAAAAACTCTTGCTGCCCAACAATAGCAAAAACATCGGAAACTCCACTAAGTATGTACTGAGGTAGCAACCAAAATATAGTTAAAGGGATTGTAGATTTAGGTAAATCCAGTAATCCATGATCTCTTGATGCATCAATCCTTTTTCTCTCAATTATGGCTGCAGCAAGCATTGACAAAATGGACAAAAACAAGCCCACCCCAATCCTTTGCAGCATAGTTATGCCGCGTTGAAGCCCAGTAAGTCGGCTTACAATTGGAACAAACACTCTATCATAGAGTGGTAGCAGTATGAGGATAATGATAGTTGTAAAGCTCTGTAAAGAGGCTGCTGGGATCTCGAAATGCGAACCCATTTTTCTGTCCATGGTGCTTCCTTGTTTAGTAAAAAGAGTTGATGTTTGTGCAAATACTATCCCATATGTCAACATTGCCAACCAAACGGGCAATATGCGAATTACTAGCTTTACTTCCTCAACTTGTGTGACAGTGCATAGGTTCCAAGAGTTTTGCTTTGCCTCGAAGTCACCATGCTCTACAACAGCAGCTTTATCTAGAAACCTCAAATGATCTGTATGAGAAAGCTTTCGTCTCCCCAGATGTAGCAACTCCTTGTCATTGCTCTCATAGAGCAACTCTGAGCTGTGAGGGAGAGGCAAGCTACGCTTGCGAGTGGCGGCAACGATGACTTTGGCGATGCGCGTGAGGGGGCTTCCTGCAGGCAATTTGTGTCTGTAAAATTTGGTGCCAGCCAAAAAGGTAGCCAATGCAACCATCATGGCAATGGCAGGGATACCAAACCCGAGCCCCCAACTCACATTGTCCTGTATGTACACCAAGATTGTGACAGCTAGCAGGGCCCCCGTGCATATGCCAAAATACCACCAGTTGAAAAAAGATGCTTTCATTATTCTCTCTTTGGGGTCCTCTTCGTCAAATTGATCTGCACCAAAGGCTTCTAGGCATGGCTTCACACCACCTGTCCCTATAGATATCAAATACAATGAGAAGAAGAAAAACCCTAGTTGCCCCTTGCTTGCCCTCGGACATATTCCATCTTTCGTGCATGTAGGAGGCTTCAAAGCGCTTAGACTTACAGAAAGAGTCAAGAAGAACAATCCAACAACATAAATAACCGAGACTGTTGCAATGGTCCAATATTTTCCAAAATAGGAATCGGCAAAAAATGCTCCGACTAATGGGACGACAAGGGTTACACCAGACCAATTGTTTGCATTCTTTACGGATGAAGCCAGGCTCTCATGTAGCACTGTGGTCATATATGTTATCAAATTTGAAGCAATGCCATAGAAGGATAGCCTCTCACTAAATTCACCACCGATGATGAAGAGGGAGGCCTTCCAGCCACCAGTGAGGAAGCGAGAAACAGGGCGGCCCTTCAAGTCGACGGAGCCATCGGCTACCCGGCACTTGCACTCCTCCTCCTCCTCCTCCTCTTGCTCTTTGAGTAGCAAATCCGCTTGCGTGTTAGTCGCAAACCCAACTTGAGCGGTTTCCTGAGAGGAAAGCAGTTGCAAATCTGCTGGAAAAACATCTCTCTCCAGTGTGGGTAGTTGTGAATTCCGCATCTGCGACGGTGGTTCTGCCAGGAAGGCATCGTGGGCAGTTTCGAGTGATTGCGAAGCAAGCAATTCCGGCTGCAAGTCTGTTGTTGAACTGTCTTGGGCGGTTTTCTGAGGCGAAATCAGGGTCTCTCTAGCGCTGCAAGCTAAATGAGTTCTTCTGTGTGGCTTCTTGAAGTGCTTACCTGAAGAGGATGAACGTGCTACCTGCCACAATGGCCACAAAGGGCGCTTGGAGCTTGCTTGCTCGAGTGAACAAGTGCACTCCAGCGACGAGAAAGTCCTGCATGCCATCGAGCTCATTGTTCCCGTTCAACGAAATTGCCTTATCTACTCCTATCTAGTGGGATTGGCGCCGCGGGATCGACACCCGCGTCGATCCCGCAAGCGTCGCGCTCGC

**ADENINE NUCLEOTIDE TRANSPORTER 1 (ADNT1):**

TATTTACCATTTACCGTACATGTAGAGCCTCTTAGCAGTAGGCATTCTTGAAAACTAACCATTTACCATACATGACGAGTCTCTTATCAATAGGCATTCTTGATAGTGATAAAGAAGTTAAAAGCGGCTTTATCAGTAAGATTCCATGTACACCAAAACAAAAAAGTTAAAAGGTGCCTGCCAAAAGAGGGATGCCATTGCATTGATGCGGTTTTCACTTCAACAATCGTTAGAGGATCTAAGGCCACTCTAAACATATGCTCAATTACTTGTATGACCTAACAAATTGACTCAATATGTCACATTATACAGATTTACTTTTAGAACATTATATGAAAGTCACGGGTACTTCCCTTAAAAACCCAGAGCCTTCTATTATCATGGAAACGCCTGGATGAGCATTCCATCAGTAGCCAATGCGCGCATTTGTCCCTTCCAGATGACATCCAAAAAAACTCTTCCTGAGACACAAAAACCGGACCGAAAGATATTTGAGCACATTATGTATACTCAAAGTAGTCTGCATGTAATGGTACTCTCATGTGACCGCAAAAAAGCACCACTTCGGAACTTTATTATTACAAACTGGCCTCCACTGCCAGCAACTGTTTGAGTTTGCAAACTGCCTAGCTTACAAACACAGCAGTAATGCCTTTATTGAGCTTTTTGAATTAGGCAAAGCACATCTGAAAACTGCCCATGCCATAGCTACCTGATTACCGCACATAACCAATTGATCCAGGTATAATACTGTTCACAAGTGGAAAAGACTCCAAAAAGTTTTGGTGTATTTTTCCCATGAAGTCACTCCATTAAAAAATTGAAGAGCAGAGCTAGCAATTTGAGAGAGTTTGCTCTTTCACAATCAACGGCAGTAAAAAATTCCTGGCCTACTCCATGGCATCGTTCCACCGTTCACTCCACCTATGAGGTAAGAATTTGAGCAATTCAAGAGAAAACTACGAAAACCCTTGGAACTCCCAAAAATACATTCCGCTGCAGTACCCCTTCATTAACTTCTCTGCTACAAAACTTGAACCTTGTGAGTGAACATGTTTGCTTAATCTGAGATGCGGAACTCTACTCCAAGAAAGTCCTTCACCAGCTCATAGGTCACAAAAGCAATTGAAATTGAAGGGACCACCTTTACAGAGTTGGGAATCAGCCCTTTGTATAAAGCTCGAAAGCCTTCATTAGCAACTGTCTTTCTGAAGGCATCTGTCATACCAGTGTACTGCACTGTAGCTTTCACCTGCCCATCAGCTGTGATGATATTCGACGCGTCTCTCCATCCAACCATTTGCAGCCGTCTGCGTACTACATCCAGAGGATAAGCCACAGTCTGGCCAACCATACCTGCTGCAGCTCCACATCCAAGCCTTATCGGCACACTTAATTCCCCGTCCGTTCCTGATTCATCCATGTTTTTCTTTTTAACCCAGACTTTCAGAGACTCATACACGGAAAAGTTCAAGCCCACATAAGGAATCACTCCAATAACTGATGGTAGCCAGCCCTTGTACAAAGCCCTTGCTCCCTCCTCTCTGATGATTGTGGATGTAGCATGAACCATGCCTCGATAGCGATTTGGTGAGTCACTTGTCTGAACAGTAAGTCTTCCTCGAACCATGTCCATTGGATAGGTAGCAGACATAGCAATAATCCCAGCACATGCACCAGCTCCAAGGCGGAGAACAGGAGTTAACTCAGCATTTTCTTGTCCAGGCTGCTTTCGGTAAAGCCACAATACAGCCCTTGATGCCTGTTCGTAGCTGTAAAACTTGATTGCAGAGTTTGGAATGATTCGAGCACAATTTGCTCCGTTGCCTTTGAAAAATCCAGCAATGCCTTCATGTGCCCAAATATACCTTAAACCTTGAACTGTCCCATTGTATTTAGGATTCAGAGAATATTGAACCTGAAGAAGTATCTTGAGACGTTCAAGCGGTGCCACTGCAGTGCGTGAAACTCCACCAGCAACCCCTCCAGCAATCAGTGATTTGCAGACAGAAAGAGCAGAGTGTCCGGAGGCCTTGACATCTGGAGTGGCGAGCTTTGCCCCCTCCGCGATTGTGGCTAGTGTAGACGAGGTGGAGGCTTCTCCATTAACATATAAAGGAGGTTGTTGACAGAATGCTACACTTCCATGCCTTGCAGCATCATTACTATCAGATTGAACCTTGTCATACTCTTTCTGTAGCCTGGGAACTGCAGATGCTGCCTCAGACACCATGGATGATGGAATGCAACAACAATTAGCTGCAATTTCTTTTACAGAGACCGGGCGGCGATGGTGCTGAGTCGACTGCTCGGCAATGGAGCTTGAGAGCCGAGAAGCTGCAGAAAAGAAGGTGCGTGAAATTAGCGACACTTCATTAAGAGAAGAAGAACACGGCCCCCATTTTTTGCAAAGATTCTTG

**MITOCHONDRIAL PHOSPHATE TRANSPORTER 1 (MPT1)**:

GCATACCTAGCTAGGTGCGGTGAGGGGTAGGGGGCTGTCGACCCTAGGTATGCGGAGCCTCGTCTGCTGAAGCTCATGATGCTGAAACATAAGCTGGGAAAGGGGGCCCCTTTCCCACATTGTCAAGGAAAAATGCCCAAAAGCTCTTGTGACATACATTGGGGTCTCTCCTCAAGTCATGCCGAGATGCTCTTGCCGGACTTGAAGCCAATATGTGGAATGCCCCCTGTGTGGTTTTGTCGTCGGAGAAAAGCATCATTGTAGAAATATGCCTCTTCTCGAACCACATGCTGTGATGAAGTATCTCTAGCACACCAATATACCTTATACTTTCTACTTTGCTAGGAAACCAATAGCCAAAGGTTATCCAAGACATATAATTTTCCTTGAATTTGCTTTAGTAAACCACAAGCACCATAGAATCTGGTAAAGCCAAAATGTGGTTGCCTTTTCACTACAATTGGCCATATTGAAAGCTTCAGTGCTCTTGACAAAAACATTTGTCAAACTTCATTTGCATAGTGGGAACTCTGTATACATGATATACTTAAGAATCTAGAGTTCTAAAAACATTGCAGCAATATATATAATAATGCAAGAGAAGAGAGATGTTTCCAATTCTTCATGTCCATTATAACAGTACAAAAGACATTTTCGTCGTATGTTGTAGATAATGATGCAATATAGCATCCCATTTCTTATCACATTGTTAATTAGGCAGGATTTTTTTCTTAGACTTCCACCTTCTCTGTTTTTAATTAAGGAGTACAAATAAGTGAATATGTATGGCGTGTCGTTTCTAAGCGAGTTTGATCTCAAAAACTTGAATTAGATGATGACTTGCAACTCCAAAAAGTGTATCAAAATCGATCTGGAAAACTTGACGACGGCATGATAATTTTTGACTCCAAATAAATGACCTTGAAAACTTGAAGATGACATGATAATTTGTTTGATCAGGCGAGTGATAGATGCATATGGAAAGAGCGAACGAAGGCTCAAACAATTTGTATTGGGGTGCTCCTGTAAAACCAAACTAGTTCTTGTATGAAGTAAAGTGGGAAGCCTTTGTTACACAAAAAGGAAAGTTTGTAGAGCAGCTGCAATCATAAATTAAGGTTTTCAAAATGGATCCTGAGTGGCTAAACAAAGGGGACAATGCATGGCAACTGATTGCTGCAACCTTGGTGGCAATAATGAGCGTGCCCGGGCTGATGAGCATTTACGCGGGCGTGATGAGGCACAGGTGGGCCGTGAACTCGGCCTTGATGGCACTGTATGCGTTCGCAGCTACACTCATCTGTTGGGTGCTTTTTGCATACAAGATGTCGTTTGGAGAGAGATGGATACCCATATTGGGGAAGCCAGGGATTGCTTTGGGGCACAAGTACTTGCTACACCAGGCATATTTACCTGCATCGTCTTTCCATTACGAGAACGGCACGATCAAGACTGCAAAAGAGATGCCGATGTTCCCAATGGCGACACTAGTGTATTTCCAATTCGCGTTTGCCGCGATGGCGGTTGTGCTTGTTGCGGGCTCGCTTCTGGGACGAATGAATTTTACTGCTTGGATGCTGTTTGTGCCCTTATGGGTGACCTTCTCGTACACGGTCGGTGCGTTCAGTATGTGGGGCGGTGGCTTCCTTTACCACTGGGGGGTTATCGACTTTGCAGGTGGATTTGTCATTCACTTGTCCTCTGGTGTTGCTGGCTTCGTTGCTGCTGCTTGGGTGGGGCCAAGGTTGGAGGAGGACCAAGTACGTATGGATGAGAACTTTAGCGTGCCGCTAGTGGGGGTTGGAGCGGGGCTCTTGTGGCTGGGCTGGACGGGCTTCAACGGCGGTGCGCCCTACGCCGCGAGCCTGGACGCCTCCATGGCTGTCATTAACACTCACATCGCTACTGCCACCAGTCTCCTCCTCTGGACCATCCTCGATGTTTTCTTCCACAGAAGGACTTCTGTCCTCCACGCTGTTCAAGGCATGATAGCCGGCCTTGTCTGCATCACCCCGGCCGCAGGTGTGGTGCAAGGGTGGGCAGCTATTATAATTGGAGCAACTGCAGGAGGCATAACATGGTATACAATGATCAAGTTGGAGACATGGGAGTGGCTAGACAGGGTGGATGACACACTACGGGTGTTGCACACACATGCGTTATCGGGGTTGATCGGCGGCATTTTAACAGGGCTTTTAGCAGATCCTCACCTTTGCAATTTATTCTTATCTGTGAAAGGCTCTAAGGGTTCCTTTTATGGATCTGCTTCTCAGTTGGGTAAACAGCTTGTGGCTGCTCTATTTATAATTGTTTGGAATGTAGTTGTGACATCCATCATTTGTTTTCTTATACAAATCAAAATTCCTTTAAGAATGAAGGATTCAGAATTGAGGGAATATATAGAGGATGTTTATCCTAGGCCTAGATTTGTAGGACGAGATGAAGGAGCAGATGCAATACAGTTGTATGGTAGAAGGCTCCACCAAGCTCCATTAGTAGGTGCTGAGGAGCCTATTGCTGTGCTTGCATAAACTTATTTCTTCTTTCCTATGACACATTAAAATTCATTAGTCTTGTGCGATCCCATAATAAATAGGCTATTGAAACAATGGGAATTTTCCAAATTCAAGGCTCCTCTTCATTGTATAAGTTTGAGATTTTTGTAAAATTACCATTGGTAACCAGTTAGATTTTTGAGTTACCATACATAAAGTTTTCCAATTTTAGGGTCCTTTAAAAAAAA

**AMMONIUM TRANSPORTER 1 (AMT1):**

CGGCAACAATGGCTTCATTGGGCATCATTTTTTTGGCCTCGCGGATGTGCCAAGCCAGTTTGCTGGAATGGATTACAGCTTCTTTCTTTATCAATGGGCTTTTGCAATTGCAGCAGCGGGTATTACGTCGGGATCGATTGCAGAGCGCACGCAGTTTGTCGCATACCTCATCTACTCATCTTTCCTCACTGGCTTTGTTTATCCTGTAGTCTCTCATTGGGTCTGGTCCGTCGATGGATGGCTTAGCGCTGGTAACACCAAAGACCTCTTGTTTGGCAGTGGAGCTATAGATTTTGCTGGCAGTGGTGTTGTGCATATGGTGGGAGGAATTGCTGGCCTGTGGGGAGCCCTCATTGAAGGCCCTCGAATAGGCCGTTTCGACAACAACGGACGCTCAATGCAACTTAAAGGTCACAGTGCTACTCTTGTTGTGTTAGGTACATTCATGCTCTGGTTTGGATGGTATGGATTCAACCCTGGCTCTTTCTTGAAGATCTTAGTCTCCTATGATGGAAGCTTCAAGGGCAATTGGACTGGTGTGGGAAGGACGGCGGTGACGACGACACTAGCAGGCAGCACGGCAGCCATCACCACTCTCTTTGGAAAGAGATTAATTGGAGGACATTGGAATGTGTTGGATGTGTGCAATGGCTTATTGGGTGGCTTTGCTGCCATCACCGCCGGATGCTCTGTGGTTGAGCCGTGGGCTGCGATCATATGCGGGTTTGTCTCCGCATGGGTGCTCATCGGATTCAACATCCTAGCTGACAAATTGAAGTTCGACGACCCCCTTGAAGCTGCACAATTGCATGGAGGATGTGGTGCTTGGGGGCTGATCTTTGTGGGGCTCTTTGCAAATAAGAATTATGTGAGTGAAAGCTATGGAAGAACGGGATCGACGCCGCACGGGCTCTTTCTGGGGGGTGGAGGAAAGTTACTTGCGGCGCAACTGATAGAGATAGCAGTAATAACGGCGTGGGTGAGTGTAACAATGGGGCCATTGTTCCTAATACTGCACAAGTTTAATCTATTAAGAATATCAACAGAGGATGAGCTGCATGGGATGGACCTTACAAGGCATGGGGGGTATGCATACTATCATCATGAGGACAGTACAAGTCCTACAAAAGAGACACCACCTAATGAGAATAAGAAAGCCTCTGCTATACAGTACGCAGTCTAAGGCTTGTTAAAGTTCTGCTTGTACTTTGCATAGTTATTACTCCCCTTCCACTTTGATTCATCTTTGTTATAGGGACTCCACATTTTAAACACTTGAATGATAAACAGCCCGTCATGTATGGTTTTTTTTCTTTAAATTCCATGATATATTATTCATTATTCAAAAAAAA

**AMMONIUM TRANSPORTER 2 (AMT2):**

GCATACCTAGCTAGGTGCGGTGAGGGGTAGGGGGCTGTCGACCCTAGGTATGCGGAGCCTCGTCTGCTGAAGCTCATGATGCTGAAACATAAGCTGGGAAAGGGGGCCCCTTTCCCACATTGTCAAGGAAAAATGCCCAAAAGCTCTTGTGACATACATTGGGGTCTCTCCTCAAGTCATGCCGAGATGCTCTTGCCGGACTTGAAGCCAATATGTGGAATGCCCCCTGTGTGGTTTTGTCGTCGGAGAAAAGCATCATTGTAGAAATATGCCTCTTCTCGAACCACATGCTGTGATGAAGTATCTCTAGCACACCAATATACCTTATACTTTCTACTTTGCTAGGAAACCAATAGCCAAAGGTTATCCAAGACATATAATTTTCCTTGAATTTGCTTTAGTAAACCACAAGCACCATAGAATCTGGTAAAGCCAAAATGTGGTTGCCTTTTCACTACAATTGGCCATATTGAAAGCTTCAGTGCTCTTGACAAAAACATTTGTCAAACTTCATTTGCATAGTGGGAACTCTGTATACATGATATACTTAAGAATCTAGAGTTCTAAAAACATTGCAGCAATATATATAATAATGCAAGAGAAGAGAGATGTTTCCAATTCTTCATGTCCATTATAACAGTACAAAAGACATTTTCGTCGTATGTTGTAGATAATGATGCAATATAGCATCCCATTTCTTATCACATTGTTAATTAGGCAGGATTTTTTTCTTAGACTTCCACCTTCTCTGTTTTTAATTAAGGAGTACAAATAAGTGAATATGTATGGCGTGTCGTTTCTAAGCGAGTTTGATCTCAAAAACTTGAATTAGATGATGACTTGCAACTCCAAAAAGTGTATCAAAATCGATCTGGAAAACTTGACGACGGCATGATAATTTTTGACTCCAAATAAATGACCTTGAAAACTTGAAGATGACATGATAATTTGTTTGATCAGGCGAGTGATAGATGCATATGGAAAGAGCGAACGAAGGCTCAAACAATTTGTATTGGGGTGCTCCTGTAAAACCAAACTAGTTCTTGTATGAAGTAAAGTGGGAAGCCTTTGTTACACAAAAAGGAAAGTTTGTAGAGCAGCTGCAATCATAAATTAAGGTTTTCAAAATGGATCCTGAGTGGCTAAACAAAGGGGACAATGCATGGCAACTGATTGCTGCAACCTTGGTGGCAATAATGAGCGTGCCCGGGCTGATGAGCATTTACGCGGGCGTGATGAGGCACAGGTGGGCCGTGAACTCGGCCTTGATGGCACTGTATGCGTTCGCAGCTACACTCATCTGTTGGGTGCTTTTTGCATACAAGATGTCGTTTGGAGAGAGATGGATACCCATATTGGGGAAGCCAGGGATTGCTTTGGGGCACAAGTACTTGCTACACCAGGCATATTTACCTGCATCGTCTTTCCATTACGAGAACGGCACGATCAAGACTGCAAAAGAGATGCCGATGTTCCCAATGGCGACACTAGTGTATTTCCAATTCGCGTTTGCCGCGATGGCGGTTGTGCTTGTTGCGGGCTCGCTTCTGGGACGAATGAATTTTACTGCTTGGATGCTGTTTGTGCCCTTATGGGTGACCTTCTCGTACACGGTCGGTGCGTTCAGTATGTGGGGCGGTGGCTTCCTTTACCACTGGGGGGTTATCGACTTTGCAGGTGGATTTGTCATTCACTTGTCCTCTGGTGTTGCTGGCTTCGTTGCTGCTGCTTGGGTGGGGCCAAGGTTGGAGGAGGACCAAGTACGTATGGATGAGAACTTTAGCGTGCCGCTAGTGGGGGTTGGAGCGGGGCTCTTGTGGCTGGGCTGGACGGGCTTCAACGGCGGTGCGCCCTACGCCGCGAGCCTGGACGCCTCCATGGCTGTCATTAACACTCACATCGCTACTGCCACCAGTCTCCTCCTCTGGACCATCCTCGATGTTTTCTTCCACAGAAGGACTTCTGTCCTCCACGCTGTTCAAGGCATGATAGCCGGCCTTGTCTGCATCACCCCGGCCGCAGGTGTGGTGCAAGGGTGGGCAGCTATTATAATTGGAGCAACTGCAGGAGGCATAACATGGTATACAATGATCAAGTTGGAGACATGGGAGTGGCTAGACAGGGTGGATGACACACTACGGGTGTTGCACACACATGCGTTATCGGGGTTGATCGGCGGCATTTTAACAGGGCTTTTAGCAGATCCTCACCTTTGCAATTTATTCTTATCTGTGAAAGGCTCTAAGGGTTCCTTTTATGGATCTGCTTCTCAGTTGGGTAAACAGCTTGTGGCTGCTCTATTTATAATTGTTTGGAATGTAGTTGTGACATCCATCATTTGTTTTCTTATACAAATCAAAATTCCTTTAAGAATGAAGGATTCAGAATTGAGGGAATATATAGAGGATGTTTATCCTAGGCCTAGATTTGTAGGACGAGATGAAGGAGCAGATGCAATACAGTTGTATGGTAGAAGGCTCCACCAAGCTCCATTAGTAGGTGCTGAGGAGCCTATTGCTGTGCTTGCATAAACTTATTTCTTCTTTCCTATGACACATTAAAATTCATTAGTCTTGTGCGATCCCATAATAAATAGGCTATTGAAACAATGGGAATTTTCCAAATTCAAGGCTCCTCTTCATTGTATAAGTTTGAGATTTTTGTAAAATTACCATTGGTAACCAGTTAGATTTTTGAGTTACCATACATAAAGTTTTCCAATTTTAGGGTCCTTTAAAAAAAA

**PLASTIDIC NUCLEOBASE TRANSPORTER (PLUTO)**:

TGTGCGCGCGCGCGTGTGCGCATGAGATCCTCTTTACTGTAGATACGTGCGCTCTCCTCCACCATTGTCTTGCAAAGCAGCAGGACATGGGAAGATATGGCACGAACGAGCAAAGCAGCTGCATCACACTGCATTCTCAGCAGTAACGCTTCTCTGCTATTCATCAACAAGCACAAGCTCTGCAATTTCAAGCTCCCATGTCCTCCTTCACTCCGCTCTTGCAGAAAGCCTCCTCACCTCGTGCGATCACACAGAGAAGAGAGAGTGCCCACCTCCCATACAAACCCTACCAATGGGCCTTCATCTTTCTTTGCGCCTGCCTCCGAGCCCTTGCTTACAAGCCCCCCTCACCCCACCTTATCTAATGAAGACCTTCTTCCTACTGCTCTCTCTCAGAGAACATTCTCAGCGTGGGATATTATCAGCTTGTGGGTAGGCCTTGTTGTTGGCGTGCCTTCGTATTACCTTGCCGGCAGCTTGGTTGAGCTTGGCATGTCATGGTGGCAGGGTATCTTTACTGTAATGGCAGGCAACTTCTTACTCTTTTTTCCTCTTGTTTGGACAGGACATGCAGGCATTAAGTATGGCATCCCATTCCCTGTCATGGCTAGGGCATCCTTTGGCATCCATGGTGCACACATCCCAACTATTCTTCGAGCCCTTGTCGCATGTGGGTGGTTCGGCATTGAGACATGGATAGGTGGGCAAGGAATCTATGTTCTCTTGAATGCCCTCTTACAGGGGAAACTAGCCAACTCTTCCATAGTATTGATGGGCACGGCCCTTCCCGAATGGGCTTGTTTCTTTGCTTTTTGGCTACTTCAAGTGGGCATCATCTGGAATGGGGTTGATAGTATACGCCAATTTGAGAAATATTGCGCGCCTATATTGATAGTTATGTCTGCTGCCCTCTTGATATGGGCATACTCAACAGCCGGTGGATTTGGGGCCATGTTATCTGCTTCGTCGCAATTTGTAGCAGGAGGCTCCAAACAAGGGCAGTTCTGGCAGGTTTTTTTTCCAGCTCTTACTGCCAACATTGGTTTTTGGGCCACCTTGAGCTTGAACATACCAGATTTTACAAGATATGCCAAAAGTCAAGCAGACCAAGTTGCTGGCCATATAGGTCTTCCAGTTTTTATGGGATTGTTCACGTTTTTGGGTTTGGCTGTAACTTCATCTACTGAAATTATTTTTGGCCGTGTCATATCAAATCCTATTGAATTGCTGGGACAAATTGGTGGAGTCATACCAATCCTTACGTCTTTCATCGGTCTCACATTGGCCACACTCACCACAAACATCGCTGCAAATATCATGGCCCCTGCGAATGCATTTGTCAATTTGAACCCCACTCTCTTCTCTTTCAGAACCGGGGCTCTTCTTACTGCTGTATTAGGGGTTATTTTTGGACCATGGCAATTTATAAAGAGTAGTCAAAGCTTCATCAGCACATGGTTAGTGGGATATTCTGCCCTTCTTGGGCCACTGAGTGGCATTATTCTTGTTGATTATCACATCATTCGGAAGCGTCATCTTGATCTGGATGCTTTATATTCAACAAGTCGCACTGGAATGTACTGGTATACCAAAGGCTTCAATTTTATTGCCATAGGGGCTCTGTTGCTGGCTGTCGTTCCATGTATCCCAGGTTTCCTTTACACGATAGGTTTTTTGAAGAAGATCCCAAGCATTCTTTTAGCTGTGTACAACAATGCATGGATTTTTGGTTTCTTAGCTTCCGGGCTTCTTTATTGGGTTATGTGCATGCTTTATAACCTCTATGCAAGCTTTGCTGAAAATTTCAGCAAAACTAGTTGATTTCTATTGTGGGTTGAGCTCAGCTGTGAAATTTCTTAAAAAGAGCCCGTCAAAAAATTTCTTACTCTTTGCTACACTTTTCCTTATTGAACATAACTAAATTGAGACCAACAATTGTAAGCAGTAGATCAAGCAATGTTTGCCACCTC

**MULTI-DRUG AND TOXIC COMPOUND EXTRUSION (MATE)**:

AAGAGGAGGAGGAAGAAGAAGGAAGAAGAAAAAGGAGCCCTTCTTTTCTGCAGGCTTCATGCCCCATTCCTTCTTTTTCTAGCTTTCCATTGAAGCTTAACATGAAGTAGGTGTGCATTGTAGCTTTTGAAGCTTATGGACTATTGTATATTAGCTCAAGAAGCTGGCATTTTCAGTTCAACCCCATCGGCAATGGTCATTGGGCTTGTCTAAAGAGATAGTGATTCTGCTCATTTAAATGCTAGCTGACATGAGCAACCTACATTTATGGTCTTGTCATGCCAAAGCGCTTTTCAGCACATTTATCACGCTTCTCAATGAAGGTAGAAACACAACTGTCAAAGCTGAAAGCAAGGAAAGTAATACATGCTGCTTAAAGCTTTTCATGGGGGGTGCCAGCAATGTTTTTAAGGGTGATGAGCTTGGAAGGGAAATCGCTATCATTGCTTTTCCAGCCTTGGTTGCACTGGCAGCTGATCCTCTTGCCTCGTTAGTGGATACAGCTTTCATTGGTCAGCTAGGTTCTGTGGAGTTGGCTGCTGTTGGGGTATCCATTGCGGTTTTCAACCTGGTATCAAAGCTACTCAATTTTCCTGTGTTAAATCTCACAACATCACTTGTGGCAGAAGAAGACGCTACAGATTCAGGGAATGTGTGTGAAGTTGAGCTGGAAACTGGTTTATCTGGATGCTCCAATCCTGAAGAGAAAGATGCGTTTATCTCAATCAAAGATTCACAGAATCCCGTTGATGAGAATGTTACCAGCAAGGTAATTAGCAGTATCAAGCAGAGCAGTGGCACTGAGCAAAAGAGCTTACCGGCTGTTTCAACAGCACTAGTTGTAGGCTCCATGCTTGGATTTTTTGAGCTTACTGTTTTGACATTAGGATCTGGACGGATTTTGGATCTCATGGGGGTTACAGTGAGTTCTAACATGCGAATACCAGCTGAGCAGTACCTTGAGTTACGTGCATTAGGTTCTCCTGCTGTTGTACTATCCCTGGTGGTGCAGGGTGTTTTCCGTGGTTTTAAGGACACTAAAACCCCTCTTTTTGCCACTGCTTTTGGAAATGTTGTCAATATGGTACTGGTTCCCATTTTGATGTTTACGTTTGGTTATGGTGTTAGCGGATCAGCGGTTGCCACCGTCATATCCCAGTATATTATCGCTCTGATCCTTTTGATGAAGCTCTGTAGGAAAGTGGTGCTTTTTTCACCAAGGCTGGGTGGACTTCAGTTTGGTCTGTTCATAAAGAATGGCAGTCTTCTATTAGCTCGTTCAGCTGCAGTGATGCTCAGCTTGACTTTAGCAACATCACTGGCTGCAAGACAAGGTGCTATCCCAATGGCTGCGCACCAGATCTGCATGCAAGTGTGGCTTGCAACCTCTCTTCTTTCAGATGCTATAGCTCTTGCTGGCCAGGCCATTATCGCCAGTGCTTTGGCTAAAGGTAACTTTGAGCTGGCAAAGAGTGCTGCTTTTAGAACACTACAGATTGGTTTTGCCTTTGGCCTTGGTTTATCTGCGATACTTGGTGTTGGGCTTAGCACATTTTCCAAGTTGTTTACAAGCGATCGTGCGGTGTTGAATTTTCTTGCCCTGGGGATTCCTTTTGTGGCTGCTACCCAGCCAATAAACGCTGTGGCATTTGTGTTTGATGGCCTTCATTTTGGAGCCTCAGATTTTTTGTATGCAGCATATTCTATGATTATTGTTGCAATTTTAGCTTCGGGCTTCATGCTAGTTGCTGCTCCAATGTGGGGCTTTATTGGCGTGTGGCTTGGACTTAGTATGTTGATGACATTAAGGATGCTTACTGGGTTTTTGAGGGTAGGCACATCAAGTGGGCCTTGGAAGTTTTTGAAGACTTTGTGAATGTGGTATTACCACACAAAAACAGGGTAATGTGGTGGCTGACATGTGGCTTTGCTGAAGCTACGACCAGAGAAATCGCACTTTTGAGTAGGCGTTTCAAGTTTTCATTGCAGTCCTAGATCCTTGGTTTCTCATTCTAGGACCTGCAGCAACAATATTAGTTTACACAGTTGAGCTCTGTAGATCATCCACTGCAAAATCCTGGAGATGGTGAGATCACATGGTCAGCTGCGAGGCTGATTGTATTATCAGCTCTCAGTGTATGCATCATCCCAGTATTGATGTTTCCTCGGACTAAAACTCTGTGTTTGCAATAACTCTTGACTTGCCGAGTGTGGGAGGCAGCTCTAGCCCTGAGGGTTTAATGAATGCTGTTTTGGCATGAATAGCCTGGGGGTGTGAGGCACCTTGGAACTTCTCACAAGCCTGGTAGAGACTGAAAATACAAAAACAAAACCAGCTTCTTCTTCTTCCTCTATCACTACACCTTCACTGCCAGTATATTAATAATATACCTACATTTTCAATATATGTGGCTACACTCTCATCATTCTCCCCATTGACTTTATATAGCTCCTACAAGTTTTTGGGCTTCTGATTTGGTAGGATATTTGCATTGTATCTTTGTTTTTTAAATTTTTACCCACCATCCAGTTTAGTAGCAGAAAACTAGGTGCATTATGGGCATCAAAAATGCCCATTATGACGCTATCGGTATCATATATTTGAAGACCTCTCCTCTCTTTTACAGGTGAGGGTTTGATTCTA

**MAGNESIUM TRANSPORTER 1 (MGT1)**:

GGCACACCAACCATAGATTATCCTTGTGGGCTGATCATACATTGAATTCTTCTGTTGTTAGCACATCAAACATAATTGATAAAAATCAAGTATAACCATACTATTGAAATCAAGTTAAAAAAAAATTGGAGGAGAGAAACTTCTCCCTTGCTAAACTTAAAGATATTCATTCATTTTTTAATATCCTTCTGCCAACATGCACTTTGTGCATACCAACTTGCCTCCTTATGGCACCTCCCTACAAAGAACAGATTGGTATTCTCTCAATTTACACTTGAGGCTATTATGAGCACAGTATAATGCCATGCTCCTATATCATGATAGAGGCATCAAGCCTTTCCACTTGAAAAACAGCAGGAAAGAGAAAAATATCAACAGGCCGCATAGTCCGGCGATGCCAATGACCCACTTGAAAGCCCTCGGCTCGTCAAATATTGCAATTTCGATGTTCATACCAAACACTCCCGCCACAACCCCATAAATTGCTACAACAAAGGTAGCAGTCGTCAAAAGCAGCTCAAATTGAATGAGCTGGTTCCGAACATTGTCCAGCTGAATATTAATAAAATCTTCTGTATCATCAATGTACTGCTTCAGAGATGTTAACTTGTTGAGTGTGCTATCTACAACAACAAAATAGGCCTCCAAGAGCATCTCCAGTTGTACTATATTCTCACTATCATTCTCAGAGCTTCGTATGCTACTATGTCTGCTTCTAGTGTGACTGAACGCATTGCTGATGGTCTTCTCAAGTGTTCTATGCTCTGTAGGTGAGCAAACCGGAGATACCGGAGCTGACAACGAAGGTCCACCTCCAAGCTGCACATAATTATCTTGATAGAAGGAGGAATCCATTCGTTCTTTTTTCTCTGTTAGATACATCTCAGCCATGTCCCCATCATCATCCATTAATTGCTCAACCTCATCTCTTACCTTTTGCACACGGGAGGTCAAAGTGACGAGCTTGCTCTTGAGCCTTCGTACACGTTCCAGATTCAGTGTGTTGACTTTAGAGGTTAGCTCATCAAGCACAGGATAAGCCTCCACCTCCAATTCTGTGGCCTGACTGTCCAAAAATGTGCATGCCGCCTCTAAAGCCACCTCCAACGCTCTAAATTCAAATGGAAGGTTATCAGCCTCACCAAGCTGGAAGAGTCGAAGACGTGACATAACATCACTAGCAGTAGCACCCAAATCAAGATCCCCTGCGTGTGAATGCATGGATGTCCGCCTCTTCAAGCCTTGCGCTTCATTCTTTACTGTTAGCCTTTTTTGCAACTCGGTAACATACTGCAACACATGGCTATCGAGAGAGTTGAGAAGCAGGACCTCATCTGCTGTAATGACACATCTAATTTGCTCTAAATTCACCACAATGGCTTTCTCGCGGCCTAGTATTGTAGATGGATATACAAACAGGGGATCAAGGAGTCGTAGGTCACGGGGAGGTAACTCACATCTTCGCATTATAGTCATCTTGTCCACCTCTAGGAGTTGTGAGTTCCCAAATTTATCAATTCGAATCCAATGGCGTATGCTATGTCCTCTTTTCTTCAGATCTACTGTTGAAAACAGGGGGCGTGCAGGCTGAGAGCCTGTGGGTGGAGGGCTTGCATCGTGCATCCCCGGGCGAGGCCGCAAGAGGGCAGCTTTCAAGTCTGACATGAGAGCAAAAGAGGAGCTTTGAAAAAACCAATATACTACACGCGCTGGCGATGAAACTCATACATACATATTGACAAGGATGGGTATGCATGTTGGTGCATGCTGGGGCTGCAATATAGCCTTTTCATTGACCACATTCACTTTCTTGTTACATGCAATGAATGTGATGCGCTTAAAGGCTCTGAGTAGTGATGGTGAGGCTAAACTTTTACACAGAATCCACTTGCTTGACAAGCTCTTCTTCCTTCAAAAGACATTGAATAGCATGACATTCATCAAATTATGTTGCAAATGCTCATGCTAAGCATCTCCTGCATTCTTGCACCTATATCGCCAATGCCCTGAAATCTTACACATGCAGAGATGCAAAGTTACATGTAGAGTTCTCTGTTCAAGCTTCAAAATTGGTGTGCAAATACAGATATATGAATAGAAAGGAGGGTTCTGCGTGTACAGAATGAAGATTTGATTAAGTAGTGCCAGGAGCTTCAAAGATCATTCCGCCTATGCCCAACTAGGGTTTCGATGTGAAAAAAATAGGAGGGCTCATTCAATAAAGCTCCATGCACAGGGAAAAGAAGCCATGAAAGCACAGAGAGAGTGCGCAGACGATGATTCCTGCCAAGTGGACCAACAGAAATGCGAGGAGGAGATGAGCGAGCAAAAAGCTTGGTGGTAGCCTTT

**UDP-GALACTOSE TRANSPORTER 3 (UTR3)**:

TGCTGGTTTTTAATTGTTTTGATCACAGAATGTCGAAGAGGTCGGGACCTTCGCGTCAAAAATGGTGTTCTTGAGCTATTGATTGCTTCTGCGTTGCTTTCCTCTAGTGGGCGATGGGAGCTTCAGGTCTTGTTGCAGGGCGCGTCCTGCTTCTTGGCTTCTGCATTGTTGGCATCTTCTCCGCATACATTACTCAAGGCGTCCTTCAAGAAACTGTATCCACAAAGAAGTTTGGCCCAGATGGGAGACGCTTCCAGCACCTTCCTTTCCTCAATTTTGCACAAAACGTTGTCTGCTTTGTCTGGTCGCTATTGATGATTGCTGTGTGGACCAAAAAAAGCAACAATCAAGCGCCTTTGTGGCTGTACTGGAGTCCAAGTGTTACTAATTCTATTGGACCAGCATGTGGCATCCAGGCATTAAAGTACATTAGTTATCCTGCTCAGGTGCTTGTAAAATCTTCTAAAATGATTCCAGTAATGCTAATGGGGACGATTGTGTATGGAGTCAAATACAAGCTGCCAGAGTACATATGCACGTTCCTTGTAGCTGGAGGAGTCTCCACATTTGCTTTGTTCAAGAGCAAGGCGAAATCAATTCCAAATCCAAATGCACCTTTGGGTTATAGTTTATGCTTTCTTAATTTGGCTATGGACGGTTTCACCAACGCCACACAAGATTCTATTTCTGCTAAGTATCCTAAAACAACCGCTTGGCACATCATGCTTGGAATGAATCTATGGGGCTCAATCTTCTCCTTTTTCTACATGTTTGCTTGGAGTGGAGGTGGTGGAGTTGAAGCCGTTAGGTTCTGCAAGGAGCATCCTGATGCTGCTTGGGATATTCTGCTCTTTTGTTTATGTGGTGCAGTGGGCCAAAATTTCATCTTCCTCACAATAAGCAAGTTTGGATCTTTAGCCACAACAACCATCACTACAACTCGAAAATTCATGAGCATTCTTGTCTCTTCCTTGTGGAGTGGGAATCCATTGTCCCCCCAGCAGTGGACTGGTGTTATGATGGTGTTTGGAGGGCTTTCAGTGCAAATCTATCTCAAGTGGCAGCGCAAGAGCAAGCAGGTTGTCGAGAAACGAAAAAGTTGAGTGTATTTTTCGTTGATGTCGAAAGGCATACCCCAATTATTCTTTACGAACTCTTGCATTAATCATTTATTGCAAGTTATACAAGAGGTTAGTCTTGGAAACATGGGATCCTTCAGAGTGTGGATCTTGTGTGTGTAAGCTAGTAACAGGACATCTATGATAAAGTTCTAAATGAAAGGGATAGAAACAAGTACAGGGAGAATACATATTGAGCATGTACTATGTAGACAGACATGTACATGTTTAAAGTCATGGGCAGCATATTCTTTTTGCTGGAGCAAATCTTGTAAGCTTCCCAGCATTTTTTGTCTTAATACAACAAAGCAGACATTGCTGCCAAGGGTTGATCTTGTCAATGTGAACTTTGCTAATTACCGCTAGATCGTATGTGTTTGGTAAAACTGTACTTGATACTGATTATGAAGGAAATGAGAAATGAAGGTGGTG

**TONOPLAST SUGAR TRANSPORTER 2 (TST2)**:

CTTATTCATGGTTGTTAAAGATAAAACCAAAAGAGTAGTAGGCCCCAACAGCATCCCCTCTCTAGATTGCAACATCAAAATTTTAACCATATACAAAGTGATTGGCCCCACTTAGCATGACCCAATCAGATGGATTTACATCAGTTTACAAAAAACATGTTCCATAGCATTACTATAAGGCAGTGAGCAAGCAAGCTCACTTGCTGCAGGAGTTTAGCTCTGTCCCCCTCCCCCCCCCAAAAAATGAGATCAAGCCTAGCCCTGTTAAAGCTCAAGATATGTCAAAAATGTATGGTTGGATCCTGCTTCATGCCTCAAAACTAAGGAGAACCCTCTTATTCTGTATGTAGCCAAAACCCCCCCTCTCTCTCTTCAGCTTCTCGATCACTCCCCCCGATAAGGAAAACCGATTATTTAACTGTCAAAGCTCCTGCACACTACATTACCAACCCCTGCTCAAATGCAAGCACCTCAAGTGGAGTATTAGGGACATCATGTGGAGTATAAGGGACATGATGTAAGGCATTTGCTGCGGTATCAATCCCCGTCAGTCTCACACGATGATGCAGCCCATCTTACACGTCGCCTTCGTTTTCGATGGCTGCTGGCTTCTTTTTCTCTGCTGCCGACAAAGCAAAGAATTCCACTATGACTTCTAATGGCATGCCTTTGGTCTCTGGCACCTTGAAGTAGACATATCCCCAGGTTATGAAGCAAAATACGGCAAAAATTGCAAACGTGTTTGCAACACCAATGGCTGAATCCAAGACCGGAAAGAGCTCTGTCACTAGGATATTGGTTATCCACATGGCAGCCTGGCATATAGCGATACAAACGCCTCTCACTCTTGTTGGGAAAATCTCAGCACATATCATGTTTGGGATGGGACCAAAACCCGCCATAAAGAAGATGACATACATCACCACACTTAGAACAGATACAGTAGCAAAAACGGCATCGTTTGTAGGCACAAGGTTGATGATGATCAAACACAGAAGAGTTATCACTAAGACTGGTAAGGCGTGTAGAAGGATTTTCCTCCGACCGGCTACATCCATAAGGCGCATCGACATGATAATGAAAGGAAGTGACACCAGGCTAGTAGCTGCACTGGCTAGAATAGAAGCTGAATCGGATCCAATGCCCATTTGAGACAACACCACATCTGCACCTGTTTCCTGAAGTATCTTGGGCGTGAAGCTGAGAACAGCACTGATACCACCGAACTGCTCAAGTATCTGTAAGGTAACTCCAACAATAAGAGCTTGTTTAACACCACCTTCAAACAATTCAGACCATGCTGGACCTTTTGTGGCGGTCTCTGCCGGATGTAACAATGCAGGTCCAACAAGGTTCTCTGATAAGATCTCCTTTCCATACTGAGATGGCCGGCTAATTAAAGCTGCGGCCGGAACAGACTCTATTTCACCCACTGTACTACCAAACCGCGGAAGCGAGTATGAAGAGCCCATTCCAGTCCCAGCCATTTGTGGTTCCTGGAACAAATATATCCTCTTATAGCTTCCATGCTCTGGTTTTCCATCAGTCCCATCTGGACCTGCCAATTGAAAAGCCAACTGCCAACCGCCACCAACGCCCATATTCCCAATCGACCCAACAAAGCCTTCATGTGGTGCAACGCTAGCCGCTACAGCATCATGAAAATTTCCTTTGATAGATTTAAAAGAGTCACCTCTGTGTAGATCTGAGACATTCCAATGCTCACTGCTGCGTCTTGAGAGTAGAGGGCTTGTCAAGTCATTATCAATATCACTTGTGTCTGAATGATAGCCATTGGTTTCTTGCAGGCTGCCCCCATTTTCTTCATCCTCTCTAATGTCATCAACAGCTTCCAAGTGGTGTCCCTGCACATGGCTATGCAGTATGCCAGGCAAATGATTAACGAATGAACCCTTAAGGCTTCCCATCAAGGTAACAGTTGAATCCATAAAAGGAAGACTGTTCGCGACCACTGGAGTTCCAATGACACTAGGAGGTCTTGAAAGAAACCCACTTGATCCAGTAAAGCTTGATATAGGTTTTGCAATCCACGCTCCACCATCATCAGGAGTGAAAAGTGTTATGGAATCGTCTTGAATACCAGTCAAGTCATCAGGGATAACGAAATCAGTTGGCTGAATGATATATTCTTCCAAGAACAAATCCTGTCCAACACCCAGACCTTCCACTAAAAGAGCCATCTCTCCTGATACATCCTCCTTATTGCGAAGCCGTTGTAGGACTTGTTTAGCTTCTTTTGTCCTACCCTTACTAACAAGCCATCTTGGTGACTCAGGCAAGTAATAGAGCCCGAGAATCAAGTAGATAAAAGAAGGAATAAGCGTCAGCCCAAGCATCATCCTCCAATTGGGACTGCTTGTAAGAGAGATGCCGAAAACCATACAATATGCCAGAAACATTCCTGAAGTGCCGAGAAGTTGAGGTAATGTATTCAATTCTCCTCGAATTTCTGCTGGAGACACCTCAGAAATGTACATGGGTGCCACCGTCACAGCTAGTCCAACTCCAACACCATCTACTATGCGAGCTGCCAATAGCAAGTATACATTTGGGGACCACAGCATAAGTAAAGAGCCAATGCAATATAGCGTTGCAGAGATGCATAGTATCTTGCGTCGGCCAAGCCAATCTGCTCCTGACCCTGCAAATATCGTGCTAAGAGCAGCACCTATAAGTGATGCAGCTGCAACAGCACCTTCCAGAACTGGCTGTACATCCAAGTTGAACTCGGGCTCTATGTAGAGCAAAGCACCTGCAAGAGTTGCACTGTCCCACCCATTGAGAAGGTTGCCAACAGCAGCTACGGCAGCGACAATCTTGGTGGCCATCATTTTTGCGGGGTGCTTGTGGTTTCTAATAAAAATTGGCTAGGGGGGAACTCAGACCTCACCTTAGAGAGGGATTTGTACAGACGAGGAGCAGAGAGAGAGAGAGAGACTCTATTATTGAAGGTGAGGTCTGAGGGCAAAGGTGGGCATGGGAGGCAGTGCTCCTTGTTCCAAATGGACCAAAAGATACAGCGAAGGTTTAGAAAG

**SUCROSE TRANSPORTER 4 (SUC4)**:

AGAGAGAATGCTGATAGACGATCTTACATTTCAGTGAAATAATTGAATATCCTATACCAATTCTCCTTCCAAGGGAGCATTTCTTTACATCAATAAGGGAGCCATAAATGTCGTATAGGAGTTATATTACAAAGGCATGATAAACTTTGGCCTTTCAGAAAGAGTCAGGCAGTAAACGAAGATCAGCGAATGGTGTTATGTTGTATTGAGCTACTTTGAAAATCAGGGGGTGGCTTGGGCAGCAAAAATGCAGCCATTATGCCACTTACTAGGGCAGCTACGGCTGCCAACACAAAAGCAGGCATATTGCCCCCACCAAACAACTCATCCCATGGCCCACTTCCTACCGAAACCACAATCTGGCGAAACATAAAAGAAACACCTTAAGAACCTAAAAAGCTTTTCACATGATTTTCCACATTCCATGCCATCACACAAAAAACTTTCAGCCGCCAAAACTCCCACTTCTCAAATTAAGACAACATCGCGAGATTCTGCAGGGGAGGCACTAGCCACACTCGAGGGGCTGGCAGTCCTCGCCCCCTACCGCACCTACCTAGGTATGCAGTGCCTCCCCTGCAGAAATTGAATACTAATGGAGGGACCCTCTGCAATGTTCAAAACTTCTTGTGATGAATGTGGAATGCCTTTAATTCCCAACAAATACACAGACTGTAAAACACGGTGAGTTGTTTAGAAGCGTACCTGAGGAACGACCACTCCTAAATTGAGCACCCCCATAGAAAGCCCTTGGCCACCACCTGCAGTGTTTGTGATCATTGTAGTCACAGAGTATGGAACACTATACGTGACCGCGAGTGGGATCCCTAGAATTGCAAAGATGCAAATTGCCGACCCCAGGATACTCATCGGCAGATAGCCTGATGCTTGGTAAATCTGTTTAGCCATGATTGTGATAACAACGGTGCTAAAAAAACAAGCGGACATGATCACATTGGCTGCAGCCCATACCATCCTCTGCCCAACTTTACGACATAAAGGTTCAATAAGCAATGATGTGATACCGAGCACAACTGAATTCAGCATAAGCCCAAAAGATCCTGCACGCACACCAGAATTGTACAGCTTCTTCACGCTTCCATCTGCCGAGGAAGGGTCTCCAAGAAAAACTTCTCGGCCCATCCAATCAGTGTCAAAAAGAATGAATGGAAACCATGCAAACCAAGTCAGAGCTGTAATCAACAGGATGTGACGCATGGGCACAGGGAGGTCTTTGAGTGTGCCGAAGAGCTCCCAAAAGAAAGCCTCCTGCTCCACCTCGTTGACATCGTCATCCTCATCTTCTTCCCCACTATCTTGCAGCCCCACTTGCTTTCCTGTAACTTGAGTTGAAGACTCCTTGCACAACAAGAGAGGAACCGTTGCATTGCTGTTTTGCTTCTTATTACTGCCTGTCGACGAAGACCATGGTATCTCTGATGCTGCCGTTAAACTCAAGAATGCGGTCACAATCAATATGACAACAGCTACGAGGAAGGCCGACTTGAGATTGGCACAGCTCTTGCTGCATGCTGTTGTCAAGGTGAAGGGTAAGATTTTGTACCAGCTGCTATATGCACCAGCCGCATAGCCAAGCACATTCCCTACAGCCATGCATAGGGAGAAAAAAGCATTGGCCCGACGTGATCTGCGATGATCCTTACCTGTGAAGTCTGCAAGAAGCGCTCTGCATGGACCCTGCAGGGTGTTATTTGCCAGGTCAAGGACCCAAAAACCAACGACAAATACAAAGATTGCGCGAGGTCGAGATGTTGTGCTATCCCCAAATGCATAGCCAATATCAGCTGAGAAAGCAATGATGATAACAGCCAATACAATGAACATTGTGCCTGTGATTATGAAAGGGCGCCTTCGACCCCATGAGCTTTCGCAGTGGTCACTCAAGTGCCCAACAAGAGGCTGGATGAGCATGCCAGATATGGGCCCACAAAGCCAGATATAGCTTGCCCATGCATGGGGAATGCCAAGCTCTATGACATAAGGAGTGAGTAAAGAGAGCTGAAGCGCCCAACCGAATTGCACGCCGGTGGCAACCAATGAAACCCGAGCAAGAGCCCTCAGAGGTACTCTGTTCTTGCCCTTGTTACTTTTCAATCCTGTTACACTCACAGATACTTCTGTACGATCCATGGAAGAATCGAACCCGTCACAAATAAGATGGTATCAAGGAATTGCAATTTGCAGCTGGGGAAATTGAACAAAATTGTGGAGGTTTTGGTTTTAAAGGAGGAAAGATTCCTTCTTTACAGAAACTTGTTGCAGTTATTTACAGGGTGGCCGCAGGGCACCAACGACAGCGTACCTGAATTTTGATTTTAATTCCTTGAAAGATTATCTGTGGGTGAAAACCATAATGAGCTACAAACACACAGAGGCTTGATCCAATGAAATGGGTAACTTGGCCATAGGATTGCACCACAATTGCAGGAACCTAAAGACAGCAGAAGAAATGCGGGTAATACACCTGCTCTGAATTAGATGGATGCGCATGGACGATGAGGCATGCCGTGAACCTTAAACAAGCCCTATATGGTGAATAAACAGCAGTATGTGCCCTGCTTTCTCTGCGGATTAAACCCTAATCTTGGGCCTGCCGCGTCTTTCTAGGCCCCAGAAAACCCAAACCCCTAGACTTGCAAGGCGTCTTCTTCGGCGTCCTTCGCCCCCAGAGACAAGATGAGTCTGTGGATGATCGATGGCGTGCAGTGTGTCGTGCGCGTCCCTGATTAACAGAATCAATAATTGCCTCCCCTTAAGACATCTTTTTA

**SIGNAL RESPONSIVE 1 (SR1)**:

GATACTTTCAAGAATTGTAATGATTTGCAGCTGCTATGATTCAGCTGAGGAACTCCAACAATACAATAATTTTATCAAGCTCGTTGATTTTCATTTCAAATCCAAGAACAAAAAGCAGAAAACACATGCACTAAGCAATTGAAGTGGATTCGATATCAATGACAGCTCAGGTGGTAGTCCTCCAGCTTCATGGAAATGAAGTCTTGCCACTACAATAAAAACTGGCTACCGAAGGCCTAAAAGTGACTTTGAGCAGCGCGCACAGCGCAAAACTGATAGTGAAAGTGGCTAAAAACATGAGAAAATATACTTGCTGATTAGATGGTGCAAGCACATTTTCATAATATTTGGTTCACACCGACCCTCGGGTGATTTTTGACACCTTTTAATGAATCCAAAAATGCCATGTTTCTCTCCAACAAACCCATAATTTGCTGATTTGTAATAATGTCATCAAAACTCAAGAGTCCCATTAAAAACACCAACTTCTGCACATGCTCTAGTGCAAGACCTCACTTTTAATCAAGTTGCACTTCATTGAAGTTGAAAGTTAGTCTCACTCAAATTCTGTAAGAAGGCTTGAAAGCAATCAGACATTTATGCCTCCAAGACATTCGGACATGGATCCATCTGATAATTAATGATCAGAATCAATCATTACAAAACTTGGCAAAATTAGCTTGGTTATACTGCACTTTATACAAAGAGGAAAAGAGTTCCAAGAAACAAAGCTTCGCCCATAATATGAACTCTGCCCTTTTGGGCTCTTGCCTTCCCACTGCCCCTTATCTTGCGTGTGTTATCTAAATTGTTGTAATAAGAAGGGAAACAGCATAGCTGATGCCGAAATTCTAGCTTCCCTTTTCATGTAATTGGAGTATAATTTAGTGGTTCCTGTGAAACCCCAGTAAATCAAGCAGGCTATGAGAACTAGATAGCTTCATGGATTGCGTAACTCAAATAAGACTTGGGCTAGTGGAAAGAGTTCTAGCACACATGGAAGATTTTGCTAATATAACTAGAGGGAGGAAACTCATACAAGAGCTGGACCTCTACTATGATACATCAAATGCATCTACATGTATAAGACATCAAATTTAAGGGATGGTTAGTAAGACCTAAGGAGCTTGTATCACATCCTTGAGTAGTGTAAAGCCGAGCACTAGCGATATGTTGACATAGCGTAATCTTCAGAAAAAAATTGTTCTTTGTCCTCCGTGATAAACGAGGTATCATGTGTGTTACCTTCAAGATGGACCTTTGTCTTCTGATAGCCTTCAACCAGCCTGCGATACTGTTCTCGGGCTTCTGGAGATCTCACCATAGATTGTACTCTTGCAAGAGCTTTCTCCAAACCTACCTCGATCTGTTTTCTCCCAGCCTTCAAAAAGTCCTCATCCCCATCACTTCCACCCAGCTCATTTGCTGTCGAAGACGTAACTGATTCAAAACCACGCAAACCTCTGCCTTTTCTCCGCCACCTTAAAATTGCTTTCTCCACAATACTAACTGACCATATGATTTTCTTATACTGCTTTCGAACTTGATGTCCTCTAAAATGAGCCTGAATTTTAACAACATGCTGACGAAATAAAAGAAAGTTCCGACGCCCTTTCCATCCACGATACTTCTGCTGTATGCGAATAGCAGCAAAATGCGATAATTCATCGTCTTCATAGAGTAATCCCCCTCGGTTTTGCTTTTGGGCAGCAGCAATGTACAGAGCCTCCTCATTCGAAATCCCATATTCATCCTCCTCAAGCTGGTAAGACTCCTGCTTCTTCCGAAATGACTCCACACGAAATTCAGACTGGATGCGAGCTGCTGCTTGAGCAGCATTTCGCACAGCTGCAAGCGAGTCTTGCAGAGACAACTGCTCCTCAGAACCCCCCTCAATGGACTGCACAATACTCTTTTCTGAGAGAGATTCTACCGCATCAGCAGTAGCTGGAAAACTCTCACTTTGTTTAGACTTCAAAGTCAGTGATGAGAGATGACTTGTAAGAGAGGACTCTGCAAGATAACCTGCAATTCCTTTGTGCCGCTCAGCAGAGGCCAAATCAGCTGGAGTTTGGCCCGCCGGAAACACATTGGTAGGATCAGTTTTGGCCCCTGGAGCTGCACCACTGGCCATAAGAGCAACAACCATGGCCTCCCTGCCACAGTGGGCAGCCCAGTGAAGAGCAGTCCATCCACGCGAGTCCCGAAAATTAATATTCACTCCGGCAGCCAATGTTGGGGCGATGATCCAAGTATAACCAAGCGCAGCCCCCATGTGCAGAACTCCTAATCCATTATCATCCAAAATAGCCGCACCTCTACCTCCCTCAAAAATCTTGTGGAGCAGCCAAAACTGTAGCTTATCTTTCAGGAGCTTCTGAACAAACTCCTCTTTTAAAGTTGCTCCAAGACAGCCTGATTCAGTAAGCTGATCTTCCATCTCTTCCCACTCATCAAACAAACCCAAACAGTTCAATTCACTAGGAACACCTCTATTAAGTTTATCAGCTTGTAAGGTTTGAGATGTTGGTCCTGATACAATTGAGAGCATCCTTGCAAGACGAATCTGAAAAAGCATGTCTCTCTTGTTTAAGGGTGCCCCTTCGACTGTGTCAGCAGAAGAGAACTTGTACTCTGCATGATATTCAAATTCTTTGATTTCACTGCAAGCAATCTTATCACATCGGGTTACATAAAAAGGCACTTTGCCAGGTGGATGTGCAGGTGCATTGCATCTCAAAACCCCTGGTTGTAGAAGTTCTGCCGCAACCTCCAAATCCCCAAACATACAACTCCATTTAATTTCAGAAGGATCCTTTATCATGTCCCCAAACTTCCCACTCACAAGTACCTTGGTGTTAGTAGTTGAAACAGCACAGTTGGGTGAAAAATCAGTGATAGAAAAACAATTCTCGTGGGGGAGAGAAGGAATTGCTCCAATATCCAGTTGCATCTGCTGTGTCAGACCAGGGACTTCTTCTGCAAGTGCACTAGCCCAATATGTTGAGTCTGATAGCATCCCATCCCCTCCGATGTCTGTACCCATCCATCTTCCAAAACTTTCAAATTTTCTAAATGAATCTTGTTCATCCTGCTGAGTAGGAAAATCAGCTTGAAAGTCTCTGAATACATTTGAATAAGAATGCATTAAAGAAGCATCCTTGATGTCTGCACCGGGCATATCAAATCTACCAGAGTTAGGTTCTGAAATATAGGACTGCTCAGTGTCCTGATATGTCTGAGGACGAGGAAGTTGAGCCTGTTGGGGGTCATTCAGCAGATCATTATCAAACCCATTGAGAATATCACCCTGACCAGGACCATTATCAGCCAGCTGTGCCAAAACTGCACTCCAAGGAGTGTTCTGCTTGTCAGCCACCTGCACTGAGCCCAGTCCATCAGCAGCAAATGAATGGTATTTGTTTACTGGATTGGACAATGTATCAAGAAAGCCTTGTGAATTTGGCTGCCATGATGTTTCTGCAGCTGGAAATCTATTATCCGGTGCAGACTGGATAGCCCCCATCCCACCATTCCTGGCCATTTCAGAAGGATCCTTAAGCCATGTAGACGTATTGCTCGTTGTCACAGCCTTCTCCCTATATTGATCAGTTCCAATTGGTAACACCATGTTGGTAATTCCAGAGCTAATTCCAGAGCTTTCCTTGCTTCCCCAGCTTTTGAGCGCCTCACAGTATTCTAACAGATCAAATCCACCAGGTGGGGTTACAGCACCATTTTCCTGCTTTACAGGTAGCATTTGAGCTCCACCTGCCCCAATTGCCAAAGGATAGGCTGCTGCAGCAGAGCCATTTGTATTTTCATATAAAGATGGAAGACCTTTTTCTCCATAGCTGACACCATTTTGATTTCTTGCAGAATAAGGCATGGATCTTGGAAGGCCTCCAGCATTTGGGACAGCTACTCCTGCTCCCCGGTTTGTAGCATTACTGGCAATAGCGTCTAATTCTCTTACACGAGACTGATCCGTATCATCACTAGATTCAGGTTCTTCACCTTCAGGTGAATACGTTGGTCCACCATTCCAGTCTGCACTAGTATTAGGGCTTGAGGAACGAGGAGAAACAAGGGGTGATGTTTGAGGGGAAGAGCTTGCTACCCCACGAGAATTTCCAAATGAAGCAGTCCCATCAACTGAATAACTGTTTGGAGAACGCAAACCGCTTCTACTGCCCTCTTTAACTTCCCGATAATGTACCAGCACTATGTGCTCATTAGTATCTAGCATCCAATAGCTCCGTCGCTGGAAATTTTCGTTCTCTTCACCATGAGCATAATAACAATGTAGAACTTCAACACTTCCTGCCTTCAATCTTTCATGAGCCTCCCTGACGGTCTTTCCATCCTTCTTCTTACGCCAATTGTGACCATCTTTCCGAAAATACCTCAAAGCCTTGCGATCGAAAAGAAACAATGAACCACTTTGTGGCTTGTGAGGGGGATCTGGTGTGAGCTTGAACTTGCGGTAATTGCGAAGAATTTCACAGACTTCAGTGGGACGAAGCCATCGATGCTGTGCCTCTTGAAGTATTTGACACAAAGCAGCTTCTGGTTGATTTGGAAGCCCTCCATAGCGGCGATGGTCCATATCTTAGCTATAGGATGTACACATCAGAGCATTCCAGCTTCAAAACCCCTAAACGTCAGCAATTGGGTCGCAGAAGCTTTCCTTCAGAGGACCAAAAAACCACAATAAGGGGTTCTCGAAACAGCAAGAAGTGAAGGAAGAGGGAGGGGAGGGGGGCCGGG

**CERAMIDASE (ACER)**:

GGAATGTGCAGATAAAAAGTAATGCTTTGTGCAAGGGTGTAGAAGCTTTAAACTAGTCAAAAAATTCAGATATCCACTTGAAAAAACCCTTCTTATCACTCTTTCTCATGTACATTGCATAGCCAATGAATAAAACATTCAACAGAACTATTGCTGGAAGTCTGCTATCTACTCGTGTAGAGACGTGATATGCAGTCACATCTTGTGGAAACTGTCGATTTTACTTCTTGTAGGAAGGGGACACTTCTTTCAGCCACACGTGGAACACCCAGCTTACTCAAGGTTCACTAACATTTGCAGTGGCCCAGATCTAAGTTCAACAAACCAGTCCAATTTAATAGGTCGACGGTAAGGCAAAAAATGAAAGAGATCATGTAGTAATAAAGACATCTCTAAAACACAGCAAGCATCTAGATTTACGAACCTTTTTGCACGCCCTGTACAGGACCTGAATCAGAGATTATACAGCAAAGAGATTAAATTTAAGGCATAAGAAAAAAAAAAAAAACAAATCCCATCAAATTTGTCATAGGCCTAGGCCAAATACTAATACCAGCATGTACAATGACCTTTCATCACTGTTGTTTGGGTTTGACCACCTTCACGTAAGGCAATCCACAGAAATAATGTAGCTGGGGACTCCAGTTAAGTTGCTGGGCGCGGCAAAACTGCAGGTATGTGTTGGCAAAGAAGGCGTCGAGCCCAATGAAGAAATGCCACCATGCATGCCCATGAGGGTTAATCGCCCATGTCACTATTGTGTCGCAAAAAGCCCAATCCAACACCCAACAAGTGCATCCTAACACCATTGTGAGTATGTACATATGTGCCAGCCACTTGGCATGCCGATCTGTAGTGTGAATGTAGTATTTATACATTCTGGGCAAGCAAAGTAACCCCAAAAAAAAGTAATGAATTTGATAGCTTAAAATGAACCGAAGTTGCGAGTGGATGATAGCAAAGATGGAACCATAGATAAAAAGCACTGTAGGCATTGTGCTTCGATAATGCCAGTCTGGGGAGTAAAGCACATAGAAATACATCAGCATCAGCCACACCATTGGGGTTTCTTCACTCTGCTGATGAACATGCTGCAATGTGCCATGGAAGAACATACTCCCCAGACCAACCATCATTGTCGCAAGATTTAATGAACTGAACCTCTTTTCAAACCTTTGCCTCATTGAATTCACTAAAACTAAAAAAGCTAGAAGTATGCCAGGCCCATTCGAGATAGTATTGTAGAATTCTGCTACATATGCAGAAACAGTGTAGTGACTTTCACACGTTGTGCTTGAAGTAATAGGTCCCCAGTACCCATTGATGTCAGCTGAATCTGCCATTACTCCCCTTTGAAATATTAATAACAGACTCCAAATCTTTACTGCTCACATTTGACCATTTTTGTAGAGGTAGAACCTCCTTCACAATGTCGGTGCAGAGGCAGTAACCCTAAACCAAGACATGATGAAGACCCTGTAGGAAGCCGGTGGAAGGACTTTGAATCGAAATCGAAATTAGGGTTTTTTTTATTCGCGTCGACAAGCTTATGTCTGTGTACGCGCGCAAGTGCGTGCGTGCGTGTGTATGTGAAAGTGAGAGAGAGGGGGCTATGTGGCAC

**AMINOTRANSFERASE ALD1 (ALD1)**:

GAGAGAGAGAGAGAGAGAGTTCTCGTAGAAAATGGCCGAGCTGGTGCCGGAGAAGAAAATGGCCACTGGCTTTTTTTATTTTGCTGAAAAAAAGAAATATTGGTGGTGGTTTTTTTTTTTATGCGGCTCGCCTTAAAATTATATAAAAGTTAACTTAACTTACTGAAACACTTGAGATGCTTGTAATATGGCTGGACTTTATTATGATTAATGAGAAGATGTGACTTTTCAAAAAAATCTTATTTTGCATGCAGGCAAAGGAGTACTCATGGTGCTGTCATACAAAAAGCATGCAGGCAAAGGAGTACGGAATTGTAGTTGGAAAATTAAAAATCTCACTTTATTTTGCTGTTGGCTCGTTGAGCTTATCAGAAAGTCCAGCTGACACCAATCATTAACTGCATCAACGCCATGGAACGAGGTTTTACGACCGGCGTTCAGCGCAATGACAATCTAACACACTGTGAATCTAATTTGTTTTTTCAAACGGCTTACGTTCGGGGTTTGCTTTTAAGGGGTTCTAGATTCGTGTGGAGCTTGATGCATACCTTAAAGCCCATCCGAGTGCAAAGCTGATAAGTCTTTCTGTGGGAGATGCAACTGAACCCATTCCACAAATCGTCACATCTGCCATGGAGCAGTATGCACGTGGTTTGTCCACGCTTGAAGGTTTCTCTGGTTACGGTACGCATCAAGGGATAGAGTCCCTGCGAAATCGGATTGTGGAACTGCTCTACAAAGATCTTGGACTAAAGGCATCGGAGATTTTTGTTTCAGATGGAGCAAAAGGGTGCTTTGCACGTCTTCTGATGTTATTTGGATCTGATGTTACTGTGGCTGTTCAGGATCCATCTTATCCAGTTATTGCCCAGATCTGCAATCTATTTGGCCCATCAATTCAAGATAAAAGTCGAGTTTTTTATTGGAAGAGTGAAGCTGAGAATAACTTCTTCCCAGATCTATCGTCTGTACCACGCACAGATTTGATTTATTTGTGTTCTCCAAACAATCCAACAGGAATAGCAGCCACAAGGCAACAACTGGAAGAACTGGTTGCGTTTGCCAGAAAGAATGGGTCCATTATACTGCATGATTCTGTTTATTCTGCCTTTGTAAATGATGACTGTCCTAAATCGATCTATGAAATTCCTGGAGCAAAAGAGGTTGCTATAGAGGTTGGATCCTTTTCAAAGCTAGCAGGGTTCACTGGAGTGCGTTTAGGATGGGCTATTGTTCCTGAATCACTTACCTTTGCTAATGGGCAACCAGTTCTTAAGGACTACGGTAAAGTCGTGTTGCTAACAGCGCCAGCTGGTCCTTCAAACATTGTGCAAGCTGGTGGTCTTGCTTGTCTTACTCCGGAAGGTTTAAAGGTGAGTAACCTTGTAAGGACCTATATATAGATATGGTTGGGCGGTTAATCACATGGGTTCTTCCCTCTACTGCTTACAAACCTTATGAGCATCAACAACAAGAAGATATGCGATCCTACACTGTTCACGTTCCAAGGTAAAAAGGGTTGCTCGAAAGTGCTTTATGGCTTGTTTCTTTTGCAGGAATTGATGAAAAGAATTGAGCATTATAGGGAGAATGCAAAGATTCTTTTGGAGGCATTTGCTTCCTTTGGACTTACAGCATCAGGAGGTAGAAACTCACCATATATTTGGGTTGAGTTTCCAGGGCGGAGCTCTGATGATGCTTTTAAGGAATGGTTAGAGATTGCCAATGTCATGACTTGTCCAGGGGTCTTGTTTGGACCTTCAGGGGAAGGCTACCTCAGGGTCAGTGCTTTTGGACATAGGGAGAACATCCTCGAGGCAGTAGAACGACTGAAATTAATTTATTCTTCATGAGCAAACACAAAAAGAAAAGATGTTAACATTCATTATTTTACGAAGAGGGCCTTCTGAGCACTAAAGTGCTCTTTTCACATCCTCTTGTTGTTATGAACATGTGAGAGAGAAGACTATCCTTATGTGAGAGCCTCACATTATCACTCCCATTGTGAGGGGCAAGTTTTTTGTGCAATTCTTTAACC

**β-GLUCOSIDASE 42 (BGLU42)**:

GGGCAGCTGTTTTGTGCGTGCGTTGTAGGAATAGATCTGTGTGTGCGCGCGCGTGTTATTTTGTGCGTGTGTGTGTGTGATTTGTGCGTGCGCGCGAACTTGTGCTCGTGGGTTTTGGGTGTGTGCGCGCCGCTTGTGTGTGTTTTTCATGGCGCCTCCCTTTGGGTCGTACCAGGAAGATGAGGTATACGAGGTCTACGAAAGCCCTGCTGCTGTTGCGCCTCTCCTCTCCAGATCCCACTTCCCCCACTCCTTCATCTTTGGAACCTCCACTTCTGCTTATCAGGTTGAAGGTGCCACAAATGAGGGCGGCCGAGGCTGTAGCATTTGGGATACATTCACTCAATTACCAAATAAGATTGCAGATGGAACGACTGGAGATATTGCAACAGATCAATACCATCTGTATGAGGAGGATGCTACATTAATGCAACAATTGGGCTTCGATTCATATCGATTTTCCATCTCTTGGCCAAGAATTTTTCCAGATGGAAGAAAAACCTCATTAAACAAAGAGGGCATTGCATATTACAACAGATTGATCGATGCTTTGTTAAGGAAAGGAATCAAGCCTTTTGTAACCTTGTATCATTGGGATCTCCCACAAGCACTTCAGGACACAATTGGAGGGTGGCTTCACCCTGAGATAGTAGAACATTTTTCAGATTTTGCAGAGACATGTTTTAATGAGTTTGGGGATCGTGTGAAGCACTGGAATACTCTCAATGAACCGCTAAGGTTTGCTTTCTTCGGGCATGGTATAGGTATACATGCGCCCGGAAGAACTTCAAATAGAGCAAGGTCTGCTGAGGGAGATTCGTCCATGGAACCCTACCTAGTTGGGCATCATGCTTTGCTAGCTCATGCTTCAGCTGTCCACATTTACAACCAAAAATTCAAGGCGCAGCAAGGTGGGGTGATTGGCATTGCTGTTGACTCAGAGTGGGGGGAACCTCTTACAAACACGAAGGCAGACATAGAGGCAGCACAAAGGCACCTTGAATACCACCTTGGATGGTTTTTGGATCCTATTTTCTTTGGAGACTATCCTTCTTCAATGAGAGAAACAGCTGGTGATAGGCTGCCAACATTTACTCTTGATGAGCAAGCCCTCTTGAAAAACTCTGTAGACTTTGTTGGAATCAATCATTACACCACAAGATATGTAAGCCCAGCGGACCCAGATGCAAAGCCGACCAATGGTGGCTGGTTTGATGATATCAAAACAAAGAGAATAGCCGAAGTGGATGGAATACCAATTGGAGATAAGGGTGCATCTCAATGGATGTACATTGTGCCATGGGGATTTAGAAATCTCGTGAATTGGGTGACTAAGAGATACAACAGGCCTCCAATCTACATTACCGAGAATGGTATGGATGATGAAAATTGCCCTTCGAAATCGATTGAGGATTTTTTGCACGACACAAAGAGGGTGCATTACCATCAAGATTACTTAGCAGCACTTGCCCAATCAATAAGGGAGGGTGCCGATGTCCGTGGCTATTTTGCATGGTCTTTGATGGACAATCTTGAATGGGAGTCTGGATTCTCGAAGAGATTCGGGCTGATCTTTGTGGATTACAGCAATCTCAAAAGGTATCCAAAGGATTCCGCCCATTGGTTCAGCTCATTTCTAAAACACAAGCCATCTGGGAAGCAGACTCACCTTTATGACTTCATGATAAACCATGCACATATGAAACTTGCAAGCCCACAAGATATGGACCGCATAACAGTCATCAGTTAAAAAGCATGGTTTTTTTGGAATGAAGTGGATACACAAGATATGAAGAGCTAAGCTTCGAATTGGTGCCAGCTAGAGAGTAGACAATTGATGTGTGGGGATTGCTAAATGCCATCAGAATTGGGGGGAAATATCTCTACAGTTCTTTTCTCCACTTGCTATAGTTTATTTACCATTGTGATGGGTGAAAATTCACTTATTAATCAGCATTAACTATCTTGATTAATATTGTATATGCTGCACAAATTTGTAAAATGTGCGCGATCGATGGGAATCTTAATGCGAGAAAATTGGACGTCCGGGGGCATTACAGCGGAAAGCAAAGGAAGAGGATGGGAAGAACCATCAGTCCAGGGTAAAGCTCTTGCCCCCCACTCTGCAAGCAAAGAGGGGTTGACTTCAAAGTGCTGTTTATACATTAATGATACCTATGCAAAAAAAACTTGCATCCCACCTACACTTAAATTATCCCAATCAACAATGGCCCCTCTTGTAAATTAAATTACAAAATTTGTAATCAAGAATAATTATAGTTCTATGTAAATCTCTGTGTGTGTCTCCCTCTGCGCGCACACATGCCTCACACACACACACAATTTCACAAAAATATACCTTGTATAAATGAGATTTATTATAATCCACAGCCCCACAAGCCCTTTTTTTTTTGTGAAAATGAAAAATTTAAACAAAATAAATTTAA

**L-TYPE LECTIN RECEPTOR KINASE S.4 (LECRK-S.4):**

TTTAACTTTGAGTACCCTTGATTACATTGTTAGCAAATCCCTATTCCCAAAAACCATATGAGCAAAATGTGATATACCACAAAAAACACTTTGGGTATGTCTAGAAATATCAGATTGATCATAAAATCTAAAGGAACACAATCACACATGTTTCCAAGAAGCCACATCCATGTAAACACTAAGAGAAAAGATGAGCATAGCTGCTAAAAGAGACCTCATTAGGAATTTGGCATGCTAAAGTTCATGAAGAATATATTAAGGCAACTTCTTTGCCTTTGCGCTATAAAACTGCTAGACTCATAATTAATGTATGTGGCAAGTCTTAGCATTACATGCAGTTGCAAGTCATATGTTGCATCCTTTTTTTTTTTTCTTTTACCAATTGCGAAAGTAAACCCTCTCATACCCCCACAACACAAACCCTCTCAGATCCATAAACTAATGCTGAGAACACACTCAAGGGGGGGCAAGGATAGGCCATGAAACACTCATGGATGGACATGCCAAATAGGAGTCCGTGATGCACACAACCTTGTTAAGGTAGCTGAGAGCTACAAGGCAAGTTTTCTAAATATAATTATTTGTCTTATGGGGTTTGAACTGAAAACCTCATACTTATAAGCTTCTACCTTGGGCCTGAAAGATCATCGTAGTCTATTGTTGATGAAGAAGCATCATAAGCACTTGAAAGTGGCTTAGATGTGAATACTCCACTTCTATACACCCTATTGGTTTCAACCTGCACATATATGGCCACGGGCCTAGACATGGGCAACTCCGGCAGCTCTGTTTCTCCCCTCAATATCTGCAACACTTGCTGCGTCGGAAGCCTCGACTTGGGATCTGGGTGTGTGCATAAAAGCCCTATGTGCAACATGCATCTCATCTCATCCACATCAAACTCACCTTGCATTTGCCGATCAGCTACATCCATCAGCTCCCCCTCCTCATGCTTCCTCCACACGAAGTCCAAAAGCGGCTCATCAGTGTTTGAAAAAGGGTGTCGCCCGCATGCTACTTCTAATGCCAGAACTCCAAAGCTGAAAATGTCTGACTCCGGGGTAACTCTGCCGGACATGGCATATTCGGGTGCCATGTAACCGAATGTGCCCGCTACTCCGGTTTCCGAAGCTTGCGTGCAATATTGATCATGATCGAAAAATCTAGCAAGACCGAAATCACCAAGCCTTGCATTTAGATCAGCATCCAGCAGCACATTGCTAGCTTTAATGTCTCTATGTACAATCATCTTCTCCCAATCCAAATGTAGATAAACCAGAGCAGAAGCTACTCCTTTGACAATATTGAACCTCAAAGGCCAGCTCAAAAAATGAGGGCTACTTGAATACAAGAGCTTTTCCAAGCTCCCATTTGGCATGAATTGGTACACAAGCAAGAGCTGTCCTCTGTCACTACACCAACCCAACAAATGCACAAGATTCCTATGCCTTAGCTTCCCTATTGTCATGACCTCTGATTTGAACTCTTTCTCCCCTTGTTTTGACGCCTTGACAATCCTCTTCACTGCTACATGCTCCACCCCTGCGTTCTTAGTCGTGATCACACCTTTGTATACGTCCCCCGAACCCCCATGACCGAGCATCTCCTTGAAGTCTCCGGTTGCCATTTGGAGTTCTCTGTATGTGAATTTTTGAGGAGCGGTCACTAGGTCGACATCAAGGCGAAGACCTGTAGACATCCTTCGTTTCCATCCAATCAAGAGAATCACCAGAAGAACTATACCAGAACCTATGCTTATACAAGCTATAATCAGGTTCCTTTTCCTGTTGGATGATGGTGGTGAAGGCGAGGATAAGGGTGATAACGATGGTGTTGGCACCGGAGGAATGGGAACCGACACAAGTTGGCTTGAAAAATTCCAATCATGAATTGCATTCCTCTCTTGGAAAACCCCTGTACCTGCTGTGAAGCCAACATAGCAGCTACTATTTACATAATCCCAAATGCTTAATTTGGAGTAATTCAGAACTTTTGTTGCGGATGCCTTTTCTGCTGGTGCCACGGCAAGATACACATCCAGTAGCTGTTGTGCAGCATCATAGTCTATCCATGTGGTGAGGCGGCGGCTAAAATTGGTGGTTTTCAAGACCAATTCTGAAGGCAAGCTTCTAGCTAAAGTGGAAGTGAGGCCATTGATATCCAAGCCAACATGATGGTCATCGGGGTCACCATAGTCGGGGTTCAGGAAAGTATCGTATTCGACAGCAACAATGTGAACGAGGGATGCATTAGAGCCCTTGTATAGGTCCACATCGAAGAGGCCCAAGGCGCCTGCGGTGGCAGATCGGGTATGTGGGAGGAGGTTATTGGGCGCTAGTAAGAAGGTGAAGCCATCACCGGAGATGTCGGTCTGATTGGTAGCAATGGAGAAGGTAAAGGACGTCGAAAATGAGGCGGCGAGGCCTTGTGAAAGGTTGTGCATGAGGACAGGGGAGCTGTGCAGGACTCTTGAAATGAAATAAGGGCCAGTAGCTCCAGTACTCTTATTGTAGGTCATGTATAGCGTTCCCTCGTCGCTGACGACCGCTTCTCCGAGGATGAGGAGATCATTGGCAGAGGAGAAATTATTGAATGTAAAGCTCACATTTTGGGAGTAAAGGGGCTTAGCAAGGGCTAGAGTAAGCAACAGGAATAATGCTTGCATCTCTCTCTTATTGGGTAATGCTTGCATCTCTCTCTCTCTAGTCTCTCAGGTTGCTTGCATTCAGCTTGTTTATCTTTGGTGTACATCAGTGTGTTTGTGTGTGAGAGAGAG

**GUANINE NUCLEOTIDE-BINDING PROTEIN SUBUNIT β (AGB1):**

AGAGAGAGAGAGAGAGAGAGCTCTTTCAGCAAAATGTGTTGTTTGAATCTACAAGCAGGCAATGGTACCTGCATGGAAGCCATCAATTCAGGGGGCCTTGCAAAGCAAGCCTCTGTGTATTCGTTGAAGAGATAATTGCAAGGGTTTTCTTCTTTGTGCGCATGCGCCACTGGATATGGGGAGCAAGATTGGGAGTGTCTACGAGTGGCATTAACAACACTGATCCAGATTGCAGGCACGCGTCTGTAGAGAGTGCGCATGTGCGTCCACAGAGGAAAAGCAGGAACAGGATCAATCTCGCACACATGCTGTATAGGAAACACGAGAACAGCATTGAGGTTGTGCAAATATGCGGCACTAGAGGAACAGAGGAGGAGGATTGAAAATGACCTTCTGATTTGGCTTGAACATCTTTTCTTTGCGCTTGCAAAGCTTCTTCACTTGCAAACACTTGGATACCTTCAAGCAAAAGCAAAGGTTTTTGAAGTCACGCGCTCGAAATGACGTCTATGGCAGAGCTGAGAGAAGAGCTCAAGCAGGCAACTGAACAGGTCAACCTTCGGAGAGAAACAGTGCTACAGAAGCGCATGCGGCTTCTTGATACTAATGTTGCAGCATATGCTCGGCAGAATGGACATACAAACACCATTCGCCTGCGTCATCAGGATTTTGGCTGCTGTAGAACCCTTCAAGGACATTTAGGCAAAATTTACTCATTGGATTGGGCACAAACGATGGATCAAATTGTGAGTGCATCACAGGATGGCTATCTTATTGTTTGGAATGCTTTGACGAGCCAAAAGATACATGCAATCAAACTAGCTTGCTCTTGGATAATGACATGTGCATTTTCGCCCAGTGGACAGGCAGTGGCTTGTGGGGGGCTTGACAATGTGTGCACCGTCTATCCGTTGAGCTCTCAACAAGGGTCTGCTCGTCCCGAACCCAAGGTGCTAAGCGGGCATAAAGGGTACATTTCCTGCTGCAAATTCATACCCGATAAAGGCACCCAAATTCTAACAAGCTCTGGGGATCGAACTTGCGCTTTGTGGGACTTAGAAGGCTTCCAGAAAATCTCTGTATTTGGTGGGGATACATCATCAGGTCATAATGCAGATGTAATGAGTTTATCTATCAGCTCAGCAAATACGCAGCAATTCGTGTCGGGATCGTGTGACAAGACAGCACGATTATGGGACATGCGTAGCCCAGGCAGAGCAGCCCAATCTTATCAAGGTCATGAAGGGGATGTTAATACGGTGCAGTATTTTCGTGATGGTCTGAGATTTGGGACAGGATCTGATGACAGTACTTGCAGAATTTTTGACACTCGAACCGGACATCAGCTACAGCAGTATAAACATCCCGATCAAAGTGTTAAAGCTAACTCGATAGCCTTTTCACATTCAGGGCGCCTTCTTTTTGCATCATATTCAAATGGAGATTGCTGTGCATGGGACACTCTAACTGCCCAGGAGGTGGTTAATTTAAAAAGCCTACCAGGTGCACACACGGAACTTGTAAGCTGCCTAGGTTTAGTTCCTGATGGAAGCGCTCTGTGCACAGGAAGCTGGGACCGAAATTTGAAGATTTGGGCCCATGATGGGATAAACCAACACATTTCTTGACTCGTGGTCTTTGCAGTCAACATAAGCTATGCCTGTATTCTTTGTTAGCAAGTCCCTTGCTTGTCTCGTTGCCTCAGCATTACTTTCTAATGTCAACAGGAAAGACAGACTGTAAGAAAGATCCTAGAGTTCACCAATCTTGCTTCTAGTGGAGCTTAGTTTAATAAGCCATGAAATTACTCCCTGAAAGATGTGCGCATACGTACAAGGGTGCTAACCTTTGGAGTTTTCTCAAAGCTTGTCGATCTTGTATTGAAGAATTGGATTTAGGCTTATCAAGAGTTGTACCTCGTATGAACGGATGTAATGATTCTTGATTGTTTTACGATTCTCACGTTAGCTAATCACATGTCATCAGCTTTCATTCATAAAATCTCTGATACATGGAACTTATTTCTGTTTGAGGTATGAAGAGTACCAAACACGGCTGATTCATGTACCGTATTCAAAACTTTTTACTCGCGTGGTTTTCATCATGTCATCAACTTTGGAAATTCAAGCGATTTATTTGAGATTTCAGTCTA

**HISTONE MONO-UBIQUITINATION 1 (HUB1)**:

CAAAGTGTAGCAGCGACCTTTTTTTGGCTCGAACCCTCGTTCTCAGCCTAGTAGCGTTTTTCCTGTATTGTGTGCGCGCGCGCGAGCGAGCGAGCTGCTGCGGAGATAGAGAGAACGTGCGCTAACCCTAACTCTGTGTTTACCCTCCTCCTCCCTCCTTCCCCTCTCTCGTGAGCAGCGAACTGGAGTCTGTGGGCAAGTATCATGCCCCTTTACCGCACCGGGGCAGCAGCATCATGGGAAGCACAGAAGAGCCCGAGCGCAAGCGACGCCATCTCAACCATAGCTCCATCTCTCCTGCTCTTAAGAAGCAGGCTACCTTACCTTCTGCAGACGAGAAAAAGGTGGATGCAGCAATGCTGCAATACCAAAATCAGAAGTTGGAGCAGCAGCTTGATGTTCATAAAAGTGAAATCAGTGCACTAGAGAGCAAATGCAGTCAGCTCAAGAACAAACAAGCTTCTCATGAAAGTTCTTTGCAGTCAGTTCGGAATGCATGGGGCATGTTGTTGGAAAAGGTTGAGCATCTTGCAGTCCGAGCTGCCCATTCAAGTAGAGGAGAAGCATCAGATCACTGCATCTTGTCAAAAGATGCTACTAGTACCCGTGCTCCAGCAGAGGAAGCTTTCCTTGAAAGGTTACTTGAGACTGGTGCAACTGAGAGCAGCAATGAGAGCAATGACACTGCTTCTGCCGAGCTAACTTTGAAATCGTGCAAAGTGGCCACACTGAAAAGTGTAGAGCATCTCGTTCGTGCGATTGATTCCCAGAGAGCCCGCAGAGAGGAACTTTTTCTGGTGCTCAAGGGCAAGCTGCGAAATGATGATAGACTGCCAGTGCAACAGATTGATGAAAGTATAAGTGAAGTTGTTGCATTGAGGTCTGCTATGGATACTATACATTTGAAGCACAAGGATCTAGCATCACAAGTTCGCTTGTGGTGGAAGAACACAGCACGGGATCAAGCCGAAATAAAGCTTTTAACTGGCGAACTGGAAGAAACGATCATGGACCTGGACAGCACTAGGCAGAAGATTGCAGCTTTAAGAAATGAAAAGGATACAACTTCAGGAAAGCCTCTTCCAGTAGTGGGGGTCAAATGTGAGGCTAGTGAAAAAATGGAAGAGAAGCCTTTCCAGGATACTAAGGCCTTGGAAGGTGCATTGGAAGAGGCCAAAAGTTTAGCTTCGTTGAGGTTGAAAGAGCTAGAGGAGGTACAAAGAGAGAAAAGTCGTCTGGCTGAACAGCTCCAGAAAGTGCAGGAGGCACTAAATGGTGATAGGCATATAACAATGTCGCACCTGTATATTATGTTGAACGAGCAAGTCGAGGGCCTGAAAAAAGAATTGGAACAGTATAAGTCTCTTATGGCTAATTCAATGATAGAACAAGATAGCGTTTCAAGGGCTAATGGTGACACAGATGTAAAAGCTGAGGCCAGTGAAGCTGGGGAGGATGGAAACACAGTGTGTGAACAGAAGCTGTTGGAGTTGCAGTCAAAGTTGCTGGATAGCATATCGAAGAGGAACAACCTGAATATCAGAATGGATGAAACATCTCATGCCTTGGAGCGACAAGATGAAGTCGCGGAGTTTAAAGTCATGGTATCAACCTTAGAGAAGGAGATGGGCATGATGCAGCAACAGCTGGACAAATACAAAGAAGCTGCCTGTGGAGCTGAGTTGCTTAGGGCTGAAGTGGAGTCAACAAGCTTGTCTTTGGAAAGCAAGACAAGCGAATGCAAGTTCTACTCTGATACAATTGCAGACCAGCTTCTGGAAGTCAAGTCTCTTCGAGATGAGGCAAGGCTACTGCGTGAAAGTGAACAGGAGCTCAAGCTTATTCTTGACATGTATGGTAGCGAATCAACAGATTCTAGGGACTTAATGGAATTGAAAGAAGCAGAATGTTGTGCATGGGCTGAAGTGGAAAAGCTGAAAGCCGCTCTGGATGATCATGGTCTTGAGCTAAGAGTGAGAGTTGCAAATGAAGCAGAGACAGCTTGCCAACAACGGCTTGCAACAGCAGAAGCAGAAATTTCTGATTTTAGGCAAAGATTGGATGCATCTGAAAGGGTTGTGATGGAGTTAAAAGAGGCTGCAAATATGAAGAGCGAGGAGCGTGATGCTTACCTTGCAGAAATTGAGACGATTGGACAAGCTTATGAAGATATGCAAACACAGAACCAACGGCTTTTGCAGCAGATTTCTAAGCGAGATGATTATAACATAAAGCTTGTTTCTGAGAGTGTTAAATCAAAGCAACTTTACGCCACATTGAGCGAAGAGAAGCATGCCCTTCTAGAGTATATACAGCATGCGTCTAATTTTGGAGAAAGCCACAAACAAAGAGTGGCGAGGTTGGAACAGCAGGTTAAGTTGCATCTGGATCAGCTTAATAAGACTACAGAAGATGGAAGGCAAAATGCCAGCGCATTGGAGGTCTTGAAAAGAAGGCTAGCTGATGCTGATAAAGAGTGTCTTTCGACTAAAGTCTCTCTGGAAGCTGTTCAGAAAGAGATTGAACAACGCAAGGGTAAGACTGGAGATATACAGGCACAGCTGGAGAAGGAAAGGGTGGCAAAGAAAAAAGTTCAGGAAGATCTAGCATTATTAAAGTCAAAATTGGCACGCACTACTCCACAAAATGACGGGAGCTCCCTGATAGAGAAGCTTCAGGATGAAGTCAAAGAGTATAAAGCAATTTTGAAGTGTAGTGTATGTCATGACCGCCCCAAAGAAGTGGTCATTACAAAGTGCTATCACTTATTTTGCAACCCATGCATTCAGAGAAATCTGGAAATCCGACATCGTAAATGTCCAGGTTGTGGCATCACTTTTGGACAGAGCGATGTGCGAGCAGTTTACATCTAAGTGGGGCATTTGATTGAAGCTTGGATGCATCAAATTTCATGGGTTTTGCCACCCATTAAACTCTAGAGTCCAAAGAGACTTTTGGGTAGGAGTTTACCGTGCAAGATATCCTCGAACATTTGTCATGGCATGGTAGTGGCTCCATTTGGCATGGCACGGTAGCGTCCCACGCAGCTTTGTCCTGCTTTGGAGATCAGCGGGACTTGTATATTAATGATGGATCAAGGTTTTGGCTCCACTTTTGGGGAAGTCTCCTTGTAGCTATTAGGTGCACATTGAGTGTAGAAGCAGGAAATTTTGCTGGTGCGATACAATTCTCTCAGGTGATTACTCAAAGATGCGGCTCCACGATCATTGCCCAGCATGCAGCCGCGTGCATGTTTCTGCCTGAGCTGCTTGCTGTACAAAATTTTTCTGAGTGTTGAAAGGCACTTTAGTACCTTTTACAGGTGTATCACTCCAGTGAGCCTTCTTCAAATCAATTTAAAGTTGATTCATTTACGAAAGCCATCCCTTTGGCCCATTTCCAGCATGCAGCTCCACACGGGTGTGTATGTGCGTGTGTGCTTGCTTTTGTGTTCTTGTGTGACAAGCTGTTGTACATAATTGTTGTGGAGTGTAACAATCGATATTGTCTAACAGGCCTTAAGAAAAACTGCTCACAGCTCTCTGATTCTTCAGGGTTTACCACTAGAAGAGGCAGTTTTTTGTGTAAAACTGATGTGGGTATCAATCTTATACTGAAAAGGGAAATGAATAGAATATTCTGGCAAAAAAA

**INTEGRIN-LINKED KINASE1 (ILK1)**:

GGACTCGGGAGCTTTCATTTTCTGTCTCTGGAACCTGCAACAGCCCACCCACCCTCCATTTTTGAGGCGGCCTATGGTGGTCTTCGTCTTCTTCTCCTGGCGCCTCCTCCCGCTGCATCCTCTCCTGAGCTCGTGAACTCCATCGCCCTTCTGCATTTCCTGGCTCGCATTCTCCATCGCAGGCTTGGATACTGCATTTTGAGAATTCCTGGTTTCAAGAAGGCCAGTAAATCAGCACAGTGTCCATCCTTCAGAAATAAGAACCTCCATGGATTCAGCTAAGCAGAGATTCACTCTGGGAAGGCAGTCTTCTTTGGCGCCTCCCAACAAAGATGGTGACACAGATTCATCAAGCACAGTTTCACTCCCAGAGACATTGGACTTGGCCATGCAGCTACTGTTTCTTGCCAGCAAAGGGGATGAGGATGGCATTAGGGCTCTACTTTCCCAAGGTGTTGATGTGAATATTGCTGACTTTGATGATCGCACGGCTTTGCATGTGGCTGCCTGCGAAGGCCATGCCAAGGTTGTCAATCTGTTGCTGAAAAAAGGAGCAAATGGTAATGCACGAGATCGATGGGGCAGCACTCCACTTTCAGATGCGAAACATTATGGCAACAAGGAGGTGTGCGAACTCTTGGAAAAGGCTGGTGCTAAACTTCCAGAAAGTTCTAAGAAGGCTCCGATGAAAGTTGCAGATCCAGCTGATGTGCCAGAATATGAAGTCCGTCCAGAAGCTTTAACACACGTGAGGAGGGACCTGAAGATGCCTGAGTATATGGTTGCTACCTGGCATGGAATAGATGTTTCTGTGAAGGAGATTCCAAGGGAGTTATCAACTCAGCCCCAAATTGTTGCTGTTTTCCGAGAGGAGTTGGCTTTGTTGCAAAAGCTGCGGCATCCAAACATCGTGCAATTTCTAGGGGCCGTAACTCAAAGCACTCCAATGCGTATCATTTTGGAGTACTGTGAAGGGGGGGATTTGCATAGCTACTTGCAGAGGAAGAAACGACTTGAGCTTGGCAAAGCAGTGAGATATGCTCTTGACATAGCAAGGGGTTTGAATTACCTACACGAGTGGAAGCCTGAATCCATTGTGCATCGTGAATTGAAACCAAGTAAATTGTCAGGAGCATTCTACGAGATAGAGTTGGACGTTTGAAGGTTGCTAACTTTCACTTGAATCAGATTCTGCATGGAGCCTCTGAAGCTCGTTCAGATGCCAGTTTATGTGGTCCCTCTCGTTACATTGCTCCAGAAGTATTGCGAGGAGTTGCCTTCGACAAAAGTGTGGATGTGTATTCATTTTCCATGATCTTGTACCAGATGATTGAGGGCTCTGCTCCATTTGCGGACAAGAGTGAAGATCAAGTGCTTCTTGTTGTTACTCAGAAGAAAAATGAGAGGCCACCTTTCAAACTCAAGGCTAAGCACTACCCAGAAGGATTAAAAGAGTTGATCCAGGCTTGTTGGCATGAGAATCCGTCACAAAGACCAACGTTTGGAGTTATCATTGATGCCTTGGAAACATTGAAGCCTTTGTGCGACAAGAAGAGCTTTTGGAAGCGCATTTTTAAATCACCTAAAAGATCTCTCAGTATAAGGTCTCTGAGCCTTCAGTAAGTTTTGCTGCTGGTTGGTGTATAATAAGGCTGATGGCACCTACAGGAAGTGGGAAGAATGGTTCTAGTTTGGGATTTACTGTAGAACACACAAAACGCAACAAAGATTTTGCGCATGGCTATACATTCAACCTGTCATCATCAAGGGTTGATTAGGACACTTGCTGCCTCATTTTTGGAAAAACCACACATTCCCTATACCGTCATTTTTAGCTTAGTCTGAGTCGAAAACTGAGCTCAATCAATGTAAAGGTTTTACTGCTTCAATACTTTCAAGGTCGACATTCAGGTTCTGAGATATTGCAAAGCTAGTCTAGTACATTTCTTGTCTTATTCTATGCATAACAGAAGCAGTGGTTTGTGCTAGGTGGATA

**FLAGELLIN-SENSITIVE 2 (FLS2):**

GAAAAAATGTCAAAAAATCAAATTGAACTATATTATGGGAGTGATACATTAATATACTTTGTGGCCGCGAAGCGGCCACTAAAAAGAGTGGACCAGGCTGACCGGTAGGCCGGCCCCGGCCTCGGCCACTCGTGAATTTATAACGAAAGGCTTGTTATGGTCGATGTTAAAGAGAAATGCACGCACTTCTTGCATTGTGGGGCGGTCATTGGGGTGGTCTTGTGTACATGCCACACCCATTCTTAGGAGCTGCACTGCCTGCATGCGGGCATTATCAACATTAATGACACTCGATGATGATGATGATGTCTCACAATCTGAGATGGAGGCGTTGAGCAACTCCTCATAATTTAAATCTACCCCATAGTTGGGGAGCTTGGCTTTGACCCATTCATGGAGGCTTGTTGATCCAGGCAGTTGTGTAGGCTTTCTCCCCGTCACCATCTCTAGGACCACCACTCCAAAGCTGTAGACATCCACCTTAGTTGTGATTCGTGACTCATATGCATACTCTAGCAGCCAAAACCAGCATTAAACGACTAAAATTAAGAATGTGATGTCTATATATCATTACTAATTAGAATGTATTTACATATTTCTTCAGCAAACTCAACGAAATCATTGAGTTTCGAGGGGTCAGCTCCCCTCAGCTTTATCTCCCACGGGATGGCATGCATGGGATCAAGGGACCTTATGAAAAGGCGGCCTCCACGCCTTATTGGGGCCGTTGATCTCAAAATTAATCTCGGGATAAAAAATTAATCTTAGGATCCTGAGATTAAATTCCATTCCGAGATTAATTATAGGGGAAAACAATGTCCGGACTCCAAAACTTGTGGTGATATTATCGCATTTGAGCTATCAAAAGACAGACTTTAGTACCTACCTGGTGCCATGTACCCAATTGATCCCTTGAAGTTGGAAGTTAAAGAGACAGCTCCCTCTTTAACAAGCTTTGCGAGCCCAAAATCTGCAAGATGTGCCTCGAGAGCAGAGTCTAGAAGAATGTTGTTGGGCTTGACATCCCCATGAATAATGGGTACACCATACTCGTGATGCAAGTAGGTCAAAGCATTGGCAACACCCAGTGCTATATTTGTGAGACTCATGAACCCATAAGATTGATGCCCATCATGATGATGCAGATATTGCTCCAGGTTCCCATTGGGTACATATTCTAGCACCAAGGCATTCACCTCTAGGTTGAGGACACACCCCAAGATTTGAACAAGATTTCTGTGCCTTAGTTTGCCAAGTATCTGCACCTCTTGTAGCAATGCTTCACGGAAACAGACCTCCTTGAGCTTTTTCACAGCAACAACCTTGCCATTTGACAACACTCCCTTGTACACAGAGGCGAGCGCACCTTGTCCGACAAGGTTGGCCTCCGAATACCCTTGGGTGGCATTTATAAGCTCCTCCGCTGTGTATCGTTTTGTTTCTGCCAACAACTTGATCTCTTTGGGCACCACTTGAAGACATGATTGACTCCATTTTGAGCAATAACACAATGCCAATAGTATCAAAGACCCGGCTATGGCTGACGAGGTTAAGACAATACCGAGAGTTGCTTTCCAGCTGATCTGACCACCATGGTGATGTTTCTTCGAGGAATATTCATGAATGCATATCCTGTGAATGACAGGGCCGCAAAGCCCAGGATTTCCTAGAAAGGAGCTTGCATTAAGGTGCTTGAAGACCCCAATAAGAGGGATGGTGCCAGTGAAGTTGTTGAAGGAGACATTGAGAAATTGTAGGCGATGTAATTGACTCCCCAGGACTGATGCAGGTAGCACCCCTGTAAGGTTATTGCTAGAAACATCGAGCTTTTCGAGGTTGACTAGCTTGCCTAAGTGCTCTGGGAGCCCTCCAGAGAGGTGGTTGTGAGACAAGATGAGAGCAGTAAGCCCTACACAATCACCAAGGCTGTCTGGTATGGTGCCAACAAAGTGATTGTGGGACAAATTCAAACTCTCCACCTGTACTAGCTGCTTTCCTAACCAAGAGGGTAAAGCCCCAAAGAATTCATTGTGGGACAAGTCCAGATCTTTTACCGTGCGCAGCGCTTGAGCAAGACTTGATGGCAAGGAGCCATTTAGTCGATTGAAAGACAAGTCTACTTCAAGCAAGCTAAGGGATTCAGAAGACTCAAGCTCCGGAATACTGCCCTGAAGCAGGTTGCCGTGCAGATATAGCACTTGCAACATTTGGAGCATTGTAAGACTGCTTGGGATTCCACCAGTTAGGCGATTATTGACTAAGCTAAGATAATATAGAGAGCTACAGTTGTACAAGGAACCCCATGGTATCTCATTTGAAAAGTTATTGTCTCCAAGGTCTAGAGCTTGAAGCTGTTGCACTCGCCCCAGCTCTATTGGTATAGAGCCAGTTAGGTGATTATGGTGCAAGACTAATTCAAATAGACTAGTGAAGTTGCTGAAGTGGAGCAGCCTGAAGCTGGACAGATGATTTTCACTTAAAAGAAGGTATTTTAAATTTCCCAGCATTGAGAGCTGGGTGGGAATGACTCCTGTAAGTTGATTAATTCCAAGGTTAAGGTTAAAGAGATGTGTGCAGTTTCCTAGGCTGGATGGGATGGTTCCCCTCAGGTTGTTTCCCCATGCAATTATCCTCTTGAGGCCAGATAGCGAGCCAAGCTCTGTGGGGATAGGACCGCTAAACTGGTTGGATTCCACATCCAGAATGATGAGGTTTGTGCAGTTGCCCAAACCCCCAATAATGGCTCCTGTAAGGATGTTCTCATTCAAGCTTAGCCTTTCAAGCTTACGCAGTGAACCGAGCTCCATGGGCATGGCACCAGTGAGCTGATTGTAGCTGAGCTCGAGGTCGGTGAGATGTGCACAATCTCCCAGACTTCGTGGAATGGATCCGCTCAGGTTGTTTTGCCACAACCATAAGCCGGATAGATTGCTTAAAGATCCCAGCTCTGATGGGATGCTGCCACTGAGCTGGTTGGAGCTGAGATCCAGCTGTAAGAGATGAGTGCAGTTTCCTATGCTCCGAGGTATGCTAAATCTGAGGTTGTTTTCCCGCAAAAGTAGCGTTAGGAGATGGGTTAGCAATCCTAGCTCCCAGGGGATAGAACCAGGCGTCAGCGGGTTAGAGTAGATCTGAAGATCAACGAGGTTTGTGCAGTTGCCGATGCTCGGAGGGATGACTCCTGTGAGGTTGTTAGTGTACAGGAGTAGTCGCTGAAGATGAATGAGCAAGCCAACCTCGGAGGGGATGGGACCGCTAAGCCAATTGGACGAGAGCTCAAGGCCGATGAGGCCAGTGCAGTTGCTGAGCAAGGTGGAAGGGATGGCTCCACTGAGGTTGTTACCCCACAGAAGTAGCTGCTGAAGAGGACTAGCGATGCTCGCTGGGGCTGGGATGGAGCCTGTGAACAAATTGTTGCATAGACCCAGACTTCCGAGAGAGGTGCAATTTGTGAGTAGGGTGTGAGGGATGTGACCGCGGAGGAGGTTGTTGCAGAGATCCAGAGCTATGAGAGAAGAGAGCGAGCTGAGGGATGAAGGGATGAAGCCGGAGAGGCTGTTGGTGTTGAGATCCAGCACTTTGAGAGAAGCACAGGCAGAGAGCTGCGGGGGTATGGAGCCGGTGAAAGAGTTGGAGACGAGAAAGAGGGCGCGCATGGAGGTGAGGTTGCCGAGGTGGGGCGCGAGAGAGCCTGCCAGCCCAAAGCCTGTGAGGTTCAATATATCGACGTGAAGCTGGCCAGCAAGTGAGCTGCTGCTGCTGCATGTGATCCCCACCCAGGATTCACAAACGCTCCGTGTGTTGTTTGGATCCCAGGTGCTGAGCAGCGCCCGTGATTGTGTGGAGAGGTGAGCCTTGAACTGCAACAGAGCATCCACTTCCTCTGATGGAATATTTAGCGGCACTGAAGTGGAGTACGCCGGCTGCATTACCAAGCTTAGCTGGCAGATGAGAATCAGAATGAGAGCAGCTCCCTGCTCCATGGCTGATGTTCTTGCGCAG

**HOPW1-1-INTERACTING 1 (WIN1):**

TTAGAACCTCCTCTTCTTCTAGCTGCCAGAATGATGCGTTGCAAAGTCAATTGATTGTTTGCATTGGCGAAACAAGTTTAAAACATGCGATAAATCTAGTTCTGATTGCTAGGTAACGAACACCCGGGCAGGCAGTCACCTGCAAGACTGAAGAACTAGAAGCATGATAACTAGTTCCTCACATGTATAAGAGATGCTAATGTTTATTTTGTTGGCTTCATGCAAGTAGGTGCCCTCGTAATAACATAAGCACTTGAGGTTTATACTAGAAATCTTGTTAATTACCTAAACATGCATACACAAGAAGTTTTGGGCATGCTACACCGACCCCCCTCCCCACAACCGCACCTAGCTAGGTATGCTACACCGGCTCGCCTGCTAAAGCTCAAGATGCTGTTGTAATCCGTAAAGGGGGAATTACAGTGGGAAAATGTTTAAAACTTCTTGTGGCACAACCAAACTTCCTTGGTTCTGTCAACTTCTGTGGAAGAGACGTGCCATTCTTTTTAAACTCTGGATGCCGCAAAAGTGAGTAGTGGGCGCGAGTCTGCACGTTGTTAAAATTTAGAAATTAGGGTTTAAGGCTTCAACTGTGCATCTTTTACATGGCTGCCCTGCAAACTTCATTGTGCCATGTCTCAAGCCGCCCATCAACAGAGAGCTTTTCCTACAGCCCCATTAAGCCTCCATGGCTTCCCAAGATTCCATCCATTTCATCTGCTAAGGCCATGGCTTCTCAGCACCCACATGCCATTGCAGACATTGACGCTCGCACAGAAGAAGGAGAACAACAACAGCATGCAGCTGAATTGGCGGCTGCCGTCTTATCTGTGCAACCCACAATGCATGCCTCCACCCAGGATATCATCAATGCTGAGTCGCGCTTGTTTGTCAAGACTTATGCTCGGTGTCCCGTTGTGTTCGTGAAAGGTCACGGATGCAAGCTTTATGATGCAGAGGGTAAAGAGTATCTAGACATGACGGCGGGCATCGCTGTGAATGCACTGGGCCATGGTGATGAAAGCTGGGTCAAAGCTGTTGTAGAACAAGCATCAGTTTTGGCCCATGTCAGTAACCTTTATCATTCAGAACCACAGGTGAAACTAGCCAAACGTTTGGTGGAATCATCATTTGCTGACCGCGTGTTTTTTTGTAATTCTGGTACGGAGGCAAATGAAGCTGCTTTGAAGTTTGCTCGAAAGTACCAACGATACAAGGCAAAGGAGAAGGGTGAGTCACAGTGGTTCAAACTTGGACAAGATCATCCAGCCACAGAGTTTGTAGCTTTTTCCAATTGTTTTCATGGGCGGACAATGGGGGCACTTGCTCTCACCAGCAAAGAGAAATATCGTGAGCCATTTGAGCCCATCATGCCAGGGGTGCAATTTGTAGAATTTGGAAACTTACAGGCAGCACAAGTAGCCATTGAACGAGGGAAAACAGCAGCAGTCTTTGTGGAACCAGTGCAAGGAGAGGGCGGAATCTATCCAGTGATGGTTTCTTTTATCAGAGGCCTTCGAAAATTTTGCGATGAAAGTGGAGCTCTTCTTGTTTTTGATGAGGTGCAATGTGGCCTTGGTCGCACAGGTCGATTATGGGCACATGAAGCTTATGGAGTTCAACCGGACATATTGACTGTAGCTAAGCCACTTGCAGGTGGCCTTCCGATTGGTGCGGTTCTAGTAACAGAGGCAGTAGCCTCTACTATTGAGTTTGGTGATCATGGAAGCACATTTGCTGGTGGGCCTCTGGTATGCAATGCGGCACTCGCAGTTTTGAACAGGATACAAGAGCCAGGATTTTTGGAAAGCGTTGCAGCAAAAGGAGACCTATTGCGAAGTTTATTAAGTGAGAAGCTTGGCAGTAATGAGCATGTCAAAGAGGTTCGTGGAATGGGTCTTCTTGTTGGCGTTGAGCTTGATGTGTCTGCTTCACCTTTGGTGAAAGCGGCGCTTAATAAGGGGCTCCTAGTCTTAACAGCAGGTAAGGGAAACATTGTCAGGTTTGTGCCACCTTTGATTATATCAGAGCAAGAGATTAGGCAAGGGGTGGAGATTTTGTTGCAGTGTATGCCTGCCCTTGAGGAGACATCTTAGAAGATTTAATAAACTTGGCCTGATTGTGATGGTTAACGAGGTTCAAAATATATATAACATGCAGCGGTGCCGAGCTTTTTGGAAAGAGTAAATTATGAATGATGCACTAGGCAACTCACATTTCGTAGAGTTGTCTTGCATGAATTTTCGATTATGCCCGTTGCTTTTTAAAATTTACGCTTCACAAATAAGTAATTACCAGAAAATTTGACAGAAAAGTGTGCATGCACATGTTTTATTATAGCTTCAAAAGCTAGGTTGCAGGTGGTGCACACTAGGTGTTCAAGCACTCCCCTCAAATATGCATCCAATAGAAACCCTCTTTCTTTATCAATTATCTTTCTTTTGCTATATTTCTTCTTTCAAAGCACTGCAATGGGACCTCCTTGCAAGCCATTCCCCCTACCAATACATCTGGGATACAACTTTTTTATTGACCTTTG

**HOPW1-1-INTERACTING 2 (WIN2):**

GAAGAAATCACATTTTGAACTCCACTGATATTGATTGCACTCCCTCAGGCCACATGCAAAGCAAACTTAAAGTCTTGTGCACTCATACAAACACTCTTTTGGTCTCTTTGAGGGAGCATCGAAAGGTAACCCCCCCTTTGTCCATGTGTGCGATATCTGTCTCTTTCTCCACCTTTTTCTCCTCGTGGTAATAGATTTCCAAAAAAAAAGCACATTTTTGTCTATCAATGAATCTTCCAAGAAGTGTAACAATTGCATCTATAAATTGCACGGTGGCAAGTCCTACATCCAGATGATAGAGCACGGGAATAGCAATAAGCAATCCCAGCTAGAGAACACAAAAGTGATGTAAGCTCAAATAATGATATGATCTTTCCTCAGCCTTCCCTTATTACTACTCGAACTGCTTTTGAAAAGGGGAAGGCCTGTCTATGCAGGTTTAGTGTTATGATATAAAATGAGAGCACAAATGCCGTGGGGCTCCGCTCTTTAGACGCTTCCAGCAAAGCATAATAGGAACCTCCCTTTTGAAAAGGCTCCATCCCTGTTCAAAGCTTAGAAATAGACTCCATCCCTGTTAAAAACTAAAAAATAGGCAGCATCCATGTTAAAAGCTAACATTACTCCAATTTTTAGGCTGTGTCAATATTAAACCGCACTACAATGCATGTGATGTTGTCTGCGCTTCCTTTCCGTAAAGCCTCTTCAGTAAGCTTTCTTGCTGCTTCTTCTGCATCAGGAATAGACTGAACCATTGTAACTGCATCCTGATTGGTCACAACATCCCATAAACCATCACTAGCAAGAACCAGAAATTCAACCTCATCATTGATGATCTCCTCCTACAAACCATTCGACATGAGCTAAAAACAGGCAGACCTTCCACAAAAAGTTAGTCAGGTAGTACAAAAATCCTTACTTGTCACCACAATTAACACGCGAACTAACAAAATTTGTTACTAGAACAATCCCCTACTTTAATTTTCACAATGTCATATTTGTAATCTCTATATGCCTAGGTTTTCAAACTTAAAAGGCCAGAAGATGCTCACTATCATAAAAGAAAATATCACACCTACTGTTTCAGCAACAAAGTACCAATATGACATAGCCCTGAGAATTATCTTTCCCAGGGCAAAGAGACATCTGCCTTACCTGAAGTTCAGGTTCCGCTACCACATACTGTTTTAGCAACCGATCACCAAAAGCTCGAGAAACAGCAAGCACCCCTCCCACTCTCCAAGTGCCAGCCCACATCACAACTCCACCAGCTTTTTCAATCCGCTGACGTTCATCCTTGCGATTGGGTTTATGATCAGTTGATAATGCAAAAGCTTTCCCAGCCCTGCAAATAACTGCTCTCGAATCACCCACGTTTGCAACTAGTAGACGGTCGCCCACCAGCACTGCAGTTGACGCAGTCGAGCCAGCATCTCTGTGTTGGCTGTTCTCGGCTTTTAAATACTCTGAATCCGTCTGTTGGTATGTTTCAACTATTGCACATCTTGTATCAGTTAAGAATTTGGGATGCTTGATCAGGTTCTCAAAAAGGTGTTTTTGCACATAATCTGCAGCACGAGAGCCACCATGACCATCGAAAACGCCAAAAAGGCCAACCGTAGAATCTCCAACTTGAGCAAATCGAGCCTCGTAAAAGTCTTCCATAGAAGCTCTTTTTCCACGCAAGCTTGAATATCCAAAGCTACATTTTCCATCATGACTTTGCCCCCCACCGCTTGCTTCAATATTTTGGCACTCACTAGTCATTGTTTTGATGAGAGCGTAGTTATTCCAACAGCATTTCTGGCAAGAGGCGTTGGACCTTAAACTGGCGATTGAATCCACACCTGATAAGCCCGAGTCTTTCAAGCTGAAGGCTTACATTTGATCTGGAAGAGGAAGAGGAGAGGTTGACCACAGAAGAGGGTAGGAAACCCTAATCGAAAACCTAGCTTTAATGGAAACTTGAGCCGACAACAAATTCGCGCCAAGCGTTCCCTGTGACTGCGAAGATGCTCTCACAACTCTGCTCAAACACTAAAATTGAGATGATCCCCAGCCGTTCGATCCCTACCCCATCCAACGGCTCTTACGATGCCGAGCTCTTTCGTAATCTTG

**LYSM-CONTAINING RECEPTOR-LIKE KINASE 1 (LYK1):**

CAGACATGCAAAAGTGGTACACAATCATGCTGATCAAAGCAATCCCACATGAAAGCTAAGCAAACATGCAAAAGTGGTACACCTTTCCTATCACCATAATACGAATGGAGCCTGTATGTGAGCTGAGTAGCATAATTACACAAGAAAACGAATTGAAAACAGCATTGTAGTGTATCAGAAAGCATTAATGAGAAGCAGCTAGAAGAGGAGCTTAATTTTTCCCTAAGCTTGCAACTCTCTTTATTGTTTAAGCTTGCAATCCTCTTCATTGTTTATGACACTTTCCAAGTCTCCCTGGGTATGTAACTACTTGCGACCTCTTCACAAGGAAGCGCGTGTTACTTGACCTATCATTGTACAGAGGCTCAAGCTTTACTTATTTGTGTGAGCACTTGAAAACACCAGGGGTGCCTGAGGGTGGCTTTAAAGTTTGATGAAATTTTAAGGTAGAGCAGGATGATTAGCTCAACCCAGGCACGCACCTCAGCCACGCTCTTGATTTGTCAAAGAGACGCCCGAACCAACAGCATCCACAGAAAAATTGCATTGCACATCCACTGGCGTCGAGAAGTAATCCAAAAACCAATGTGTGCAGGAGACTTTGAAAGGTGTGGTCACGAGAAATTATAAGTAGGCTTTTCCTCCTTTTTTGTTGGGCAAAATTCGATCGATGAATTCAAGCATATACAAGAAATATCTAGAGAAATACAAGTGAGCCATCAACGATAATAGATGGATGGGTAAATAGTAAATCTTCCTCGTTTCATTTCATTTCTGTCTACATCTGAAATATTCTGTACCTATATGCTGCCTTAGCTAAGCTCTTCTCGACAGGAGATTTCTCCAATATTTGAAAACTTTATATGTTAGTATTACTGATACAAATTTGGCATTAGAACAAGCTGCCGTGTTCCAGGCATAGATTTCCATGACGGTGTCCAAGGCCTGGAAGCTGCCAGTGTGGAATTTGTGGTTAGAAACCCAAAATCTCTTCTTCAAAACTCAGCTACCACATGGGGCACACACGAAACAGTCCGAACCAGCCCTTCATCATTTGAGCTTTCTCAAACTCCTGCTATTATTTTGAGGCTGCGAAAACAGCCTACAAAAAGAAGCCATAGCCTTATTCGTTTACATAGCAGCCCCCTCATGTTTCCACTTTACAAATGGCGGGCCGTCAATTTCACCTTGCCATCAAGACCCCCTCCATGCCATCATGATCTGCATCCCATTCCTGTGTTGAAGCGGATAGTGTCATGAGGGCCACCACCACTACCCTCATGTTGGGTCGCAAAAGAGGCTCTTCTCTGGTGCAAGCGCCTGCCAAAGTTGCCATCTTCCACACAGACTCCAAAGGGTAGTCATCTCCAAGTGCTGGATCTACTAGACTTCGAAGTTTCTCTTTTCCTTTGACTGAATCACTCAAGACCCCCTCGAACAATGTAACAAGCCCATGAGACATTGCTCTTGGACCCTGGGTCGTTGATACTTGGTCAACGTGATGTACGATTGCATCTTTGGCAGAAATGATCTCAAACAAAACAACACCAAAGGAATAAACGTCGGATTTTGCTGAAATATCTCCAAAGCGTGCATACTCTGGAGCCATGTATCCAAATGTACCCACTAATCTTGATGGGATAGTACTTGAACCAGTTCCAGATTCCGTTAGTTTAGTAAGCCCAAAGTCAGCAACCTTAGCACGGCAACTGTTGTCCAGTAGAATGTTGCTTGATTTAATATCACGGTGGATATAAGTAGGCGTTGTATGTTCATGTATATATTCCAAACCTCTAGCAGAATCAAGGGCTACCTGAATTCTAGCTTCCCAAGACAAAGGTGTGGTCCCTTCTCTTCCTGAGTATTTCAAGTGCTGGCTTAGGGTTCCATTTTCCACAAACTCGTAAACAAGAAAGAGAAATTCATGTGTACAAAAGCCAACAAGGCGGACCAGATTTGTATGATGAACGCGTGTCAACACGTGCAATTCAGCCAAAAACTCTTTGGTGGCCTGCATATTCATCATTTTGATTGCTAGCTTCTGCCCATGTAACTCTCCATAAAAAACAGAGCCATAGCCTCCTTGCCCTATCTTGTTGGCAGTACTAAAGTTGTCAGTGGCATCAGAAAGCTCCTTGTAAGAAAATTCGACAGACCTGTCTGGGCTGTAATATGAAAGCCCCGAGGACACTGTAGTATTTGAATATGCACTCCCTTCCGAAAAAATCCTTGGTGCTCCGGGTGACACAGTAGCAGTCTGAGGTGTTGCAGGATACTTAACCATCTCAGTGGCTTCTCTGTGTTTCTTGGGGCGATAGATACACACATAGAACAGAAAAGCTCCTGTCATTGCTAGGAAGAACACAACTGCACCGACCGTTGCTCCTACAATGGCTTTTGTATTACGGTGGCTCCCAAGTCCCGTGCTGTTGAAAGGTGGGTACACACCATTTCCATCTTGTGCTGGCACAAAAAGTATGCTGACATATTGCTGAAGAAGATTGAGGTTGAGGTTTTGATTATATCTCTGCAACAAATCAACAGAGATATTGAACTTCTTTGAAAGCGCTGCAGCATTGTCATTCTCCACGGCAACATAGGTAAGAAAGAGTCCGTATGATGAGTTGATATTAGGGTCACCACAGGAGCAATTAACAGGGATAGTGAGTACGTTCTGGGGATAGATAGCGCTCAAATCCTTGTCGTTGTTGACGCGCAACATTGCATTGTAGGTGGTGAGGAACCCATACTTGTCATTAGCTATTTCTATGGCCGTGTCACCGGTAACCACTGTGTATGAAAAGTTGTGCCAAAGGAAGTTGTCTGCGCATCCACAAGCAAAGGGGATGCGAAGCGGTTGGTCCGAGAGTATTGCAGACGATGCATTGAGCTGGCTGTACCAACGAATGTCCCTCTCATTTGCCCCAAATCTCTTCGCTACATTGGACAAGTTTCCATCAGCAGCTGAAACAAAGGTGTAGTATGCTAATGCGACACTACATCCACTTGGGGGCATGCAATCAGCATGTGCAAATCTAAACATTCCTGGTAACAAAACCACTAACAAGAGGCAAACAAGGTTCATCCTTCCACCCTTAAGCATCTCAATAGTCCTTACTAAGGGCACCAGCTTATGATTTAGCCCTTTCGCGCTCACATTAAATCCGTTTGTAGAGGGTCAATTTCTCCCTGCCTGAAAATTATTCAGAGGCCCTGCTTGTTTGGCTGCCAATGTGCAGCTACATCATAGCCATGACCTCCATCTAAGAATGCGAGAAATCTACTGCTGAATCACAAGATTTCAAGGTGCCCATATTTGATCAATGCGCTGCAACTTGCTAGATTGAGCAGTAATTCGAATAAAGTGATGGGTATCTGCAAAGCCTTCTTGTAGAAGCTCGTCTGTATCAAGAAGAAATGGAAAACCCACAAGAACCTGCTCAACTTCGCAGAATGAAGCACCCAAATTAGAGCTTCTGAGGCCTCGATTAAACAAGGAAGAGCTTGACTATGGCGATGAAGAAGATGAAGATGATGGGGAGACGCGATTGCTGAAAGCGCCGGCGTGACGACGAAGACCCTGAAGAAAGAAGAGGCCAATTCGAAATAAAAGTCAATATCTGTAGGCAAG

**RNA-DEPENDENT RNA POLYMERASE 1 (RDR1)**:

TTTTGAATTGCAACATCAATAAATGTGATTAGAACAATAATGTCTCTATATTTTTATAATTACATTTTCCTGAATAAGGTGTTTGAGATGATCAAATAATTACATAGGGATAGTATATATGTAGTTTTGATCGCACGGGCTTCATTTGATGTATGATAAATACAAGGTACATTTCCAATGTTTCTCTTAAGCTTTAAGTTAGTAAAGGAGCAAGATGACAGCAGGAAGCCAGAGGCCATGGGCTGGGGATTTTCCTCGCCAGTCCCCGTCCCCATCGCGCCTAACTAGTTATGATGTGCCCCTTCTACAGAAGCTCATGATGTCGAAGTTTTCTCACCCTTGCAGCCAAGCCTCTAGGCTATGCTTACACTGGATAAGCATGAGCTCAATTTGCATGCTTGCACTGTGCCTTGGGTTTTCTAATATTCCGAAAAAGGGGTTCTTGTGCATGTTCACAAACATGGTCTTCCCTACTATGCCAAGAAAGGGCTCGCTGACACGACACGGGCCTTAACTGTGCCAACTAAGCCCATTCAGATGAGCATAGCCCTATTTCACTATTCAGCAACTCCCATGTTGCTAGCAATTCGGCAATTCAGCACTTGCCTGCAAGTGGCCCTTTCAAAAACCAATCTAGTTGCTCTTGCTTTACCATCTTATGTCCGGATCGTACATTACAATGCCACAAAGTGGCTTTTCATATTATACAAAATATCATCGAACGGTGTGATGTTTTTGATTCCACCCTTTAGCAGATTGTCATCTGCAATCTGTATCCCTTTTTGATGGCTGAGAGCCTGTCATGTACCACCCAAGGGAAGCTGAGAAGATGCTCGTGTTCAAATCCATCATAAACATTGGACTCAATAACATAATCTGGATGATAGGTCACATGATACCAAGCTGAGGCTTTTGCCAACTCTTCATGAGTATACTCTGGGGAAACATCATCATGATTATCATGCATACTCACTCGGTGACTCCTTAACCAGCTTCTCGCCTCCTTCTTTAAAGAATTCATGCTTTTTTGGATCCTTTCTTTTACTTCTCCGGGTCTTTTCTGATAGAACCTAGACAAGCTACGGATGTTTCCGGTAAAGATTTCCGCCTCGCTTTTCACATCGTATTGATACATCAAAGTTAGAAGCTGTTCATCATAATAGCCTTTCTGTGCATGTGCTTCTTCCACATACTGCTCGAATCCACATATTTCTAGTTCTCTGTCATAGAATTTTGCTGCTTCATGCTTAGTAAAAGGTACTATCATTTTGGCATCACAGTCTTCTGCCACTGCCCTATAAAGCTTTCCAAGCACCCTCTCTGATGTATACGTGGTTTTATCATCCTTTTCCATAAAATCAGGGTATGTCCTTGGCCGAAGCTCAACAGGTAATTTTGCAGGAACTCCAGTCTTTGGAAAGTCAACAGCTATAGAGCATAACTTTGCAAGGGCCAAACACTTCATATCATAGGCCTGTGCTTCTTCAGTATCGGCAAAAACTGTGTGCGCATTTGATATGAGACCTAGGCTGTCATTTGTCATGAAATCCACAAAGTATTCTTGGATTTCCTGGATTGTCACTGGATGATCTTGCACTTGTGTTGGGGATGCCTGGTAATCCATTGGTGGAGCTGCTACTGGGGGGATCAGCTCCTTGTCCCAGCTCACAAAATACAAATCTCCATCAAGATCACTGCCTGAACATTCATTTGGATGAGGCCTGTGACCCCGTTGTGGGAATACAATGCAGTCAACAAGATGATGCAAAGCTGGAACATTGACGGCAGCCAGCACTCGCACATCTCCAGGATGAAGACAGGGATTCTTTGCCACAATTACCTTGCCTTCTACAATGCACACATTTCGAGTTGCCCCATTCATCCCATGAACAGTTATGGGACGAAGTCCATCATCAAAATACTTGTAGGGGCATCCAGCCCTGGGTGAAACACTCACAAATACTTCCCCATAATTCAAGACTGCTGCCTCATCCAAGCAACCCATTAAAAGCCGACCTTTCGCAACAAATATGCGTGTCTTTCTACGAAGCTGCATTAGTTGTACCACTTTAAAAGCATACAACATCCTCTTTAGGAAGGGCTCCGAACTAGGTTCATAACCCGCAGAAATCATGTCCAGTATAACCCTTTGAGTATCACCTCGGCAAAAGGTCTGCATAATATCCATAGCCACCTCACCATCCTCTAGCATTTCATCTAACCGTTGCATTACCTTCAGCTGCATTCTTTCAAAAACCAAGTCATTGACGCCTAGCGTGGACAAAAGAGTGATGATTTGGCGATTAAGGTAGCTTGGTATAAAGTGGCTCCAGTTAAGAATATCAAGACCAATTTGTTTAGAAGGAAACTTTTCCATACTATCCCGCAATGACAGTTTGTAAGTGCTACTGGGATCAACAGCTACGACTCCTTTGTAACCACTATATCGCACCTGATATAATGATGGCATTTCTCTGTTGAGTCTTTTTAGCTTGCACTTTTTGGCCAACTGTGTAGCAAAATCTTCTGAGATTTTTCCTATGCCATCAGAGAAGCAGTATGTACATGGGTCATCCCAAATATCAGGGATAGTTTCAACTTCATTTGGGTACACATCCACTGCTTTTGTTGAAGAACTGAAAGATTGCCCCATTCTGGCTGCACATTTTGCCACATTTCTTATACCTGAAAAATCCCCCATCCATTCTCGAATGTTGGTTGCTGATGTTTTGCCATCAGATGCAAACATCCATATGCTATTTTCCCGTAGTTGACTGCTAGAAAATGCCAGAAATTCAAATCTTTTGCTGCCTATACTGATACCATCCCTGAGAATAGCAATGATGCGCTCATACACCTTTGTGTGCTGCTTCTGCTCAGAGAATGGCCATTCCCATGACTTTGATTTTTTTAAAATCATAAGCACACTGGAAGAGAGACTGTCATAGTTCTCTTCAACAAAAGAAACTCTGAGAAAGTTCTCAATGCTGTCAATAAATTGCCGGGTTACTCGATTTGATACAACTGCTTCTGGTCCCAAAAAATAGACTCTTGAGGGAGTCACATAAACCCTTCGAACCTCTACGAGGGCATCATTTGACCAGCTCTTCTTTGTTCTTACCTGATACTTTTTCCTTAAAAGTTGATGGGTTTGTTTCTTGAACCATACCTCTGGATTAAAGCAGGTGTCTTCTCTCAGATTTAACTTACTCAACACATGTATGACATGCTTTAGTGGCAACTTGTCAGAAGGTTGGATTAGCTCAAAGAATGTCTGATTGAGGTTAGGCCCACTTAAAAATCCTTGCTGCACAAGCGAATTCACCAGGAAAATCAGTTGAAAAGGAAGACTACAAGGTAAGTTGAGTATGGGAACATTGATATTGGAAACACTGTAGGTAGCCCCCTGTTTTATTGACAAATATTTCTCCTTCGACGAAAGATCTCTAAGCTCTGAAAGCTTTCGTAGCAAAGTCTTCAACTCCTTCCCACGTTTTTCAAACATTTCTACACAATAACTGAAAGATTGGCCAATGCTAGCATCAAGAGAGAAGTCAGTTGTTCGAGCCCATGCAATCTTGGCACCCTTGCCAAAGAGCTCAAATGCATCCTCAAAAAATATATCAGATGCACATGGAAGCTGTTCGTATATTTGTGGTGCAAGCCATACCTTAAGAAGTAGGCCTTGGCTGTTTTTCTTGTCTTTGATCTTGCATGGATGGATATCGAAGATGTCACGGAGGCGCCATTCAAGCTTATAATTATTGCCAGACAAATGCAACATGATTGCAAGCTTTTTAGTGCTATCATCCACTGTGAGTTTCACATTCTCAGCTGTCCACAATTCATGGAGCACATCATGGCTGACAAGACAGCCCGCATGAAGAGTTGAGTTAGCAGCAAGGGTTGCATTGTCCTGTCTTATATGAATTGGGTTTTCCCTTTTCCTGTGAGCGGGCTTGATGGCAAGCTTGGTGCCCATGAAGCCTAACCGCTTGAAGCTTGCAAGGTCACATGCCCTCCTTGCTGCTGTCTCTGTTGTAAACTCCACTACGGCTGAACCCCTCAGTTTCCCGGCTCTTGTGATGGACAGGAAACGACAATCTGTAACTGTCCCTTCCCCAACAGCATTCTCCAGAAAGGCAGTGAGTTGCTCAACTCTGTCGATCTCTGGAACACGATGCACATGTACAGTGCATCCATGCTTGATTTCGCTAGAAAGGATGTTACTGTTTTCAGCATGCGAGAGCTTGAAGAGCTCCTGAAGAAGAGAGTAGCCCTGCCCTCCTTCATGTTCCAACGTGTTCGCACCTTGAAGTAGCTCTGTAAGCTCTGCATAACCTGTGGGCTCCATGCGCAAGCACGCTATTTTACTGTCTATGTATGAAAAGCATCATCAAGAATGAGAGCACTGACTGTACACTAGTCAAGGCGCGAGTCAAGCAGCCTGCCCACTCCAT

**SUPPRESSOR OF GENE SILENCING 3 (SGS3)**:

GAAAATATAAACAGTACCAAGCCTATCAAGTGTCATGTACCAATGGTACCATACTAGAACTGAAGACAGGTATGCTTTACATGTTGGCCCCAGACAATATGAGGTGCTCATTTGGTAATTGATGCAAACAATTAAGTGAGCCCAGAACATTAAAGAAGGCCTAAAGCTTCAAACAAGACCCAAGAGCAGCACTTTAGCGAACTAGTATGACTGAAGTTTTCAAAATTGTGTTATCCCAAAGCTCCCTTTTGGCAGGACACACACGTCATACAAGATCAATTCCCACAGATTGATGTACTCAATAAAGACCTACAAAAGGGTCTTACCTAACATTCCACCAAACATGCAGGAAAAATAAAAAAATAAAATAAGGCGATTGGGCACATGTAGGGGATCCTTATTGCTTCCATGGCTTAGATGTCTGATGAGAGTCTAAAACTTTAAAATCCACCTATTCTCTAAATAATCGTGGATATTCAAGCTTGACAAGCATGCTCACTATACTTGTCTAGCAACTGTTGCCTCTCTTCCTCAAGTCCTTTCTCAAATTCAAGAGCTTCTTCATACTGCTTCCTCATAAACTCCCTCTTTCTAATGTGTTGCTGCTTCAAAAGTTCTCGCTTACTGCACTCATAATCCTCGGACTCCCGCAAACTCGATTCAAGAATCTGAGTCTGGTGTGCAATCGCTTCTTCCACTTTGGCCTGCTTGGGCTGTTGTTCTTCACCGAGCTTCTTCTCAAGCTGTTGACAGCGCTCTATGTGCTCTTGCTTGAACTCTTCTTGTTTTACCTGAATCTCTTTTTCTCTTCTTGCACTGTTTTTCTGGAGCTGAATTATTTGGCTCTTGTATGAGTTTTCAAGTTCATCCATTTCTCTTTGATTCTCCTCGTGCTGTTCTTTAGCTCGCTGTCGTACAACAGAAATTTCGTCCTCTCTCAGCTCCAGCTTCCTCGTGAACATGCTCATAGATTTTTCCAAAGTTTTACTATGTTCCTTCTGTTTCTGCACTTTAATTTTATAATAATGGAGTTGTTGATTATCCTCATCCATTTGCCTCATGGGCTGGGCAACAGCCTCATGGTATTTTTTCAGATCCCACTTGAGCTTGGATTTACCTGAGCTGTGCCTGTTAAAAATCTCCAAGTCTTCAACAGTTGCCATGTATCCATACAAGATTCGGTCACCTCCTGGGTGGAAAATGCTTTTGCTCGGTTGATCCCAATGCTGCCTTCCTTTACGAGCATCCTTGAAATTCTTCACCAAACGTTCAGCCGAATAATATCCTGCTGGTGAGTCTGCAAATATGACGATACTCATTCCACGATGGCCTTGAGGACCATATGCATGTCTGGCTTTCAATGGGTTATATTCCTGAAACATATCTAGCAGCTCCTTGTTTCCCATCCCAATCCACTTTTCTTGCTCATCTATCTCAAGTTGGGTATTTTGTATTACCACCATAGGAGGCCAAACAATCATCAAATCTCTAGTAGCATCGTCGTTGTGCAGCCCTTTCCACTTTCCAAACTTAGTTTCACCCATAGATCCTGTCCCTGCTTTTCGCCTTTCAAGATCCTCTTCCAATACTTTGGCAAACTCTCTATGCAGCCCAATTCGTTTTGTTCTATGAGTTTTTGCATGAGCTATGAGTGGTTGCAAGCCTCTGTACCAATCAATTGCACCAACACCGCCATGGCAGGCAGGACAATGCCACTGACGATCATGCTCTACAATCTGCTCATTGGTGAAATCATCGAGCGTGTCAAAGAAGGACTGAAACCACTTATTCCGCTTTCTAGTATCGAGGCTTTTCACACTTGCATCAGAATCACAAGGGTCACTTCCAATCCAGTCACTGTCATCATCCAGATCAGCATCCGATAGCGCACTGTCGGAATCAGCTCTACCTGGTACCTCATCAAGGTCCGGAGGAGGATAAGAAACTCTACTCTCTGTTCTTGGACCTGAATTTCCTGATGTACTAGTGCTTGCAAGTTTTGAAATTTCTCTTTTCTCAGAGGGCAAAAGTTGTGACAAGATGCTGTCAGTTGCCTCCTGTCCGTTCTTCACTACTTTGGCCACAGAGGGCATACCCCACGCTTTAACCTGAATCTGTCCTTGAGCATTTTCTGTCAAATTCTTTCTGTTCATTCCCTCACCGTGATTGAGAATTTGAGGAGCAAATCTAGAGTTGTCCTGAAAGTTTCTCGGTCTTTGAGCAGCCACATCGCTAATCCTCCATTGTTGAGTGCTGCTACTCATGCATCCAAAACCAGCTCCTTCACTATGCCCGCCTGTGCTGGATGTGCCAGAATAACCAGCTCGATCGCCTTGTCCATTAAAGGAGCCTGTTTTCTCATATCTATGAGTAGGAGAGAGGTGGCCTGTTCTACTGGGTGAGGAGGGGAAACCATGGTTAAGCGAAGCTGATAGATGATTGGTGGGTGGAGGCGGATTAGGGTTCAATGAGACCTGCCTAGACTTCCGAGAGCTTGCAACTTGCCACCCGTCGTCATGATGCATTGTTGCATGTGAGCTATGGCCGGGCATTGGGCCTGCTCCTATGTTGAGGTTCTTGAACTCGCCTTCAAAATGGGCTCTCCCCATACTATCTTGCCTTGCTGGTACGCTGCCATTACTCTTGTTCTTCTTCTTCTTCTTGATCTTTCTTGGCTTTGATGGGCCTTCGGGATATGTAGGCATGGAGATGCCACCAGTAGGACCATCTTGCTGTGAGCAACCTGTTGCCTCGTCGTCATCTTCTACCTCGCTTGCCCATGAAGGCTTCCTGTCACTTACCGAGTCATGGGCCGCCATGGGAGCCTGCAAGCTTTTCTGGGAAGACCTGATACAAACTTCAAAGTGCACACGAGAAAGAAGAAAGCATACTCACAGCTAATGAAGCTCGCCATACATTATAAGAACGTGGCTAACTACCTGAAGTTTCATCTTCTCAACTACTTCATCCGCCGCGGTGTGTAGAAGTGTAAGCGCGCGCCTACAGAGAGAGAGAGAGAGAGAG

**EMBRYO DEFECTIVE 2728 (EMB2728)**:

GAACACAATAACGGAACCACGTGTGACAATACCGCATGTGTATATGGCAAGGATGAGGCCATGTCAGCATCCAGCGCGTGGCATAGTATGAAAGGTGGGATAAGGCATAGGAGGTTTTCAACCAGGAGATGGGGAAGCCACGTGGCAGAATGAGAGGCTGAAAGAAGCCTTTGGAGAGCACACTCTTGCGATCAGAAGAGGAGAGCTTCGACCGCATCCTTCGCTCCTCCATCTTCTGATCGCTCCTCCATCTGCAAGCAGGCAAGCAAATTAGCAATGGCGGCGGCCCTCTCCTCCTCCTCCTCCATGGCTGTCTCAGTCCCTTCGCGGAGTTCTTTCCTCGGAAGCAAGGAGCAGCTCTGTAGTCCCTCCTCCGTCTCCATCTCTTCAAGTAAGATATCTCTCCCTATCGTGAGAGCAACATCCCGTGTTGACAAATGCTCAAAGTCAGACATCATTGTGTCTCCTTCCATTTTGTCTGCAAATTTTGCCAAGCTGGGCGAGCAGGTGAAAGCTGTGGAAGATGCTGGATGTGATTGGATTCATGTTGATGTGATGGATGGGAGATTTGTACCAAACATTACAATTGGCCCACTGGTGGTGGATGCATTGAGGCCTATAACTGACTTACCCCTTGATGTGCATCTTATGATTGTTGAGCCAGAGCAAAGAGTTCCAGATTTTGTGAAGGCGGGCGCAGATATTGTCAGCGTGCATTGTGAGCAGTCATCCACCATTCATCTTCATAGGTCTATTGGTCAGATCAAGAGTTTGGGCGCCAAAGCAGGAGTCGTGCTAAACCCTGGTACCCCTCTGACTGCCATTGAGTATGTCTTGGATGTTGTGGATTTGATACTCATCATGTCTGTGAACCCGGGATTTGGAGGGCAAAGTTTTATTGAGAGTCAAGTAAAGAAGATCTCGGACTTGAGACGGATGTGTGCAGAGAGAGGCGTGAATCCCTGGATCGAGGTTGATGGTGGGGTGACACCTTTGAATGCTTACAAGGTGATTGAAGCTGGAGCTAATGCGCTGGTTGCTGGGTCTGCAGTATTTGGTGCGAAAAGCTATGCAGAAGCAATCAAAGGTATCAAAACTAGCAAGAGGCCAGCTGCCGTGGCTGCTTGAAGTAATATACAGGCGTCATGTTTGCAGTGCACATGTTCCTAAGTTTTCCAGTGAAAGGCGATGTCTTCGTTTTGCATATTCCCAAGCTGTAGGCTGGATGGTCTACCTTGAGCTTCTTCCCCCTCGGAGGGGGACTCCTGAAAAACAGAGGATATAGATTGGCAGCTTGGAAATTTTCAGATGGGTGATTGTCAGAGGTAGAGATATGGTCATTTTTAGGCTGCCTGGATCTTGAGAGTATTTTTTCTCGAGTATTTGTAATAATACCAAAACAAGCAACACCCTCATCCCTGTACACAATCAATTTTGTGCTTTAAGTAAGAAATGCCGTGCCTGTCGATGTATGTTTGTCATTAAAGTATGCCTGTTGTTACTCTCCAAAAA

**BRASSINOSTEROID-SIGNALING KINASE 1 (BSK1)**:

GAGAGAGAGAGAGAGAGAGGTTAAATCTGTGCTTCCTCAGTTCCGGAAGTTTTTCCTGCTTTTTCTGCTTGGATACAATCAAAGGTTTCTTCCGCCAAAGTCAAACTTGTGATGAGCTGACAAGTTTCTGCTTGTGCTATCTTCCTAAGCGATGGGCTGTTGTTTTTCACGGCTCCAAGTCTTGCAACATTCTGATGTTGATGCGACACCCCCGGAGTCAGATACTGTTGGTGCCACGGAAGGTCGTTCTGTGCCAGCATTTCGCGAATTTTCTATAGATCAATTGAAGGCTGCAACAGATGATTTTAGCCATGAAAATATAGTGTCGGAAGGAGGTGACAGAGCACCAAACATGGTTTTTAAAGGATGTTTGGATAACAAATATTGGATTGCAGTAAAGAGGTTTCCAAAAACTGCATGGCCAGATCCCAGACAATTTGCAGAAGAGGCTTGGGGTGTTGGCCAGATTCGCTCTAATAAAGTCGTCAATCTTATTGGGTGCTGCTTTGAGGGCGATGAACGGTTCTTGGTATCTGAATATATGCCCAACGAAACCCTTGCCAGGCATCTCTTTCATTGGGATCAGCGCCCAATGGAATGGGCAATGCGGTCAAGGGTTGCCTATTATATTGCTCAAGGGCTGGAACAGTGCAGCAATAACGGCCGCCCACTGTACCATGATTTGAATGCGTATCGTGTTCTTTTTGATCAGGATGGCAATCCACGTCTTTCATGCTTCGGTCTTATGAAGAATAGCAGGGATGGGAAAAGCTACAGTACAAATTTAGCATACACACCTCCTGAATTTATGAGGACTGGAAGAGTGACTCCCGAAAGTGTCGTTTTTAGTTTTGGTACAGTCCTCTTGGACCTGTTGAGTGGCAAGCACATACCTCCGAGTCATGCTTTGGACCTGATAAAAGGGAAGAATATGCTGTCATTGATGGACTCACATTTAGATGGTCAATTCACAACAGAGGAAGGAACGGAACTTATACGCCTGGCTTCACGCTGTCTCCAATATGAACCAAGGGAGAGACCAAGTGTAAGAGCTCTTGTGGCATCGTTGTTGCCCCTTCAGAGGAAAGTTGAGGTACCTTCCTATGTAATGCTTGGTGTACGAAGAGAGCCACCTTCTCCACCCCCACCTGTTGTTACTCGTCCTTCGGCACTGCGGGATGCTTGTTCAAGGATGGATCTTACTGCAATACATGAATTCATGGTGAAACTACACTACAAAGATGATTCTGGTGAGAGCGAGGTAAAGTCAATGGCATCCTTAGTAACCTTTTTCTGTGCAAGCTCTTGGTTTCTGTTTCTTGATTGCTCAAATGTTTTCTGACTAAACAACGTTTGTTGCTTCTGTATACATATGATATGTTTTGTGCGCTTTCTAGTAATAAACTAAAGAAGACTATGGAACGGCAAGGGCTTGAGAAAGACCATAACCCTCTCAATAGCATCAATTTTCACATTTTCAATATATCTTTCTTAGCTGTTAGGAGTCTCTCTGATTAATGATTTATATATCCGGCCCGTCTAGTCGTTGGATGTAATTGAAGTTGTGCTTTTTGGTCAGCTTTCTTTTCAAATGTGGACAAAGGAAATGCAGGACATGCTGAACTCCCGGAAAAAGGGCGATCTAGCTTTTCGCGAGAAACATTTTACGACAGCTATTGAGCGGTATTCCGAGTTTGTACAAGGAGGTGCCATGGTGTCTCCAACGGTGCTTGCAAGGCGTAGCTTGGCTTATCTTCTGACAGATCAGCCTAACCCAGCTCTTCGGGATGCAATGCAAGCCCAGTATATCAATCCAGAATGGCCAATAGCATTTTACCTGCAGTCCGTTGCGCTAGCGAAGCTCAACATGGAGAAAGAAGCAACAGACATGCTCGAGGAGGGAGCAAGTTTGGAGGCAAATAGGCAGAGCAGGACGTAGTTGTGTAGTGTTTTTTGAAAGCAAATCATAGTTGTGTGTATTATCTCCTTGTGTACAATGTCAATAGAGTTTCGATTGCCATGCAGATAAAGCCTTGTGTGACCACAGCATTTCGTGGTCGCAAAGAGTCGATGCGCGAGAGCAAAGCTAAATCTTTATGCAGGTTTATACCCATTCTTTGGAATTGTGTAGGTACGATGCATTGCAAACAAGCTGTTAGCTAAAGACATTCCAACCAGGCCTCTCTCCCTCTCTCTCTCTCT

**NON-RESPONDING TO OXYLIPINS 7 (NOXY7)**:

CGCAACAAAAGAGATTTAAGCAGCAGCAGCGACGACATGGCTTAGCTACACAAGCTCGGATCTCTGCAGAGTTTTAGTTTACAGAAAGAAGAAGAGCAGAGTTTCCCAGCAGCTTAGATCTTCATGGCGCTCACTTTGCAGGCTGCATCTCAGAAATTAAGTACCTCCTCCACCAAGTTGCGTGTTGCCATTTTTAGGGCCAGCGTTGTCTCTCTTCTCGATAACCCAGGTGAGAGTATTACAGAGTATGCTGCATCCCTGGCAGACTTGATCTTTGACACACTGCACCTATACAATGATCGGGCTTCTCAGTCAGCAGTTGAAGTTGTTCTTGTTAGAGCCTTGAAAGAACCATCATTTGTAAAAGTATTTACCGGGATGCTGGTACAAACAGCAGATAAAGCTACAAAAGCTGGAACCTTCTCTGTATATTACAAACTGTTAAGGTGGTCTTGTCTATTATTGAGATGGAATCCTGGGATCTTAACAGCAAAGGTAGCATTCTCAAAACTGGCTGCAGTACAGGGGGCTCTACTTTCCCCTCTGCTTCAGGCTGTTTTTCGAGTAAAAAGAGCAACTAGGCGAATATTTTTCCATACATTGACAGAGGTTCAAAGCCTTTTTGAGCTTTACATGAATGAGTTGAAGGGATCACATCTTAGCACAGAGTCTTTATCTGGATTGATCGCAATCTTGCTAGACTATTGCTCTTCAAAGTCAACAGAACTTTTTAACAACCATAAGGACCTGTTTTTAGACCTCTATATTAAAACTGTGCTAAATTCGAAGGAGAAGCCTTCGACTGTTTTAAGCTATGCATTTACACCTTTGTTGAAGAAACTGGCACATGACGACTTCGGAAAAGTTCTTGTACCAACAGCTGTAAAAATGTTAAAGAGAAACCCAGAACTGGTTCTGGAGGCAGTTGGTATTCTTTTGCAGTTGACAATCCTAGACCTGAGTAAATATGTTGGTGATCTTTTACCCGTCTTGCTTGTTCAAGCACGGCACAATGATGAAAGCAGGAGAAAAGAATCGGTGGTGGTAATTGGGAATCTTGCTTTGCAGTCTAGTGATCCAGATGCAATATTATCAATGTTTCAGATCATAAAGTCGGTCATTGGGGGGTCTGAAGGAAAGCTTTCTTTTCCTTATCAGAGGGATGGAATGATCAATTCTTTGCGTGCCTTATCCAATGCGCCTGGTGGAAAGACACTGACTCAACTGGCTGCAACCATATGTCTTTTCCTTATGTCCTACTATAAAGAAGATGGAAACGAAGAGGTGCGCACATCTGTACTTCTAGCTTTAGGAGTGTGGCTTTCCCGAACAGGAGACTTGGTCCCTGAAGGTGCACCAGCGTTTTTCTTAGCTGGGTTAAAAGAAAAAGAAACCTTACGCCGCTTTCATTTGCGATCCCTTCGTTTGGGCTTTAGAAATTCACACTTTTGTTTGAAGATGCATTCGTTGTCAGAGACTCTACTTACTCTCAGTAAGAATTGGGCAACGAAGTCTTCTCAGCGAGTGGACAGTATTTATGCTCTCTTGATTTTGTTCCAGATTGCTTCAGTTGATATCAAAGCTGATGATAGGATAGCAAAGGAAAAGCTTTGGGCATCCGTTTTACAGAAAGATTCTATCATTTTTTCTACATCCTTGATAGGAAAAGTGCAAGTTGAAGACTGTGTTGCCCTTGTAGAGCTGATTGAAGTGATCATTCTTAAGCATTCTTCAAGATTGTCGGAATACGAAATGGGGAGCCAGCAGCTTTTTCAGCTCATCGTCCACTTTCTATGGCATTCGTCAGGGGATACAAGAAGACTCGCTTTTGAAGCTGTTCGTCACATTCACTCTGCTTCTGCAAAGTTTTCGGAAAGCCTGTTTGATGGCTTTCTATCATGGCTTACTTTGTTTGAAGATCAGATAGGGGTTTTTAAAATGAGCGAAACAGAGAACACAACAGATGGATCGTTACCTCATAGTGAAGCAATAGTGAAGGCTTTGATTGCTTTAGCTGTGCCAACCATGTTCAAGGATTGCAAACTGAGTGCCCGAATTTTGCTGGCCTGTCATCATCCTTGTGTTGTGAGAGGTCGCCAAGCAAACGCAGTTTGGAAGGTGCTTCTCAGAAGGCTGAAGGGACAAGGTTTTAATGCTGTTGATGCTCTAGTCCGTGACACAAGCTCTTTGTGTGAGGTATTGTTGGGACATAATGGGTTACACAGTGAGAAGTTATTTGAGTCACAAGCTTCACTTCGTACGCTTCATACTATCATGCAGACTATTCCTATCGAGTTCTTTCCGAAGTTTCTAAATGCTTTTGAAGGTCTAATTGATAGGGGTGCTCATGATGGACTCAGTGAAACTGATATTAAGATCTTCAAAACAGAATCGGGAACCCTTTCAACGGAACAAGGTGTGTATATTGCAGAAGTTGTCACAGACAAGAATGTAAAGCAAGCAAAAGGACGTTTCAAAATGTATGAGGATGATGATCAGGAGCCTGTCAAGACTATTCTTCAACCCCGACAAGCATCCTTGAAGGAAGCACCAGGTCCTGGAAAGAAGGATGCTTCTAAAGCTCCAAAAAGGCCTGGTGAAGCTGAAAAAACAAAGTCAGCAAAGGAAGAAGCAAGAGAGTTGCAGCTCAAAGAAGAGGCGGTAATACGGGAAAAGGTTGGCTATATCAAACAGAATTTGTCCTTGGGTCTCAAAGCTCTTGGTGAAGTGGTTTCAGGGAACCCGGTTATCGCCCATGAGCAGCTTCCCTGCCTGGCAGAAAAAGTTGCACCATTGTTGAAGTCTCAAATTGTCAGCATAGAGGCATATGAAGCAATGGTAGGGCTTGCAGGTTGTGTTGCACCCTCATTACGACACCTGGCAACTGATTTGGCAGCATCTTTAACTATCATATCCACATCTCACATTTCCATTCTTCAAGATCTCGTCAATGTTTCCAAAGGACCTGATTCACAGAAGAGAAAGCCAACTGTCATAGAACGTGTTGTTGCTGGTTTAGTTGTTGCCTGCAAAGAAGGTCCTCTACCCGCGCCTTCTTTTGCCTTTGTATTCCCGATCATGGAGCAAGTTTTGGGATCACTTAAAAAGACAAATCTTCATGATGATCTGCTGGAAGTTATTTCTCTCCACACCTCCCCTGTCACGACCTTACCAAGGATTAGGATGCTGGAGGTGTTATATCATGTACTTGGGTGCATACCCACATATCAAAACGATATCCAGCCAGTGCTGTGTGAACTGTGTGCAGGTTTAAAGCAAGAACATGTTGTACAGGCACTGATAGGGATATACTCTCAGCATGCACATGTTCGAGCGGCATGCCTGTTTGCGATTAAGTTTGTGCCAGCTTTGAAGAAACGCGCAGTGTCACATGATCCAGTTTTGGCAACCACCCTATGGATTACACTTCATGACCCTGAGCAGTCAAATGCAGAGGCAGCAGAAGAGATATGGGATTTATACAATCATGAATTGGGCACTGAGTTCAGTGGCCTTTTCACAGCTTTAGCTCATGTTAACTACAATGTAAGACTGGCTGCTTCGCAAGGCTTGACTTCTGCTATGGATGACTTTCCCAGCTGTGTGCAGGAAACCCTATCTACATTGTTTTCCCTTTATGTGAAGTACCTGCCCTCTATTTATGGTGAATCAGATCCATTTTGGACGGCTAGGCAAGGGGTAGCTCTTGCCTTAAGTGCAGCTGCAGATGTTCTGACCATGAAGGATCTTCCTGTGGTGGCAACATTTCTGATCTCGCGCGCTCTGGGTGATCCGAACCTGGAGGTGAGGGCAAGAATGGTAGATGCTGGCATTGCAATTATTGACAAGCAGGGAAAAGACAATGTTGGGCTTCTGTTTCCCATTTTTGAGAATTATTTGGACAAGAAGGCTGCTGATGAGGAGCAATACGATTTGGTTCGAGAGGGTGTGGTTGTTTTTATGGGTGCACTTGCCAAGCATCTGTCAATGGATGATCCAAAAATTTCAGTGATTATCGAGAAGCTTTTGGAAGTTTTAAAAACTCCATCAGAATCTGTACAACGTGCTGTGTCGGACTGCTTACCTCCTCTTATGCAGTCTCAACAGGAAAATGCTCAGGTTCTTCTTTCAAGGTTGTTGAAACAAATGATGCAAAGTGAGAAATATGGTGAGCGTCGTGGCGCGGCTTTTGGTTTGGCTGGAGTAGTGAAAGGTTTAGGCCTTCCCAGCCTCAAGAAATATGGTGTAATGGATGCACTCAAAGCTGGAGTTGAAGACAGAAACTCAGCTAAGGCACGCGAAGGAGCCCTGTTTGGCTTTGAATGCTTGTGCGAAAAGCTTGGAAGATTATTTGAACCGTATGTTATCTATATCTTGCCTCTCCTGTTGGTTTGCTTCTCTGACCAGGTAGTTGCGGTTCGTGAAGCAGCTGATTCCGCTGCACGAGTAATAATGGCTCATCTTAGTGGGCAAGGGGTGAAGCTTATCCTGCCAGCACTACTTAAGGGACTTGAAGACAAAGCTTGGCGCACAAAGCAAGGCAGTGTGCAACTGCTCGGAGCGATGGCATTTTGTGCTCCTCGTCAGTTATCTCAGTGTCTGCCATTAATCGTCCCTAAGCTGACCGAGGTTCTGACAGATACACATCCAAAGGTACAGGCTGCTGCACAAACTGCATTGCAGCAGATTGGAAGCGTTATACGAAATCCTGAGATTTCAGCGCTTGTCGCAACTCTACTTTTGGGGATCTCAGATCCTAATGAGAATACAAAGCACTCTCTGGATGTCCTTTTGCAGACAACATTTGTCAATACAGTTGATGCACCATCATTGGCCTTGATAGTTCCTATAGTGCATAGAGGACTGAGAGAACGAAGCTCTGAGACAAAGAAAAAGGGGGCACAAATTGTTGGAAATATGTGCTCTCTTGTCACTGAGCCGAAAGATATGCTTCCTTATCTTCGATTGCTACTTCCGGAAGTTAAAAAGGTTCTCGTAGATCCAATCCCCGAAGTACGCACTGTTGCAGCTAAAGCATTGGGTTCACTCATCAAAGGCATGGGAGAGGATAATTTCAAGGATCTGGTACCGTGGTTACTTGAAACTTTGAAGTCAGAAGCAAGCGGTGTAGAAAGGTCTGGAGCTGCCCAAGGATTAAGTGAGGTACTTGCAGCATTAGGTACAACACAGTTTGAATCCTTGCTTCCAGAAATAATCCAGAATTGCTCTCACCAACGAGCTGCTGTTAGAGATGGATATTTGACACTGTTTAAGTATCTGCCAACTGCTTTGGGGCCTATGTTTCAGAAGTATTTACAACGTGTTCTGCCTGCCATTTTGGATGGGCTGGCCGATGAGAATGAGTCAGTAAGAGATGCTGCATTAGGTGCTGGCCATGTTCTTGTCGAGCATTATGCAACCACATCACTTCCCCTGCTTCTACCTGCTGTGGAAGAAGGCATTTTTAATGACAATTGGCGTATTAGGCAAAGCTCAGTAGAGCTATTGGGTGACCTACTGTTCAAGGTTGCAGGAACTTCTGGGAAAGTTGTTTTGGAAGGTGGCAGTGATGATGAGGGCGCTAGCACTGAAGCACAAGGACGTGCCATTGTGGATGTGCTTGGTAAAGATAGGCGTAATGAAGTTCTTGCTGCAGTGTACATGGTTCGATCTGATGTAAGCCTTACAGTCAGACAGGCTGCCTTGCATGTATGGAAAACTGTAGTAGCTAACACTCCTAAGACCTTAAAAGAGATTATGCCAGTTTTGATGAGTACCCTTATCTCCTCCCTTGCATCTACTTCATCAGAACGTCGGCAGGTTGCCGGTCGATCCCTTGGTGAACTTGTCCGTAAATTAGGAGGCAGAGTGCTCCCATCAATCATTCCTATCCTCTCTGAAGGATTGCAAAAAGGAAACGAAAGTACTCGGCAGGGTGTTTGCGTTGGTTTGAGTGAAGTGCTGTCTAGTGCTGGGAAAGACCAGCTCTTTGCTTACATGAGTAGTCTGATACCAACTATTCGGGATGCTCTTTGTGATAGTGTTCCCGAAGTGCGTGAAGCAGCAGGGTTAGCTTTTAGCACTCTTTTTAAGAGTGCAGGAATGCAGGCAGTGGATGAAATTGTTCCTAATCTCTTGAATGCTCTAGAGAAGGATGAAACCTCAGAGACAGCTCTTGATGGCTTGAAGCAAATATTGAGTGTCCGTACAGCAGCTGTGCTTCCTCACATATTGCCAAAGTTGGTGAAACCACCTCTAACATCCTTGAACGCACATGCTTTAGGAGCCTTGGCAGAAGTTGCTGGCACTGGGCTAAATTCTCACTTAGCCACAATACTACCACCTTTGATATCTGCAATGGGTGACAAAGAACAGAGCATAGAGCCCATTGCAAGAAAAGCTGCGGAAACAGTTGTCATGGTCATAGATGAAGAAGGTCTTTACCTTCTTATTTCAGAACTTGTGCGAGGTCTTAGTGACACACAGGTGTCAGTAAGAAGAGGCAGCGCATATTTGGTTGGGTTTCTGTTCAAAAATAGTAAGATGGACCTTTCTGAGGAGGCTCCAAACCTGATCACAATGTTGGTGGTCATGTTGACGGATCCAGATAGCCTAACAGTGACAGCTGCTTGGGAGGCTTTGTCTAGTGTAACTGGCTCGATAGCAAAGGAAAATCTTCCCAGTCATGTCAAAGTTTTAAGAGATGCTGTGTCAACAGCCAGAGATAAGGAGCGACGAAAGCGGAAGGGAATGCCTATTACCATTCCAGGTTTTTGTCTCCCGAAGGCATTGCAGCCCGTGCTTCCAATTTTCTTGCAGGGCCTGATGAGCGGATCAGCTGAGTTGAGAGAGCAGGCAGCGGAAGGGTTGGGAGAGTTGATTGATGTGACAAGCCCTGAAGCATTGAAGCCATTTGTTGTTCCCGTGACTGGTCCTTTGATTCGCATCATAGGAGACAGATTCCCTTGGCAAGTGAAAAGTGCAATCCTAGCAACCCTAGGAATCATCATAAGCAAAGGTGGTATTGCGCTCAAACCATTTCTTCCCCAGCTGCAAACGACTTTCATTAAATGCCTTCAGGATAATTCAAGACCTGTCCGATCTAGGTCAGCGTGGGCATTAGGGAAGCTGGCATCTTTAAGCACCCGAGTAGATCCCTTAGTAGGCGACCTTGTGAATGGACTACAGGGATCAGAAGGAGGTGTGAAGGAGGCCATGCTTATTGCTTTGAAGGGTGTATTAAAATATGCAGGGAAGAGCCTGAGTGCACCAGTTTTATCTCGAGTAGTTTTGACCCTGCAGGAATTTCTTCCATCTGACGATGATGAAGTGAGAGGGTCGGCAGCAAGAGCTCTAGGCATGGTTTCACAGTTTCTTAAAGAGGAGGACCTCAGAACATTGCTAGGTGGTTTGTCAACAATGGGGCAAGCTCAAGGATGGGCATTACGACATGGTGCTATTCTTGCAATATCCTCAGTATTGCGTCATACTGCTACTCAGATTTTTTCATCATCATTATTGCTGTCAACAACAGTGGATGTAATTAAAAGCAGAAGTCGGGATGATAAGGTACCTGTTCGTGACTGCGCTGCTAAGGCATTAGGCAGGCTCATGGTTCATCAGGCAAAGACACATCATCCTGGACCTTCATTCACAGACTTGATGCCTCTTATTGTGACTTTTCTGAAGGATGAGTCAAGTGATGTGCGAAGACGAGCTTTAAGTAGCTTGAAATGGATTGCAAAGGCAAATGTAGATATGGTCAACCCCTTCATGAACTACTTAGGTCCAGTTCTTGGAGAGCTAATAAAAGATGGCAGTACTCCGGTTCGAATGGCTGCAGAGCGCTGCGTTTTCCATGTCATGCAACTCAGTAAAGGACCGGATGCTGTGGCAGCATCTCAGAAATATCTAACAGGTCTTGATGCCCGGCGCATCTCTAAGCAGCAAGAGCAAAGTGATGCGAGTGAAGACAGTGAAGATGATACTCCAAGCTAGTTAGTCATTTGATACAGAGCTTTGGTGACTATATTTTTGCCTTGTGGATCGGCTTTGCATCAAATAGGTCAAGCAGCAGCTTTAACTCCTTTGCTGTACATACACAGCAAGCTCTTGACATACGAGTTCTTCAATTCAGAGAACATAAGAGGGCTCAAGGAAAATGGCAGGAACAGTAGAACCCCATAAGAACGAATGCGTACTCCCTACAGCTGGTCATTGAGCTGGAGTATAGCTTCCATATCATAAAGATGCCTGCAAACTGCTGTCATTGCCCGAGTATTCATGGCGAAGAGCACCTCTGCTTCTCTAGAAAGGCAAAGCTGCAGTATGGGGAAACCCTCTCCTCGGGTAATTCACAATCGCTGGTACGAATTTTGTTGGCAGAGTGTAATTCAGCTAGACTATTTACAAGCTTCTACATCTATCAGTTTTTCAGAGATTTCTTGATCTGGTAGTAGGAGGCTGTAATATACTGGCTAGGTCGATTTTCTTCAACAGTCTTATAGTGGAGGCCATGGCATTTGTGCCATTGAGGTTTTGTGACTAATGTTGGTTGCACTTATGGTCGTTTGTATCATATGAGGCTATACGGTAATGGTGAGGAGTTGTTGATTGACTTAATCTTATTCAATGCTTGTAATGTCTCAAGGGGATGCACTCACAAATTATGGAGTAAGTTAAAACTGGAACAATGTATCTAGTGTCTTTCTGTTTTAAAAATCATTGAATAAATGCATACATGAGGGGGGCTTGCTAGTCTCCACCTCATAGCCGGAGAGTTGTATCTCCAGTAAAGACTTTTTCCCTTGATCATTTTTGGAAGGTACAAGATGATTAACCTTTTACCCTAGTGTGCAAAGCAAGCAATCGGAATATGTTATTACCAATTAGATGTTTAATGTTGTGAAACGTGGATCGGTGTATTACCATATATGTCCCATAAGGT

**SERINE/THREONINE-PROTEIN KINASE/ENDORIBONUCLEASE IRE1A (IRE1A):**

AGAGAGAGAGAGAGAGAGTTGATCACCGCCGTCTTCTACGCTTACAGAGAAAAAGAAGGCATAGCAGAAAAGAAGACGAAGTAGCAGCACCAGCAGGAGAAGAACAAGCACAAGCAGCAGCAGCAGACTTGGACGAAAGCTTCTCAAAGCCCAATTTCATCCCAATCTAGCAGAGAAATGCAGAACAAGGTTCATTAGAAAAGAGGCAGACATGGGCAAATTCTTCCGCAAGCCCTTTGTGCCTGCATTTTTTCTTGCATTTTTCTGCTTCACAGCATCGTTCACTGCAGCTCAGCCCGTCTCCGAATCTTCTGATGCATCACCATCTTCATTGCCTCCACCATCCCTGCCTCTCCCTCTTGGAGAAGATCCTGACAAATTTCAGGTGACTTTGGAGGTTTTGTGGGATGGAATGGTGACTGCTCGAGATTCTCTTACAAAAGAAGTGAGGTGGTCATTTTTTTCAGGAAATCCTCTAAGCTCAGCTCATGGGATTTTGCTGGATGCTTATGAGGGTACAAACTCATCCATCGGTGTAACAGGCGATACTCTTATTTCTGAGGACGAGTTTCTTTTCTGTGGCACAGACTGGAAACTCTACTCTTTTGACAAACAGAACGGCATAGAGGACCTTCCCATCCCACCTCATGAGCTCATACAAAACACTCCCACCATCCTGAATGGGGATCTCATTCTTGGCACCATGTCAACTTCTGTGTTTCTTTTGGATGCAAAGTCTGGGAATCTGGTACGCACGTTTGAATCAAGGAATGAATCGTTTTCAGCTGATCACTTTGGCGAAGAAGACATATGGCGCTTTTTAGAACTTAATTTTGAAGAGCGAAGAAACTCAGACTACCTCTTCGTATTTAGGAAGGATTACACAATTAAGTCTGTGTCAATGGCATCAGGAGCACTTTTGTGGAACGTAAGTGTGGCAGAGGTTCAGCTGCATCACATCAATTCTGATGGCATTCTTCGTCTTCCAGGAGGAGGGGAGAGAAGCAAAGAGACATTCAGTTTGTTAAGCCAAGCTGTTCCACCACCCGTTGCGTTGCCAGCCAGCTATGAAAGGAATGATCGGTTTCTTCCAGCTCAAGTGCAGGTATCTCAAGATAGTGCATTGCCCTTGGCACCAGCTCAAGTGCACACATCGCAGAATAGTGCATTGCGGTGGCAAATTGGACTTGGATTTACTGTCCTGACGACCATGGTATGCGTATTTGTCACACGTTTCTTCTGGAAAAGATCTACACGAAAGCAAAATGAGGAGGAAAAGCAGGCTCCAAGGAAACGGAAGGCCAGGAAAACAAAGGCAGTGGCCGGAGTGACAGCACAAGAGCCTCCACAAGGTGAGATTGGTCCTCTTCAAGAAAATACTAATAGCTTGCAATCTCATTCAAAAGCAGATCCACTCATGGATATAAATCAGCCAAATGTCAAGTATCATTTTTTAGGGGGCCTGTGTGTGACTGATAAGACTATTGGTTGTGGAAGCAATGGAACCATTGTATTTGATGGTTATTTGGAGAAGCGACCTGTTGCTGTCAAGAGGATGCTTACGCAATATTATGATAAAGCTCAGAAGGAGATAAACCACTTGATAAAGTCTGATGAACATCCTAATATCCTGCGCTATTTCCATGTGGAATCAGATGCACATTTTGTCTATGTTGCCTTGGAACGTTGCCGGTTCAGCTTGAATGATCTAGTTCTTGCCCAAAGTGCTCAACACTTAGGAGCATACGAAATCAATGTTATGCCCGGAGAACATAGCAGGGAGGAGTTGAAAGTTGTTCTAGAAAAGGTTGGAGACACCAAAGATTGTCAACTATGGGATGAATGGGCTCGTCCATCTGCAAAACTTTTGCAGTTCATGAGGGATATTTTGGCGGGACTTTCTCACTTGCACGAGGTGGGCATAGTGCATCGTGATCTGAAACCCCATAATGTGTTGATAAGCGGTGGGCGACTGTCACAAGCAAAACTATCTGACATGGGGATTAGCAAAGAGTTGGCTGATGGAATAACCTGTCTAGACACGCAGTCTACAGGTTCGGGAAGTTCTGGCTGGCAAGCTCCTGAACAGCTTATGCGGCAGGGACCACAAACGAGGAGTGTAGACTTGTTTAGTCTTGGCTGTATACTGTTTTTCAGTATAAGTGGAGGCTACCATCCATTTGGTTCACGTTTGGAACGAGATTTTAATATTGTCAATGAGAAACTTGACCTTTTTCGAGTTGAGCATGTGCCTGAGGCGACAGATTTAATATCAAAATTACTTCAGCATGATCCACATAAGAGACCCCAGATGAGTACAGTGAGGATCCATCCTCTTTTCTGGAGTTCTGATATACGATTAAAGTTTCTATGTGATGCTAGTGACCGCGTGGAACTAGAGGATAAGGAAAACCAGTCGCCTATTTTAGAAGATCTTGAGGAGGTATCACAGCTGGCTCTTAGTTGTTCCTGGGCTGACAAACTAGATGCTGTTTTTCTACAGAATCTTGGAAAGTATAGGAAGTATAACTACAATAGTGTTCGAGATCTGCTTCGAGTCATCCGTAACAAGTCCAGTCATTTCAGAGAGCTACCTCAAGAAGTTCAACAATTATTCAATTCATATCCAGAAGGCTTTGAGACTTATTTTAGATCCAGGTTTCCCAGGCTTTTGCTTGAGGTGTTCAAAATTTTGTACAGACATTGCGGCAATGAGGATGGATTCAGAAAATATTTCTCATATGATTTGTCACTTCTGTGAGGGGATGCAGTATCATATTGTTATCGCTTGAGATTCTGATACTCTTTGTAAACTACAGCAGGTATCAAATTAGACAGCTCAAAATACCTCTCTTTTCATTGCTAGCCTGTCATGTCTGTTGTTACTCTTGTGAAGCATATCATTGATCGCTGTCCTTTTTTTGCTAGAGCGAAGACTCTACCTCCTGGGACCCTCAGAAAGTCACATGTTTAATGTGACGAGGACGTTTAAGTTAAGTTGTGCGTTAAGCTCCATGTGTCCAAGGATGTCCTTTGAAAGATAACATGACTAGTGGGAGCAAGTTGCATGATACTTACTAGTGCTGGCTTTGGCAGTATTTGATCTCTGAAGTCACCCATTCCTTCCATTCTCATTGCCCATGGCAGCACCAGTAGCAATGGTTGTATAACAATCTGCTCATGGAGCCTTGAAATCAGCTTCTGTTCGCAGTTTTATTAGTGACACAGGAAAACGGTTAAATTTGTGTTTGTGTGTGTGTGTGTTTGTGTGTG

**PLANT U-BOX 59 (PUB59):**

TGCTGCTTGCAGGGTTATTGCAAGATTGAAGAAAGAAAGGGATGAAGCACATAGCTTACTTGCTCAGGCAGAGAGGCTTTTACCTGCTCCGTCTCCTATGCCAACTACACTAGCACTGAGCAATGGAAAACGAGCTGCTCCAGAGAGTGATCAGGAATCTGCAGCTGCCAAACGGGCTCGGCAAGGAATTACTCCTGAAATGATCAAGGAGCTGACCGATTGCAATACCAGGCTATCAGCGCAAAGAAAAAAGCGCCAGGTTTCTCAAGATTTGGCACCCCCAGAAGCTCTGGAGACCTATACACTACTCTCCAGTCATCCTCTTCACAAGACAAACAAGCCTGGCATTTTGTCAGTGGATGTCCACCAATCAAAGGATCTAGTCTTGAGTGGTGGAGTAGATGCCAATGCTATAGTGTTTGACCGATCTTCTGGAGAGATAGTATCAACCCTGAGCGGACATAGCAAACGGGTCAATAGTGTTAAGTTTGTTCCTCAGGGTGATTATGTTTTGACTGGATCGGCTGACAAGACAGTCAAAATCTGGAAAGCAAACGAGGATGGAGTTTATGAGTGCAGGCACACTGGTCGAGATCACACTGGAGAGGTCCGTGCTGTGACTGTACATGCAACTCACAACTATTTTGTAACGGCATCATCTGACAAAAGCTGGTGCTTCTATGATCTTACCTATGGAGATTGTTTGGTTCAGGTGGCCGAAAACACAATGCAGGAGGGCTATACATCAGCTGCATTTCACCCTGACGGGCTTATTTTAGGCATGGGAACAGCAGAATCCTTGGTACGAATTTGGGATGTCAAGTCTCAGGCCAACGTTGCACAATTTGAAGGGCATAGTGGACCTGTGACTGATATATCTTTCTCGGAAAATGGCTATTTTCTTGCTACAGCTGCTCAGGATGGTGTGAAGCTGTGGGATTTAAGAAAGCTGAAGAATTTCCGCTCCTTCTCACCATATGATGCTGATACACCGACAAATGCAGTGGAGTTCGACTTCAGTGGAAACTATCTTGCGATAGCTGGATCAGACGTAAGGATTTATCAAGTTGCAGGTGTGAAACAGGATTGGAACACCATCAAGGTCTTACCTGATCTGTCTGGGAGAGGAAAAGTAACATCTATCAGATTTGGGCCACTCGCGAGCTACATGGCGGTTGGGACGGAGGACCGGAATTTAAGGGTACTTGGTGTCCCCATTGCATCAGTGGAGTCTTGAACTGTAAGGCACACATTGACTGGTGTTTGGGAAGAGCTATTTGCTTACTTTGTTGGAAGGCCTTTCGTGTATGGGTCACTGACCATGGGGAGGACATCGTGCGTGTGGGGACTGTAGAGACTTGTTTAAGGCTTAAAAGCTCCTTGATGGTTTGCGTTAACTTGGTGAGGGCAGTCCAAATCTCCTCCTCCTATCAGTTAAGAGATCTGAGTTTGCTCATCTAAAAAGTGCTCTTTCACTAAATTTTGTGAGCTAGTCTCGTACGAAGTGAACCAGTTTGAAACCTAAAACCTTGGCGACGTACTGCTTTCAATCATCAGTTTAACAGATCCCATTGCAGCAATAGAATGCATGTACGATTATTTGACTATTTGAACTAGATGGCACAAGCGATAGAGTGGATGTACTAGCGAACTGGTCCAATTCTTGTAATTCAATTCAAAAGTTTACCTCCACTTCATTATAGTGGAGGCTCTCTCTATTAGCACTGCTAATTCTTGACACTAAAATGCTGCAGCTTGTGCCATCTACCGTTTCTTCTCTCTTGGATCTTTGCTTGAGCTTTGGCCTTGGGTATGTGATATTCATGCGGAGGATCAATGCATAATTTGTTTGAAATCTTCAAAGGGGGAATTGAATGCAGCAGTTCCCACGCTCTTCATGATACAAGCTTTAAGCTCCTTGCTCAGCACATGTTGCTTGAGAAGAACTGCTTGTGTCAGAACAGAGGTAGTGATATTACTATCGTCCAAAGGTAGGTTAATTTACTATGGCCTTTACAAACATCAATTCTAGGTGGCGAGCTTGTTGATACAGCTGCTCTGGCTACCTTTTTCAAGTGGTAGTATATCCTTTCATTCACCAGGGTACATCTATTCTGGGTGGCAAGCTTGTTGATACAGCTACTCTGGCTACTTTTTTGCGTGTGCATGTGCGTACGTGTTCATGCAGATACACACAGTAAGTTGCTGGAATGGTTGAGACCAAGTATGACATTGGAATACTGATATGAAAAAGTTCATTATTCTGGAGGATGTTTGACCGACACATGTCGAGGTAATAGGCTTGGAGAACTTTGTGGGCCCTAGGCACATCCACCAGATGGAGTTCAGCTGCCAACAAGACCCAAGAACACGTGTCTGGCATTCATGTTGTTCACCTTCTTATGGAGGAATACCTCTTTTGGCAATGAGCTGATTGAGTTGCTTACGCTTCGTTGATGGAACATTTTTATTCTCAGTCAGCTGATCTGTTGATGAGGTTAATCTTTTGATGTATCTCTATATGATGAGATAATATTACACCCTGGATGATGCCAATGTAAAGACAGACTAATTTTTTTTATTGTTTTTGGCTGAGCCAGAAAAGGTTGGGTGATTTACATGGCCCACATATTTACCTTATTTTATCGGCAGAAACAGGCCCTTCTGTTATGATTTTTGTACTTGAGATGGTCCATAGTTGCGTTAGCTGCTATTTTGAAAGATGATGAATACTTTGAGGCTCCTCCAGGAGAGATTTTTA

**SUPPRESSOR OF CPR5 44 (SCPR44):**

TTAAGTAGTAAAGATCGTACATTAGGTGAAATATGAGAAAAATAGAACAAAGAGTAGCAGAGCAGGAGACGTAGAGGATGCCGGTGTCTGTAGAGAACGCGCAGCCCGTAGAGCCTCAATCGCTCAAGAAGCTCACCCTCAAGTCCCTTAAGAGATCCTTCGACCTCTTTGCCTCCTGCCATTCCCTTCGGCCTCCTCCCAACCCCGACAGTATCAGGATTCGAAAAGCTTGCAAGATTCTTGAAGAATATAAGTCTGTTAAAGATTCAACAGTCTATCCCATACACGATGGGGCGAGTGCTACCCCTCACACAACTGCAGATGGTGGTCCTCCACTTGCTCTACCTGCACCAGCTGAGGCAGACAACAAGGTTTCAGAATTTGCTTTGGCTATCGTTCCTTCTCAACCTTCAGGAAGGGCTGATGGTGCAGTGGAGTCTTTGGCTGCACCGACATCGTCCACACTGTCAGAATACAAGCCCCCAGAAAGGTATCAACCAAGTGCTGCAGTTATGAGAAGGCTTCCAAGCAAGTGGCCTCGTCCTGTGTGGCATCCTCCTTGGAAGAATTATCGGGTTATCAGTGGCCATCTTGGGTGGGTCAGGTCAGTAGCATTTGATCCCGGAAATGAGTGGTTCTGCACTGGATCGGCTGATAGAACCATAAAGATATGGGATCTTGGAACAGGGCAGCTGAAGCTGACTTTAACAGGGCATATTGAACAAGTTCGAGGCCTTGCAGTCAGTCCAAGGTTTCCATATATGTTCTCTGCTGGAGACGATAAACAAGTGAAGTGCTGGGATTTGGAATATAATAAGGTCATACGCTCATACCATGGACATCTTAGTGGAGTTTACTGCTTAACCTTACACCCGACCTTGGACATTCTCTTGACTGGGGGACGTGACTCTGTTTGTCGAGTCTGGGATATGCGGACAAAAGCACAAGTCCACGCTTTATCTGGGCATGAGAATACTGTTTGTTCAGTTATTACTCAAGCTATGGACCCTCAGGTTGTAACTGGGTCACATGATACAACCATTAAGCTTTGGGATCTTGCTGCAGGAAAGTCCATGTCGACCCTCACGTTCCATAAGAAGTCGGTCCGAGCACTAGCAATGCATCCTTTTGAGCATTCTTTTGCATCTGCATCTGCTGACAACATCAAAAAGTTCAGACTCCCGAGGGGAGATTTTCTGCATAACATGCTGTCTCAACAACGGACAATAGTGAATTGCATGGCAATAAATGAGGAGAATGTGATGGTTTCAGGAGGTGATAATGGAAGCATATGGTTCTGGGATTGGAAAAGTGGCCATAACTTCCAGCAAACACAGACAATCGTTCAGCCTGGATCTCTTGACAGCGAAGCGGGGATATATGCCTCGTCCTATGATATTACAGGGACTAGGCTTGTCACATGTGAGGCTGATAAAACAATCAAAATGTGGAAGGAGGACGAAAATGCAACCCCTGAGACGCATCCAATTAACTTCAAGCCTCCCAAAGAAATGCGGCGGTTCTAAACAGAGACGGCGCTTTTGCTTAAGGATTGCGATGTACGATTCTTTTGCCTGCTGCGTGAAGAATGTTAGAAATGGAGTCTCCGCAAGGGATATAGTCTAAATTTCAAAGCTTTGGATTAGTGGAAAGTCGGCTCATGAAAATACTTGGCTGTGTAAAGGCACCAGCAAAGGCACTTGAGGATTTTTCTTTCTGCAAGTTTGTGTTCATAGTCTAATAGGCAATGGCAAAAGACTTTGAGATGTTTCTTCTAGTGCTTGCAGTGTGGGTATTAATATGGGATTACGACACTTACTTTACAACGATATGGGTCCACATGCAGATTAATCTGCATAAAATGCAGCAGCAGTCTCTGTGGATGCTGGATTTCTATTTTTGATTTTTTTTTTTTTAAATGTGGGCAGCATAATATATTAGAGAGAAGCATACAACCATGAGAAGCATACAACACTGAGTTCTAGCTCAGTTGGTAAGGTGTTTGACTGGTTATTCCCAGGTCTTGGGTTCAAGCACCGGCTACGCTCAGAAAGTTACCTTACCCCTGCCTTGTGGGATTTTTTTGGGGGGGGGGTGTCCCATCCTTGTGGTCATTTTGAGAAGTGTTATGCCAAATTTACATACTTTTGACTAACAGGTTTTCATTACATTAACTTTCGGAAGATCACCAAAATTTTCGACATCTGTATTATCGATAATATGCTGTTATTATTTCCAAGTCAGATTGGTTTTCATCGAAGCAAATATTGAAAGAACGACTTGTATTATGTTCAATTCAAGTATGGGTGTATGAAGTCTTAGATTAAGGGGCCTTAAGACCCTGCTTTTATGTTTGCTGAAACAACTTTAGCTTTTTAACCTGCTCTTCTAAATTAAA

**ASPARTYL PROTEASE APCB1 (APCB1):**

CGTGAATTAGAATTGCCAGTCAGCCTCCCGAGGGCGAGGAGGAGCGCGGGGGAGATGGCGCACTCTGAATGCAGCTCTACAATATTGTAACCCTCTGCTTCTTTCTGAGTTGTACTCTTCGTGTAATCACGATTCACGCTTACATTTTCCTCTTCATGAAGGAACCCATTTTTTCAGCCTAGATTTTCTGCTCATTTTGCCCCGCTATGGAGGATGGGGGTAACATTAAAGGCATCTTTATCATCAAGCTGCCACCGCCTAATGATCCCTCTCAGAAGAAGACGATCACAGGAGCTTTAACCTTTGCGGAAAACCCTCATACTGAGCTAAATCCGTATCACCCTAGTCTGAATCCACCTGATGTAGGCGATGACGAAGACTACAGGCTTCCTTTGCACACAAACCATGAAACCTCAACACCTCTTCGACCTGCAAGGCGAGGGTTCCGCTTTCTGAAAAGGAAAGGGCTCGCTTCGGTTCTGTTTGTAGCTGCAGTTGCTTACTTCCTATGGCAGTATGGACCTTCAACTCTCAAGGAAGAAGTAGAACTTGACTCTGATAACACAGAGGAGTCGCAGCGAGCTATGGTTTACAGCATCTACTCCAAGGTTGCGTTCTACCCCAATTCAAACTCTTCTTGGGCACCTTCTATTCCTGCTTACCCCAGCAACACCACCCCCATTTCTCCGTCTGCCATATCCAAGCGCGCCGAGAGGGATATTGTTCGTAGCAGTACCAGGGTTGGCAGTGCAGCAGAAGCAGCGCCTTCTGCCGATTCTAAGGAAACATCCACCACCATTTTTCCTCTGCGGGGCAACATATATCCTGATGGGTTGTACTATGTATTGGCTAAGATTGGCAATCCTCCACGGGAATACTTTTTGGATTTGGACACAGGAAGTGATCTTACTTGGTTACAGTGTGACGCCCCTTGCTCCAATTGTGCCAAGGGACCTCATCCCCTTTACAAACCAAGGAAAGCCAATCTTGTGGACTGTAGTGAAGCAACTTGTGCATCAGTGCAAGCAGGAAGTAAGTTTGGGTGCGGTCAGGGCTCTGGCCAATGTGATTATGAAATCATGTATGCAGATGGGGGGTATTCCTTGGGCGTTCTTGTCCGCGATCAACTGGCCATGATGCTTACCAATCGCAGCTTGGGTTACACAAAAGCTGTTTTCGGGTGCGCTTATAACCAGCAAGGTTCTCTAGCTAAGTCTCCGACTTCCACCGATGGTGTGTTGGGCCTGAGCTCTGCAGCAGTATCTCTTCCATCTCAATTGGCAGAACAAGGACTTATTAAGAACATTTTTGCACATTGTATTGCTGGAGATGGTGGGGATGGAGGTTATATCTTTTTAGGTGACAAGCTGCTGCCGTCGCGCGGCCTAACGTGGGCACCCTTGCTTGGGAAACCAGCAATGAAGAGTTACATGTTAGCAATTGAGGATGTCCAGGTTGGAGATACCTCGATACGTTTGGGTTCTAATGGCTTTGGAACAGCCATCTTTGATAGTGGCACTTCTTACACTTACCTGATCTCCTCCTTGTATACAGCCATTGCCACAGCTATCCAAAACAGCTTAGTTGGCAGTGGCTTTAAGCATGATTTGTCAGACAACACTCTACCGCTTTGCTGGAAACATGGTGTTCCTGTCAGTGCACTCAACGATGTGCTGCACATTTTTAAACCTATCATATTTGAGTTTAGAAACAGTGGTGGCTGGTTCAGGAGAAACAGTAAAATGCAGTTAACACCGGAAGGATATCTGATAGTCAATTCTAAAGGAAATGTCTGCCTTGGGCTCCTAGATGCAAATGTTATGGGCAATCATGGGTCAAGCATAATAGGCGATATTTCAATGCGGGGAAGCTTGGTGGTCTATGACAATGAGAACAATCGATTGGGCTGGATGAAACAAGATTGTAGGAAGATCCCGAAAGATGCTGTTTTGTCTCCATCAGTTTGACAGTGGTCGTAAGTTTTCATGGTCTTTTGCTGAGTTGTTGGGTGTTTGCTATTTTTCATGAGAAGATAAAAGACCTTGTCAAGAAAGTGGATTGCTTTTTTACCTTCATAAACTATTCAAAGGATGGATACTCGAGCATTGGTAGGTTGGGTTGGTTATTGGAATAGCCACCAACGATTTCATGTTTGCAGGCCTTTTTAACACATGGCTGACATAATGCAGGTCCGTTTTGGATGCCATACATTTGGCAATGCTATTTGAGATGACATTTACTGTTTCTTTGCCTGAACGCTCCTGCTGACGAGGGGCTGCCTAGGTATGGTGCTACAGACTCACCAGCTTCTTACAGTGGTCCTTGATGGGTGTTTTTTGTCAACCAGGGGGTCTATTATTAGAATATACACGCTCGAGTATATCACTTCATAAGTAGCAAAGATTTGTGTGTGAATATTAGCCTTGGGCTCAGGAGGTGCTTATTTGATTTTACTTTTGGCAATTTGAAGAAAGGCACATGAATATTTGTATTGAGTACTGTACTATCATGAACCAAAATATGATGTATGATATATGTTATATGATGTCCATATACAATGAGTGAGTTTCTTGTATGATATATGACATATGATGTCCATATACAATGAGTTTCTTTTAACTATCTTTACCAGGTG

**ASYMMETRIC LEAVES ENHANCER 3 (AE3)**:

TCCCAGACTATGCTAAATTGAAAAAAGGACCTAGAGACCAGGTAGAGAAACAAAAATATGGTGCTCGTCACTATAATTGGCATCATTTTTTGAATAAAATCTTCAAGGGCTTCAAGCAAAAGGAGCCAAGTATTACAATCCATAGCCAACCTGTTCCATACCTCCGATACACCTTGCTTGGATGTTAATGCAACTTCAATGGTAGACTTGGCCTAACATACTTTTTCTAGTCCTCTTGAAATATCTTCATCTACACCATAAGTATAGCTTTGCACCATTGTATTGAGACGAATAAAGACTACCTAGCACATAAACCCAAAAATTTGCTTGCTGCACTTTTCAAACCCTTCCCCATCAAATTACATCTTTAGACATGCAGGAGAAGCTTCGCAGAGAAGCTTCGCAGAGTTGTGATACAGGCATGCACAAGAGCAATCACGTTTCTTTTCAAATTAAGGCATTCATATAGAGTATCTAGTCTTGTTGTGAACAACACTCATCCTATTTTCATATCAACATTATTCATCGCAAAGTTTCCAGTGCAGGGTTAAACACTGCAATCTGCACCTAGAAAAAAAAACCTCAACCGCTGCCAGCATGCACTGGACAAGTGTTCATTGCATTCACTATCAAGATGAATAGACTACATTCTCAACTCATAGGTGACAACAGATCTGTTGTGAAAACAAAGGTTAAATTCATCTATACCAGCCAAACACAGATGATGCATGTGTAAAGAAGCATGCAATCTGGAAAATGGCTTACAAAAAAGGTAATCTGTGGTTTTAGTGTGGCTTTGCTACCCAGCAGCTGAAATGGCGGACCCTGAAGATCCATCTGCAAGCTTCTCATGCTCCTTGTTCAACACCTTGTTGTTGATGAGATTGTGCAACGCGATGACGCTTCGAATAAGTGAAGCCAGATAAATGACCAGCATCATGTCATTTGTTTTCACTGCGAAGGACTTAACAAGCTCATTGACATTCAAGTTTGGCAGTAGATTGAACACATCCTGTAAATGGTAAAGAATTTCATGATTCAGAGGCAACTTTCCATCTATAACGTAGTCCAAGTACGAATGAATCTCCTTCAAACGTGACTCAAGACCCTTCAAAGCAGTCAATTTTCCACTAACCTCTGTTGCAAGCGTGCTTATAGTTGTGTCCTTCACATCTCTCAAAAGATGTTCAACACCAATTTCTTCTGCCTCATATGCACCAATCTCAGAAGCTACATGCACAAAAACTTTTTGGCTTTTCTGAGTGGAATCCTCCTTAACTTCTTCAACTGCATAGTAAGCCTTAGTTGGAATTCCGAGCTCTTTAGGCTGAACATCAATGATTACGAGCACTGGATTGGGGACATAGTCTCTAAAAAGCTCGTGTACATTTAAATCGTTCTCGCGCAACTTAGGCCCGGTACTATACCAGCCTACAACGTGTTCTTTGGCATTGATTCTTTTGAACATGTCAAACATAGCCTCATGGTAATTGTGGTCCAGGAACCAAATGCTGGAATCCTTATCATCCTCCTCAAAAGGAACTGCATAGCTGTTGGTAACATCGACACGGCCACGAAAGACAGCTCCAAGAAGAACGCCAACAACGCGCTTGCGAGTCTCCTTGGCCACACGATTGTAGTTATCAACAATGCTGAGCAGAACGAGAGGGTGAACGACGACCTTCTCGATGGTTCTCTCCCCTATGTGGCTCGATGTAACCACGTCCATCTCCACCTGCTCCTTCTTCTTCTCTTCTGCGCTTCTCTTTCTTCGTCGTCTCCTCGCGCTCTGTGTGCGTGCGTTCGTTTAGCTCCCCCCAAAAGCACGCAACAAAGAAATGTGGAGTTGTTGAAGCAACACGCCTTCTCTGTACAGACCTTACTTCACTTCCCAATGTTGCCCTTTATTTGCTCACCCGGACAACCTTCGTCTTCAAAAGCTAGATTCCTA

**NATB CATALYTIC SUBUNIT (NAA20)**:

TTTTGAGTTACGCATCAGTATGCTCCTCCGCAAAAACGGCCAAATCCCACATTGCTTACTCCCGTAAAGAAAGAAAGAATGACGACCATCCGGCGTTTCTGCTGCGAGGACCTCCTTCGTTTCAGCTCCGTTAATCTCGATCACCTCACAGAGACTTTCAACATGTCATTCTACCTCACATACTTAGCCAGATGGCCAGACTTTTTTCATGTGGCAGAAGGGCCAGGCAATCACATCATGGGCTATATTATGGGAAAAGTAGAGGGACAAGGAGAATCTTGGCATGGTCATGTTACTGCGGTAACTGTAGCACCGGAGTATAGGAGGCAACAACTTGCAAAGAAGCTAATGAATCTTCTGGAAAACATCAGTGAGAAGACGTACAATGCATTTTTTGTGGATTTGTTTGTCAGAGCCTCTAATGCACCGGCTATAAAGATGTATGAGAAGCTTGGATATGTCATATATAGGCGCGTGCTAAAATATTATTCCGGTGAAGAAGACGGCTTAGATATGAGGAAAGCTTTACCTAGAGATGTCGACAAAAAGTCGATTATACCATTAAAAAAGCCAGTCACTCCTGAAGAGCTGGAGTATGACTAGACACTATGTTGGCTAAATATATAGAGACTGCTGGTCATAATACTCTGGCAGTATGGTTCTTTTCTACGAGCTTATGGCATTAAGTTTTATGACCTTGTCATAAAAAACATTTGCTTAGAAACCTGTGTGTATACTTGCAGCTGCTCTATAGTTTTGTGAGTGTGAGTATTCTCTCTTAGGCTACATATACTTTGCTCTGTAATACAAGCAGCCTACCACCATTCATTGATTTCACCTTGCCATCTTTACTCTTCGCAACTGCTTCCGATATATACGAGCTTTCTGGTTCTCAACTCCACAAATAGTTAATGATATGCATCATGATGGCAGGGAGATGAGTTTTTATAAGTTGTTCGTTTTCAACAAGGACAATTAGAGCTTTAACTTCTTTCATGCAAAAAAA

**MITOGEN-ACTIVATED PROTEIN KINASE KINASE KINASE 1 (ARAKIN)**:

TCTCTCTCTCTCTCTGTGTCTCTCCTGTGTGCGCGTCTTCTCTCTCTACCACGAACCCAGTCCTTGCTCCTGAAGGCGCAACAGCTTCGACCATCGGGATTCATGGTCGCCTGCACAGAATTCCCTCTCCTTTTGAAGCTTGCTAGCCATGGCTTCCTTATCCAAGCAATCCTCCGCAGGTAAACCCAAGCTTGAAAGACGCAATGCAAGGAAAATCTCTGATTATGAGCCCTCAGATTCTCCACGTACAGCAGCTGTTCCCTCTCCCTTCAGAGCTCTTGACAAGGGCTCTTCTAAAAGCTTCAGGGTCTTAGGGATTGACAGTCAAGGGGAGATCGACAAGATATGTGAAGAGTTGGGCATTGACGGTGGCGCTGATGCATTCAGCATAACGCCAGACGCTTGGGAAGCAGGAAGGGCAAAATCAGGCTCGCTCGATCGCTTCATCCTAGAGCACAAAACTCGTTCGAGGTCCTTTGATGAGCCGCCGGGCAGCTTCTTAGGTCACCGTGCCACAGGAGCTGGGGATGGTGAGCTTGCCAGCTCCGTCCTGGCAAGCATGCGCAGCCTTAGCCCCAGACGGACCAGCCCATTTTACAACTCAATGTCTTCTTCATCTGGGGACTCGTTCGCTTCCGACAGCAGCCTTGTCACAGGGGAAAAGGGTTTTTTTGGATCTCTCGATACGCCAAGAGAATTGGTGCCAGTTATGGTTGTTAGTGAGACTCGGCATCATAATGATCGCCAGCAGCCTGGCAATGTGATTCAACGCATGCACAACCATGGCATACATGAGACACAACATCTCGATGATTGCAAGCAGACTGGCAATGTTGCCCGTGACACCATTCATGAGACACGGCCTCAAGATAATCGTCAGACTCGCACACACCTTGGGAGTGGCCATGACGCGCATGAGGCACAGCATTTAGTTGATCGCAAGCGGACTGACAATATTGCTCACGACCTCTCCATTAATGAGACACGGCGTTGCGACAATAGTCAGACTGGCAATGAAATGGAAAGGGTTTCAAGAGTGTTAGAGCAAGCGGCTCGCTTAACGATTGAGACTAATCGACAAACGTTACTACCTCCACCTGGATTTCGGGTAAAGTCGTACCTTCCTTCTCTCTCAACTCAGGAGCTGATTGCGAGCTTCGGGCCAGATGAAAGTGATAACAATGACCGAGCAAGCGATCATGATGACGCATCCCATAGTGATCCTGAAGACTATGAAGCTGATGTAATAGAAGACAAAGCAAGAGAAGAAGTTGACTTTGTGTCAGAAGTAGTGCAACAGCACGGGGTTTTGGCCAAAGAGCAGATCCTTGTGGAAGACACAGTGGCTACGGTTCCCTCCACACCGGAACCTGCCAATATGATAAGCATTCAGTCTTGGAGAAAGCTTGACTTGCTGGGCAGTGGCTCATTTGGAACGGTATATGAGGGGATGTCTGAGGACGGAGTATATTTTGCAGTGAAAGAAGTGTCTTTGTCCGAACATGATAGCAAAGCGCAGCAATGTGTTTTACAGCTTGAGCAGGAAATCGAGTTTTTGAGCAAATTTCAACATGAGAACATTGTACGATATATGGGTACTCAGAAGGAATCTGATAAGCTGTACATTTTCCTCGAACTTGTCACCAAAGGCTCACTTTCTTCACTTTATCAAAAATATACTTTATTCGAGTCACAAATACGATCCTATACACGGCAAATTTTGAATGGTTTGAAGTATCTGCATGATCGGAAAGTTATGCATAGAGATATTAAATGTGCCAATATATTGGTGGATGTCACCGGAAAGATCAAGGTTGCAGATTTTGGATTAGCAAAGCAGATATCTCAACTAGACGAACTTAAGTCCTGCAAGGGAAGTGCCTACTGGATGGCGCCAGAGGTCATTGACCCAAAGAAGACGTACAGTTTTCCAGCAGACATATGGAGTCTGGGATGCACAGTACTCGAAATGGAGACAAGGCGTCCTCCTTTTGGTGACATGGAATGGCATCGTGCATTATGGAAGGTTGGGCATGGGGAAGCACCTCCAATACCTGCGACTCTTTCTGAAGATTCTAAGGATTTCATCAGAAGGTGCCTAGAAGTTAATCCTGCCAGACGGCCCACAGCCAGGCAGCTTTTGGATCATCCTTTTGTGCAGTATGTAGGTGGAACAGGCCTTACACCTAGAAATTCCTTCACTCAGGACACTCCGGAGCTTCCAATCATTGCAGAAGACAAGACTTTTGATGGTTGACCATGAGTCCTGAATGTGCCAAGTTATGGCCTACAGGTTTTTTTTTTCCAGAGGTGATTGAGATGTGTGAAGTGCTGTGGTTAATGTGTTCGATCATGCTTTGGAGACGAGCTCTGCAAAAGTACAAGTGCACAGGCCGTAGTCGCAATGCCGAAGTCTGCATTGGATAGGAGGAAGATGCATGAGCTAAAGGAGCAGCAGCATCTCTTCACAGAGACCATTTGTACAAAAGCTTGCAGGCCTGTTCTGCAAGAGATGAAACTGTGGAAACAGCATGACTTTGTACTTAAGCCTGGTCATTTATCCAAGCCCAATTCATGGTAATTTCGGACTTATAGCTGATGCTCTGATTGCTTTTATGGAGGTCTTGTTTACCAAATAAGATGACCGTACATCTTTCTAGATTCTTTAAGGAACTATCGAGCCATAATACTTGGTCATATATGTATTCAGAAGGATTTGAGATGCGACTTTTGTACGAGTGTTGTTAGGGAATGAGAGCATTTCTCTTTGGCAAACCTGTGCCTTGTATAGAATTGATTGAGCAGTAAATGGCTCCGCCTGGAAAATCCATCAACTGGGCAGAAGTGAGGCAGTACGGATGTCTTGTTTGCACATTCTGTATATCTGGTTTTCTGTGACAAAGCTGCACGATTTCTCTTGTAATATTTTTGATTGGCCAAGAGCACTCCAAGCAGTTTATGGTAGTAACGGTGTGAAGAAGAAGTTGGAGATTAATACGGCCAAGTCACGCTGTGGAAAGCGGTTATCAGTCATTAAGCTTAACAGCCGGTAGTTCCTAGTTCAGGAAACTTGCAATAATGAGAACTGTTCATAACAGCATGCAGACATTTTTTAGAGACATGGCAATGTAATTTTGGTATTCTTTCTTTAAATCTCTACTTCATAACGCTGCTGCGAAGGTCATGTGCATAGCAGTTCCAAAGATGCATGGTATGAAGGCAAGAGAGACTCGACATTCAATCTGTATGAAAAGAGTGTATCTATTTCTAGCACCTTTTATTGTATTCTATTGTATATACCAGCTCTTGCAATAGTTGGCTACTACTTACTAGCAGCAATACAATCAGCAATGTCTTTCTTTGGAAAAAATAAATGTGTATCTTTGGAGTTGACATGCATCTTTAGAGTTTACAGGTTTACAGATGAAAGACTCCTTCAGAGACCCCCTCTACTTCAGCTCATCCACCTTCTTCTAGCTTGGAGCCGGATCAATAGCAAATTAGCACAAGTGCAAATGGAGATAGTTCCTCATATGAGAAACTATTTCTTACTTTATTAGAAGCCTATAGTGTGCCTCCTGTTAAGGAGTTGAGAAGCTCTGCCAAGAATAAATGTAAAATATAGTTCTGTATTTAGGCAATCCCTTAAGAGCTTCGAGCCCGGGATTGCCTTTTTTTCTTTTGTTGGTGATACCTCTTGGTTACTTTCCTCTACATTGTAACCCCTTTTGTGTATTTTACCTTGGGTCTTGCCATCGAGGTAAAATAGAATGATTTATTCTCTTGGGAAAAAAAAATTACATTGATATAACGTATCCAATACCCTACTGTTCAGAGTCAAATTACCTACGAACCAAGTGCAAGACAAAATAGGTCAAGGGTGATGAAGAACCTTTCACAAGCCCCAAGTGTGATACATTTTAGCAATAATTAGACCAAGACAATCAAGTTCAGGAATTTGGAAAACATTCAAGTAACATAATAAGGCTAAAAGACACAACTGCTTGTCACACCCGTAATTTCCTTAGCTAAATCCATAAAAAATAACAAACCTGGAAGTATTTAAAAACATCTCAGATCTGTGACCTACATCAGCTTCTTACAACACTCAGCATGCAACTATCAACAGAATCTCAGAGTCATCAGCTGTGGTTAACACCGTCTAGAAATAAAGGTACTCCAATGACACTTATAGATCATATAGTAAAACTTTTCGTATTTGAGATTTGCTATTAAAGTAACCCGATCTAAACAGAAAGGGAGATTTACAGACTGCTTGCCACACCCATATAGGGCTAACTTCAGCCATAAACGGCAAAATTACAAGCATACATGGCATAGCAGTGTATGTTCGCACACAAAATGCAGATATGATAGCCTTCTATGCCAAAATCTTACATCAAAGCATGCCAATGCTCTGAAAAGAAAGTTTTGGTCTATCAACCACGACTCTTATGTGCACAGCAAAAAAATAAATCATAAATAAAGTATATCCAGGCGTAAGTACTTATACCGACCTTATTTAAGAGCCTTCCAATCGAGCCTTAAGCCGGACATGAAACAGAAAAAAAAAGATCCCGATGCCCTTCTTACCTGAAGATAATATCCCATAGCGGCATGCTATATTTAATCAAAGTGTTCTTGCCTTGCTTCAGACATATCAGTATCCCCATCTTCATCTGGCCTGGAACCCGTAAGAACTTGCCCGCTAGCCTCATTAGCCAACCCACCAGGACCACTAGAAACTGTTTCCGCACTATTGGATTGGCTAGTATTCCCAGCTTGTGCATTGGCAGGCGACATGAAGGTGGTCGTAATCCCTCTAAACCATGGAAACGTGATTGAGAACCGTCTTCCCGTTCCTTGCACCCCCGCTTCCCTGCCCCTGGTGTTACTATCTTGTGAGACAAGGGTCTGTTGTTCCTGCTGCTGCGAGGCAGCATTGGTGTCTGTAGGAAACCTCCCCACAAGGAACTGCCTTGGAATACCTTGGATCGAAAAACCACCAGGCCCTCTTTCACCCCCTCCCTGCACACCTCTAGCCATTGCAACATCAGAAGTGGAGCCTGTTCTTTGTCCGCGGGTACACCCTTGCTCGTAGTCAGGATCATCTGTGGGCAATTCAAATCGACACACAGGACAAGAATTGTGCTGTGCCAACCATGGCAAAATACAATCTTGGTGATATAAATGCTTACATGGCATCTGCTTTGCTTTGGAATGCAGTTCAAACTCATCCTTGCACACTGCACATTGGGCAGCATCCGTTCCTAGGTGTTCTTCAGATATAGCAATAGTAGGCAAGGCTTCAACAGCTGATTTTGAAGCCGGGGGAGCTCCATATCTGTTTGGGTCATTTTCAGCAAGCTGTTGGATGAGCTGTTCAAAACCTGGTCCAATGAAATAATCACCAATGCTACCAGGCAAGCGGTGAGGTGTGCCACCTGCAACATTGTCAAGCAGAAATTCACCATTGCGCATGTTGAGGAGGTTTTGTATTCGACCTCGTAGGAGAAGTATGGGATTGATGTTTCCAGGTTCAGTGCCGAGGCTTATATGAGCCCCTGAGCCTCCACCTTCTGCTGCCTGAACCATGTGCATGTGGGTGAGGAAAGATGACATGTCTTCGAGCAGCTGGCCAAAGCCGGGAGCCATGTAGTCCCATGCATTCAAATCAATGCCGGGGGCATTGGTGGCCAAAGGAGGCGCATCGGCTGCAGGGCCGGACCATTCCTCAAGGAAATCACCATTACAAGTTGGGCAGATCAACTCCCCGGGATTCGTGGGTCGTACAGCCTGGGAGCACTGATGGCAATAGTAAAGCTGCCCAATATCAGATGCCATAGAGAAGCGGCGGAGTTGGTCGCCGGTTGCAAGCTACTGAATCGATCTGTACATAGAGATGTGGCAGTTTACTACCCTAACCCTAGGAAAGACCGTCGTGTGCAAGTGATCCTGAAAAATGTGAGAGAGACACTTGCCTTATCGATCCTTAGGAGTAGTGGGAAGGGTTATTCTGTGCAGAAACGAGAGTTTTGGCGATGGCGGGTGTAGAAAGCAGCGATCCAAGGCCCCGCGTGAGGGGCTAGGAGGCACCCATTTCGTGAAACGACATATGCTCGCCATGGCTCATTATAATCCAGTACAGCTTTATGGGCGCGTGCACGTGTGATACGTTGGCCACGCCTTTGGCCGCCCACGCCTCTGCTCTCGATTTGATTTTATCTGATTTCCTCCTAGGCTAGCCTCTTT

**PBS1-LIKE 39 (PBL39)**:

AATTGATCCAACATATTTCTGCCATGCTATGATGTGTACTTGGTGAAATCTCCAATCTAGAAAAATATATTTCCCTCTTAAAATACCTCATTCAAATGGTTCCACGTAGTGGCCCCGGTAGAAATCACAGCAGAGGTTATTGTTTTGCATCTAAAGTGAAGGCACTTATACCTCTATGACAGCTATCGACATAACCTTTACATTGAATTATAGACATTGTTTGTGATCTCCATAGAGTCTATGCCTCACTGTAAGAATAACATATGGGTGTAATAATCTGGGATTCTAGTTTGTTCACACACACATCAAAATTATCTATGCTTATGAGTCATAACTGGTTCATAAACATCTCACATCAGCTCTTTGACAGGGCCATTCAAGCAGGAATAGGTTCTGAAGCCCACTCTCTTTTTCTGACATCTTCACCAAAAAACCCCCTTTTCTTTGAAGGCTTCTTGTGCTTGGATTTCGAACCACTCTTTGCCTCAACACTCAGGTGTGCATCACTGGCATGGCTCTCGCCTCCCTCGAGTATATGTTTCAAACTCCCCACCACAACACTCATTGATGGGCGTGCTTTTGGAGCTCTATGAGAACAATGGAGAGCTAAATCCACTATGTGTTGTGCTTGAATCAAAGGGTAATCTCCTTCAAGTCTGGGATCCATGATAGCGTACACTTTCTTCTTACCATGCAGGTAAGGTCTGACCCACTCAAGCAGATGTTGCTCGCTTTTTGGACGATTTCGATCCACAGACCTCCGTCCTGTAAGAACTTCAAATAGCACCACCCCAAAGCTCCAAACATCACTTTTCAATGTAAGACGGCCTGTCTGTACGTATTCTGGAGCAGCATAGCCTATTGTCCCAACCACCGAAGTTGAAACATGGCTTGCCCCTATCTCTGGACCCTGTCTGGCTAGCCCAAAATCCGACAATTTTGGAGTCCAGTCGTCGGCTAGCAAGACATTCGACGTTTTAAAATCACGAAATATGACCTGAATACCTTCTATCTCTTCATGCAAGTAAGCCAAACCGTGGGCAGCACCAAGCATGATGTGAAGCCGCTGTTCCCACGTGAGAACAGGATATTCTGTGCGAAAAAGATAATCCTCGAGGCTTCCCCTTACCATATACTCATACACAAGCAGGCGCTGTATTCCCCTCTCATCATCTTCTGCACAATATCCCACCAGCTTCACCAAGTTGGGATGCTCTACCAGCCCAAGAAAGCGAACTTCTGCCAGCCACTCCTTGTGCCCCTGTTGGCCATTTCTATTCAAAAGCTTCACAGCCACTTCCTTTGAAGATTGTGGGTCGTTTGGATTGCTGCCCTTGAGAGTGCCTTTGTAGACACACCCGAACCCACCCTCGCCCAGTAGATTGGAACGCGCGAAGTTGCCAGCAACCGCCCGTAGCTCCGCATACGTGAACACCTGAAGCTCCAATGCCGTCCCCGACTTGCTCTGCTCCAAAGACGAAGTCTCTTCGCCTTGGCGCATTTGCTTTAATTGAGACGATGCTTGTTGCTTTGCATGTGATGATGAAGAAGGCGATTGTTTCACTGCATTGGCTTGTAACGAGGAAGAAGGAGGGGCTGATTTTGATTTTGAGGATCTCTTGCCCTCTGAAGATGAGCCATTGCTGAAGAAGCCCCAGGAAAATCTGTGCTTTGAATCAGGCCCGCATGATTTAGAAGAGAAAGGAAAACACCTCATGGCACGAGCCAAGAGAGAGAGAGAGAGAGAGA

**TRANSCRIPTION FACTOR TGA2 (TGA2):**

TTTTTTTTTAGGAGAAGCAAGCTTAGCATTAGGAGAAAAACAAAGAGCTACATGTAGAGAGCAAGGAGCTACTAATTTGATGACAAATTCTCCTACAAATGGCATGGTGAAGCCTTTTCAAGAGCAAAGATTTTTGTGCTAGTATGCCCTTGCAAAGACTCTTAATCATAAGCATAGAGAGAGAAGAAATAGACATGGTGAGAGCTTAGCATTTAGATGAGATATATGCATAGGAATTTTGGGGGGGGAGGGGCAAAGAGAGTGATGATGGTGTCTAATTCAAGATCTAAATGAGTGCTAAATTACTCTAAGGTGAAGAGAAAGGAAGCTTTACTAGAAGAGAGTGGGAGAATAGTGAGAGTTAAGCATGGAGAGCTTCATGGCAAGAGAGGGAGGAGAGTGGCACTTCTTCTTCATAGTGGGCAACATGAGAGGCTCAAGAGGGAGAAGGGAGAGGGCAAGAAAAGAGGGTAAGAAGAAGGCCCCAAACTCGTGGAAGTGGGGGGTATTTATACAAAAAGATGGAGGGGTGAGGTGTGAGTGAGTGTGTGTAGTGAGGATTAAATTGAGCACATGGTGTGGGTGGATTGCAAGGGTAGGGTGAGGTGGAAGGACTAATGTGATGTGATTAGGGTGAGGTGGAGGGACTAGCCATGTCATGGCTAGAGAGTGAAAATAATTACAACATTGGCTATGCAAAGCCATGCATGGGCATTCCATCTACAAGATGTGATGATGTGATAATTACAAGGGGTGATTTTTGTTGTCTTCACTCACTGCTGGCGTCAATACTTTGCTCCCGCTGCTGCCACTAAGGGTGGATGCACACGCTTTATGTACAACTACTCATGCCACGAAGCTTTATAACATTTCTCATCATATTGAAGATGTATGCATATGGTGTGAGAATGTGAGTGCCCAAGTATGCTTGCAAGAGTGCACATAAGTGTATGTGCCCGTGTGTGCATGAACTTGCAAGTGCCAAGTGTATTCATGTTTCAAATAAAAATTATTATGGGTGCATGGTGGCATGCAATAGTATGCATGTAAACATGTCTATGGGTGTGCATGCTTAAAACACAAAGAAATGTGTGGTCGACCTTGCTGGGATGTCGTTCCAAAAGATCAAAGAGGGGTTAGTGGAAATGGGTGTGTGTACATGTGTATGCTTGGCATGTGTGTGATTAAAGGAAGAGAATATGTGTGCCGTGTGTACACATATTCATGTATGGTTGCGTGTATGTTCATATTCATGCGGGGGTGTGTGTGCATATAAGTATGTGCATGTACTTGTGCACATGCGTGTGTGTTTGTAGACGAGCAACTAGTGTGTGTGTGTGTTTGTGTGTGTAGACCCATTCCATGTCACTTTGGGCTTTCTGCCTCTGCGCGTGATAGGGTGTTTGCGGTGTGGATCAAGTTGCGGGCGCTACAGCTGGTGATTCAAGCTGCCATTGATCTTCTTTTCATGTTTTTGCTTGTATAGCACTCTCAGGACTGAGGTGAAGCAGTCCAATGGCTGACAGCAGGCACAATCCCGGGCTTCCGAGTGTCCTTGGTCACGGCATGCAGAGCTTTCTCCAACCATCCGTAGAATTCAGAGACACCAAGGGTTCAGATATAGTTAGCCTTGCAAAGATCAATCAGCCTGTGAATGCAGGCTTGAATGAAGGAGTAAGCTTTTTTGACAAAGGTTTCTTTCCAACACCTTATGTGGGCAATTCTAACGCACTGCCTGCCATTCCTCTTCAGTACAACTGTCGGCAAGTGGACAAACAGCAAGCCTCTCACTCGAGCTCCTTATCGGAAGCCAGCTCAGAGTCAGAATCTGCACAGAACCTGAACAGACATATGGGCTTTGATCCACCACCAGGTCGGCCCTTCCAAAACTGGGCTGATCGTAGTCCTAGAGGAGAGACTTCCACGAGTATGGAAGGAGATGCCAAAGAAAATGACAAAATGGAGAATAGGGTAGTGGCAGCTTCTGGTTCTTCTGACCATGATCGTGGGAAGATTACAGATTCTAAGGTTCTGCGTCGCTTGGCACAAAATCGTGAAGCAGCTCGTAAAAGCCGCTTGAGGAAGAAGGCTTATGTTCAACAGCTGGAGTCGAGCCGTGTGAAGCTCAATCAAATGGAGCAAGAGCTTGATCGAGTGAGGCAACAAGGAGTGTTCATGGGAGGTTCCTCTGGAGATCACAGCCATCCAGGTGTTTTTGCCTTTGATATGGACCATGGACGGTGGGTCGAGGAGCAAAATCGGCAAATATCTGAGCTACGGTCTGCTTTAGCTGCACATGTGACTGATAATGATTTACGGGTGCTTGTGGACAATGGTGTTACACATTACGATGAGCTTTTTCGACTGAAAGGAGTGGCTGTGAAAGCAGATGTGTTTCATATCATGTCTGGCATGTGGAAGACGCCAGCTGAGCGTTGCTTCATGTGGATGGGTGGTTTTCGCCCTTCCGAACTTTTGAAGATCTTAAGTCCGCAGCTGGAGCCACTTACGGAGCAGCAGCTTATAGCAGTGTGCAATCTGCAGCAATCCTCACAACAGGCGGAGGATGCTCTATCTCAGGGTATGGAAGCTCTACAGCAAAATCTGGCTGACACGTTGGCTGGTGGGTCACTTGGAAATTCGAATGTGGCAAATTATATGGGACAGATGGCCATGGCTATGGGCAAACTCGGCACGCTGGAGAATTTTGTGCGCCAGGCTGATCACCTTCGGCAGCAGACTCTTCAGCAAATGAACCGGATTCTTACTACACGCCAGGCTGCAAGGGGTCTTCTTGCCATGGGGGACTATTTCTCACGCCTTCGGGCTTTAAGCTCTTTGTGGGTAGCCAGGCCACGAGAATGAGAAACTTACAGGTCTGTGGTGAAGTTTAAGAGGCTAGTGGGTTTCTGGTTTCTGAAATGTTGGGAAGGAAGTCATGGGTCACAAGATCAAGGGTTGTCTATAGATGTAGAATGGGGTTCATGTAGACTACAGGGGGGATATTATGCTGGTTGGTCATGTTCATAATATTGGATATATGTTACATGGGGCTGTAAATGAACTGAAGTTACTGCAAATCCATGAAGGGCAAAATTTGTATCCAAGAAACTGCATTATTGTAGTGTAGTACTTGCCCATCCCTGTTGGTTGTCTCAGTGTAACCCTGAGCACTTGTACTACAATGTGCTTCAGTAAACATGCTAGGAGACTATAAGACTTCATCTAACATAAAGAGGTGCCCAGGTTTAGACGAAACAGACATGCTGTCTGTTTCCTCGATTGCATGATTCTTATGACTTCTATATTGAGCATTTGTTGTACCTTTATCGGAGAGTTTTCGGACTCTTTGGTGGTAACAGGCCTTGTACTTACTAGTTTCATGGGTGCCAGTCTAAAAGGTCAGCTACCATTACATTCATGGATGATTTGCCCAATGAATTTCGCCTCCATCATGGTCACTTGTCTTGCAGATCCTTGGTTGCATGATTCTTATTTCTACTGGAGAGTTTTTTTTGACTCTTTTGAGCAAAAAAACCTTGTTTTTGTGGGTGCCAGTCTAAAAGGTCAGCTACCATTACATTCATGGATGATTTGCCCAATGAATTTCGCCTCCATCATGGTCACTTGTCTTGCAGATCCTTGGTTGCATGATTCTTATTTCTACTGGAGAGTTTTTTTTGACTCTTTTGAGCAAAAAAACCTTGTTTTTG

**FLAVIN-DEPENDENT MONOOXYGENASE 1 (FMO1):**

GTGGCTAAGGATAAGAAAAGAAAGTGAAGAGAAGGGTTAGCAAAAGTTAAGGAGAGAATTGAGTGCTAATCCTTTTCATCTTTTTCCTTGTCAATTTTTCCCATCTTAAGTCATTCATTTACATAGGCTACAAAAGCTACATGTTTCAAGTTGGTAGCATGTCCATTTAAGAAAAAAGCTATAAATTTGAGACCTCTGAGAACAACATAATTAGGCATTGCACTGTCATGTGAACACCATTACAGTCTTGCTTGTGTGAATGATTTCTTCATCAGCACAAACTTGTTTTATGACTAACTCGCTATTGCCGAACTTGATGAAGATACATCTCAGAAGCGTAATGTACATCGGATGGGTTATAGAGCTGCAACAAGCAAAGAATGTCTGCATACTCATAGATGGCCAAAATACATCGATAAACGGGTAATATATACCAGATAGGTTACAAAACTGCAATGCACTCATTTAGCAAAGGATGAGGGCATATTGATAGTGAATATGCACAATTACATAGAATGTGTAGACACCACTTGTAAGCTGTTTTAGTGAGCCTCATCTTTGTAATCCATATTACTATATGGAGCAAGAAACTCCTGAAGGAAATTGTTTTTCCTTTTCGGATTCCATCCGAGGTCCTGGCACACTTGATCTGAATGCCATATGCTAACTGCACCTATACAGGATTTGTGATAAAAGGGGGTTGAATTCTTCATGTAGTCTATCCACTTCTGTGTGATTTTTTCCATTTTGTCAATGCTTGGAAGAGTGATTTTTTGAGAAAGCAAGTGAGCCAGCCATCGTGCGCCCATTTCAGACGAATGAAGGTTCGAAAGGTTCTCATGAAACCCCATGATTGCCAAGTCGGGAATCCGAGGATGGATCACACCTCTGTAAAGAGGTAAAACATTTCCCAGACCTTGCAGTTCCCCGCATTCTCGCTGTAATAGTGATCTGAGCTTTTTTTCCCCATCGTATCCTGTGCCTAGAATTATAACATCAGCTTCTATGTTAGTTCCATCTGCCAGTACCACTCCAGTTTTTGAAAAGCTCCAATTTGCTGCCTTGTGCAAATTGATGTGGCCAGTTTTCACTCGTTTGAAAAACGCTTTGGGCAATAGAGGAACCTGGCAAGAAGCAAACTGCTCAAAAAAGGTCTGGTTTGGTATAAGGCCAAGCTCTCTGAGAGGGAGGTTAAAGAGAAGGTAAAGCTCGACAAACTTAGAGATCGCCCATCGCGCTGGTGTCAAGAGAGTTGAAATGACACTGAGCAGAAACCCCTGATCAGGCTTTGGAATCATCAATTCAGAAAGCCTCGTTGCATAAAGGTATGACAAAGGCAAGCCGAAAGGTTTGTGATGAGGGAGCATCCAATGAGCACTTCTGAATATTACGGTGCACGGGTGACCATGGACACCTTGGTTTGCATTTGAAGCCTCCATGGCAACGTCGAGTGCTGATTTCATCATTCCAACAACCACAACACGCTTGTCTTTGAGCAATTCTCTAGCTGCCCAATCATCCAGCATAGCGTACTCCATGGAATGCAACACTTTGCCTTCGAAAATTTCAGGCCCTTTGCTTGGTGGGAAAATTGGCAGCTTGGGAATGTCTCCATATCGACCAATGCATAATATAAGAAAGTCAAAGTAAAGCCACTGCAAGCACCAAGGATATCAATCTGGTAATCCTTTTCTGAAGCTTCCTGTTTAAAGTGAAATAATACATGCATCACACCCGCTCGTGCACAAGACCCAACAACTGAAAGGGAGCAATGCCCTTCCAATATTGGCCTTTCCTACAAAGGAAAATGTAAAATTATTTGATTTGTGCTCAATAAACTAGAAGTTCAGTAAATATAATTCACATCTCAGAAAATGCATATGTTGCACAAAAATGGAGGGCAATCAGAGCTGCAAAATCCATTAAAAAAAATCTATTTTTCCCTCTTTCACAATTTCAAGACTTCCAATTGCTGAATAATGAAAGAAAAAGTTAACATTCCAGCGAAATCTCTTCTCCATAGAGTTTCGACAAGACTCAAGCAAAGAAGATCCGAAGAAAGGATCCTTCAAGAAGACTGTCACCAGATGGAATAGAAAAAAAACCTTTCCATTGAATAACTAATCTTTCCTTACTAAAAGACATTGGCAGACAAGCAGAAACAGGGTGACAATTGAGTTGCCCTAGAATTGATACGTGATGTTCTCCTGGCGGTTGTGACCGTGAAGAGTGTGTATGCACGGCCATTTAGAGCATTTCCAAAGCTTCAATGTCCTTGCAAATTCCACTACTTGTGCTTCGTAGCGCATTCTGTCCGCTAAAACTCTCTACTTCGAAAACTCTACACATATTTGAAAGTTGGAAACTCCATTCCGCGGCATACTTACAAGTTACATACCACCCTGCCATCATGTAAAGAAGAAAACAAAGCTTCCCTGCATCCTTTGGTTTCCAAATATTCAGGCAAGGTCCCTCGAAATGTAACTAATGTGATGAAGCATCAATTCAATCTAGCCAAATGGCATGGTATCGTCATGTCGAATTTAAACCTGCCCATTGTCTATATAGAAAAAATATGATAGGTTGCAAAGTCTAAAGAAGAGAAGGCCAATCAATGATTGGTGGTCAAGAAAACTACCTCTGTATCATCATCGTGGTTAACATGCGGGGGCCTCTTGTCAAGGGAACCCTTTCGAATGCGTACAGACCAACGAGAACCTTTGCTGATTTGTTCACCATCATAATCAGCATCATCACTTTGATTATCATAATGGTCATGCTGTCGTCGAATGTCGATCACTTTGCAGTTGAATCTGATGCGATCCATGACATGATATTTGCATGCATAGCTATGGAGGTACTCCATCACATCTGTGTGGGAGGGATAATCAGGGGTTCCCGGGGGCCACGAGTAATCCGAGAATTGGTAGGCGTCCCGGGGGGTCTGCAGCTTGGTGGTTGCAAAGGTGTGCCTCCAGACACCGCCGACTCCCGGTTGCGCTTCAAGGACAACTGGCTGGAAGCCATGCTGCAGAAGGTACTTGCATGCACAAATGCCGGTCATGCCGGCGCCTACAACGCAAATGCGAGGGCTCTTTGTGTCATGACAGGAAGGGGGAGAACCCAATTCGGAGATGGGAGGTGGGCTGTGTGTGTATATGTCCATGCTTGAAGTTTAGCGAGGTTTGTCTGTGCTTTTCTGCGTGGGTGAATCCATAGCAGAGACAATGTTGAATGGAGCCTTTGACAAAGGAGGATTTTAGAGGATTTTAAATTCAAATATCCTTTTGTTCACTTTTGTTTGATATTTG

**SUPPRESSOR OF FATTY ACID DESATURASE DEFICIENCY 1 (SFD1):**

GCTGCTTCTTTGGATAGAGTAAAGACCACCTTGCTTCAAACTGTTTAAAGGGGAGCGATGGCTGCATGTCCTCCTCCCCTCTTCCCTCGCCTTCAGCTGCGGGCCATCTACTGTGCTACTTCACGCAGCTTGCCTCGCCCCTTCACTCCCTGCGACCACATTCAACTCCTCTGTCAAACTTCCTCCTCAAGATCCTTCGTTCCTGCCTCCTCAAGCTCCAACGTTTGCATGCACGAGGGCTCTCCTGTTGAAATTAGGGTTTCCGAGCTCTCCTCGAGGGTCATTGAACCGTCCATTTCTTCAGATTTCATAGCACGGGAAGCTTCCTCTGCAGAATGTAGGCTTGATTGCTCAGCTGGATATGAGAGCCAGCGGACTTCTGCAGAATCTAGGGTTGAAGATGTTGCAGATAGAAAGGTCAGCGACACTTTTCTTTCTGATGAGGCATCCACAACCTGCTCCATAACGGACACACATGCTTTATCTTTGGCTGGAGCCAGGTTGCTAGGGGAAGCTTCTCCTTCCAATGAGATATACAGAACAAGCTACTCTATGCCCCAAGAAAAGGCGTCATCATTGCGGGCTTCCTCGGATGCTGTAGAGAGCTCACAGAGTTTGTCATCTGCATTGGCTGCATCTACACAGACTCGAGATCGGAGTAAAGTTGTAAGAACTGCATGGAAGGCCCTTGTGCGATGGTCCAGGTCATGGCAGCTCCTGAATGCGCGCAAGGCCACCAACGTGCTGCAGAAAACAAGAAAGGTTGTCGTGATTGGTGGAGGCTCTTTTGGAACAGCCATGGCTGTATTATTAGCAAGGAATAAAGCTGAGATGCAGGTCAGCTTACTTCTTAGGGATTCCTCTTTATGTGAATCAATAAACAATGATCATATTAACAGGAAATATTTTCCGAAGCACATTTTACCACACAATATTGTTGCAACTATCTATCCACAAGAGGCGTTCAAAGATGCTGAGTATTGCATTCATGCTGTTCCAGTCCAGTACAGTGCAGAATTCTTGAAAAGCATATCAGAACATGTTCCTCCCAGTCTTCCATTCTTTTCAGTCAGCAAAGGTTTGGAGTTATCCTCTTTGGAAATGATGTCACAGGTCATCCCGCGAGCGCTTGGCAATCAGCGGCAGCCTGTAGCAGTAATTTCTGGGCCTTCCTTCTCAATTGAACTTATGGATGAGCTTCCTACTGCAATGGTTGCTGCTTCAAAAGACAAGGAACTTGCAGAAGCTGCACAGCAACTTCTGGCATCTCGATATTTAAGAATTAACACTTCCAGTGATGTTGTTGGTGTGGAAATTGCTGGAGCACTGAAAAATGTGCTTGCAATTGCTGCTGGTATTGTGGAGGGTATGAATCTTGGAAATAATTGCATGGCTGCTCTTGTTGCCCAAGGTTGCTCTGAAATCCGATGGTTGGCTGAAAAGATGGGTGCAAAATCAACAACTCTTGCAGGGTTGTCCGGTACAGGGGATATAATGCTTACATGTTTTGTGAACCTATCAAGAAACCGGAGTGTAGGAGTAAGGCTTGGTTCTGGAGAATCACTAAGTCATATCCTTTCCTCAATGAATCAGGTCGCTGAGGGTGTGGCAACTGCAGGAGCAGTGATTGCCCTTGCTCGTAAGTATCGTGTTCAAATGCCTGTTCTGACGGCAGTGGCACGTATACTTGACAATGAGCTAACACCAAAAAAGGCTGTTCTTGAGCTGATGAGTCTTCCGCAGGTGGAGGAAGTATAGCCAGCTGGTTTGAAAAGAAATGAGTGCTATATGGAAGGGTTGGGTTCTGAAAGTCCTCTGGGTCAAAAGAGACCTGTTGTGCAAGGGGCTGCATCTGTTGTATGAGCTGACCTATGGCTCACTTCTGCGTCACAACATCATTTTGGCAGAAAATCCTGAGAGGAAGATAGGGGACTGTGAGTTACTGACTGACCATACACTCATTTCCAATATAAACCGTTCCACATCAACATCCCTTCAATCAAAACAATCACCGAGGTACTTCAACTGCTGTATCTCCTTGGAGAGGTCTAACACATTTTCTGGTGGATTTTGGTACCACTAGTATCAAGCTTTACAGAAATTTCGCAACTCAGTCTAAGAACCCCTCGTTCTGGAGGTCCAAAATAACTTGTAACTATCAGTGTAACCAAGTTGTGAAAATTTTCTATGTGAGGCTCTGTTGTAACTTGTAAGTGTGCCTTAAAGCATCATCCACTTCTCACCAGTGTAACCAAGATTGGGATGTGAGAGAGCACCAAGATTGGGATGTGAGGCTCTCTGCTCTTCCTTCGACGCGTTATTTATTTCTGTGATGTCATCTTTGCACACGGTTGCAAACATCATAAGCATTTGTCTCCGAAATTGAAAAGAGTATTTACCAAATTGCCATCATTTTTCTTGTACAAAAAGGA

**PBS1-LIKE 34 (PBL34):**

GGTGGCGCCCCGCCCACCGTTACAGGAAGGAGAACTCACAAGCAGGAAAGAGTTGGGAGCCTTCCGAGAATGTATCAACTGTCGAAGACCTGCCGGGGAATCAATTTTCAGAGCCGTGATGCCCCACCTTGATCCCTACGAGCATGTATCGAATGTCGAAGACCTGCAGGGGATCAATTTTCAGAGCCACCTTGATCCCTACGCACCCGATCGCTCTCGCGGAAGCTTTTCTACTACACAAAACAGACTCGGTGAAGCAAAATCGCTGCAGAACTCTCTCTCTTGCTCTCCCTCTCTACACACACACACACACATATACACGCACATTCCATTATGGAGGCAATCTTGCGAGGTGGAGGCAATCTTGCTTACCTCGCTGATTTGCTTCCATAGCCAAAGCTTTGTCTTGTCTGAGTGGTGAACTAGTGTTGTTTTGGGCTGTTGGAGAGCTTTGTGCTGCTTTCTGCATACAGATCTTGCCTTCTGGTAACCCTAATGCTCCTCATTTCTCCTTCAGCTCATCATCAGGCCTTCTTCACAGTGCAGGTTATGGTGGGTTGGCTTTGATCCCACATGGGGTTGGGCGCTGCAGGGGCCCAAGTGCAGGCTTGGGAGGAAGTGCGCAAATCGTCACAGGATGCTAAGCTCGCTAGCAAAGGTGGGGACGCAAACACGCACAAAGGCTGCCACTTTTTGTCTTTTCTATTGGGCTGTGCAAGCTGCTGGTCTTCAAATCCTAACCCTAGAGCTTCCTCAGTAGACCCTCACACCGCTCGACCAAGAAGCTCATGTGCAGGCAACTCTAATAACTCCAAGCAAAATTTCGCAGCCTCATCTGTACCGCCCACGGAGCAGTCCACTAAATCGGACATTTCAATAAAGTCCGATGCCACAGACTCGTCAATCATCTCCAAAACGGAGGAAGAGTTGGTGGCATCCTCAAAGCTCAGGAGGTTTTCTTTCACTGATCTGAAGCTGGCCACAAGGAATTTTCGCCCGGACAGTCTTTTGGGCGAGGGAGGCTTTGGATGTGTCTTCAAGGGCTGGGTGGATGAATATGGCACGGAACCTGTCAAACCAGGAACAGGCCTCACTGTAGCCGTCAAGATACTCAATCCTAACAGTCTACAGGGTCACAATGAATGGCTGGCGGAGGTTTATTTTCTGGGCTTGCTTCACCATCCCAATCTTGTGAAGCTAGTTGGGTTTTGCATGGAGGAAGATCATCGCTTACTGGTGTATGAGTTTATGCAGAGAGGGAGTCTTGAGAATCATTTGTTTCGCCGGGGAGCCCTGCCTCTATCATGGGATACTAGAATGAAGGTTGCCCTTGGAGCTGCGGAAGGTCTTGGTTTCCTTCATGGACGAAGCAAAGCTGTTATCTATCGGGACTTTAAAACATCAAACATTCTTCTGGATGTGGACCATAATGCAAAATTATCTGATTTTGGGCTAGCAAAGGATGGACCAGATGGTGACAAGTCTCATGTCTCGACGAGGGTCATGGGAACATATGGGTATGCAGCTCCTGAATATGTCATGACAGGCCATCTTACGGCGAGAAGTGATGTTTACAGTTTTGGTGTGGTATTGTTAGAGATTGTGACTGGGAGGAAATGTATGGATAAGTCAAGGCCTAGTGGGGAACACAATCTGGTTGAGTGGGCCCGGCCATACTTGAATGAGAAGAAGAGAGTGTTTAGGCTGATTGATCCTCGCCTTGTGCAGTATTCTGAGAAAGGGGCTTACAAAGCAGCTAGTTTGGCTTTTCGATGTCTTAGCCGAGATCCAAGGGCGAGGCCGTTGATGCCTGAAGTAGTACAGACTCTAGTACTGCTCCAGAATATGAAAGACATAACAGAAAACCCTTTCTACCAACAAGCTTACAATTCTGGGTCTGCAAGAAATTTTCACTGATGCAAGGTGTTACAGTTTTGGGAACCCATTTTGTGATAGCAGTGTCTTACCTTTCATACATATGTGACGCAAACCATATGTACATCCTGAATCCTTTTTGACTGCCTCCTTATGCCACATTAGCATTTTCTGTATTTGCAGTTTGTGTTCAATTGTTGGTGTTTTAATAAGCAGCATACTTTCTTGCAGTCTATATGCATGGTCTTGCAGTGGCATTTTGAGAGTCGAGACTTAGGGATGGTTGCATATTCCCATGTCACTCTATAGGCACAGGGGATGGCCATTTTGGACTCTTTTTGTTCCCCGTCCTTGTCCCCGTTTCCTGTACCATATGACTATGATGTCAGGAGGCTTTCATGTTTGTAGTCATTGTAAAATTGGAGTGTGCGTCTCCATTTCAGAAACAAAATCAAATGTGAAGAAGGTTGCCATGCATTGCTATTTGACAGGTATTGACAGTATCTTGTTGCTACTCGTTGCTTCTCAGCTCCTTGGAGTTGCCAAGGTGCAGCTAAAATGTTCTCACCAAGGATCTTGTCTTGAAAGCCTTGGGAGTTATTTGTCAAGTGCAAAGGAACATGTGATGTAGCCGACACATGAGCGGGCTAAATCTCAGGAATGCTCGTGTAATTTTTCCTATGCTGACAACTAGCAGCAATGACATACCACTAAACAGGATTATTGCATATCACAGAGCTTTATTGTACATGGACGTGTACTTGAGAGTAAACAGCCTCAGCTGAAGATGGCATTCTTATCCTGTGAGCAGTGGCAGCCTTTATGCCTGCCTCAGTGGCAGTTCACACACATCACGAGTCAGAGTCAGAGTCAGAGGGATCTGAAATGTGGCAGAGGGAACTGTCAGATTGCAGCCTGCGCGGAATGAGGGCCCAGTGAGATTAATACTCTCACAACCACAGACCGTACGTGACGAGAAGGAATTCCCAATGCCCATGCTCCTCTCCAGAAGTGCCTGACCGGTGCTGGCTGGGTTGCAATGTTTCTACAATCACAACAAGCTGGCGACAGCTCTTCATTGCAAATTTACACGAGTCACTTCTGATTGATTGTAGAGTCATAGGGTAAAGGGCCCGCCTGTCCTGTCCGACAGTCCGAGTAGCCTTGCTTTACTTAAGGTCTTTTTTTGAGGGCTACTCTCTGTATCTCGGGAGTGTTTGAGTGGGACTTGTAAATAGGCACCCTCAACCCCTTGAAGAAAGTCTAATGAAATTGGATGGCCATAATTCAAATTAGTTTTTGAAATAA

**RESPIRATORY BURST OXIDASE PROTEIN F (RBOH F):**

TTATAACTTACAATTTAACAAAATGTTCTATTAATAAATTTTGATATAAAATATGGCACATGCCATTCAATTGCACAATACAAAGGAAATGACTATCATTATGCTGACAAATGAGGTCATCTGCACTCCAAGACTCTTGAACCATCAATGTTGACTCCAATCGTCGCTCAATCTTGTGACATTTTAACTTGAAATGACAATTGTCCATATCTTCAGCACCCATAATTTTGAATTTAAAGTGAAGCCCCTTCGGTCACCTTCAGAGCGAAGATGGATATCTGAAAGAGAATAAACAAACTGCACAATAGCCACCATGCCTTCTATCTTCTATTTTTTGAACCCAAGGAGCCTTCCTTTCCAATGTTGATCAAGAACACTACGTTTACAAACACGGTGAGGACCTATAGCAATAGGCTACATAAAGATCCAATATCCCTTGTGAAATCGCTCCCCCACTCACTCTTCATTCTCATTCCTTGTGTTGTATAGGACTAATTCGCCAATCTTTTGTTTAGGCATCTTGTGGCCCTGTGTGGACACAATTGAGAGGTGAAGAGAATGGGTATGTGTAGTTTTGAGAAAGAAGAAAGAGAGAAAGTTGTAGTCTTTGGTTGTTTAGGAGAGGGAGAATGGTTTGCAAAGCTAGTTTATTGCAGATATAGAGAGGTTAGGAGGGATTTCTAGAGAGTAAATTGGATAAGAGAGGAAGAGTGTGTTACAAAATCTGGTTTGTTGTAGATATGGAGAGCTTGGGAGAGGGATTGTAGAGAGAGTATTGGGAGGAGGACATGAGGAAGAGTGTAGTCATGGAAAGATTTGGAAAGAGGAAGAGTGAGTTAGAAAAGTAGTTTGGGAGATGAGTAAGAGAAGTGTGCAACCATGGAAAGTCTGGGAGAGAATGAGGAGAGGATGAAAATCTTGGGAGAGGAGGATGAAAAGTTTTGAGAGAGGAGATGGAAAGCTTGAGAGAGGAGGAGGAAAGATGGAAAGCTTGAGTGCTAGTTTGTAGCAGCCACAAAAAGCTTAGTAAAGGGATTTTAGAGAGAGAAAATGATTTTCTCATTATAATATAGATTGGGGGAAGGTTTGAGAGGGAAAAGAGAAAGTTTTGGAAAATTTTAAAATTTTAGGCTTTTTTGACCCACAATATTCTTTTGGGATGGGACTAATCTATATGGAAAAAAGTAGGAGAAGCCTAAAAGCAAGGGACCCTCCTTTGCCCTCTCTTACTAGGGACAAGTAATAAAGAGGGAACCTCCCCTCTACTTTTAATAGGAGCCCAATTGGCACCACCACCCTCTTCGTATTGGTCCATTAGTTCTTAATCAAAAGAGCCCTTTCATAGTCATGATTGATCAAATGATGTGACGATTTTTGATTCCCTAGGTTTCAAATGATGTTTATAAATATGCTGTTGTCCCAATGGTGGGAAAGGTTCCTAGGGAGGTTGTCCAACAGACTATTCCTAGTCAAACGAACTCCTAGTTTTACTTTTGCAAGCATGGCTTGTGTTCAAATGCTTAAGAAGCGTGGTAATTGCTTAAACTCGGATGAAAGATAGAGGCAATTTACAAGAGTCCAATAGTAGAGAGTTTCAAGCTATTGTAGCAAACATTGTATTCCTATCAAGTTCCTTATCAGCCCATAAAGGAGAAGTGCTCCTTGTAGCTTGTAAAGAACAACTTAGGTTGTGTGTTTATTGCCACATTCAAAGTAAGGTTTGTTTGAATGTGTAAGAGTAGGCTCTCTAAGCAATAGTAGCAACTCTATTTAACCTTATTTTGAATGAGAGGGGCAACAAAAGCAATTATTCTATTTCTGAGCCAAGAAAATCAATGTCCAGCTCATATTATTGCTACTTTTTTCCTCCACATTTGTTGTTGGATAGCAACATTTTTGATAAATATCATGTCCAATATTGAGAATGCCTAGCCACATTAGATTGAGAATTGTGAACCAATGAAAATGCCCATTTGAATTTTTAATTTGCCAGCTTCATCATGATTATATGCTTTGCAAAGGCATATGTTAAGAACTATTCAGTTCACCAAATACAATTTTTACATTTGCAAAATTTTTATGATATCTTGTTGAAACTTGAAGATAATTTTTTCATATTGCTGTACACTTTTAATGAGACTCTCGTCGCTTTCATCATATTTGATGATGATTGGAAGCAAGATAATGGCTGTTATAGAATTACAATGTACACAAGCTACATATGCTCCACCCATCCATCTTTGCATTTAGAAGTTCTCTTTGTGAAAATGAAATTTGGTGTTGCCTCTCTTGTTGTATTCTTGCACAAGTCCTCTCAACTCCTTTGCGAGCATTACAGGCCCACAGTAGAACACCCCAACTTTGGACCCTTTATGTTTAGATGCTAGATCTGAGAACACCTTTCTCCAATTGGGTCTTGCAAAATGAGTTCGTACCCGACTTCCCGAAACAACATCCAAGCCATTCTTCGCGTTCTGGAGCGCTTGAACCATGGTGATAAGAGCTGATCTTGCATCCCCCTCCTCGTATACACTTGTCAAGTAGTTGTGCATCTCAATCACCGACTTTGTATCATAATCATGCACTTCATTCATGACCCCTTTGAACCACTCAAAGGACCCTTGTTCTCTGGTGACCCAATAGAAATAAGCGTTCTTTGGGCCAACATGCTTGTGCTTTTTCTCATGATTTGAAGAGTGGTGGCTAGATAAAGCAGTTGATAAGTCATCTTGGGACATATCAGATGTGCTATATTCTTCTTCTGTTTTGATCCAATTGAGCATATCTCTCAATATGCTGATGAAAGGTGTTGCTCCAATGCCAAGGCCAATGAGAAGTAGAATGTCGTAATGCTTATAGTTCTGTGCTGGTGCACCATAAGGTCCATCTATTGATAGTCTGGGAAATCTTGAAGTGCGATCGACAACACTGAGAACTTGGTCAGCTCTTAGGAGCCCACTCCTTCCTTGCTGAGGGGATTGACACACCTTGGAGAAAATGTTCTTCATCTCATTAGTCCAATCTCCTAAAGTCCGTATATGAACGCTTAGAAAATCGTCTCCAGGAGAGGAAGTGATGGAAAAGGGGTGCCATTCAAAAGGGGATATATCTCGGCACTTGACAAACATGTACATTCCACTTTGGTACTTAAAAGATGGCGGTTTTGTCATGTGGACAGCCAGCACATCTCCTGGATATATGGCAGCCTGGGTTTATCATATAACAACAAGCCATTATACACACACCAAGTTTATTCATGTGATTAGTATGGAGCCGTCTGTAAAAAAACCAATTTTTACTTAACCTATCACTTGTTTTTACGCCAACATGTGTGTGTGTGTGATTGAGAGAGAGAGAGAGATGGGTTGCCCACAGATACCGTAATGGGCAGTTTATCCATGGCCATTCCCCCGCAAGGACATGCTGCACATCAGGTGAACTCGCGCATTGCCCCTGCCTCCGGAACCCTCCAAGCCCGCCTTCACAATTTTGTCAAGTATGCTCTCCTTCAACAGCTCGTACCGCCTGGTTCGACAGTTTGTGATTTGTATTGTGGTCGCGGGGTGGACACTGAGAAATGGGCACGCACTCAAATTGGCCGATACATAGGCGTGGATTTTGCAGCATCAACTCTGGAAGAGGCGCGAGATCAATGGAATAACCTCAATAGGCCTTACTACGCAGATTTTTGTGAAGTTGACCCAAGCTCGGCCAATCTGGAAATGGAATTGAGATATATGGGGTTACCTGTGGATGTTGTAACATGCATGGGGCATTTACAGGATTGCTTTTTAAACGAAGAGATGGTTAGGAGGTTATTACAAAATGTTGCTTCAATCTTGAAAGCTGGTGGTTATTTCTTTGGCATTACAGCTGACTCTTCCACCATATGGTCTAAGTTTCAGAAAGCTGTTGAGCACAGGGCGGGGGCGGCGCGCGCCCTTCCTCGTGTACACTCCGAGCATTATGTTATCAGTTTTGAAGATGACAGGTTTACACCGTTTGGGACAAAATACACTCTACAGTTCACAGACGGGGTCAGTTTTGAAAGTCAGCAGCTGATTCATTTTCCAAGTTTCATAAGGCTAGCAGAGGAAGTGGGTTTGGAATACATAGAGCTGCAAAATCTACAGGAGTTTTTTGACGATTATCGTCTACAGTTTCCAGACATCTTGCGTACAACGTGTGCCAATCTAGTGGATCCTAAAGGCAAACTACCACAATTTGCTCATGATATACTAAGCTTATACACGACATTCATTTTCAAAAAGGCAGATTATCTGGATCCAAATTACATGTTAAGCCCTTTTCCACCTCCAGAAGAAGAGCAATTAGAAATTTTGTTTGAAGATGCCTGCAATGGCAACAAGGCATCTCTCCAGGATGATCTTTCACCACCAGACGAACTCAACATTGCACTGAAGAAATTACCAGTGGCATTTCCCCTTCAGACTGAGAACAAAGAAGCTCCTGTTCCCACATCAGAGGAAGATCAAAGAGCTTGTCCAGCAGTTACGCCAGTCGAGGGCAACTTACAGTTGTCTCCATTAGGTGATAGTTGTAAAGAAAGGGCACGTGATCTGAACTTTAGGAGTGACTCAAAGCAGTCTGATGAAGGCAAAGAGGAGGTGAAACTTGATGTTAAAGCTGCTGCTGAGATTGCGATGACAAGCAGAGAGATAGAGGATTCTGATAAAAGGAAGGAGTTGGAAGATGTAATAATGACACAGCCGCCGGGGTTTCCTTGCAAAAGTGGTCTACCTGATGAAGATCACCAAGTCCCTGATAAAGAGTGCAAAGACCCCTTGGAATCCACAGAGGAGCAGATTCCCCCCCCAGAATCTAGCGCACATCCTGTTCTTGACCCAGAAAGTCCACTAGTTGATAATAAACAGCAGCTGGGTGTGGCTGATGAGACTAGTGTGTCACAGCTGCACATTGCTAATGGAAATATATCGATGAAAGAGGCTTCAATTTTAGAAGATGTGGTCATGGAAGAAATCAGAGAAGAAGAACAAGAGCAGAGTTGTTCAGAGAATCAAGCTGTTGAGGGAAAATTATCCAAGCAGTGTATTGTGCATGCTTCCTTATCAAAGAAGCGCACTTCAGGAACCAGGGATGAAACCCCTCCTGAGGATCTCCCCACCCTTGCTGTGAAGGATGCAAGCTTGACAGAACTTGATTTGAGGGAACAGAAAGTAAACGACGAACAGCCAGATGAGATGAAAGAGGTACCTAAGCATGACTCTGAAATTTTGGTGCTCACTGATGTACAGAAGGCAGCGTCACCTGCCACTGGTCTTGACATGGATGGGGATCGGCCTGCCACTGGTTTTGACACGAATGGAGATCGATCTTCGATTAAACGGTCTCTAGGGAGGTTAAATATAGCTCTCAGAGAAGAGAGTGTCAATAAACGCTTGCATTCAGGGCAGCTGCGGTATCATACCAGCAGGCTACCTCATCTTTATCATTTACCGTACTCAGAGGAGAAGCAAGAACAGTCACATGCACCTCGACATAATTCACTTTCTTCTCTCAACCCCAAGGCTTCAAGGGACAGGGTTGGAGATCATAGAAGTGAGCATTTATCCCCTAACCAGAGAGGTGATAGTCACAGAGGGCGCTCTCGAAATGCTCACCATAGCCCCACTTTTGTCCCATTTAACCATTTTGTTCCCCAGACCGAATTCCCTATTTTCCATAGTCCAAATACCCCAGGTATCCTAGGACCAGGTCCTCCACCACCTGTCCTGTTGATAGAGCCCATTTGTCCTTTATGGCCAATCCCTCCACCTCTTGAAAGAAGAGTGTTTCCCAAATCCCGTACACCACGCAAATCTGGGACTGAGCGAGAATTTGGAGGCCGTGGCCGAAGAGAGTCACCAAAAAGACATTGGAAAGATTCCCCACCTCCGGGAACAGAAAGCTAATGGCTGTTGGATGAAGAGAGAGACCTTTCTTCTTAGTCTTATAAACCCTTCTTTGAAGAGAATTCTTTCGGTAGTTTGGGAGTTTTGTGTACTATAATGCCTTTTCCTTAGTTAAACCTACAAAGTGTGATGGGTAGGAGGAAGATAGAAAACAGGTAGCATGAATCCAACATCACACGCAACCCCCATTTTCTATGGGGATTGACAACTTGGTCAGTATTGTGCCAAGTCATTGTGATCTCAATGCAAGAGCCACGAATGTGGTTGAAGCTTGACTACTCAACTCACACAGTCAAAATCATCGATTCTTACTCTGTCTCCAGAGACTGGTGATTTTACAGATAGGATGAATTCGACATCTTTCACAACCTTCATCTGTTTATCGGGTTTGACAACTTGGTCGGCAATGTGCCATAATCATCATGATCTCAATGCAAGAGTCACAGATGGAGTCGAAGCTTGACAACTTGATTCTTCAAAACTATGGCTTCTCACTCTGTCGAGAGACTGTGGTTTCACATTTCTTTGTGAGAAACGAAGCACTCACCTTAACAGCTTTGACTTCAAAGCTTCTACACTCAGGAAGGACGTAGAAAAGATGATGGTTGTAG

**E3 UBIQUITIN-PROTEIN LIGASE NLA (SYG1):**

GAGTGGGTGAGGTCACGAGTGTAGTTTAACTAAGAGTTTACTTTCGAATTTGTATACGCGCCGCCGCCCGGGTCTCTCGTTCTCTCTCAACTTCCTCAAGTAAGCCGCTTGCGAAGTTGCATTGTGCTTCTTCTTCACAAAATATATGACGAATGAAATTCAACAAGCAGTACTCGGCGTTTTTAGAGACGCAGCACAACCTGCTGCCGGGGGTCTCTCATGTAGAGTTTAAGCGTCTCAAGAAGCTCCTCAAGCACTGCCCACATCACGATTCTGCCTCCTCATCAAGTAACCGCTGTCCAGAATGTGATCAAACATTTTTTCCTCTCTTGATGAAGGAGGTGGCTGATGTGCTTGGATGCTTTAATGCTAGAGTTCAGCGCCTCTTGGAGATTCATCTTGCTTCAGGATTTCACAAGTATGTGCTTCGGTTGAAGTATAGGTCTTCTTTTGGCAATCGCTCTGAGATGATTTCTGAGGGGCAAAAACTTGCAAGCTATGTGTCCATGAATGCCCTTGCCATAAAAAAAATCTTGAAGAAATATGATAAGGTTCATTTTTCCAAAAATGGACAAGCTTTCAGAAGCCAGCTTCAAACAAGGCATGTTGAATTGCTTCAGTCACCATGGCTGGTGGAACTGATAGCCTTCCATTTAAACTTGAAAGACAAGGCTCTTTCAGATACGGAAATGTCTTCTCGTTGTTTGTATGACTTTGAGAACTCAACGCTATCATTCATGTTATCTAGTTCGATCAAGCTAGATATTGATCTCACTTGCTCGATATGCCTGGACATGGTGTTTGATCCTGTATCTCTAAGGTGTGGCCACATGTTTTGCAATGGTTGTGCTTGTTCTGCCGCGTCAGTTTCAACAGTGGAAGGCTTACAAAATGCAGCTGAAAGTGCGAAGTGCCCTCTCTGTCGAGAGGAAAGAATGTATGGAAGTGCTGTCCGCTTGATAGAGCTAAGCACATTAATCAGTACAAGGTGCAAGGATTTTTGGGAAAAAAGGCTTCAGATGGAGCGTGCAGATCGACTTGAGCAAGCTAAGAAACATTGGGATGAGCAATGCCGGACTGCGCTGGGGTTGTGATTTTTGTTTTTGTTCATGTACAGCCAGTGAAGCATTACAGCCAGTGAATCATTATCTAAGTAGAGTGCTTTGTATCCCACAACGTGTACTAATGATTTGGGAGTTTTTCAATGTTATTATTTTCACTCACAATCACTCCAAAAAGCTTAAGCATTGAAAGTTGAATTTGGTTAAGGGGAAGGAGCTGATATAAACCACTCCCAAAGTAAAGGATTTAAATTGTTTTCTTTTCATGATTAAAGGTAATTGCATTGTGCTCCCTAGTGCTAAAAAATTTTGTAGTTTGCTCCAAGGCTCTCATAACTTTCCACCTTGGGAGGAGC

**GREENING AFTER EXTENDED DARKNESS 1 (GED1):**

TAAAATCCCCACGTAAGCTACTGTAATAATCTTACAATTCGAGGCATGACAGGAATGCACAGCTAGCAATCACAGAGATGTTAAAAGATCGTATGGCGCAATGCTCCCAGCAATTTGCTGACCAAGCCAAATGCATTAAGTTTTCTGAGTATACTCAATGATTTTTCGCACTAAACAGTGTCAATACCTGCCCAGATCATTCAGAATTGTTAACTGCATGCAGACTCAGCTATAAGCACCTGTATCTCTCTTATTCAGCATTTCGAAAAAGGACCTTTTCTAGAAAATTATGCCAGAATACTCAATATCAAATTTCATCCCTGAATGTACATCAAATATATTGTACAAAGGTTCCACAAGCAATATCTCTCATCAGATTACTTTCGGCTTACATGACCGTCAAGATTATATACAGGAAAAGTGCTTACAAAAAGTGATGACATAAAATACAGGTGGGACCTCTTTCAGGAACCAAATTTTAAAAGCATACTGCTCTTTTGGAAGGTCCTTCGTCAAAGCTTCATTATCTCTTTCAGCCCCATATGACTGCTTCCCATGTTGTATGAGAGAGAACCAAGGAGCTGTGATCAAGTCCATGTGATACGACCATTGCTGTAAGAGCGGCATAGCGCTCTTCATTCAAGTGCAAAGGCTTTCCTCTAAGCATGTCAATGTCCTCCTCGCCAAAAGAATCTAGGTATGGTGAAGGCCAGTAGGCCTGCCTAGCACACCTCTGAAGAAGAATATTGGTTCTTTTTATCATCAAGAAGACTCCCACACCACCATTGCAAGAAATTGCATGCCTGAAGCATTCACTTTGCCTATTTTTACTGCAGCACGAGCGGCGACCAAGACCACAACAAAGAGTCCCACAAAGGAGGCACAAAGCAGGCTGTTCAGGTACCAACGAGCATACGGGACATCGTTCTCTTATGTAACGTTGTAACAATGTCTGATATAAAGGTGGAAGATCCATGAGCTTGAAAGGAGCTGCCATGGCTGGCCTGAGTACAGACAGAAGCTTCCTTGGCCCCAATTCCTTGGACAAGTGAGAAGACCATTTCAAAACAACTTCTTGGTGATCTTCCCTCTCCAGAATAGTTTGCAATGAAGGAAGCATAAACTTTCGCTCCAGTACTTCCAGCTCTTTCAAATCCTCAAAAGCATCAGAAACAGCCTGCCTCTTTTCATTGTCGATTAAACTAACAGCATATTCTGTTAATGTTTCACAGTTAGTCTCCTCAAAAGTTAAATTGCAAAGAAGCTTTCCAAGCAGTGCACATCTTCGTAGGAAGGGCAAAGTAAACCTCCGTATTGCTAGCAATACTGGCCCTGATAAATCAACTGAAGCCTGATCACTAAGCGGTTGAAAAAGTGGAATGCCTGTCAAGTTTTCCAATATAGTTTGTAAAAATGCACATGCAGAGGAACTGAGCTGTAACTCTGACAGGTTTGGCTGTATTATATCCCCCATCAATACAACAACCTGGATAATGCACACAAGATAGAAAAGATGAACCAGTGAAATGAATGGCTCCACTGAAGATGGGAAAGAAAGAGGAAGGCAGAAGACAATCCAAAAAAGGGAAGAGAAAGGATCATGTGCCAGTATTGGATCGGCTGTTCGTCTCCAAAGACAAATGTCTGCAGCCTCCTTGACATTGTCTATAAATTGCAAAATTTTCAAAACACTCCCTGGCATGAAGGGTTCTGCTATAGAGTCTTTTGATACACCAAAACAAATGGAACCCACAAACAACTGCATACCCCTAGCTCTCAGTAACAGAGACTGACGATTTTGATTCTGTATAGCTTTTGCAGCTCTTAGCAACACTGGTAACACTGACCCTTTGTCAGTTGCCTTCCAAAGCGCTCCTAGGTTGTTTGGATTTGAAAAATGGGAATCTTTCTGAGCTCGACTTGCCAGCTCCACAGATATGATAGTATACCTAAGGACATCCCATAACAGCAAGCTCTGACTTACGCGGTTATTATTGTGCCTGCTGTTGCTCACTTTTCCAGGAAAGTATAAACCATAGAGTCTCCAAAAAAGGCTGTCAAAGGTTGATTTCAACGACTTTGACAACGGCTTTGACACAGCTTTCCTGAAGCCCGATTTCCAGACCAGTTCCTCAGCATTACAAAGCAGATGGATGGCCCGAGGCACATGAGAAACTAGACCCAAAGAAGTCGCACCGAGAGGAGATCGCAGGCCACTTTCAACGACAGTCGCTTGGTCAGCGCTGGGTAAAAAGGTTTCCGCAAGGTAAGGAAGAACAGAGTTGGCGAGTCGTCGACATACAGGGCACAGGAACTCTCCCTGATTAGGATCAACAATTTGGAGCCCTTCAAACAGGGTTCTACTGTGGTACCTCTGCACTAGAGATGAAAGATATCGATCAAGACATTCCTGGTGAACAGCATGGCCACATGAAGACACGTGCACACCAACACTATCATTAGAGCCAACATGATCATGCAGCAATCCTTCCATTGCATTTGTATGTGATCTGGAAGAATTCTGTCTTTGAGAAGTAGCTGAATTTTCAGTCTCTAGTTGTTGTTGTTGTTGTTGGTGGTGCTTGCGGCTTCTGGAAAGTATGTTGGCATACTCAAGAAGAACAGTAATCATAGAACTTTGTCTTGTTTTAGCTTTCTCATCCTCATCCCCTCTCGCTCGATCCTTGCAAGAGAATATTCCAGGGTATTCTAGAAAAGAATTCGGTACTTCACAACTCGTGATTATATTTTCCTCCTCTACAGAATCCTCACTCCCAGAATCAGAGCTTGTTGTTTCCGACATCCTTGCATCTACAGAAGACTGACTGCCTTCAATAGAGTCTGTGAATCCCTCATCGCCAAGAAACTCTAACAAGGCTTCTACTTCAATAGGTCCTTCTCGATCTACCTCCCCATTCAAGGCCATCTGTATCAAGTTCAAAAGTTCAACGCGTATCTGTGATTCCTGTACTTCAACATCTACTACAGATGATCCATTATCAACCTCTCCGTTGAAAGTTAAAGCTTCAACATTCTTAGACCTGTCTCTTCTTTCCCACGAAGGAAGACCTTTTTCAGATATTTCAATCAGCCTTGATCTCTGAACAAATGTCAAGAAAGATGTAGGGCTGTTGGAGCTAGAATCCCTGCAAAGAGCACATGCAACTGCAATCGAACTTTGCTGACATTGTTCATTTTGAGACCTGGATCCATTGGATTGTCTACACCCAGGACTTTTATTTTCAGGTTTCTCTTTACTCATGTCACTGTGTTCTTGTGATAACTGAAAACTAGCGACAAACTTCTCCTGTGCCGCTTTCATTTTTGCCATAACTGCTGCTTGACGTTCCCGACCTAGCACTTTCTTTCTGTCCACTTCAGAAAGGCATGATTGCTCTGTCTTATCAGCCTTATCATCCTGACATGATGACACTCTATGTAAAATCTCTGGTGCAAGTGATTCTATCTCATACATACACCCATTATGTAGCTTTGAAAATTTCTCCAATAAATTTTTTATCAAAGCCCCCATGCTGCACAGATTTGATTCTCCTGCAATCCCTTGAGCTTCAGTGGCACTGTATCTCCGCATTAACAAGACCAACAACGAAAGCAGGCTCTGGCGATTCGATATCTCTGTCTCATCAACACCCGCGAAGTATACCCTTTCTGTTGCATTGCCCAGCAATGGAGGCGGTTCTTGTGTATCTTGCATGGAGCTTGTTGTTTCACCTTGAAAACCACTAGATTTGTATTCCAAGCGCTTTCTATATGCAAACATCGAGCATACATCAAGCCCAAGTGCCAGCAAATGCAAAGCTGTGATTAAGACATTCTCGGGAGCACGAGACTCAGAAAGCTTTTCTGAGAACACAGCATGAAAGAAAATCGAGCGCAAGATATCATGCACTTGTTTGCAAGTAGCTAATCTAGAAAGATTTTGGAAAGGAGTAAAGGGTTTCTGCCATTGGGGTAATTGATTGATAACTGATGATGCCTTGCATGCACGGAAATATCGATCCTCTGCAGACTGCAATTCCCTTGCATTCCAACGTGGATGGTAAAGATCGAGCTCTTTCCAGCATTCTTGCCGTAAGCTAAACCTGCCCTGCTGCATTCCCGATGGCCTGTAGTAAGTCGCAACTAAAAGAAGAGCTTCTTGCCAATGTTTATTTTCCTGAAGACGAGGTGGCAAGGCTTTCATTAGCTGGCTATGGGTCGCATCACTAACAGCAAGCCTCTTTATTAATTCTCGTCGGCAGGCCTGGGTGGCAGAAAAGCCACAAAAGCCGCGCTCGACAACAAGTCGAATTAAAAATGCCAGCATATGTTGAGCCAATGTGGTCTCATATTCATTTGGCCGAAGAAGAACTAATGAAAAATACTCTGTTAAACCAAATCGAGACACTATCCTTTCCACAAATTCTTCAGTAGGAGCCATGACCGCACAGCATTGCAAAAGAAACAAATCAAGATCCAAGCTTCCTTCGCACCACTTGACAGAATTGTAATGTTCGCAAAGGGCTTGAACGGAGTGGCCATTTTTGCGCCACATACCAGCACGAACTTGAGCACAAAAAACCTGAATCCGCAGAGGGTGCTCCATAAGGCAGGATGTAAAAGCAGGCACCTGATAGCGTGGTTGAAACAACTGCAGCAGAAATCCCTTTTCAATGACAAGGTGGTCCTGGCAGTGCCAACTTCTCTTGTTGAACTCCTTCTGATTGAGGTTGGGACTTGCCTCTAAACTCTTTTGCAACGCCACAGCCAAAAATCTGTGCAAAGGAATATGAAAAGATACTTCCTGCTGAGTGACATCAAAATCAATCTCCAACCAGTCCTTATCATTTCGTACAAATCCTGCCATGACACCTGTATTTGGTAATTCTGCACCCATCCACCATTTGATAGATCCATCCTCACCTTCCCTCCTTTCTTGCATATTCACAGGCAATGAATCATTCTCAAGCAACACGCTAACACCACCTAAGTTACCTTGCGCTGCTCTTAACCTTTCTCTATAATTCCCTTCGTGCGTAGAGTCCACCTCTGTGTCAAACACCTCATCCTCTAGATTGTCTAAAGCAGGTCCAGCAAACCTCCACAATGTGGCTCTACCTCCTCTCCCTCTCAAATGCATCCCTCTCCGATGAAGAATTCCACCTTTTGGTGCACTTGACCTACTTGATTTGACTGTTTCTCGATGTGCATCAAGTAAAAACCAGATGTCAAGCACCTTGGCACACTCAGCAATCAACCATAAAAGCACAGCAGGAACGCCTGGCGCTCTTGATGAAACATTGGCAAGGACACCACCTGTGGAAAAGTGGTTGTTACCTTGACCTACCACCCCAACTGTATAATCAACATCCATGTGAGCCAAACTATTGCTTGTTCCCACCTTCCCTGACACATCACTTTCCATGGAAGTTCTACCAATCTTAGTGTGCCCATGGACAACGTCCTCATCAACAGTCCAACCATGGGTCGAACTTTCATCCACAGGAAGTAAAGAATCTATTCTTGCAGCTCCATCAATGAGTAATGGATGGATGCTAGCCATCTGCACTTCTAAAATATATGCAGTAAGCCAGTTTTCTGTCTCTTCCTCTACATGATTTCTTGTCTCCCTTCGCTGTGACTCCATACCCTGAGTAAAAGCTAGTAAATGAAGCCAGGCCGTAAGAAGATCCAATCGTTCTCGAGCAACATAATGTGCCACCTCAGGATGCCTCAGCACAAAACGGATGTCATCAATGATACGGTAGTAAAAGTCTGATACCGCCTGTCCTTTGCTCGCTAATTGCTGTCCATCACCCCCAGAACATGAAAGAAAGGCATCCTTTAGAGCTTCCAGCAGCATGTCCAAAAGTTTCAGATCTAACACCAGCTGTGGAGTAAGAGTCGGAACTGTAAATATCTGCACCGAAAAGCTGCTGAGTGCAGAACTTTCCCTTGCTCGAACTCCGTTAGTTGACTCGAGATTTTGCCTCAAACCGTCTCGAATAAAATGCGGGTATTGACGAATAAATGTTTTTGCAAATTGGCATTTGAAATTTGGGTTTCCCATTAGCTTGAATAAAAGCTCGTGCAGTTTAGTCAGGACTTTCACTGGAAGAAAGCACTCACTCTTAAGGAAAAGGTCAAGTAAGCCAATCTTGTCACACGCAATTAAACTACCAGTACATGTCAATAGGCTCTCACTGCAATTACAGAAGCTAAGCAACATCTCAACCAATGCTAGAGATGTTTGAGTAGCAACTCGCTGTATAACCTGATCACTCAGCTTTGTCAATAGGTCTCCCGAAAAACTTGCCGCTGCTTTAAGTCTTTTCCCCCACTCATTCAGAAGAACCTCCATTACTGGCTCAAAAGTAGACATCAGTGCCTCTGAAAGGGAAGGTGCTTCACCCGTTCCATTGTGTTGAGAGCAAAACCCAGATTCCTTCCATGCAGTGATATCACCACAATCACAGCAACCCCCTCCTGTGTGAATCATTGCGTAATCATGATTCACGTGATTACCAGATTCAAAACAAGAAACACAAATAGCACACGTTGGATCATTTTCACAAGTGCGGCACCTGTAAGCAATGTCATTGGAACCCCATACAGCTCCACAAACACCTCGAACGTGTCCTGTCTCTTGCTGTATCCCTTTCATAATTTTCTCTGGGTCTCCCCCAAACATTAACCATTGTGCCCATCTTACCCCTTCTCTGCAACCTTGTTTATACTCATGCCACGCTAGCTTTGCTTCATCTGACCCTAGTAATTGCGCCTCTTCAGCACCCGGCATCAAGACACTCACAACATCCTGCAATAAAGCCTTATTTTCTTTGACAAACTGTATAAGCCCCATTTCCATGGCATCCAGTTTATGCATTGGCACGCCAAGTTTGCAAAGTCTCTGGGCAACCATGTCGGCTGGAGGGAGGCTATTGGGGGGCAGGGGCGGTGAAGCAGAGGACCCATCACCCTCCATCATTGGAAGATCCAAAAGAGAAGTTGGGTTTTGAAAGCTGAGGAGGGCTACTTCTTCATAAGGACATAGCCAGGCAGGCGTCAAGAAGAAAGCACCCTTTTTGTGCAACTGCAAGCAATGCGCATCTGATAGTGTCGAGAGCTCGAAGAAACGCACTCTCTCTCTCTCTCGCTCTCT

**NRL PROTEIN FOR CHLOROPLAST MOVEMENT1 (NCH1):**

TCTTTACAATTTTAAAAAAAAAAATCAAACATACATTGTTTGAGCTCATCTGCAAAACACTGCATAAATTAAATTCAAGGTGAGCAAATCAACAGTGACATTTTCTAGCTCACCACTATCTATATATATAACACCTGCATTTTTCAGCTCCTTGCTTCTCTCTGTAGGTAGCTTTTTAGCTTGATTTCTTCAAGCTCTTTAGTTTTTTTACTTCTTTAGTTCTTGTACTTCCTCCTCTAGTTGCTTTTATAACTACCTTGCTTTCTCTGCTAGATCTTACCACTGTAGCTCAAATTCTCTATTTATATATTTTCCTTAAATCTCATGCTCATTGATTTACTTCTACAATAATTGCTCACAAGATTTTTTAAAAGAGAGAAATTTTGTTTTCATGCTAAATTTGCACCAAATTATGCTGCATTTTTGAGATCTAAGTGGTTCTGCGTTCGCGCGCGAGAGGTTCATTTTTAAAGAAGGGTGTAGCTATGGCTGTAAATGGCTTGAAAACAGATAAATTTTCTTTTCCCAGGAGAAGCTCCAACTTCATTCCTCCAACCAAAGGGATAAATTTTTCCTCAGATGTGCTAAGTGACATAATTTTGAATGTGGATGGAGTCACTTTCTCTCTTCATAAGTTCCCCCTAATGACAAAAAGTGGGAGGATACGAAGGCTTGTAGGGGAAGAGCTGAAAGAAGGGGACTCCATAGTGAGCAGCCCTCCTGTAATAGAGTTTTCCAACTTTCCAGGAGGCCCCGACACCTTCGAGCTGGCCGCTAAGTTCTGCTATGGCATCAACTTTGAGATTACAACAGCCAAGGTCGCATTCCTGCGCTGTGCTGCCGAGTTCCTTGAGATGACTGATGACTATGGTGAGGACAACCTTGTCACTCGGACCGAAGCCTTCCTTACTGAGGTTGTCTTCAAAAACTTGGACAAATCTGTAGAAGTCTTGCATTCATGTGAGCCCATCCTCCCCTTGGCCGAGGAGGTTAAGATGGTAAACCGATGCATCGATGTCATTGCATCCAAACTCTGCAGTGATCAGCAGAAGCTTTGTAGCCTTCTGCAAATGGAAGGAAATCATTCCGGCCGATGGAGCAATGGTCCTGTGCCTACTAATGAGCAAGACTGCACCTCCCCACCTACAACAAATTGGACTAGAGATGGTGACGCAACTCCTCCAATGCCACCCTCTGCCATAATCAAAGCAAGGGTTGAATGGTGGGCTGAGTATCTTTTACCTCTTCGCATCGATCTCTATCAGCGTGTTTTTGCCGCCATGAAAGCTAGTGGTGTTAGAAATGAGAGTTTGGCTGGCTCCCTTATGCACTATGCGCAGACATCACTCAAAATGCTCGGTAGTCGAAGCAATGGACCCTCTGGAAAGGTATCATCCTCTACGACAACAGCAATTCGTTACAAAACCAAACATCATTTGGACATCAGCTTTGGCACCAACCAGAAATTGATCCAGCAACATGAGCAGAGATCTCTTTTAGAAACCATTGTGAGCTTGCTACCTACAGAAAAAAACACATTCCCCACAAGTTTCTTGCTATCTTTACTGAAGAATGCCATCACATTGGATACAACAGTCGCTTGCCGCCTTGATCTAGAACGCCGCATTGGAGTGCAACTCGAACAAGCTACTGTCGATGACCTCCTTATCCCAGCTGCGTCACCGGGTGCAGGCGAGGCACTGTTTGATGTTGATGTTGTCCAAAGAGTACTTATGAGCTACTTGCAGCAACAAGAGGATAAAGTGAAGCGTGAGCAACATTTTAGCCTCAACACAACAGGAAGTGACTATGAAGATATGTTCAGCTCCCCAAGCTCCCAAAAAGGGATTACTAAGGTTGCTAAAGTGATGGACATGTATTTGGCAGAAATTGCCGCAGACCCAAACCTAAAGGCCTCCAAATTTGCCTCCCTTGCTGAGCAGCTCCCCAGCTATGCCCGAGTAGTCGATGATGGACTCTATCGTGCAATTGACATTTTCCTCAAGGCTCACCCTACTGTATCAGACACGGAGAGGAGGAAGATTTGTAAGGCGTTGGACTGCCAAAAGCTCTCCAAAGAGGCATGCATTCATGCGGCTCAAAATGAGCGTCTACCTGTTCAGGTTGGTATGGTAAAAAGTTTGAGAGCCATCATTTATGCTCTGTAACACACTATTCATGCATTTGCAATCTGTACAGCATCATGCATTCATGTGCCCACATCTTATGCTTTACTTGGGTGTAGATGGTTGTGCAAGTGCTCTACTTTGAACAAATTAGGATGAGAAATGCCATGGGTGGGGGCGATAACCTTCATCATCATCCTTTCTATCACCAGGACAGCGGACCAGGGGGCACTTTTGCAAACCGCAATCCCTTCAAGGCAAGCGGTATGATTTCGCCTAAGCCTGATACGTTCTCCAGTCTTCGGCGAGAGAATAGAGATCTCAAGGAGGAGGTGGCAAGATTGCAGCTTCGTGTCGCAGAGCTCCAAAAAGAGAACATGGCTCTCAAGCATGACACAGAGAAATCAACGGGTGCAAATGGCAATTCAGGTCGATTCTTTTCCTCCGTTTCCCGGCGCCTTGGAAAGATGAACCCCTTCTCATCCAAACCAAAAGAGACTATCAAGCAGTCCGAGGCTCGTCTTCCAGCACGACAAAGACGCCATTCTGTATCATAAATGCGCATATACCGTACAATCCTTTCAGAAAAAATTAAAATGTTAAAGTTCTCTCTGTTTGTTAAAGAGCTGGTTGCCTGCACTAGTTTGTTTGCCCCAAAAGTATTAAATAAGTGTGTACTTATCAAGTTACATCTAGTATTGTGTACATGGTTTACTCATAAAAGTACATCAATTGCCAAGCATTATACCCATCTCCATGCATAAAAACCATGGAGTGCATGTATGCCGCCCTCTTTAACGATTGTATATTCCACTTGATCTTTGTGCAGCAAAAAATCACTTGCCAATAATCTACTGAAAAA

**ORNITHINE-Δ-AMINOTRANSFERASE (Δ-OAT):**

CTCTCTCTCTCTCTCTCTTCACGCGCGCTCATTCACTCACCCACCCGCACTCACACACACACACACACAAGCGCAAGCAGCTATCCTCCGTGGAGGTTTACTCGTGATTTTCCAGGATGTGGCAGCGAGCAAGATGCAGCTTGGTCTCCTCTTCCTCCTTGGTCTCCCGTCTGCGTGTGCGTGGCCTCTCAGCTCTGCCTCAGAGCAACTCCTCCGATAGCCTCTCTTCTCACGACTATATGCAACTCGAAAATACACATAGCGCCCACAACTATCATCCCATACCAACTGTGTTTGCGCGGGGGAAGGACGTGAAGGTGTGGGATCCTGAAGGCCGAGAGTATCTCGACTTCCTTTCTGCCTACTCTGCTGCAAACCAGGGGCACTGCCATCCGAAGGTCATAAAGGCATTGAAAGAACAAGCAGACACTCTGACATTGAGTTCAAGAGCTTTCTACAACAATAAATTTCCTCTTTTTGCCAAAACACTCTCGGACATTTTCAAGTATGACATGGTGCTACCTATGAATACTGGGGCGGAAGGTGTCGAGACTGCACTCAAAGTGGCTCGGAAATGGGGCTATGAGAAAAAAGGCATTCAGTCAGACAAAGCTATCATTATCTCATGTTGTGGTTGCTTTCATGGGCGCACTTTGTCTGTAATCTCGATGAGCTGCGACAATGAAGCTGTTGATGGGTTTGGTCCTCTTATGCAAGGCCTTCTCAAAGTGGAATATGGCAATGCAGATGCGCTTGCAAAACTGCTGGAAGAACATGGAGATAATGTTGCTGGGTTCTTGTTTGAACCAATCCAGGGGGAAGCTGGGGTTATTATACCTCCAGATGGTTATCTGAAAAGAGTTCGCGATTTGTGTTCCAAACACAATGTCCTAATGATCGCTGACGAAATTCAAACCGGCTTAGCACGGACAGGGAAGATGCTGGCATGTGATTGGGAGGATGTACGTCCTGATGTTCTGATTCTGGGTAAGGCACTTGGCGCAGGGGTATTGCCAGTTAGTGCAGTGCTTGCTGACAAGGAGGTCATGCTTTGTATCAAGCCAGGCGAACATGGCAGCACATTTGGTGGCAACCCTTTGGCAAGTGCAGTAAGCATTGCTGCACTAGAGGTTATTCAAGAAGAAAAACTGGCAGAAAAAGCTGCAAAGTTGGGAGAAGAGTTTAGAGGGCTTCTTTCAGAGTTACAAAAATTGCATCCAACTCTTATCAAGGAGGTAAGGGGGAGAGGCTTACTGAATGCAGTTGAAATGGAACCAGCTGGCCTTGGGACTTTATCTGCTTATGATGTGTGCCTAAGCTTGAAGGAGAGAGGCATTTTAGCAAAACCAACGCACGACACAGTAATCAGACTCTCTCCTCCATTGACAATCAGGTCAAGTCATCTGCAGGTTGCAACAAAGGCTTTGCAGGATGTCTTGGAGCAAGATATTCCCCACTCTGCAACGCAAGATGCATCTAAAGCAGGGAAGCATTCAGAATCCACCACTAAAGATGCATTCAAAGCAGGAAAGCCATGCCATCGCTGTGGGCTTTATCGATGAAATCCTTCACCATGACAAACAGAGAGATTACCTTGCAACTTTACACACAGCTCATTATGAATTAACTTTATATTGTTAAGCATTCAACTTATGCAAGGATAGGAAGCACGTTGTCCCTCTTAGTTTGAGGCAATGGCTTGGTCAGCTTGCTTATGCACACCAACCTTTACCGACTTGTTTACCAAATGTAACATTAAGTGGCTGGAAAATGCCCTTCCTGCAAAAGTTTAGATTGGAGCTTTAAGTTTACGGCTGAAGCTGAGAGTTCTGCAATATTTTTTGGCAATCTCTTTGCAAAAGCCAAATTTGCATAAAATTCTTAAACTTGCTTATGTAAAGTTTTCAGAGTTGCTTTTGGCAAATAAGGAAAACACAAGTATACTCATTTGTGCTGGGCATGTATCCAGCCACCAAGCTTAACTACGATAAGATACTCTCTCAGAGTTTGAATATACATGGGATTTTTTTTGATTGTGA

**PLEIOTROPIC DRUG RESISTANCE 8 (PDR8)**:

GGAGCTTCCAGCACAAGGTTTGAATAAGTCTTTTGTGCAACCTTCTTTAGAGAGAGAGAGGGGCTGTTCATGCCTCTCATCATAAGGTATTATATTAGCTGTACTGTGCACACCCTTGTACTACTTCTGTGTAATCTACTCCCCTAATTGTTTTTTAATCAGCACACCAAAGTTTGTTTTTCCTTTTCTTCCTTTCATCTTTGTTTACACCCACTTTTCTGCTTGATTTGTCAAGGATGGTTACCACCCTCTCTCTCTCTCTCTTTCAACCTTAATTGAGCTTCCAATTTAATTTATAACCTTTACAATTAGGCCCCCATTAAGCCAACTGGTACAACTGTAATTGCTCTCCTAGCACACCTACCTTTTTCTCAAATGCCCAATTATCTGCACTACCCATTTGCCTTAGCACAATTTTGGCAAGCACATTTGTTTACAGTTACCTTGCCAAGCTCATAAAATTCCGCATCCCCTCCCCTGTTGAGCAGAGTTTCCTGATTGAATCATCAGCCAACACAGCGCGATTTGGAGTTTCGCTAATCGATCAAGACCCACTTGTAGAATTTGGAAAAAAACTGCTAGGAATTTTGGAAGGGAAAAGAGAGGTTAAGGAGCACCATGGCGCAAGCTGTGGAGCTCGAGAACATAGACATATGGGGGACACAGCTGAGGAATACTTTGAGCAGGAGCATGAGCAGGACAATAAGGCAAAATCCCAGCATAAGGAGCTGGTCATCCAGGGGGCCCGACAACGTGTTCACGCGTTCGCTGCGCTCTCGGAGAGACACCGAGGATGACGATGAAGAAGCACTCAAGTGGGCAGCCTTGGAGAAGCTGCCCACCTACGATCGCCTCCGCACTTCCATCCTCAAAAGCTTCGAAGGGGGCAAGGTCATCACGCAAGAAATCGATGTCAGAAACCTTGGCATGATGGAACGGCAGATGCTCATTGAAAAGCTTTTGAAGGTCACCGAAGAAGACAATGAGCGATTTCTTATCAAGCTTCGTAACCGAATCGACAAAGTGGGCATAGAGCTGCCAACAGTCGAGGTTCGGTTTGAGAACTTTACGATAGATGCGCAATGTCATGTAGGAGGTCGGGCTCTACCCTCACTATGGAATACTATCCGGAATTTCGTAGAGGGCTTATTGGGTGTGCTCCACTTATCCACAACACGAAAAGCCAGTCTGACAATTCTTAAAAATATCAGTGGCATTATCAAGCCTTGCAGGATGACACTTTTATTAGGGCCTCCTGGATCAGGAAAGACCAGCCTTCTCTTGGCTTTAGCAGGAAAATTGGATCCTGCTTTGAAAACAAGAGGAAGTATAACATATAATGGACATGCACTTCATGAATTCGTTCCTCAAAAAACATCAGCGTACATTAGCCAACATGATTTGCACATTGGGGAGATGACTGTGAGGGAGAACTTGGATTTCTCTGCTCAATGTCAAGGAGTCGGCACAAGATATGAAATGCTTTTGGAGCTTGCTAAGAGAGAAAAGCAGGCTGGGATATTTCCAGATGCTGAGATTGATGCATTTATGAAGGCAACTTCATTGAAGGGGATTCAGGGTAGTCTTCAGACAGATTACACATTGAAGATTCTTGGCTTGGACGTCTGCGCAGACACCATGGTAGGCGACCCAATGCGAAGGGGTATCTCGGGAGGGCAAAAAAAGAGGGTCACAACAGGAGAGATGATTGTAGGGCCTACCAAGACACTCTTCATGGATGAGATCTCTACAGGTCTAGACAGTTCCACAACTTTTCAGGTTGTGAAGTGTCTGCGGCAGTTTGTACATGTGTTGGAAGGGACCATCTTGATGTCGCTTTTACAGCCACCACCAGAAACATTCGAGCTTTTTGATGATGTGCTTCTTATTGCTGAAGGGCAAATTGTTTATCAAGGCCCACGCGAGCAGGTCTTGGAGTTCTTTGAGAGCTGTGGTTTCAAGTGCCCAGAGAGAAAAGGTGCTGCCGACTTTTTACAAGAGGTGACCTCCCGCAAAGACCAAGAACAGTATTGGTTTGACAAGAAAAAGCGCTACCATTATGTCACAGTGAAAGAGTTTGCTAATAATTTCAAACACTTTTATGTGGGAAGGCGTATCGCAGAGGAGCTATCTCATCCTTACCCTAGAGAGAGGAGCCATCGAGCAGCTTTGGCCTTTAGCAGATATTCTGTCAGCAAAGTAGACATTTTCAAAGCTTGTTTTGCTCGGGAATGGCTCCTCATGAAAAGAAACTCATTTGTTTATATTTTCAAAGGTGTACAGCTAGTCATTGTGGCCTTTATTGCAATGACTGTGTTCCTGAGGACAAGGCTTAACCAAACTGTAACAGATGGGAATTACTACCTGGGAGCATTGTTTTTTTCCTTGGTTATTATCATGTTCAATGGTTTTGCTGAGATGTCTTTAACAATCTTTAGACTCCCAGTATTCTTTAAGCAACGAGATCTCCTCTTCTACCCTGCATGGGCATATACATTGCCGACTTTTGTCTTGAGGATTCCACAATCGGTCATCGAATCAGGCCTTTGGGTCTCTCTAACCTATTATGTTATTGGGTTTACTCCTGAGCCTACCAGGTTCTTTAGACAGTATTTGCTTCTATTACTGATGCATCAGATGTCATCCTCCCTATTCCGATTGATAGGTGGTTTCTGTCGGTCTATGATTCTTGCCAACACTGGTGGCTCCTTTGTGCTCTTGATAATCTTCATGCTTGGTGGTTTTATTATTCCAAGAGGTAAAATCCCAAGCTGGTGGATTTGGGGTTACTATCTTTCACCGCTTGCCTATGGAGAGTCGGCCTCTACTGTTAACGAATTTCTGGCTCCTAGATGGAATACAATTGCACCAATGAGCAATGTTACCACGGGTGTGCAAGTGCTTATGAATCGTGGTCTTCGCACCGAAGCCTATTTTTACTGGATTGGTGTTGGTGCGTTAATTGGATTCATTCTGGTGTTCAACCTCCTCTTCACATTGGTGTTGTCATACTTCCACCCGATTGGAAAGCCTCAGGCTGTGATTTCAGAGGAAGCTCTTGCTGAAATGCAAGCCAATAATGAAGATACAAGTGGGGAAGTGCTGCTTGACCAGAAAATGCGGCGCTCTAGACGGAATCATTTGCCCAGGTCTTTGTCTTCTTCAGATTCATATGAATTTCAGATGCACCGTATGACAACTAATTCAACGCGCCGTGATAGCTATAATCACATGGGGGGTGGTGAAATTGCTGCTGTGAAGCATGGAATGTCCTTACCTTTCCAGCCATTATCTATTTCTTTTGAAGATGTCAAATACTTTGTAGACATGCCTGTGGAAATGATGGCTCAAGGGATTGTCGACAATAGATTGCAATTGTTACGCAGTGTGACGGGATCTTTCAGGCCGGGTGTGCTCACTGCCCTTATGGGTGTGAGTGGAGCTGGAAAGACAACTCTCATGGATGTTCTGGCTGGTCGAAAAACAGGTGGCTACATTGAAGGAGATATACGGATATCTGGGTACCCTAAAAGGCAAGAGACATTCGCTAGGATTTCTGGTTATTGTGAGCAGACAGATATTCACTCCCCCCAAGTCACCATTCGGGAGTCTCTGATATTTTCTGCATGGCTCCGTCTTCCAGCTGACATTGATATCGAAACCAGAATGCAATTTGTGGCCGAAGTCATGGAGCTTGTTGAGCTTGACAACTTGAGCGGCGCCTTGGTAGGACTGCCAGGTGTAAGTGGTCTTTCTACAGAGCAGCGAAAAAGGCTTACTATTGCAGTAGAGCTGGTGGCAAACCCTTCTATTATCTTTATGGATGAACCTACATCAGGTCTAGATGCCCGGGCAGCAGCCATTGTGATGCGGGCTGTAAGAAACACCGTGGACACCGGGCGGACAGTCGTGTGCACAATTCACCAGCCAAGCATTGACATCTTTGAATCCTTTGATGAGCTTTTGTTGATGAAACGAGGCGGTCAAGTCGTATATGCTGGTCCACTTGGTCAGCATTCTCAGAAGATAATAGAGTACTTTCAGGCTATTCCAGGAGTCGCTAGGATCAAAGAAAAATACAACCCTGCAACATGGATGCTGGAAGTCAGTAATGTCTCACAAGAACAGCGCCTTGGATTGGACTTTGCAGAAATCTATCGAAAATCTTCTCTTGCACAGCGGAACAAAGAATTGGTGAAGGAGCTCAGCTTGCCAGTACCAGGAACAACGGACTTGTACTTCCCCACGCAATACTGCCAAACAACATGGGGGCAGTTCAAATCCCTGTTATGGAAGCAGAATTTGACATATTGGCGCAGCCCAGGATATAACTTAGTTCGATTCTCATTCACCTTCATATCAGCTTTGATGTTTGGCACTATATTTTGGCAGCTTGGCACAAAGATTACTGCACAGCGAGATCTTCTTACTGTTATGGGGGCAATGTATGGAGCAACACTGTTTTTGGGAATAAACAATTGCTCGACGGTCCAGCCTGTTGTGTCGGTCGAGCGCACAGTTTTCTACCGAGAGCGAGGGGCCGGGATGTACTCGGCTATGCCATATGCTTTATCTCAGGTTGCGATTGAGTTGCCGTACGTTTTCGTCCAAACAATCATGTATGCATTCTTGGTGTACTCTATGATAGGGTTTCAGTGGGTAGCATCCAAATGTTTATGGTTCCTATTCACAATGTTCTGCACATTCCTCTACTTCACATACTATGGGATGATGGCTGTGTCCATCACACCCAACCATCAAGTGGCAGCAATTGTGGCATCAGCTTTCTATGCAGTTTTCAACCTTTTCTCGGGCTTCTTGATGCCAAGACCTAAACTACCTCAATGGTGGCGTTGGTATTATTGGCTATGCCCGACGGCATGGACTGTGAATGGCCTTATCACCTCCCAATATGCTGACTCACAAAAACTTATACAGGTAACTGTGCCACCAGGAGAAATAGCCAATGTAACTATAAGTGATTACTTGAACACGAATTTTGGATTTCACAAGGACTTCCTTGGCGAGGTGGCAGCAGTACTAATAATCTTCCCAGTATTCTTTGCTCTGGTTTTCACCTTTTGTGTGAGGGCCTTGAATTTCCAGAGGAGATGAGCCAAATATAGGAGCATGACAAGGAAATGAATCCTATTTATTTGGAATTAATGAGCAGAAGGTGGGCAATGGGGAAGGTGACACGTTCCATGTGAAAATTGGACATTCACTAATTTTGCGTGTTATCAAAGGAGGCATGCTCCCTGAGGTCTACGAGAAAAAGCAAGACGTGGGTTGTGGGGTGAGGCTTGTTTTTTCGACCTAAAGAGGAGTCGTGAAGCTGCTGAGTATTGCATTCCCTGCTGTATAGTGCATTCTTGCCTATGATCTCTCTATACTTTATACATGACCTCTTCAGCTGAAGGCTAACTGTCAACCAGCTAAGCACATTGCTTTATTGCAGTTTTAATCAGAGCCCCATTACAACATTTACCTGTGTATATAAGTGCAGCCATAGGATTTCTTTGTCCTGCTTCAGTAACGCCTTTCTGAAGTTGCAGGACTTGGAATTTACAAATTTTGTCTCAATTGTATCATTGGCATTTTGTACAACCCTGTAATGGCATCCCTGTCAAGGCAGGCTGTGAGCAATTGTATAAATCTGCAGTTTTGTCCTCTTTCGCCCTTGTCTCCTAGTATGGATTTTAGCATAAGGAGAGAGAGGAAGACGGAGAATGATCAAGTTTGGGGTTTGGTTTT

**CALLOSE SYNTHASE 12 (CALS12)**:

GTCAGCGATGAAGCAAAGACTGCACTGCAGTGCTGCAGCAGCCGCCGAGAAAAGCAAGCGAAACAGAACTCAGAGGTCGCCATGGCCAAGCTACAGGCTACCGTCCCTCAATTCCACGAGCGCAAGAGGAACATGGCGCAGAATTCGCCATAGCCAAGCTCTCCCTTCCGCTTGAGCATTTTGGGTGCCTTTTGAGCTCTGCTCCCCGTTTCACTTGGGCGCCTTCTAGAGAGGAAGCTCGCGCATAACAAGAGAAACAGAAACAGCCCTTCTCGCACTTTCAGCCCAAAACAGGGGCAATTCACTGGGCGCCCTTTCAGGGACCCTCGCCCTTTGTTGCTTCTAATTCCTCCATTCTTTTAGTCTCCCTGTGGAACCTGCTCACAAGAAGCGAAGTGAGGAAGATTTAGCTCCTGTGATGCACCAAAAAGAGATGATGATGACAACCTTCTGTAACTTGGAATTGAAGCTTTGAGCTATTTAAAAAGGGTTTTAAGCTTTCCAGCATTTGATAATGGCGGACAGAGAAGATTTTGAAGATTCGTACACATTCAACATCCTGCAGCTCAATGAGGATATGATAGCAGATGATGATGCTTTGCGGTTTCCAGCCGTTCAAGTAGCCATGCAAGTGTTGAAGAGCGTGGAGGACCTCAGGCATCCACCAGCAGAGGTCTTGGTATGGCATGTAGGGGTCGACATGCTTGATTGGCTCGGTGCCTTTTTTGGCTTCCAGGCTGATAATGTCAAAAACCAGCGGGAGCATGTGGTACTTCTGCTTGCAAACTATCAAATGCGGATGCAACCCCCTCCGGAGCCCATGGAGAAGCTGGACCCGAGTGTGGCTCGTCTGCTGAGGAAAAAACTGCTGAAGAACTATAAGAAGTGGTGCTGTTTTTTCCACAAGAAACAGAACCTCTGGCTTCCGAAGAATGTGCGCAACGTCGATACTCGCAGAGAGCTTCTCTACTCTTGTCTTTACCTTCTCATATGGGGGGAAGCTGCCAATCTTCGTTTCATGCCCGAGTGCCTCTGCTTTATATTTCATCACATGGCCATGGAGCTCAACCACATCATTGAACGCTACGTGGATGAAGAGACGGGACAGCAAAATCTTCCATTTTCTTACGGTTGGAACAACTTCTTAATGAAGGTCATCGAGCCCATCTACAAAACGGTAAAAGGAGAAGCAGAAGCTTCCAATAATGGGAAAGCGCCTCATTCAGCTTGGCGAAATTATGATGACATGAATGAGTATTTCTGGTCCAAGCGGTGTTTTAAGCAGCTTTCATGGCCATTGAAGCAGTCTTCTAGCTACATGGTTGCTCCAAAGAAGGGCAATAAGAGGGGAGCTCTACTGTCTGGCCAAGAGAAGGTGGGGAAGACGGGCTTTGTGGAACAACGCTCATTTTGGAACATCTTTCGCAGCTTTGATCATTTGTGGATAGGTCTCATTCTCATGCTGCAAGCCATGATGATTCTTGCTTTCAACGGAAATGGAGAACTAGGGAATATTGTGTATAAAAGAGATGTTCAAGCGGCTCTGCTGACTATCTTCATCACGTGGTCAGGTTTGCGTCTAATGCAAGCAGTCTTAGATGCAGGGATGCAATACAGCTTAATATCAAGTGACACAAAGCTTGTTGGCATACGGATCCTTCTGAAGATAGTGGTAGCCATTGGCTGGACCACCATATTCTCTGTCTTGCTTATCCGGATTTGGTGGCAGCGAGAAGCAGATGGAAGTTGGTCAACACGAGCAGATCAGAGACTTATTCAGTTTATTCAAACTGCAGTAGTTTTTGTCTTACCTGAGGCTCTTGCACTCACCCTTTTTGCCCTGCCTTGGATAAGAAACTTTGTAGAGCAATCAGAGTTTGGCATTTTTCATGCTATAACATGGTGGTTTCAGACAAGATTGTTTGTTGCTCGCGGCTTGCGCGAAGGTGTCATTGACAATCTGAAATACACTGCATTCTGGCTGGTTGTCCTCTTTGCAAAATTTTCCTTCAGCTACTTTCTGCAAGTGAGGCCCTTGGTAGCTCCGACTCGAGAGCTTCTGAAACTTGAAGTAGATCAATTTCAGTGGCATGAGTTTTTCAAAAATCACAACAGGTTTGTTGTGGTTGTGCTTTGGGCACCAATTATCTTGATATATGCAATGGATCTGCAAATCTGGTACTCTATTGCCTCATCAATAGTAGGAGCATCTGTTGGGCTCTTTGCGCATATAGGCGAAATCCGCAATGTTGAGCAATTACGATTGAGGTTTCCATTCTTCGCAAGTGCAGTGGCTTTCAATTTGATGCCAGCTGCAGGACTGCTGGTGACTGATGCCTTACCCCACCATCGTCTTGTTGCTCTCTACAGAGATATGGTTAGCAGGTTAAAGCTCAGATATGGGCTTGTAAAAGCCTACAAGAAGTTTGAGAGAAGCTCCCTTGAGTCTTGGAAGTTTGCACGCTTGTGGAATGAGATTATTTCCTTTTTCAGAGAGGAGGATCTAGTAAGCAATCATGAAGTAGAATTATTAGAGATACCTCAGGTTTCATGGAATGTGAGTGTTATGCAGTGGCCCTCTGTGCTTCTTTCCAATGAAGTTCTTGTTGCCTTAGGATTTGCCCGTGATTGGCAAGGTGTAGAAGATAGAAAGTTGTGGTGCAAGATTTCCAAGAATGAATATAGAAAATGTGCTGTATTAGAATGCTATGACAGTTGTAAACACTTACTGTTAAGGATTCTCAAAGATGACAGTACGGAACACAGCATTGTTCAGCACTTGTTCAAGGAGATTGAAACAGATATTGAACTTCAGAGTTTCTGTCAGCACTACAAGGTGAATGAAATAGCTGAGGTTCATTCACGTGTTGTGGGCGTCGTGGAAGTTCTATTGAAGAGGCCAGTATTAAAAGACATGCAGAAGGTTGTGGATGCTCTGCAGAACTTGTATGATGTGGTGATCCGAGATTTTCCCATGCGCAAGCGCCAATACGAGTCACTGAGGGGGGCTGGCTTGGTTACCGATTGCAATGCGCTGTTGTTTGTAGACGCTGTTGACCTGCCGAAGCCAGATGATGAAGTGTTTTTCCGAATGTTGAGGAGGCTGCATACCACATTGAGCACTCGAGAAGCCTTGAGCGGTGTCCCCAAAAATCTGGATGCAAGGAGACGCATAGCTTTTTTCAGTAATTCTCTGTTCATGAATATGCCCCACGCTCCGTCAGTAGAGCGAATGCTTGCTTTCAGTGTACTCACACCATACTACAGTGAGGATGTTATGTATACCAAAGAGCAGTTAAGAACTCCAAATGATGATGGGGTAACAATTCTTTACTACCTCCAGAGCATTTATCCAGATGAGTGGAGTAATTTTCTTGAGAGGATAGGCTTGAAAGAGAGCGCAGACCCCGAGGGTGAGCTTTGGGATGGCAAAGACCAGGCAAGACAGCTTAGGCTATGGGCCTCCTACCGTGGTCAAACACTTGCTAGGACTGTCAGGGGAATGATGTACTACCAGCGTGCATTAGACACCCTTGCACAACATGATGCAGCATCTGAGGCTGATTTCAAAGAGGGGCTTGCCACTGCACAATTGGAGAGCGAAAGCAGCAGCCACAGATCCTTAAGCAAGCTTAATGTGAGTGGCTTGAGTGCCAAAGGTGCCATGCACAGAGAAAGATTAGCTAAAAAGGTAAAAACGAGGAAAGCAATAGCTGGACTGAAATTCACATATGTTGTGGCATGCCAGATTTATGGTGCTCAGAAAGCTAAGAAGGAAATCAGAGCAGATGACATTTTATTCTTGATGAAGAATCACCCTGCAATGAGGGTAGCATATGTTGATGAAGTTCTTTCAGGTCAAGATAAGAAGTACTACTCAGTGTTGGTCAAATATGATACATACATGGAGAAAGAGGTAGAAATATATAGAATTCAACTGCCAGGTCCCCTCAAGCTTGGAGAAGGAAAACCTGAAAATCAAAACCATGCCTTGATCTTTACAAGAGGGGATGCAGTACAAACTATTGATATGAACCAGGACAATTACTTTGAGGAAGCTCTCAAAGTGAGGAATTTACTGGAAGAATTTAACAAGTGTTATGGTTTGAAGAGGCCACAGATTTTAGGGGTTCGAGAGCATGTTTTTACAGGCTCTGTCTCTTCGCTTGCATGGTTTATGTCTGCCCAAGAGACAAGTTTTGTCACCCTTGGTCAGCGTGTGCTGGCGAACCCTCTTAAGATAAGAATGCATTATGGACACCCAGACGTGTTTGACAGGCTATGGTTCTTGACTAGGGGAGGCCTTAGCAAGGCTTCAAGGGTGATCAATATCAGTGAGGATATCTTTGCAGGATTTTACTGTACTCTTAGAGGAGGAAATGTCACTCATCATGAATACATCCAAGTGGGTAAAGGTAGGGATGTGGGCCTAAACCAGATAGCTCTTTTTGAAGCCAAGGTCTCTAGTGGCAATGGCGAACAGATCTTGAGCAGGGATGTTTATCGTCTTGGACACCGCCTTGATTTTTTCCGCATGCTCTCATTCTACCATTCAACAGTGGGGTTTTATGTTAACAATATGATTGTTGTTTTGACAGTCTATGCCTTCTTGTGGGGTCGTGTTTATCTTGCTCTCAGTGGTATAGAGGATAAGATTAATCATAGTTCAGTCAATGGGAGTCTAGTCAACAAAGCACTCACAGCCTCACTTAACCAGCAGTTTATTGTTCAGCTTGGGTTGTTTACCGCATTGCCAATGGTTTTGGAAAATATTCTTGAGCTTGGCTTTAGCAGAGCAGTGTGGGATTTTATCACCATGCAACTAGAGCTGTGTTCAGTTTTCTTCACTTTCTCATTGGGCACAAAATGCCACTACTTTGGTAGGACTCTACTTCATGGAGGGGCTAAATATCGCGCTACTGGTCGTGGTTTTGTGGTGAGACATGAGAAGTTTGCCGAAAACTACAGGCTCTATGCAAGGAGCCATTTTGTGAAAGGAGTTGAGCTGATAATTTTGTTGATAATTTATGAGGCATATGGAGGTATAGCTTCAAATACTACTGCATATATCCTCTTGACAGTCTCAAGCTGGTTTCTGGCTCTCACATGGATCATAGGGCCCTTTTTGTTCAACCCCACAGGGTTTGATTGGTTACAGACGGTTTATGACTTACAAGACTTTCAAAACTGGTTGTGGTACAAAGGGGGGGTGTTCACAAAAGCAGACCAAAGCTGGGAAGTGTGGTGGAATGAGGAGCAAGATCACCTAAGAACAACTGATTTCTGGGGAAGATTGCTGGAAATTGTCCTTAGCGTGCGTTTTTTCCTTATTCATTATGGCATGGTCTACAGACTGCATATTGCTGACAGCAACAAAAGCGTTCTTGTGTATTTTGTGTCATGGATATACATTATGGGTGCCATGGTGCTGTACATGATTTTGGCATATGCTGGAGAGAAGTATGCAGCAAAGAATCATCTCTATTACAGAGCCATTCAGTTACTGGTGGCAATGTTTCTAGTAATGGTCATTGTACTCTTGCTGGAGTTGACAGAGTTCAAATTTGTGGACCTTTTTCTCAGCTCCCTAGCTTTTATACCCACTGGATGGGGGTTGATTTGCATCTCCTTGGTCCTAAAGCCTTTTCTGGTAAATACAGTTGCATGGCCAGTTATTGTAGCAGCTGCACGGCTGTATGAATTAGCAATTGCTGGTGTAGTGATAATTCCCCTAGCAATACTGTCATGGCTTCCTGGTTTTCAAGCAATGCAGACTCGCATACTGTTCAACCAAGCTTTTAGTCGAGGTCTTCAGATTTCTCGAATATTATCAGGAAAGAAACAAAGTCAAAGCAGTTAAGAGCCAAGGTGGAGTATTTCACAAGTTTTTTGGGTCTTGAAGATGTAATTATCTGCAGTTATTGATGAACGGAATCCACTTTCTTAACACACAAGCTTAGCATTCAGCATGGAGCCTGTGTAATGCCATAGTCGACCATTGTGGAGCCTCAAGTTATCTATCATTAAACATCAAGGGGTTTGATGTCGCTGTGCTTCTTCTGCCGAGTTCTCGCACTGAAGTCAGCACCTGTGGTAAAGATAACCTTGTTGTGCAAGCTCAACGAATGTTTTTGGAGGCTGTGTGTATCATAGCCAACCTAGTCACATTTTTTGCAGCCATTTATACAAGTTTCCAATCATTTTTATCCAAGCTAGGTCTGTCATCTGTTTTACTGCCTTCATGTACTTCAAAATTTTCTTGATAAGATCGGAAACTTTGTTCTTTTCTTTTGTGCTCAACTAAAGGGCACTGATTGGCCAACTCTTAAAAGTGGAGCTCCTTTATGTTCCTTCTTTTGCTCGAGTTGATACGTGATTATTATGTAACAGAAGAACCAATCATCATACAAAACTCAGGGAGTAGGCATGGAGCCGAAAGCTGACAGCTTGAGAATATATGCTGGGCAGTAGGTGCATGATACAAGACACTACCTTTTGTTTGGAGCCAACATTCAGGTATAATGCAGTGATATGGTGTCTTAGGTAGGTGTCCATGAGATGACGTCTAAGGTAAGCTTGAGTGATGGTGCTTGCCAAATAGAAACTTGTCAAGTTCGACTCGTGGTGGTCGTCAGAGAAGGGAATGTAGCTTAATCGCACAGAAACAAGCTTCAGAATATCTGAGCAAAACGCTGAGTATGCTACAGTTGGGATTTTGTTTGACACCAAAAAATATCAAAAGCAAGGGTGCAGAGAAGGCAACTAAAACAGTCTCATAAGACTGGTGCCACTTTAAGACCTATGACACGAGCACGGGTATTTGATAAGTCTGATGGAAGAGTTCTGATAATTTGCCTACTTGCACTGGATCTATTACTTCATAATTTAGGATTTTGTTAATTTGCCCACTTGCACTGGATATATTGCTTCATAAGCTAGGATTTTGTTTGTGAATAACAGAAGACTTAATGAGCCAAGCTCAAGCAAAACCCATGTCTATCAAGGCACAGATCTACGAACTAGAAGAGAATCAACGATGCCCGCAGCCTTGCGACAAAAGCTGGCTGCTGCAAGGGAGCTCTCTAAATATAAGAGCCAGAGCCACACAAGCTGTTCAAAAAGGTTTTTTTGGCAGTTTTGGCTACCTAAACCAAAGGTTTTTCTTTATGCTGTATCAGCCTCTTGTTAAAAATTCCTGTTAATGTCTTATCAAACAAAATTCATATTTTAC

**PYROPHOSPHATE-ENERGIZED VACUOLAR MEMBRANE PROTON PUMP 1 (VHP1)**:

CACCAATCTAGCTAATAGCCTACTCCCTCTGCTGGACCCATGCATGACACAAAAATGATTTAGCACTGACATACAAACCCTCTTCATAAGCGATTTCTTAGAGACTACCATTTTAAACGTCTACATTGCATGATGATTGAAGCGTGATCAGCGTTAAAAGGAGATTCTGGTATAAAAATGAGAGTTTATCCTTGGTAATGAATGGACTGCTTGAGCCGTCTTCAAGTCGCAGCATGCCGTTTGAATCTCAACAGACCTCATCCTAACAAATTTTAAGCCTTAACAGGAACTAGTTGGATCTGTCCCTACAAGAACTAGAGAAGGAAAAAATTGCATTGGCTGAAGAGGCCAAGATCTCCAAGATCCAACTGCATCTATATCTTGAAAAGAAGGCCACCATGGGCTGCAAAAAATGGAGCAAAAACAAGCGACTCCACTGCCATGAGCTTGATTAGAATGTTGAGAGAGGGTCCAGATGTGTCCTTCAATGGGTCACCAACTGTGTCTCCAATTACAGCTGCCTTGTGTGCATCCGAGCCCTTGGGACCAAGGGACCTTGCATGATTTGAAGCTCCAGCCTCAATGTATTTCTTGGCATTATCCCATGCCCCCCCAGTGTTGGATGCAGAGATCGCAATCTGCACACCAGATACCAAAGCACCAGCAAGAAGCCCTGCAAGAGTTTCCACTCCAAACAAGATGCCCACAATCAAAGGTGTCAACATAACCAAAGCACCTGGAGGAATCATCTCTCTGAGGGAAGCATCCGTAGATATCTTCACACAGGTGGCATAGTCTGGCTTGGTGGTACCCTCCATAAGGCCAGGAATGGTGTTGAACTGCCTCCTCACCTCCTCCACCATGCTCAACGCTGCCTTTCCCACACTCTTCATTGTCATAGCAGAGAACCAATACGGCAGCATGGCTCCAACAAGCAATCCAATGAAAACCCTAGCTGACAAGACATCGACTGTAGAAAGTGATGCACGGCTCACATAGGCTCCAAACAATGCAAGGGACACCAAAGCTGCTGACCCAATTGCAAATCCCTTCCCAATTGCAGCAGTCGTGTTGCCAGCCGCATCAAGGGCGTCAGTCCTCTCACGAATGTGGTGACTCATGCCAGCCATCTCCGCAATGCCACCAGCATTATCACTAATGGGTCCATAAGCATCAATAGCAAGGCCAGTTGCAATGGTGCTGAGCATGCCAAGGGCTGCTACGGCAATACCATAAAAGACAGCCAATGCATAGCTTACATAGATCGCGGCTGCTATGGAAAAGATTGGGATAATCACAGACTTGTAGCCCAAGGCTAAGCCAAAGATGATATTGGTGGCTGCTCCCGTCCTGCACGAGTCTGCTACATCTTGTACTGGACTATATGCATTGCTTGTAAAGTACTCTGTTGTGTACCCAATGATCAAGCCAGCCCACAGTCCAACGCCCACACAAAAGAAAAGTTTCCACCACTGCACCAGTTTAGGGGTGCTACCGAGGTCATATATGTGGAATGATGCCGGCAGTGCAAGCCAACAAACAACAAGAATGCCGACTGTCATAAGCAAAGTTGAAACAATAAGCTGCTTCTTCAGAGAAGGTTCAATTTGCTTGACGTCTGTAATTTCATAGAAATCCGTGGCAAAAAGAGTGGTAATTAAACAGACGATAATGCCCATGGAGCTGATGAGCAGTGGAAAGCACATTGCTGTGAAATTATGATCATTCCCAAATGAAGAGATTGAAGCCACGACCAGAGCTGCACAAGAGGACTCTGCAAAGGAGCCAAAGAGATCTGAACCCATGCCGGCAATATCTCCCACATTGTCACCAACGTTGTCAGCAATAACCGCTGGATTCCTTGGATCATCTTCAGGAATGTTTCTCTCCACCTTCCCAACAAGATCAGCACCGACATCAGCAGCTTTGGTATAGATACCTCCACCTACTCTCCCAAAGAGAGCCACAGAGGATCCTCCAAGCCCATAACCAGTGATTGCTTCAAAGAGACCCTCCCACTCCGTCTTGTAGTAAAGCTTGAACAAATTGATGGTAATGTATAGCACCAAGAGGCCATTGGCAGCAAGAAGAAAGCCCATTACTGCACCTGATCTGAATGCCACAATGAAGGCCTTGCCAACACTCTTTCTGGCCTCTAGAGTTGTGCGTGCATTTGCATATGTAGCAATTTTCATCCCAAGATAGCCGGACAGCACAGATGTCACAACCCCTAGAGCAAACGCTGCTGTGGTGAAAGCTGCTGTGATGAGCGCTGGAGCACATTCCTGTGTCGTGTCATACTTGCATTGTTGCGGAGACAAGCTGAAATTTTCCACTGACCCCAAGAACAAGAAAATCACAACAGCAAACGCGCTCAAGAAAATGCCCAAGTATTTGTACTCGGTTAGAAGAAAAGAGTTGGCACCAACAGAAATGGCTTCTTGGATCTCAGCACATTTGGAAATAACCGCAGCACCATCGTGATCTCCTTCTTCCTCATGATGCAGCAGGTAGTCTTGCCCGTTATCTACGCTTCCATTCTTCTTGTCCTTGGGCCCACTCGAAGAGGGCAACACAGAAACAAAGGAGACGACATACCACTGGAAGAGCGCAAACACGATGCCTATCACGGCTGCCACGGGAATCAAAATCTCCGTTGCAAGCTCAGACAGCACCGTCATGGCGAGCTTTTCAGCAGAAAATTCCAAGCTGTTATGTGAACAAATTGCGCGAGAGAATAAGCTCTACAATTGACTACGGAGTGCTTTTCTTTGCGGCGCAGAACGCCAAATCTATTTTTGGACTTGAGTTTCCAAATTTAGAAGAAGTGGGCGCGTGAAGAGTTTTGATCTCTGCAGAAAACGAGCTCTAAGGTGGGCGATGGAGAGCTAGAAAGGCGGGATTTTGAAAAACCCAAGAGCGCAATCAGCGATGGCGAGTGCTGTGTGAGGGTGATGCTTATGAAAAGGGTGATGGGTGCGTGTGTTTGAAGGCAAGGGTGTGTGTGTGCGTG

**SUGAR INSENSITIVE 7 (SIS7)**:

GAGAGAGAGAGAGAGAGCTCTCTCACACACACACACAGACAAGCGCGGGCGCGCAGAGACACTCTCAAGAACACATAATAAATATGGCTTCTTTTATGAGTTCCTGGCAATCACAAGCAAGCCCCATTCCTGTACTAGCCAGGGAACAACATAAAAACAAACCAATGAAAGTGTGCATGCAACCATCCCACCTATCCTCACAAAAACGTATGAATGATACCTTAGCAATGCTAGCCTCATATAATGATAATGATTTGCCGATACCACCTTTAGAACCTCCACCTTCCTCACCCCTACCTAAAAACTCTTATAGCCCTCTTTCTCTAGTCTCCCACATTGCCTCCAATCTTTGTAATGCTCTTGAAGACCTTGTCAACTCAACAACATCACCTCCACATGATGCTATCAATAGCCACCTTGCCTCCAACTTTGCCCCTTTAATGGCTGAGTCCCCCCCCACTCCATGTCCCCACATTCATGGCACCATCCCTCCCTCCCTTAATGGCATCTTTCTCCGCAATGGACCCAACCCCTTGCACCCCCCAAAGGCTGGCTACCACTTCTTTGATGGAGATGGCATGCTCCATGCAGTCAAGATAAGTAACAACCAAGCTGTGCATGTGTGCCGCTTCATACACACCAATCGCTACGTCCACGAGAAAGCTGCCGGACAACCCCTTTTTGTCAAGGCACTGGGAGAGCTCAAAGGGCTAAATGGCATTGCAAAAATCTCAATTTTGATTCTCAGACATGTGCTTGGCCTTCTTGACATGATGAAAGGGTTTGGCCCTGCCAATGCTGGCCTCGTTTACTTTAATGGTAAACTGCTTGCCATATCGGAGGATGACATGCCTTACTGGATCAACTTGTCTAACGATGGAGACATCCATACAATATGTCGCTTCGACTACTCCGGAAATCTCAAGTCCAACATGACTGCCCATCCTAAGGTGGACCCCAACACAGGGGAAATGTTTTGCTTTAGCTATAGCTTGTTAGCTCCACCTTACCTAAAGTACTTTAGGGTAAGCCCTAATGGAAAAAAATCAAATGATGTCTCTATCACAATCAAAGAACCCACATTATTACATGACTTTGCCATCACAGAGAACTATGCGGTGTTTTATGAGCATCAAATTGTGTTTAGATTGCACAACATGTTCAAAGGAGGGTCACCTGTTGTGTGTGATACCAAGAAGGCACCTAGAGTGGGGATCCTACCAAAATATGATACTAATGAGGCCAACATGAAGTGGTTTGAATGTCCCCTAAGCTTTGGAACATGCATGCACTTTCTAAACGCATGGGAAGAAGGTGATGAAGTTGTGATGCTAGGATGTGTGAATTCACCGTTGGAGCTCATTTTCACACACTTGGAGAAAATAGAAAGTAAGCTAATAGAGTTTAGGTTCAATATCAAAAGTGGACGAACAACAAGTGAGCAAGTGAGCTTTGCAAACATTGACATGGGTAGAATAAATGATCAATACACGGGAAAAAAGACAAGATATGCTTACTTATGTACATTTGGACCATGGCCTAAGTACTCCGGTGTTTGCAAGATAGATTTAGAGGCAACACGTCTTCCCCAACCAAAAGAGTGGAGCATTGGTCAGAAAAACGTGCACTTTGATGAACCATGT

**CALCINEURIN B-LIKE PROTEIN 1 (CBL1):**

GGCCAAAACATAAGCTTTTCACATTGATGAGATAACTTAAAAAATTCAACAGTACAAGACCAGCAAAGTCAAAAACGCTTGACTATGTATTTTATATTACAAAAGAGCTTTCTGGAGCTCCATACAGATGATTGCATGGGATAAACGTACAAAAGCAATCAAAGTCTGGTCTATGGCACATACATCTATTTGATTCCTTGCATGAAAATGATTGCCCAAAGCCATTTCACCCTTTACTGTAAAGGCATGGGTATCCCATCTTGACTGACATCAGCAAGTAAGCATTTAGCATGTAAGAAACATGAAGGGACCTTCTGCCTACGGTCCCAAATCTTTCTGAAAAGCATGCATAATCGAAGTTTGTTAGAAACTTCTATAGATCTGCACGATCATCTTAATGCCCTCGCATATAGGATACATAATATTGACCGGCACACTAAAAGTGTTGACTAAGTCGCCATATCCTCAATTTCAGAGTTGAAAACAAAGCTTGGAAATGCCGTTGTTATATCCTTAAGGTAGGGCAATGTCATTATCTTCATAAGAGATGGATTGCGGGACACAAATCTCCTCCATTCGTCAATATCAATCTTGCCATCTCTATTTGTGTCTGCTTCAGCAAAAGTCTTATCAAGAATAACTTCGACCACCTCATCTGCTAATCTCATATCTGATTCCATCAACAACGCATCCACCATCATTTTTACCTCTTCTCGCTCAATGTAGCCGGTGCTTCTCAGATCATAAAGCTTGAAAGCAAAGTCCATTTTTTCATTGGCAGCTGTATTTGGGTGAAAGACGCTAAGAGCTCTGACAAACTCCCCAAACTCTATAACCCCATTTCTTTTGAAATCAAACACATCAAACACCCTATCAGCAAATAGACTCTGCTTTTTACTGTTCTGAAACAAGGCAAGCTGAAACTCCTCCTTATGTATCAAGCCATCATCAATGACAGCACTGCTAATCTTCTTGAAAAGCTCATACAAAGCCTCAACTTCACTCACACTGAATGTAGTCTCTGCTGCAAGATCTGCAGGATCTTCATACCCTTGAGGCTGCTTGACAAGTTTAGATTTAAAGCACCCCATGCGAAGCCCACAGGCATACTCCAGGAGAAATAGGCGAGTCACTCTTGCTATGTAGCAAGCATAAATGCATGAGCTACTTCCAACTGGAAAGAAGTAGCACTGGTTTTATTTGAGGAGCATTCTAGAGCAGCTCTGACTCTTGAGTATGTCAATGGTCGCCGTGCAGAATGCAATCTCTTCAGCATTTCCAGGTTGCAGAAACATAGCTATATAAAGACCTGCGACGAGGCTATGTCTGCAAAACCTAGAATAATCACTGGCATTCTAGCATCAGGCGGGCGATGGCGCGCTAAAATATCGCCTCTCTCGCAAGCTCGCAGGCTATGGCTCTCTCTCTCTCTCTCTCT

**METHIONINE SULFOXIDE REDUCTASE A4 (MSRA4)**:

TACAAGAACACACTTTTAACCAAACAGGGGTTACATTTAAAGGTCTCCCTTCCCTGTAAAGAACCTGGTCCTTTAGTCCATCACGCACGCACCACAAAAGAACATCCATTGCTACAATAGCTGTTGCATGTTCAAGAAGGGCTTTCAAAAAAACTAGATACTAGTAGTGAGAAAATTTCCACTTTAGTGATAGGCTCTCTTAACAATCAAACAGATGTTATAAAGATGATACAAAATTTGATATACAAGTCCTCGAATGTTGTTATTCTGTGTACAACAATAATGGTCCGTGCTCCTGAAAAATCAGTCGAACGAGTAAATTTACTTGTGCATGACCTAGCCATAGCAGCGAATGGGATCATTGCAGCCTTTCTCCGCTGACTGCTTGAAACCAGAGCGGCCCCCTTTTGCTAAATATTGCTGGTGATACTCCTCTGCCCTATAAAACTTCTTGGCAGGTAGAATTTCTGTGACTATCTTTCTATTTACCTTTTGCTGATGCTTTTGCATAGATTCAATGGCAGCTTTCTCTTGTTCAGGCGTGTAGAAATAAATCCCCGACCGGTACTGTGTGCCCACATCATTCCCCTGACGGTTCAACGTAGTAGGGTCATGGCGATCCCAGAAGCAGGTAAGGAGGTCATCATAAGCGCAAGCTTTTGGGTCATACTGAACACGTACGACCTCCGAGTGGCCCGAGTTGCCTGCGCACACAGCGTTGTAGGTGGGGTTGTGAAGGGCACCTTGTGTGTAACCCACCTCAGACTTGGTAACCCCCGGGACTCTTTGAAAAGCAAGCTCAACGCCCCAAAAGCAGCCAGCCCCAAATTGGGCAAACTGCTGCCCAGGGGCAGGAACATCATCATCAGGGCCTTGGGCTATGCTCATGCCGGCAGTGTCGGCTTGCGCGCCCACAACACCACGAGCATTGCCCCGCCCAATACCGAGCTTCTCAAGCCATGATCCCATGGTGCGCAGACCGGGATTGAAATGAGGGTTAGGTCTAGGGTTAGAGGATAAGCTCTGGATATGGCGCTGGCGATGGCGAGGGAGAAATGGGCTATGCAGGCTCTTAAAATTGGAGATGGATGCAATGGCCACCTGCGTTATGGTGGCCTGCTGCAAACAGCTTGCACTGTGTGTATAGCCCACCTTCATTTTCTATCTTTTGAGAGGGTAGGCAAGAATTATCAACCTTGCAAGTTCACAAGCAACCTGATTCTTCCCCACTAATTTGAAAATG

**ACONITASE 3 (ACO3)**:

CTCATTACAAGATTCAACGGAACTTTGATGCGTCGCATTGTACCTTTGCAACTGCATGTCAAACGAACAAAGATAAAAGGGATATTTGTGACAGATTTACATGCTGATCAAGCAATAATGCATGAATACATTATAGCCTCAAGTACATGCTCTTTGGCATAGTTTTCATAAACTTAAAAACACTCCTTAGAACATTTACTGCATAAAAACCACATTATTATCTGAACAAAGCCTTTGTTAGCCTCACTGAATATTCCTTTTTGCAATTGTTTTCAGCTTCTACATGGCTAAAACTACAGACAACATGAACAAAAAAAGCCAGTACTCTCTTCATAAAGCTAAAACAATTGTAGGAGCAAAGCGTCAAGGGAACCTTTGTAAGAAAAAGAATCCATATTGACCAAGTTTATTCATTCACTCGGCATCGTGCCATAGCACAACTATACACCATCTCTTGCCTTGTTTAAGGCCTCCTTATTTTAACTCAACCAATAGAACATGATTGGCACCATATCTGATTCCTGATTGTTGATATCAAAAAACTCTTGCCCAAGCTACAAGCATACTGAATGGGATCACTGCTATATGTGAACTAAATGAGTTGCCGAAGCACATATTGAAGAATTCCACCATGTTCAAAGTACGTTAATTCAACCTGTGTGTCAAACCGGAGAGTACAAGTGAAACTCTTTCCATTATCAGTTGTCACTTCAATATCCTGCCCGGGAGTGATAGACTTGACGTCTGAAGGCAAGTCAATGGTGTACCTTTCGTAGCCTGTCAAGCCCAGAGAGTCTGCATCTTGACCAGACTTGAAGCAAAGAGGAATGATACCCATGCCCACCAAATTGCTGCGATGAATGCGCTCAAAACTTTTTGCAATAACAGCTGTCACCCCCTGCAGGTAAGGACCCTTTGCAGCCCAATCTCTGGAGCTTCCACTACCATATTCAGCTCCTGCTAATATGACCGTATCCAAACCTGCTTCCTTGTATTTCATGGCAGCATCATACACATAAAGCTTCTCTCCAGAAGGTACATGAATTGTCTTGGGCCCTACTTCACCCTTCAAGAACTTGTTGACAATACGAATGTTTGCAAAAGTCCCCCTGGCCATGACTTCATCATTGCCACGACGGCTACCATAGGAGTTGAAATCTCTCCGATCCACACCCCTCTCCATCAGGTACTTGGCAGCTGGACTGTCCTTATGAATGTTTCCAGCTGGAGAGATGTGATCGGTTGTGATGCTGTCTCCAAAATTCAAGAGACAGTATGCATCCTTCACGCCATGAGCACCAGGAGGCGTGCGAGTCATAGTCTTGAAAAAAGGCGGATCGTGCACATAAGTAGAGGTTGGATCCCACGAATATAATGCATCGCTAGGAGCATGCAACTCATTCCACATCGCATTACCCTTTGTAATAGATTCATAAGTAGACTTGAACATCTCTGGTAAGACAGAGGATTCCACTACCTTTGCAATTTCCTCACTTGAGGGCCAAATATCTCTTAAAAACACTTTGTTTCCATCTTTGCCAGTACCGATTGGTTCCGATTCAAAATCGATGTTAACAGTTCCAGCAAGTGCATATGCCACCACAAGTGGTGGAGAAGCCAGATAATTTGCACGAGTTAATGGATGCACTCTGCCCTCAAAATTACGATTCCCAGATAGGACCGCCGCAGCAACAATATCTTTCTCAGCAATTGCTTCTGCCACTGACTCATTTATCTCTCCCGAATTGCCAATGCAAGTGGTGCAGCCATACCCAACCAGACTAAATCCTTGCAAGTCTAGAAACTTGTTTAAACCACTTTGATGAAGATACTTTGTTACTACTCCAGAGCCAGGTGCTAAACTTGTCTTAATCCAAGGCTTTACCTCTAGGCCTAATTCATATGCTTTCTTTGCAACAAGGCCAGCTCCCAGCATAACACTCGGATTGGAAGTGTTTGTGCAGCTAGTGATGGCTGCTATAACAACATCACCGTGCCTTAATTCGGCAGGCTTACCATCAAATGCAAACTTTGCAACTGCACCTTGCTCATCTTTAGGTATAGCAAAACCCTTGAACCCAACTTTGTTGTTAAGGCAGGCATGCCAATCCGCCTTCATGTCTTTCAGAGAAACTCGATCATGAGGCCGCTTAGGTCCAGAGAGAGAAGGCTCTACTGATTCAAGGTCCAGCTCAAGGTACGAAGAATATGTCTTCTCTTTTTGAGGCTCATCATAATCAATAAACATCTTGTTCGCCCGTAGGTAAGCCTCAATCATTTTCACCTTCTCTTCTTCTCTTCCAGTTAATCGAAGATATTGCAGTGTAACATGATCAACTGGAAAAAAGCCCATGGTGGCCCCATATTCCGGGGACATATTCGCAATGGTTGCACGGTCTGCTAGAGACAGAGCAGACATTCCGTGACCATAAAACTCCACAAACTTGCCAACCACACCATGTTTCCTAAGGATTTGGGTGATCGTGAGAACTAAATCAGTTGCTGTAACCCCTGTTCGCATCTTTCCATTCAACTTAAACCCGACAACACCTGGAAGAACCATACTCATGGGTTGGCCAAGCATGGTAGCCTCTGCTTCAATGCCACCAACCCCCCAGCCAGCCACACCAAGCCCATCTATCATGGTTGTGTGTGAATCAGTCCCAACCAGACTGTCAGGGTAAAGAAGGCCATCAGAATTAAAAACAACTCTGCCAAGATACTCCAAGTTCACCTGATGTACTATACCAGAGCCTGGAGGAACAACAAGCATGTTCTTGAATGCAGTTGACCCCCATTTCAGAAACGCAAAACGCTCTTTGTTGCGATTGAACTCATACTGCATGTTAACCTGTAAGGCATTTTCTGTCCGTGCAACATCAACTTGGACTGAATGGTCAATAACAAGATCTACTGGAACCAGTGGGTTAATCTTATTAGCATCCCCTCCAAGCCTTTTGATAGCATCACGCATAGCAGCAAGATCCACAACAGCTGGAACTCCTGTGAAATCCTGCAGAATAACCCTAGCAGGCTTGAAAGGTATTTCTACTTGTTTAGGTGCAGTATTTTCCCAGTCAATAATTTTCTCCACGTCATCTTTCATTACTTGAAAATTGTCGCAGTTCCTTACAGCAGACTCCAACAGAATCTTGATAGAGTAAGGAAGAGTTTCGATTCGGGGATCCTTCAATGCTGCAAGGCTGTAATATTTACCAAAGATCCCACCCTCTGACTTTGGCAGGTCTGTCAGCGTGAACTTGAAAGGATTTTCTGCAGCCATCGTCGTGAGGCTCCTGGCAAGCCGCTGCAGAGGGGATGTGGAAACAGCCGACGTGCGAACAGGTGAAAAGAAAGAGGGCGAAGGTGGTGATGTTGAGAGATCCGCGTTACGAGTTGCCCCAATATGCAAGAACCCCCGCTGCTGCACACACGAAGCTGCTTTGCAAGAAACGGGAGCCGAAAGCACAAGATGGGTATTAGACGCTCCTCGGATCAGAGAAGATATATGCATGCTTTTCTGCCCAAATTAGCGGATCAGTGTAGTAAGCTGCCACCGTTCTTAATTTGCCATTCTAGGCCGCTTTTTCTTGTTTGCAATTGACTCGCCCACGAGCTGAAGCTCCGGGTAAATCTCCTTGAACTTCGCACAGAGATGCTTTTTTCTGAGCTTCTACACCTACCCACCTCGCGCTTGAAGCTGATGACCTAGATGATGACTTCGATTGCTCGCAAAATGCAGAAAACTACCCAACAGAAAGCTAAATTTAGCAAGTAAAGCTTCCCAGAACCCAAAATTAAAAGTAGCTTCAAGCCCAAAATCTAATTTTAATTTTAGATATGGTGCTTCGACAGGCAAGAAACCCTACGCTTGCGTGCTTGGCAGCTGCCCTTTTCCACGTGGCAGTTCAAAATCACACATTTCAGCCCATTGTTCTTGTTAGCACACATGAATCATGATAACTAGGAAGATGTTGAAATAG

**LIPOCALIN IN THE PLASTID (LCNP):**

TCCTTAATCACAAAAAAACATGGCATAAGCCCATCACACCAAAAATATAAAGAGATTAGAGTGAGTGAATCACCACTTTGCCTTAGTCTAGATATAAGTAAAAGCTAACTACGTAGTACAACTTTTCCCTTGAGAGTGCAACTAATGCAAATGTATGAGCCAACAGAATTGAAGAACAAGTCAAAAGCTTTCAAATGTATGAGCCAACATGAACTTAGAGCAAGTTACTCCGAAATGTAGAGGGGTGTTGAAGTGGCCAGCTCATGCATCACTTAATTGGCTAAACTAGTTTTTGCCATGTGCAAAGCCAATACACTTTGTTGTCAGAACATTGTTTTTTTTCCATAAGCACATCCAGGAATAGTTGCGACGACACAGGCCTTCATTATTTCCTTGTATAATTGACTTCATCAATGCAATATTAGTGCACATCCGAACAAGTCAATGCAGATGTGACTTCAGTCAGACCAAAGCACCCTTTTTTTTAAAAACAAAAACAAAGTGTTAACAATGTGTAAGTTATAGCATTGTTAGTCACAGCATCCTCCTCCATATCAAGAAATTGAAATCGATTTTTCCTGACTGAGAAAATATTTAATACCGATTATTTAAAGTAGAGTTCGACAAGCTTCTTGAGGGTATCAAATACACTGGTAAGTGGGTTTAGCTGCACAGGACGCCTTAGCCCCAAGTCCGGAAATTTGTTGATCAAAGCCTCCTGCATCCCTGGCATGGTCATCATCTGTTCCAGTTGGTCCATAGACATGCTTTCACAATCTTGAGGTGTATCCTTGATTTGATTTGTATTATATCCTACTGTTCCTAGGTAGTTCTTATATTTCTCAATGAACTCTGGCCCTGGATTTGGCGTTCGGGAATATATCTGCACAAAGCTTTTATCTTTGGCACCTGATACAAGAGCAAAAGTATCATAGTCAGTATTAATGACATCATATGGCTCCTTTGGAATAAATGGTAAAAAGGGAAACTTCAGGTAGCACTTGGAGGTAATCATCTCCTGTCTTTCTTCTTCGGTTTCTGCCATCGAAATCTCAGCATCACCCACGCATCTTACAACTCCTCTTATGCCTGTAATGTATCCATCTGGACTCCCATGTACACAGAAGGTATCAACTTGTATAGCTTTATTCTTCTCGTCCAAAGTATATACACCCTGGGTGCAATGGCAATCTTCTTGACCTACTCCTGCAAACCCACGCTTAAGTGAAGCAACTTCAAACCATCTTCCAGCATATCTTGCTGGATTGAAGTTCTTGGTAGTCATTCCCCCCATCACAAAAGGAGGCGCCTCACCATCAGATTGAGCATCGAGACTTAAAATCGTGTCAATTGGAGCTGCTGCTAGTTGACAGAAGTTGCCAGCAGAATGAGTAAGCACAGACTCAGCTAATGCCTGGTTTGAAATGGAGCTGACCAGCAAGGCTGTGACTACAGAAGAAGCAAGGACTTTTTGTGAAAAACTTTTCGCTTCTGACAGATCCATAGGCATATCCCAAGAGCAGATGATTGTTGTGCCACTCCCTCTTTCTTTTGTGCACCCATACCTTGGTTTGCAGAGAAGATTGTGGGCGCTGGAGCACGATAGCGCGGACACAGAAGCCATGAAACTGCGGGCAGCTAATCACTCATTACTCTTCTTGAAAACACACACACACTCACGTGAAATTGCCAATTGGTAGGCCAGACATTTATCTGAATTCCACTTCATTTTTCTTATATTAAAG

**EXECUTER 2 (EXE2):**

ACGAATTAGAGGAAGGCACGTGCTCACAGGCCATATCATCATTCGTGGATATGATGTTGGGGAGGCTGTGTGCTGCTGCTTGAGCAGAAGATTTTCATCGCGAGGCAAGGCTGTGTTGTTCGCTGCATCGAGACGGTAGCATTCGAAACCCTCTTCATCCCCGCGGGCTCTCTCTCCTTCACGAGGCCTTGGTCCCTAGATTGAGCACTACTCGTTTTCCCCGGCGCGGTGATGGCTGGAGCCGGCATGTCCACCGTTGGCCACATTTGCTCCGCGTATGTGGGCGATTCTCACGTTTACTCCCCTCCAGAGCCTCTCAAGCTTCGAGGCTTATCCCTCAGCTCTCACCCCAGGAAGCCCCCCCACCAACGCCTTTTCTGCCCCCGCCTCTCTAGGTCTCGCAGCGGCACCCGCTCCCTCAACAGAATGATTTCTGTTTCATGTAACCTTGATGGTCCCAGCGACCCATCTTCACCCTCATCATCATCATCATCCAATCATAGTCTACCTGAAGGCCCGCCACAGAAAGATGAAGGGGGCGAGGATCTTGTTTCCGAGCCACCTGTTATTTGGGACTTGCAGCAGCTCAAAGCTGCTGCTGCTCGAATTGGGGATCCTCTGCGCGAAATTTTAAGCAGTGCTATTAGGAGGTGGCAAACCTGCTTTGACAACTACAGGAAACAAAATCTCGAACAGGAAATCGAGATAGTTGAACCAGCAGAGCCAATTGTGGTGCAGGAGGTACCATGGGACTGGGAGCGATGGCGAGAGCATTTTGTTCAAATTGAAGAACATGAGAAGCTGGCATCCAATCTTAAGTTCCAATTAGAAGATGCAATTAGTGAGGAAGATTTCAAAGAAGCTGCACGGTTGAAAGGGGCTCTTAGTGCAGCAACTGCAAGTGACCCAGTTTCTGAGGTTCTTACTGAATTAAAGACGGCACTGGAAGATGAAAGTTATGAAACAGCTACAAGGCTCAGAGATGAAGCAGCTGCTGGGCTGTTAGGATGGTGGGTTGGTCTAGCAAAAGGTGAAAACGATCCTTATGGTCGCATTGTCAACATAAGTTCATCGTATGGAAGATTTATGGCGAGGGGCTATACAGCAAGGCAATTGGCATCTGCCTCTTCAGGCACTCCATTATTTGAAGTTTTTGTACGTAAGGACCCTGAAGGTTATCACCAACAGGCAGTATACTTGCAACGAGGGGGAACTGGAGAATCTCTATGGGGGGCACCAAACTCAATTGATGTCAATATTGATGTAAACATTGTGGACATGAACTCAATGACTGAGGATGGCAAGGAGATCAAAGCGAAAGATTTGACAGAGGCCTTTGGAGATGAAAAGGGAAAGAATGGTGATGCGGCGGTGTTGGATGATGGATTAAATGGCATCATAAATTTCATTAAAGATCGAATACCTGATGTAAAACTGAAGGTCTTCAGAGTTATTGCCCAGAATGGTGAGGATATTCCTAAGATTGTGGAACAGATTTTAGAAGAGGTTGAAACACGGGAGGGTGAGGGTATGAAGCAAGAGAATGAGGGAGATGCCCTGCAAGAAGCGAAGGAGAAAATTGGAGACACTGGTGAGGCGGTACAGGAAGTGCAAAGGGAGATACCCATCAGGTTGATTGTAGGTGGGTCGCTGCAGAATGCTGCTGATGATAAACTTCCCAAAGCTCCCACAAGGAGGCGGGCAAGAATTGAGACTCAAAGCCGAGATTCTTTTCTGTTCCACATTGATGACATTACTGAGCAAATGCCTATCAAAGAAACAGTAGTTTCATCAAAAGCTGCTGCAGTTGTGAATACACAGGCTTCAGCAGAACTCCCTCCTGAATTCTCAAAAGTAGTCTCAAATCTTGACAAAATTCCAGTGAAGGTGTCCAAGGATATGGGAGAGCTTCTAAAGCTTGCCATCAGCCAAGTGCAAAGACGACGTGGCTTGTCTAAGTGTACTTCTTTTGAGCGGATTCAGCTGACAGGCTCCAGTACAGACCCTATGATAGGCTTGTATATTGGTGCTTTTGGTCCTTACACTTCCGAGCTGGTTCAATTACGGCGGAAATATGGGCAATGGCGTGAAGAAACTGACATAAGCAACATAAAAGATAAACTTGAATTTTTTGAATATGTCGAAGCTGTAAAACTCACTGGAGACTTGAATGTTCCTGCCGGTCAGGTCACTTTCCGGGCTAAGATTTCCAAGGATAGTAGGCTTTCTCACCGTGGTATATATCCAGAAGAGCTGGGAGTGGTTGCGAGGTACAAAGGCCAAGGCAGGTTAGCAGAACCTGGGTACAAAAATCCCCAGTGGGTAGATGGGGAGCTTTTGCTTCTTGATGGCAAGGGATTTGGCCATTTAAATGGTGCAGAGCTTGGCTTTGTCTATTCAGTTCCTGAAAGGCATTTTCTGGTCCTGTTCAATCGATTGAAGTTGCAAGTTGAGTGATGAGTGTCCTCGACAAATCCCTGGCAAAATCAACACCTCGCTCATTTGCAGTTCTCATCACATCACTTATATGTATGGAAGTCTGGTGCGTGTGACCGTATCATGTTAAGAGAGAACATGTACAGCTTAGACCCTTTGACAAGGTTACAGAGTATCACTTTCGATATGTAATGGCCATACCTTTTGCTGCAAGTGAGCTTCTGGGCCACATTACACATGTTTGTTTCCTTGGCATGACGAAGGCCTCTAGTGACAACACCCGATCTCAAAGGTTCACAATAGCTTGTACCAGAAGGGGTGGTTAATGGGGTTGATTTACGTGAGCTTAAGTCATCCGCTGGTGATCAAGTAGTAAGAGTCAAAATGTATAGAGCTGTCATACTCCACATTAGACTGAGTCTAATGGCACAGTGAACTTGTACTGAGTTTCACATTTGCAGAGTTTTCCGTGCGACATCCATAATAAATTTTAGGGGAATCTTTCCACCTGGTCTACAGTTCTCTATTTTGAAAGAATTCCTCAATTATGGATTACTCTGTCAACTGCATGAAGGCTTTCGAGGAAAAAACGAATGTACAGGGACTATCTTCTTTTATGGTATAGCTGTATTATGTCTCAAATGTTTGAGACACTTGTCATCTTTAGTAGGTTTACAGCCCTGACACTGAAAATTTTATCAATACTGAATGTCATGAGTTACAACGTGCATTTACTGATTTTTCTGTTCTATAGAGAAAAATGTGTGAATATGGTTTCTTTCAGGATGTAATTTGGCTCGGCATCAGCGATATTTAATAGAGCTGTTGCAACTATCAACAAGTCGG

**ACTIVITY OF BC1 COMPLEX KINASE 7 (ABC1K7):**

CAAATGCAGACTTAAATTGCTTGTGGACTGCCTGCGGCCCATAGCCATCTCCTCGCTCTCTTGTGAATCCTCCTCTGTGTATTCATGGATTCTGTTTGCGTCTCTCCTCATTCATCCCCTGCCCACCTCAGCTTCTCTTCCCAGTTTTCAAATGGAGATAGTTCTCTCAGAGTGCAAATGCGCAGCAGGCGACAGCGCTCTCGTTGCTCTGATGGGGATGGCATTACCAATGGGTTAACTCGAAACTTAAGGGTCTCTGCTAGAACCAACAGTAGCAGTGGAGAGGTCGCGTCGTCAACGCCCACTTCAGATAACCCTGCAGTCAATGGATACACACAGAATGGTAGGAAAAGTAGTGATGCTGTTGGTGTTAATGGGGCCTTGACTGGAAAAGGAAAACAATCTCTAGCTATCAACGGACGTCAGTCCCGCATGGTTCCTATTGCAGAAGCATTGAAAAAGGAATCGGTAGGAAAGGTGAATAGAATGAATGCGAATGGTGCTACTCGTAATGGAGCTGTTAGTAGCCTGAGTACGGGCAACAAAATGAGCTTGGCTCCTAAGAATGGTGCAGTTAATCAGATAAATGGGACTAAGGTGGATGCACTCTCCAAACGACCTTCTCCACGACGATTGCAATCCCCCTTGGAAAATCTTGCATGGGATGACAATGTCAAGGTTTTACCTTCTGATGAGTCTTTTAGCTGGTCCAAAGAGAACTACAGCAATCTACAACGGACGATTGATGTTTGGAGCTTTGTTCTGTCTCTAAGAGCACGCATATATTTAGATGGTGCCAAATGGAGTTTCTTTGGCGGTTTCTCTGAAGAAAAGCAGAAAAGCAGAAGGCGCAAGACAGCTTCTTGGGTGAGGGAGCAGATTTTACAGTTGGGCCCCACATTTATAAAGCTAGGGCAACTTTCATCAACGAGATCAGATCTTTTCCCAATTGAGTTTGTTGAGGAACTTTCTAAGCTACAGGACCGTGTCCCTGCTTTTTCTCATAAAAAGGCTGTCGCACTTATCGAAAAGGAGCTGGGGCAGTCTCTGGATGCACTCTTTGTGCTGTTTGAAGAACAACCTATAGCTGCAGCCAGCCTTGGACAGGTGCACAGAGCAGTTCTACTTAATGGTGAAAAAGTAGCCGTAAAGATACAAAGACCTGCGTTGAAAGAGTTGTTCGACATTGATTTAAACAATTTGAAATTGATTGCTGAATATTTCCAAAATAGTGAGACCTTGGGAGGTCCAACCAGAGATTGGCTTGGTATTTACGAGGAATGTGCAACTATCTTGTACCAGGAAATAGATTACATAAATGAAGGACGCAATGCTGACAGGTTTAGACGTGACTTTAGAAATTTCAAATGGGTCAGAGTCCCACAAATATTTTGGGACTATACTTCGCGTAAAGTTTTAACATTGGAATATGTTCCTGGCATCAAAATAAATGATTTGGCATTGCTGGATGCGGGAGGTTTTGATCGTTCTAAAATTGCAAGTAGAGCTGTAGAGGCCTATCTCATTCAGATATTGAAGACTGGCTTCTTTCATGCAGATCCACATCCAGGAAACCTAGCTGTTGATGTGGATGGATCTCTGATTTACTATGATTTTGGTATGATGGGAGAGATAAAGGCATTCACAAGAGAGAAGCTACTTGAAATGTTTTATGCAGTGTATGAGAAGGATGCTAAAAAGGTAATTCAGTCCCTCATTGATCTTGGGGCCTTGACACCGACCGGGGACATGGCATCAGTGAGAAGATCAATTCAATTCTTTTTGAATAATCTGATGAATCAACGACCGGATGAGGCAAGCACTGTCTCGGCCATTGGTGAGGATCTGTTTGCAATAGCTATCGATCAACCATTCCGGTTTCCTTCAACTTTCACCTTTGTCTTGCGTGCCTTTTCTACATTAGAAGGCATTGGGTACACCCTTGATTCAGAATTCTCATTTGCAAAAGTTGCTGCCCCATATGCACAGGAACTCTTGGATGTTAAAAGCACGCAGAGAAGTGGGAATTATTTGATTCAGGAGTTGCAAAGACAAGCCAGTGATGCAAGAGATGCAGCTGTTGCAATGCCTACGAGAATTCAACGAATTGATGAGCTTTTACAACAACTGGAGGCTGGTGATTTTAAGCTTCGAGTGAGAGTTCTTGAGGCAGAAAGGGCTGCTAGAAGAGCATCAATTCTTCAGATGGCAACAGTCCACGCAGTAGCTGCAGGAACACTGTTGAATGTAGGAGTTTCTTTGAATGGTCTAGAGCTGAATCCCTTTGCAGTCTCCTCATTTATTGGATCAGGTGTGTTTACATTGTTTGTGATCCAAGCCATGAGGAGAATTCAACGCCTAGACAAGTTTGAGAAGATGATATAGTTACTTAAATCACTTTTGATATTTAGCAATTGTATGATATGTAAAATGTTGTGACATCACAAAATTTATTGCTAACTCTTGTTCTTTTCGGGTTAGCTTGTAACTGCTCTCAACCCCTATGGGCAAATGCCAAAAACAGGGGTTGAATTCCTTATTTTTTACATTCATAAAGAAAAATTGTTCTCATCTCTATACACCAATTCCTTTGAACATTTGATTCCTTTGCTATGATATTGGAGTGGATTTGATTACTCCATACACCAATTCCTTTGAACAATTGATCCTTTGATAATATTGAGAAGTAAGAAGGCCACCTTCTTGGGCCATTGGACAATTGATTCCTTTGCTATGATATTTAGAATGCTTGTGAGTT

**ACTIVITY OF BC1 COMPLEX KINASE 8 (ABC1K8):**

GGCATGATTAGACATGAAGCTTAAACCTCAACGCACAACCCAATGCCACCTTATCTCTTCTATATACCTGACCTTGGTTGAGCTCATTGAACCTTTTTACAATTGGCTACAAAATGTGCATAGAGAGCACTAATATTACAATAGTTGGTTGAATGCTTGGAGGAACCAACTCTGATATGCGAGCTTAGATTAGACGGATTACATTCAGAATAAAAAGTCAATTCTAAGCTGTTCCTGTTATAAGTTTCTCCTGTTTGTCCAACTTGTTCACTTTAAGAAGTCCTACCAATACTTTCAATCCAAACAATGCACTCCCAATAAATGCAGCAAGAGCAGGTACCCGAACAGCATTTATGTGAAGCAATGTTGCCAGGTTAAGTAAGGTTCCCGCTAGGATAGCCTGTCCTATGGTCTGTTGCACTGTAGCCACTCGCTTGAAAGCTCTTTCCGACTCCAAAGTCCGAACTCGAAGCTTCAAATCACCTTGTTCAATCCGTTGAATAACTTCTGCCAGCTTCTCCACTCGATCAGCTTGACGAAAGAGATTGGAGAAGGCCTGAGCCTGCCTGTTCCATCGTTTCTTAAAATCCCTTATTAAGACATCATAGCCAAACTCCCGGAACTTCAGAATCTCCAAAGCATACGGTTTAGCGATCTCTGATATGTCGAACCTGGGATCTAGGCCTTTACCTATTCCGTCTAATACTGAAAAGGCTCGTACAACAAATGTGAAGGTTGCAGGAAATCGGAAAGGTTGGTCCACAGAAATAGATAACAGATCCTCCCCTATAGCAGCAAGGCGCAATTTCTTCTTTTCGGTTTTCTCCTCCTTCGATAAAGGTTTCTTGAATCCTAATTCAGTAGTGGCTGCTTCTTTCTCATCCCTTTGGGCAGCTAGACGATCTTCGAAACTGTTCAAGAAGAATTGAGCAGTTCTGCGAACTGCAGTCATGTCCCCAGTTGGGACCAGAACTCCCATTTGAACCATTGCTTGAAGAACCTTGTCAGGATCTTTCTCATAAACTCCATAAAATGCCTCAAGGAGGCCCTCCCTTATATTTGGGCTAATGCTGCCCATCATTCCAAAATCATAAAATATCAATCGCCCTCCATTTGCATCGTCAACAGCAATGTTACCTGGGTGAGGATCTGCATGAAAAAATCCATGGCGCAGAATTTGTTCCAAGTATGATTCTACCGCATACCTAGCCAACTTATTTCTATCTACACCAAGTTGATCCAAAGCTGCAATTCGATTTATCTTGATACCCGGAACATACTCCATGGTAAGCACCTGCGGAGTAGTAAATTCCCAACATATTTTGGGAACCTTGACATAGTCTAGGTCTTTAAAATTGTTTGCAAAGCGCTCAGCATTGGCTCCCTCTTTGCAGTAGTCAATTTCCTGATACAATACAGAAGCACATTCGTCATAAATTGCCACCCAATCCCGCTTCGCTCCATCAGACTTCGGATCAAGCTTTTGTAAGTTTTCTGCAATGACCCTCAGATTTTTCAGGTCAATGTCAAACAAATCTTTTAAGCCTGGCCGTTGAACTTTAACGACGACTTCTTCACCCCTAAGAACAGCGCGATGAACTTGACCAAGACTTGCCGCAGCAATAGGATCAGGGTCAAACTGATCAAAAATTTGACCAACGGGAGCACCTAATTCCTCCTCAACAATCTTCAAAGCAGTCTCAGATTCAAACGGAGGCACTTGATCCTGGAGTTCTGCAAGCTGGTCCACGTATTCTTGTGATAGGATGTCTACTCTGGTGGAAAACTGCTGCCCAATTTTAATGAAAGTAGGGCCCAATCTTAAAAGGCCCTCTTTCAGCCACTTTGCTAGAACTTTCCGCTTTTCCTTCTTTTTTTCCTCTGTTATCCCTCCTTTGTAAGAAAATTTTTGGTTGATGAGCCAAGCCTTAAACAGAAAGGTAGCAACAAAGAACCAGATCTCACAAGTTCTTCGAATGGTTGAGTAGGTCTTGAACTGATTCCATCTACCACCTGCTGCAACAGAAACCGGAATTTTCTCTCTGCCTGCTTGCTTGAACCAAGGATCCTCTTGACCAATGGATTCTACTTTGTCCTTAGTTCTGAATTCTTGTGCTTGAGGAGTATCAAAAGTAGGTGAAAAGAGCTGCAGAGCTGTGCCTGAGCGGCCCTGAAACTCCATATCCAGAGATCCTGCATTGGCTCTAACAGAAGCCCCATTGGTGCCAGCAGCTGCAGGTCCGTTCTCTGCAGTGGCATCAGCCTCAATAGCTGCAACTGAGGGCTCCTTGCATACGACTGCAGTTGCAGATCCGTTCAATCCTTTTCTGTCATCCCTGGATGCAAAAACCACATTCCCAGCACCATTGTTCTCTCTGTAAATGGTGCCATTGTTTTCACTACCTGAAGCACCATTACCTCCATCGTATGCTGCACCAGTGCTTCCGCTGGACGTGGCACCATTGCTTCCTTTATGTGCCACACCATTGTTTTCATGATATGCAGCACCATTGCTCCCAAAACTAGCGGCACCATTGCTTCCATAGTTAGTGGTACCATTCAAGCCCAGTGAAGAGCTTTCTTCATTTGCTGCAGTATGAATCCCTTCATGTAATAATTGATCATCTACACCTACAGAAGCCACTGCTTCATGTGGGTTGCCTTCTTCTTCCACAGTTGCTACCATCTGATCAGTTAGGGCCCTTGCTAATGTCCTCCTGTGGTTTGCCCTTTCACTTCCTCTTGAATGAGTCAATACATTTCGGGAAGACAACCCTAACCCTAACCCTAAACCGCGCCCTGGAGCTAGCTGTGTCCTTGACGCTTTCTTTACAACAAGCCTCTGCAGAAACAATGAATTGGCACCATGGGAGCTAGGAAAAGGCAGCAAAGAATCTGTTGCAGCTTGAACAGAATGAAAAACAGGGATAGAAACCATGAAAGCAGCAGCCATGGCTACCGATGCTGGAGAGAGGCGGAGCTCTCATGCAAAATTGGATACAGGGGTGATTTGATCAAACTGCTTCAGTAGAAGGATCGTGAATCAATCGTACCGTCAATCAACGATCCTGCAGGCTTCTACAAACAATCGATGAGCCTGCAGGCTGCGATAGAAACCTGCAGAACTCAAGCTCTGCAGCACAACTTCAC

**ALKALINE/NEUTRAL INVERTASE A (A/N-INVA):**

TTTTACTAGAAAATGAATATTTCTATTAAGTGAAGTAAGTTGTCTTCGCAATAATAATCAAGATACAAATTTTATAAATCCCATCTTACAAATTTGGGATCTGTTTGGAGCTTTTCACCAGAAAAATATGGAAAAGCCTCTCATGGCAGAGACCCACTCTACTGGGACCTGCCCCCAGCCAACCCTTCAGTTGGTGAAAGAATGATGAACCAGCATTTGGTTCTATATAATTGAGACTATATGAAAAGCAAATAAGTTGTTAATTTCTCGTTTGTGACTGTACTTTGCAAGCTTTTAAGCTCTGCTGCTCTATGTACAAGTATTCGTACACGACTACTGCCCAAAGCCTCAAGATGGTTAGACCTTTCGGCTTCTCTTTCGCCGAGGGTTGGAATCAAGAGGGCAGGACATAGCCTCGAGCAATGACAAGTCTTCTTCAAAGGTGAAGATTTTAACTGCATCAGGGTTATCCAAAAGCATTTTTGCCACTACAAACCCTGCTATTGACCAGGTCTGGAAAAGTCTAGCTTGCTTTCCTATGAACCTTCCCGTGCGTGTGTCGTAGTATTCAGGCCACTTATCCTTGGAGAGCCGTTTCTCTGCCACTTTCAGCGCCCTTCTAGCAAGATCAGCCCTTCCCACCTTCAAGCAGGCCACAGTGAACTGTGTATACACGAGTCATATTTATCAACATTGAATGACAGTAGAAGTCATTAACAAAGAAGATGTGTCCACATCATTAAAATGGCTCATACTTGTGCATGCATCCATGCCATTGCATATGAAAGTTCATAAGAAATCAATGCACTTGTGCACGTTATTATGTGCAAGGGTCGGAGCCTGGGCATTCACACAAGGGTATACAAATTCATCACACTAATAAAACCCACCTGCCATATTAGAGTAGGCCAAGATCCACCATTATGATAAGACCACGGCCTGCACAACAAATTCAGGTTGTGAATACACTGATGATTGGCAAGAATAAGACAAAAGCTCTTCTGCTGAGCAAATTTTAAAAAAAGATCATACGTATTCTTTGGATCACTGCCAGTAATAACCCGCCACTCATCATATTCCAGTGCTGGAAAAGCGATTTTCAAAGGCATTGTGCCAATAAGATCGTCCCATTTCTCCTCAATGAAATTCAATATTTCCTCTGATTGATCTTGAGTGGCCAAAGAGCCAACAATGGCCCATAAATTGCCAAGAGAGAAGAATCTGAAATCCATGTGAGCTGGCTGCAAATTCCCAAGCATGTAACCTCCTTGATTGGGCATCCAATCAATCAACCAAAGGGGAATCTGTTCTGGGTAGATATTGAACTTGTTGACAGCATCATGTGAGTACTCCTCCGTCTTGTATCGATAAATTTCATTTAGCTTCCTCATGTCTACCCAATAGTGTTCCCGTACATGGTACGAAAGAGCACACAATCTGCTATTTAATGCCCTGAAGAGGTCACCTGAGCCATCTTCTGGAAGAAGCATCTCTCTAGCGCATCGGAGCGCTGAATAGAATAATGACTGCAAAAGTAACAAGCAAGCAACAACAATTATTAACATTCTGAAGACAAAACAGCAAAAACTTCTTAAACAACTTACTTTAACCACACACCTTGTCACATTTTCACAAATTCTCGGAATCAGAAAAAGAAGTTAGTTTGCCAAACCTGAATTTCCAGTGGGTGGCCGTGTATACCCATGCGTCGATCTATCATGCAGGACCCATCAGTGACCAGTAAGGTTGGAAACATGTCAAATCCATCTGCTAAGCACAGCTTGAGGATGAGCTTTATTCCAGTTTGAACATCTACTCTTTCTTGCAATCCATAATCGCCAGTATGTTTGCAGTAAGCTCTCAACAGGATAATCCACCAAAGTCCTGAATCAACAGGGGCCACACGACCTATTGCAGCCTCGCCAAAATCAGGATCTAACACCTCCTCAGTTGCACCATCTTCGCTCCCTTCTACATAAACTGTTTTTACTTTGAAACTTGCAGGCATTAGACCCCGTCCAGGGCTGTAGCAATCAACTGTTTTCTCCCAGCTCTGTAATTGTAGTGTATGGAGAAGAAAATTTCGGACAATATCAGGCTCTCCCTTCAGCAGGAAAGCCAGAGCTGAAGGGATAAAATCTCGTACAAAGACTTGATCATAATTGAGAGGAGTGGTATCACTAGGATCATTCGCTGCAACAGTTCCAACTGGATTGCCACAATAGTTTACAACAGCGCCCCTCAAAAGAGTCCACGCCTCTTCCTCAGTTGTGTCCTCCTTGGACAATCTTGACTCCATGAATGATTCCTCAAGAACAGCATCTGTTTGATTCTTTGAACATGTCTTCTCTGGAGTCACAGCAAACTCTATAGTGGCAGGCTCTCCAGAAATCCATAAAGGATTCACGGTAACACTGGACACATAGAAATTGTCTGATAGAGGCTCTGCAATGTGTATGTGCTCCACTTTACTCAGCCTTGTGGCCTCAGAAAAGGCAGCACTTGAAAATACGCTGCCATTTCTGCTACCTTGATCCGCTTCGCATCTAACCTTACAAGATTTCCTTTGACCGCGCCTCTTCGATTTGCATCTTTGCACAGTAAAAGACAGACTAACATGCTGAAGCGAGGACCTTAAAGACCTCCCAGAGAGGTCATTAGACAAACGATTCCTTGGACGCGTGGACTGATCACCAACACTCTTGCGACTCTGCATAGTACCATCCAGAGAGCTATGCACAGTAAGGGGTATGCTTCTAGTGGTATGTCGGCCATGGTGTTGTAAACTGCCCAAACAAGCTGTATTATCCATGAGTACACTCTCAGTTCTGCTAAGAACCTGCAGTTCTAGCCCCTCCTAAAAGCCTGCAAAACTTGATTGAAGGATCGCTTTGATACGGAGGAAGGCTGCTGCAAAAGGCTGTAGATGGAGTTGTAACAAATCTGAAATTTCAAATGCCGATCTGCTACAAATTAGACTGTTTGTGAGGGTGAGCGGGTAGCAAGGCGTGGCGTGTATGTGTGTGTGTGCGTGAGCGTGTGCGCGTGTGTGAAATAGAAAGAGCGTATGTGTGTGTGGGTGTCTGTGTATGTGCAACAGGTCTTAAGGTTTAGCGATGGCGGAGAGAAGTAGACGCGATATCGAAAAGGAGAGAAGAGTGCTGAAAGGAG

**ENLARGED FIL EXPRESSION DOMAIN 1 (ENF1)**:

AACATCTTATTATAAGATGTAGACAAAAGGAAGTAGTGAAGATGTTGCGAGTCACTGTGCTTTCGCGCAACACAATGATGCACTCTTGCCTGCGCAGGCTGGGTCGCCCTCGCCCCAAGCTCTTGGCCCAGGTGCGAGCATTGACATCATCACCTCAGGAGGTAGCAATAAGCAAAATCAAAGAGGCAGGCTTGCTGCATTCCAAGGCCTTAATTGGAGGAAAATGGGTGGAGGCCGAGGATGGCAGAACCTTACCGGTGCATAATCCTGCTACAGGAGAGGTTCTTGTAGAGGTTTCATGCATGGGTGAAGTTGAAACCAAAGCTGCCATTGATGCAGCATCAAAGGCATTTCTAAGCTGGCGTGAGCTTACTGGATCTGAGCGCAGCAAGTTGTTACGGAAGTGGTATGATCTTCTTATTGCCAACAAAGAACATCTGGGTCACTTGATTACATTAGAGCAAGGAAAGCCTCTTACAGAATCTTTTGGAGAGGTGGTCTATGGATCGAATTTTGTGGAATTTTTTGCTGAAGAAGCCAAACGCATTTATGGAGATGTCATTCCAGCTCCAGTTCCTGATAGACGACTTCTGGTTTTGAAGCAGCCTGTGGGTGTTGTTGGTGCAATAACACCTTGGAATTTTCCGCTCGCTATGATTACTCGCAAGGTTGCTGCAGCACTCGCAGCTGGGTGTGTTACTGTCATCAAGCCCTCTGAGTTGACACCACTGACTGCACTAGCAGCTGCCGAACTTGCAGCTCAAGCAGGCATTCCCCCTGGTGTGTTGAATGTTGTGATGGGGGATGCTCCTCGAATCGGTGATACACTCATGGAAAGCACTCAGGTGAGGAAAATCACCTTCACGGGCTCTACAGCTGTTGGAAAGAAACTCATGGCTGCTGCTGCAGGGACTGTTAAGAAGGTGTCTTTAGAGCTGGGTGGAAATGCACCTTGTATAATCTTTGATGATGCGGACATTGATGTGGCAGTAAAAGGCACAATAAACTCAAAGTTTCGCAACAGTGGGCAAACATGTGTTTCTGTTAATCGCATATTTGTCCAAGCTGGTATCTATGATAAATTTGCTTCTGCATTTGCAAGTGCTGTCAAATCTTTGGAGGTTGGCAATGGCTTACACAAAGGTGTTAGTCAGGGTCCTTTGATCAACGAGGCTGCAGTTGGCAAGGTTGAGAGGCACCTTGAGGATGCAATCTCAAAGGGAGCAAAGGTGTTGCTAGGTGGAAAGCGGCATAATCTGGGTGGTACTTTTTATGAACCAACGGTTGTCAGTGAAGCAGCAACCAATATGCTTATCTTTAGGGAAGAAGTTTTTGGACCCGTGGCCCCTTTGATACGATTTACAACTGAAGATGAGGTTATACAAATGGCCAATGATACTGTGTTTGGTTTGGCGGCATACTTCTTTACCCAAAATATAGCAAGAGCATGGCGTGTTGCTGAGGCATTAGATTTTGGAATGGTTGGTGTTAATGAAGGTCTCATCTCTACAGAGGTCGCTCCATTTGGTGGTGTAAAACAATCTGGCCTTGGCCGGGAAGGATCAAAATACGGCATTGAAGAATATATAGAAATGAAGTATGTATGTATGGGTAACATTCAGAAGTGATGGAAGAACAGCAAGTCTAAGGTGACTCTATCTCAAACCCCTTGTCCATTTTTCGCAATCACAGATTGACCAAATTAACATACAGGTCTTGAAAATACACCAATGAAAATCTGGTGCATGTGTATCAAAATATATCATCCAAAGTGTAGTGTAAGTAAGCTTGAAACTTTGTCATCCATAAAGAAGAAAATACTTCTAGGAATTCTTTGTCATTGATAAGGAAATTGCAGTGCCTATCCTTGTAATTAGCCAGCAGTATCAGCTATGTTGTTGTTCAGTATCAAAAACTGTTTTGTAGTTCTTTTTTAACTCTTTTGTGAACAGAACCAAACAAAACAAATATATATATGTTTTGCATTGAAGAAGACAAGAACATGTAAAAAATTGCTCTATGTCGGGAAGCTTTCTTCCCTCTAGATCGTACACAAAGAGAGCTTACACACACACACACACA

**Γ-GLUTAMYL TRANSPEPTIDASE 1 (GGT1):**

AGAGAGAGAGAGAGAGAGATTTGCTACTTCTTGTCATCTGCTTTGCATCTGTTACAATGCCAACATGTGCCTCTACTTTGTGGGTCCTTCTCCTCATGTTCCTTGCCACAGACACCCTTGCACTTGCAGCCCGTGTCCCATTGAAGAGTGGGACGCCCGGTCGCGGATTACAAATCGGCAGCAGGGTTGCTGAAGGGCTTCTAGGTGCTGTCACATCTGATGATGGGCGGTGCTCCGACATTGGCCGTGACATTATGGCGGATAAGGGTGGAAACGCTGTTGATGCTACGGTGGCCGTTGCTTTATGCATCGGAGTGGTGAACCCAATGGCTAGTGGCATTGGGGGCGGTGGGTTTCTCCTCATTTCCCTTGCAAATGCCTCCTCCACTCTGGCATATGACTTTCGAGAGGTCGCCCCTGGTGCGGCTTCTGAGAACATGTACTCTGGAAATTCAGACTTGAAGGAGAAAGGGGCTTTGTCGGTAGCTGTTCCAGGCGAGATAGCGGGCCTGCACTTATTGTGGCAACATTATGGTACACTGTCTTGGAAGACACTGATGGAGCCAGCTATCCAGCTTGCTGAAAATGGTTTTGAGGTAAAGCCATACGTAGCAGCAGCAATCAATACATCTTCATCTTACATTCTAGCTGATGAAGGCTTGAGTGAGGTGTTTGCTCCCAATGGGACCTTGCTTGTTGCCGGTGACATCTGTGTCTGCACAACTTTGGCGGCTACGCTTACTTCCATAGCTCTTGATGGCCCTGCTGCCTTTTACAATGGTTCCATTGGGGAAAACTTTATTAATGACTTGACAAGTGCTGGTGGGATTATTACTAAGGAGGATCTCGCAGGATACAGAGTCAAGATTCGCGACCCTATTGTGCAAAATATTCATGGGTTTACAATTTATGGTATGCCTCCGCCATCTTCGGGGGGTGCTTGCGTTGCTATGGTGCTTAACATCTTGGCTGCTTATCTAGACCCTATTCAAGCAGTGATGGGGCCTCTAGGGCTGCACCGAATGGTTGAGGCGACCAAGCATTCCTTCGCACTTCGAATGAACCTCAGCGATCCAGATTTTGTTGATGTGACAGATGTTCTCAACAGCATGCTGAATGCTACCTTTGCAGCTGAACTTCAAAAGCTCATCAATGACAACAGGACATATCCTCCTACATATTACGGTGGCAGCTGGAGGCAACTAGATGATCATGGCACCAGTCATTTCAACATAGTGGATAAAGACCGCAATGTGGTATCCATGACATCAACAGTCAATTACCCATTTGGGGCCAAACTTCTGTCAAAATCTACGGGAGTGATGATGAACAATGAGATGGATGATTTCTCAATGCCCAGCAATGACTCTGGAACTACTCCACCAACCTCTGCTAACTTCATCCAGCCTTACAAGCGTCCCCTATCATCAATGGCGCCTACTATTGTCATGAAGGATGGGAAGTTTCTAGGAGCCCTTGGAGGGAGTGGAGGTGTAAGGATCATCACAGGGACAATTCAAGGGTTTTTAAATAGCTTTTGGAAAGGTTATGATCCCTTAACATCTGTGTCAAGTCCAAGGATCCACCATCAGCTGATGCCTGATGTACTGGAGTATGAGGATTGGACAACAGTGAGAGGAGAGGTGATCGAGGAGCCAGAGGAGGTGGTGGAATATTTAGAGAGCAAAGGCCACAATATGACAAGCCATAATGGGGCTATTATACAGCTGATTGTGCAGAATCTTAGTGAGCCACTTACCTTACACGGTATCCACAATGGCCGTGTTATGAGGGCGTTGGGGCCTTGGGTTGGCACACTCACTGCTGTGAGTGACCTTCGAAAGGATGGCTACCCATCTGCCTTCTAACTTCATTGGGACTTGGGGCATACTCAGTGCATTCTTAATGCATTCTTAGCATTGAGACCATGTTTTTGCTAGAGTCCAATTTGGGCACACACTAGACAGAGTCTTGGCATTGGGGTAAGGCCATTTTTTCTTTCACATTGCAGATGCATATGTATACTACATTGTGTAGGAGGGGACTAGTGTACTTGAATGAATTGGGCTAGCTCTACCTCACCATTCGGTGAAGGATCGGATATTATGCAGCTTTCTAAAATCATTAGTGAACTCAGCGCCTAAACTTTTCGCCTGCTATCTGTTTATTCTCATCAAATATCTGTGCGGCAGTAGAGCGTGAAGGTCAGGGCCAGTATTGAGGCTTTTTTCACCTCTTTGCCACGATATGATGTGCCATTTGAAGGGGTGGGAGGGGGCTCTTGAAAGGCGGGTGCCTCTTAATCGGCCCCTTTACCCTGGTGGGTCCATATCTTTGCCAGTGTGCAGTTGTGCCATCAGCTGCACCGTCAAAGTCCAAAGTGGTCTGTTCAGCTATTGCTGCTGAATATCAGGCTACACAGACTCATTTGTATTGAAAGGTAGCTTAGTCGAAATTTATTATGCGTCAGTCTTAGCTGTATATAACACATCATTGGCATGCAAGTTACATGATACCTGTAATTACCTGAATGGACACTTATAAGTTAATCTTGTCTTTGCATTTGGGAAAATCCACTGCTTGCTTCCAATGTAGCTGATTGGTTGCCATGGAGAACAAGACATTGTTGGAGAGACTCGTTTCCTCGCACGTACGGCATTGCACTTTCAGAAGGCACATGCGCAGAGTAGGACATGGTAAAACATGTGGGTTTAACAAGCTTACTAGTATCAGTAGAAGCTGTTTCTATATTTCTTGGGCTGGAATTGTCTGTGAAGCAAAGCGCATGAGACGTATTTTTCGCAAGCTG

**GLYCERALDEHYDE-3-PHOSPHATE DEHYDROGENASE C SUBUNIT 1 (GAPC1):**

CAACTGCCTTGCCCCTCTTGCCAAGGTTATAGATGACAATTTTGGAATTGTGGAGGGTCTTATGACTACGGTCCACTCAATAACAGCAACTCAAAAGACAGTGGATGGACCATCAGCTAGGGATTGGAGGGGTGGCAGAAGTGCTGGCTTCAATATAATTCCCAGCTCTACTGGAGCTGCCAAGGCTGTTGGCAAGGTGCTGCCATCATTGAATGGAAAGCTCACCGGCATGGCATTCCGTGTGCCAACAGCTGATGTCTCGGTTGTCGACCTTACCGTACGGATCGAGAAAGCAGCTTCCTATGGAGACATCAAAGCTGTAATCAAGCGAGAAGCAGAAGGCAAGCTGAAAGATATTCTTGGCTACACAGAGGATGACTTGGTTTCATCGGATCTTATTGGGGACAGCAGGTCAAGCATTTTTGATGCTAAAGCAGGCATCGCACTGAATGACCACTTCATCAAGCTAGTGTCGTGGTATGATAACGAGTGGGGCTACAGCAACCGAGTTGTGGACCTGATCAAGCACATCGCATCATCTGCCCAGTAGTGAATTTCAGGCGAACATGGAGAAATTGTCATTTACTCATGTTTCGAACTTTGATAAGATGCAGCACTAATTAGAAATGTCAGAAATAAATGAGAGCGTTTTGAGACAGCGCTTACCAAATGAAAATGTAGTCTTTTTTGGCATGCCCGTAGGGAGGTCTATGCAATGGGGGGGGGAGGAAAAAATTTCTTAAGGAAAAAGCATCATTAGCTTTGGATTCCTCAAAATAGCATTTAAATGTCTTGAGATGTGTCCTATGATGGTTGGACATTTTTTTACTTGCATTGGGTGAAGAAACATTTCTTTATCATTTTTTTTTTTTTTTTTGAGAAAAGACCCTTGCTTTGATATCAATATTACATTACAACATGTACGCGCGCATGTGCTCCATAATTACAAGGCTTGAGAGAATTCTCGGGAAAAGGCACATTTAACCAGTGAAGGGACTAAAAAAGAGAAACCTCCACCACATTCCCTCCTCTCCTATCTGGGAGAGATGCAACAAACATTGGAAGTTTCAGAATGGTGGAGGGAAGATTTATACAGGTTCTCTTTCCATAGGTATCAGAATGATGAAGTTTCATCCCAAAGCTGAGTTTTGTGACAGATGGATAGGACATATTGAAGATTTACAGCATCAAGATTTACAACATCTGATCAGGCATATAGATGTTGGTTCTTTTGGTGCATTTGGTGCACAAGGAATACGTTGAAGTTCGATGGTTTAACTACTTATCTTCATTTCCCCAGAATCACATTGAAACAGAGAGTATTAGAGGATGCTATGCCTCCCCAACAGATTTAATAAATTTACATTGTCTGCTTGCTTTGATAAACAGATTTTGTGTCAAAAAA

**GLYOXYLATE/SUCCINIC SEMIALDEHYDE REDUCTASE 1 (GLYR1):**

AGAGAGAGAGAGAGAGAGATAAAGAGAGAGAGAGAGGGAGGCAGAGGGAGAGCTAGAGAGATGGAGATCGGCTTCTTGGGGCTGGGCATTATGGGTGTAGGCATGGCCCGGAACCTGCTCAAGCACGGTTTCAAGGTGGTGGTCTGGAACAGGACTATTTCCAAATGTGATGATTTAGTTAAAGATGGTGCCACGCTTGAACATAGCCCGGCTGATGTTGTGAAAAGGTGCAAAATTACAATCGCGGTGCTGGCTGATCCCAGTGTAGCTCTTTCTGTAGTTTTTGAGAAGGATGGAGTTTTGGAGGCGATCGGACCTGAGAAATCATATGTAGACATGTCGACTGTGGATTCAAAGACTTCGTGTACAATTAGTAAGGCCATTATAGAAAAGGGAGGACGTTTTCTGGAGGCGCCAGTATCAGGCAGCAAGAAACCAGCTGAGGATGGGACATTGATAATTCTAGCAGCTGGGGATGAGTCTGTCTATGCAGAGGTCCTACCTGCCTTTGATGCAATGGGGAAAAAATCTTTTTTTCTTGGTGAAATAGGGAATGGTGCAAAATATAAATTAGTGGTCAACATGATCATGGGAAGCATGATGAATGCATTCTCAGAAGGGCTTGTTCTAGCTGACAAAAGTGGTCTCAGCCAGCAAACGCTTCTGGATGTTTTGGATCTTGGAGCCATTGCAAGCCCCATGTTCAAGATGAAAGGACCATCGATGATAAAGGATAAATTTGACCCAGCATTTCCTTTGAAACATCAACAAAAGGACATGAGACTAGCTTTGGCTTTGGGCGATGAACTAAGCGTTTCCATGCCTGTCGCAGCAGCTTCTAATGAGGCTTTCAAGAAGGCAAAGAGCTTGGGACTCGGGGATTCAGATTTTTCAGCAGTTTACAAGGTTGTCAATTGATGCATCTGCTGCTTTATGAGGGGCTCTACTTCTATAGGAAAGGACACAATGATGCTGCTCACATCTGAAAATTTCTTTCCCAATTTAAGTGGGTTGTATACTTGAAGGACATCAACAGCATATTTTTGATCTATAAGCGAATTTTTAGTGTTTAACAATGTTAAAAATCTGTGTGTACCAAATTTTCTCACGAGTATCTTATGAAGGAAATGCCAATGATGTTGCATTTTTTTCATTCCTACAAAACCATTCAAGTCTCATTTCTCAAGCTAATATTCAAAGAACAATGGTGTCCTTGTAATGGCATTGATGTTATTCCTCATAGGCAAGCTTGTAATATTAAACGTGCATTGGTGTGTTT

**NEUTRAL CERAMIDASE 1 (NCER1):**

GTTGCCATCATCTACAAGCGAGTTCCCAAACACAAAGAGAGCCGCCGGGTAGCTGCTTGGTTCTCGCTTCCACTATCTCTCTCTCTCTCTCTAGGTGCCTTAGAAGCTTGCCATCTATTGCTTGGAATACAAAATGTCGTCGCTTTCACTTGTTGTGTAAAGATTGGAATGATGACGGGTCTTGTACGACCATAAACAGAGAGCTCTGCTGTTTTTCAGCGTTTTGGAGCACGCCATGGCTGATCTGCACAAGCTGCGCTGCGCTCTATTTTTTTTTCTCATTGCAATCTACATGTCAGTATTATCATCAATAGAAGCAGGACAAACATCATCATCATCATCAAACTACCTCATCGGCATGGGTAGCTATGACATCACGGGGCCTGCTGCGGATGTGAACATGATGGGTTATGCGAACCCTACCCAGAATGCTGCAGGTATCCACATCCGTCTCCGCGCCCGGACCTTCATTGTTGCGGAGAGCAGCAGCACGAAGGGCAGCAGTGACAACCGCATCGTCTTTGTGAACCTTGATGCCTGTATGGCTTCCCAAGCTGTGACAAACAAAGTCCTTGAACGCCTCAAAAACAGGTATGGGGACATGTACACCGAAAAGAACGTGGCAATTAGTGGAACTCATACCCACTCTGGGCCTGGAGGATATTTGCAGTATGTGCTTTACATCATTACATCAATTGGTTTTGTCCGTCAATCTTTCTATGCGATCGTTGACGGCATTGAGAATAGTATCATTCAAGCCCATGACAATCTGAGACCGGGTTCCATTTTTGTCAATCAAGGAGAGCTGCTTGATGCTAATATAAATCGAAGCCCTAGTGCATATCTCAATAATCCATCAGAAGAGCGGGCTCGCTATCAGTACAATGTTGATAAGGAGATGACCTTATTAAAGTTTGTTGATTCAGAATGGGGCCCTGTTGGCTCTTTTAACTGGTTTGCTGTGCATGGGACCTCTATGAGTAGAGAAAATAAACTTATCAGTGGTGACAATAAAGGCATTGCTGCTCGCTTAATGGAGGACTGGTATGATGACCAGCGATCTCAGAAAAATGAATGGTCAATAAACTCTTCTAATGCTGATGATATCCTCTCAAGCGTCATCTTTAAATCTGATAAAAGGACAAATAGTCGTTTTGTTAGACTTGGTAGCACAAATGATGGTAAGGGACCAGAAATTATACGACGACGGGTCCGAAGCTTGTTTCGAAGTCAATCTAGGCCACGCTTAGTGGCAGCCTTTTGCCAATCTAATGTGGGGGATACAAGCCCAAATGTACAGGGTGCAGTTTGCCTCGACACAGGCATTGCTTGTGATTTTAACCACAGCACTTGCAACGGAAGAAATGAGCTTTGTGTTGGTCGTGGTCCTGCATACCCAGATGATTTTGCAAGCACAACAGTTATTGGTAACCGGCAGTTTAAAAGGGCAGCCTCCCTGTTTGATGATGCCACAGAGCAGGTGTCAGGAACGATAGGCTATCGCCACAAGCATTTGGACTTGTCAAATCTTGAAGTGAGCATGTCAAACGGTGAAAAAGTTCATACCTGTCCTGCTGCAGTGGGTTTTAGCTTTGCAGCTGGAACAACAGATGGTCCAGGGGCTTTCAATTTTAAGCAAGGAGACCATCAAGGAAACACATTCTGGAGAATAGTTGGAGCGGCACTAAAGCCTCCTGCTAAAAGTCAGATCAGCTGCCAACAGCCAAAGCCAATTCTGATTGACACTGGAGAAATGTCACAACCATATGCATGGGCTCCTTCTATTCTTCCAATTCAAATCTTGCGAGTGGGACAAATGGTGATTTTATCTGTTCCTGCTGAACTCACAACTATGGCTGGACGACGCCTGAGGGAGGCTGTAAAGTCAACACTTATTTCTGAAAGTGATGGCGAATTTAACAAAGATACGATAGTTGTGATAGCTGGGCTAACCAATGCTTACTCACAGTACGTGACCACATATGAGGAATATGAGGTTCAGCGATATGAGGGAGCTTCAACTTTATATGGTCCTCATACTTTGAGTGCTTACATTCAAGAATTTCAAAAGCTTGCAGCAGCTTTGGCAAAAGGCGAGGCTGTGGAAGAAGGTCCCTCTGCTCCACAATTACTTGAAAAGCAGTTGGAGTTCCTTCCAGGTGTTGTTGTTGACTCCACACCGTCTGGTTTGAGCTTTGGAGACCTAATTGAGGATGTACCTTTGAACTCTTCTTATACGAAAGGGGACATTGTAGTCGTCAAGTTTTGGTCTGCCTGTCCTCGCAATGATCTGTTTACTGAGAGCACATATGCTCTTGTGGAACGTCTTGATAGCAACGGAAGTTGGAGTGCAGCATATGATGACGATGACTTGTGCCTCCGTTTTATGTGGGGACGCCCTTTCTCAAAGAGTCCTTACAGCCATGCCACCATTAGATGGGAAATTCCAGACACAGTTCATCCGGGCATGTACAGGGTACGGCATTTTGGTGCTGCAAAGCATTTTTTTGGTTCTGTCCGGCACTTCATTGGAACATCCAGCACACTTGTGGTGAACTAACAGAAGAGATGACTTTTGTACAGCTCTCATATCAGAATACTTGATGACTGTAAGCTCACAACACACAGCTATAAATTTTGTATCTTACATATTGTATATTTGCTATGTTTAAACCAAAGAGCAAGTGGGGGATGCCCAATCTTGTGGCGATTGTACCCCATACTCATGACACGGTCCAATTTGATGATATAGACAGCTATTATGACGCATTACTCCTATTTGTGCCATTGGAGCACAAGCTTATGTAAATATGGTTTTCTGTTTGCTAAGAGAAGTTGATATATTTGTCAATTTCAA

**α-MANNOSIDASE 2 (GMII)**:

TTTTTTCCTTTTCAAGCTGAACTTCTCATATAAAGTAACAATCCACATTTGAGGTTGTCAAATACACACAACTAAAACGCCAAAGAAAATGTACACTTGCTACTTGAGTATAATAAATTTAGTCTTTGAGCTTGCAAAAGTAAGATCTAAGCACTGGATATGACTTTTAATCCTTTCCAGAACCTAGAGCCCAGAAGGCACTTGTATCATCACTTGTATGCTGCTAGATTTTTACTTTTTGAAGCATTCTATCACTTGGTTGCAGCTGTGAAGCCCATCCAGACAAGGAGGAGCAGTTTTGTTTATAAGCTCCCTTCTAGTAAAGCTCTGGTCACAAAAAAAGCTGCTGTAAATGAGAAGATTTGAAAGCAGCTGCTTACCCCCCCTTCACAGAAGTAGCAGGCAAGTTCCATCCAAAACACACAATCACCATACACTCCTTTTGGGCATCTGTTGAAAATATTCTATTGGCGATCCTGAGAAGCTATGAAAACTCCCCAGCCTGTAAAAAATACAAAATTCAAGTAGCCTGCAAAAACTGCTGCTTCTTGTCTAATGACAAATCCAGCTTCATTGACCGTATCTCCATTGGAGCCAGCATGATGGGTCCTTTCAAATCTGTAAAACCTCCCCTTTCCTTCCTGCCAGAAGGCTCTACATAACCAAGTTCCTTAGGACTGTCATGTAACAAGTTCAAAGATGAAAGCTTTGCAGTAGATGTATGAAGTTCATCAAAAAGGCCATATACATCTACGATTTGATCAGCAAACCTGCTGCACTGTGTTTGACCCTTTCTGCAATAAGAAGAATCCCATCCCCTGCACTGTAAAGTAAGAGCGTATCTTGCCTGCTGAGTAGATCTGGACTCCAAGAATTTTGAAGGCTGTGGCACTTTCAAGCTTACTACGTGCAAGTCACAAGGAAAACTCACTGCCAAAGGTGCAAATCTCTTTCGCAACATGGAATGTTGGGCATCTGAAGCATTCTCAATCTTCCCAAAAAACAAATGCATGGGATAGCTGAGCTGCGCCAAGACTCGATGAGAAAGGAGAGATGGGACACGGGGAGTGGAAGCAGCAGTATTTGACAGAGATGTGATATTAGTTTCTAAAAGCAGGTGGAAGGTGACATCCATTGGACGATTGTCAAGCACTCCTTGTCCCAGACCTCGCTCATCATCCCGTGTCAATCGCCTATCAAGCATCACCTCTAAAGCCCCGGTTTTTAAACTTGCCACACCCAGTGATTGGCGAGAGTGAATAGAGAATCTTTTCCCCTCTGCTCCCTGTAAAAATGCTAATGACGGCATTGGATAATAGTTTCCTTGAAGTGGTATCTTGCTGCGAGTCTCTCTAGCTATAGTTTGAAAGCCGTTTAGGTCAGTGAAGAAAGTATTGCCTGTATCCAGATCTGTTTCAAATCGGACAATCAACTCTTTGTCATTGTATGCATTATCGAGTAATTCCACATGATAAATCATCTCTACCATGAAAGCTTGAACAGAGTCTACCCCACTATAAATACAGGCCCTCCGGCTGGCTGGAGAAGGATTCCAGTTGCTTTTTGGCTGTGTATACACTTCCTGCATTATGGACCCTTGAGTTACAATCATCAAACCACCTGGACTAACCATAGAGCTCGCTTCTCCTGCCGGTTTAAAGAGGTATGCACCACTTCCACGACTAGAGTAGAAAGCTATGTCTTCACTTATGGCAACTCCAATGCCATCCTTCTGGCTCTTCACATGCCGCAGAAGCCCATCTTGGATTGAAAAGGCCAGTGTTAGATGTTGGTTGTGTATCTCAACCACCTCTTCTTCTGGTTGAGAGCATTTATAAGGCGAAGGACAAGTGAATGTATCTGGTGCATGAAAAAGCTTAACTTGAGCAAGTTGTGCAGACCTGCAAGACTGAGAATCCTTGGCAGAAGAGACGAAGAATGTTTGAAATCCCATGGGGGGAATCCGAGCCTGCCAGTTCAGCCTATGTCGGCTGCCAGCCTTCTCCATGGAAATCCCCCATTCAGGGCTAACTTGACTTTCAATAGGTTCCCAGTCAAACCCCATTACGCATACATCTGTCGTCTTTACAAGGACCATCACAACCTGATTTACTTCCTCCTCAAGGGGATTGTATATAACCACTCTTTGTACTTGATCAGAAACCACCTCGACTAGCCTCTTAATTGGTAACCCATTGTGTGATGTTTGCTCTTTCTCAGGTATGAAAATGTTTAAATCTGCATCTCTGCATGTGCCTCGCTCAGCATACCTACCCAACAGGGCACCCACACCTTGTGCCATGAACTGCTGCAGATCTACAAGAGCAGAGTACATTTTTTGTGCATAATCCACCACTACATGGTCCTTTGCAGTGCCAGTAATTCCATCATGGTGCTGAAACAATGCAAGTGTCCTGCGGGCAGCTATTAGCTTCTCACCAAAGGGCAAAGGAAAGCTACTTTGATCCGATGGTTGACAGTATGCTGAGGAGAGGGAAAACAAGACATCTGCTGCCCGCAATGTCTCCTCCAGAACCCGATCTACAGATTTATAGAACGGCCGTGACACATAATAACCACTCCAATAATCATGCACCCTGTCTGCATAAGTAAAGAAATCTCCTGACAAAGAGGGAATCTTTAACTTGGTTTTCTGTGCATCCACTTCTTCCTTAACCGCTAAAAAATAATCTTCTAGTGTCCCAAACTGTATTTCAGCTTTAAGCCTAGAATGTGCATTGATGTAGTCAAAAATCTTTTGATAGTTTTCAAACTGCAGGTCTGCTTCAGCAATTGTAATGTAGCGAAAATCATCCCCGAGAGGCACAAGGAGTGTGTTTGTCTTGTACAAAGTAGATTTCTTCCTATATTGGTCCAGTAGCTTCTCCGACCTCTCTTGCACATTACTTGAAGTTATTTGTTGCGGATCAGAACCCCATGGACAACGCTGTGCGGATCCAAACCCTGGCAGACGCCAATAGTCGTACTGACAGCAAATAGCTGGTTCAGGACCACAAGTGTGAGGAATGTCATATGAGTAAAATGGCATCATGTGACAGAAGATGTCAGTAGTATTTAGTGTGTCCCAGCTCTGTCTCCAGAAAAACTCCAAATTTTGTTCCTGTGCTAAGACTTTCTTTACTTCATAGTGTGTCCTTTGAATGAGCATATTCTGAAAGCCCATTCGATTTAAAAGGTAAGCCATCGTAGGTGAATGTCCAAAAGGATCAATAGCCCAGGAATTTTTTGGAGTCACACCCACTGTTTCCTGGAGCCACAAGTTTCCCTCCGTTATTTGTTCAATGATTGCATAGTAGTGAGAATTTGCCTCATCATTCATGACCCAGCCACCACCAACAATTTCCAGCTGTCCGTCCTTGACAAGTGTGATAAAATCTCGTTTCTCAGCTTCTGATGCATCTTTCCACCACCTCTCAAGGTAGGACATTTCTTCCCATATAAATTTTCGCTTTTTGTCCTTTCGAAGAGCCTGCACAATGGTGGACAATATTCGGCGTGATCTTTCTTGGTAGTACTCCTCGACAGTCCGAATCCAGCCGGGGTCATTGTGGGAGTGGGGGACAACAAATACACGAAGTTTCTGCTTGTCCCATTCGTTTCCTGAGTACTGCACATTCCAGCCTTGCGTCCAAACCCCCCCAGCAATATCCTTGAACTCAATGGCATCATAGAGCTCATTCATCTTGATATCTACTTGTGATTCTGACAAATTGAACCCAATGTAGGGGGTGACAGTGCTCGTCTTTGCTTGCAAGGTCCCCGGTAGCCCCACTTTGAACATGGCAATTACAAAAATGGCAAGGGCAAAGAGCACCAAAAGCCTCATCTGCTTCAATACGACCTCTTGGAAAACACGCCCTCGGCGACGACCCTTGCCCTTGGATTTGGGCGATGGCGAAGGCAGCTTTTGCACCGGGTTTCCGCGTCGCGATGCTGCACCTGAAGCAGCCATGATGCTCCACTCAAGATCTACGCTCCCTTTCTTTTTATGTTATACATGTATGGTTATGACATAAAGTGGGTGTGTGTGTGTGTGTGTGT

**BEACH-DOMAIN HOMOLOG A1 (BCHA1)**:

GAGAGAGAGAGAGAGAGAGATGATATTATCGTCCTCAGAAGCCGAATACACATAAGAGACTAGACCAAGACCTTTTCAATCAACTCATACGGCATAGATCCACAAAATTAAGGTATAAGACGAAGACTCCAACCTTGGATACACTGTTTCTCTAAGTAGGTTCTGCCAACAAAAACACAAAATACAAGCATAAGGATGAGAGCTCACAATCTATTCTGATTGTAACCTTCTACGGAAAACCGAATACGATTTTATCTGTATTGTGTAAAAACAGTTATGCCCCTCAAAGGTGGCACTATACATGAGCCAACAGTGTGAATAAGCTGAGCATGTGTTCTAATTCTACATGTGCACTCCTGCATCCCTTGGAAGGAGCTTAACTGACCAAAAATATGATACATCCAACTTTACAAAACACCTTATCTTGCAAATATCTGGTTAATGGAGGCAGCCTACCCAATATGATGTATAGTTAGTGCTAGAATGGTTAGTGACAAAATTGGAATCAAAAAATCAGCAGAGGTGTGAAATAGCCAACCTCTCCCTGCCAAAGCGACGAGCAGCGTTTTGCGGCTGATGTCCCTGGCTAAGAAGGTGACCTTGCACCAATGGCATGTTTTCAATCTGGCAAATCAGCCATGGTGCTCGAGCATGGGGTGGCACATGCAAGGCATTGCAAAGAAACTGTAATCCCTTCTAGTAGCCCAGCACCATCTTTTTACAGCTCATGCAGCAGTGGCTTTGATTGCATCCTTAACAATGATCAGCCCGTTTTTGCTTTTTACATAGCATTCCACAATTCGACATCATTAACTTCATCACTGTTGGTGCTGTCAAACTTTCAAAGAGGGAGGCTGAGCCTCTACCCCCGCAGAATTTGACGAGTATCAAGATTGAAGCTCCTTTTAAGCTCCCCGCGCAACCAACCAAGACCAAGCTTCATAAATCATTGCGCCTGAGTTTACAGACAATCGGGTCTTTGAAAATTATCATCATGACTTGTCATCCATCTCCTGGGAAGCTGCATTCTTGACTGTCTCATCTGGGAGCGACCAAGAAATCAAATGCCCTCCTGAATCACCACTCAACAATTGTTTCAAATCACTTGTCAAATGTACACATGTGACGGACTGCTTATGCCACTTTAGCACTTTGAAAAGGACGAGGCGATACTCTTCTGTGCCGCTTTGTTGCTGCAGAGCTGATTGTGAGAGGCTATCTTCCCTCCATTTTTGTTGAGTACTCCCAGCTGGTACGCTTTGGCTTCGTATACGCATTCTAGATACACTATCTGCAGAATTGTGCTCCATGTGCCAAAGCTTAATTGAGCCACTTTGATGACCAGTGATGAACCAATTGGTTTCCATCCAATCGGAAAAGCAGGAGCCCGTAACAGAGACGATGGTATCAGAAGGGAGCTGAGAAGTATTTACAATAGCCAGGCAGTCACCATTAATGCTCCAGACTGCCAGAGTGGTGCCTGCTGCTGTCACAACATCCCCCGTCATGTCGTTTGCATACACTGCAGATGCTGCTGCAGGAAGCGCGGGCAATTGCCGTATGTATTCCAAGCTACTAAGATCCCAAAAAATGACACTACGATCCTCAGAACCGCTAACTACAAGGCTATAAGGCTGGCATACAGCAAGGCAGGTGACCCTCTGTGTGTGACCAGACAATGCCCTTTGTAGCTGTAAATGCCGTTGCCTTCGAGCACCCTGTTTGCGGAGGCGCCAGACTGCCACCACACTATCCTCACCCCCTGTGACCAAAATTCGACCATCTTTACTCAAACCAGCACATTGCACTTGCCCACTTCCATGCAATCCTTCATGGGTTGACATAAGCTTGTCCTGATCATAAGTAAAGAAACGCAAACTTCGATCTGGAAAACCCCAAGCTATATACTTGCTGTAAGAGCGCGGTTTCAACAGACGATTAACTCCTGAAGCTAATACTTTGTCATGAACAACAGCTATCTGTGAAATGGAAAATGTGACTGATCTTATCTCTTGAGGGAGTAGAAGATTATGCTGTCTTAAAATATGCATAGGTGGAAACTTCTGCTGCCATTTTCGCTTTGGGTGTGGCTTTTGGAAAAGCTGTCGAGGCGTCTGCCCAAAATGATTGATCTGGGCTAAAATGGACGCTTTCATAGCTGGATCATGTATTGAGTCTATGTCAACAGCCCCCTCATATGTCAAATAGAAAAAGACATTGGTTGCATCCACAGCAGCCTTGCCTTGCTGCCGAAACCCAAAAATAAGGTCTATCCAATGGTGAAGATTCTCGGAAACATATTGCGACTCTAAAGCTTCTCGGTGCTTTTGAATAAATTCCCTAGCACTGCCGTTGGCCCATGGTGGAAGAAGAATGTCACCAACCTTTTCCCCTGACTGCTTGACCCCAAAATCCAAACTAAAACGGTTTTCCAGCATTTCTGGTAAGTAAAAGAATTCTGGAATGAGCTCTTTCACATCTGCTGTGTTGCCCTGACTGGCACTTAACCATGTGTCCCTGATGCTGTTGAAAAGGCGATCCGCATGATCAAACTGACCTCCTTGCAGCTTCTGATTTTCCCCACTAAAGGGTGGCAATCTGATAAGATAAAAGAGAACTGTTCCAGCACTTGAGTAATGAGACCCATAGTGAAAGCTTGGAATCTCTGGGTCATCCCAGCTATCAAACCTTCTCCGAAACTCCTCTTCCCTTTCACGTGTTAAAGCACCCATAGGTTTGTCCAGCTGACGGAAAGTGTTTGGATCCTCAAGCTTTAGATCATCACTCTCGTAATCAGCCAAGACCCATGGAAAAACAGGATATTGTGTGAGATCATTGTAGCCCCGTCCTGCGAGTGTGTTTAAATGCATCAAGTATTGGAAGTTGCTGATTTCCCCGTTCTGCCAGCGTTTGCTAAATGACTTTGCCATCATTTTGAATAGCCGTCCTCCTTCATTGCTCTCTTGTTTAGACGCACCTGAGATTGTGGCATCCAACATGCTGTTCCTTGGAAGATTCAAAGCAAGAAGGTTTTTGAAGACCTCATCCCGCTCATTTTTGTGGAAGACTAAGAGTTCATTGCAACCATCCATACTAAAAAGCTCAATAGCAACAGGGCGGAGCTGGTAGCGCCGTTTCAAAAGTTCATGCACGTTCTCCAGTTTCCACATGTGCCAAGGATGAGCCATGTGGCTGACATTTTGTACCTTCTCTCTCCCCCAAGCACCTCCATTGTATGCCCAAGCTCGGCCCCCTAGCCATTCACCTGCAGTCTCCTGCCAAGAACCAGTTGGATTCAGCTCTTGGAAAGAAGCAACACCTGAGCTGCTAGACTGGACTCCAAGTGCTTGATCAATAACGGACAGCTCATGCTCTCCAAACTTCTCACATATGCGGCCCGTATCATCAATAAAGAAGTTATCAATAACATACAAGCACATTTCTCCAATGAGAAAAAGCCCATCCTTCTTTTCTAATCCAACAACTCTTTCACAATTGTACCGGTGCCTTAACTTCTCCCCCGGCTCCAAGAAAGGCCTTATCAGATACTCACCATCATCATTAACATCCTTGTCTGTCTTGCCATCCTCATACCTGAGGTCTTCACTCAACTGTAGTGATTGCTTAGGAGAGCTTTGATGAGACTGTGTAAAGTCTCCAACAGTACTAGTTTGAGCTGAGAGAGTGCTAAATGTCCTACCGGGACCTTGGGAAAGACGTCCGGCAAACCCATTTTCATTCATTACTTGAGTAGTATTAAACCCTGTCCTAGCTGAGGAGGATGTGCTTCCTTCTTCCCTCAACTTCAAATCTTCTTCTTCATCCTCATCAACTTCTAAGAAATCTCTATTACTGTAGCTAAGCGTATTCTGGTTTGTATTGCCAACTGAAAACAGATGGAAGAAAGAGTCATAATCTGGATCTGTGATATCCACCCCAGCACCATCTCTCAATTTTAGCTCATGTGCAGCTTCCAGCTGCAGTCTGTTTTTAGTGGTAGAAGCCATGCTGACCTTGATCTTAATGCGTACCAGCTTTTTGCGCATGCGATATGGACCCTCTGTTGCACACAGTTCCCAGCCAGGTTCGTCTGTGGATGTGACCTTAAGCATGGGTAATATGCCACGTTCATGCACAAGATGTTGAACATGTATCTCCCATTCACTTTCTGCATGGAGAACCCAGCCATATTTGTCCTGTCTCACAACCCTGAGCTCTGTTGCCATTGCATCCCTTACAACATCAAGCGCATATCGCCGCTCATTCATCTGTTCCCAATGGCGATCTTCAAATTTTGAAATGTCCCTGAAGCGCCTGGACATCTCCCGTTTCCGTCGGCTCTCTCCCAGCTTCATACGAACTCCAGGGAACTTCGCTGCATTAGCAACATACTTCATCCACATGATTGAAGCACATTGCTCAAGTACTTTTGATATGACCTCTTTGGAGTTTCGAAACCACACATAAAACTGGGCCAATTCATTCATTAGAAGCTTGTCAAGTCCACCTCGGAGTACATCCAAGGCTTCACCTTGACTCGGTCTGATAATAAGCAGCTCTTCCAATATTGGTTTTCGGAGGGCTATCAGACTCTTCATGACCTCTACTGCAAGCATGCGAACACTTGGCCTTGAATCCCACAGCATAGTTATGCTGTTAATACAAAGACAGCATACCAAATCAGTGTCCAGATTGATAGGGCAAAAGATCAGTTGCCTGTTGGCTATCACCAGTTGAAGCACAGTGCAGATATCCACTGAAGCTGATAAAAAGCTTCCATCTCCGTCGTCCAGCCTCTTCACTTCTGTCTTTGGCCAATTAGAAGAGATGAAGTCCTCTTCCCCAACTGCAGAAAGAAATGGAGGAGTGAAGCAGTACATAATCATGCGATTCATGTTCTTTAACAAACCCTGGACATAAGCTTCAACTTGCCGACCTCCCCCTCTGCTCCGTGATAGCAAACCCTTCCCAGTTAGAGTAGATTCCTCCACTCTCCTGTCCTTGTTTGCAAGTTGCAACATTGATAATAGGAACTCCAAAATCTTCAGCACTGATCCAGGCTCGGCAAAAGCACCCATATACACCCGATCCACTAGCAGAGTACTGAAAGCATCTAGATTTGAAGACCACCTGTTCTTGTCTAGCCTTTTGTTGACCTCTTCATCATCACGCAGCAGCCTGCGCTCAAAGTATATCATCAAACGACCCAGGCACAACCCTTGAAATGTAAGTGCAGTTTCTGCAGTAACATATAGCGGTGCTGCTTCAAGGACTGCTTCCACAAGTGAAATGGCCTTAACTTGTTCAGACAAACCATCTGCAAGCACCTCGGCGATCAGGTCTAGCACAGCTACAGATCCTGCAGAGCAAGGACCACCACCTGACCCCACAGAATCAAGTTTCAAAAGCCACTCCGGCTTTATTTCGACAAAGCTGCTCTCTGGATAAGCTCCTCCAGGTGTACTTATTCTGTAGTAAACCTCTGAATTGAGACCACTCATGGATGTTGAAGGCGTGGCCAGTGACGAAGCCCCCACATTGAACTCACTGCGAGATGAGCTTCCCATCCATGATGCTGACAGATCAAATGGAGTGATGGACATTGAATGTGCTGGTTTGATGCCCGATAAAAGTCCAGATGCTTCAGAAATAGGTGATCCTGGAGCTAATACCAAGCGCGATCCTACAGACTCTGCATCTGCATTTGCAAACGAGGAGGACTTTTCATCACGCTCCATCTTCTGAGCAGGTGTCGAAATAGCTGCAAGGGCAACCAGATGGTCAAGACTGTCCATTGCAAAAACATTCACATCGCATACAATATCCTCCGATGGCAAGGTATCTTCTTCCAAGTTTGGTATTATATTTATTTCCTTGGATGCGGAAATTGCACGTTGAATGCTGACAACAGTATCCGCAACGCCCAAGTTCCTGTGCTCCTGTAATTTTGGGCCCACATTTGTCAATTGCGGCCCGTTCATTGAATCTGAAGATGTGATGCGAGAACTAAGTGCATCTTCTGTATCCATGGAATTCTGCCGCTCTGCTAAAGCCTTATATTGAGAAGATTCAGCAGCATAAGCAGATCTAACACATGAAAAGTAGAGATCTGCGCAACTCTCGAGAACTTCTTGACGCCGCATCGCAGCTGAAAAAGGCTGACATATCTTTGTCAGGTCTACCATGTACCTTATTATGGAGGTGACCACTGCAGGAGCCGCAGCTTCACCACCTAGCAGCCTTGCTGCATAGGTTTTGTGCAATAATGCCTCCCCCTGCAACTGTTCTGTGGTATCTGTAAGTCCCCTAAGAATTTCTTCCATTGAATAACCTGCTTGCACCTGATCTCCCATCTGCTCCACTGATTGCAATCGCAGAGTTAATCTATCAAAGGCAGCTTTTGACATGGCTATAACTGCCTCCAAAAACTCAATAAACTGTAGCTCTCCAGAGCGTCCATCATCAGGCATCAGTGCATGGAAGTCTAGCAGTCTTACTTCATGTTGCCTTGGGTACACAGACTTGCCAAACATGAGACAAAAAAGAATATAATACACTTCTGGGGAGTCAAAAAAACTTGGGAGGACCTGGACAAGGCCATTAAATCCACCACTGCTGCGAAACTTTGACCAAAATGTGGTTGAGGAAGAAAGACATACACCAAGCAACGTCATCACCCAACGCATGCTTGAAGGGTGCACAGCCTCATCAAGAAGAAATGTGATCAGCTTGGATGATACAATTTTATGCCATAACTCGAGGACATCTTCTGAAGGGATTGTTACTTGAAGGTCAATTAACATCTCAAGTAGCATATTGCGCACTATAACCTGCACCCCATTTGACTCCCTTGGAATTGTAGACCTTTGTAGCACTTCAGGGGGTTCAAAAGTCATAACCACAAAATCAACCACACTCTTGAGTTCAGAAGTTAAAAATCCATCTTCTAGTATGATTCCCAACAAAACCACTAGCTTTTCCAGCACGGTAGTCTCCACATCACCCCTCTGCAATGTAACCAACAAGTGCTGCACTACATTGACACGACGTAGCATTGTCAAGTTGTGGTTTCTATATCTATGCATTGACACAAGCCGCTCAATAAACCCAAGTAGAGCAAGCTGAACTGCAACAGGAGCAGCAACCCACAATGTCCAGTCCATCAGAACATACTCCATCATTTCAGGGTTAGCAAGGATGAGGCTATTACAGACTTCTGTGGAAGGCTCAGTAAGCTCCATATCTGACACATAACTCATGGTCTCGTGCTGAAAAAAATCAGCACTGTCTATTTGAGATCCAAAAGGTGACGCCTCTTCTCCAGTCACTTTAATACTTACTGAGTTCTCCAAGTTATTACTGCCTGAAATAGACCGAAGTGCCAAAGAACTGTTGGTAGGTTCTGGAAGTGTTACTTTGCCTCTGCTGACCATGGAAGCCTCGCATGCAGCAACCTGAAAGAGCATGTCAAGATTTTGCATTTCAAAGAAGCACATCCGATGTCGTAAAAATAGTGCAAGCAGGTGGTAGCCTCTGCTAGCAGACATGTCCCGAACATTGTATGGGCTCCAACGAAGAATGCTGGTTAGCAAAGCAAGGGCTAAATGAAGCATCTCTTTACTTTCAGCAGCTTCGACTAACGCCAGTACTACTTCAACTCCACCGATCTTCCTAATATTTTCACCAAGACTGCACGGGGAGCATATGCTGATGTCTCCATGCAGCCTTGCAATGCGGGGTAAACCACCAAATGGAGAAGCTGCAGCTGATAATGGATCAACCAGATTAATTACAGCCAAAGACCCTGAAGCAGCAATGTCCACACAGGTTCCATCAAAGCAAAAAATGAGCTTCTTACTAGACAATTGAGTTACAAATGTGGCTAGCCTATCCAAGTCCCACACAACTCCACTACCCTCTGTCTTGCTTGCACTTGCTTTCAAACTGCTCTCTACCTTTTGATTACTGGCACCTAAAGCTAGTTCTACTTCTAACGCCTCAAAAGTTGCCAGATTTCCTCTACCACAAGCTTCATACGGCATGAACCTTAGAAGATCTGTGTCTTGAAATAAACCACGATAGCTCCTTCCTAAAGCATACATCAAAAATATTGACGGCGCCGAGAGAACTTCCTCGTATAAGTAACAGGAACCAAGATTCCAACACAAGGGAAGAATCTCAAGCTCATTGGGTGGTGTCCCAATCGTTCCTTGAAGAGATTTACCACCTGGTGAGGCTGAGTAACCCAGCTTTCCTGTTTGCCTCAGTCTTCCATTCACATAAAGGCAGGCACTGCTTGATTGAAATAAACCTGCTAAGGCATTTGGTTTGTTGTGCACAACCACCACATGATGCCATATGTCCTCCTCAAGATACACTCCTTTGAAGGCCAAGAAACTGGTGGCGCTTGTTGCAAGGGCCAACATTCCAGAATCTTCCATGTAGAATTCAGCATACAAGTGGCTTTTGTCCTCTGCAGAACCTACTGAAAAAATCTTGATTTTGGCATGTTCACGCCCTGTTGTGTTTGCCCTAGCCTTTGAAAGTGGTTTTCTTGGAGCTGCATCATCTAGTTCCACACTTGAATTTTTTGAAAGGCTTCTATACCTCAACCAGCATGCAAATGAATATCCTGCTGCTGGTGGCCAGGTCCTGTCACCTAGCGAAATACGCATGCAAGAGTAGCCGGACTGTCCCATTGACATTTCTACAAAGGGTGCAGCCTGGCTATTCTCAGATCCTACAGTTTCCAAATGAATCAGTCTCTTCATTGAATCAAGAAACATCTGGCCGAACTCTTTGTTAGTAGAATCTCTACTCAGCCAGACCCACCTCGACAGTGTCCTGAGTTCAGTCGAGGATAATCGGTATGATCCAAGAATTTCCACGATACTTAACACAGGTTCAAGTAAAGATGGATTGCTACTAGCAGATTTTGCAATGTCCAGCAACAAGCTCACACACCCTGCAGCTGTAATACAATCTTGGTTGTAAGCACAAGCCTGCGAAAGTCGTTTCACGACTTCCAGCACTTTCAGTTGCAGTCTTGATGACAAACATTTTAAGCAATCAATCAGCACTTCAATAGCTCCTGCATTAGGCACAAGTTCAGGTGACTCAGTGTCATGTCTCTCACTGTCCAGAAATGTGACCATAGCATTGTTAGTCTCCTCGGGTGATTGACTACCATCAGATTGCCTACGAGATGATCTGTTTTCATGTGAGAGAACAAGCAACTCTCCAAATGGTCCTTTCTTCTCTAAAGCAAGACTCAGAAGCAAGTCTGCAACTCGTTCCTCGTATTCTGCAGAGAATAGGCCAGAGTCACATAACAGCTCTTTGAAGGTCTGTGATGAGATTACTTTATGTAACCGCCCCCTGTTAACAGTGTTCCCTGCTACTGCTGCCTTCACCACATTCATTAATGCATCAAACACTTCCATGTATAAGACCATCATACATGGCTTTTCCTCATTATTTTTATCAGAAGCTGTGTCTTTGATAAGGTCTTGGCCAGCTTGAAAACTGTGCAACACTGTGAAAAGTAGACTGAAACCACTAGCTGCACCAAAAACATCCTTGCAAGCACTATTTACACCCAAGATGCGACAAGTTGCCCACAAAACATCTCTTTTTGTCCGAACATCCACCTTGATTTGCTGACCCATTGTATTTGCAACAACTCCACTTTTTGCTACATCAATAAGTACACCAAACTCCTCTGAATGGGACTGGTTGACATCTTCACAAATAAGACAGCTCAGCAATCGTAAAACTCCACCACGATGACAGGACACATTCAGCAGCGGAAGGGCTAAAGCAACTCCATTTGCCTTCCTAAAAGCGGTCTGATTTCCGTCACCATTTTTAAGCAATGTAACCAAACAGTCCCAAGCAATACTCAGAGTGCTATCCTCATTGAAAAGTTGAGCCTTTTTTGATAATTGCCCTGCATCTGGATTGGAAAATGACAAAGTCCTCAAGCTTATATTTCTGGTCACGCCCTTGCCACCTGCCCTTTCCACAATACTCCTTGTTTGGGGACTTCGAGAATCGGGTGGCGATGAAGTCTCCAGCTTCTTTATATCATCTACAAGGACCTCCAATGCTCCAACTTCTCTAAGAATTTTTTTGAATTTCTTGTCAAATGAAAGCAGCTTTGTGAAAAATGATAAAATTCTCCGAGCCATGGAAGCAGGGAGGGGCTGCTGCAGCAAACAACAAAGTGAGAGCAACTCTTGCTCTGGAACGTAGTTCACCACAGTCACAGCATACTCCAAGATCTTCAGCAAACGCTCTTGAAGTACTTCAGGAAAGCTTGCCATGTTTAAAATGAACAGGGGAATAGTGCGCAACTCTTGGCACAAGCCATAATTATCTAGGTGGCTAGCAAATATGCGCAACAGCCGATCCAGAACCTCCAACTGCAATCGAACATTGTTAGCTTTGAGAAAAATATCTTGTAACACTTGCACAGCTTCCAGATCCTTTACTTTTCCTTCAACTTGTCTATCATCGTTGGATGAAACCCTCCCATCAGAAAACGGGAACGAGCGATTAACTGGTTTTGCTGCTACAGCCTTCCCTCCCTTAGTTATTACTGGAGTCCCAGATACTCCAACCTGAGCCAGATTAACAAGAATGTCAAGCAGGCGTGTAAGAACAGGAGGCAAGCTTCCTTCTGTATTATTAGCAGTTTCTTCCATTTTAATGCTCGAGGCCAGTGACCCCTTGTCTTCTGAAGTACAGTCAGCTGTTTCTACTTCTCCTGCACTGAGAAGGGGATGTTGTTCCGGAATATTCAAAGCGAACTGGACCAGAAAACGGTACCCATGTGCATTATGTAGATCGTCCCTTAACCTTACAGTTGACTCTGCTCTGCAAGAAAGCTCAATGCATTCAAGCAGCAAATCTATTACACCCACAGCATATTCAGAATCTCGAACACTGCTATCAAATTCACGGATAACTTTAAGAAGAACTTTAACCAGCTGATGTTTGTGAATATACTGAGCTGAGCTGCCATTATCATTGTTAAGAAGAAATCCCAAGATCTGCAAAGCCTGCCGGTAAAGATGAGCTAAGGAGAAAGTTGGCTTCAAAGCATCAGAACCATCAACAGAGGTGCTAGGATTCATTGCCACAATGTGAAACAATAACTGCAAGGAGTCATCCTCAGCAAGACTTTGAGTTGCTCCTAAATGCTGAGCCAGAGCTTTCATAATATGTATAATGCTTCCTTCCACCACAAGTTTATAAATGTGCTCCAGATCAGCATCAACGAAATGTTCCTTTAATGTTACAGCAGGTACATCTTCCAATCTAGGTTGTCCCAATACAGTGTGTAGCACCCACACAAGGCAAGAAAGAAGTCCAGCATCCAGCAGTGGTTGCACATCCAATGGTGGGGAAACAAGGCATTCAAGGGCATACAACAAGTTTGCACCGATACTCATTACATCCTTTGGGGAGTCCATGAAAAAATGTAAGATCTGCTCTACATCCTGGCTCTCGTTTGTGCTAAGCTTCTTCAACTTGTTAACTTCTGTGACCAAGGCTTTAGCCACCACAAATGAAAACACCCGTGCATCAACCAGTGTATTTACAAGCTGGATGGCATTTCCTGTTCGCTTTGCAAGTCTACAAAAGACATCAACTGCCATCTCCAACGCTTCTTCTTTCTCCTTGTCAGAAGCCGAGGAGCGAAATTCCTCCCAGAACCTTTTGAAATCCAATTCTGATTCATAATCGGTTCTGGACTTGAAAGAAGACCCATTTGCAGAGTTTGCAAATTGGTCCCGGCCAGTTGAAACTGTTGATTGTGCCTGTTGAAGGCCAACCTTGTCCTTAAACCGTGCCACCCATTTCATTGTTACTTTGCTTGACACTGACAGAAGGTTAGCATTGGTAGTTTGCCTGTGCCAGACTACTTCAAATAACAGCACTAAACAATCCTGAATGGTGATCCTTCGCCCTTCTTCAGTGTATAACCCACAAGACCACCTTTTATCTCGCCTACTACTGCATTCTTAGGCAGATATCATTTTCTTATCCATAATGAATCATTTTAATCTGAGCGAATCCTTCTTTGTACTTGTTTAATACACTGATGAGAAGACTGTACAGACTTGAAATGAAATGGCCCTCTTGATGTTTCCGCTATGACTGCAAACGTTCTGATCAAATCGATTAAATGGCCAGCCCAAAAACCAAAATGCCGTAATGATGATCGATTATCAATGCTGAATTGCAGGGACCAGTAGATGCCAGATCGGCCAACGCTCTGGAAAGTCAATTTCTTTCGCCGATACTGAATAAAAATTGCATAGGCAAGCCTCATTCAGCTCCTAGTGTCTTGCAGTCACAAAAATGGATTGACTGAATTAAAGGCAGACGCATTTGCAAGGCGGGCTCTTAAATCTGAGTAAACCCTTTCAGCACAAAACAAGACAAAAAAGAAGCTGAACCCAAAAAGCTCTATGTCACCGCATACAATATAGAGACGCAAAAAATCAGGGAGAATTGAATCCAACGCAAACCCTCTTCAGTCCCAAAAAAGAAGCGCCAATGCAAATCTGCGCTACCCTCTTCGGTCCAATGCAATACAGAGCGAGCCCAACCATTACGAGGAAGGGCGCCTATAACAAGAATACAAGCGAGGCTGTAGATCTGCAAATGGGTTTTGAAAGGAGACCTTCCTTTGCTGCACTTTGGGCGCCTTATGATCGTGAGAAATTCCAAATCTCTCTGCGGGGGTTTCCTGGGTTCTTCTTCTTCTTCTTTAGCTGGAGAGGGATCAAAATAGTGTTGATCGCCCACAAACGCTGTTTTAATGTATAGAGAGGAGGAGGGAAAAGGCCTCGCCTGCTGCTGCTGCGCTTAGTTTTGTCGCCAAAGAAAAGAGTCACGAGGTGTCTCTCTCTCACATGAAAGATAC

**DEAD-BOX ATP-DEPENDENT RNA HELICASE 3 (RH3):**

AAAGCAAAATAAACTCCCTGCCCTCTCTCGCTCTCTTTCTTTTGTTGCAAGACGGTGGGGAGGATTGGGCTTTAAACAGAAGCTCTCTCGCGGTAGCTTCCATACCCTCTTCAACTCCACCGCCGCTGAAGGTTATGCTCTTCTGAGTATTGGGTAAGCTTGTTTTATTCTTGAAGATAAACCTGCTAGCTCTGCAGTCTCACTGTTTTTCTATGAACCCGCGTCTCTTTTTTGCTTCGCCTCCTGTTGAGTAGCCATAGCCATACTATATTGAAAACTTATCCTCTTCTTGAGCCAGTTTAGCATAGCTAGTAAGCCATCATGGCTTCTTCAGCTTCTATTGGTCTTACACATACATTCCAAGCTCTCTCTTTGGAAACTGGAAGCCCAAGAATGTCTAAGCCTAACATGGTGAGCATGCTATCTTCTTCTTTCCTGGGAGAGTCGACTCAGTCTTTCATAGCCCCTTGCTCTCTTGGGCTTGGACAACCATCCCGCTCGTTTCATGCAATGTGCGTGCGAATGGACTCACGTCTGTTTTCCAAAGAAGATGAATGGAAGCGTCCTGCAGTTGCGCTCGCAGACTATGCAGTTGCCGAGAGTGACCTTGATTTCTCAGGACTTTCAAGGAGGGGAGAGAGGAACGGGACATTGGGTGGAGTTCCAAATTTTGAGAATGGCGCACCCCCTGTTGTAAAAGCTGTACCGGCGGACAAAAAGGGTCTTGACGTTTCTACGCTAGGTCTTAAGCCGGAGCTAGTGGAGGCTCTGCAAAAGAGGGGCATCACGAATTTGTTTCCAATTCAGGAGGCAGTATTGAAACCTGCCACTGACGGTCTAGACATTATAGGTCGTGCTAGGACTGGGACTGGTAAAACTCTGGCTTTTGGTATCCCGATCATCCAAAGCCTTTTGGAAGAGAACGCGCGAGAGCAACGCCGGTTCTCGCGCAATCCAAGAGTGTTGGTGCTGGCCCCAACAAGAGAGCTCGCAGTGCAAGTTGAGAAGGAGTTCATGGAATCAGCCCCTAACTTGTCTACCCTTTGTGTCTATGGGGGAGTTTCCATTGTGGCACAACAACGGAGGCTGCAGAGGGGTGTTGATATTGCTGTTGGAACTCCAGGCCGTGTCATTGACTTGATAGAACGAGGCTCCTTGGCACTTAATGAAGTCAGATACCTTGTCCTTGACGAAGCAGATCAAATGCTTGCTCAGGGATTTGAGGAGGATGTAGAGAAAATTTTAGATAATGTGCCTTCCGCTCGGCAAAGCATGCTTTTCTCAGCAACCATGCCCTCTTGGGTTAATCAACTTGCTAAGAAATATTTGAAGAATCCCTTGAGGATTGACCTGGTAGGAGAAAATGAAGAAAAACTTGCAGAGGGCATCAAAATCTTCTCAATTGCATGCCCTTCTGCAGCCAAGCGATCCATCCTAAGTGACCTAATAACGGTGTATGGAAAAGGTGGGAAAGCTATTGTTTTCACCCAAACAAAGAAGGATGCCGACGAGGTTGCCCTTTCGATGAATCGAATGCTCGGCACTGAAGCCTTGCATGGTGATATTTCTCAGGCCCAAAGAGAGAAGACTTTAGGTGGTTTTCGAAGTGGGAAGTTCACGGTGTTGGTGGCAACAGATGTCGCAGCTCGCGGGCTTGATATTCCTAATGTGGATCTCGTTATTCATTTCGAAATACCAAATGATCCAGAGACGTTTGTTCACCGGTCCGGAAGGACGGGGCGGGCAGGTAAAGCTGGACAAACAATTTTGATGCACTCAGACTCCCAGATGAGGACAGTAAAGACGATCGAACGGGATATTGGCTGCAAGTTTGAGATCATTGGGGCACCAAGTCCTCACGATGTTCTTAGAGCGTCGTCTGAGCAAGCCCGTACTGCCCTGACTAAAGTGCACCCTGAATTGGTGACTGTATTTCTTCCTACTGCAGAGAGCCTTCTGCAGGAGCAGGGGGCTAATGCTTTGGCAGCAGCAATTGCTCATTTGAGTGGATTCACACAGCCTCCGGCTTCCAGATCTCTTATTACACATGAACCGGGATGGATGACGCTGCAATTTACTCGATCTCAAAAATGGGGTGCTGGTGGTGGCGGTCTCACATCTCCAGGTGGTGTTATGGGAGCACTTGGTGCAATTTATCCCCGAGCTGCCGATGAAATTGGAAAGATTTCAATGCTTGACCCATCGCAGGGCATGGGTGCTGTGTTTGATCTTCCTGAGGATATTGCAAAAGAGCTTCTTAACATGCAGCTGGGACCGGAGGATTGTATTCAATCAATCACTAAGCTGCCTAGGCTTCAATATCAATATGATGATTCCCCGCGAGATCGGTATGGGAAGTTCTCATCAAGGGGCGGTCCTCGCTCGGGTGGTGGATCAGGTTATGGTGGAGGGATGCGGGGCAGGTCATATGATCGCTCTGACCGCACTTATGGATCAGGTGGATCAGGTGGATCAGGTTATGGTTCTAATACTTGGAGTAGCAACTCTGGCTCGAATCGGATGGATGAGCAGCGTTCTGGTGGATGGAGAGGAAGTAACCAAGGATATCAAGGGGAAAGTGGTGGATATCCAAGGACCCAGCAAACGTGTTTTATATGCAATGAACCGGGGCATAGAGCTGCTATGTGCCCTAGGCGTTAAAGATGAACAACCATCGGACAACAGTTTTATCTTCTTCAACAGTGCTCGCATATCCTAGCACAGATTATATTACTGATGCTGAGTGCTGCAGATATTGCAGCTTTATCCATTTGATAATCTTTGTAAGCCCTATGTGCGTTTTGGCTGCTTCTGGTCTCTATGATGAACTTTCCCGCTCCTACCAACTGACATCTCTCCAGCTCATAATTTACTTGGGCTAGGTTTCAATTTTTAGTATCTGTGCTAGTTGTTCCAAGGTCTTTTTCTCAGGTTTACATTAATAGCTGGTGCAAGCCATTGGATATGCGGTTGTTTGTTGGGTAGAGGAGATTTTTCTGTTTTTTTTTGTCTTGGTTTCTGGATTGCATGTGCTGAATATGGCTACAGGTTTGGTACGCTTCCCATCATTTTGTTTACAATTTAAGCTATAAGTAGTTTCTCTCTCGTCTCTCAACTTAAAATTGGGATGTCGAATTGAAAATCATTGTAAAAGTTCTCGAACCAAGTACATGCTTGCCAGTGAAATACCAGGGCTAGGGGTCTCTCTCTCTCTCTCGGCCTTTCTGCTGGGAGCTTGCACAGTTCCATGTGTCAATGGACAAGAAATCAAAATTTGTTAGTTCCTATG

**MA3 DOMAIN-CONTAINING TRANSLATION REGULATORY FACTOR 3 (MRF3):**

CTTCTCTCGATTCTTCCTGTCCATGGCTGAAGTAATAAGTTTCTTCACAAAAACTGCGTTTAGTGCCGGAGCTGTCAAATCTTCTAAGCTCCTAATTACTTCACCTACATCATCAGAAAGGAAGTACTCTTGAATGATGCTTGCAGCCTGTTGCTTAAAGATTGTAACTTCATCTGTCTCCATGCTATCAGTTCGTTCTGAAATCACTAGGGTAGTTGCAGACGATGGACTTAGAAATCCAGCATCAATAGCCTTGGATACTAAATGCAAAAGCATGGATTTTGCATTGGGTATGTCCAAAGACAGGTCATCGACAGCATCAGCAAGACGTGCAAAACCTTTAGACATTTGGCTTGAAGTTATCAAACCTTCATCATCAGCTTCCTTTAAAAGAGACAGAATGAGGGAGTCAGAATTTCTCTTCTCCATTGCAAGTGTCAAAGCCCTCTTCACCAAGTCATGATGGAAAAATGGAACATTAAGCTGCCGGATGCACCGGCATGCTTCAACTCTGTCTCCACTCTCAATGTATTCATTCAGTAAAGCAGTCATCTTTTTTCGCACTTCCTCTACCGTACTATGAGTAGAACCACCCCATCTGCGTTCTACAATCTCAGCATGTAAAGGTGGTAGCAAGTAGTTCTTTTCCGCTTTTTTTATTACTTCATGTCCCTTTGCCCCTTCAGGTAATGTCTGTACAATCTTTTTAAGAAAAGCAGGGGGAAGGATATCATCAACAACTGCTCTAGCTACAAACAGAGCCAGAACATCCACAGCATCAGGAATGTCAATTGCCAAGTCATCAGCTGACTCCACAAGTCTGGCAAACCCTTTGGCAACTTGATCAGGTTCAATTACATCAGCATATAGAGCAGAAAGTAAGACTGCCGCCATTTCTTTCTCACGATCATGTCTGTCCAAGGACATGGACACCAGCTTCTTTACGAAGTGGTGATGATACATTGGAGAGCCCAAGTCACGAAGGTCAGCTGCAGTCTCAGCAACACGATCTGTTGTGAAGTACTCCTCCACAATTTGAACCACCTTGCCTTTGTAATCCTCAAGAGACTCTGAAACGGAAGCACTAACAAGCTTGTAGGGTTCCTCTTCACTGTTGTAATTGGGATCATTCCGATCTACACGAACTTCTCCACCTGTCTCTAGTAATGAGCCCCATGTACCTCTGCCTCCAGCACCACCTTTCTTGGGGCGGCCGAGCTTTCCAGAGTGGGAGCGCCTGTCCTGCTTGAGACCGCCCATGGGCTTTCCCACGACAGAAGATGAAGAAGAAACATTGCTAGGTTGCCTCACCAGCTCTGATGAACCTCTAGGAGATGAAGCTTCTGGCGGAGACCCATTGCTGCCGAAAGACATGGTCTTCAGCAGCACCCTCTGCTGTTCAGTGAGAAAGCTTTCCTGTTGAGGAGAACCCATAAACCCTGATCAAAGAGATCGAGAGAGAGAGAGACCCGCTGCCGCAGTTGCTGAGAAACCCTAATGCTAGATACCCTTACCCTGATGAGCTCAGAGAAAGGAAGGGCTATAAATGCGCTGATGGAGATGAAAACCCTAAATTAGAAACAGAAATCGATGATGGGTCTTGACCAAAGAGGGCCTGAAGATCACAGCGAGAAACCCTACAAATAGCACACAGAGCTCAAGAATGGCGAGACGGCGATTCGGCTTCCACTCTCTTTGCTGTATTTGTAGGCCTCTCTCACATTGTTATTGAGCTAAAGGCTACTTACTATGTGTATGGCCAAAATGGATATGTACAGCATGTCACCTGAAAGAGACATCCTTAGTGTGTCTAAGTCTCTTAGATAAGATCAACAAGAAGTGAGCACACACTTATTTTCGCATCGTCATT

**PRE-mRNA SPLICING FACTOR SR-LIKE 1 (SRL1):**

GGGACACTTTTCGAAAAGGCATTTAACTAGGGTTTCGACTTCCAGCATGAGGAATGGGCATTCGAACACGCTCTCACAGTTGATGCCAGATGCGTAAAGCAGGAGAACTCGTTTGAAGCAGGAGAACCCTGACTCTGAAGCCCTAAAGCTGGAGCAGGTGAAGCCCAACAAGGCAAAGCGGTGGCCTCTCACTCTCTAGCGAAAGCAGTTTTGATAAGCGAAATCGTAACCCTCTAGCCAAGTGAAACCCTATTACGGCAGCAGGTGAAAACCCTGCCGTAAACCCTAGTACCGGAGCAGGTGAAAGCCCCCCTACCGTGAAGCAGGTGAAACCCTAATCTGGAAGCAGGTGAAACCCTAACCCTCACGCTGGTGAATCACCAAGCACAAGATGATGGCTCGGTGGAGACCGCCATTACCATTACCCTCTTCAGCATCAAGAGCTTCATTACCACCATCATCATTCTCATCTCCATCATCTGCGTCATCTTCGCCATCCTAACATGTCTAACATGGAGGTGCAAACATGTGGAAAACCCATTGATACGCTATTGGAGAAGGTCCTGTGCATGAATATTCTCTCTTCTGACTACTTCAAGGAGCTTTACAGATTGAAGACTTTCCATGAAGTTGTCGATGAAATTTACAACCAAGTGGACCATGTGGAGCCATGGATGACAGGCAATTGCAGAGGCCCTTCTACTGCTTTTTGCCTTCTCTACAAGCTCTTCACGATGAAGCTCACTGTGAAGCAGATGCAAGGCTTGCTCAACCATGGGGACTCTCCCTACATCCGAGCGGTGGGTTTCCTTTATTTGCGCTATGTCGGGGATTCAAAAGCTTTGTGGGGATGGTATGAACAATATGTCAAGGACGATGAGGAATTTGCACCTGGTTCAAGTGGGAAGAAGACAACCATGGGCGTCTATATTCGTGATCTATTGCTAGATCAGTACTATTTTGATACACTACTTCCTCGTATTCCTGTCCCTATTCTCCGCCAAATGACAACACACTTAGAGAGATTGAAGCTTTCAACAAGCTCTGCTGGTGTAACGGGTACTGGTGGTCGTCATGCAGCTGATGATACAGCCCGAAGGCCCCCATCTGTTAAAGCTTCACTGTCTGTTTCCTTTGGACAACGTGCTCCACACAGGGCCTCTACAAGGGACTCCTCACCTGTGCGTCGCAATGTTCCTGGACCCCGTGATTCATCACATCATAATGGAGATGACCGTGGGGAAAGAGATCCTGAAAAGAGCAAGGAGTATGAACGTGCGAGGGGTAAAGATCGCGATGGGGAAAAGGCAAGGCAAGCAGACAAGGATAGAGAGCGAGACAGGGAGCGTGAGCGGAGCAGAGACTATGACCGGGATGGTAGAAGGAAAGATCGAGACAGAGAGTACGATTCTCGCAGGGAACGAGATCAGGACAGAGATAGAGATCGCGACAGGGATAGAGATAGCAGAGACTACAATCGTGACAGGGAAAGGGACCGTGAGGGTGGGCATCGAGAAAGAGGCAGGGACAGAGAGAGGGATGCAGAAAAATATAGGGACAAGGATAGAAGGTCAGACCATAGAAGTCGAAGCCGTAGTCCACGCGGGCGAGTGTATCGGAGCAGAAGCAGAAGTCCTCAGAAATTTAGTAGAGGTGGCAGTGCCGATGTGCAGAGGTCAAAGAGCCCTGCCAAGGTTGCAGTAAATCTAGCCAAGCTTCGTGACATGTATGGAGATGCTACTGCCAAGAGCAATGATAATGATAATAGTTACCGTGGAAAGGACACAAGTGGGGAGGATGTTATTCGTCTGGGTGGGAATTCCTCATGGCGATAGGCCCTGCTGCCTATTATATATTGTGTTATGAGATCTTGCAGCACTCTGGCCACTATGTGATGAGGTCTTGCAGTGCATTGGTACAGTCTTTGATTTTTTTTGTTTGTAGGACATTTTTTTGCTCTTTACACGAGAGGGTTTAAGATTTTGTAAATGTGGAGTGGCAGTCTTCTATACCTTTAGTCTATTGGTACTCTTCCACAACTGAACCTACGAGCTTGTGTGGGTCCAGTCTTTGCCTGAAGTTCTGTGAAGTGTATCCTAAAAGTCATTGGTCCTTCAAATGGATGATACTTTCCAAATTGGTAAGAATTGGATCTGGGCCTATTAATTGTCTTTCAAAGGTGTCCTTTCAAGTTTTGTGAGATCCTGAATCTTACCAAATGGGGATGCCTATTTCTTTCCTAATACATAAGTACTAGTATAATGTAATAATGTCTGAAACTTATTGAAGAATTGAAGCTAGTTTTTTGAATTA

**PROTEIN PHOSPHATASE 2C GROUP 1 (PP2CG1):**

TCTCTCTCTCTCTCTCTCTGCGTGCAGGCTCAAATCATCTGCTGCGAGTCGGACGTGCAATTGCAGGCTGGATTGAGGGTTGTCTCCCACTTTTTGATCACAAGTGTTGTATGGAAGCATAGAGGAAAGCGTGGAAAGCAGGCTTGAGGACACATTGTGATCATTAATTATTGAAGCAGGCAACATAGCCAGCATCTTCATGCACAGCAGGGACATCTCGGATGCCAGCCGGTACCGGAGTAACTCAATCTGTTGCGAATGTTTTCGAAGGAACAACCATGGAGGGAAGCTCTGTTGGAGATAACATCGCCAAAAGCTCATCAACATCCTTAGCAACTTCATTGTCGTTAGAAAAATATGATGCCTTCAATAAGGCAAAAACTGTATCTCTTCCCACAATTCTTGAGGTTTCAGTCAATGGGGGTTCTACTCCCAAGGCTGAACCAGAAAAAGTTGAGGTTATTAAGGAGGACTCTATTGCGGTTGAATTCAAGCCTGCAATTCGCTCGGGGGTGTGGAATGACATAGGGTCGAGGCATGTCATGGAGGATGAGCATGTACTCATTGATGATTTGGTGGAGCATTTGGGTTCTATGTTCGCAGGGGAAGCCTCAGGATCATATTTTGCAGGAGAAGCCTCAGGATCATATTATGGAGTATTTGATGGTCATGGTGGAAGGGATGCTGCTCAGTTTGTCAAAGAAAAGCTGCTTAACTTCATCGTAAGAGACGTGGCTTTTCCTACTGCAGTTGAGGAAGCCGTGCACCATGCATTTTTGGATGCCGATAGAGAGTTTGCAGAGGCTTGCTCATTGGATCAAGGCCTATTTTCTGGCACGACAGCGCTAACTGTTCTTGTGCTTGGAAGAAAAATGGTGGTTGCAAATGCTGGAGATTGCAGAGCTGTTCTTTGCAGGAAAGGCAAGGCAATTGAGCTTTCGCGGGACCATAAGCCTTCTTGTGTTTTAGAAAGGAGCCGCATTGAGGCTCTTGGAGGCTTTGTAGATGATGGCTACTTGAATGGACAGCTTGGTGTTGCGCGGGCTATTGGCAATTGGCACATCAAAGGTTTGAAAGGTGTAAATTGTCCTCTTAGTGCAGAACCTGAAGTCAAGCAAGCACTTTTGACAGAAGAAGATGAATTCATTATCATCGGCTGCGATGGCTTGTGGGATGTTTTTTCAAGCCAAAATGCAATAGATTATGCACGTAGAAGACTTCAACAACACAATGATCCGGAGCAATGTTCGCGTGAACTTGTGGATGAGGCTCTGCGAAGAGAGGCAGGCGATAATCTAACGGTGGTAACAGTCTGTTTCCAACAGAATCCACCTCCCCAGTTGCTAACAAAATTTTTTGTCCGGAGAAGCATTTCTGCTGAAGGACTGCGCAATCTCCAGGATGTGCTAGATAAGTCTGCTTAATTGGCAGGCTTGATTTGTATTAGGCTTTCCTGGCACGATTGGGACCAGGTGTTGTGAACTACATTTTTGAACAAGACGACGTTTTCAGGAGAGTCAATTATTAAGTTATGATGGATCAGAAAGGTATCAATTTTTGTCTAATCAAGAGAACTTTTGAGTTGTTGATGACCTGAGTATGCAAATTTGGTTCAGTGGCTGCGGAAGGATGGACTCCAATACGTGGCCACAGTTTTTTTGCCAAGCCAATGTCAGCCTTTGTCATTGATTTTGGACGACTTCCGGTTTGTCATCCGTTTTATACGACTTTCGTTTTGTGCAGAGCTCAAGTTTGTCTAGGCAACAGCAGTTTCAAAAAGAAAAGGGGGAAGGAGGTCCAATTTCATACATGACGTGACCTGTTTATAAGAGAGCCCCGAGGGCCTGTCAAGGTCCTTCAGGCTTGAGTGCAGAAGCTTCCATTGCACTTGTTTTGGGCTTGGTTTACCTTAGTGGATTTTGGTTCGAATGCATTTTCTTTCATTGAAATGTCAAAGTCTAGCTTGCATTTCTCCCTCTGCTCTCCCGCTGAGGTTCTGTGTCCCCTCTCCAATGTATTTATCCTGTACATGAGCAGATTAATGTATGTAGGAATGTTTTGCTGAGCTATGCAACCAAAGCTGTGTGTGTACAATGTGACTGCACACATTCTTGCTTGGCAAACCCTGAGCTTTTAGTCTAAGGTGGACATAGCCAAACAAAATGTCATAGGTAAACTAGAAATTCTACGTTTCAAGTTTTTTCGTTGGAGTGATGCGAATAGTTAAGAGTTAC

**SUPPRESSOR OF ZEAXANTHIN-LESS 1 (SZL1)**:

GAGAGAGGAGAGAAGATGGTCTTATCTTTTTCTATGGCTCAAGATACCATGTAGTACACGGTTGTTAAAATGTGGAAGCTCTCAGAGGATTCTCTGTGACAAGAAGGAGATATGCCCTTTTGATTTTGGCCTCCAGTGCATATCATCAAGGGTTTTTCTTTAAGGACTCAACATTCCTTCAGAAGTTGGAAATCTTCACCACAACTTCTCAATTGCCCTGCCTTCCCCAGTATAAGATGGATGGGGTCATCTCCCAGAAAGCTGCCAGTTTTTGCAGCCCTCGAATTCTCCACTCAGGCGAAAAGGATCATGAGCTATTGGGCTTGAGCTCTCGTGAATTTACATGTGGTAAACAGAAATCAATTTCTGGTGTCAACTTGAGAAAAGGGCATGCTCTGCACCCTTTGTCTCTTCGGGCCAATGCCACCATGACCCCTCTTGAGCTTGCTCCTCAGGTCAAGAAAGAGAGCCTCTCCTTTGAGCTACCCAAATATGATATTGCAAAGCAGAGCAAAGTAGATCTAATTGTCGTGGGTTCGGGCCCTGCTGGCCTCGCCGTTGCAAAACAAGTTTCAGAAGCTGGTGTGAGCGTTTGTTGTGTTGATCCTGCCCCACAAAGCATTTGGCCCAACAACTATGGTGTTTGGGTCGATGAATTTGAAGCTATGGACCTCTTGGATTGCCTTGATACCACATGGGCTAGTGCAGTAGTCTACTTGGATGAAAATCAAGAGAAATCCTTGAAAAGGCCATATGGTCGTGTGGATCGCAGACGCCTGAAGGCTAAAATGATGGAGGGATGCATTAAAAATGGGGTCAAGTTTCATCAAGCCAAGGTGTCGAATGTTGTACATGAGGATGAGTACTCATCTGTTCTTTGCAGTGATGGCTTAACCATCAAAGCCAATGTGGTTCTAGATGCAACGGGTTTCTCAAGATGCCTGGTGCAATACGAGAAGCCTTACAATCCAGGGTATCAGATTGCTTATGGCATTGTAGCCGAGGTCGATAGCCATCCATTTGACGTTGACAAAATGGTTTTTATGGATTGGCGTGATTCTCACCTCAAGAATGACCTGGAGCTGAAGAGCAATAACAAGAAGCTGCCCACCTTTTTGTATGCAATGCCGTTCTCGTCAAACAGGATTTTCTTGGAAGAAACCTCATTAGTAGCGAGGCCAGGTGTGCCGATAGAAGAGATTAAGAAGAGGATGGTTCAAAGGTTGAGGCATTTGGATATCAATGTGAAGAGCATTGAAGAGTATGAGAAGTGTGTAATTCCAATGGGGGGACCACTTCCTGTGATGCCACAGAGGGTGGTAGGTATAGGGGGTACAGCAGGTATGGTTCACCCATCTACGGGGTACATGGTTGCACGGACTCTTGCGGCTGCCCCTATTGTTGCAAATGGAATTGTGCAGAGGCTTGATAGCAATAAAAGCAACCTCACGTCTCTCTCAGCAGGAGTTTGGAGGGACTTGTGGCCTATTGAGAGGAGGCGGCAACGGGAGTTTTTTTGCTTTGGCATGGATGTTTTGCTTTCTCTTGATTTGGATGGCACCAGGCGCTTTTTTGATGCGTTCTTTGATCTGGAGCCGTATCAGTGGCATGGCTTCTTATCATCAAGACTCTACCTTCCGGAACTCTTTGTCTTTGGTTTGTCGCTCTTTGGAAATGCATCCAATACTGCTCGGCTCGAAATAATGGCAAAAGGCACTCCTCCTCTGGTAAGAATGCTAGGAAATCTTGCAAGCATGAGAAAGTACAGGAACTAAAGTGGTAACTGTTCCTATATTTCTCCTTGAGCTTCCAATCAGTTTTTCGGCATTGTTCATCACCTACTTAGTACTGACATGCATGCATATTATTTCAAGGGACTGCAAGGTCCTTGTGTATAACAAACAGCAAAGACATCGCATCTACTGGATTGTGAGCAGTTGTTTTGTCATATGTATATTTGATTTGATTGTCTTTTCTGCAAATGCATTCTCGATGACACTTTGTTTTTGGATGCCATACCACTTCAAGTATGTCATACCTATGGAGTTTTAATATGTAGCTGCAAATTCTTTCAAGGATGTTAACTATTGTTCACAGCCGGTAGCCTTGGAAAGGGTGTTGTATGTTTGTCAGTGAAAGGAAAGCCATGAAAGGTTGGCATATAGGCCTGGCTTCACTTTGGTATGGTGATAAACGTTTGTCATTTAGAGTGGATGTAGTGACTTTGACTGATTTGGTATTGCCTTGTTCCTCTCTCAGTTGCAACACGCTCTTCGTCTTTCACATTTAGTCCACGCATGTATGCTAAGCTGTTTACCCTATCTTTCCTTCACGGAAAAGAAATTCTAGTTTGTTCTCAGAGCGTTCACAAGCAAAGACGTTTGATTGTAAGTATTGGACAGGCAAGGAGACTCAACTAGTATATGTATGTTTTCATGTATTTGTTCTATGCTTTACACGGGTATCTTGGGCCAGAGTCCACACTTTCATATTGAAATTCAACATCTGAGAACCTGTTTGGGTTATGTGCATTTTGTCATTCCATTTCAGCTCAAAGGCTGGCTTGTGTCTGCTGTAGGTCAATTCTAGTGCAATATTGCGATATGTTACTGGGTACAAATACCATCTCGCATGGAAGCTTCAAATTAGGCACACTACAGTCTACCTCACATAGCAACTCAATAACAAGAGCCTTTCTGATGCTCCACGCTTCAGTCCAGCGTGATTGGTGCTGCCTTCCAATGGGCCTTTCTTCTCTTAGGGTTGGCAGAGGGAGAACTTGAGACCCTTAATGAGTTCTCTAGTAAAAGCTGATAGAGCCAATGATTATCGGAACGGCAGCAGCTGGATTTTTTTCCCCAGCCTGCTGAGTTGTCTACAGGAGCATAAGTCAGTCATACTGCAGAAGCTGATATGTCCAGGTGAGATTAGTTTAGCTGTGAACATCCTCCACCTCCGCTTGGCCCTGATGGCCAGGAACGACTCTGCACTGGTCTTTTCCAGTTGGGAACAGGCTGTACGCTCACGCCATTGCTTGTTTTCTTCCTGGGAAAGGCCAGTACAGGGCTTTTCTCCACTTCCACATCACTGAACTGAAGGTTTCTCAAATGACCTGCACATGTAGTTTTGGCCAATAAAGATGATGCAAAGACTTGAGCAGATACAGTGAGAGCGTTCGATCTTTAGCAGTTTGAGTTTGTGAAGGGCTTTCTTTATTTCCTCTCTGTTTTGAAGATACTGGCAGGGCATGTAGAACTTGTTTTCTTTGGATCTTGCTTGTGGGTTTATTTGGCACGTTATTTTTGGGTATACCTCATGTAAGGTGCATGTTTGCCTTAGGTAAGGTGCACGTTCATGGGAGAAGGGATAAACAAGATGGTGCTAACTTGCCAAGGCCGTCGTAACACTAGCAGATGGCAAGTCTATCTGAAAGCATCCCAAGATGTTGTGGGTTTGCAGGTGAACAGCCTCTCCACATTGAAGCCTGAGGAGACAACTCACCTCTAATCATGCTTTTATTTATTTATTTATTTTGTGGGGAAGGTTGGTTTTACATTGGGTGGACGAGCTTGAAAGAAGCTCTATGAACTGGTGGAACCTTTTTTAAAGACTCCACTATGAAAATAATGGTCATATGCATCTTGAAAGTACATCAAAATAAGCTACAGATAAGAGGTGGTTTGCGTTCTCTGTACTCTGCACTTTCATCCTTGCAGGTGGTAAAGTTGCAGATTTGACGGTGTTTGATTG

**PROTEIN KINASE 2 (PK2):**

ACACACACACACACTCACTCACACCGATATCCCTCCTTTTTGCTTTGTTCTCACATTCTGTTCCTTACAGCTTCTTTCTGGGTTTGTCCCGCTTCTTCAATTACTTGCCAAGAGATCCTTCTCATGAGTCTGATCAGCAGACACGACTCTCCGTACGCCACCATCTCTGTGGCATGCCTCCCCTTCAAGACAGGACCACCCTTCAAGACAGGACCACCGTTTTCTGTTAAATCTCCCATTAAGAAATCTGCAGCAGCATCGGGCCTTGTGTTTCTCTCTCAAATTGTGCACAGGGTTGCGGTGCTTGGAAGGTAGCCCTTTCTAGTAATCTTGGCTGTGTTGGGACAGAGCGTGTGCGCGTAATGGCGGGCTTACATGCCTCTTCTAGCATACAGATTGTAAGGAAACTGAAGCCATCAAGCAAGCTCACCATTGTACCTCCTGATGATAATGAATGCATCGAAATGCACATAGGATCTGGAGAGTTCGATTTTTCTGAGGTATTTGGTCCTCTTCCTTCTCCTCATCATCACCTTGGGTCAATTACGGAAACCGAAACTGGGCCCAAAAGTTCGCCCACCTGGTGTGTAACAGAGCTTGCCGCCGGTCGCAATCCTGCTGTGATTTGCACACGTTCGCATTCATTTGTGGGGCCATCACCCCGTCCAAACATTGTGCGACAAACAAGTCTTCAATCATGGGATGATACTCGTGGATTAAGATCGTTTGATGCTCTTCGGTTTGCCAAGCAGGGTGTTGCGGTCGTGAGCGTCTCTCCTCCGCGCATCTCTTTCAGTGGTTCAGATGAGGATAGAGATGATGTAGAGGATGGGGTGTGCCACTCACTTCGTGACAGCGAGGGCTCGGTGTCAGATGGCTCAATCCCTACACAGCTTGATAATGATGATAATACAAAGAAATTGGGCCCTCAAGACTTCAAACTTCTTCGTGTTGTTGGGCAAGGCGCGTTTGGTAAAGTGTTTCAAGTGCAGCAGATTGAAACAGAAGATATCTTTGCAATGAAAGTCATGAGAAAAGACACGATAATGGAGAAAAATCATGGGGAATACATGAAAGCAGAAAGGGATATCTTGACCAAAATTGAACATCCTTTTATTGTTCATCTCCGGTACTCCTTCCAGACTAAATCAAAACTTTACTTAATCTTGGATTTTATTAATGGGGGGCATCTATTTTTTCAACTTTATCGCCAAGGGACATTTAGTGAAGAGCTGGCTCGGGTGTACACTGCAGAAATAGTCTCTGCTGTGGCTCATCTTCACAAGAATGGCATTATTCATCGGGATCTCAAGCCGGAAAATATTCTGCTGGATGCAGAAGGTCATGTGAAGTTGACAGATTTTGGCCTTGCTAAGGAGATTGATGATTCTAAACCAGCAAATTCACTATGTGGAACTGTGGAATACATGGCTCCTGAAATTGTCCTTGCAAAAGGCCATGGGAAAGCCGCTGATTGGTGGAGTGTTGGCATTTTGCTCTTTGAAATGCTTAGTGGTCAGCCCCCTTTTACTCACAAGAATAGACAAAAGCTGCAACAGAAGATTGCAAAAGACAAAATAAAGCTTCCATCCTATTTGACGAGCGAAGCGCATGCTCTTTTGAAAGGGTTGCTCCAAAAAGAAGCTGGTAAGCGGCTGGGGAGTGGAGTAAGGGGGAGTGATGATATTAAGGAGCACAAATGGTTCAAGCCCATCAATTGGAAGAAGCTGGAGGCGCGCGAGATCCTACCCAAGTTCAGGCCTGAGGTTAGCGGTCAAGACTGCACTGCCAACTTTGATGAGAGGTGGACCAAGATGCCCGCGCAAGATTCACCCGGCTCTACTCCAAAGGCACAGGACATGGGTGTTTTTGAGGGTTACACATATGTGGCTCCTAACCCATGGCTCACCTTGCTTGATAAGGCAAACATTAGTGATGCAAAGGGTCTTTGAATTGGCCATGCTCATTTTGCGTATTTTGCCCCTCTTTCATGGTTTGCTTCATCGGTGATTCTAGATGACAGGTTGTAACTTACTTTATTATTTGCTGACATGACTTTAGGTAATCAATCATTTTGGACGTTTTAACTGTTTTTGACAGACTCTCATTCAGCCATTTATCAGACACCTTGCGTTCTAGTGTAATGTTTGTACATTTCTTGCAAATATGTCAAATGATTTTTTTGGGATACTGCGTGTGTATGCACACACATTAGTGATGTCCATAGTTGTCGGGTCTTGTGTTTCAAGAGTAAGCCAAGTGCACTGCTTGTGCCCATGTGAGTGCCGGAATTGGCATGGAACCCAGCTAGGGCAAACGCTCTAATGGCATGTTAGGAAAAAGACACGATAAAAATACAGGCTTACTGTGTGTTAATTGTTGTATAACAGATATTTTCAACCAAGATCACATTGAGTT

**CALCIUM BINDING GTP-ASE (CBG)**:

ATTTCAGGTCAAGTGCTTCAATGCCCGTTTGCAACCATCTGAGATTGTGGGAGTGAAGAAAGTTGTTGCTGAGAAGTTTCCTGAGGGTGTCAATGACAATGGCTTGACATTGGCAGGCTTCTTGTTTCTACATGCACTTTTTTTAGAGCGTGGTCGTCTGGAGACTACTTGGACAGTTCTTCGAAAGTTCGGATATGATGACGAGCTCAAACTACGCAATGACTTGCTGCGAACTCCATCATTCAAACATCTACCTGATCAGAGTGTGGAGCTAACTGACACTGCGACTGATTTTTTGAGAGAGATCTTTGCTTATTACGACCTTGATGGGGACGGGGCATTACATCCTCAAGATGTCGAGGAGCTTTTCTCTACTGCCCCATCAAGCCCTTGGGAGGATGTAGTTTATCAAGACTCGGTTGAAAAGAACTCCACTGGTGCCCTCACCATCAAGGGTTTCTTATCGCAGGCTAGTAGGGCAAAATCTGAACAGAGTTCCTTCTCTGGACTGTGGCAATGTAGAATGTTAAAATTTTGGTACTCCTTGGTGCGGTTGTTACTTTACAGAGTTTTATTGACATATGCAGTGGGCATTGATGACCTTGCTGGATCCTGTCAGAAGCCTTGCTCATTTTATCTATCTGGGTTACCCCGGGGATTTGACCACTGCGTTTCAAATTACTCGACGGCGATCTATAGATCGAAAAAAGCAGTGCTCACTGCGCGTGGTGACACAATGCTTTGTTTTTGGTTCAATCAAAGCAGGAAAAAGTGCTCTTCTTCATTGTCTAGTTGGGAGGCCCTTTGCAGATGCATATTTGCACACAAATGAAGAACATTATGTTGCAAACTTTGTCGAACAGCTAGATATCCCTATAAGAACTGCCCCCTTGTGACTTTAATGTTTACAAATGCTGTAGGGCTCCAATAAGGTTCTTGTTCTTAAAGAGATTCCAAGGGATGCGGTCTCAAAGCTTTTGGAGAACAAAGATGCATTGGCCGCCTGTGATGTGGCAGCCTTTGTTTATGATAGCTCGGATGAGTGGTCTCTAAAAGATGTGGAAAACCTTCTGCTTCAAGTGGCAACCCATGGTGAGAACGGTGGTTTTAAGGTGCCATGCCTCGTCATCGCAGCGAAAGATGAAATAAACTCAAATGTTGCAGCTATTTCTTCAGAGTCAAGGAGGGTGTGCCAAGAGATGTGTGTGGATGCTCCCATTGCAGTCAGTATGAAGGTGGGACATATCAACAACATGTACGCATGTATGGTTTATTTAGCATGCCATCCCCATTTGAGTATACCCATGACAGAGGCTGGAAAGAGTTACCAACATTGCAAGCGAATTGTGGGACGCTCTCTCTTAGCTCTTTCCGTGGGAGCTGCAGTAACTCTAGTGGGAATTGCAGCCTACAGGGCTTATTTGGTGCATCGACATGCACACAGTTGAAGCTCTAATAGTCAAGTATCTATGCTTTGCTTAAGGGATTTATGTAATTTGCACACTTGGCATTTCAAAGAAGCACTTCACCATGCAGAAATGCAGTCTTAATATTTACGTGATGAAGTCTAAGAATTTGGTAGCAATCGCAATGATTAGGCAGAAAGAACTCATACCCAAAATTGGCGAATATGTGTTGTGATAATCAACGCCAAGCTTCTATGTGAATCCACAG

**NF-X-LIKE 1 (NFXL1)**:

CACAGCAGTGCCGCGCAAGCTTCCTTCCCTTCATAAAGGTTATTAGTTAGACAAGACGAAGCAGCACGCACACAAGACACACACACACAGGCTACAGCAGTGCCGCGCAAGCTTCCGCCATAGCCTTGTACAAACCCTAATCCAACGCCCATTTTCATATTTTCAAGCGCTCCCTTACCGCCATTGGTCCTTGTGGAGAAACCCATTTTTCTCTTATAGAGAAAGAACTCATGGACGATGTCTCCTTGGGCTCCCGCAGAGGAGGAAGATCTTCGCGTCGCGGTCAGCTTTGGCGTCCTCAACAGGGTCGGGGTGCTGTTGCTTTTCAAACCGGGTATGCAAACCCTTTTCCTGATGCACCCAGCCTTCACCTTCAGCACATGCACAATGATGCACGTGTGAATGGGGGTTTCATGTTGAACCCGGGTGCAGAAAGCTCTGGGTCAATGAGCTTTTCTTCTCAGGATGTGAGGAGTTCTGGTTCACATGCACAAGCGGATGTACGCAGAAGGGAAGATATTCATAGAGAGGAAAATTTTGTGCTCCAGAACATTGATATGTCTAGCAGACGTGTACACAGAGGGGGTCACCCATTTATGCCAAATGAAAATGCTGAAAGGCCAATGGTGGCAGAAGGAAATCCTTTGAGCAATGGAGATCACAGGAGAAGGAATCATGGCAGAGGAGATGCCAGGATTCCCAGGCATTCAGAAAATGATATGTTGAGAGACAATAGACTGGTGGATGCAGCTGAATTTGCAGAGATCAATGCAAGTGCACCTGATTTTGTCAGAAGGACACAGATTGCAACAGGTGGGCGTACTGGAAGGGTGGAATCCAGAGATTATCATCCAGCTTTTTCTGGTAATGGAAGGCAGCAATGGAGACCGGCTTCTTCTCAGGGTGCTTATCCAGGTAGCGATCAATCTACTGGCCAAGGCATGCTCACTTCAGAAGCTCTGGAGGCATCCCGAGGGGCAGTTGATGAGGTTCTACAGCACCTTTCAATTCGCAGAGATCAACAAGGAGGAGGATCCAATAGAACAGTAGATACACATAGGCATTGGAGCAGCCAGAAGCATTTCAGAAATCGTGCAGAACCAACTAAGCAAGTGCAGGCTTTTATTCCTTCGATTGTCGTCGTACCTAAAGACAAGACAAACACTCCACAGCTTGTTCAAGAGCTGGAGGACAAGCTGAGCAAAGGGAAGATTGAATGCATGATCTGCTATGATATGGTAAAAAGGGAGGCTGCAGTGTGGTCATGTGGAGGGTGTTATGCAATCTTTCATATTGCCTGCATTAGGAAGTGGGCAAGGGCACCCATATCTGCAGACTTCTCTCTTTCTTCATCTAATGAAGTGTCTGTGGGGAATTGGAGGTGTCCTGGCTGCCAGTCACCTCAGTTTTTAAGCGCTGAAGAGATTAAGTACAGATGCTTTTGCGGTCAAATGTTAGAGCCTCAGGTTGATTTTTATCTAACTCCCCATTCCTGCGGGGGACCTTGCAAGAAATCTTTGAGCTCAGCAAAGAGTCCTCATTGTAAACATCTGTGCACAATGCAGTGTCACCCAGGACCCTGCCCTCCTTGTAATGCCCTAGCTCCTCCTCAATTTTGTCCATGCGGAAAGACCAGTTACACTCAGCGTTGCTCGGAACTCAAGAGCGCAAAGAGTTGTGGGCAGATTTGTGGCCATCAGCTGCAATGTGGTAGACATTATTGTCAAAAGATCTGCCATGAGGGCCCTTGTGGAAGCTGTGATACTTTACTCACAGTAAATTGCTTTTGTGGTAGGAAAAGAGAAACTATGGCATGTGGTGCAATGGTCTTACAAGGTGAGCTTGACTGGGAACATGGTGTGTTCTCGTGTGAAAACCCATGTCTAACCAAACTGCAGTGTGGAAAGCATGCATGTGGCAAGAAGTGCCATCCTGGACAGTGTGGAGAATGTGAATTAGCTCCAGGAAAGGTTCAAACATGTCCTTGTGGCAAGGCTTCCATTCAAGAGTTGTTGGGAGCGGGCAAGCAGAGAGAAAGTTGTACAGATGCTGTGCCATGCTGTGGCCAGATTTGCCAGAAGCTTCTTCCTTGTGGTAAGCACGTTTGTCAAAGCTTGTGCCATGTGGGAGGTTGTCCAGCATGTGAAGTTTTGGTGGATCAGAAGTGTAGGTGTAAATCATCTTCTCAAACTATGCCGTGCCATAAAGCTTTGGATCTTGACAGAGGCAGTAAAGAAATTTTAGCAGATGAGCCTGGTGACGGATTGTTTCTGTGCAATCGCAAATGTGGGAAGAAGAAGAGCTGTGGACGGCACAGGTGTAGCAATCGGTGTTGTCCACTTGCCAATGAAGATAACGCTTTTCTGATTGTAGAGGATGATCCTCACCTTTGCTCGCTGCCTTGTGGGAAAAAACTGAGGTGCGGGCAGCATACATGTGAGGAGTTATGTCACAATGGACATTGCCCTCCTTGTCTAGGGTCTACTTTCACGGAGCTGTCATGTGCATGCGGGAAGACATCTATCCCTCCTCCTATACCTTGTGGGACATTCCCGCCTTCTTGTCCCCACCCTTGTTCTCGTCCCCAACCTTGTGGTCATGCTGCAACTCATCAATGTCACTTTGGAGAGTGCCCTCCATGCATTGTTTTGACAGCCAAGGAGTGTGTTGGAAGTCATGTGATTTTAAGGAATGTTCCTTGTGGTTCAAAAGACATCAAGTGCAACAAGCTTTGTGGAAAGGCCCGTCAATGTCGACTGCATGCATGTGCGCGACTTTGTCATCTTCCTCCCTGTGAGGTAGTGTTAGATACTGATGATGATAAAATTATGAGCTGCGGACAACAGTGTGGAGCTCCTCGGAGGGATTGTGAGCATGTATGTATTGATCCTTGCCACCCTTGTTCCCCATGTCCAGAAACAAGGTGCAAGTTTCCTGTTACAATAACATGCGTATGCGGTCGGCTGTCACAGCAGGTGCCTTGTGATGCTGGGGAAAACTCAAAGGGAAATGATATACTAGAGCTTGCATATATTTCAAGTTTATTATTACAGCCAGTTGATGTGAAGGTCAGAATTCCTTTGGGTCAGAGAAAATTGGCCTGCGATGAAGATTGTGCTAAGCTGGAGAAGAAGCGTGTCCTTGCTGATGCATTCGGCATGACAGGTGTGCCAGTTGATCCATCTCTCTCGGCTGAAAGTGTTCAAACTGCCTCAGAGGTTTTGAAAGAACTTCTTATAAGGGATCCACAATGGATTCTAGCTATTGAAGATCGTTTCAAATATTTAGTGCTAGGGGCAAAGTCCACGACTGCACCACTTCGAGTACATGTATTTGGATTTCTGCCAAAGGAGAAGAGAGAAGTGGTTCGACAGATGGCTGAAAGATGGAATCTGACAGTGAGTGCTGTTGGCAGGGAACCCAAGCGCTTTTTGACTGTCCATGTGAACTCAAAGTCTCGGGCTCCACACATGAGAATGTTGTTTGGAAGATCGGCCCTTCCGCTTCCAGGCCACGCTCAGGCTCCAGCTTACAATCCGTCTGTAGACATGGATCCAAGGCTTGTTGTTGGGCTGTTTGAGCTCCCGAGAGAAGGAGACATAAGCACTTTGGTGCTGAGGTTTGGTGGGGAGTGCGAGCTCGTCTGGTTGAATGATAAGAATGCCTTAGCTGTCTTTTCTGATATGATGAGAGCCGCAACCGCTATTCGCCGTGTTGATCACGCATCTGCTTACAGCGGAGCTGTTCGAACATCTTTTACGGCTGCTACTACAGCCAACGTCCGGGGGTGGGAGGGTGTGACAACTTCCAAAGGCCCATCCCGCAAGAAAAATGTTCAAGAATCAAGCTGGAGTGAAGATGCCTGGGGGGAAGAATGGGGAAGGTCGCAGGCGAAGACAGCGTGGGATACAAAAGGTCCTGCAATTACAACTTCTATCAATCCATGGGGAGCTTTACAGCAGGATGAGACTGTGACTAGGAACTCCATTAAGGTCATTGAGAAGCCCTCCTCCTCTACCACAGCTCCTAGTGCTACCGAACACTCGAAGAGCCCTAGACAAAGTGTTGCTGCAGACAGTTGGGAAGATTTAGACAGTTGACCAGCAGCTTGAAGCACAAATATGTGCAGAGCATTTTGGGAGCCTGGTTGAAGAGCAACTGGTGTGGAGGCTTATGATCCCCGATTTTGCGCGTGATTCTTTGATTATGACAGGCAAGAATTTCTAAATGGATAATGTATTTGAGAATTTGTAAGCAGTTTTATTACCATCCTTCGAAGTTTTTGTGGTGGCTGTATTGTTGACTTAGCTCATAAGTATGTTTTTAATGGCGCAGTTTCAGTGTTTTAAATTGATAGTAGTATGTGCTCATGTTTTTAATGGCGCAGTTTCAGTGTTTTAAATTGATAGTAGTATGTGCTC

**PEROXISOME UNUSUAL POSITIONING 2 (PEUP2)**:

GAGGGCTTACGGACATTATAACCCTCTCAAAATGTAAATGCCTAAGACAATCATAAAAGTTTTTTTTTCTCCTTAGTATACATGTGAGGAGAGCGTTTTGAAAAGTAGATACTTTGCATCTTTACAAACCTGTCAATGATCTATATATCGTTGTGCATCAACACCTCAACTACATGAAAGATGATACATAAACCACGAGGTTGATTGTCAAAAACAACAAAGAGGTGCTAAAATTCAGTTCTACTCTAATTACACTAGTCTGAAATATCAATGTATGAGACCGAGGCCCACATTGCCACGGGCTTTCCTGCGGAGCCACAAGACAAAGCTAATTGCATACCTGAAAACTCCTGTCCAAGATTGTGTAGATACAAATAGGCCGTATCAGTCTAAACAATGGGATACTCCAGAGTCCAGACATTATCAATTAATAGTGCTTATGTGGCGCAGGTAAAGAATTTCAAAAAGAGAGCTTATATACAGACTCTGTGCTGAAGTGGCCAACAGCATGAGATACAGCCCCTTTGAAGAGAACGTGATGGGAAAAAAGGCAATTGCAGCCAGACCTTGCACCAAGCAACAAGAAGACCACATGCACAGATGGCAAAGAAGAAGACCTTCACACAAAATACAAGGTTTTTCTGGAATCCCATATTTTATATGCAGACTGCTTTGCCCATGGCCATTTCCAGTGTTAAACACAACATGCACTTTCCCTTCCCCAAATCATTGCTCAAGGCATCCCTTCATTCTCTAATCTCAAAAATCTAGCAGATTCCTGTTGTTTCATCTCGCCTCCATGAACAGGATCAAAGGAGCACCTGTAGAAGCTTCCATCCAAGCCAACGATAACTACTGTAGTCTTATCCGGCCCAAATCCAACCAGTGTTCTCACATCTTCAGGCAGCCGATACTGCGCAAACGACCACTCTGAACTGAAATATCTTGGAAGCACTCCTTTCAAGAAGGATAAAGTGGAACTAGGATTGCTGCCAGAAACAGAAGAGCTACTACCAGGCGGGCTATTACCAGTGTTTCCATTCCCAAATCCCGAACCTTGCGATGATCCAGAACCATTGTGTCTCGGTTCATCATTGAGTATAGCACCTGTAGGACCCCTCAAAGTGAATACATGCACTGTCCCCTTGTCACTTGACAAGGCAAGCCATTGGGCAGTTGACGAGAATGCAATGCTGTAAATCTCTGCCCTGTCTGCCCCACGTCTAACCTCTCGTAATCGAGTCCCATCAAGACTGTTGAAAATCCGTACGAGGGTACCCTTAAAGCTAGCAGTGGCAAGAAGCTGTCCATCAACTGTGAGTGTAAAGCAAGCTAGATGGGAATCATGTGCTGAGATAAACTTTGTCTTCTTCAAACCATAATGCTCCACCCTCACTTGTCCACGATGCAAGCCTAAGCAAGCGAGTACAATCGAATTTGACGCTGCAGAAAGTGCACACAAGCCTCGAGGGTTCGCAATGGTCTCGATTTGATGGAGCAGTTTGAGGTCAACAAAGTTGTAGACATAGATTTTGTGCTCCAGGATGACGACAATGCGATCACGGCGAAGGCGGACAGCACGAACTTCCGAGCGAAAGGAAAGCTCCCCGATGCAGCGGTTCTGGTGATCGTCCCAAATCATGACCTTGTTGGGCGGGTACATGGGCGTAGCACCTCCTCCCACAAGGGCAAGAATGTTGCACCTGAAGAGCATCTCCACCATGCCCACACCACCCTCTTTGAAGTCGCGCTGGAAGGTCTTCTTGAAAGGATCGCAATTGTAGATGCGGAAACCCTTGCTGGTTGCACACGCGATGCATCCATAGTCTTGATTGAAAGAAACAGAAAGCAAATCGTTCTCTACATGGAAAGAGGAAGAAACTTCGGCTATGGTGGGCTTTGAGGCCGGCATTTCGCCTGCATCTGATGCAGAATCTTCGGTTTCTGAAAGCGGCGACATCACACACAAACACACACACACACAGACAGAAGAGGGCCTTGCGCATGCGCAAGAGGTCTCCACTTCACTTTGAGTTTCGCGTTCCTTCTGCTAGTGAGCGTCTCAAAATTTCTGCTCGAACAAATCGCGCTGTGTGAGAGAAAGATGTAATACAATTA

**2-OXOISOVALERATE DEHYDROGENASE SUBUNIT β 2 (DIN4):**

GAGAGAGAGAGAGAGAGCGCCTCGTTTTAATAACAGGGTTAACTGATACGTCATACACGTCCTTGTGCATATCTTGTTTTCTTCTCCGCCTGCCCACTCTGTCTCATCGGATTCAGTAGGGTTTCAGTGCTTCTCTACTCATGCCAATCCTAAAGCAGTTGAACCATCTTGCCCAAAGATTCCATGGCAAGAGGTCCCTCACAACCCAAGCGCACAACGTGAAGAGGATGAACCTCGTCTCCGCAATCAACAGCGCTCTTCAGATTGCACTAGAAAGCGATCCCAGGTCTTACGTTTTTGGAGAGGATGTGAGTTTTGGTGGTGTCTTTCGATGTACAACCGGACTTGCCGATACGTTTGGCAAGCATCGTGTTTTTAATACTCCATTATGCGAGCAGGGTATCGTTGGCTTCGGTATTGGCTTAGCAGCAATGGGGAACAGAGCCATAGCAGAGATACAGTTTGCTGATTATATTTACCCAGCATTTGACCAGATTGTCAATGAAGCTGCTAAGTTCCGGTACCGAAGTGGCAATGAATTCAATTGTGGAGGTCTCACCATTCGGACACCGTATGGTGCTGTTGGGCATGGAGGACACTACCATTCACAGTCACCAGAGGCCTTTTTTTGTCATGTGCCAGGCATAAAGGTTGTCATACCTAGAGGTCCCAGTCAGGCAAAGGGGTTGTTGCTTGCATCAGTTCGGGATCCCAATCCTGTAGTATTTTTTGAACCTAAGTGGCTCTATCGCCAAGCAGTTGAAGACGTGGTTGAGGATGACTATGTGATACCACTTTCTCAAGCAGAGGTGGTACGCACTGGAACGGATATAACCGTTGTTGGTTGGGGAGCGCAAATGACCATACTGGATCAAGCATGTTCTGATGCCCTAAAGGATGGAATTTCGTGTGAGTTGATTGATTTACAAACATTGATACCATGGGACAAGGAAGTCGTGGAAGCATCTGTGAAAAAAACTGGACGTCTCCTTGTAAGTCATGAGGCACCAATAACGGGAGGATTTGGCGCTGAAGTAGCAGCTTCAATAGTTGAGCGCTGCTTTCTTCATCTCGAGGCACCGGTTTCGAGGGTGTGTGGCCTTGATACCCCATTTCCACTTGTTTTTGAACCATACTATATGCCGACAAGAAATAAGGTGTTGGAGGCAATCAAAACGTCTGTAAAATTCTAAATCTTCACATACAGCTGTAAATATCAAAATTCTATATTGTTAACGTCCATTGAACATGACATACCAATGTGTTAATATACACAGTTGTATGTATATGTATACTGCCTAAGAGGCCTACAATTTAGGCTGGCAGCTTTGTATAATATACACTCATATGAAAGACCCCAATGGGTTTGGTTTGTAGCTGAATGAAGCCACATTATTGAAATGCAAAGATCACTCATGTACAAGGGTAAGTTTTGGAGTGTGACTCCAATTGC

**SOS2-LIKE PROTEIN KINASE 17 (PKS17):**

TTATAATAGACATCCTGAAAATCCGACTGCCTATCCTGCAAAGCAAGCTGTCTCAAATTTGATGTCAAGAAATTTGGGGATTTTTTTTGGAGAGAGAGAGAGTCTACCTTACATATACCAAACTTCACCAAGATGAGTTCAACACCATGACACATAGCAAGTCAACAGATAGTAGGCTACAAAATAATTTGCCATCGTACTTTACAACATCGAAAGCTAAATTTTATGGATGACATTGACAAGGCTGAAAATATTTTTTTACTTTGAAATTACGCTTCCAATTCCAATTCCAATTCATTTTTCCACACTATGTCCTTCAATCCTCTCGAAAAATTCTGGTAAAAGTTATGGTACTCCAGAGTGTCACCGCCAGCCTTTTTCACTTCCACCATGTACAAGGAAGGAGCCACTTCAAACACCTCAGTAGCTACCGAAAGATGGCCTTTACGCCCCGTCTCTGCACCATGTAACCTCATCTTGTAGTCTTGTCTTTGGACATTGAAACCAAGAGGCTTAGCCGTCTCTTCCATTTTTGAAATGATTTCTGGAGCTGGATGCTTTGATGTGAAGTGTAACTCTTGTTTTACTTCCTGCTTCGTCTCAAAGAGTCCAGAGAGATTGAGGCCTTTTGATAAGGAGATAAGCTCAAATGCATTGAGAAAAGCTGGTTTTGAAGGTTTATGGTCTTTACATTTTGCTACAAGATCTTCTTCACTCGTTTCATTGAAGACTGCATTTACATCCTCTAAATTAGAACCTTTCTCCTCCACAAACTTGGGTAATGTGTAACCCTTCTTGAACCAAGAATCTTGTTGTATTTGAGGAATGGTGATTCTAAGTTTAGGATTTGGATTCAATATCCGTATTATTAGCTTGCGTGCACCAGACGAGAACCATGGAGGAAACCTTACTTCTGCCCTGTGTATCTTCCTGTACAGATTCATGAGATTGGAATCATCAAAGGGCAAATACCCCGCCATCAAAACAAATAAAATTACTCCACAAGACCATAAATCCGCTTTGGCACCATCATAGCCCTTGTGGACAATGACCTCGGGGGCAACATAATTTGGAGTGCCACATGTAGTATGCAAAAGCCCATCTTCCCTCAACTGTTGCGGAAGTGCACTTAGTCCAAAATCAGAAACCTTTAAATTAAGTTGAGCGTCAAGAAGAAGGTTTTCTGGCTTCAAATCCCTATGATACACACCACGACTGTGGCAAAAATCTACAGCGTTTATCAACTGCTGAAAATATTTCCGAGCTTCATCCTCTCTCAACCTTCCTTTTGATGCAATTCTGTTGAATAACTCCCCTCCTGAAACATATTCAAGAATAAAGTATATTTTTGACTTGCTTGCCATAATCTCATGCAGCTGTACAACATTGGGATGCTTTACAAGTTTCATTGTCGATATTTCGCGCTTGATCTGTTCAACTAGCTTGTCCTTGAAAATCTTCTCTTTGTCAAGGATCTTGATGGCCACACTCTGCCCGGTTTCAATGTTCCTGGCAAACTTGACCTTGGCAAAGGTTCCTTCGCCAAGGACACGGCCAAGTTCATATTTGCCAACCCTTGTTCGTAGATTTTCAGAAATTTCCATGCTCATTTTATATCAATTATTGAAATATCTCCAGGCCTTGTATGAGACAAAAAATGCCTCAAAGTCGAAAAACCCAGAAAAAAGGGCAGGATCTGAATAAAATGAAAGGAGAGTATGCTTGCTAAAAGACCAACATGAATGATGCATTGCAGGCTGCTTGTAATATGCATTCAAGAAGACCCAAATTGATATTAGTAATGTGGATGGCCTGCAACCCAAGGATGATATAGTGAACAGTAAAATCCGCTGCAGATGTTCAAATTTATAGGTCAAAGTGCCAAAGGTAAAGCTTTACAAAAATGGGTTTTAATTTAAGCCCTGATCAGCAAGGGCTCGAAAGTAGTTTGCCCCAAATGTGCAATATCTGTCTGTGCGGCTGGCGACTTGCGATAGAAAAATAGTAGGAGCTCAAGAAGCCTATACCAAACGAGGAAGGCGTTGTCGAGTTGCCTGAAGAAGAAAATGGTTTCCAAGATGGCGATTTACTGGATCGCTCAAAAAGCAAAAATGTGCTCCGTTAAGATGTTCGCCTGAAGAGAGAATCTGCCCTGCTATGATGTACGACCAATATGATGTTCGCCTGAGGAGAGAATGTGGCACAATGTGATGTACGACCCGTTGCCCGATGAGGAAAATGAGCCCAATTAATGCAAAACCTGCAGCTAGATGTTGTACGACACTTGCCCAAAGAAGACAGTGCGCCCACTTTGCTATATGCAACAAAGAAATATTCACAAAGGGGGGTGCGCTAGGAGGATCGCAAAAAACCCAAATAATGTGTGCAGAAAAAAGGAAGAGATTTTGTTGATTATTGGGCCCTCTGTTTCTTGGTGAGACAAGGTGGAAATGAACTGCCTCGGACCCTCTTCTTCTCTCTCTCCCTCCTCTAAGAAAAGATGATACAGGGACGCCCTAGGAGAGGAGGAGGAGGAAGAGGGAGAGGCAACCACACAGCAAACGGAGGGAGAGGGGAGAGCGAGGGGAAAGGGAAGGCAGGGAATGAAAGGCAAAACGCAGTGAGAATATAGGCACTGGCAGCCTCAAGGAATCCAGATCTTCCGTATAATTGGGGGGTCGCTTGCATATCCTCCTCAGTGCTCACTGAAGCATGCATATGT

**PROTEIN KINASE 6 (PKS6):**

TTTTGGAAGAATTTCTTGCATTGAGGATAAAATGTACAATTTTTCTTCCTTCATTCAATTAAAAAAAAACGTCTGATTTACATAGATCAGCCATTGCTAAACAATGTAAGCTGTGGGGTGAGCTGATCCGCAAAGAAGCAAGCCATGAGTGATTGAAACCAGGAGAAATTTTATAGAGATACTCTGGTCCATAAATTTCATGCAGCTTTACATTGCTTCAAGGCTGAACAATGATGACGGTGTAAAAATGTAGGGGATGTAAAGCATCTTCTTGCCTCTGTTTCAAGGAACTAAATAAGCACTTCCACCACAGCCCCAAGGATAATTTACAGTGTTGTAGAGTAAGGAAACATGGCATCAGCGCTTGCAAAAGATTACGAGCTGAAAATCAATGACAGATGCTAGATGAAGATTTTTTTCTCCATGTAATTGAGCATCGATCTAGAAATGCAGAGAGTCATTCTACTCACCCTTTTCCCTTGCGGCAAAGGGTGCAATAAACTAAGTCATGGGCCAAAGAATCCAGTGTAAATCTAGCTAAGGATTTGATAAGCTCGGTGCATACAAATAGGGCACACTAGGTTTGATTTCACTATTCTTGCCACATGTAACCAGGGTTTAGTCGGACATATCCACTTCAGTCTTCCACACTATATCTTTCAATGCTGTTGAAAAATTGTCACAGAACTTGTGATACTCCAACGTATCGCCACAATCCTTATGCATTTCAACCATAAAAAGTGAGGGAGACACTTCAAACACCTCAGTTGCAATTGATAAGTGACCTTTCCTACCCGTCTCCGCGCCTTGCAGCTTCATCTTATAATCCCTCTTGTGAACATTAAACCCAAGTGGTTTGGCAGATTCCTCCATCTTCGATATGATTTCGGAAGCCGGATATGTAGATGTAAAACGGCTTTCACGCTTTGCCTCCTGCTTGTTCTCAAATAAGCCAGAAAGATTGAATCCCTTGTTTAAAGCAATTAGCTCGAATGCATTAAGCAGAGTTGGCTTTGAGTTCTTAGGCTCTTTTTGTTCCGTAACCAAGCATTCACATGATGGATTGAAAACAGCGTTCACATCGTCTAAATTGATGTCATGCACTAAAGCCTCCTTGTCTGGTTTAAACCCTTTCTTGAACCAAGAGTCTTGGATTATTTGCGGTACTGTAATGCGAAGCTTAGGATTCGGTGTAAGTATCTTTGAAATCAATTTCTTTGCTCCACTAGAAAACCAAGAAGGCAATTGAAAATCACCTTTATAGATCTTCTTGTAGAGGTTCATGAGATTCTTATCATCAAATGGTAGATACCCCGCCATGAGAACATAGAGAATAACTCCACAAGACCATAAGTCAGCTTTCGACCCATCATATCCTCTATCAACTATGACCTCCGGTGCAACATAGTTAGGTGTACCGCAGGTTGTATGGAGAAGTCCATCTTCCCTGCATTGCTGTGGTAAAGCACTCAGGCCAAAATCAGATATTTTCAAATTCCCATGCGAGTCTAGCAGCAGATTTTCGGGCTTGAGATCTCGGTGATAAACTCCTCTGCTGTGACAATAATCCACTGCATTAATAAGCTGTTGAAAATATTTCCTAGCATCATCCTCCTTCAATTTGCCCTGCCGTTTAATTTTGTTGAAAAGCTCGCCTCCAGTCACGTACTCCATCACAAAGTAGATCTTAGTTTTACTTGCCATAACCTCAATTAGTTGGACAACATTCGGGTGCTTCACTAATTTCATGGTGCAGATCTCCCGTTTAATCTGCTCCACCATCTTGTCCTTGAAAATTTTGTCCTTGTCGAGGATTTTGATGGCAACATGCTGGCCGCTAAGGGTGTCCTTTGCGAACTTCACCTTCGCAAAGTTGCCTTCACCAAGTACCCGACCGAGCTCATATTTCCCGACTCTGGTTCGCACATTCGAGTTGCTCTTATCACTCATGGAGGCGACCAAGGCGCACACACCAAAGAAAGGGTGCAACCCTCTGTTTACACCCCGTTACACCGTGCACATACAGAACAGAGCAAATCTGTCTGAGAGAGAGAGAGAGA

**GENERAL CONTROL NON-DEREPRESSIBLE 2 (GCN2):**

CTCTCTCTCTCTCTCTCTCTACAGCTAAGCTGGTATATACATAAAAAAGGGCACTTGCCAGCGGTTTTGGGTTCACCAATTCACGTGAGGCTGTGCATGTTGCTCACAACCAACTAAGCACCATTAATGCAACCCATTATTTTATCCAGGAGGAATATATAATAGATTGACTACTTTGCAGAATAAAGAAGCCGATTTTTCTTATTTGTCATTGTTGCTGCCTGTTCGGAGAAGAACTTCACCAAATCTTCTCTCGGAACATCTTCAACTTTTCGGGTCTCCATATTCCTTACCTTGAGACTGTTACCACGAGCAAGATCTGTCTCTGTTATCATGACCAGCCACTTAATCCCGTGCTCATTTGCAAAGTCATATTGCTCGGTAAGACTAGGTGAAGGCGAGTGTACAAACTCTGCCTTAATACTTCCATCCCATAACTCAGAGACAATCTCCATACGTTCTCGAAGTAGACCTCCCCCACCCTTGGAGCACACCAGCACTTCAACACTAAATGACTCTGGCTCACTGTCAGACATTGCTGCCAAGATGAGTTTCTCTAATGCAATGCTTACTCCAACAGCACCTGGGGCAGAGAGAGCCATGACATTTGTCCAATGCTCTTGAACAAGATGATCATAACGACCGCCAACAGCTAGAAGCATTCCTGCTGATGAAGCCGCGAGGCCACCTCTCTTGCGTAGATAAACCTGGAAGAATATTCCTCTATGGTACTCTTCAACAGGTATCATCAACGCATCAATGTGGACATGTTTTCCAATACTCCAAACCCTCAAATAACTTTGAAGAGCAGCTAGTTCATCAATTGCACTAGAAATCTGAGCACTGTCTGCCAGCCCTCCTCTAAGACGAGAAAGAGCCACATCTGGATATCCATTCCATCTGCGCTCTACTGTTTGCAGGCGGTCCACTACAGTTTCTTGCAGTTGAAGTCCCTGCACAAGTTGTTGTCGAATGTGGACCCATCTTGATTTCCTGTCAGATGATTGAGGTGGAGCACATCCAGCCAAAGATAATAACTTTGCAACATCTCTTCTTTCGTGCTTCTTTACACCACTCCAGGACCAGATTGCTTCAAGAATACTTGAATGATTGATGTGTATTTCATATGATTCATTCAATGCACATTTGGATAAGATTTCAACAACAACCTTGATAGATTCAGCTTCAGGTAAGCCAAAAGAGTCTCCAATGATATTGAAGTCTCCCTGATAATACTCCTCTGGAGCATTAGGTCCTGCAGCTGTTCGATAAACTTGTGAAATCTCAAAGTGTTTGATTGAAGGGACCTGGTTTGCAGCAACCCAACACACAAAACGTGTCCGCATCTCTGATCGCAGCTCCAGCATTCTTCCAGCTGAGTCGATCATCATCACCATCTGCGTTGTCTATCAACCAAACTCAAGGTGCTCGTGTTGAATCCTTTTGCTCCATGGGAAGTAAAAATGTCTTTGGCAATCCTGAGAATTTGATCCTGGAAAATTGACTTTGCAATGCAGGAGTAACTCAGTTGCTCTCTTTCCATCTTTTGAGCGATCAAAGAAGTCCTTTCTTCATCAAACAATGCAGAGACAATTCGATCTAAAACATATGAATTTTCTGCTCTCTGGATAGTTCGAAGCACGTCATCAAGTGCTTCATCTTCCATTCGTGGTGGCAGATGATTTCTCAAAACTTGCATTGCAGACGGACGATCTGCAGGATTGGGTGACATGAGGTCCTGTATCAATGCTGCTTGTCGAGGATACTGAAGTGCCCATGATGACGGTGCACCTTTACTTTTTAGCTCGTTCAAAATTGTAAATCTCTCCATGGCTGTGCCAAAAGGATGCCAAAGTTCCAGTAGAATCACTCCAAGACTATACATATCAGCCTTTTCATCAATCCGTGGCCATCCCTGTCCAACTTCTGGGGCAGTATAGAGATATGTGCCGACTTGACCAGTTCCATCCAACGAGCTGCAATGTAAATCAGAAGGAAGAAAAGCTTCGCGATCTGTTTGTTCTAAGCTCATAAATTTTGCAAGTCCAAAATCTCCTATTTTGATATCATTATAAACATCAAAAAAGATGTTATTGGGAGTGAGGTCGCGGTGGATTATTCCTTGGCCATGAATATAGGCCAAACCTTCCACAATCTGGCGAAAAATACTCCAGGCAGTCACCTCATCCAAAGTATCTTCGCTAGAATCAAGGAGTTCCCTTAATGTTCCGGGGCAATACTCCATTTGAATATAGAGATATGTTCTCTCCACCACAGTATCCTCTGTATAATAAAGAACTGGGTCTTTGACAGATAGCGGCATAAAACTAGAACCTGTTTTATCCCACTCCCCAGATGTTTCCAGTTCTGATGCAACATGCGATATACCAGCCACCGTCTCAAACCAGGCCTGATAATATCGTACAATATGATGGTGCTGCAACCTAGAGAGTGTAGTGACTTCTCGTAAAATTTTGTCGTTAAGAGATTGGCTTTTATCTGTCAGCCTTATCTTTTTCACCGCGTATTTCCGTCCATCAAGGTTATTTTTGCATAGAACCACATATCCAAATCCTCCCTTTCCTAGCATAGACAGTTCCTCGAAATCACTTGCGTAGCGGGAAGTATGCCCATTTTCCTCACCAGACAGCAACTCTTTACTGGCGCTCCAGAAGCTTTTGGTAGAGGTAAGTCCCCTAGATACATCAGTTGCTTCTCTAATAGCTTCTTCAAAAATCTCTTGAAATGCCAACTGAAAATTTTGAGGGTGGCTGATTACGAGGTCCATAACGCATTGTGGAATAACATCAAACTTTTGTATCTGCAACGCAAGGGCTGGCCATGCATTGGGCAAAGGCCCTCGTGGGCAGCAAACAAGACGCAGCAAGTGTGCAAGCAGAAAGTCCTTGCGAATTAAGTTCTCGGGTCTGTCTTGCAAACTAATACACATGTGTTCTCGTGATAATTGGCTGTCACTTGATGCCACTGAAGCTGAACTTTGTTCTGCTGATATCGGATTCTTGTGGTTGGCAAGGTAAAAGCCATTATAGACCTTCTCGAGGAGCTCAGAAATGGAAGAAAGCGGGATTGCTTTGCCCAAGCCAGCACCATCCTCTTTGTCATCAGAAACTGGAAGCTTATCACGTACTACTTTCAATAACTCTGGCTCAGCAACGCTTGAGACTAGATCATCAAGAAGCAGCTTTTGTGAAACTTTTTTAAAAGGCCGTGAATTGAGGTTCTTCGCAGGAAAAATATTTCTCATTGCAGCATTTCCAACACGCTTTTCCAAGCCATTCTGGTTTAACTGTTGCACCTCTCCTGTCCAAAGGCTTTCATCTCCAAGTTCATTGAAGAGATCTAAAAACCCATAGACAACTGGGCCTGTGATGTCCTGTGAAGTGCGTGCTGTACTGTCTAAAATTACTGCTTGAGACTTGTCTACTGCCTCCTGGTCATTCTCAAACTCCTGCAAAAGTCCACTTCCGATATGTTCAGATAAGAACTCTTGTGCAGCTTCAGCCAGATTGTAGATCATTACCCTACCTTCTCGTGAAATAGCACTTGCTTGATCAATAAGCAAGGAGTGCAATGAAAGTAGTTCGTCATGAGAAAGTCCCTCCATGCATTTGACTTGTAATTTGGGAGGCTTGTATGGATATCCTGGTAAACATCTGACATGGAGCTCTACGGAAACATTACTTCGACCTCCATCAGCATGAGGATGCAAAGTTATGGAGAATTCCGTACAAGAATCTTCAGAAATGAGAGTGAAATCTTCTTGGAAGATGGAAGACAGCGCTGTGAGCTCTTCAGAGAGAGACGAATCATTTTGAGGATGAGAGAGGGAGTGATCTTTGGCAGGCTGCCGATTCGCCCTGATGTTCTTCTTTTGGCTTCCTCGTTTTCGGCGGCCCCCTCCAGCCATACACTCACACACACACACACACA

**14-3-3-LIKE PROTEIN GF14 PSI (GRF3):**

AAAAAAAGGGGCAGGCAAAGAATAACCCTTGAAGCATTTTTAAAAAGGAGGCATTAGGTTATGATAAAGGCAGGCAACTCAGGACGAGTGACCAATGTTTTTTTGCGTTTTCCAGCTGTCTAACATATTGTGGTAGCCTATGTAAAGAGAGTTCTGCGAGCTGTATGTAGTATGTACCTACTAGCTTCTCTCTCGCATTGCAGGTCTGTGTAACCCTAATCGCAGCTATGGAGGAGCCCTCTCGCGAAGAGAGCGCGTACATGTCGAAGCTCGCTGAGCAGGCCGAAAGATATGAGGAGATGGTGGACTACATGGAGCGCGTCGCCAAGAGCGCCAGCGGCGAGGAGCTCAATGTCGAAGAGCGCAATCTTCTCTCTGTTGCTTACAAGAACGTGATCGGAGCTCGCAGAGCTTCATGGAGGATCGTCTCCTCCATTGAGCAGAAGGAGGAGAGCAAGGGCAACGATGATCATGTCACTCTCATCCGCGACTACCGGGCCAAGGTGGAGGCCGAGCTCTCCAACATCTGCAATAACATCCTCTCCCTCCTTGACGCCCACCTTATCCCCGCTGCTGCCTCTGCAGAATCCAAGGTCTTCTACTCCAAAATGAAAGGCGACTACCACCGCTACCTTGCTGAGTTCAAAACTGGCGCTGAGCGTAAAGAAGCTGCTGAGAACACCCTCATTGCCTACAAGTCTGCGCAGGATATTGCTAGTGCGGATCTTGCCCCTACCCATCCTATTCGCTTGGGCCTTGCTCTAAACTTTTCGGTTTTCTACTATGAAATTCTGAACTCCCCTGATCGAGCTTGTGCGATGGCGAAACAGGCCTTTGATGAAGCCATTGCTGAACTCGACACACTGGGAGAGGAGTCTTACAAGGACAGCACCTTAATTATGCAGCTTCTGAGGGACAATCTTACTCTATGGACGTCTGATATGCAGGAGCAGTCTGATGAACCATAGAGGGTAGTTTTTTTCTTTAGGGATTGACCATGCTGGTGGTCATAGCTTCAAATCAATTATGGCTTTGTTATGTCAGGAACAACAGACTATGTGACTCCCCTTCTGTTCGGATTCTGACCTGGTATTTGTTTCTGGCGCATGGGGTACTCTTCATTGTCCAAGAGTAGAGTCTTGGAAGCTTATCCTTGGCTGTATGTAATAAGATTTTTTAAGGTTGGTATCTAGTTAATTTTGCAAGTAGCTGAGGAACTGCTACTTGTTTGGAATGCTCGTCGCATTTTTATTGGGGAGCATCGGTTATCAAGAGCCGGCCTCGGGTTATGAGCCAGCTTTTGTGACATTGTGTCATTATTATTTAAGATTTATGTGGCTTGCTGTCTCTACCTTGTTGATCAAGTCATAATGTTCTGTTATTTTTTTCACATCAATCTCAGTACTGTTTTCAATAA

**KIN17-LIKE PROTEIN (KIN17):**

CTTTGTATATGATAATGTGTGTGTTCATCTTTTGAAGTGTCGTTCAGGAGGGTTCGACAGTCGGATATCGCGACATCACAGTTTTGCAGGTTGCCATAGCCGGCCGGCTTTGTTATTCTCAGGTTGCCATAGCCGGTTTGAGGGCTGAGAGAGAAGCGAAGGTCTGCAGGTATTCGGTTCTGCTATTTACTTGTGATTTGGCTTGGCGTTTTACTGGTTTGCAGGTATTGGAAATTAGGGTTTTAACAAGCCCCAGCGCCCACAATCTTTGGGCTTGATCATCAACGCCTCACCCACACACACCGGATTGTTGTCGCATTCATGGGGAAGAATGACTTCCTCACGCCCAAGGCCATTGCGAACCGCATCAAGGCCAAGGGTTTGCAGAAGCTTCGATGGTATTGCCAGATGTGTCAAAAGCAATGCCGCGATGAAAATGGCTTCAAATGCCATTGCATGAGTGAGAGCCATCAGCGCCAGATGCAGGTTTTTGGTGATAGCCCGCATCGCTTTGTCGACGGCTTTTCACAAGAATTTGAGACTTGCTTTCTGGAGCATGTGAAACGTAGTCATCGGTTTAGCCGAGTTGCCGCAACCGTTCTTTACAATGAGTACATCTCTGATCGCCATCACGTCCACATGAATTCTACAGAATGGGCCACACTCACTGACTTTGTCAAATACTTGGGAAGAACTGGTAAGTGCAAAGTAGATGAAACAGAGAAGGGATGGTTCGTTACCTACATTGATCGCGAGCCTGAGACGCTACTACGGGAGCGGCTCAAGAACAAGCGCGAAAGGGCTGAAGTTGCTGATGAAGAACGGCATGAAAAAGCCATTGAGCAGCAGATTGATAGAGCTGCAAAAAATGCACGGCCTATGGTTGCGCCTGAGGATTCTAATACTGAGGAGAAGAAAATTCTGCAGAGAAATGGGGAAGAGGGGAAAGTTGTATTTGCTATTATGCCTTCCCGTTTCAAAAATGCAAAAGAAGGTGAAGAAATCAATGTCAATGGTGGTGCAAGGGCTGGTCAAGGAGTAAGCTCTGTGTTTCTGGAAGAAGGAGAACTGAGGGGCTCTAAGGAAAGCTCTAAACAAAATGGAGGCACACGAGATGGTGGTCGTGACATGGCTGCTACCAGCAACAATGCATCTGCGCGTGGGGAAAGTGGTAAAAGTGGTTCTGCAGCTCTTGCTCAATTAATGGAAGAGGATGATAAAGCAAAGGAGAAGGCAAATCGGAAAGATTATTGGCTTGTTGAGGGCTTGGTTGTGAAGGTCATGAGCAAGAGCCTTGCAGAGCAGGGTTTTTACAAGCAGAAAGGTGTTGTGCGCAAGGTGATTGGCAAATATGTAGCGGAAATCAAGATGCTCGAAAGTGGTGCCATCCTTAAGGTGGATCAAGAGGAGCTGGAAACTGTCATACCTCAGATCGGGGGCCTTGTGAAGATAGTGAATGGAGCATACAGAGGGTCTAATGCAAAGCTTCTTTCTATTGATACTGCGAAGTTTATTGCAAAAGTCCAGGTAGAGAAGGGCATTTATGATGGAAGGGTATTGAATGCCGTGGATTACGAAGATATTTGTAAAGTCTCTATGTCTAGCTAGGTCTATGCCAAAGGAAGTTCTTGTTTAACCTCTTTTAATCAACTGTAGTGGTTCTCATCATGCTCTTAGTGGGCTATAGCTTTTTGCGATCTTTGAATGTTGGAGATGGTAATGGACATGTTGCAATTTAAGTCCCTTTACTCAATGAAAGCATTGTGAACGAGTCCCTGATTGGGCTCCCCATGCACCCTTGCACTTCCTTGTTCCTTTCGAATCATTGAGCAGGTGTGTTATTTCGTCCCCTGTATCTCCTGTCTACTTTACAGCAAGGATATCTGAGTGAGCAAGGCTTTTTTTTTCAAAGCAAATTGAAATCCTGATAGAGCAAATCAGGGATGTTGGTAATGCTGCCTATGTACACAGACACAGTTGGCGATTTGCCAATCACTGATCAAAACTTGTGTAGTTCAATCTGGCCACAATAGTTAATGATACGAATTTTTAAAGTTCTGGCAGTGCAGGTTTGTATAAGTTCTTGGAAGGTTACACAGAGAAGGAGATATGTCAATTAAAAATGTTTCGTTTGTCCGTAAGACTTGGGCCTCATAACTGCTATTAGGTTGTGGTTCCAAGTCTCCCTAAGCTCAATGGGAAGGAACATCTATGGGTCTTGCCCTTGACGGATGCTCCGCCCTCAATCAAGCGTAGGCACTCTCCCTCTTCCTGGCGAGGTTTCATGTGTTGCAGACTCTGCAAGCTCATACGCCTGTACCACATCCGGGACTTCAAAAAAGGGGAGACATTGCTGTATTCTGTCATTGTAAAGAGATTTCACTCTGACAGGACCCTTTTCCCATCATTGTTCGTGATCTTAGTGATCTTGGGCTGGATAGTCGTAGTGGTCGATCTCTGCTGCTACTGGAGTAATCTCTCTCTTTGTCGTCATATCTCCAAATTGTACTTTGTCAAAGATCAAAAGATCTGGTTGCTGGATCCATGGTTGATGATGGTTTTGTTGTTAGCATTCTGAATGGATGTGATTCAAAAACAATGCAATGCATGGTCGCCTTCAAATGTGTTTCTTCATGAGCTCCCTCTTTCCATGTGACACTAGTATTCTTGCTGGCATATGGCGTCGGAATTCGCTCTCATTTTCTGAAATTGCTCTTCTCAACATGTTAAAGGGGTTTTTATTTTCTTTCGGTTCGACAAGTTTGGCTGGACTCTCAAGATGTTCCAGGCTGTCTCCG

**PHOSPHATIDIC ACID PHOSPHOHYDROLASE 1 (PAH1):**

CTGACATACGTGGCGAAAATGGTGGATGGGTTTGATGAGTCCACCCGTCGTGCTGACGAGGAGCGAAGAGAAAAGGGTTTTATGATTTTCCGCTTAATCAGAGATTCAGAGTCGTCTCGCTCTACATCGCTCCCTCGCTATCGCCATCGCTGCACAACTTTTCGCACAGGTAATTTTATAAACTTATCATCCATTGGGTTTTTGTTCTGTTAAGCCCCTATTGGCGCCCCCTCTTGTGCTTTTCTACCCATACGGGCCTCTTGATGGGCGGTTCCGCTTCATTATGGAAGGCGTTCTTCTCCTTTAATTTACTCTCCAAACCCTTTGCCATGCTCGGGCATCCCATCATTTGCATGTTTCTTGATATGGGTTACTCAAAGCTGCCTGCACATTAGGCGGAGTGCTAGCAACGTCACAAGGCTTACAAAAAGGAGAGATGGCGCTTAGCTACCGGGCGGGGCTTTGATCTTTTTCGCTACAAATCGGTTGATTTTTCTTGAGTTCCTAACCAATATTCATGGTAACCAGCCATTTGCAAACCTGAGCATCTGCTATGATCATTCGTCATTGCAATTGCTGAATGTACAGGAGGGCTTGATGTGGTGCCAACATGTATGCATTTGGGATGCTGGGCAGCTATTTCTCACAGGGTGTCTATGCTGTGGCTGGCCCTTTCCAAGGCGCTGTTGACATTATTGTGGTGCAGCAAGAGGATGGTTCCTTCAAAAGTTCACCTTGGTATGTGAAGTGTGGCGATTTTCAAGGTGTGTTAAAGAGAAGGGAGAAGATAGTAGACATCTCAGTCAATGGCATTCCAGCCAATTTTTACATGCTTCTTAATAGCAGAGGTGAAGCCTATTTTGAAAAGGAAGTGGAGGATGCAGTGCTGGCCTTGCCTAGCACCTCATCATCAGCAGAGGGTTTTGAGGAAAATGAAGGCCTTGGAAGTTCAAAGGCCTTGGCGGGTTCTCAAGTGCAGAGTATAAGCACTGCAGATACAAATGGGTCTACTCAAGGCAATAGCATTGATCAGATAAAAGATCAACAGGATGACCTAGTCTCGAAAACTGCTTCAGAGGTCCGGGAACCACAAATCACAGTAGATCAATCGTCTGATTCTTCAGTAGATCAATCATCAAATAATTTGGAACATCAATCCTCAGGTTTTCAGGAGGCCAAAGATGTAGAGTATGCTTGGAGAAATGCAGCTAGTAGTCGAGACACGAGCAGCTTGAATGCAGCTGATCAGTCAGACTCGCTCTCCTTAACGGCAGAAAACACATATGAGCTGCGACAACCGCTCGATTCTGGCTTAGAAGTGAGATGTATGGTCATAAATGTCAGTGGTGAGAACCTGGTTCTAGCAGAGGCTGACAGTATTTTGCCAAAACAGATGGCTGATGAAGAAAGGCTGCAATTGCTGCCAGTGCGAACTGGTGAGGATGGATTCTTATCAGAGGTGGATGATGAAGCTCTCAATACCGACACAAGCTCATTAGGGACAGATGGTGCTGTATTTTGTAAGTTGAATACTGTGATAACAGATGATTGGTTTGGATTGCAGAAGGCATTTAGTTTTGAATCAGGTGATGAAGCCACTAGCATACGGAAGGATTTTGAAACTTCCGATGGTAAAGTCATGGATGAACTTATATGGAATGACTTCAAGATTGTAACAGCTGAAGAAGAAAAAGAGCTACATGATGCACTTAGAACTACCCCATTACCAGGTGTAAACATTGACGGTAGTAGTCTGACACCCGACACGGCAGATGAAAAGAACTGTGAATTGGTTGGTTCATCTGTTTGGGTATCAGATGCACTGGTGGATGACGCTGCTAGTAATGTAGCAGTTGCTAGTACGGAGGGTAGCATTCCAATGCAAGAAAACAACGGGGATGCGAGCAAGGATGGTGTCTTATCAGACAAAAGTGTGGTGTGCAAGGTGGAAGAGAGCCCTCATACTGAAGGTAACAAAGCTGCAGAAATTTCAACAATTGCTGTAGTCCCTCCTCCGTCTAGGTGGTGGCCCCTTTCGCTCAGGAGGTCCAGAGCTGAGAAGGCGCATGTGTCCGAAGCTGCAGATACAAAGCACGCCTTGTTGGCAGCCACCAATATAGCTGTTAAAAGTCCTTTAACAGCAGAAATTCTTAGAAAAAATGGCTACTATGAGTGGACGCAAAAACCAAAGCTACGCAGTTATGTTCCAACATCAGCACAGATTGAATCTCTAAAACTGAAGGATGGGCCAAACAAAGTAACTTTTACATTTCCCAGGGTTCTTGGTAGAACGCAGGTAGATGCTAATATCTATTTATGGAAGTGGAAGACCAGGATTGTGATATCTGACGTCGACGGAACTATCACCAAGTCAGATGTTCTTGGCCAATTTATGCCTCTTGTTGGGAGAGATTGGAGCCAATCTGGAGTGACTCGTCTCTTTTCTGCAATTAAGGAAAATGGTTATGAGGTTTTGTTTTTGAGTGCTCGTGCCATTTCTCAAGCATACCTTACACGACAATTTCTTGTTAATCTGAAACAGGATGGGGAAACACTTCCAGAAGGGCCGGTTTTCATTTCCCCTGATGGTTTAATTCCCTCATTGTACCGAGAAGTTGTGAGAAGAGCGCCTCATGAATTCAAGATATCTTGTTTGGAGGGTGTCAGGAGGCTTTTTCCAAATGATGTCAATCCATTTTACGCGGGTTTTGGCAATCGTGATACAGATGAAATTAGCTACCTGAAAGTGGGAATACCGAAGGGAAAGATTTTCATAATCAATCCGAAGGGAGAGGTAATGGTGAATCATCATGTGGATGTCAAGTCATATACAAGCATTCATACATTGGTGGACGACATGTTTCCGCCAGTGTATTCATCTGAACAGGAGGATTTCAATACCTGGAATTATTGGAAGCTTCCCCTTCCTGATTTTGTCGATGAGCTGAGTGCAACAAGCACGAAGCCCAAAAAGGTGGAGCTTCCAAGAAGTAATTTTTTGGCATGCAATCTCCTTCGCAGAGATGAAAGGTAGACAGTATTTTTACCTGTGGGTTGGACAGGTGCAGGTGGTCACTGTGTGATTCCTAGTCTAATTATTCAAATGGTCATTTTTTGCCGCCCCCAAGGGGTTGGCTCATTAATGTAAAGAATCTAGGGATGCTTGTACATTATGTGAATATTGTGGGAGGTTACTGGGCTGCTTTTGACAGGGAGTGTGTTGAGAAAAGAGAAACCTGTAGGAGCCTTGTGTAGCATCACACATGACAGCCAACACACTTATGAGATGTAATGGACATTCTTGTAAAATAGTTGAACGTTGACAACATTTAGAGACATTTGTGAGTTTATTATCACGTATACAATAGAAGGCTTTCTGTATGTATAATTTGACCAAGGAAGAAGAAATGGGTTGGTTCTGCCAGGATTTTTACTGGAATCGTACATCGATG

**β-KETOACYL-ACP SYNTHETASE 2 (KAS2)**:

TTAAATAATTATTGGCCTACCCATCTCTCTCGCTCTCTCTCTCTCTCACATGCCCTCCACGCCTTTGCTGCCCCTATCGGGTGGCCTTTGTTTATACATTTCTCCACTCTCCATGATTTCTGCGTGACCCCGATTCAACTTGCAGCGGGAGCTCCTTTGTGTGCTGCCAGAGATCTCTTTACCCTTCTTTCTCTCTGCCAATTAACGACACAATACCCAGTTTAGCAATTCTCCCTTGAAGCAATTCTTCACAGAGAAGTCATGGCTATGGCAATGGTAGCTCCCCTTTACACTTGGCTGGTAGCAGCAGCTTGTTCATATGGAGCTGTAGGGGGAGAGGAGAGAGGCATGCGCGCATCTCCATTGGGAAGGCGTGCGCGGCGTAGGCTGCAGAAATTGGGCGGGGCAAGTTCGGCATTGGTCTGCAGCAACGAGAGCTACAGCAGCATGTTACCTGGAGGAAACCTTGTCGCATCTTTCTGTGGATCCAGCCTTCAAGGCCTTGGCTCTTGCCTCTCTTTCGAAGCCTGCCCCGAGTACTACAGCTCCAAGCAATCCCCCCTCTTTGGTGATGGCGTTTTCGGTCTTGGATTTGGATCGAGGCCTATCAGAGCTGCCCGGCGGCATCCAAAAGCAGCAGGCGCAGGCAAAGCTTGGGCGATCGCAATCCAGCCCGTGAGGGAGAGAACTGAGAAGAAAAAGCTCACCACCAAGCCCAGGAGGGTTGTGGTGACAGGCATGGGTGTGGTGTCGTGTCTAGGGCATGACCCAGATACTTTCTACAACAATCTTTTGGCCGGGAAAAGCGGCATCAGCGAAATAGAAAGCTTTGATTGCAAAGATTTTCCAACTAGGATAGCTGGGGAGATAAAATCCTTTTCAACAGATGGATGGGTTGCTCCAAAGTTTTCTAGGCGCATGGACAAGTTTATGCTTTATCTTCTTACGGCTGGTAAAAAAGCTTTAGAAGATGGAGGGATTTCACCGGAAAGCATGAATGAACTTGATAAGGCTAAGTGTGGCATATTGATTGGTTCTGCTCTTGGTGGAATGAAGGTTTTCAATGATGCAATAGAGGCTTTAAGAGTGTCTTACCGCAAAATGAACCCTTTCTGTGTCCCCTTTGCAACAACAAATATGGGGTCTGCCATGCTTGCAATGGACTTGGGTTGGATGGGACCAAACTATTCTATATCGACGGCTTGTGCGACAAGCAATTTCTGTATTTTGAGTGCAGCAAACCACATTCTGCGGGGGGAAGCTGATGTCATGCTGTCAGGTGGTTCAGATTCAGCAATCATTCCAATTGGGCTGGGAGGTTTTGTTGCTTGCAGGGCCTTGTCAGTGAGAAACGATTCTCCTCAAAAAGCTTCGAGGCCTTGGGATATTGATCGAGATGGATTTGTCATGGGTGAAGGTGCTGGAGTTCTGCTTTTGGAAGAGTTGGAACATGCCAAGAAAAGAGGAGCGACTATACATGCGGAGTTTCTTGGTGGAAGCTTCACATGTGATGCCTATCATATGACAGAACCTCATCCTGAAGGCACTGGGGTCCGACTTTGTATAGAGAAGGCACTGGCACAGGCTGGAGTTCGGAAAGAGGATGTGAATTATGTAAATGCCCATGCAACTTCGACGCGAGCGGGTGATCTCCAGGAGTATAAGGCGCTGATGAGAGTTTTTGGTGAAAACCCAGAGTTGAAGGTAAACTCCACAAAGTCGATGATTGGCCATTTACTGGGAGCAGCAGGAGCAGTTGAGGCTGTTTCAACTGTACAGGCGATCAGAACTGGGTGGGTGCATCCAACTATAAATGTTGAGAATCCAGAGAAAGACATAGACATGAGCATTATTGTGGCAGATAAGAAGGAACGCTTGGACATAAAAGTAGCACTATCCAACTCATTTGGTTTTGGAGGCCATAATTCCTCTATATTGTTTGCACCATACGTTGAGTCTAATGAGAACTAGCTGAAATACCTACTTTCCCTATCTTTACAGCTTTCATGTTTGAGAGCCTTATCTTGTATGAGATAAATGCGTTTCACCAACAATGTGCAATATATTGAAAGGCGCACATAACTCCTTCAATGAATCTGGTGTTTCTTGTGTACAGTTCTCTCTTGCCATTGTCCTTTCCTTTGATTTTTGCACTTGTTAAGAAGCAAGCAATTGATAAAAGTGTAAAAGCAACCAAATGCGCAAATGATTATCACCATTGAATAAAAACACAGGCTTCTTTCGAAAAAA

**PALMITOYL-MONOGALACTOSYLDIACYLGLYCEROL Δ-7 DESATURASE (ADS3):**

CTGGGGAGCACTTGGCCAACCCGTGCTCGCCATAGCTTTCTTTGCACAGACGAGCTATGGCTGCCGCAGCAGCTGTCTCTATGAAATCTGTAGCCATGGCGCGTCCAGGTCTCCTCTACACCAGGCAAGGTCTCGCTTCTTCGTTTTTGCCTCCCCTTCGCTTCTGCAGCAGCAGAGCAAGCAGCAGGTCTTTCTCCAGCAGTGCCCAATTCTCTTCTTGCCGACCGCTATACATATCAGCCTGCGCCGCGCAAACCGTGACGCCTCCTGAACCCGGCAAGTCTCCTGCACATATTTGGCTCTCCGATGTTGCACCCCTCTGGAAGAAACCCTGTTTTAAGAACAGAAAATGGTCCGGAATAGATGTGGGCATTGCTGCTTACATGGTTGTTATCCATGCACTATGCTTATTGGCGCCCTTCACCTTCACATGGCAGGCATTTGGGGTGTTTGTGCTCACCTACCTGATTACTGGCATGCTTGGCATTACCTTATCGTATCACCGCAACCTCTCCCACAAAAGCTTTACGCTGCCTAAGTGGCTTGAATACACATTTGCGTATTGCGGTGCCCTCGCTATTCAGGGGGATCCTCTTGAATGGGTGAGCTCTCACCGCTATCATCACCAGCACTGTGACACCGATATGGATCCCCACTCTCCATATGAAGGATTTTGGTACAGCCACATGGGATGGCTGCTTGATAACAAAACGACTTTGGAAAGGGTGGGTGCCCGTGGGAATGTTGGAGATTTGGCGAAAGATCCCTTTTATAAATTTCTCCAGAAAACATATCCTCTTCATCCCATCCTTTCAGCAGTTGGTCTGTATGCAATGGGAGGTGTTCCATTCCTGATATGGGGTGTGGCTGTTCGGGCAGTTTGGGTGTACCACATTACTTGGCTTGTGAATTCAGCATCCCATGTGTGGGGCTCTCAAGCATGGAAGACTGGAGATTTGTCCAGAAACAACTGGTGGGTTGCAATGCTAGCATTTGGGGAGGGATGGCACAACAACCACCATGCATTTGAATACTCTGCACGGCATGGCCTTGAATGGTGGCAATTAGACCCCACATGGTATTGCATTCTTGCACTTCAGAAATTGGGACTTGCAACAAAGGTGAAGCTACCGACTCGAGAGGCCATGAACCGCCTGGCTCTAGAAACTTCTTAGTTTGTCGGAAAGATTTGATATTTCTAATTAAAGAGCTCAGCATCTACCCTTGTCAAACAAATTTTTTTGGGTAAGGCTTTCGTATGCTCTGTGTGAGTGTGTACATACCAAAGACCTAGTCCAATTGAACTTTAAGAGGAAGAATTCTATGAACAGAAGAGGATCCGATCATCAGAAAAGCTTCTCCGCATTATTCGAGACTTCTGCTGTCAGAGTGTTGTATTTTTTGTAGAAGTGTCATTACCTCAATGCATTATATGGCTTGAGAGAAGTGAAGGAAGCTAGCTGTAAAGGCATGAGACCAATATGTTGTTAATTTCGATATCTACTTTTCGATTCAATTGCAAAGTCCGACTTGTTTCATATGTAGTCAGTATGGGTTGATGAGGAGCAACACCATCTGTTTTGAGTTGCTTTCAATGTGTCAAATTTCTATTTAGCCAACTTAAAATTAAAATATCCATCTTAAAAC

**PALEFACE 1 (PFC1):**

TTGTAGCTATGCCCAAAGAGGCGGATTACATGACTCCACCTTTGATATATAAATCAAACGAAAAAATTATGCTCCCTGTATTCTCTCCTTGTCTAAGGCCACTATGAAGAGTGGTTGGTTATGGTGTGCAAAAATTTAGATATGCACACAAGTGCTCGTTTAACCTTCTTCCCAATAGTCAGGGTATGGGGCAGATTTTTGAGCCAGTTAAAGCCAATAGCTTATTGTTTTCATTCACAAACTTAAGCCCAAATAAATCCCACATGGAAATGCACATGTGTGAAGCCTGGGTGCTTAAATCATCTCTTCTATACAGCATTCCCTCTTGTCCACATGTAGAGCATTGAACAAAGCGACGAACTGATCGAATGTAAGCTCATCTGGCCTTGCAGTTTCTAGAAGATGCACCGACTTTAATGCATCTCGAACCTCTCCCGAAGGATATTGGTGCTGAAGCGAGTTACGTAACATTTTCCTTTTCCCACCAAAAGCTGAGTTTACTAGTGTAAAGAAAGATCTTGTAGATGAAACTGGGGGGTATTTTAGTGCCTGCTTAATCTCAAATGAGACAATAGCTGCATCAACATTGGGACTAGGGAAGAAGTTATTGCGGCTGACTTTCAGTTTGTACTGAGCATCTGAGAAAAACTGTACAAATAAGCTGATAGGTCGATAGTTGTTAGCCTGTGGAGACGTATCTATCAGACGAGCTGCTGCTTCATCCTGTAACAAGAGAACCATGGAAGAAAAAGTTTCTCCCATTGGAAGAATACGTTTGACCACTGGTGTAGTTATACTAAATGGTAGGTTGGAAACCACCTTTGCATGCTTAAGAGGCATGCCCCTTTCTGAAAATGATTTTTTCAATTGATCAATGGAAGTTTTGATAGGCCACTTCATAAAATCTTCCTGTACCACCTCAACATGGCCATTCCCTTGAAACCTCTCCAGGATTAATTCCGCCATCGCCGGATCTTTCTCCACCGCAATAACATACGCGCCTGCTTCCACTAAATAATTGGTGAGAGCACCTGTGCCCGGCCCAATTTCCAAAACTAAGTCATCTTTTTCAACATTTGCAAACTCCACAAGCTTCTCATTGATTTCTTGGTTCACCATATAATGCTGGCCAAGCCATTTCTTGGGCACTCGGTTTTTGGACTGGAGAGCCTGCAAGGTGCCATGAAAGTCGTCCTTGTTCTTGTGCTTCTTCGTAGGAGGAGGAGGAGATGCGTGCCGGTCAGCGGAAGCCCGGCAAAGAGAGAGCTTGTGCAGAAGAAGAGATTGAGGCCGCAAGCTCAGCTCTTCTAAAGAGCCCAACCCACTGCAAGCTCTCGACATAGCCACTGGAAAAAACTGATTTTTGTTTTCCCCCAAGGATAAGATAGGAGCTAGTAGGTGCTAAGAGCCCCGTCCCGCCCCGCGCCGCGTAGTGCTA

**β-AMYLASE 3 (BAM3)**:

GTTTTTTTTTTTTTTTTTTTTTTGCAAACAAATAAAATATTCTAGTTTGAGAAATTCAAACAAGTGAGTCCTTGCAACAAGCAGTATGATGGAAGGCACATTACGATGCTAGTGCCTCTGCAGAACACGAGTGCTACAAAAAATTACAAGGATGTTCAAATCCTGAATGATATACATAGCTAGTCAGCACCTGAATGATATACATAACTACTCGGCAATAGAAAAACACATTTACTACCTTGGCATCAAAGGGTTATGATATCCGATTAAGAATCCACTAGTAATCAAATTCACTTTAAAAATTCTTACCAGTATATTATACATATATATATACACTGAAAGGCAGGCAGCTTCCAATCAAGATGCTGCTTGAACATGCTATTTGCACATCATTGCAGCTTCTGCTTCCTGGATCAAAGGTCTTGTGGCACGCACGTGAGTTTCTGTTGCTTGGTGTTTCTCCTCCCAAGGATTAAATGTTCGCCCTTCGGCCATCTGCTGAACGAACTGGACAAAACTGTGCCAATTATCACTACGAAAGAGCTGCTCACTCATCCTTAAAAATGTGAAGGAGCACATTGGCTCATATGCAGCTTCCCATCCATTACTCTGATCTGCCAACCGGGAATTCTTAATAATTTGGTTGTGAGCCCCTTCGTCAAATCTCATGAGTGCATTCTCACCAGCTAGTGGCACACTCGCTTTTCTTGCAGCAAGCGTGACTTGCTTCAAAAGGCCTTCAGGGCTACAAAGGGCACATGCCGGATACTGCTCCTCGTCCTTCATCTCAAAGCAAGTAAAATTCAGTGTCACTCCATGCCTGCCAAACATTTGTGCGACAGGAAGGTAGCCATCCCTTGTTCTGGTATTGTAGTAGCCTGCAGTAAGTTCTGCTGAATGAGAACGAGTGCCATAATGCCAATGAATACCTGCTACTTTGCCAGAGAGTTTAGAGCCCGTGCCCCTAAAAATGCCCTCTGCTGCTGTCAGTATCCTCTCACCATGATTCAAAAGCATTCCCGAGTACCATGCAAGAAAAAACTTTCCATATGGTGTATTCCATGCTCCATCGTGATTGAAGAAACCTGTTTCATCTGGAAACTGCTTGTAATGCCCTGCATCATGTGGGCCAGCTTTACCCCACTCCACTTCTCCCACTTTTTCAGCGCATGCTTGCAGATTTGCCAACATGTACTTGTCATAGCATTGGAATTCTCCAATACCAGGAAACTGCCAGGTATTATTGCTTTCAGGATATGCTGGGTACCTAAGCTCCCCGGCAGGTCCTAACCCAACTTGAATTTCCACAATAACATCTCCAAGATCACCTTCAAATTCATCTCGAAAGCTCCTCATGAAGTCTGCATACACCTGGACAGGAGACCTTCCCTTGAGCACCGGAAGCGTATCACAACCCAGAGACAAATACTCATCGTTTCTTCTGCCATCCTTGTCCGTATAAACAAGGTCTGGGTTGCTGTCGACCTCCTCCTTTACCCATGGTGGTAAAGGGATCCAGCAGCTATCTCCAACATTCCCGCCGCATTGATGAAAGGACATGACCGCTTGCATCTTGAGACCATACTCTTTGGCCATATCCAAAAGCTCTCTATATCCAGACCAATTGTAAAGCCCGGGTCCTTCTTTTTCCACAATACCCCACCATACATCCACCATGATACCTTCTACTCCAGCGCTTTTCAATGCCCTCATACTCGCATTCATAGCCCGGGGCCTGTTCAATGTGCTGTTAGGGTTTACAGAGTCCAAAGGGAGCATCACATACACAGGAACTCCCCCTTCTGTGCCTCTGCTTGTTGGTGTATCTATATAAACATACTCCTTCTCTGCAGGAGCTTGTGTTGTACTTTCATGGAAATCAAAAAGCTTGTCACCTTGAGAATGCGTGGGTGAGGAACTTGGCCATATGCTGGCTGAGGGTTGTGAAGTCGACCTGTCAGAATTCTTTCTCCACCATTCATCAGAAGAGATGAAAGATGGGGGAGAGGAGCCGAAAGGAGCTGCTGTATCAGGATTGCAGTTGAAATTATGGTTGTGGGCAGGCAAACTGTGGAAGCCATGGCTGCTCTTGCGATGCCCCAGGGCTTGCTTTTCATGAGCATCAGGCCAGAACCCCCTATGGGATTCGGGAGAGCTTCCATAGAGAGAGCCGGAGATTGTAGATGAAGGAAATGCCATAGCTCACAGGCTAAGCTTCTAGCAGAAAGAGAATATGAGGGTTTCTGTACAATCAAGTAACCTCTTGAAGAAACAATTTGTACAACAGAGAAGCAACTTATAATCTTAAGCAGAATAGAACTCTTCACACAGAATTTAGGGTTAGGGTTAAAGCGCAGCTAAGGTTCTTAACATAGGAAAGCTACCTTACACAGAGTTAAAGCGCAATGATATTTCCGCGAGAGGAAAAGGAAGACGATGAACGAGCCCGCTGGCTATGGCGATGGAGAACTCGGAATAAGGATATACTTACACATCATCTACGAAATGG

**FUMARASE 2 (FUM2)**:

AGAGCTGGAAAGGTGGGCTTTGATGAACTGCGATTAGAACGGTAGAAGTAAGAATAAAAGACAAAACAACCCCTTACTTATTAAATTCCAAGGCATCACTCTTACGGATCAACAATTACCAGCCATCCTCACATGCTGTGAAATCAATTGTCATGACTGAGATTATTGAAATTGACTGCCCGTAAGCACTTGAGCACAAGACAAGCTTGCCTTCTCCTCAATAGCTTTCCATTATTGAGAAAAGGAATTATCCAAAACTGTAACTTCCTTGATTATAAGCCTGAAGGTGAAATTTACAATTACGTGCCACAGATACAATTCATACAACATCTCTTTAATTGCACCTTCTGAGCCGGGAGAACGTGACTCCTTATTACATATAATGCTCCTTTACTACCAAGAATGTAACCACCAAGAACTGAAGCTACCTCTATGAACCTCCTTGAAGGGTTTCTTATGGATAAATATGGGAACCTACTTGGTGTACCACTTAAGGTAGCAAGAAGGGGCTGTAGCAAATTTAACCCCCCCCCACCAAAAAAAAAAAAGAAGAGCTAAGAGGTACCGTCCATGCGAAGAATGGGGATTAGAACTAATAGCTAGGCCTCCTTTACTTTATGATGGTCCTAGCATTTTCTCTGGAACCACATAGTTATCGAACTCTTCCTCTGTGAGCACTCCCAATTTTAATGCTGCCTCTTTCAGTGTTGTCCCTTCCTTATGCGCCGTCTTTGCTACCAAGGCTGCCTTGTCATAGCCAATTATCGGGTTCAATGCTGTCACAAGCATCAAGGACTCACGTAATATTTGTGAAATCTTGCGCCGATTTGGTTCTATGCCAACAACACAATTTTTCCTGAATGATTCAGCTGCATCTCCAAGCAAGCGCAATGACTGAAGCACTCCATTCGCAATTAAAGGTTTGAACACATTCAACTCAAAGTGACCACTTGCACCTCCCACTGTTATTGCCACATGATTTCCAATGACCTGTGCGCAAACCATGGTAAGTGCTTCACATTGAGTTGGGTTCACCTTGCCAGGCATGATGCTGCTACCCGGCTCATTTTCTGGAAGAAGCAGCTCTCCTAATCCACAGCGAGGGCCACTTCCCAGAAAGCGTACATCATTTGCTATCTTATTTAATGACACGGCTACTGTATTCAAAGCACCACTGACCTCAACGATGGCATCATGCGCTGCCAAAGCTTCAAATTTGTTTTCAGCCGTCACAAATGGTAAGCCAGTCTCAACTGCTACAGCCTCTGCTACCTTTTCATCAAAACCTCTTTTTGTATTAAGCCCTGTACCAACTGCTGTTCCACCTTGTGCAAGCTGATAAAGTCGTGGTAGCGTACTTGTTATACGCTCAATCCCATACTTTACCTGTGTAACATAGCCACTGAATTCTTGGCCAAGAGTTAAAGGGACAGCATCTTGTAGATGTGTGCGACCAATTTTCACAATGTCCTTGAATTCCTCCATCTTTCTGGCCAAAGCCTCATGCAAGCCCTCCAATTTGGGGAGGAGCCTTCTGTTTATCTCTATTGCAGCTGCAATATGCATAACCGTTGGAAAAGTGTCATTTGAAGACTGGCTCATATTCACATGATCATTAGGATGAACAGCTTTATCCCCCACTCGTTTGCCAAGAATTTCAGCAGCACGATTTGCAATAACCTCATTTGCATTCATGTTTGACTGAGTACCACTTCCTGTCTGCCAAACGACCAAAGGAAAATGGGCATTAAGCTTGCCATCAGCGACTTCTTGAGCAGCCTCCATTATGGCAGTTGCTATCTTAGAATCGAGACCATAGTTTTCATTGACCTTGGCTGCTGATTTCTTCAGGATGCCAAAAGCCTGGACTATTTGATCTGGCATGCGTTCTCTGGCCCCTCCAATTTCAAAATTCTGCAACGAGCGTTGTGTTTGCGCTCCCCAGTACTTGTCTGCGGGGACTTTGATGGGACCAAAAGTGTCCTTCTCTTCTCTGAATTCAAGGAGGAGGGAGATGGATCTGGAGGGAAGGGGGTTACGTAGAAGGGAGAAGAGGGTAGAGGGGCGGAGGAGCGAGAGGCGGGGCTTGACGAGGCGAGAGGTGGCCATGTCGTTCGTTCTCCTGATGAACAAGAAGCACAAGGGAATTTCGCCTGACACAAGCACGTGCTATCATGCAGCAAAAAATGGATATGTTTTTTGAATGTTCTACTGGGCACAGCAGGTGGGAGCTGTTTGTTGACCTTAAAGTTGCCTGTAAGAGGAAGAGGTTTCTGGGCTTGAGCGCTAGAATCTGAAACCGATAAGTGAATCTGCCTTACAAAAAATCTTGATTCAGTAGAGAAAGACTGGAGCTGCAGCTGTGGAAAGACCCAACAAGAGCGTGCCCAAGAATCGAACTCTCGTCTACAAATCCGGTGAACGGAAAGAAAAGAGTTGTTTCCCGGGAGACCGAGGAAGGGGACTGGGGAGGGGCGATGGAGTTCTTGGTGTGCAAGCAGAGGGTGGCGCAATGCCCTAGAAGGCGAATTAGGTGTAACGTTGTTGGTGTCGGAGAGCAAGCGAATGGTGATTCTCCACATGCCTCCTTGCGCACTGATTGAAGCCTGGTCCTCGCTCTCTCAGCAATTTGGCGATGGCGCTTGCATAGAACATCGAAGCTCACACACACACACTCCCACAGCTGCTGTGGCGGTGGGACGAGCCACCAAAGAGAAAACCCAAAACCTCCTCCCGCTGCTTAAG

**3-HYDROXYISOBUTYRYL-COA HYDROLASE 1 (CHY1):**

TTTTTTTTGAATCATAGAAGCAGGCCTTTTAGATCGAAAACCATTACATAAAGTTCTAACTGAAGATAATAAGCAGACCTCGCTGATTTGCACTAATTTTTTGTAATAGACTTTCATTGTACATCATTTACAAGATAAGGAGCAAACTCATCGGGCACAATGGGTTCATTCCTGGTGTAAGACAATGGGAATGGCAGGATAATCTTGATTAAGCACAGAGAAAGTACTGAGAAGGTCGTGCCAAGACATTTGTTGTTGAGGGCCTTGACGGTGCCCCTATTAAGCTTTTGAGAAACCATAAAAAGCCAGCTTATCAAAGCCTTGGATGAACCATCCTTATGGCACTTAGTTGCCGATGAGCACCCGGAAGTTGCAATTCCTCATTCGATTTCTCGAACGGGGAAAAATAATGATCTACCATGGCTGGTGTGATTGATGCAAGAGTTGCAGGGTTCCACTGCAGAAAATTACAAGAGATAAGAACGGAACCTTTGACCCCACAACATTATGAGAGGTTCCTTCAAGCCACTAAAAAAGAAACAAAGCGCCCATGAATTGGCATAAAGAACAAATATCCTACACATTCAAAATATGAACTATATTGCAAGCTCAACCAAGCTGGCCGCAACTCTTGCATGATAGGCATACCTTGGGGGTCCTGTCTTTGTCAATCAGGATTGCACGGCAACCCTCGTAGAAGTCATTTGACACCTTTCCCAGAAGGCATCTTACAGTCAACCTGTACTCGCGCATGAGGCACTCATAGAAAGTCTGGTTTCTGCCATCACGTATAGATCGCAAAGTTATTTTCAAGCTCATTGGTGATGCCTTCCTTAATGTTTCGGCAGTTGTTGAATACCAATCATCTTTAGTTTCTTCCTGCTGCAGGTCAAGAGCTTCAAGAATATCTTCCACGGTGGTTTTAGAAAAGCAATAATTTACGTCTTCTAATCTATGAAGTATACTTTCTTCCTTGAATTGTGGCTTCTCAGAAAAGCTGCTAATGACTTCATCAACAACAGCCGGATTCCCCGAACCCAAAGATTTTTTTAATTTCTCCTCCACGTAAATGAGCCTCTCTGATGGTACATAATGGGTTGCAAGGCCAACAGCAAGAAGTTCAGTCCCATCTAGTCGAGCACCAGTAAGACCAACATATTCACCCAAAAATCCAGGTAGGCGTGATAGAAAGTAAGAAGCACCAACATCAGGGTGAAGACCCAGACCCGTCTCTGGCATAGCAAAAACTGATTTCTCTGTGACTACTTTAAAGCTTCCATTAAGGGAGAGACCAGCACCACCTCCCATCACAATGCCATCAATCAGAGCAACGTGAGGCTTTTTATAGGTTCCAATCAAATAGTCCACATTATATTCTTGTGAAAAGAAATGCATAGATCTCTCATGCAAACCTTGCTTTCCAAATCGGTACACTTCTGCAACATCGCCTCCAGCACTAAATGCCCTACCAGAACCTTTTACAAGCACAACTTGAACATCTGTATCTTCTTCCCACTGCTCATAGATTGTTTTCAAATGCCCAGTCATCCAATAAGATATTGCATTCAATTGCCGTGGGCGGTTCAGTACAATAGTTTTTGAGTGTTTGGTTTCTTCAACAAAAATCTGATCTTCTATACGTGTTGGAGTATGATTGGTTGACATTGCCAAAGCCATTTTGGCAGCCGCAAGTATATTTGGATTTCCAACCAAGGTTTTGATGGTGCGCTTTGACGAAGAGGGAGCTCGGCAGGCACGAAAGAGCTTGGAGAACATGTCCCGTGATAGAAGAAGTCGATTATGAGAAGGCGGGGGATATTCCAA

**SENSITIVE TO FREEZING 2 (SFR2):**

CAATTAGTTAGGTAGGTACCTTCTAGCAATTAAGTAGCTCAAAATTTACCACAAAAAAGGGTAGAGCATTTGGCAAGAAGTAAATTTGGCGCGCACACTAACAATTAATAGCTTAGCTTTGGGCCTCTACTCCTCAAGTATTGATGTATACAATTAGTTAGGTAGGTACCTTCATACCAAGTGAACTACAATGAAGTACCTTCATACCAAATGAACTTCAATGAACACGCAGTGTATTTCATTCAAGAGATGAAAAGCACTTCGTCGCAATGATTGAAAAGTTGTTATGGATCCCTCACTCGGTGGATTACAGCTTACTGTGTACACCAGGAATTCTACAGAGGATAGAAAGCACGTTTCAAAGCTTAAGAATGTAGTTTCGCCGTGACAATGAGGAATGAATATCTTTCCAATCTGTACAAAGTTAAGCTACAGCTAGGGCTGGAACCCTTGTTTCTGTTCCTTGCGACTCCGTCTCTTCACTTCTTTCCACATCAGACTCGACGTACCTCTTGCTACGAGTGTTGGAACGCTGCCTTTTGAACAGAATCAAACAGCTCAAGAATCTGAAAAGACGGCTCAGTGGGTCATGCAACCCATCAACTTCATAATGTCCAAATCTCCAGTCTCTCTTAACGAAGGGCCTTTCTATGGGAATATCTAGACCTCCTGCATACATCAAACCACTGCTGTCTACTGCACGACAAAAGAGCCGCTTTTTCTGCTGTGATGCTGCCGCCTGCAACTCGGACCAAGCCTGGTCTCTTTGACGTTTCGTTACTTTACCCGTTTTGCAAATCTCAGAGAACAAGTAATACGAAGGCCGAATAATCCTCTTTAGATTATTTTCACGATCAACTGCCACGAGACCAAACTTTGGGCCATATCCATCAGCCCATTCCCAGTTATCACTGGTAGTCCAGAAACAGTAGCCCTGGACTTGAACACCCTTATTCATGGCTGCACGAACGGCCAAGAGATGCTCAAGAAGGTAAGGTCGACGAATGTAATCAGTGGCGTCCGAAACACCATTTTCAGTGATGATGTACGGAATGTTGCGAGTTTTATATCGATCATGGAATGACATAAGGACACGGTACAGGCCGTCAGGATAGACTCCTCGCCCTGATTCGCTGTATTCATCATTTTCCACGAGCTTCAGCCCCGGTGCAGAAACTACTTCCTGACCATAATAGTTTATGCCAATAAAATCAAGCTTGTGACAAATCTCATCTATATAAGGAAAACGAGTCATCTGGTCACTTAGTGCCACTGCTGTGATGTCAAAAAGACCATAAGGTCGCATAAAGGAGACATGATGGGAAACACCAACTTTTGCCCTTCTCTGTCGACTCTTACAGAAGTCGTGGATAATGTCATAAGCTTTTAAGTGTGCTTCAGCCATCAGGTGCATCGCTTTGTTGAACGCTCCTTGCGGTAAGACTGATGCTACCGTCTCAAGCAAGTCAGGATTGCCACCTGGCCACGCTCCCGCACAATATGTCAGCATAGCGAATACATGAGGTTCATTGAAAATGATCCAAAAATCCACCATGCTCGAGAACTTTTCCACTGCTATCCGAGTGAAGTCCAAAAAATAATCGAGTGTCTTCTCCACTCTCCAGCCACCATATGCCCCTGCCCATGGAGGAAGAGAATGATGAAACAATGTAAGCATCACTTGCATGCCACGGGCTCGCACCTTTTCAATAATCTGTTTGTATCGATCTAGAGCTGCCCAATCTACAGATTGCTCTACTCCTTCCACAGGTTCTGTTGGCATTACTCTTGTCCAGTCGATGCCCATTCGATAAACTTGAACATTTGTTCCTTGAGCCAAATTGATTTCCGTATCAGGATCAGACCAAAATTTTAACCTTTGTTCCGGTTTAGAAGCATTGTGCCACGCAGCCACATTCGTTTTATCATCCTCAACAAGTTTCTCATAACCTCTAATCATGGCCTCCATTGCAACTTTCATCAGCCTCTTGCCTTGTTTTGGAACATTGAATTGCAAAAGAGGATCTTCAGTGGTTCCCTTGCCTCTACTAAGCTGTGACTGATCCGCATTTTCATCTATTTTGATAGAATCACTGCGTTCAAGCATCTCATCTGTATTGGCTCGTGCTATTGCCTGCTGGTCGACATTCTCCTGCAGACTTTCACTTGCTATGAGGCTAGCTTTATCTTTCTCCTGACCTTGGTTTACAAGTTCCCACTCCTCCACTGAAATTGGGCTGTTCGACTCTTGTTCCTTTCCAGGACTACCTTCAGTTGATCCTTCTTGCTCTCCTGAAGATGCCTCTTTTTCCACATTATCCGCTCGCTCTTCCTCACTTTTATTCTCTGATTCAGCATCAATTTCTTTTGATGTATCACCTAGAAAACGTTTTAGGGCTTTTTTGGTGTCGCTATCTTCCTGCTTTGGAGGCGTATCTTTCGCAAATTGAAGCCAAGCATCATCCAATTTGTCCTCCACATGTGCGGGAGCAGTTGCAAGAGCAAAGAAAAATCCACCATCGTCCACATCTCTGTCAATGGGAAACTCTGCAAGGATATCTTGATCCTCATCGATGGGGCTGTCGAAAGGGGCGAGGTTCTGTTGTCTATACCGCCGATAAGAGAAGACATTTGCAGCAACACCCAGCACACCCACGACGCCAATCAGCTTTGATACGTTGAGAAAAAGGAATACCGACATAATTCTGAGTTAAACCACCACCCAAATGGAGCCCAAGGTCAGGGAGAAAAAGCTGGAGGAAGACACAGAGAGAGGCGTCAGCGTCGACGGCGAGATCAAGTGAAAGAATGGCTCTCAAACTTCAAAGGATGCGCACCG

**ENOLASE 2 (ENO2):**

AAATTCTATACCTAATTTTATTTTATTTTTTTTAAATTTTTTTAAAGTGGGTGTATTCTATACCTATTTTGAGAGAGTTGTATTCTAAGAGAGTTGTGAGGAACAGTAAATTCGGTTTAGCTAACATTTCAAGCTCGCCTGCAAGCTTCCATGGCAACCATCTCCTGCGTTAAGGCCCGCCAGATCTTCGACAGCCGTGGCAACCCTACTGTCGAGGTGGATGTGATTCTTTCAGATGGAACATATCACAGGGCTGCCGTGCCAAGTGGTGCCTCCACTGGTGTATATGAAGCATTGGAACTGCGAGACGGGGGGAAGGACTATCTTGGCAAGGGTGTATCAAAGGCAGTCAATAATGTGGTGAAGATTATTGGCCCAGCTTTGGTTGGCAAGGATCCAACAGATCAAACAGGCATAGATAACTTTATGGTACATGGACTTGATGGAACGAAAAACGAATGGGGCTGGTGCAAGCAAAAGCTTGGAGCCAATGCAATTCTAGCAGTATCATTGGCGGTGTGCAAAGCTGGGGCAGCAGTCAATAAAATTCCTTTGTACAAGCACATTGCCAACTTGGCAGGGAACAGTACTCTTATACTTCCTGTCCCTGCCTTCAATGTCATCAATGGCGGGTCACATGCAGGAAACAAGCTTGCAATGCAGGAGTTTATGATTCTTCCTGTTGGGGCTTCCTCATTTAGTGAGGCTATGAAGATGGGTGCAGAAGTGTATCATCATCTTAAGAGCGTGATAAAGAAGAAGTATGGGCAAGATGCTACTAATGTTGGAGATGAAGGTGGTTTTGCGCCCAACATTCAGGAAAACAAGGAGGGTCTTGAGCTTCTCAAGGTTGCAATTGAGAAGGCTGGTTACACTGGCAAGGTTGTCATAGGAATGGATGTTGCAGCATCAGAGTTTTACATTGAGAAGGAGAAAAAATACGATCTTAACTTCAAGGAGGAGAACAATGATGGAAGCCAGAGAATTTCTGGGGATGAGATAATCAACATGTACAAATCCTTCACAGAGGAGTTTCCCGTTGTGTCAATTGAAGATCCCTTTGATCAGGATGACTGGGAACATTACAGCAAGTTCACCAAACAGATTGGTGAGAAAGTCCAGATTGTTGGAGATGATTTACTTGTTACCAACCCGACGAGGGTGGGGAAGGCAATCAAGGATTTGTCATGCAATGCACTCCTCTTAAAGGTCAATCAGATAGGCTCGGTAACTGAGAGCATTGAAGCTGTGAAGATGGCCAAGAAGGCAGGCTGGGGAGTCATGACCAGCCACAGAAGTGGAGAGACGGAGGACACTTTCATTGCCGATTTGGCTGTGGGGCTGGCCACAGGCCAGATCAAGACTGGTGCGCCTTGTAGATCAGAAAGGCTTGCCAAATACAACCAGCTCCTTAGGATTGAGGAGGAGCTGGGTGATGCAGCTGTCTATGCTGGAGCAAGCTACCGGCAACCAGTAGCACCTTATTGAGTTAAGGTTGTGAAAATCTGCACATGGAGTAGTAGAAAAAGCACTTCGAAATAATTCTACAGTTTTTCTAATAATGGATGCTTGCCCTGGTAAGGGCAGATTTTCTACACCCATCCTCCCGAAGACATTAGACCATCTTCAAGGTAGAGACATTTCGAGGCTTACAATTGTAATAAAAAAACGGAGGCTTTGCCACTTTTGGACTTATGTGGGTTCTTTTTGGAAGTATCAAATTTTATGTCAAAAATACTAGCAATTTTCAATTTAAGCTTTTTGATTTAAAAAAAA

**Δ (8)-FATTY-ACID DESATURASE 2 (SLD2):**

AGGCGATCTGCCTGGGTTACGGCCTACAAGGACGTACATTGTCTTTTCCAGCCTGCGGCCCTGAGTGGTTATGACGATGTCGAGGTATATCTGTGTCGCTCTTTCAAGAGCGCGAAACAAACCCAAGTGATCTTTTGTAATTTTGAGAGGTGGGTCTTCAGGCAGGCGGGGGCTTGGAAATTTCTGGCTATGGCGGCTCTTAGGCAGCAGGAGCAGTTTGAAGAGAAGAAGGCACACGAGCTTCGTTCCGTGAAAGCAGAGGAGCTTCAGAAGCACAATTCTGCAGATGATTTGTGGATCTCTATCAATGGCAAGGTCTATGATGTGACACCTTGGCTTTCCTTGCATCCTGGCGGCGACATCCCTCTGCTCAGCCTTGCTGGCCAGGATGTTACCGACACCTTTCTTGCTTTTCACCCCCCCTCTGCGTCTGCATTCCTCCCCCGTTTTCTTGTTGGCACCCTTGCAGATTACCATGTGTCGCCGCTCGCTTTCGACTATCGCAAGACCCTTACAGATCTGAAGAAAGCAGGTCTCTTCAGAAAAGATCTTTCTATCTATCTATGGATCTTTGTCGCCATCGTGGTCATGCTGGGCTTATCTGTGTCAGGTGTGTTATTGTCTAGCAGTTTTTCAGTGCATCTTGTTAGTGCATTCCTATTGGGATGTGTGTGGTCGCAATGTGGATGGATTGGTCATGATGCAGGGCATTCTCCGTTGCTCGAGAAGCCCTCCTTGGATAGGTGCATTGCGTTGCTTGTTGGAAACTGCATTTCAGGGATTAGTATCAGCTGGTGGAAACGAAATCACAATGCACATCACATCTCGTGCAACAGCCTTGAATATGACCCCGATCTTCAATACATCCCCCTTTTTGCAGTCTCTTCCAAGCTCTTTTCATCGTTGCATTCATATTTCTACGAACGCAAGATGGGCTTTGACCCCATTGCACGCTTTCTTGTCAGCTACCAGCACTGGACATTCTACCCGGTCATGGCAGTTGCAAGGATAAACCTCTTTGCACAGACTTTCATCTTGCTTTTGTCAAACAAGAAAATTCAGAACAGGGGGCTGGAGATTGCGGGCGTTTTGTGTTTTTGGGCCTGGTTTATTGCGCTTCTTGCGCACCTGCCAAACAATTTGGAAAGAATGGCTTTTGTGTTAGTGAGCTTTGCAGTTACGGGCGTACAGCATGTGCAGTTTTGCTTAAATCACTTCTCTTCACCAGTGTACAATGAGCGGCCATTGAGCACGGCATTTGTGCAGGCGCAGGTAAGAGGGACATTGAATCTTTCGACCCCAAGTTGTTTGGATTGGTTCCATGGGGGCTTGCAATTTCAAGTGGAGCATCATCTCTTTCCGCGCCTACCACGGCACAATTTGCGAAAGATCTCTGGTGTGGTGAAGTCACTTTGCGACAATCATGGGCTGCCTTACATTTCTGTTAGCTTTTGGGAGGCCAATGCCATGATCATACGCACATTACGAACTGCTGCTCTCCAAGCCAGAGATTGGTCGAAGCCAGTTCCCAAGAATCTCGTGTGGGAAGCAGTAAATGCGCAAGGGTGAGGGTCTCTCGGTGATCTATTTTTTGAATAGTGTGTTTCTGTCTCTTGCTCGTGTGCGTCTTTGTAACAACTTGTTTTATGCTACTGTTATTTCTTGGGCATAGGTTTATTGCATTCACCATTTGTAATCGAATCACACAGACCAATCTCTGTCAAACTGTGAGGAGTTATCTTTCTACCTTTTAGAAATTTAATTTTCTTTAATCTTAAAAAAA

**LONG-CHAIN BASE KINASE 2 (LCBK2):**

CACACACCAAGAAAAGATGATCACCTCTTTTGGCATTCCTCTCCCTAGTACAGACAAGGACAAGTTCCAAAACAAAAGCATCAAAAAAGGGGAGAAAGCCTGGCCAAAAGTCTCCATGTTTAAACACATTCATTGACAAATACAAATCTGCTTCTGCAGTTGCTGACCCTTGGCTATCTGTATACCACTATGAGACCTCTTCAAATACGACAGGCTTCTATTATAATGCCGTATGATTCATCCAAACTCACAACCATGCCATTTGCTCTTGATCATCCTTGCCATATTGCAAACACAGAACTTGGAGATACAGCAGATAACCCAGTCTCTCCTGCTAGACAAGTAGATCAATGGCTCCAGGAACTACTGTCATGGTTGCAGGAAGAAACCCAACATGCTCTCCATCAGCTTGAATAAAGACATCTCTCTTCAAGGGTTCACTCCCACCCTGATGTACCTCCCTCATTTCAATTTTATGGGCTCTTTCAGACAACACGTTTTTGACTGATGTATGAGAGCCGTTATAAAGCTTGTGAAGTTTAAGAAGGAAGTCAAACCACTTGAAGTTCTGCAATGTTACTATTTCAAAATCTCTGCTTGAAGGATCTGCACTAGGAGTAATCTTCATTCCACCACCAAAATACTTGGCATTGCCTATAGCTATAGCTGTAACTTGTGAAGTTTGTTTCCACTCTCCATCATCAACTCGGATTTTCAAGTCTTCATTTTTATGACTTGTAAATGCTTGCAATGCGCCTATTACATAGCACAGGTTGCCGAACTGTTTATATTTAGCAGCATGATAGCCAGCTTTTGCACTTCTGCCTTCCAACATTCACAATATCTTCAGCTCATAAAAGTTGGAACAGACCTACATCCCTTGCTCAGCTAGAATACTTGTACTGCAAGCTTTGTCATCAAAACATAATGCCCACTTACAAATGAACATCTGCAACATTCACAAAATAGCGGCCAGCATTCTCGTTATCTGCTTTTGCATACCCAACATCAATCCTTTGTCGTCGACCTTTCATTATTCGATCCACTGCCTCGATGGGATTGTTTGTCCAACCAAAAGTTCTTGCAAAATCTGAACCAGTACCAAGAGGAATCAACCCTAGGGCTGTTTTTGGTGAACTTGATCCATCTTTTGATTCAACAGGCCTTCCATCTTCAAAAAACCCATTGACCACCTCATGAAGTGTGCCATCACCTCCGACAGCAACAACAGCATGAGCTCCACTCCGGATAGCCTCTCTTGCAATTTCTACTGCATGCTGTGGCCCTGAAGTAAAAGCTTCATGTACCTTGCAATCTTTATCGAGCCTCGTAATTAGCTTTGGAAACAGATTCTTCCATTCCCTACCCGTGTGACCATTTGCACCATTTGGATTGACAATGAAGACGTAATCTCTGGGCATGGCCACTGGCCGAGTTTGCAGACACCTCTCTCCAGCTGCTGATGATCTGCAGGGAAAAGGAGAAAACGGAGAAGCAGCCATGAAAGTTAGAAATGGAGTTAGGGTTTTGTGACAGGGTCATTTTGGACAAATTGAAATGCAGAAAAGCGCAAGTGGCAGCATCATGACAAGAGAGAAATGGACACACAGTCG

**FORGETTER 1 (FGT1)**:

GAGATGGTCGTTTCTCCTCCAGCACCAGCAGGTGCAGCGATGGTCCCGACAGCATGTCAGGTGCGATGTGCGGGATGCCGAGGTATCCTCAACGTGCCTGCAGGAATTCTGGAGTTCCGGTGCCCTAAATGTAAGCTGCCACAGATGCTCCCGCCTGAGCTCCGGCCACAAGCGGCAGCATCTCATCAGGCTATCGACCCAACGAAGGCGCAGCTCCCGTGTGCCAATTGTAGGGCCCTTCTCAATGTACCTCATGGCCTCGAGAGGTTCATCTGCCCGCAATGCCGTGTAGAGCTTGCTGTCGACCTCACCAAGCTCCATCAATACATTAGCTCCCTTGGTGCTGCCAACAAGGACTCGGCGCCGGTTACCCCGCCTGCTGAAGTCGAGGATGTGAATGAGGCTGCCCTTGAGGTCGAACGAGAAGAGGATGAAGGAGGTATAGTGGGAGAGACTTTTACAGATTATCGTCCATCAAAGCTCTCTATAGGACCCCCTCACCCAGACCCTGTTGTTGAGACATCATCCCTTTCGGCAGTACAACCCCCCGAACCTACCTATGAATTGAAATTATTGGCAGATGTGGAAGAAGCGAAAGTTCTTTCATGTCTGCAGCTTGAAACTATTGTGTATGCCTCTCAGAGATATATGCAGCACCTTCCAGATGGGTTACGGGCAGGCTTTTTCATTGGAGATGGAGCAGGTGTTGGAAAAGGACGCACAGTTGCAGGCCTAATTTGGGAAAATTGGCAACATCAAAGATGCAAAGCGTTGTGGCTATCAGTTGCATCTGATCTGAAATATGACGCCCGGCGAGACCTTGATGATGTTGGCGCTTCCTCTGTGGAAGTGCACGCACTTAACAAGCTGCCGTACGGAAAACTTGATTCCAAGGCTGTGGGTGTGAAAGATGGTGTTTTGTTCTTAACTTACAGCAGCCTCATTGCATCTTCAGAAAAAGGAAGGTCAAGGTTGCAGCAACTAGTCCAATGGTGTGGATCTGATTTTGACGGTCTCTTGATTTTTGATGAGTGTCATAAGGCCAAAAATTTGGTTCCTGAATCAGGAGGTCAAGCTACTCGTACCGGTGAAGCTGTTTTGGAAATTCAGGCTAGATTACCTGAAGCTCGAGTTGTTTACTGCTCAGCAACAGGAGCCTCCGAGCCTCGCAACATGGGATATATGGCTAGACTTGGGCTTTGGGGAGTCGGCACTGATTTTCCCAACTTCCAAGCTTTTTTGGGGGCATTAGAAAAACGTGGGGTTGGAGCTCTTGAACTCGTGGCCATGGACATGAAGGCAAGGGGCATGTATGTGTGCCGGACGCTTAGTTTTAAGGGTGCAGAGTTTGATGTGATTGAGGTGCCATTGGAGACAGACATGCAGAATATGTACAACAAAGCAGCAAGATTGTGGGCAGAAGTGCGAAAAGAGTTTATGCTAACAATAACAACATTGTACGATTGTGAAAAGCCCAGCTCAAGTCAGTTGTGGCGTCTTTATTGGGCAAGTCATCAGCGTTTTTTTCGACATATGTGCATGTCCGCAAAAGTTCCTGCATCTGTGAGATTGGCAAGGCAGGCTTTGGAGGACGGCAAGTGTGTTGTGATTGGTCTTCAAAGTACTGGGGAAGCTAGGACTGAAGAGGCTGTTTCTAAATATGGCAATGAACTGGATGATTTTGTGTCAGGGCCAAGAGAGCTTTTGCTGAAACTGGTGGAAGAGAATTATCCTCTTCCCCAACCTGCAAAAACATTACAAGAGGAGATGGGAAAGGAAGGCTCAAAAAGGAAGCATTTTAGAGGTCTCAATGTGTCAGGAAGAAGAAGATCAAGGAAAGGGAGAAAACCTGGTGATGAGCTTGACAGCGACGAAGACTTTGAAACTGAATCTGATGCAGAATCAGGAGAAAGCACTGAGGGCTTGAACATTTGCAGAATCTGTGGCTCTGATGATTCCAAAAAACTCTCTATAGAATGTCAAGATTGTGATGCTTCGGTTCATTATGTATGCCTTCCGTTGGCATCAACTGGTCCTACTTCCGAATGGAGATGCCCTCCTTGTGATGAAGCTCAACGTGCATACATCAATCAACTTCAGGTGAGGTATGAAAAAGCCCTAGAGAGAAAGCTCAAGTGCTTGGAAGCTATAAGGTCATTTGAGCTTCCAAACAATCCACTGGATGACATTATTGACCAGCTTGGTGGTCCTGATGCTGTTGCCGAAATAACTGGTCGCCGTGGATTTTTAGTACGAGCTGCAAATGGCAAGGGAGTTATCTATCAATCTCGAAGCACGAAAGATGTTGCTATGGAAATGGTCAACATGCACGAAAAGCAGCAATTTATGGATGGGAGAAAACAAGTTGCTATAATATCCGAGGCTGGGTCTGCTGGTGTATCTTTACAAGCTGACAGACGAGTCAGCAATCAGAGAAGAAGAGTACATATAACTCTGGAGCTTCCCTGGAGTGCTGATCGGGCCATTCAGCAGTTTGGGAGAACGCATCGTTCCAATCAAGCATGTGCACCCGAATACAGGCTTTTATTCACAAACCTGGGGGGTGAGCGGAGATTTGCATCAATTGTAGCAAAGCGTCTGGAGACTCTTGGAGCATTGACCCAAGGGGACCGCAGGGCTGGTCCATCATTGAATGCTTACAACTATGATAGCATGTACGGAAAGAAGGCTTTGGCTCTAATGTACCGATCAATTATGGAGCAGTGTGATTTTCCTGTACTACCCCCTGGTTGTTCAAAGGATGGTGATTGGCTACCTCGAGACTTTTTCAAACTTGCAAGGGCCTCATTGCTGGCGGTTGGGATTATCAGAGATGCAGTACTTGGTGATGACCCGGAGAAGGATTTTGGAAAGCCTTCAGGAAGGATTCCAGAAGCTGATACTCATGATGTGGCACGCTTCCTGAATCGTCTATTGGGTTTACCCCCTGATGTACAAAACAGGTTGTTCGAATATTTCGTCAGCATCTTTGATCTCTTGCTGTCTGATGCTCGTAGAGATGGACAGTTTGATTGTGGAATTGTAGATGTCAAAGCTAATTCTATTGAGCTTCAGGGCACACCAAAGACTGTACATGTAGATCCAATGTCTGGAGCATCAACAACTCTTTCTATTTTAAAGGTCGATCGAGGGCTTACTTGGGAGGCTGCCGTGACGATTCTAGAGCAAAATGACACTGAGAAGAAATCTAACAATGGGTTTTACGAATCAAAGAGAGAGTGGTTAGGAAGAGCACACTACTTGCTTGCAATAAATTGGCCCAAGAGTGGATCTGGAGACTATCGAGTTTTCCGACCTGCCAGTGGAGAAGCTATAAGGGAAATGCCAGAAGATGAACTTCTCAATAAATACTCGAAGCTGACCTCATTGGACATGGCTGAGAAGAAGTGGCAAGAGGAGTACAACATCACAGAGGAGCAGTGCATGCATGGGCCAAAGTGCAAAGTTGGATCAGCTTGCACAGTGGGCAGAAGAGTTCAAGAAATGTTTATCTTAGGTGGACTTATTCTTCCAGTTTGGGGCACAGTAGAAAAAGCTTTGTCCAAACAGGTACGCATAAGCCATAGGAGACTCCGAGTTGTATGGTTAGAAACATCTTCAGATAATCAGCGCATAGTCGGTCTTCACATCCCCAACCCAGCTGTAAATGCTGTTATATCTGGTTTATCTCAAATGCAAGGTTCACAGCGCAATGATGCCATCCAGCGCAACGATGCTGCTGAAGCTTAGTTGTACAATCAGAAGTTGTTTGCACTGTTTGAGATCAATATCCCGAAGATGCTTTGCACATGAATTTTAGAAACTCAAATGTAATGGTGTACTTTGGCGCTTGTAAATACTGCTACCTAGCAAAAGGTGTATGCATTCATTATTTAATGTACAAGTGTCTTAAGCATCATACCTAGGTTCTTGCACCGAACCGCACCTAGCTAATAGGCTAGGTATGTGGTGCCTTCTTCGCAGAAGCTGATGATGTCGTTGTAATCCGCCAAGTTCAAATTATGGTGGGAAATCATCAAAAATTTTGTGTGACAAGACCTGGGTTGGAGTTTCATGTCAGAATCCTTTAAAAAATAAAATTCCAAAAAACATATCATTCTATTAAAAAACCCTCAAGGGCAAATAAGAGGATTGCG

**RNA-BINDING KH DOMAIN-CONTAINING PROTEIN RCF3 (KH28)**:

GTAGAAAGCAAGCAGAGCGACAGACGAAACAGGGGAACGCTCAGTATCAAGCAGGGCTCTTTTTCTGCTCCCATCGCCATAGCTGCGCCCCCAGTGTTTGTTTAAGTGGAGATGGATGATCCCTACTACAGATCCAGAGCGAAACGGAGTTATTCTTATCGTGAGCGAGAATATCTGGAACCTAGTGGAAGGCAGAAAAAATATGTCTCTGAAGATCCATTTAGTGGTCTTCCCCCAATTCTACCCAAGCCTGCTCCTGGTGACACAGTTTTTCGGTTGCTATGTCCGGAGCCAAAAGTAGGATGTGTCATTGGCAAGGGTGGCAGTATTGTGAAAAATTTGAGACATGAAACGGGTGCTAAGATTTTCATTCCAGATTTTTTTGAAGGTGCAGAAGAGAGGGTCATATTTATATCCTCTTCTGAAAGTGATTGGGAAAGAATTAGGGACATGGAGAGGGGAAGGCGTGATAGGGATAGAGACAACAGGGATGTTAGGATCTCACCTGCGCAAGATGCACTCATGCAAGTTCATGCAAGAATTTTAGAAGGTGGGTTTGCAAGTGAGTCAAACACAAACGATGAAGATTCTTCTCGGGGTGTCACAACACGCTTGTTGGTGCCTAACAGCCAAGTTGGTTGCTTGCTTGGGAAGGGAGGGCATATAATTGAGCAAATGCGCCAAGAGAGCAAGACACAAATTCGCATTCTGCCTAGAGATCAGGCTCCCTTTTGCTCGATGCCTTCGGATGAGATTGTACAGATAGTGGGGGATGCCAATGCTGTCAAAAAAGCTTTGCAACTTGTATCCACCCAGCTCTTTGAGAATCCACCGCGTGAGCGTCACCCTGGGGCATCTCGCTCCCCAAGCCGTGGTGCTTTCTTATCAGAAGATTTGAACGCTCCTTATCAGCCAGCCTTAAGTTCACGTGGAAGTTACAATGCTGGACAGAATCAGAAAAGTCTTCCATTATCGCATTCTGATTATGGGGCTGGTGCTAATGTCATGGGGGCGCACTTTCATGGGTCTGGCAAAACACATAGAGATGAGAAATGGGATGCTCCAGTTACAAACGATGATTTGGTGTTTCGCATCTTATGCCCCAGAGAAAAGATTGGAGGCATTATTGGCAAGGGTGGAACTATCGTTCGTAGTTTAGAGGATGAGATAGGTGTGAGCATCAATGTTTGTGGTCCTGTTGCCGGGTCAGATGAGAGAGTTGTCATGATTTCTTCCGCTGAGCATCCCGATGATAATATATCCCCTGCTCAAGAAGCATTGTTACATGTGCAAACCAAAATTGCTGATTTAGGCCCGGATGAGGATGCAGTCATTACATCAAGGCTTTTAGTTTCTCCAAATGAGGTCGGTTGTCTTCTTGGCAAAGGCGGCAGTATTATAGCAGAGATGAGAAGGTTGACTGGTGCAAACATTCGAATATTGGGCAAGGATGATCTTCCAGCGTGTGCACTCCCCACTGATGAAGTTGTGCAGATTGTAGGTGACATAAGAAATGCTCGTGAAGCTCTTGTGCAAGTGACCAAAAGGCTCAGATCTAATCTGTATTTTGAGAAGTCGCTTGAAGCAGGAGGTGCACCATTATCCTTGGGCAACTCCACGCCTCAGAATAGACATGATTTTCTTTCTCCAGGACGGATTGACTTGCCAGCATCAGGAGTTCAAAGTGCTGGACGATCAGCTTCAAGCTATGGCAAAACATCTCCCCCTGGTTATTGGACTGGACAGGATCTTCTCGGTGGAAGTCGCCTTGCAGGCTACGAAAGCCCAGTTCCTCATAAAGATTCTGAATTGTTTAGGACAACAGGAGCTCTTGTGACAAAAACAACTGTTGAGGTTGTTCTACCTAGCCGAGTTGTCGGATTGATTTTTGCGGATGGTTCGGCAATCGCTCAGATTCGCCAGATATCTGGTGCACAAGTAAACTTTCTAGATGTGAGACCTGGAGCGGCTGAGGCTGTCGTGGAAATTTCGGGCACACCTGAGCAAACACAATCTGCTCAAAGATTGCTACAAGCGTTTATTCTGAGGGGGCAGACATCCTCTCTCGGTTATGCATGATAAGCCTATCAACTTGGAGAAGGGTTCTTAATGCTTTACAGGAATGCTTTGTCCTCGTGTACGCAGAAATTCCCTGTTGGGGGAAATGGCCTAAACACAATGATCTTGAAATATCCCATCATAAATCTGACTACTATCCAATTCTTTTCTGTCGTCCGTGCTTGAAAAGCCCCTTAAATGGGAAATGTCAGTCCTCCCAGGCTAACCTTCAAAGACTCCTGTTTTTTGATTCAGTGCTCTACAGCAAATTTAATATACATTAATTGGGGCTCTTGCTACAATACCAGAGGGGCTTAACTTTCTTGAATGCATGTGCATTGTGTACATGGTCCCTGCGTTTTTCTCCTTGTAGGGATGAGCCAATTTGTACACAAGTCTCTCATGACTGCATTATAGAAATTTTCTTCCTCGAGGGATGAGCTAAATTTTTACACAAAGCTCTCAAAGTTGCATTGGAGATGCCATGAAATTTTAATCAACGAGTGAAGAGAAGCATGAATTGTTAATCTTTTACTACCTCTGGTATGCTGTATTGAGTAATGCAAGTGACTAAGTTTCTCATTTGGTACTCAACTGG

**HEAT INTOLERANT 4 (HIT4):**

GAAATTCATTTGGCACACAAGATAAGGCTAGTACAGCATGTAAAACTTATGCAGAACCTTGGCTTTGCTTAAGAAAAGACCAGTATCATTCTTGATACAGAGGCATTCTTGATCCTTTATCACAAAAATAGGTGTGCTCATGAAGCAATTTTTAGAGTTCAAGGATTCAGCTTTGCAAATAACATTTTGTTATGTACTAAGTTAATAAAGCCATACATCAAAGCTGCGCATACTCAATGGAACGACAGAGACAATTTAAAAAACAGACTAAGTCTCATATAAAAGAGACCAACAAATGAAACAAACCATCAATGAAATATACTGAATGTAATGTACACTACAAACAGAAACACCAAAATTTTGAGGCAACTCTTGTCCCTTGGCTCCTCGTATCAGCTGCTTCTAGCTCTTTGATGTACACTTTGTTGTAGCCTCGGTTTTTCTTCACAACTGTTTGTAACCCAAATTACACTTGAAGTGCAGTCTTTAGGTCTGGTACCAAACTGAGCTTCTATAGCACCTCAGAGGCTTTACCGTAGTATCTATTGATAAACGGACTCTTCAGGGCAGAGATGTCTGGGGTGCCTGGATTTGGACGAGGATAAAACTTATAGAAACGTATGTTGTCAAAAAAAGCCCTAGTCTCTGCAGTCATCCCTTCAATTTTTTTGCGCCGCTCCTCTCTGGCCTGCCGTTGCTGTTTTCTCTCTTCCGAGACTCTTTGCTTCACATAAGTCTTAAAGGCATCCTTCCCTTCTACAGTTAGGCTCTCATCTTTCACCAGGTTATCAGCCATTTCCTCAAGCTCGTCCAGCTCCCAATCAAATTCTGTAAGGAGTGGTGGCCTTTCATCGTCAAAAGGATAGATTATGGAGACGACTGTATCTGGCTCTTGCTCTTCTTCTTTGAAAGGATTGTTGAAATAGGGTAAGCAATATTCATATTTCTTGAGCCTTTCTTCTTTGAGATTTCTTAAAGCAACCCTTCGTTGGACACACTTCAAGGTAAAAATTTGAGATTTGTACCTCTCCACAGAAATATGGTCTCTATCAGCCGGTATGTATGGGACCCAATCCATCTTCATCTCTTTCATTGGCACAATGAATTCACCTTCCATTTGAACAGACTTTATCCCTATTTTGTCTGAAGGGGCATGTGATGCTACAACTGCAACGACAGCTGGAATATGCACCACCCTTAATACATCTCCCAAAGCGAGAAATTGAGGTTCTGTGCATCCAAATAGAAAGACCTTCTTCCCATGTAGAACTCCATTCTCTTCGAAAGCTTCTTCTAGATGAGAGAAGTCCCAATCCATTTCGTAAACACGGTCGTATTGGTCCCACTCTGTGCCCACTGGAAATGCCTCCTTCCATAAATCTCTCAGGGGCGCGTGAGGGGGCACTTCCTCTTGCACGTAGTGGCGCCTGCGGGGAGGGGAGAGAGGCAATGCGGCAGCAGCAGCAGCAGGGGCAGGGGCGGGGGCGGGGGGAGGGGG

**SERINE/THREONINE-PROTEIN PHOSPHATASE 7 (PP7):**

AAGTTGAAAAAGATGAGCTAGCCAAGTAATGACATTGGCACGCACATTTGAATTCACACTCTTCCCAATGAATCTTATGGGAAGAGTGTGAGTTCAAATCCCGGTAGACGCATGTGCAATTGAAAGGTTCAATGGCTATGGAGGAAGCTGCAGTGGTGAGGTGGCCAGGGGACGGGCTTCTTTGCAAAGAGTGGCTGGATGCCTTCACTTCCTCTCTCGACACCGCCTCCAGAGAGCCGCCCTCTTTCTTAAGTCGCCTCCTGCCCGTTCCCGTCATCGACTCGCTCGTCTCTTCTGCTTCCAAGCTCCTCCATCGCGAGCCCAACTGCTTGCAGGTGACTGTTGAACTGTCTTCCTCTGTTGTGCTGGTGGGTGACCTTCATGGCCAACTCCATGATCTTCTCCACCTTCTCCGGCTCGCCGGCCTCCCTTCCCCCTCCCGGATCTTTGTGTTCAATGGTGACTATGTTGACCGTGGGGCTTGGGGCCTTGAGACCTTCCTTCTACTTCTGGCTTGGAAGATTTTCTTACCCCATAGAGTGTTTCTACTTCGTGGAAATCATGAAACAAGGTTTTGCACAGGGGTTTATGGTTTTGAGCAGGAGGTTATGGTGAAGTATGGTGAGCATGGGAAGCATGTTTACAGAAAGTTGTTAGGATGCTTTGAGGGTCTTCCTCTTGCTGCCATTATTGCAGATTGTGTATATATGGCACACGGAGGTCTATTCCGCCACATTGAAGCAACACAGAAGAAGGAGAAGAAGAAAGGCAGAGCTCGGAGAGCCACAGAAAAAATTGAGCACGGCAACTTGCAGCTAGGCACTCTCAGTGAACTGGCAATGGCTAGGAGGGGCCTTTTAGACCCGCCAGAGAAAGGCAACAACGTGCTAGTAAGTGACATTTTGTGGTCAGACCCTGCACCTAAGCATGGTCTTTATCCCAACCCTGCTCGCGGCATAGGTCTTTTGTGGGGGCCCGACTGTACTGAAGAGTTTTTGCAGAAGCACAAATTGAAGTTGATTGTAAGGTCCCATGAAGGGCCTGACGCACGAGTGAAACGTCATGACATGATAGACATGAGCAAAGGCTACACAGTAGATCATGTTGTGAATTCTGGAAAACTGATCACACTTTTCAGTGCTCCCGATTATCCACAATTCCAGGCTAGCAATTGTAGATACAACAATGAAGGGGCTTACATTCTGTTGGAGGCTCCAGACTTTTCATCACCAAATTTTTTCTCCTTCACTGCTGCTTTACCAAGACCAAAGGTCACGGCGTACTATGACTTCGTGAATGCTGTGGATTCAGATGAAGAAATTGATCTAGGAGCAAGCTCAGATAGATCTACTGATTGGGACAGAGGAAACAAATAGACATAGGTGGGCTGGCACTTGCTGTTTGTTGGCTGCAGAAGGTTATCAGAAATGGCTATGGCATGGTCATGTATTAGCTTGTGGATTGGCAAGCTACTAGCTGTACCAACAGATTGATTCAATGTCAAATGTACACTAAATTGCCCTTTTGAAGCAAAGCTCTACTTGAACACCCCTAATTGAAACACTCGAAACCTTCATGAAGCAACCAGAGTTGTTCTTTAACAAAGTAGAAACTCAATAAATTTCAAGAACTAAGATAAAGCCAACTACATGCGAAACTACATGCGAGATGTGAGATTATTTACAAGCTAATAGTTAAATGCCTGAATATTTTGTTTTGATGTGATATTCTTATATGGTTGTTTTTATCACATCTTTGGAGGTAAAGGGAAACCTTTAGCCATGATAAAGAGTGATGCCAAAAACATTGCTAGCAAATTTTCAGTGTTGATTGAGCTCAAGCCTCCTTTTTGTAAAGCTATCCTTGG
