## Supplemental Table 2 for "Comprehensive analysis of proteins associated with light responses and stress tolerance in the gametophyte of the fern *Dryopteris affinis* ssp*. affinis*"

**Supplementary Table 2**. Selected proteins found in gametophytes of *Dryopteris affinis* compared to fern species with sequenced genomes.

| **Protein** | ***Adiantum capillus* protein** | **E-value** | ***Alsophila spinulosa* protein** | **E-value** | ***Azolla filiculoides* protein** | **E-value** | ***Ceratopteris richardii* protein** | **E-value** | ***Marsilea vestita* protein** | **E-value** | ***Salvinia cucullata* protein** | **E-value** |
| --- | --- | --- | --- | --- | --- | --- | --- | --- | --- | --- | --- | --- |
| PGR6 | ADC00041 | 0 | [Aspi01Gene45250.t1](https://fernbase.org/tools/blast/match/show?blast_db_id=63;id=Aspi01Gene45250.t1;hilite_coords=1-699) | 0 | Protein kinase superfamily protein | 0 | [Ceric.02G050500.1.v2.1](https://fernbase.org/tools/blast/match/show?blast_db_id=67;id=Ceric.02G050500.1.v2.1;hilite_coords=1-696) | 0 | [Mvestita_S5g09409-RA](https://fernbase.org/tools/blast/match/show?blast_db_id=71;id=Mvestita_S5g09409-RA;hilite_coords=1-689) | 0 | Protein kinase superfamily protein | 0 |
| LFNR1 | [ADC09596](https://fernbase.org/tools/blast/match/show?blast_db_id=59;id=ADC09596;hilite_coords=18-375) | 0 | [Aspi01Gene25421.t1](https://fernbase.org/tools/blast/match/show?blast_db_id=63;id=Aspi01Gene25421.t1;hilite_coords=18-374) | 0 | Ferredoxin-NADP reductase | 0 | [Ceric.04G111900.1.v2.1](https://fernbase.org/tools/blast/match/show?blast_db_id=67;id=Ceric.04G111900.1.v2.1;hilite_coords=22-378) | 0 | [Mvestita_S6g00197-RA](https://fernbase.org/tools/blast/match/show?blast_db_id=71;id=Mvestita_S6g00197-RA;hilite_coords=18-368) | 0 | Ferredoxin-NADP reductase | 0 |
| LFNR2 | [ADC14692](https://fernbase.org/tools/blast/match/show?blast_db_id=59;id=ADC14692;hilite_coords=16-379) | 0 | [Aspi01Gene13178.t1](https://fernbase.org/tools/blast/match/show?blast_db_id=63;id=Aspi01Gene13178.t1;hilite_coords=16-379) | 0 | Ferredoxin-NADP reductase | 0 | [Ceric.32G074200.2.v2.1](https://fernbase.org/tools/blast/match/show?blast_db_id=67;id=Ceric.32G074200.2.v2.1;hilite_coords=16-377) | 0 | [Mvestita_S10g02304-RA](https://fernbase.org/tools/blast/match/show?blast_db_id=71;id=Mvestita_S10g02304-RA;hilite_coords=20-377) | 0 | Ferredoxin-NADP reductase | 0 |
| KEA3 | ADC24416 | 0 | Aspi01Gene40991.t1 | 8.51·10^-144^ | Azfi-s0006.g009901 | 0 | Ceric.10G024200.3.v2.1 | 1.03·10^-148^ | Mvestita_S3g03339-RA | 0 | Sacu_v1.1_s0013.g005687 | 2.6·10^-80^ |
| HCF136 | [ADC12171](https://fernbase.org/tools/blast/match/show?blast_db_id=59;id=ADC12171;hilite_coords=2-413) | 0 | [Aspi01Gene72238.t1](https://fernbase.org/tools/blast/match/show?blast_db_id=63;id=Aspi01Gene72238.t1;hilite_coords=1-418) | 0 | Photosystem II stability/assembly factor HCF136 | 0 | [Ceric.10G037400.1.v2.1](https://fernbase.org/tools/blast/match/show?blast_db_id=67;id=Ceric.10G037400.1.v2.1;hilite_coords=1-409) | 0 | [Mvestita_C41g20867-RA](https://fernbase.org/tools/blast/match/show?blast_db_id=71;id=Mvestita_C41g20867-RA;hilite_coords=1-423) | 0 | Photosystem II stability/assembly factor HCF136 | 0 |
| HCF173 | [ADC04721](https://fernbase.org/tools/blast/match/show?blast_db_id=59;id=ADC04721;hilite_coords=1-594) | 0 | [Aspi01Gene21050.t1](https://fernbase.org/tools/blast/match/show?blast_db_id=63;id=Aspi01Gene21050.t1;hilite_coords=30-554) | 0 | NAD(P)-binding Rossmann-fold superfamily protein | 0 | [Ceric.24G028300.1.v2.1](https://fernbase.org/tools/blast/match/show?blast_db_id=67;id=Ceric.24G028300.1.v2.1;hilite_coords=1-596) | 0 | [Mvestita_S8g05336-RA](https://fernbase.org/tools/blast/match/show?blast_db_id=71;id=Mvestita_S8g05336-RA;hilite_coords=93-606) | 0 | NAD(P)-binding Rossmann-fold superfamily protein | 0 |
| HCF244 | [ADC00113](https://fernbase.org/tools/blast/match/show?blast_db_id=59;id=ADC00113;hilite_coords=1-406) | 0 | [Aspi01Gene42667.t1](https://fernbase.org/tools/blast/match/show?blast_db_id=63;id=Aspi01Gene42667.t1;hilite_coords=1-412) | 0 | NAD(P)-binding protein | 0 | [Ceric.07G004000.2.v2.1](https://fernbase.org/tools/blast/match/show?blast_db_id=67;id=Ceric.07G004000.2.v2.1;hilite_coords=1-405) | 0 | [Mvestita_S5g09502-RA](https://fernbase.org/tools/blast/match/show?blast_db_id=71;id=Mvestita_S5g09502-RA;hilite_coords=26-398) | 0 | NAD(P)-binding protein | 0 |
| CYP38 | [ADC11434](https://fernbase.org/tools/blast/match/show?blast_db_id=59;id=ADC11434;hilite_coords=250-473) | 0 | [Aspi01Gene55127.t1](https://fernbase.org/tools/blast/match/show?blast_db_id=63;id=Aspi01Gene55127.t1;hilite_coords=254-477) | 0 | Peptidyl-prolyl cis-trans isomerase CYP38, chloroplastic | 0 | [Ceric.32G007000.1.v2.1](https://fernbase.org/tools/blast/match/show?blast_db_id=67;id=Ceric.32G007000.1.v2.1;hilite_coords=252-475) | 0 | [Mvestita_S10g02797-RA](https://fernbase.org/tools/blast/match/show?blast_db_id=71;id=Mvestita_S10g02797-RA;hilite_coords=225-448) | 0 | Peptidyl-prolyl cis-trans isomerase CYP38, chloroplastic | 0 |
| EVR3 | [ADC07600](https://fernbase.org/tools/blast/match/show?blast_db_id=59;id=ADC07600;hilite_coords=4-566) | 0 | [Aspi01Gene63102.t1](https://fernbase.org/tools/blast/match/show?blast_db_id=63;id=Aspi01Gene63102.t1;hilite_coords=12-314) | 0 | Peptidase M50 family protein | 0 | [Ceric.08G062200.1.v2.1](https://fernbase.org/tools/blast/match/show?blast_db_id=67;id=Ceric.08G062200.1.v2.1;hilite_coords=58-649) | 0 | [Mvestita_C264g21182-RA](https://fernbase.org/tools/blast/match/show?blast_db_id=71;id=Mvestita_C264g21182-RA;hilite_coords=104-591) | 0 | Peptidase M50 family protein | 0 |
| LHCB2 | ADC09525 | 4.75·10^-177^ | Aspi01Gene25498.t1 | 4.22·10^-171^ | Azfi-s0013.g013388 | 1.21·10^-164^ | Ceric.05G101100.1.v2.1 | 5.75·10^-172^ | Mvestita_C45g19797-RA | 3.42·10^-167^ | Sacu_v1.1_s0086.g018412 | 2.01·10^-164^ |
| LHCA1 | ADC27960 | 2.54·10^-134^ | Aspi01Gene51505.t1 | 1.03·10^-137^ | Azfi-s0549.g076444 | 3.53·10^-128^ | Ceric.07G087800.1.v2.1 | 1.89·10^-128^ | Mvestita_S6g00577-RA | 7.65·10^-134^ | Sacu_v1.1_s0079.g017788 | 1.22·10^-124^ |
| LHCB4 | ADC11351 | 5.41·10^-170^ | Aspi01Gene47726.t1 | 7.27·10^-160^ | Azfi-s0173.g055756 | 2.84·10^-153^ | Ceric.05G075200.1.v2.1 | 2·10^-165^ | Mvestita_S14g07746-RA | 7.04·10^-150^ | Sacu_v1.1_s0067.g016319 | 5.37·10^-141^ |
| SOQ1 | ADC23954 | 0 | Aspi01Gene46305.t1 | 0 | Azfi-s0061.g034958 | 0 | Ceric.02G025300.2.v2.1 | 0 | Mvestita_S9g06198-RA | 0 | Sacu_v1.1_s0015.g006621 | 0 |
| EVR3 | ADC07600 | 0 | Aspi01Gene63102.t1 | 0 | Azfi-s0230.g059204 | 0 | Ceric.08G062200.1.v2.1 | 0 | Mvestita_C264g21182-RA | 0 | Sacu_v1.1_s0119.g021268 | 0 |
| FTSH1 | [ADC01449](https://fernbase.org/tools/blast/match/show?blast_db_id=59;id=ADC01449;hilite_coords=17-708) | 0 | [Aspi01Gene32354.t1](https://fernbase.org/tools/blast/match/show?blast_db_id=63;id=Aspi01Gene32354.t1;hilite_coords=12-701) | 0 | ATP-dependent zinc metalloprotease FtsH | 0 | [Ceric.14G083900.1.v2.1](https://fernbase.org/tools/blast/match/show?blast_db_id=67;id=Ceric.14G083900.1.v2.1;hilite_coords=19-711) | 0 | [Mvestita_S5g09983-RA](https://fernbase.org/tools/blast/match/show?blast_db_id=71;id=Mvestita_S5g09983-RA;hilite_coords=12-699) | 0 | ATP-dependent zinc metalloprotease FtsH | 0 |
| HPR | [ADC10194](https://fernbase.org/tools/blast/match/show?blast_db_id=59;id=ADC10194;hilite_coords=1-390) | 0 | [Aspi01Gene63220.t1](https://fernbase.org/tools/blast/match/show?blast_db_id=63;id=Aspi01Gene63220.t1;hilite_coords=1-390) | 0 | D-glycerate dehydrogenase/hydroxypyruvate reductase | 0 | [Ceric.01G038500.1.v2.1](https://fernbase.org/tools/blast/match/show?blast_db_id=67;id=Ceric.01G038500.1.v2.1;hilite_coords=1-390) | 0 | [Mvestita_C161g21474-RA](https://fernbase.org/tools/blast/match/show?blast_db_id=71;id=Mvestita_C161g21474-RA;hilite_coords=1-390) | 0 | D-3-phosphoglycerate dehydrogenase | 2·10^-25^ |
| HPPR2 | [ADC30901](https://fernbase.org/tools/blast/match/show?blast_db_id=59;id=ADC30901;hilite_coords=4-314) | 0 | [Aspi01Gene21035.t1](https://fernbase.org/tools/blast/match/show?blast_db_id=63;id=Aspi01Gene21035.t1;hilite_coords=6-319) | 2·10^-157^ | D-isomer specific 2-hydroxyacid dehydrogenase family protein | 8·10^-167^ | [Ceric.1Z009100.1.v2.1](https://fernbase.org/tools/blast/match/show?blast_db_id=67;id=Ceric.1Z009100.1.v2.1;hilite_coords=3-314) | 0 | [Mvestita_S4g08520-RA](https://fernbase.org/tools/blast/match/show?blast_db_id=71;id=Mvestita_S4g08520-RA;hilite_coords=1-342) | 1·10^-164^ | D-isomer specific 2-hydroxyacid dehydrogenase family protein | 7·10^-168^ |
| GGAT1 | [ADC20190](https://fernbase.org/tools/blast/match/show?blast_db_id=59;id=ADC20190;hilite_coords=1-483) | 0 | [Aspi01Gene54649.t1](https://fernbase.org/tools/blast/match/show?blast_db_id=63;id=Aspi01Gene54649.t1;hilite_coords=1-483) | 0 | Alanine aminotransferase 2 | 0 | [Ceric.25G051100.1.v2.1](https://fernbase.org/tools/blast/match/show?blast_db_id=67;id=Ceric.25G051100.1.v2.1;hilite_coords=1-480) | 0 | [Mvestita_S3g03766-RA](https://fernbase.org/tools/blast/match/show?blast_db_id=71;id=Mvestita_S3g03766-RA;hilite_coords=6-481) | 0 | Alanine aminotransferase 2 | 0 |
| BCH1 | ADC18418 | 1.9·10^-123^ | Aspi01Gene69868.t1 | 9.05·10^-133^ | Azfi-s0305.g063924 | 1.31·10^-110^ | Ceric.25G022000.2.v2.1 | 1.37·10^-121^ | Mvestita_C33g20625-RA | 6.66·10^-123^ | Sacu_v1.1_s0147.g023174 | 1.14·10^-125^ |
| LUT5 | ADC13264 | 0 | Aspi01Gene03690.t1 | 0 | Azfi-s0006.g009961 | 7.05·10^-122^ | Ceric.11G037600.4.v2.1 | 0 | Mvestita_S10g02346-RA | 0 | Sacu_v1.1_s0095.g019289 | 0 |
| VDE1 | ADC13807 | 0 | Aspi01Gene33475.t1 | 0 | Azfi-s1476.g103495 | 0 | Ceric.1Z326600.1.v2.1 | 9.22·10^-175^ | Mvestita_S9g05938-RA | 1.77·10^-131^ | Sacu_v1.1_s0102.g019899 | 0 |
| LAF3 | [ADC06034](https://fernbase.org/tools/blast/match/show?blast_db_id=59;id=ADC06034;hilite_coords=7-563) | 0 | [Aspi01Gene42034.t1](https://fernbase.org/tools/blast/match/show?blast_db_id=63;id=Aspi01Gene42034.t1;hilite_coords=66-241) | 4·10^-84^ | Amidohydrolase family protein | 8·10^-92^ | [Ceric.20G061700.1.v2.1](https://fernbase.org/tools/blast/match/show?blast_db_id=67;id=Ceric.20G061700.1.v2.1;hilite_coords=6-568) | 0 | [Mvestita_S17g16304-RA](https://fernbase.org/tools/blast/match/show?blast_db_id=71;id=Mvestita_S17g16304-RA;hilite_coords=6-569) | 0 | Amidohydrolase family protein | 0 |
| PHYB | [ADC23537](https://fernbase.org/tools/blast/match/show?blast_db_id=59;id=ADC23537;hilite_coords=27-1104) | 0 | [Aspi01Gene59146.t1](https://fernbase.org/tools/blast/match/show?blast_db_id=63;id=Aspi01Gene59146.t1;hilite_coords=27-1088) | 0 | Phytochrome | 0 | [Ceric.29G025400.1.v2.1](https://fernbase.org/tools/blast/match/show?blast_db_id=67;id=Ceric.29G025400.1.v2.1;hilite_coords=27-1139) | 0 | [Mvestita_S16g10910-RA](https://fernbase.org/tools/blast/match/show?blast_db_id=71;id=Mvestita_S16g10910-RA;hilite_coords=26-1139) | 0 | Phytochrome | 0 |
| PAT1 | ADC24158 | 0 | Aspi01Gene48189.t1 | 0 | Azfi-s0011.g012766 | 0 | Ceric.39G021800.2.v2.1 | 0 | Mvestita_S2g11451-RA | 9.67·10^-171^ | Sacu_v1.1_s0003.g001584 | 0 |
| VOZ1 | ADC21586 | 0 | Aspi01Gene54808.t1 | 0 | Azfi-s0007.g010801 | 0 | Ceric.12G039200.3.v2.1 | 0 | Mvestita_S4g09318-RA | 0 | Sacu_v1.1_s0076.g017481 | 0 |
| TOPP4 | ADC09925 | 0 | Aspi01Gene72566.t1 | 0 | Azfi-s0065.g035769 | 0 | Ceric.15G041500.3.v2.1 | 0 | Mvestita_S13g01322-RA | 0 | Sacu_v1.1_s0002.g000877 | 0 |
| HT1 | [ADC01744](https://fernbase.org/tools/blast/match/show?blast_db_id=59;id=ADC01744;hilite_coords=1-452) | 0 | [Aspi01Gene03716.t1](https://fernbase.org/tools/blast/match/show?blast_db_id=63;id=Aspi01Gene03716.t1;hilite_coords=21-461) | 0 | Protein kinase | 6·10^-96^ | [Ceric.01G066500.2.v2.1](https://fernbase.org/tools/blast/match/show?blast_db_id=67;id=Ceric.01G066500.2.v2.1;hilite_coords=1-490) | 0 | [Mvestita_S2g12283-RA](https://fernbase.org/tools/blast/match/show?blast_db_id=71;id=Mvestita_S2g12283-RA;hilite_coords=104-391) | 2·10^-106^ | Protein kinase | 5·10^-89^ |
| KAC2 | [ADC10406](https://fernbase.org/tools/blast/match/show?blast_db_id=59;id=ADC10406;hilite_coords=1-1318) | 0 | [Aspi01Gene65584.t1](https://fernbase.org/tools/blast/match/show?blast_db_id=63;id=Aspi01Gene65584.t1;hilite_coords=53-1290) | 0 | Kinesin-like protein KIN-5A | 0 | [Ceric.22G049000.2.v2.1](https://fernbase.org/tools/blast/match/show?blast_db_id=67;id=Ceric.22G049000.2.v2.1;hilite_coords=1-1349) | 0 | [Mvestita_C105g20621-RA](https://fernbase.org/tools/blast/match/show?blast_db_id=71;id=Mvestita_C105g20621-RA;hilite_coords=1-1346) | 0 | Kinesin-like protein KIN-5A | 0 |
| PHOT2 | [ADC25029](https://fernbase.org/tools/blast/match/show?blast_db_id=59;id=ADC25029;hilite_coords=8-1435) | 0 | [Aspi01Gene53689.t1](https://fernbase.org/tools/blast/match/show?blast_db_id=63;id=Aspi01Gene53689.t1;hilite_coords=255-1040) | 0 | Phototropin | 0 | [Ceric.24G066200.8.v2.1](https://fernbase.org/tools/blast/match/show?blast_db_id=67;id=Ceric.24G066200.8.v2.1;hilite_coords=206-974) | 0 | [Mvestita_S13g01808-RA](https://fernbase.org/tools/blast/match/show?blast_db_id=71;id=Mvestita_S13g01808-RA;hilite_coords=177-949) | 0 | Phototropin | 0 |
| CRY1 | ADC04082 | 0 | Aspi01Gene21917.t1 | 0 | Azfi-s0004.g008357 | 0 | Ceric.03G029200.2.v2.1 | 0 | Mvestita_S12g15622-RA | 0 | Sacu_v1.1_s0055.g014444 | 0 |
| CRY2 | ADC21408 | 0 | Aspi01Gene71250.t1 | 0 | Azfi-s0229.g059128 | 0 | Ceric.09G012600.1.v2.1 | 0 | Mvestita_S12g15622-RA | 0 | Sacu_v1.1_s0187.g025130 | 0 |
| GCR1 | [ADC08606](https://fernbase.org/tools/blast/match/show?blast_db_id=59;id=ADC08606;hilite_coords=1-312) | 0 | [Aspi01Gene38560.t1](https://fernbase.org/tools/blast/match/show?blast_db_id=63;id=Aspi01Gene38560.t1;hilite_coords=111-233) | 1·10^-80^ | G-protein coupled receptor 1 | 2·10^-144^ | [Ceric.26G026500.1.v2.1](https://fernbase.org/tools/blast/match/show?blast_db_id=67;id=Ceric.26G026500.1.v2.1;hilite_coords=1-311) | 0 | [Mvestita_S16g10448-RA](https://fernbase.org/tools/blast/match/show?blast_db_id=71;id=Mvestita_S16g10448-RA;hilite_coords=1-310) | 2·10^-180^ | G-protein coupled receptor 1 | 1·10^-173^ |
| **Protein** | ***Adiantum capillus* protein** | **E-value** | ***Alsophila spinulosa* protein** | **E-value** | ***Azolla filiculoides* protein** | **E-value** | ***Ceratopteris richardii* protein** | **E-value** | ***Marsilea vestita* protein** | **E-value** | ***Salvinia cucullata* protein** | **E-value** |
| ADO1 | ADC29615 | 0 | [Aspi01Gene42456.t1](https://fernbase.org/tools/blast/match/show?blast_db_id=63;id=Aspi01Gene42456.t1;hilite_coords=46-626) | 0 | Galactose oxidase/kelch repeat superfamily protein | 0 | [Ceric.14G002300.1.v2.1](https://fernbase.org/tools/blast/match/show?blast_db_id=67;id=Ceric.14G002300.1.v2.1;hilite_coords=2-719) | 0 | [Mvestita_S9g05434-RA](https://fernbase.org/tools/blast/match/show?blast_db_id=71;id=Mvestita_S9g05434-RA;hilite_coords=21-633) | 0 | Galactose oxidase/kelch repeat superfamily protein | 0 |
| FB | [ADC15866](https://fernbase.org/tools/blast/match/show?blast_db_id=59;id=ADC15866;hilite_coords=1-1163) | 0 | [Aspi01Gene56730.t1](https://fernbase.org/tools/blast/match/show?blast_db_id=63;id=Aspi01Gene56730.t1;hilite_coords=264-1400) | 0 | Gigantea | 0 | [Ceric.25G015300.2.v2.1](https://fernbase.org/tools/blast/match/show?blast_db_id=67;id=Ceric.25G015300.2.v2.1;hilite_coords=1-1165) | 0 | [Mvestita_C25g19600-RA](https://fernbase.org/tools/blast/match/show?blast_db_id=71;id=Mvestita_C25g19600-RA;hilite_coords=132-1272) | 0 | Gigantea | 0 |
| SPA1 | [ADC09893](https://fernbase.org/tools/blast/match/show?blast_db_id=59;id=ADC09893;hilite_coords=62-1139) | 0 | [Aspi01Gene53620.t1](https://fernbase.org/tools/blast/match/show?blast_db_id=63;id=Aspi01Gene53620.t1;hilite_coords=193-762) | 0 | Transducin/WD40 repeat-like superfamily protein | 0 | [Ceric.12G047000.1.v2.1](https://fernbase.org/tools/blast/match/show?blast_db_id=67;id=Ceric.12G047000.1.v2.1;hilite_coords=1-1083) | 0 | [Mvestita_S1g06894-RA](https://fernbase.org/tools/blast/match/show?blast_db_id=71;id=Mvestita_S1g06894-RA;hilite_coords=1-1124) | 0 | Transducin/WD40 repeat-like superfamily protein | 0 |
| UVR2 | [ADC13113](https://fernbase.org/tools/blast/match/show?blast_db_id=59;id=ADC13113;hilite_coords=46-517) | 0 | [Aspi01Gene04059.t1](https://fernbase.org/tools/blast/match/show?blast_db_id=63;id=Aspi01Gene04059.t1;hilite_coords=35-530) | 0 | CPD photolyase | 0 | [Ceric.22G034500.2.v2.1](https://fernbase.org/tools/blast/match/show?blast_db_id=67;id=Ceric.22G034500.2.v2.1;hilite_coords=91-581) | 0 | [Mvestita_C471g21198-RA](https://fernbase.org/tools/blast/match/show?blast_db_id=71;id=Mvestita_C471g21198-RA;hilite_coords=81-556) | 0 | CPD photolyase | 0 |
| UVR3 | [ADC19713](https://fernbase.org/tools/blast/match/show?blast_db_id=59;id=ADC19713;hilite_coords=1-377) | 0 | [Aspi01Gene18806.t1](https://fernbase.org/tools/blast/match/show?blast_db_id=63;id=Aspi01Gene18806.t1;hilite_coords=1-377) | 0 | (6-4) DNA photolyase | 0 | [Ceric.20G004300.1.v2.1](https://fernbase.org/tools/blast/match/show?blast_db_id=67;id=Ceric.20G004300.1.v2.1;hilite_coords=57-433) | 0 | [Mvestita_S6g00622-RA](https://fernbase.org/tools/blast/match/show?blast_db_id=71;id=Mvestita_S6g00622-RA;hilite_coords=59-434) | 0 | (6-4) DNA photolyase | 0 |
| MSH6 | [ADC21641](https://fernbase.org/tools/blast/match/show?blast_db_id=59;id=ADC21641;hilite_coords=1-1361) | 0 | [Aspi01Gene71892.t1](https://fernbase.org/tools/blast/match/show?blast_db_id=63;id=Aspi01Gene71892.t1;hilite_coords=1-1016) | 0 | DNA mismatch repair protein Msh6-1 | 0 | [Ceric.09G018600.1.v2.1](https://fernbase.org/tools/blast/match/show?blast_db_id=67;id=Ceric.09G018600.1.v2.1;hilite_coords=1-1356) | 0 | [Mvestita_S8g04982-RA](https://fernbase.org/tools/blast/match/show?blast_db_id=71;id=Mvestita_S8g04982-RA;hilite_coords=16-1373) | 0 | DNA mismatch repair protein Msh6-1 | 0 |
| HAM1 | [ADC03065](https://fernbase.org/tools/blast/match/show?blast_db_id=59;id=ADC03065;hilite_coords=1-447) | 0 | [Aspi01Gene08937.t1](https://fernbase.org/tools/blast/match/show?blast_db_id=63;id=Aspi01Gene08937.t1;hilite_coords=1-457) | 0 | Histone acetyltransferase | 0 | [Ceric.08G064500.1.v2.1](https://fernbase.org/tools/blast/match/show?blast_db_id=67;id=Ceric.08G064500.1.v2.1;hilite_coords=1-445) | 0 | [Mvestita_C856g22136-RA](https://fernbase.org/tools/blast/match/show?blast_db_id=71;id=Mvestita_C856g22136-RA;hilite_coords=66-459) | 0 | Histone acetyltransferase | 0 |
| REV1 | ADC07240 | 0 | Aspi01Gene66531.t2 | 0 | Azfi-s0197.g057315 | 0 | Ceric.1Z081200.9.v2.1 | 0 | Mvestita_S6g00169-RA | 0 | Sacu_v1.1_s0015.g006707 | 0 |
| CSA1 | ADC05917 | 1.85·10^-174^ | Aspi01Gene04195.t1 | 3.14·10^-91^ | Azfi-s0002.g001083 | 2.98·10^-157^ | Ceric.16G038600.3.v2.1 | 0 | Mvestita_S10g02859-RA | 3.39·10^-168^ | Sacu_v1.1_s0043.g012914 | 6.61·10^-153^ |
| UVR8 | [ADC24639](https://fernbase.org/tools/blast/match/show?blast_db_id=59;id=ADC24639;hilite_coords=1-453) | 0 | [Aspi01Gene24339.t2](https://fernbase.org/tools/blast/match/show?blast_db_id=63;id=Aspi01Gene24339.t2;hilite_coords=1-453) | 0 | Regulator of chromosome condensation (RCC1) family protein | 0 | [Ceric.38G035100.1.v2.1](https://fernbase.org/tools/blast/match/show?blast_db_id=67;id=Ceric.38G035100.1.v2.1;hilite_coords=1-454) | 0 | [Mvestita_S7g18127-RA](https://fernbase.org/tools/blast/match/show?blast_db_id=71;id=Mvestita_S7g18127-RA;hilite_coords=14-450) | 0 | Regulator of chromosome condensation (RCC1) family protein | 0 |
| RUS1 | ADC10496 | 0 | Aspi01Gene68596.t1 | 2.02·10^-135^ | Azfi-s0001.g000057 | 0 | Ceric.11G021800.1.v2.1 | 0 | Mvestita_S2g11812-RA | 0 | Sacu_v1.1_s0040.g012276 | 0 |
| RUS2 | ADC12874 | 0 | Aspi01Gene01109.t1 | 0 | Azfi-s0232.g059330 | 0 | Ceric.28G019600.1.v2.1 | 0 | Mvestita_S16g11234-RA | 0 | Sacu_v1.1_s0001.g000678 | 0 |
| CSN1 | ADC12973 | 0 | Aspi01Gene01185.t1 | 0 | Azfi-s0102.g044475 | 0 | Ceric.22G023200.1.v2.1 | 0 | Mvestita_S10g03007-RA | 0 | Sacu_v1.1_s0136.g022425 | 0 |
| CSN2 | ADC07771 | 0 | Aspi01Gene52642.t2 | 0 | Azfi-s2058.g108930 | 0 | Ceric.33G045900.1.v2.1 | 0 | Mvestita_S17g16200-RA | 0 | Sacu_v1.1_s0146.g023111 | 0 |
| CSN6A | ADC23703 | 0 | Aspi01Gene11611.t1 | 4.7·10^-90^ | Azfi-s0026.g023426 | 2.28·10^-142^ | Ceric.02G112400.2.v2.1 | 0 | Mvestita_S8g04782-RA | 3.1·10^-171^ | Sacu_v1.1_s0006.g003047 | 5.94·10^-143^ |
| DDB1A | ADC13804 | 0 | Aspi01Gene33457.t1 | 0 | Azfi-s0382.g067410 | 0 | Ceric.30G020500.1.v2.1 | 0 | Mvestita_S9g06276-RA | 0 | Sacu_v1.1_s0201.g025613 | 0 |
| EMB168 | ADC13131 | 0 | Aspi01Gene04009.t1 | 0 | Azfi-s0013.g013098 | 0 | Ceric.22G021600.1.v2.1 | 0 | Mvestita_S9g05972-RA | 0 | Sacu_v1.1_s0034.g011295 | 0 |
| SPA3 | ADC20024 | 0 | Aspi01Gene13976.t2 | 0 | Azfi-s0004.g008529 | 0 | Ceric.23G014900.3.v2.1 | 0 | Mvestita_S1g06894-RA | 0 | Sacu_v1.1_s0008.g003993 | 0 |
| JK218 | ADC19278 | 0 | Aspi01Gene09339.t1 | 0 | Azfi-s0088.g042460 | 2.13·10^-150^ | Ceric.02G094700.2.v2.1 | 0 | Mvestita_S3g03505-RA | 0 | Sacu_v1.1_s0070.g016653 | 1.2·10^-173^ |
| SGR1 | ADC16159 | 0 | Aspi01Gene47444.t1 | 0 | Azfi-s0807.g087717 | 1.13·10^-161^ | Ceric.13G041400.1.v2.1 | 0 | Mvestita_S10g02236-RA | 0 | Sacu_v1.1_s0014.g006404 | 0 |
| SGR2 | ADC14628 | 2.24·10^-86^ | Aspi01Gene63751.t2 | 0 | Azfi-s0103.g044625 | 0 | Ceric.30G063100.1.v2.1 | 0 | Mvestita_S5g09872-RA | 0 | Sacu_v1.1_s0006.g003290 | 0 |
| ABCB1 | [ADC16643](https://fernbase.org/tools/blast/match/show?blast_db_id=59;id=ADC16643;hilite_coords=149-1250) | 0 | [Aspi01Gene14156.t1](https://fernbase.org/tools/blast/match/show?blast_db_id=63;id=Aspi01Gene14156.t1;hilite_coords=159-1205) | 0 | ABC transporter B family member 15 | 0 | [Ceric.06G076400.3.v2.1](https://fernbase.org/tools/blast/match/show?blast_db_id=67;id=Ceric.06G076400.3.v2.1;hilite_coords=44-1150) | 0 | [Mvestita_S4g09028-RA](https://fernbase.org/tools/blast/match/show?blast_db_id=71;id=Mvestita_S4g09028-RA;hilite_coords=84-1189) | 0 | ABC transporter B family member 15 | 0 |
| CRM1 | [ADC01727](https://fernbase.org/tools/blast/match/show?blast_db_id=59;id=ADC01727;hilite_coords=1-5123) | 0 | [Aspi01Gene46863.t1](https://fernbase.org/tools/blast/match/show?blast_db_id=63;id=Aspi01Gene46863.t1;hilite_coords=70-4343) | 0 | Auxin transport protein BIG | 0 | [Ceric.14G095600.1.v2.1](https://fernbase.org/tools/blast/match/show?blast_db_id=67;id=Ceric.14G095600.1.v2.1;hilite_coords=1-5147) | 0 | [Mvestita_S10g02311-RA](https://fernbase.org/tools/blast/match/show?blast_db_id=71;id=Mvestita_S10g02311-RA;hilite_coords=78-5126) | 0 | Auxin transport protein BIG | 0 |
| AUX1 | [ADC12642](https://fernbase.org/tools/blast/match/show?blast_db_id=59;id=ADC12642;hilite_coords=11-472) | 0 | [Aspi01Gene19841.t1](https://fernbase.org/tools/blast/match/show?blast_db_id=63;id=Aspi01Gene19841.t1;hilite_coords=1-466) | 0 | Auxin transporter-like protein 2 | 0 | [Ceric.24G007000.1.v2.1](https://fernbase.org/tools/blast/match/show?blast_db_id=67;id=Ceric.24G007000.1.v2.1;hilite_coords=9-468) | 0 | [Mvestita_S1g06384-RA](https://fernbase.org/tools/blast/match/show?blast_db_id=71;id=Mvestita_S1g06384-RA;hilite_coords=2-470) | 0 | Auxin transporter-like protein 2 | 0 |
| ABCG40 | [ADC02115](https://fernbase.org/tools/blast/match/show?blast_db_id=59;id=ADC02115;hilite_coords=867-1227) | 0 | [Aspi01Gene64035.t1](https://fernbase.org/tools/blast/match/show?blast_db_id=63;id=Aspi01Gene64035.t1;hilite_coords=859-1220) | 0 | ABC transporter family protein | 0 | [Ceric.19G065600.5.v2.1](https://fernbase.org/tools/blast/match/show?blast_db_id=67;id=Ceric.19G065600.5.v2.1;hilite_coords=863-1224) | 0 | [Mvestita_C89g20687-RA](https://fernbase.org/tools/blast/match/show?blast_db_id=71;id=Mvestita_C89g20687-RA;hilite_coords=856-1216) | 0 | ABC transporter family protein | 0 |
| ABCD1 | ADC11794 | 0 | Aspi01Gene50290.t1 | 0 | Azfi-s0048.g030599 | 0 | Ceric.01G050100.1.v2.1 | 0 | Mvestita_S9g05930-RA | 0 | Sacu_v1.1_s0138.g022570 | 0 |
| ABCC5 | [ADC27606](https://fernbase.org/tools/blast/match/show?blast_db_id=59;id=ADC27606;hilite_coords=1-1506) | 0 | [Aspi01Gene09680.t1](https://fernbase.org/tools/blast/match/show?blast_db_id=63;id=Aspi01Gene09680.t1;hilite_coords=7-1524) | 0 | ABC transporter family protein | 0 | [Ceric.02G017300.3.v2.1](https://fernbase.org/tools/blast/match/show?blast_db_id=67;id=Ceric.02G017300.3.v2.1;hilite_coords=1-1485) | 0 | [Mvestita_S16g10768-RA](https://fernbase.org/tools/blast/match/show?blast_db_id=71;id=Mvestita_S16g10768-RA;hilite_coords=4-1486) | 0 | ABC transporter family protein | 0 |
| CNGC2 | [ADC21625](https://fernbase.org/tools/blast/match/show?blast_db_id=59;id=ADC21625;hilite_coords=3-715) | 0 | [Aspi01Gene71929.t1](https://fernbase.org/tools/blast/match/show?blast_db_id=63;id=Aspi01Gene71929.t1;hilite_coords=47-746) | 0 | Cyclic nucleotide-gated channel 2-like | 0 | [Ceric.09G023200.3.v2.1](https://fernbase.org/tools/blast/match/show?blast_db_id=67;id=Ceric.09G023200.3.v2.1;hilite_coords=11-716) | 0 | [Mvestita_S19g17296-RA](https://fernbase.org/tools/blast/match/show?blast_db_id=71;id=Mvestita_S19g17296-RA;hilite_coords=71-735) | 0 | Cyclic nucleotide-gated channel 2-like | 0 |
| DND2 | [ADC21625](https://fernbase.org/tools/blast/match/show?blast_db_id=59;id=ADC21625;hilite_coords=224-715) | 0 | [Aspi01Gene71929.t1](https://fernbase.org/tools/blast/match/show?blast_db_id=63;id=Aspi01Gene71929.t1;hilite_coords=261-746) | 0 | Cyclic nucleotide-gated channel 2-like | 0 | [Ceric.09G023200.3.v2.1](https://fernbase.org/tools/blast/match/show?blast_db_id=67;id=Ceric.09G023200.3.v2.1;hilite_coords=225-716) | 0 | [Mvestita_S19g17296-RA](https://fernbase.org/tools/blast/match/show?blast_db_id=71;id=Mvestita_S19g17296-RA;hilite_coords=261-735) | 0 | Cyclic nucleotide-gated channel 2-like | 0 |
| VDAC1 | [ADC04384](https://fernbase.org/tools/blast/match/show?blast_db_id=59;id=ADC04384;hilite_coords=1-276) | 2·10^-135^ | [Aspi01Gene61361.t1](https://fernbase.org/tools/blast/match/show?blast_db_id=63;id=Aspi01Gene61361.t1;hilite_coords=90-369) | 2·10^-159^ | Mitochondrial outer membrane protein porin 1 | 6·10^-136^ | [Ceric.14G020600.1.v2.1](https://fernbase.org/tools/blast/match/show?blast_db_id=67;id=Ceric.14G020600.1.v2.1;hilite_coords=1-276) | 2·10^-146^ | [Mvestita_S11g13705-RA](https://fernbase.org/tools/blast/match/show?blast_db_id=71;id=Mvestita_S11g13705-RA;hilite_coords=1-276) | 1·10^-138^ | Mitochondrial outer membrane protein porin 1 | 4·10^-111^ |
| MSL1 | ADC13954 | 2.23·10^-155^ | Aspi01Gene10781.t1 | 4.71·10^-78^ | Azfi-s0051.g031345 | 6.42·10^-178^ | Ceric.31G030700.1.v2.1 | 0 | Mvestita_S12g15967-RA | 0 | Sacu_v1.1_s0020.g008025 | 8.19·10^-178^ |
| NRAMP4 | [ADC04754](https://fernbase.org/tools/blast/match/show?blast_db_id=59;id=ADC04754;hilite_coords=1-506) | 0 | [Aspi01Gene39825.t1](https://fernbase.org/tools/blast/match/show?blast_db_id=63;id=Aspi01Gene39825.t1;hilite_coords=14-455) | 0 | Metal transporter | 0 | [Ceric.22G080000.1.v2.1](https://fernbase.org/tools/blast/match/show?blast_db_id=67;id=Ceric.22G080000.1.v2.1;hilite_coords=3-506) | 0 | [Mvestita_S9g05861-RA](https://fernbase.org/tools/blast/match/show?blast_db_id=71;id=Mvestita_S9g05861-RA;hilite_coords=9-500) | 0 | Metal transporter | 5·10^-95^ |
| BOR1 | ADC23661 | 0 | Aspi01Gene46823.t1 | 0 | Azfi-s0030.g024428 | 0 | Ceric.29G013800.3.v2.1 | 0 | Mvestita_S3g03932-RA | 0 | Sacu_v1.1_s0074.g017183 | 0 |
| APC2 | ADC29008 | 0 | Aspi01Gene15715.t1 | 0 | Azfi-s0034.g025432 | 0 | Ceric.39G024700.9.v2.1 | 0 | Mvestita_S9g06250-RA | 0 | Sacu_v1.1_s0058.g014909 | 0 |
| CHX17 | ADC08150 | 0 | Aspi01Gene31851.t1 | 0 | Azfi-s0006.g009548 | 0 | Ceric.1Z070100.1.v2.1 | 0 | Mvestita_S15g12932-RA | 0 | Sacu_v1.1_s0003.g001982 | 0 |
| HMP51 | ADC02959 | 0 | Aspi01Gene32243.t1 | 0 | Azfi-s0514.g074936 | 0 | Ceric.04G057700.1.v2.1 | 0 | Mvestita_S6g01115-RA | 0 | Sacu_v1.1_s0039.g012091 | 0 |
| PIP1;2 | ADC22680 | 0 | Aspi01Gene24709.t1 | 0 | Azfi-s0007.g011084 | 0 | Ceric.25G031200.2.v2.1 | 0 | Mvestita_S15g12740-RA | 0 | Sacu_v1.1_s0064.g015869 | 6.31·10^-174^ |
| PIP2;1 | ADC17985 | 0 | Aspi01Gene65236.t1 | 6.56·10^-170^ | Azfi-s0020.g015473 | 3.08·10^-164^ | Ceric.05G082100.1.v2.1 | 9.05·10^-172^ | Mvestita_S5g10093-RA | 1.47·10^-168^ | Sacu_v1.1_s0006.g002971 | 1.05·10^-159^ |
| MIZ2 | [ADC10687](https://fernbase.org/tools/blast/match/show?blast_db_id=59;id=ADC10687;hilite_coords=1-1438) | 0 | [Aspi01Gene41929.t1](https://fernbase.org/tools/blast/match/show?blast_db_id=63;id=Aspi01Gene41929.t1;hilite_coords=1-1457) | 0 | SEC7-like guanine nucleotide exchange family protein | 0 | [Ceric.02G006600.1.v2.1](https://fernbase.org/tools/blast/match/show?blast_db_id=67;id=Ceric.02G006600.1.v2.1;hilite_coords=1-1407) | 0 | [Mvestita_S6g00831-RA](https://fernbase.org/tools/blast/match/show?blast_db_id=71;id=Mvestita_S6g00831-RA;hilite_coords=1-1443) | 0 | SEC7-like guanine nucleotide exchange family protein | 0 |
| COG8 | [ADC18598](https://fernbase.org/tools/blast/match/show?blast_db_id=59;id=ADC18598;hilite_coords=1-580) | 0 | [Aspi01Gene08972.t1](https://fernbase.org/tools/blast/match/show?blast_db_id=63;id=Aspi01Gene08972.t1;hilite_coords=84-639) | 0 | Conserved oligomeric Golgi complex component-related protein | 0 | [Ceric.05G079700.1.v2.1](https://fernbase.org/tools/blast/match/show?blast_db_id=67;id=Ceric.05G079700.1.v2.1;hilite_coords=1-546) | 0 | [Mvestita_S14g08035-RA](https://fernbase.org/tools/blast/match/show?blast_db_id=71;id=Mvestita_S14g08035-RA;hilite_coords=13-557) | 0 | Conserved oligomeric Golgi complex component-related protein | 0 |
| **Protein** | ***Adiantum capillus* protein** | **E-value** | ***Alsophila spinulosa* protein** | **E-value** | ***Azolla filiculoides* protein** | **E-value** | ***Ceratopteris richardii* protein** | **E-value** | ***Marsilea vestita* protein** | **E-value** | ***Salvinia cucullata* protein** | **E-value** |
| TRAPPC9 | [ADC16438](https://fernbase.org/tools/blast/match/show?blast_db_id=59;id=ADC16438;hilite_coords=416-1185) | 0 | [Aspi01Gene37553.t1](https://fernbase.org/tools/blast/match/show?blast_db_id=63;id=Aspi01Gene37553.t1;hilite_coords=92-857) | 0 | Trafficking particle complex subunit 9 | 0 | [Ceric.18G035400.1.v2.1](https://fernbase.org/tools/blast/match/show?blast_db_id=67;id=Ceric.18G035400.1.v2.1;hilite_coords=432-1199) | 0 | [Mvestita_S8g04826-RA](https://fernbase.org/tools/blast/match/show?blast_db_id=71;id=Mvestita_S8g04826-RA;hilite_coords=427-1156) | 0 | Trafficking particle complex subunit 9 | 0 |
| VPS53 | [ADC11538](https://fernbase.org/tools/blast/match/show?blast_db_id=59;id=ADC11538;hilite_coords=213-749) | 0 | [Aspi01Gene55502.t1](https://fernbase.org/tools/blast/match/show?blast_db_id=63;id=Aspi01Gene55502.t1;hilite_coords=120-368) | 6·10^-114^ | Membrane trafficking VPS53 family protein | 0 | [Ceric.34G024900.7.v2.1](https://fernbase.org/tools/blast/match/show?blast_db_id=67;id=Ceric.34G024900.7.v2.1;hilite_coords=236-747) | 0 | [Mvestita_S10g02737-RA](https://fernbase.org/tools/blast/match/show?blast_db_id=71;id=Mvestita_S10g02737-RA;hilite_coords=244-716) | 0 | Membrane trafficking VPS53 family protein | 0 |
| MAG5 | ADC13045 | 0 | Aspi01Gene00895.t1 | 0 | Azfi-s0781.g086689 | 0 | Ceric.11G037700.2.v2.1 | 0 | Mvestita_S16g10790-RA | 0 | Sacu_v1.1_s0034.g011231 | 0 |
| SEC23G | ADC24704 | 0 | Aspi01Gene59433.t1 | 0 | Azfi-s0288.g063252 | 0 | Ceric.37G050300.1.v2.1 | 0 | Mvestita_S14g07677-RA | 0 | Sacu_v1.1_s0037.g011867 | 0 |
| NUP155 | [ADC17333](https://fernbase.org/tools/blast/match/show?blast_db_id=59;id=ADC17333;hilite_coords=1-1471) | 0 | [Aspi01Gene68733.t1](https://fernbase.org/tools/blast/match/show?blast_db_id=63;id=Aspi01Gene68733.t1;hilite_coords=1-934) | 0 | Nuclear pore complex protein Nup155 | 0 | [Ceric.28G038900.1.v2.1](https://fernbase.org/tools/blast/match/show?blast_db_id=67;id=Ceric.28G038900.1.v2.1;hilite_coords=1-1473) | 0 | [Mvestita_S2g11990-RA](https://fernbase.org/tools/blast/match/show?blast_db_id=71;id=Mvestita_S2g11990-RA;hilite_coords=1-1462) | 0 | Nuclear pore complex protein Nup155 | 0 |
| NUP160 | [ADC06993](https://fernbase.org/tools/blast/match/show?blast_db_id=59;id=ADC06993;hilite_coords=19-1362) | 0 | [Aspi01Gene23935.t1](https://fernbase.org/tools/blast/match/show?blast_db_id=63;id=Aspi01Gene23935.t1;hilite_coords=548-1650) | 0 | Nuclear pore complex protein Nup160 | 0 | [Ceric.32G022400.2.v2.1](https://fernbase.org/tools/blast/match/show?blast_db_id=67;id=Ceric.32G022400.2.v2.1;hilite_coords=1-1477) | 0 | [Mvestita_S7g18016-RA](https://fernbase.org/tools/blast/match/show?blast_db_id=71;id=Mvestita_S7g18016-RA;hilite_coords=8-1437) | 0 | Nuclear pore complex protein Nup160 | 0 |
| TIC100 | [ADC20479](https://fernbase.org/tools/blast/match/show?blast_db_id=59;id=ADC20479;hilite_coords=90-744) | 0 | [Aspi01Gene10078.t1](https://fernbase.org/tools/blast/match/show?blast_db_id=63;id=Aspi01Gene10078.t1;hilite_coords=1-582) | 0 | MORN (Membrane Occupation and Recognition Nexus) repeat-containing protein | 0 | [Ceric.28G005300.3.v2.1](https://fernbase.org/tools/blast/match/show?blast_db_id=67;id=Ceric.28G005300.3.v2.1;hilite_coords=83-863) | 0 | [Mvestita_S16g11000-RA](https://fernbase.org/tools/blast/match/show?blast_db_id=71;id=Mvestita_S16g11000-RA;hilite_coords=78-785) | 0 | MORN (Membrane Occupation and Recognition Nexus) repeat-containing protein | 0 |
| TIC55 | ADC02252 | 3.18·10^-156^ | Aspi01Gene65972.t1 | 8.38·10^-39^ | Azfi-s0065.g035849 | 6.03·10^-43^ | Ceric.30G058900.1.v2.1 | 0 | Mvestita_S5g10126-RA | 0 | Sacu_v1.1_s0069.g016507 | 1·10^-30^ |
| TIC56 | ADC17185 | 0 | Aspi01Gene55665.t1 | 0 | Azfi-s0018.g014910 | 1.17·10^-104^ | Ceric.26G055700.1.v2.1 | 0 | Mvestita_S13g02080-RA | 0 | Sacu_v1.1_s0037.g011825 | 0 |
| TOC75-3 | ADC05958 | 0 | Aspi01Gene04281.t1 | 0 | Azfi-s0013.g013142 | 0 | Ceric.11G026700.2.v2.1 | 0 | Mvestita_S11g13586-RA | 0 | Sacu_v1.1_s0044.g013034 | 0 |
| AAP2 | [ADC13641](https://fernbase.org/tools/blast/match/show?blast_db_id=59;id=ADC13641;hilite_coords=128-494) | 0 | [Aspi01Gene07906.t1](https://fernbase.org/tools/blast/match/show?blast_db_id=63;id=Aspi01Gene07906.t1;hilite_coords=131-497) | 0 | Amino acid permease 3 | 3·10^-158^ | [Ceric.17G072500.1.v2.1](https://fernbase.org/tools/blast/match/show?blast_db_id=67;id=Ceric.17G072500.1.v2.1;hilite_coords=117-483) | 0 | [Mvestita_S20g19030-RA](https://fernbase.org/tools/blast/match/show?blast_db_id=71;id=Mvestita_S20g19030-RA;hilite_coords=112-478) | 0 | Amino acid permease 3 | 0 |
| AAP3 | [ADC00732](https://fernbase.org/tools/blast/match/show?blast_db_id=59;id=ADC00732;hilite_coords=1-290) | 6·10^-168^ | [Aspi01Gene56576.t1](https://fernbase.org/tools/blast/match/show?blast_db_id=63;id=Aspi01Gene56576.t1;hilite_coords=58-345) | 2·10^-175^ | Amino acid permease 3 | 6·10^-155^ | [Ceric.39G036100.5.v2.1](https://fernbase.org/tools/blast/match/show?blast_db_id=67;id=Ceric.39G036100.5.v2.1;hilite_coords=1-287) | 1·10^-154^ | [Mvestita_S16g11299-RA](https://fernbase.org/tools/blast/match/show?blast_db_id=71;id=Mvestita_S16g11299-RA;hilite_coords=14-292) | 3·10^-139^ | Amino acid permease 3 | 8·10^-121^ |
| LHT1 | [ADC12312](https://fernbase.org/tools/blast/match/show?blast_db_id=59;id=ADC12312;hilite_coords=54-389) | 0 | [Aspi01Gene59630.t1](https://fernbase.org/tools/blast/match/show?blast_db_id=63;id=Aspi01Gene59630.t1;hilite_coords=6-333) | 4·10^-175^ | Histidine amino acid transporter | 3·10^-115^ | [Ceric.1Z032700.1.v2.1](https://fernbase.org/tools/blast/match/show?blast_db_id=67;id=Ceric.1Z032700.1.v2.1;hilite_coords=12-335) | 0 | [Mvestita_S10g02904-RA](https://fernbase.org/tools/blast/match/show?blast_db_id=71;id=Mvestita_S10g02904-RA;hilite_coords=2-313) | 1·10^-172^ | Amino acid transporter | 4·10^-160^ |
| CAT1 | ADC07786 | 0 | Aspi01Gene52622.t1 | 0 | Azfi-s0093.g043307 | 0 | Ceric.29G039300.1.v2.1 | 0 | Mvestita_S5g10175-RA | 0 | Sacu_v1.1_s0033.g011077 | 0 |
| NPF8.2 | [ADC05540](https://fernbase.org/tools/blast/match/show?blast_db_id=59;id=ADC05540;hilite_coords=1-1039) | 0 | [Aspi01Gene48172.t1](https://fernbase.org/tools/blast/match/show?blast_db_id=63;id=Aspi01Gene48172.t1;hilite_coords=1-1066) | 0 | [Azfi-s0329.g065351](https://fernbase.org/tools/blast/match/show?blast_db_id=36;id=Azfi_s0329.g065351;hilite_coords=1-755) | 0 | [Ceric.01G007300.3.v2.1](https://fernbase.org/tools/blast/match/show?blast_db_id=67;id=Ceric.01G007300.3.v2.1;hilite_coords=1-763) | 0 | [Mvestita_S2g11367-RA](https://fernbase.org/tools/blast/match/show?blast_db_id=71;id=Mvestita_S2g11367-RA;hilite_coords=1-899) | 0 | [Sacu_v1.1_s0017.g007238](https://fernbase.org/tools/blast/match/show?blast_db_id=40;id=Sacu_v1.1_s0017.g007238;hilite_coords=1-762) | 0 |
| ADNT1 | [ADC20972](https://fernbase.org/tools/blast/match/show?blast_db_id=59;id=ADC20972;hilite_coords=54-434) | 0 | [Aspi01Gene32954.t1](https://fernbase.org/tools/blast/match/show?blast_db_id=63;id=Aspi01Gene32954.t1;hilite_coords=1-391) | 0 | Mitochondrial substrate carrier family protein | 0 | [Ceric.05G062200.2.v2.1](https://fernbase.org/tools/blast/match/show?blast_db_id=67;id=Ceric.05G062200.2.v2.1;hilite_coords=24-380) | 0 | [Mvestita_C66g20194-RA](https://fernbase.org/tools/blast/match/show?blast_db_id=71;id=Mvestita_C66g20194-RA;hilite_coords=1-393) | 0 | Mitochondrial substrate carrier family protein | 2·10^-139^ |
| MPT1 | [ADC23011](https://fernbase.org/tools/blast/match/show?blast_db_id=59;id=ADC23011;hilite_coords=79-401) | 0 | [Aspi01Gene37727.t1](https://fernbase.org/tools/blast/match/show?blast_db_id=63;id=Aspi01Gene37727.t1;hilite_coords=72-364) | 2·10^-115^ | Mitochondrial phosphate carrier protein | 2·10^-174^ | [Ceric.39G017000.1.v2.1](https://fernbase.org/tools/blast/match/show?blast_db_id=67;id=Ceric.39G017000.1.v2.1;hilite_coords=8-330) | 0 | [Mvestita_S2g12305-RA](https://fernbase.org/tools/blast/match/show?blast_db_id=71;id=Mvestita_S2g12305-RA;hilite_coords=10-325) | 7·10^-172^ | Mitochondrial phosphate carrier protein | 5·10^-174^ |
| AMT1 | ADC15977 | 0 | Aspi01Gene15779.t1 | 0 | Azfi-s0034.g025388 | 0 | Ceric.39G029000.1.v2.1 | 0 | Mvestita_S18g14283-RA | 8.01·10^-157^ | Sacu_v1.1_s0018.g007623 | 0 |
| AMT2 | [ADC00236](https://fernbase.org/tools/blast/match/show?blast_db_id=59;id=ADC00236;hilite_coords=18-455) | 3·10^-143^ | [Aspi01Gene33315.t1](https://fernbase.org/tools/blast/match/show?blast_db_id=63;id=Aspi01Gene33315.t1;hilite_coords=18-442) | 1·10^-141^ | Ammonium transporter | 1·10^-133^ | [Ceric.04G027400.3.v2.1](https://fernbase.org/tools/blast/match/show?blast_db_id=67;id=Ceric.04G027400.3.v2.1;hilite_coords=15-465) | 5·10^-144^ | [Mvestita_S11g14249-RA](https://fernbase.org/tools/blast/match/show?blast_db_id=71;id=Mvestita_S11g14249-RA;hilite_coords=6-426) | 1·10^-140^ | Ammonium transporter | 3·10^-140^ |
| PLUTO | [ADC16371](https://fernbase.org/tools/blast/match/show?blast_db_id=59;id=ADC16371;hilite_coords=10-334) | 5·10^-157^ | [Aspi01Gene28634.t1](https://fernbase.org/tools/blast/match/show?blast_db_id=63;id=Aspi01Gene28634.t1;hilite_coords=22-491) | 3·10^-172^ | Permease, cytosine/purines, uracil, thiamine, allantoin family protein | 0 | [Ceric.07G045000.1.v2.1](https://fernbase.org/tools/blast/match/show?blast_db_id=67;id=Ceric.07G045000.1.v2.1;hilite_coords=26-566) | 0 | [Mvestita_S1g07378-RA](https://fernbase.org/tools/blast/match/show?blast_db_id=71;id=Mvestita_S1g07378-RA;hilite_coords=85-545) | 0 | Permease, cytosine/purines, uracil, thiamine, allantoin family protein | 0 |
| MATE | [ADC18100](https://fernbase.org/tools/blast/match/show?blast_db_id=59;id=ADC18100;hilite_coords=24-479) | 0 | [Aspi01Gene64885.t1](https://fernbase.org/tools/blast/match/show?blast_db_id=63;id=Aspi01Gene64885.t1;hilite_coords=7-523) | 0 | Protein DETOXIFICATION | 0 | [Ceric.1Z024500.3.v2.1](https://fernbase.org/tools/blast/match/show?blast_db_id=67;id=Ceric.1Z024500.3.v2.1;hilite_coords=22-547) | 0 | [Mvestita_S18g14746-RA](https://fernbase.org/tools/blast/match/show?blast_db_id=71;id=Mvestita_S18g14746-RA;hilite_coords=25-514) | 0 | Protein DETOXIFICATION | 0 |
| MGT1 | [ADC08862](https://fernbase.org/tools/blast/match/show?blast_db_id=59;id=ADC08862;hilite_coords=22-477) | 9·10^-94^ | [Aspi01Gene52438.t2](https://fernbase.org/tools/blast/match/show?blast_db_id=63;id=Aspi01Gene52438.t2;hilite_coords=28-298) | 1·10^-119^ | Magnesium transporter MRS2-1 | 1·10^-172^ | [Ceric.17G066000.4.v2.1](https://fernbase.org/tools/blast/match/show?blast_db_id=67;id=Ceric.17G066000.4.v2.1;hilite_coords=1-456) | 0 | [Mvestita_S20g19218-RA](https://fernbase.org/tools/blast/match/show?blast_db_id=71;id=Mvestita_S20g19218-RA;hilite_coords=25-456) | 9·10^-177^ | Magnesium transporter MRS2-1 | 5·10^-180^ |
| UTR3 | [ADC08304](https://fernbase.org/tools/blast/match/show?blast_db_id=59;id=ADC08304;hilite_coords=4-324) | 0 | [Aspi01Gene38247.t1](https://fernbase.org/tools/blast/match/show?blast_db_id=63;id=Aspi01Gene38247.t1;hilite_coords=1-329) | 0 | UDP-galactose/UDP-glucose transporter 3 | 0 | [Ceric.05G068600.1.v2.1](https://fernbase.org/tools/blast/match/show?blast_db_id=67;id=Ceric.05G068600.1.v2.1;hilite_coords=6-324) | 0 | [Mvestita_S16g11339-RA](https://fernbase.org/tools/blast/match/show?blast_db_id=71;id=Mvestita_S16g11339-RA;hilite_coords=1-328) | 0 | UDP-galactose transporter 1 | 4·10^-34^ |
| TST2 | [ADC01767](https://fernbase.org/tools/blast/match/show?blast_db_id=59;id=ADC01767;hilite_coords=5-656) | 0 | [Aspi01Gene13510.t1](https://fernbase.org/tools/blast/match/show?blast_db_id=63;id=Aspi01Gene13510.t1;hilite_coords=1-769) | 0 | Tonoplast monosaccharide transporter 2 | 0 | [Ceric.14G096100.1.v2.1](https://fernbase.org/tools/blast/match/show?blast_db_id=67;id=Ceric.14G096100.1.v2.1;hilite_coords=1-761) | 0 | [Mvestita_S8g04360-RA](https://fernbase.org/tools/blast/match/show?blast_db_id=71;id=Mvestita_S8g04360-RA;hilite_coords=8-740) | 0 | Tonoplast monosaccharide transporter 2 | 0 |
| SUC4 | [ADC18527](https://fernbase.org/tools/blast/match/show?blast_db_id=59;id=ADC18527;hilite_coords=94-574) | 0 | [Aspi01Gene39806.t1](https://fernbase.org/tools/blast/match/show?blast_db_id=63;id=Aspi01Gene39806.t1;hilite_coords=21-497) | 0 | Sucrose transporter | 0 | [Ceric.21G082000.2.v2.1](https://fernbase.org/tools/blast/match/show?blast_db_id=67;id=Ceric.21G082000.2.v2.1;hilite_coords=22-487) | 0 | [Mvestita_S6g00487-RA](https://fernbase.org/tools/blast/match/show?blast_db_id=71;id=Mvestita_S6g00487-RA;hilite_coords=1-493) | 0 | Sucrose transporter | 0 |
| SR1 | [ADC22781](https://fernbase.org/tools/blast/match/show?blast_db_id=59;id=ADC22781;hilite_coords=1-1139) | 0 | [Aspi01Gene48770.t1](https://fernbase.org/tools/blast/match/show?blast_db_id=63;id=Aspi01Gene48770.t1;hilite_coords=1-1138) | 0 | Calmodulin-binding transcription activator 3 | 0 | [Ceric.31G070500.1.v2.1](https://fernbase.org/tools/blast/match/show?blast_db_id=67;id=Ceric.31G070500.1.v2.1;hilite_coords=1-1127) | 0 | [Mvestita_S6g00432-RA](https://fernbase.org/tools/blast/match/show?blast_db_id=71;id=Mvestita_S6g00432-RA;hilite_coords=1-1122) | 0 | Calmodulin-binding transcription activator 3 | 0 |
| ACER | [ADC25736](https://fernbase.org/tools/blast/match/show?blast_db_id=59;id=ADC25736;hilite_coords=1-246) | 2·10^-151^ | [Aspi01Gene58567.t1](https://fernbase.org/tools/blast/match/show?blast_db_id=63;id=Aspi01Gene58567.t1;hilite_coords=1-246) | 2·10^-164^ | Alkaline ceramidase 3 | 7·10^-147^ | [Ceric.29G067700.3.v2.1](https://fernbase.org/tools/blast/match/show?blast_db_id=67;id=Ceric.29G067700.3.v2.1;hilite_coords=1-244) | 1·10^-129^ | [Mvestita_S3g03474-RA](https://fernbase.org/tools/blast/match/show?blast_db_id=71;id=Mvestita_S3g03474-RA;hilite_coords=1-245) | 6·10^-147^ | Alkaline ceramidase 3 | 3·10^-35^ |
| ALD1 | [ADC13213](https://fernbase.org/tools/blast/match/show?blast_db_id=59;id=ADC13213;hilite_coords=27-303) | 3·10^-136^ | [Aspi01Gene56657.t1](https://fernbase.org/tools/blast/match/show?blast_db_id=63;id=Aspi01Gene56657.t1;hilite_coords=78-470) | 2·10^-126^ | LL-diaminopimelate aminotransferase | 1·10^-112^ | [Ceric.11G029400.2.v2.1](https://fernbase.org/tools/blast/match/show?blast_db_id=67;id=Ceric.11G029400.2.v2.1;hilite_coords=27-304) | 2·10^-132^ | [Mvestita_S4g08704-RA](https://fernbase.org/tools/blast/match/show?blast_db_id=71;id=Mvestita_S4g08704-RA;hilite_coords=27-311) | 1·10^-132^ | LL-diaminopimelate aminotransferase | 1·10^-121^ |
| BGLU42 | [ADC13038](https://fernbase.org/tools/blast/match/show?blast_db_id=59;id=ADC13038;hilite_coords=1-547) | 0 | [Aspi01Gene28917.t1](https://fernbase.org/tools/blast/match/show?blast_db_id=63;id=Aspi01Gene28917.t1;hilite_coords=1-536) | 0 | β-glucosidase | 0 | [Ceric.10G035600.3.v2.1](https://fernbase.org/tools/blast/match/show?blast_db_id=67;id=Ceric.10G035600.3.v2.1;hilite_coords=1-509) | 0 | [Mvestita_S11g13552-RA](https://fernbase.org/tools/blast/match/show?blast_db_id=71;id=Mvestita_S11g13552-RA;hilite_coords=1-524) | 0 | β-glucosidase | 0 |
| LECRK-S.4 | ADC26389 | 0 | Aspi01Gene49703.t1 | 0 | Azfi-s0602.g078920 | 8.22·10^-101^ | Ceric.03G001400.1.v2.1 | 0 | Mvestita_S6g00927-RA | 0 | Sacu_v1.1_s0006.g003408 | 0 |
| AGB1 | ADC20921 | 0 | Aspi01Gene33094.t1 | 1.48·10^-167^ | Azfi-s0356.g066765 | 2.8·10^-174^ | Ceric.05G079300.4.v2.1 | 3.8·10^-173^ | Mvestita_S5g09728-RA | 1.1·10^-170^ | Sacu_v1.1_s0117.g021146 | 1.12·10^-166^ |
| HUB1 | [ADC03574](https://fernbase.org/tools/blast/match/show?blast_db_id=59;id=ADC03574;hilite_coords=1-876) | 0 | [Aspi01Gene28102.t1](https://fernbase.org/tools/blast/match/show?blast_db_id=63;id=Aspi01Gene28102.t1;hilite_coords=1-879) | 0 | E3 ubiquitin-protein ligase BRE1-like 2 | 0 | [Ceric.14G031800.2.v2.1](https://fernbase.org/tools/blast/match/show?blast_db_id=67;id=Ceric.14G031800.2.v2.1;hilite_coords=1-875) | 0 | [Mvestita_S10g02475-RA](https://fernbase.org/tools/blast/match/show?blast_db_id=71;id=Mvestita_S10g02475-RA;hilite_coords=28-899) | 0 | E3 ubiquitin-protein ligase BRE1-like 2 | 0 |
| ILK1 | [ADC20615](https://fernbase.org/tools/blast/match/show?blast_db_id=59;id=ADC20615;hilite_coords=1-287) | 0 | [Aspi01Gene17576.t1](https://fernbase.org/tools/blast/match/show?blast_db_id=63;id=Aspi01Gene17576.t1;hilite_coords=1-177) | 2·10^-88^ | Protein kinase | 0 | [Ceric.38G043200.1.v2.1](https://fernbase.org/tools/blast/match/show?blast_db_id=67;id=Ceric.38G043200.1.v2.1;hilite_coords=1-283) | 0 | [Mvestita_S17g16421-RA](https://fernbase.org/tools/blast/match/show?blast_db_id=71;id=Mvestita_S17g16421-RA;hilite_coords=3-283) | 0 | Protein kinase | 0 |
| FLS2 | ADC21071 | 0 | Aspi01Gene15273.t1 | 0 | Azfi-s0001.g000479 | 1.91·10^-104^ | Ceric.28G009600.3.v2.1 | 6.62·10^-123^ | Mvestita_S6g00270-RA | 1.94·10^-120^ | Sacu_v1.1_s0040.g012326 | 4.29·10^-115^ |
| WIN1 | ADC14659 | 0 | Aspi01Gene12187.t1 | 0 | Azfi-s0003.g007568 | 0 | Ceric.32G040400.1.v2.1 | 0 | Mvestita_S14g07827-RA | 0 | Sacu_v1.1_s0033.g011089 | 0 |
| WIN2 | ADC21942 | 9.49·10^-117^ | Aspi01Gene69065.t1 | 2.5·10^-123^ | Azfi-s0037.g025917 | 1.01·10^-116^ | Ceric.02G112900.2.v2.1 | 2.46·10^-113^ | Mvestita_S14g07614-RA | 4.86·10^-110^ | Sacu_v1.1_s0033.g011052 | 1.03·10^-119^ |
| LYK1 | ADC19183 | 0 | Aspi01Gene58625.t1 | 0 | Azfi-s0275.g061394 | 0 | Ceric.01G020400.1.v2.1 | 0 | Mvestita_S14g07891-RA | 0 | Sacu_v1.1_s0077.g017650 | 0 |
| **Protein** | ***Adiantum capillus* protein** | **E-value** | ***Alsophila spinulosa* protein** | **E-value** | ***Azolla filiculoides* protein** | **E-value** | ***Ceratopteris richardii* protein** | **E-value** | ***Marsilea vestita* protein** | **E-value** | ***Salvinia cucullata* protein** | **E-value** |
| RDR1 | [ADC29079](https://fernbase.org/tools/blast/match/show?blast_db_id=59;id=ADC29079;hilite_coords=17-1143) | 0 | [Aspi01Gene33143.t1](https://fernbase.org/tools/blast/match/show?blast_db_id=63;id=Aspi01Gene33143.t1;hilite_coords=87-1231) | 0 | [Azfi-s0003.g007701](https://fernbase.org/tools/blast/match/show?blast_db_id=36;id=Azfi_s0003.g007701;hilite_coords=43-1170) | 0 | [Ceric.03G095200.1.v2.1](https://fernbase.org/tools/blast/match/show?blast_db_id=67;id=Ceric.03G095200.1.v2.1;hilite_coords=2-1125) | 0 | [Mvestita_S17g16685-RA](https://fernbase.org/tools/blast/match/show?blast_db_id=71;id=Mvestita_S17g16685-RA;hilite_coords=38-1176) | 0 | RNA-dependent RNA polymerase | 0 |
| SGS3 | [ADC17603](https://fernbase.org/tools/blast/match/show?blast_db_id=59;id=ADC17603;hilite_coords=1-789) | 0 | [Aspi01Gene48646.t1](https://fernbase.org/tools/blast/match/show?blast_db_id=63;id=Aspi01Gene48646.t1;hilite_coords=151-478) | 1·10^-153^ | Protein SUPPRESSOR OF GENE SILENCING 3 | 2·10^-21^ | [Ceric.1Z005500.1.v2.1](https://fernbase.org/tools/blast/match/show?blast_db_id=67;id=Ceric.1Z005500.1.v2.1;hilite_coords=5-774) | 0 | [Mvestita_S5g09588-RA](https://fernbase.org/tools/blast/match/show?blast_db_id=71;id=Mvestita_S5g09588-RA;hilite_coords=9-774) | 0 | Protein SUPPRESSOR OF GENE SILENCING 3 | 0 |
| EMB2728 | [ADC13480](https://fernbase.org/tools/blast/match/show?blast_db_id=59;id=ADC13480;hilite_coords=60-321) | 1·10^-162^ | [Aspi01Gene18480.t1](https://fernbase.org/tools/blast/match/show?blast_db_id=63;id=Aspi01Gene18480.t1;hilite_coords=23-284) | 2·10^-161^ | Ribulose-phosphate 3-epimerase | 3·10^-158^ | [Ceric.30G051300.1.v2.1](https://fernbase.org/tools/blast/match/show?blast_db_id=67;id=Ceric.30G051300.1.v2.1;hilite_coords=20-282) | 2·10^-164^ | [Mvestita_S4g09197-RA](https://fernbase.org/tools/blast/match/show?blast_db_id=71;id=Mvestita_S4g09197-RA;hilite_coords=54-316) | 1·10^-159^ | Ribulose-phosphate 3-epimerase | 7·10^-163^ |
| BSK1 | [ADC01436](https://fernbase.org/tools/blast/match/show?blast_db_id=59;id=ADC01436;hilite_coords=1-372) | 0 | [Aspi01Gene60341.t1](https://fernbase.org/tools/blast/match/show?blast_db_id=63;id=Aspi01Gene60341.t1;hilite_coords=1-315) | 0 | Kinase family protein | 0 | [Ceric.29G078500.4.v2.1](https://fernbase.org/tools/blast/match/show?blast_db_id=67;id=Ceric.29G078500.4.v2.1;hilite_coords=1-371) | 0 | [Mvestita_S15g12679-RA](https://fernbase.org/tools/blast/match/show?blast_db_id=71;id=Mvestita_S15g12679-RA;hilite_coords=1-377) | 0 | Kinase family protein | 3·10^-178^ |
| NOXY7 | [ADC10159](https://fernbase.org/tools/blast/match/show?blast_db_id=59;id=ADC10159;hilite_coords=1-2610) | 0 | [Aspi01Gene37227.t1](https://fernbase.org/tools/blast/match/show?blast_db_id=63;id=Aspi01Gene37227.t1;hilite_coords=1-2611) | 0 | Translational activator GCN1 | 0 | [Ceric.34G010800.2.v2.1](https://fernbase.org/tools/blast/match/show?blast_db_id=67;id=Ceric.34G010800.2.v2.1;hilite_coords=1-2610) | 0 | [Mvestita_S10g02631-RA](https://fernbase.org/tools/blast/match/show?blast_db_id=71;id=Mvestita_S10g02631-RA;hilite_coords=12-2605) | 0 | Translational activator GCN1 | 0 |
| IRE1A | ADC06107 | 0 | Aspi01Gene29557.t1 | 0 | Azfi-s0092.g043029 | 0 | Ceric.20G085200.5.v2.1 | 0 | Mvestita_S12g15467-RA | 0 | Sacu_v1.1_s0083.g018160 | 0 |
| PUB59 | ADC17189 | 0 | Aspi01Gene51389.t1 | 0 | Azfi-s0019.g015186 | 0 | Ceric.06G040200.4.v2.1 | 0 | Mvestita_S8g04310-RA | 0 | Sacu_v1.1_s0777.g027689 | 0 |
| SCPR44 | ADC00795 | 0 | Aspi01Gene05033.t1 | 0 | Azfi-s0100.g044340 | 0 | Ceric.03G030600.1.v2.1 | 0 | Mvestita_S7g18163-RA | 0 | Sacu_v1.1_s0016.g006776 | 0 |
| APCB1 | ADC23306 | 0 | Aspi01Gene40899.t1 | 0 | Azfi-s0001.g000679 | 0 | Ceric.02G086000.11.v2.1 | 0 | Mvestita_S3g03361-RA | 0 | Sacu_v1.1_s0017.g007091 | 0 |
| AE3 | [ADC15992](https://fernbase.org/tools/blast/match/show?blast_db_id=59;id=ADC15992;hilite_coords=1-298) | 0 | [Aspi01Gene15797.t1](https://fernbase.org/tools/blast/match/show?blast_db_id=63;id=Aspi01Gene15797.t1;hilite_coords=1-207) | 2·10^-102^ | 26S proteasome non-ATPase regulatory subunit 7 family protein | 7·10^-169^ | [Ceric.31G022000.1.v2.1](https://fernbase.org/tools/blast/match/show?blast_db_id=67;id=Ceric.31G022000.1.v2.1;hilite_coords=1-298) | 0 | [Mvestita_S18g14654-RA](https://fernbase.org/tools/blast/match/show?blast_db_id=71;id=Mvestita_S18g14654-RA;hilite_coords=78-375) | 0 | 26S proteasome non-ATPase regulatory subunit 7 family protein | 0 |
| NAA20 | [ADC22914](https://fernbase.org/tools/blast/match/show?blast_db_id=59;id=ADC22914;hilite_coords=1-158) | 2·10^-109^ | [Aspi01Gene57123.t1](https://fernbase.org/tools/blast/match/show?blast_db_id=63;id=Aspi01Gene57123.t1;hilite_coords=1-174) | 6·10^-128^ | N-α-acetyltransferase 20 | 4·10^-124^ | [Ceric.29G026600.1.v2.1](https://fernbase.org/tools/blast/match/show?blast_db_id=67;id=Ceric.29G026600.1.v2.1;hilite_coords=1-171) | 4·10^-116^ | [Mvestita_S11g13513-RA](https://fernbase.org/tools/blast/match/show?blast_db_id=71;id=Mvestita_S11g13513-RA;hilite_coords=1-174) | 5·10^-123^ | N-α-acetyltransferase 20 | 1·10^-119^ |
| ARAKIN | [ADC02569](https://fernbase.org/tools/blast/match/show?blast_db_id=59;id=ADC02569;hilite_coords=14-673) | 0 | [Aspi01Gene30774.t1](https://fernbase.org/tools/blast/match/show?blast_db_id=63;id=Aspi01Gene30774.t1;hilite_coords=186-586) | 2·10^-178^ | Mitogen-activated protein kinase kinase kinase 1 | 2·10^-145^ | [Ceric.24G062200.1.v2.1](https://fernbase.org/tools/blast/match/show?blast_db_id=67;id=Ceric.24G062200.1.v2.1;hilite_coords=11-623) | 1·10^-177^ | [Mvestita_S12g16064-RA](https://fernbase.org/tools/blast/match/show?blast_db_id=71;id=Mvestita_S12g16064-RA;hilite_coords=291-600) | 1·10^-164^ | Mitogen-activated protein kinase kinase kinase 1 | 8·10^-157^ |
| PBL39 | [ADC19702](https://fernbase.org/tools/blast/match/show?blast_db_id=59;id=ADC19702;hilite_coords=102-472) | 0 | [Aspi01Gene59207.t2](https://fernbase.org/tools/blast/match/show?blast_db_id=63;id=Aspi01Gene59207.t2;hilite_coords=123-421) | 9·10^-112^ | Kinase superfamily protein | 3·10^-152^ | [Ceric.23G007000.2.v2.1](https://fernbase.org/tools/blast/match/show?blast_db_id=67;id=Ceric.23G007000.2.v2.1;hilite_coords=90-409) | 4·10^-166^ | [Mvestita_S10g02473-RA](https://fernbase.org/tools/blast/match/show?blast_db_id=71;id=Mvestita_S10g02473-RA;hilite_coords=63-378) | 2·10^-150^ | Kinase superfamily protein | 2·10^-152^ |
| TGA2 | ADC15791 | 0 | Aspi01Gene61998.t1 | 0 | Azfi-s0288.g063204 | 0 | Ceric.34G009100.1.v2.1 | 0 | Mvestita_S4g09172-RA | 0 | Sacu_v1.1_s0017.g007351 | 0 |
| FMO1 | ADC07644 | 0 | Aspi01Gene40270.t1 | 0 | Azfi-s0087.g042257 | 0,000156 | Ceric.23G070500.1.v2.1 | 0 | Mvestita_S18g14860-RA | 0 | Sacu_v1.1_s0103.g019960 | 1.78·10^-161^ |
| SFD1 | ADC21877 | 0 | Aspi01Gene23026.t1 | 0 | Azfi-s0001.g000343 | 0 | Ceric.12G048500.1.v2.1 | 0 | Mvestita_S10g03092-RA | 0 | Sacu_v1.1_s0101.g019781 | 0 |
| PBL34 | ADC02957 | 0 | Aspi01Gene09239.t2 | 0 | Azfi-s0012.g012929 | 0 | Ceric.04G073700.4.v2.1 | 0 | Mvestita_S18g14472-RA | 0 | Sacu_v1.1_s0173.g024449 | 0 |
| RBOH F | ADC08330 | 0 | Aspi01Gene72366.t1 | 0 | Azfi-s0250.g060197 | 0 | Ceric.08G028900.2.v2.1 | 0 | Mvestita_S10g02190-RA | 2.95·10^-150^ | Sacu_v1.1_s0083.g018180 | 9.75·10^-133^ |
| SYG1 | ADC17416 | 9.01·10^-145^ | Aspi01Gene39519.t1 | 2.9·10^-161^ | Azfi-s0002.g001574 | 7.6·10^-160^ | Ceric.35G011000.1.v2.1 | 1.95·10^-135^ | Mvestita_S19g17078-RA | 1.51·10^-151^ | Sacu_v1.1_s0037.g011896 | 2.47·10^-146^ |
| GED1 | ADC02237 | 0 | Aspi01Gene67141.t1 | 0 | Azfi-s0022.g015984 | 0 | Ceric.29G053500.14.v2.1 | 0 | Mvestita_S16g10412-RA | 0 | Sacu_v1.1_s0119.g021264 | 0 |
| NCH1 | ADC08585 | 0 | Aspi01Gene38498.t1 | 0 | Azfi-s0280.g062313 | 1.77·10^-173^ | Ceric.26G037900.3.v2.1 | 0 | Mvestita_S16g10484-RA | 0 | Sacu_v1.1_s0108.g020426 | 7.57·10^-166^ |
| Δ-OAT | ADC20863 | 0 | Aspi01Gene00959.t1 | 0 | Azfi-s0001.g000450 | 0 | Ceric.29G037300.2.v2.1 | 0 | Mvestita_S7g18464-RA | 0 | Sacu_v1.1_s0014.g006339 | 0 |
| PDR8 | [ADC21287](https://fernbase.org/tools/blast/match/show?blast_db_id=59;id=ADC21287;hilite_coords=1-1498) | 0 | [Aspi01Gene20765.t1](https://fernbase.org/tools/blast/match/show?blast_db_id=63;id=Aspi01Gene20765.t1;hilite_coords=1-1500) | 0 | ABC transporter family protein | 0 | [Ceric.10G030800.1.v2.1](https://fernbase.org/tools/blast/match/show?blast_db_id=67;id=Ceric.10G030800.1.v2.1;hilite_coords=1-1500) | 0 | [Mvestita_C87g20575-RA](https://fernbase.org/tools/blast/match/show?blast_db_id=71;id=Mvestita_C87g20575-RA;hilite_coords=1-1500) | 0 | ABC transporter family protein | 0 |
| CALS12 | [ADC18226](https://fernbase.org/tools/blast/match/show?blast_db_id=59;id=ADC18226;hilite_coords=99-1877) | 0 | [Aspi01Gene07129.t1](https://fernbase.org/tools/blast/match/show?blast_db_id=63;id=Aspi01Gene07129.t1;hilite_coords=1-1766) | 0 | Callose synthase-like protein | 0 | [Ceric.12G070800.2.v2.1](https://fernbase.org/tools/blast/match/show?blast_db_id=67;id=Ceric.12G070800.2.v2.1;hilite_coords=1-1768) | 0 | [Mvestita_S18g14908-RA](https://fernbase.org/tools/blast/match/show?blast_db_id=71;id=Mvestita_S18g14908-RA;hilite_coords=3-1728) | 0 | Callose synthase-like protein | 0 |
| VHP1 | [ADC07688](https://fernbase.org/tools/blast/match/show?blast_db_id=59;id=ADC07688;hilite_coords=2-769) | 0 | [Aspi01Gene56107.t1](https://fernbase.org/tools/blast/match/show?blast_db_id=63;id=Aspi01Gene56107.t1;hilite_coords=1-767) | 0 | Pyrophosphate-energized vacuolar membrane proton pump | 0 | [Ceric.06G049000.3.v2.1](https://fernbase.org/tools/blast/match/show?blast_db_id=67;id=Ceric.06G049000.3.v2.1;hilite_coords=1-770) | 0 | [Mvestita_S15g12567-RA](https://fernbase.org/tools/blast/match/show?blast_db_id=71;id=Mvestita_S15g12567-RA;hilite_coords=1-775) | 0 | Pyrophosphate-energized vacuolar membrane proton pump | 0 |
| SIS7 | [ADC22118](https://fernbase.org/tools/blast/match/show?blast_db_id=59;id=ADC22118;hilite_coords=46-475) | 0 | [Aspi01Gene05062.t1](https://fernbase.org/tools/blast/match/show?blast_db_id=63;id=Aspi01Gene05062.t1;hilite_coords=61-496) | 0 | 9-cis-epoxycarotenoid dioxygenase | 3·10^-109^ | [Ceric.36G058200.1.v2.1](https://fernbase.org/tools/blast/match/show?blast_db_id=67;id=Ceric.36G058200.1.v2.1;hilite_coords=105-537) | 2·10^-127^ | [Mvestita_S20g18856-RA](https://fernbase.org/tools/blast/match/show?blast_db_id=71;id=Mvestita_S20g18856-RA;hilite_coords=135-520) | 6·10^-122^ | 9-cis-epoxycarotenoid dioxygenase | 3·10^-121^ |
| CBL1 | ADC20220 | 3.47·10^-132^ | Aspi01Gene14439.t1 | 3.89·10^-130^ | Azfi-s0047.g030262 | 2.51·10^-121^ | Ceric.34G029300.3.v2.1 | 2.58·10^-118^ | Mvestita_S18g14415-RA | 8.07·10^-112^ | Sacu_v1.1_s0028.g009880 | 5.44·10^-124^ |
| MSRA4 | [ADC26617](https://fernbase.org/tools/blast/match/show?blast_db_id=59;id=ADC26617;hilite_coords=29-296) | 1·10^-150^ | [Aspi01Gene11617.t1](https://fernbase.org/tools/blast/match/show?blast_db_id=63;id=Aspi01Gene11617.t1;hilite_coords=1-275) | 4·10^-150^ | Peptide methionine sulfoxide reductase | 2·10^-117^ | [Ceric.33G048500.1.v2.1](https://fernbase.org/tools/blast/match/show?blast_db_id=67;id=Ceric.33G048500.1.v2.1;hilite_coords=9-280) | 1·10^-184^ | [Mvestita_S14g07955-RA](https://fernbase.org/tools/blast/match/show?blast_db_id=71;id=Mvestita_S14g07955-RA;hilite_coords=47-318) | 2·10^-139^ | Peptide methionine sulfoxide reductase | 2·10^-125^ |
| ACO3 | [ADC14585](https://fernbase.org/tools/blast/match/show?blast_db_id=59;id=ADC14585;hilite_coords=1-978) | 0 | [Aspi01Gene64046.t1](https://fernbase.org/tools/blast/match/show?blast_db_id=63;id=Aspi01Gene64046.t1;hilite_coords=1-977) | 0 | Aconitate hydratase | 0 | [Ceric.10G044400.1.v2.1](https://fernbase.org/tools/blast/match/show?blast_db_id=67;id=Ceric.10G044400.1.v2.1;hilite_coords=1-983) | 0 | [Mvestita_S20g18823-RA](https://fernbase.org/tools/blast/match/show?blast_db_id=71;id=Mvestita_S20g18823-RA;hilite_coords=63-975) | 0 | Aconitate hydratase | 0 |
| LCNP | ADC31767 | 0 | Aspi01Gene03135.t2 | 0 | Azfi-s0114.g046130 | 2.05·10^-157^ | Ceric.12G059300.1.v2.1 | 8.71·10^-165^ | Mvestita_S15g13306-RA | 1.66·10^-160^ | Sacu_v1.1_s0125.g021654 | 3.1·10^-161^ |
| EXE2 | ADC13464 | 0 | Aspi01Gene35610.t1 | 0 | Azfi-s0008.g011467 | 0,037 | Ceric.30G051700.1.v2.1 | 0 | Mvestita_S7g18423-RA | 0 | Sacu_v1.1_s0002.g000943 | 0 |
| ABC1K7 | ADC28122 | 0 | Aspi01Gene64383.t1 | 0 | Azfi-s0027.g023543 | 0 | Ceric.31G066400.1.v2.1 | 0 | Mvestita_S17g16623-RA | 0 | Sacu_v1.1_s0013.g005902 | 0 |
| ABC1K8 | ADC09692 | 0 | Aspi01Gene70662.t1 | 0 | Azfi-s0102.g044540 | 0 | Ceric.29G083900.5.v2.1 | 0 | Mvestita_S10g03061-RA | 0 | Sacu_v1.1_s0050.g013803 | 0 |
| A/N-INVA | ADC11476 | 4.77·10^-150^ | Aspi01Gene55297.t1 | 0 | Azfi-s0010.g012047 | 1.46·10^-93^ | Ceric.30G074500.5.v2.1 | 2.47·10^-156^ | Mvestita_S10g02958-RA | 6.94·10^-147^ | Sacu_v1.1_s0024.g009176 | 1.15·10^-123^ |
| ENF1 | [ADC16403](https://fernbase.org/tools/blast/match/show?blast_db_id=59;id=ADC16403;hilite_coords=7-530) | 0 | [Aspi01Gene37467.t1](https://fernbase.org/tools/blast/match/show?blast_db_id=63;id=Aspi01Gene37467.t1;hilite_coords=196-432) | 3·10^-108^ | Aldehyde dehydrogenase | 0 | [Ceric.07G031600.1.v2.1](https://fernbase.org/tools/blast/match/show?blast_db_id=67;id=Ceric.07G031600.1.v2.1;hilite_coords=35-532) | 0 | [Mvestita_S7g18565-RA](https://fernbase.org/tools/blast/match/show?blast_db_id=71;id=Mvestita_S7g18565-RA;hilite_coords=50-548) | 0 | Aldehyde dehydrogenase | 0 |
| GGT1 | ADC07019 | 0 | Aspi01Gene13594.t1 | 0 | Azfi-s0007.g011140 | 5.05·10^-164^ | Ceric.10G003000.1.v2.1 | 0 | Mvestita_S11g13432-RA | 0 | Sacu_v1.1_s0008.g004284 | 0 |
| GAPC1 | ADC10633 | 8.42·10^-115^ | Aspi01Gene23189.t1 | 7.14·10^-110^ | Azfi-s0010.g012122 | 2.96·10^-105^ | Ceric.1Z012200.3.v2.1 | 2.51·10^-107^ | Mvestita_S13g01679-RA | 4.77·10^-106^ | Sacu_v1.1_s0059.g015030 | 5.96·10^-97^ |
| GLYR1 | ADC17932 | 8.23·10^-174^ | Aspi01Gene52549.t1 | 5.55·10^-166^ | Azfi-s0288.g063215 | 6.4·10^-177^ | Ceric.38G018500.1.v2.1 | 3.51·10^-171^ | Mvestita_S15g12560-RA | 1.8·10^-170^ | Sacu_v1.1_s0189.g025223 | 1.1·10^-170^ |
| NCER1 | ADC06915 | 0 | Aspi01Gene23715.t1 | 0 | Azfi-s0006.g010345 | 0 | Ceric.14G034200.1.v2.1 | 0 | Mvestita_S7g18647-RA | 0 | Sacu_v1.1_s0107.g020391 | 0 |
| GMII | [ADC02861](https://fernbase.org/tools/blast/match/show?blast_db_id=59;id=ADC02861;hilite_coords=1-1148) | 0 | [Aspi01Gene19432.t1](https://fernbase.org/tools/blast/match/show?blast_db_id=63;id=Aspi01Gene19432.t1;hilite_coords=1-1154) | 0 | α-mannosidase | 0 | [Ceric.24G053700.2.v2.1](https://fernbase.org/tools/blast/match/show?blast_db_id=67;id=Ceric.24G053700.2.v2.1;hilite_coords=1-1147) | 0 | [Mvestita_S1g06990-RA](https://fernbase.org/tools/blast/match/show?blast_db_id=71;id=Mvestita_S1g06990-RA;hilite_coords=36-1170) | 0 | α-mannosidase | 0 |
| BCHA1 | [ADC10001](https://fernbase.org/tools/blast/match/show?blast_db_id=59;id=ADC10001;hilite_coords=1-3612) | 0 | [Aspi01Gene37984.t1](https://fernbase.org/tools/blast/match/show?blast_db_id=63;id=Aspi01Gene37984.t1;hilite_coords=1-3648) | 0 | Beige/BEACH domain containing protein | 0 | [Ceric.12G054200.1.v2.1](https://fernbase.org/tools/blast/match/show?blast_db_id=67;id=Ceric.12G054200.1.v2.1;hilite_coords=1-3624) | 0 | [Mvestita_S7g18523-RA](https://fernbase.org/tools/blast/match/show?blast_db_id=71;id=Mvestita_S7g18523-RA;hilite_coords=1-1933) | 0 | Beige/BEACH domain containing protein | 0 |
| **Protein** | ***Adiantum capillus* protein** | **E-value** | ***Alsophila spinulosa* protein** | **E-value** | ***Azolla filiculoides* protein** | **E-value** | ***Ceratopteris richardii* protein** | **E-value** | ***Marsilea vestita* protein** | **E-value** | ***Salvinia cucullata* protein** | **E-value** |
| RH3 | ADC14450 | 0 | Aspi01Gene57148.t1 | 0 | Azfi-s0166.g054407 | 0 | Ceric.05G054100.2.v2.1 | 0 | Mvestita_S8g04877-RA | 0 | Sacu_v1.1_s0230.g026421 | 0 |
| MRF3 | ADC02232 | 0 | Aspi01Gene63305.t1 | 0 | Azfi-s0520.g075252 | 0 | Ceric.27G043500.1.v2.1 | 0 | Mvestita_S20g19240-RA | 0 | Sacu_v1.1_s0109.g020494 | 0 |
| SRL1 | ADC16567 | 1.79·10^-134^ | Aspi01Gene27067.t1 | 1.59·10^-167^ | Azfi-s0185.g056662 | 0,00000236 | Ceric.07G036200.3.v2.1 | 2.06·10^-156^ | Mvestita_S19g17065-RA | 5.78·10^-148^ | Sacu_v1.1_s0207.g025788 | 2.54·10^-153^ |
| PP2CG1 | ADC05224 | 0 | Aspi01Gene69503.t1 | 0 | Azfi-s3252.g115111 | 0 | Ceric.09G079600.1.v2.1 | 6.16·10^-147^ | Mvestita_C25g19591-RA | 0 | Sacu_v1.1_s0001.g000643 | 0 |
| SZL1 | [ADC27530](https://fernbase.org/tools/blast/match/show?blast_db_id=59;id=ADC27530;hilite_coords=1-501) | 0 | [Aspi01Gene05110.t1](https://fernbase.org/tools/blast/match/show?blast_db_id=63;id=Aspi01Gene05110.t1;hilite_coords=1-515) | 0 | Lycopene β-cyclase | 0 | [Ceric.10G080900.1.v2.1](https://fernbase.org/tools/blast/match/show?blast_db_id=67;id=Ceric.10G080900.1.v2.1;hilite_coords=1-507) | 0 | [Mvestita_S18g14468-RA](https://fernbase.org/tools/blast/match/show?blast_db_id=71;id=Mvestita_S18g14468-RA;hilite_coords=1-494) | 0 | Lycopene β-cyclase | 0 |
| PK2 | ADC21965 | 0 | Aspi01Gene58133.t1 | 0 | Azfi-s0541.g076357 | 4.63·10^-69^ | Ceric.15G073900.5.v2.1 | 0 | Mvestita_S18g14660-RA | 0 | Sacu_v1.1_s0020.g008211 | 3.38·10^-66^ |
| CBG | [ADC04972](https://fernbase.org/tools/blast/match/show?blast_db_id=59;id=ADC04972;hilite_coords=372-640) | 0 | [Aspi01Gene10892.t1](https://fernbase.org/tools/blast/match/show?blast_db_id=63;id=Aspi01Gene10892.t1;hilite_coords=290-558) | 0 | Mitochondrial Rho GTPase | 0 | [Ceric.32G015700.1.v2.1](https://fernbase.org/tools/blast/match/show?blast_db_id=67;id=Ceric.32G015700.1.v2.1;hilite_coords=372-640) | 0 | [Mvestita_S6g00686-RA](https://fernbase.org/tools/blast/match/show?blast_db_id=71;id=Mvestita_S6g00686-RA;hilite_coords=378-646) | 0 | Mitochondrial Rho GTPase | 0 |
| NFXL1 | [ADC26894](https://fernbase.org/tools/blast/match/show?blast_db_id=59;id=ADC26894;hilite_coords=19-1305) | 0 | [Aspi01Gene27607.t1](https://fernbase.org/tools/blast/match/show?blast_db_id=63;id=Aspi01Gene27607.t1;hilite_coords=77-1427) | 0 | NF-X1-type zinc finger protein NFXL1 | 0 | [Ceric.20G009600.2.v2.1](https://fernbase.org/tools/blast/match/show?blast_db_id=67;id=Ceric.20G009600.2.v2.1;hilite_coords=104-1240) | 0 | [Mvestita_S2g11856-RA](https://fernbase.org/tools/blast/match/show?blast_db_id=71;id=Mvestita_S2g11856-RA;hilite_coords=298-1378) | 0 | NF-X1-type zinc finger protein NFXL1 | 0 |
| PEUP2 | [ADC23426](https://fernbase.org/tools/blast/match/show?blast_db_id=59;id=ADC23426;hilite_coords=177-578) | 0 | [Aspi01Gene65700.t1](https://fernbase.org/tools/blast/match/show?blast_db_id=63;id=Aspi01Gene65700.t1;hilite_coords=34-293) | 1·10^-159^ | Homolog of yeast autophagy 18 (ATG18) D | 0 | [Ceric.14G026400.1.v2.1](https://fernbase.org/tools/blast/match/show?blast_db_id=67;id=Ceric.14G026400.1.v2.1;hilite_coords=11-412) | 0 | [Mvestita_S14g07639-RA](https://fernbase.org/tools/blast/match/show?blast_db_id=71;id=Mvestita_S14g07639-RA;hilite_coords=34-424) | 0 | Homolog of yeast autophagy 18 (ATG18) D | 6·10^-141^ |
| DIN4 | ADC18809 | 0 | Aspi01Gene63710.t1 | 1.9·10^-154^ | Azfi-s0257.g060580 | 0 | Ceric.33G059300.1.v2.1 | 0 | Mvestita_S12g15352-RA | 0 | Sacu_v1.1_s0015.g006453 | 0 |
| PKS17 | ADC21863 | 0 | Aspi01Gene71542.t1 | 0 | Azfi-s0329.g065362 | 0 | Ceric.33G018700.2.v2.1 | 0 | Mvestita_S20g18854-RA | 0 | Sacu_v1.1_s0037.g011806 | 0 |
| PKS6 | ADC00670 | 0 | Aspi01Gene68009.t1 | 0 | Azfi-s0102.g044556 | 0 | Ceric.06G029500.1.v2.1 | 0 | Mvestita_S6g00482-RA | 0 | Sacu_v1.1_s0037.g011842 | 0 |
| GCN2 | ADC19360 | 0 | Aspi01Gene29301.t2 | 0 | Azfi-s0158.g053868 | 0 | Ceric.19G008700.4.v2.1 | 0 | Mvestita_S4g08887-RA | 0 | Sacu_v1.1_s0156.g023677 | 0 |
| GRF3 | ADC14843 | 3.42·10^-153^ | Aspi01Gene47077.t1 | 3.32·10^-156^ | Azfi-s0531.g075854 | 4.8·10^-143^ | Ceric.37G003300.1.v2.1 | 2.11·10^-143^ | Mvestita_S10g02293-RA | 3.11·10^-151^ | Sacu_v1.1_s0021.g008364 | 2.58·10^-129^ |
| KIN17 | ADC31842 | 0 | Aspi01Gene60127.t1 | 0 | Azfi-s0020.g015536 | 0,00205 | Ceric.16G010000.1.v2.1 | 0 | Mvestita_S5g09848-RA | 0 | Sacu_v1.1_s0093.g019017 | 1.71·10^-130^ |
| PAH1 | ADC01543 | 0 | Aspi01Gene31422.t1 | 1.86·10^-165^ | Azfi-s0042.g026916 | 1·10^-168^ | Ceric.08G000300.2.v2.1 | 0 | Mvestita_S16g10443-RA | 7.62·10^-158^ | Sacu_v1.1_s0002.g000957 | 3.23·10^-157^ |
| ADS3 | ADC20692 | 3.4·10^-159^ | Aspi01Gene17758.t1 | 0 | Azfi-s0034.g025363 | 0 | Ceric.37G050600.1.v2.1 | 8.95·10^-167^ | Mvestita_S18g15019-RA | 1.09·10^-173^ | Sacu_v1.1_s0059.g015051 | 1.59·10^-170^ |
| PFC1 | ADC13607 | 2.48·10^-152^ | Aspi01Gene27022.t1 | 3.2·10^-93^ | Azfi-s0396.g067875 | 7.82·10^-158^ | Ceric.30G050300.1.v2.1 | 5.98·10^-163^ | Mvestita_S1g06946-RA | 3.02·10^-149^ | Sacu_v1.1_s0003.g001849 | 2.1·10^-134^ |
| BAM3 | [ADC05334](https://fernbase.org/tools/blast/match/show?blast_db_id=59;id=ADC05334;hilite_coords=1-605) | 0 | [Aspi01Gene69400.t1](https://fernbase.org/tools/blast/match/show?blast_db_id=63;id=Aspi01Gene69400.t1;hilite_coords=1-605) | 0 | β-amylase | 0 | [Ceric.19G057400.1.v2.1](https://fernbase.org/tools/blast/match/show?blast_db_id=67;id=Ceric.19G057400.1.v2.1;hilite_coords=1-598) | 0 | [Mvestita_S6g00244-RA](https://fernbase.org/tools/blast/match/show?blast_db_id=71;id=Mvestita_S6g00244-RA;hilite_coords=90-592) | 0 | β-amylase | 0 |
| FUM2 | [ADC17481](https://fernbase.org/tools/blast/match/show?blast_db_id=59;id=ADC17481;hilite_coords=3-499) | 0 | [Aspi01Gene64302.t1](https://fernbase.org/tools/blast/match/show?blast_db_id=63;id=Aspi01Gene64302.t1;hilite_coords=25-284) | 3·10^-156^ | Fumarate hydratase 1 family protein | 0 | [Ceric.28G057900.1.v2.1](https://fernbase.org/tools/blast/match/show?blast_db_id=67;id=Ceric.28G057900.1.v2.1;hilite_coords=3-499) | 0 | [Mvestita_S3g04271-RA](https://fernbase.org/tools/blast/match/show?blast_db_id=71;id=Mvestita_S3g04271-RA;hilite_coords=46-514) | 0 | Fumarate hydratase 1 family protein | 2·10^-66^ |
| CHY1 | ADC15564 | 0 | Aspi01Gene09646.t1 | 0 | Azfi-s0091.g042901 | 4.38·10^-175^ | Ceric.13G015500.1.v2.1 | 6.3·10^-98^ | Mvestita_S10g02707-RA | 2.75·10^-170^ | Sacu_v1.1_s0022.g008725 | 0 |
| SFR2 | ADC04500 | 0 | Aspi01Gene33715.t1 | 0 | Azfi-s0048.g030426 | 0 | Ceric.33G055200.9.v2.1 | 0 | Mvestita_S12g15673-RA | 0 | Sacu_v1.1_s0021.g008312 | 0 |
| ENO2 | ADC20480 | 0 | Aspi01Gene10045.t1 | 0 | Azfi-s0013.g013345 | 0 | Ceric.31G055000.3.v2.1 | 0 | Mvestita_S4g08684-RA | 0 | Sacu_v1.1_s0007.g003787 | 0 |
| SLD2 | ADC23332 | 0 | Aspi01Gene37001.t1 | 0 | Azfi-s0062.g035143 | 0 | Ceric.25G067500.1.v2.1 | 0 | Mvestita_C154g21216-RA | 0 | Sacu_v1.1_s0144.g022965 | 0 |
| LCBK2 | ADC17803 | 8.14·10^-47^ | Aspi01Gene61173.t1 | 8.61·10^-102^ | Azfi-s0028.g023891 | 1.03·10^-128^ | Ceric.32G057000.1.v2.1 | 2.1·10^-125^ | Mvestita_S18g14615-RA | 3.29·10^-124^ | Sacu_v1.1_s0068.g016364 | 1.56·10^-39^ |
| KH28 | [ADC01782](https://fernbase.org/tools/blast/match/show?blast_db_id=59;id=ADC01782;hilite_coords=1-598) | 0 | [Aspi01Gene68976.t1](https://fernbase.org/tools/blast/match/show?blast_db_id=63;id=Aspi01Gene68976.t1;hilite_coords=25-551) | 0 | RNA-binding KH domain protein | 0 | [Ceric.17G032000.2.v2.1](https://fernbase.org/tools/blast/match/show?blast_db_id=67;id=Ceric.17G032000.2.v2.1;hilite_coords=1-645) | 0 | [Mvestita_S10g02999-RA](https://fernbase.org/tools/blast/match/show?blast_db_id=71;id=Mvestita_S10g02999-RA;hilite_coords=72-720) | 0 | RNA-binding KH domain protein | 0 |
| HIT4 | ADC03227 | 1.58·10^-161^ | Aspi01Gene39881.t1 | 6.95·10^-114^ | Azfi-s0998.g094968 | 2.07·10^-105^ | Ceric.34G002200.1.v2.1 | 7.59·10^-165^ | Mvestita_S7g17986-RA | 6.71·10^-137^ | - | - |
| PP7 | ADC20843 | 0 | Aspi01Gene54672.t1 | 0 | Azfi-s0383.g067449 | 2.54·10^-136^ | Ceric.25G051200.2.v2.1 | 0 | Mvestita_S3g03772-RA | 0 | Sacu_v1.1_s0016.g006976 | 2.64·10^-166^ |
